# Connectome-constrained deep mechanistic networks predict neural responses across the fly visual system at single-neuron resolution

**DOI:** 10.1101/2023.03.11.532232

**Authors:** Janne K. Lappalainen, Fabian D. Tschopp, Sridhama Prakhya, Mason McGill, Aljoscha Nern, Kazunori Shinomiya, Shin-ya Takemura, Eyal Gruntman, Jakob H. Macke, Srinivas C. Turaga

## Abstract

We can now measure the connectivity of every neuron in a neural circuit, but we are still blind to other biological details, including the dynamical characteristics of each neuron. The degree to which connectivity measurements alone can inform understanding of neural computation is an open question. Here we show that with only measurements of the connectivity of a biological neural network, we can predict the neural activity underlying neural computation. We constructed a model neural network with the experimentally determined connectivity for 64 cell types in the motion pathways of the fruit fly optic lobe but with unknown parameters for the single neuron and single synapse properties. We then optimized the values of these unknown parameters using techniques from deep learning, to allow the model network to detect visual motion. Our mechanistic model makes detailed experimentally testable predictions for each neuron in the connectome. We found that model predictions agreed with experimental measurements of neural activity across 24 studies. Our work demonstrates a strategy for generating detailed hypotheses about the mechanisms of neural circuit function from connectivity measurements. We show that this strategy is more likely to be successful when neurons are sparsely connected—a universally observed feature of biological neural networks across species and brain regions.

## Introduction

Electrical signals propagating through networks of neurons in the nervous system form the basis of computations such as the visual detection of movement. The propagation of neural activity is shaped by both the functional properties of individual neurons and their synaptic connectivity. In addition, multiple additional factors^1,2^ including electrical synapses^3,4^, neuromodulation^5^, and glia^6^ are known to further influence neural activity on multiple time-scales. Volume electron microscopy can now be used to comprehensively measure the connectivity of every neuron in a neural circuit, and even entire nervous systems^7–24^. However, we do not yet have the means to also comprehensively measure all other biological details, including the dynamical properties of every neuron and synapse in the same circuit^2^. For these reasons, there has been considerable debate about the utility of connectomic measurements for understanding brain function^25^. Is it possible to use measurements of only neural connectivity to generate accurate predictions about how the neural circuit functions? Even in the complete absence of direct measurements of neural activity from a living brain?

There is considerable evidence from computer science and neuroscience that there is not usually a strong link between the connectivity of a neural network and its computational function. Universal function approximation theorems for artificial neural networks^26–28^ imply that the same computational task can be performed by many different networks with very different neural connectivity. Empirically, there exist many classes of general purpose artificial neural network architectures which can trained to perform the same large diversity of computational tasks^29^. Such differences in connectivity can correspond to qualitatively different computational mechanisms^30,31^. Similarly in neuroscience, there have been competing proposals by theorists for example, for the neural circuit mechanisms of the computation of visual motion^32,33^, and for the integration of eye velocity commands into eye position signals^34,35^, with each proposal suggesting different neural connectivity. Further, circuits with the same connectivity can function differently^5^. Thus in general, neither the connectivity of a circuit alone, nor its computational task alone, can uniquely determine the mechanism of circuit function^36–38^.

Here we show that the connectivity of a neural circuit, together with knowledge of its computational task, enables accurate predictions of the role played by individual neurons in the circuit in the computational task. We constructed a differentiable^39, 40^ model neural network with a close correspondence to the brain, whose connectivity was given by connectomic measurements and with unknown single neuron and single synapse parameters. We optimized the unknown parameters of the model network using techniques from deep learning^41–43^, to enable the model network to accomplish the computational task^44–47^. We call such models connectome-constrained and task-optimized deep mechanistic networks (DMNs; Fig. 1a).

**Figure 1:**
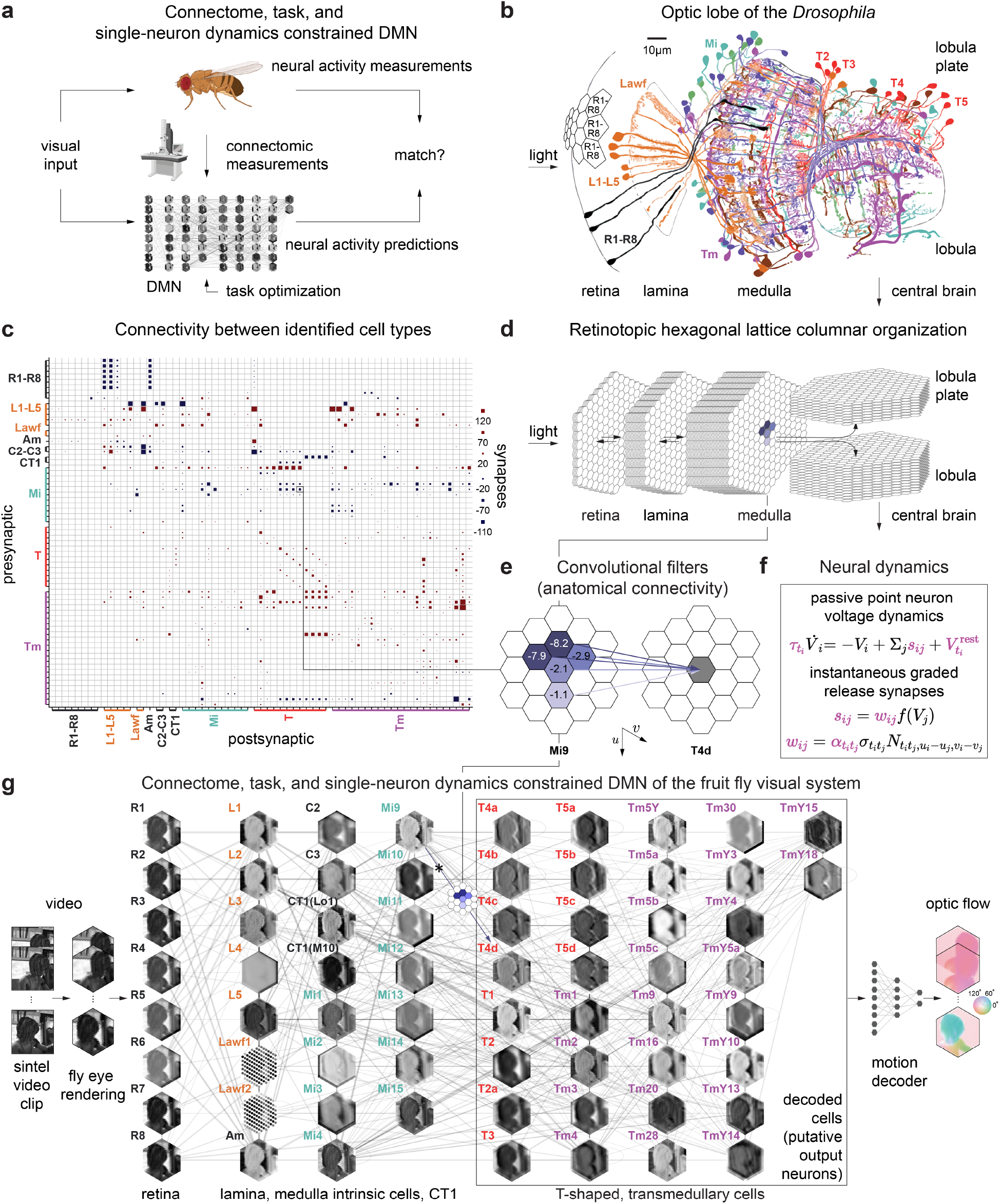
Task-optimizing connectome-constrained models of the fly visual system. **(a)** The ‘deep mechanistic network model’ (DMN) aims to satisfy three constraints simultaneously: Its archiecture is based on connectomic measurements (see b-e), single-neuron and synaptic dynamics are given by simple mechanistic models (see f), and free parameters of this network model (see f, magenta) are optimized by training the model to perform an optic flow estimation task (see g). **(b)** Schematic of optic lobe of *Drosophila* melanogaster comprising several processing stages (neuropils) and cell types (adapted from^65^). Our model includes retinal cells, lamina monopolar cells, medulla intrinsic cells, transmedullary cells, and T-shaped cells, i.e., all columnar cell types. **(c)** Identified connectivity between 64 cell types, represented by total number of input synapses from all neurons of a given presynaptic cell type to a single postsynaptic of a given cell type. Blue color indicates putative hyperpolarizing inputs, red putative depolarizing inputs as inferred from neurotransmitter and receptor profiling. Size of squares indicates number of input synapses. **(d)** Retinotopic hexagonal lattice columnar organization of the visual system model. Each lattice represents a cell type, each hexagon an individual cell. Positions of photoreceptor columns are aligned with positions of downstream columns. The model comprises synapses from all neuropils (SI Fig. 1) and downstream and upstream projecting connections from the retina, lamina, and medulla. Convolutional filter between Mi9 cells and a T4d cell (see Panel d) is highlighted in the final lattice of the medulla. **(e)** Example of convolutional filter, representing Mi9 inputs onto T4d cells. Values represent the average number of synapses projecting from presynaptic Mi9 cells in columns with indicated offset onto the postsynaptic dendrite of T4d cells. Values indicate connection strength derived from electron microscopy data. **(f)** Single-neuron and synaptic dynamics are given by simple mechanistic models. Free parameters of this network model (magenta) are optimized by training the model to perform optic flow estimation task. **(g)** Visualization of DMN performing optic flow estimation on a video clip rendered from the Sintel dataset. Given the input to the photoreceptors (R1-R8), we simulated the response of each neuron in the system. Each hexagonal lattice depicts a snapshot of the voltage levels of all cells from the corresponding cell type. Edges illustrate connectivity between identified cell types. A decoding network receives the simulated neural activity of all output neurons (T-shaped and transmedullary cells) to compute optic flow. Parameters of the recurrent network model and the decoding network are optimized using optimization methods from deep learning.

We applied this approach to model the motion pathways in the optic lobe of the *Drosophila* visual system. We constructed a DMN with experimentally measured connectivity^48–50^, and unknown parameters for the single neuron dynamics and the strength of a unitary synapse. We optimized the unknown model parameters on the computer vision task of computing visual motion from dynamic visual stimuli. Visual motion computation in the fly visual system and its mechanistic underpinnings have been extensively studied^51–56^. Thus, we were able to compare the detailed predictions of our model with experimental measurements of neural activity in response to visual stimuli, on a neuron-by-neuron basis. We found that our connectome-constrained and task-optimized DMN accurately predicts the well-known segregation of the visual system into light increment (ON) and light decrement (OFF) channels^46,57–61^, as well as the direction selectivity of the well-known T4 and T5 motion detector neurons^53,62–64^. Our model further suggests that TmY3 might also detect motion, a prediction that has yet to be experimentally tested. We release our model as a resource for the community.^1^

## Results

### A connectome-constrained deep mechanistic network of the fruit fly visual system

The optic lobes of the fruit fly are involved in early visual processing. They comprise several layered neuropiles whose columnar arrangement has a one-to-one correspondence with the ommatidia, both possessing a remarkably crystalline organization in a hexagonal lattice. Visual input from the photoreceptors is received by the lamina, which sends projections to the medulla, lobula, and lobula plate (Fig. 1b^65^). Many components of the optic lobe are highly periodic, with columnar cell types appearing once per column, and multi-columnar neurons appearing with only small deviations from a well-defined periodicity in columnar space^65, 66^. A complete reconstruction of the optic lobe is yet unavailable, but several studies have reported on the local connectivity within the lamina^50^, primarily focused on the light increment (ON) and light decrement (OFF) selective components of the motion pathways of visual processing in the medulla^67,68^, and in the medulla, lobula, and lobula plate^48, 49, 69^. We assembled these separate local reconstructions into a coherent local connectome spanning the retina, lamina, medulla, lobula, and lobula plate (Fig. 1c, SI Fig.1). We approximated the circuitry across the entire visual field as perfectly periodi^c66,68^, and tiled this local connectivity architecture in hexagonal lattice across retinotopic space to construct a connectome for 64 cell types across the central visual field of the right eye (Fig. 1d; Methods).

We built a recurrent neural network modeling these first stages of visual processing in the optic lobe based on the connectome for the right eye. Each neuron in this DMN corresponds to a real neuron in the fly visual system, belonging to an identified cell type, and is connected to other neurons only if they are connected by synapses in the connectome (Fig. 1e). Our goal was to investigate whether precise synaptic connectivity and task-constraints are sufficient to account for neural tuning across the fly visual system. We therefore constructed a model with detailed connectivity, but simplified models of single neurons and chemical synapses (Fig. 1f). As many neurons in the early visual system are non-spiking, we used passive leaky linear non-spiking voltage dynamics to model the time-varying activity of single neurons. We modeled neurons as point-neurons with a single electrical compartment, as this has been previously shown to be a good approximation given the small size of many neurons in the optic lobe^54^. As the exception, we modeled the CT1 neuron with multiple compartments since it is an exceptionally large single neuron spanning the entire optic lobe and is highly electrotonically compartmentalized^70^. We effectively modeled the neuron as two columnar “cell types”, with one CT1 compartment per column in the 10th layer of the medulla CT1(M10) and one per column in the 1st layer of the lobula CT1(Lo1) (SI Supplementary Note 2). We coupled neurons with chemical synapses whose connectivity was determined by the connectome. We developed a simplified model for a graded release chemical synapse between non-spiking neurons: A threshold-linear nonlinear function models the nonlinearity of the time-averaged concentration of synaptic release as a function of presynaptic voltage. The resulting network model follows well-known threshold-linear dynamics and is differentiable. Such dynamics are typically used to approximate the firing rates of a network of spiking neurons^71, 72^, whereas in our network, the threshold nonlinearity results from the nonlinear voltage-gated release of neuro-transmitter.

We used the cell type structure of the connectome, and assumption of perfect translation invariance across retinotopic space to reduce the number of free parameters (Fig. 1f). We assumed that neurons of the same cell type shared the same neuron time constant and resting membrane potential. We modeled a synaptic weight as proportional to the number of discrete synapses measured experimentally between those two neurons^73^, with a scale factor representing the strength of a unitary synapse. The unitary synapse scaling and the sign of each synapse was the same across all pairs of neurons with the same pre- and postsynaptic cell type. Likewise, the synapse count between each pair of neurons was the same across all pairs of neurons with the same pre- and postsynaptic cell type, and their relative location in retinotopic space.

In total, the connectome-constrained model comprises 45,669 neurons and 1,513,231 connections, across 65 cell types arranged in a hexagonal lattice consisting of 721 columns, modeling the central visual field of the roughly 700-900 ommatidia typically found in the fruit fly retina^74,75^. Connectomic constraints, and our assumption of spatial homogeneity (i.e., the hexagonally convolutional structure of the network) result in a dramatic reduction to just 734 free parameters for this large network model. The only free parameters in our model are the single neuron time constants and resting membrane potentials (65 parameters each), and the unitary synapse strengths (604 parameters). In the absence of connectomic measurements, if we had to infer the type-to-type as well as spatial connectivity between all pairs of cell types, we would instead have needed to estimate well over a million parameters (Methods).

We further constrained the parameters of the model by task-optimization, i.e. by training the model to perform a computational task which is thought to approximate the computations carried out by the system^m44,76^. We therefore implemented our recurrent DMN using the PyTorch library^39,40^ (Methods) and used automatic differentiation to optimize the model using gradient based deep learning training methods^41–43^.

As the functional task, we chose the computation of visual motion from naturalistic visual stimuli^77^. Visual motion computation in the fly visual system and its mechanistic underpinnings have been extensively studied^51–56^. This challenging computation requires the neural circuit to compare visual stimuli across space and time, and thereby critically relies on temporal integration of visual information by the dynamics of the network. We hypothesized that training our network model to perform the computer vision task of optic flow computation^77^ could help us identify circuit elements involved in motion computation. Since our model contains many of the circuit elements which have been experimentally characterized and implicated in the computation of visual motion, we could then validate our model predictions.

To decode optic flow from the DMN, we used a decoding network which maps its output to the computer vision representation of optic flow. This two-layer convolutional decoding network is allowed to use only the instantaneous neural activity of the medulla and downstream areas as input. Importantly, the decoding network cannot by itself detect motion, which requires the comparison of current and past visual stimuli, but must instead rely on the temporal dynamics of the connectome-constrained network to compute motion-selective visual features. The resulting combination of our recurrent connectome-constrained DMN model and the feedforward decoding network was then trained end-to-end: We rendered video sequences from the Sintel database^77^ as direct input to the photoreceptors of the connectome-constrained model (ignoring neuronal superposition^78^), and used gradient descent (backpropgation through time^79^) to minimize the task error in predicting optic flow (Fig. 1g, Methods).

### Ensembles of connectome-constrained and task-optimized DMNs robustly predict known tuning properties

We used only connectome and task-constraints to construct our DMN, without any measurements of neural activity. We can therefore validate the model by computing predictions of neural selectivity for each of the 64 identified cell types in the model, and comparing them to experimental measurements. Since it is possible that connectome and task-constraints might not uniquely constrain all model parameters^80–82^, we generated an ensemble of 50 models, all constrained with the same connectome, and optimized to perform the same task. Each model in the ensemble corresponds to a local optimum of task performance. Since the models achieved similar (but not identical) task performance, the ensemble reflects the diversity of possible models consistent with the connectome and task-constraints.

The ensemble of models found a variety of parameter configurations (Extended Data Fig. 2), and models with random parameter configurations—but trained decoding networks—consistently performed worse on the task than task-optimized models (Extended Data Fig. 3). We focused on the 10 models which achieved the best task performance (Fig. 2a). We simulated neural responses to multiple experimentally characterized visual stimuli, and comprehensively compared model responses for each cell type to experimentally reported responses from 24 previous studies^53–55,58–60,62,64,70,83–97^ (Overview in Supplemental Data).

**Figure 2:**
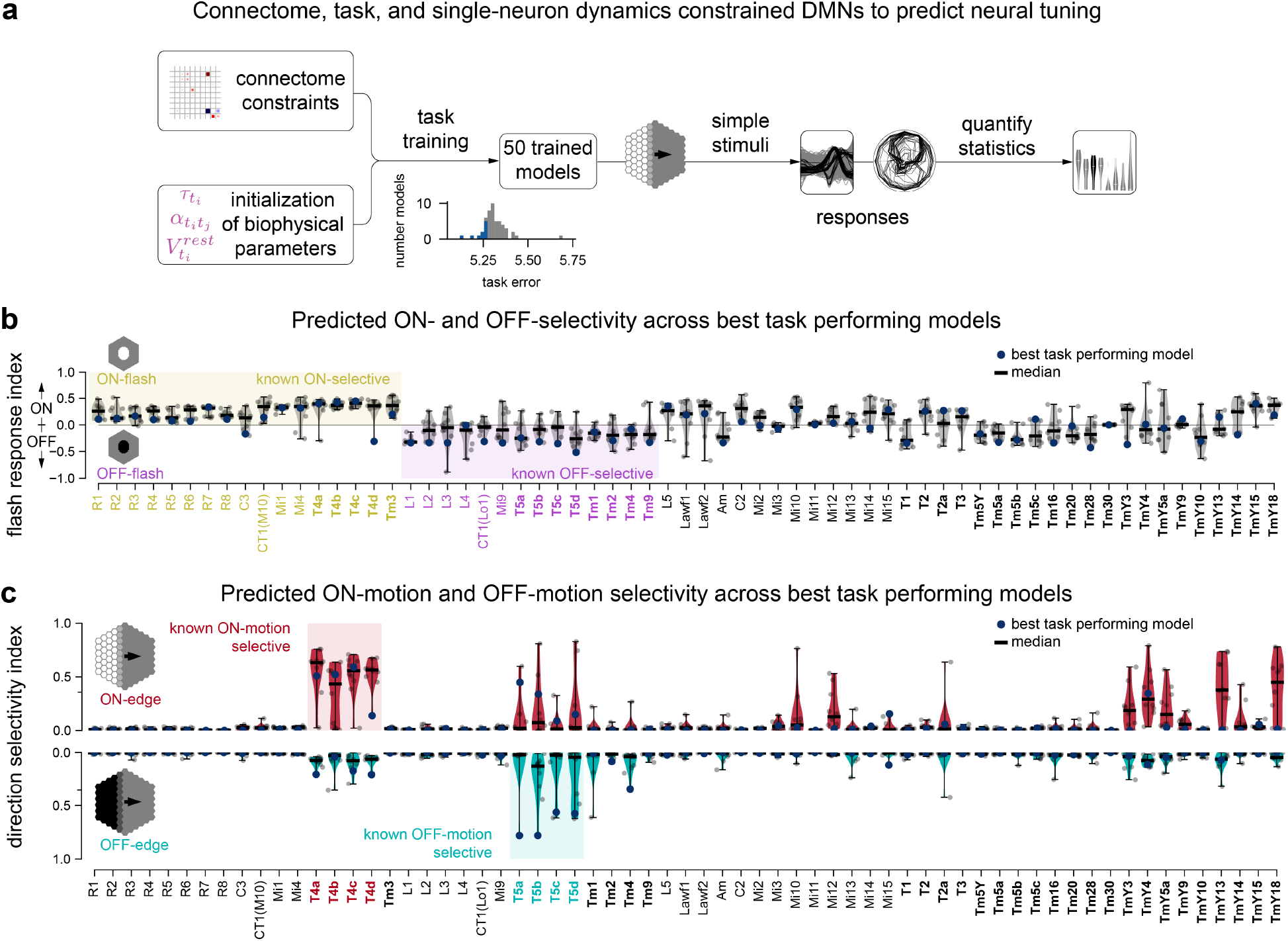
Ensembles of connectome-constrained task-optimized DMNs predict tuning properties. **(a)** We task-optimized 50 connectome-constrained DMNs, yielding different solutions for the biophysical parameters. Inset: Distribution of task errors across the ensemble, 10 best models in blue. We characterized the responses and tuning properties of model neurons from each cell type to experimentally characterized visual stimuli and compare them to known tuning properties. **(b)** ON- and OFF-contrast selectivity indices for each cell type based on peak transient responses for flash stimuli (Methods, Equation 3) for 10 models with best task performance (10 worst-performing models in Extended Data Fig. 4). Cell types with experimentally determined ON-/OFF-selectivity cell types are colored in yellow and violet respectively. Bold labels indicate cell types which provided input to optic flow decoder during training. Cell type names in black have not yet been experimentally characterized. **(c)** Direction selectivity index (DSI) from neural responses to moving edges (Methods, Equation 4) for the same 10 best models as in (b).

First, neural responses in the fly visual system are known to segregate into ON- and OFF-channels^57^, a hallmark of visual computation across species^98,99^. We probed the contrast preference of each cell type using flash stimuli^59^ and found that the ensemble predicts the segregation into ON- and OFF-pathways with high accuracy: The median flash response index (FRI) across the ensemble predicts the correct ON- and OFF- preferred contrast selectivity for 31 of the 31 cell types for which contrast selectivity has been experimentally established (p = 4.7 × 10^-10^, binomial test; note that in these analyses the M10 and Lo1 terminals of CT1 are treated separately). This is also the case for the best-performing model, which correctly assigns 29 out of 31 cells into the correct tuning category (Fig. 2b, p = 2.3 × 10^-7^, binomial test). Furthermore, the ensemble provides predictions for the remaining 34 cell types, and consistency across the ensemble provides a measure of confidence in the predictions (Fig. 2b).

Second, a major result in fly visual neuroscience has been the identification of the T4 and T5 neurons as directional selective neurons with cardinal direction tuning^62, 64, 100^. We characterised the motion selectivity of neurons by their responses to ON- or OFF-edges moving in 12 different directions. We found that the ensemble of models correctly predicts that T4 neurons are ON-motion selective, and T5 neurons are OFF- motion selective (Fig. 2c). The ensemble also correctly predicts the lack of motion tuning in the input neurons to T4 and T5 motion detector neurons (Mi1, Tm3, Mi4, Mi9, Tm1, Tm2, Tm4, Tm9, CT1; see Methods, Supplemental Data).

Our models also suggest the possibility that the transmedullary cell types, TmY3, TmY4, TmY5a, TmY13, and TmY18, might be tuned to ON-motion. We asked if our model predicted motion selectivity for all cell types with asymmetric, multicolumnar inputs, as this is a necessary connectivity motif for motion computation. Based on their local spatial connectivity profiles, we estimated that 19 cell types receive asymmetric, multi-columnar inputs (Methods), but found that only 12 are predicted to be motion selective (p < 0.05, binomial test; Methods) by the ensemble (SI, Supplemental Data). This suggests that our model integrates connectivity across the entire network, rather than simply focusing on local connectivity to determine which neurons are most likely to be motion selective. Next, we asked how well these tunings are predicted by models with random parameter configurations in comparison. We found that task-optimized models predict the known direction selectivity indices and flash response indices more accurately than random models (p =2 × 10^-11^ and p =5.3 × 10^-5^ respectively, Mann-Whitney-U test, Extended Data Fig. 5a and b).

Finally, we found that models which exhibited lower task error (Methods) also had more realistic tuning: Models with higher task performance predict the direction selectivity index of T4 and T5 cells and their inputs better (p = 5.6 × 10^-6^, Wald test, Extended Data Fig. 4b and 6).

### Best task performing model accurately predicts tuning properties of T4 and T5 neurons and their inputs

Our DMN modelling approach enables a large number of model-based analyses which can illuminate the mechanistic basis of computation in a circuit, as well as suggest novel visual stimuli for experimental characterization. We illustrate these analyses using a single model from the ensemble with the best task performance (Fig. 3), focusing on the well-studied T4 and T5 neurons. A more comprehensive set of these analyses for every cell type and every model in the ensemble can be found in the Supplement.

**Figure 3:**
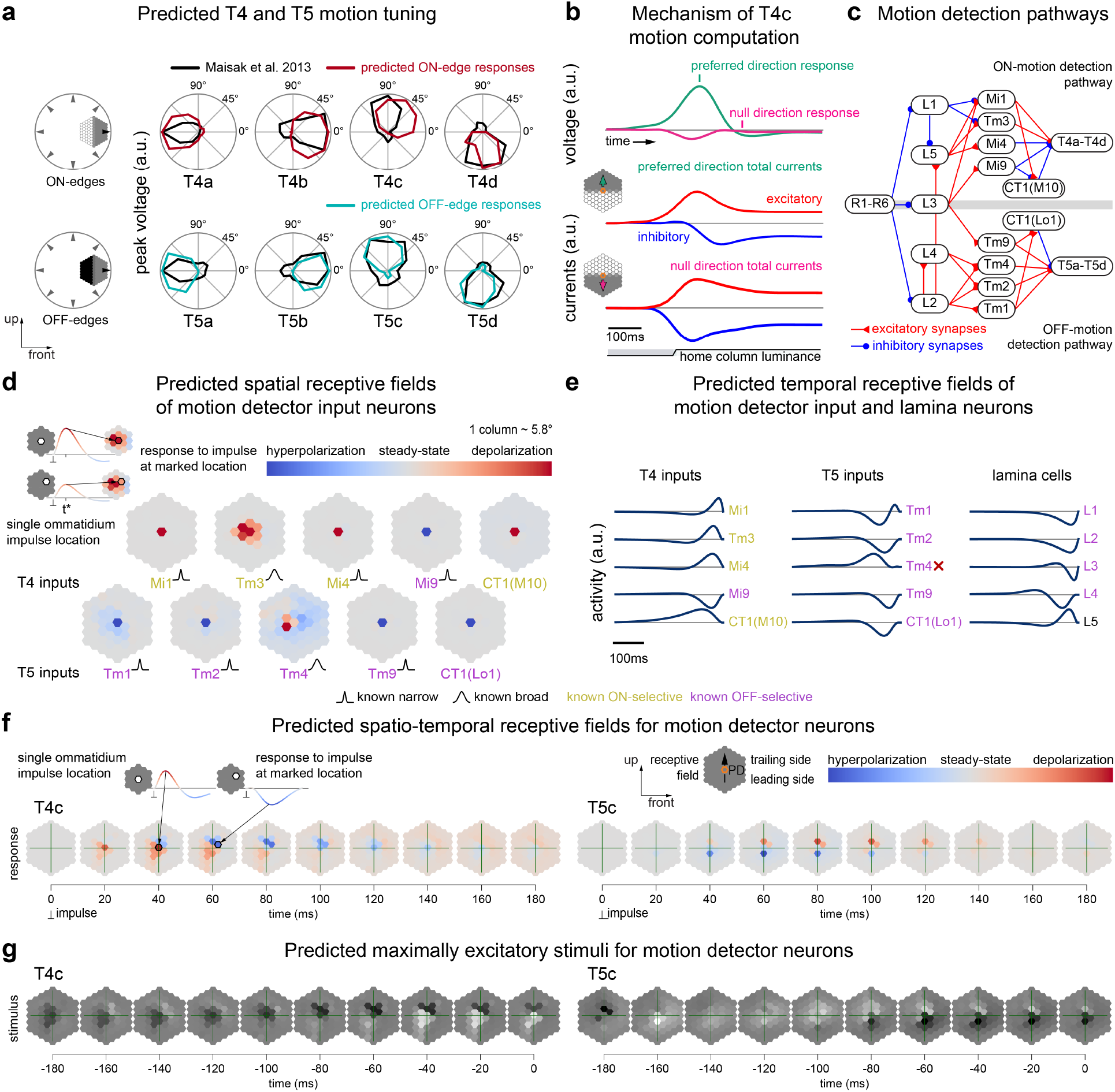
DMN with the best task error largely recapitulates known mechanism of motion computation. **(a)** Responses to moving edges in different directions for T4 and T5 subtypes from the DMN with best task per-formance. Predictions of both direction and polarity of tuning agree with experimental measurements^62, 64, 100^ (Nullcontrasts in Extended Data Fig. 8). **(b)** Fast excitation and delayed, offset inhibition enable T4c to detect motion, in agreement with experimental measure- ments^54^. An edge moving in the preferred direction elicits fast excitatory input currents (red) and delayed inhibitory input currents (blue) to T4c, leading to large depolarization (green). In contrast, an edge moving in the null direction elicits simultaneous arrival of excitatory and inhibitory inputs to T4c, leading to a null response (magenta). **(c)** Major cell types and connectivity in the ON- (T4) and OFF- (T5) motion detection pathways (simplified). **(d)** Spatial receptive fields of major motion detector input neurons revealed by single-ommatidium flashes are in agreement with experimental measurements^70,93^. For major T4 inputs, the best-performing DMN correctly predicts narrow spatial receptive fields for Mi1, Mi4, Mi9, and CT1(M10), and a wide receptive field for Tm3. Mi1, Tm3, Mi4, and CT1(M10) respond with depolarization to the ON-impulses, Mi9 responds with hyperpolarization. For major T5 inputs, the DMN correctly predicts narrow spatial receptive fields for Tm1, Tm2, Tm9, and CT1(Lo1), and a wide receptive field for Tm4. Tm1, Tm2, Tm4, Tm9, and CT1(Lo1) respond with hyperpolarization. Depending on the input ommatidium, Tm4 also responds with depolarization. **(e)** Temporal receptive fields for inputs also are in agreement with experimental measurements^60,93^, with the exception of Tm4 (red cross). For major T4 inputs, in this DMN, Mi1, Tm3, and Mi4 respond with transient depolarization. In contrast, CT1(M10) responds with a longer sustained depolarization to a central ON-impulse. Mi9 hyperpolarizes. For major T5 inputs, in the model, Tm2, Tm9, and CT1(Lo1) respond with transient hyperpolarization. Tm1 depolarizes fast followed by strong hyperpolarization and Tm4 is incorrectly predicted to depolarize (red cross). For lamina cell types, this DMN predicts hyperpolarization in L1, L2, L3, and L4 and depolarization in L5 in response to a central ON-impulse. **(f)** Spatiotemporal receptive fields for motion detector neurons agree with experimental measurements^54^. Receptive field orientation of the motion detectors T4c and T5c align with their preferred motion axis. ON-impulses on the leading side of the receptive field quickly cause the T4 cell to depolarize, while ON-impulses on the trailing side of the receptive field cause slower hyperpolarization in the T4 cell. Conversely, for T5, ON-impulses on the leading side of the receptive field quickly cause the T5 cell to hyperpolarize, while ON-impulses on the trailing side of the receptive field cause slower depolarization in the T5 cell (Receptive fields for OFF-impulses in Extended Data Fig. 7a). **(g)** Video sequence predicted to elicit the strongest responses in T4c and T5c cells. A central OFF-disc followed by an ON-edge moving upwards elicits the strongest response in a T4c cell. A central ON-disc followed by an OFF-edge moving upwards elicits the strongest response in the central column T5c cell. We regularized the full field naturalistic stimuli to show only the content of the input that the cell is responsive to (Full field naturalistic and artificial maximally excitatory stimuli in Extended Data Fig. 7b).

First, we found that in the best task performing model, the four subtypes of the T4 respond strongly to dark edges, and the four subtypes of the T5 neurons to bright edges, moving in the four cardinal directions, in agreement with previous experimental findings^54,55,62,64^ (Fig. 3a and Extended Data Fig. 8).

Second, we probed the mechanism of direction selectivity in T4 neurons (Fig. 3b). Examining the input currents to a single T4c neuron, we found that edges of the preferred contrast moving in the preferred direction of T4c cells elicit large responses through fast excitation and delayed inhibition, in agreement with experimental findings^54^. Conversely, edges moving to the null direction of T4c cells elicit no (or very small) responses because inhibition cancels excitation.

Third, we computed and compared the spatial and temporal receptive fields of the major columnar input neurons to T4 and T5 neurons. These input neurons have been the focus of multiple experimental studies of the motion detection pathways^53,60,70,91,93,94,101^ (Fig. 3c). We characterized these receptive fields in our model by computing the spatial (Fig. 3d) and temporal (Fig. 3e) impulse responses to brief single ommatidium flashes (5 ms duration, Methods). We find that the model correctly predicts the spatial scale of the spatial receptive field for all major inputs to T4 and T5 neurons, and correctly predicts the preferred contrast for all but one of the major inputs to T4 and T5 neurons.

In agreement with experimental findings^70,93^, we find that Tm3 and Tm4 have broad spatial receptive fields (two column radius, 11.6 °), while Mi1, Mi4, Mi9, Tm1, Tm2, Tm9, and CT1 compartments in both medulla and lobula have narrow spatial receptive fields (single column radius, 5.8 °). Further, the model accurately predicts the ON- vs OFF-contrast selectivity of neurons involved in motion detection, including lamina monopolar cells (L1-L5), which anatomically separate the ON- and OFF-pathways^57^, and the direct input neurons to T4 and T5. These cells either depolarize (ON-selective) or hyperpolarize (OFF-selective) in response to light increment flashes. These temporal response properties are correctly predicted for all except the Tm4 cell in this model, which is incorrectly predicted to be ON-selective by its temporal receptive field.

For the motion selective T4 and T5 neurons, the spatiotemporal receptive fields are not separable in space and time. We characterized the full spatiotemporal receptive field for T4c and T5c neurons (Fig. 3f). ON-impulses on the leading side of the receptive field of the ON-contrast, upwards direction selective T4c cell lead to its fast depolarization, whereas impulses on the trailing side of the receptive field lead to a delayed hyperpolarization, again matching experimental findings^54^. ON-impulses on the leading side of the receptive field of the OFF-contrast, upwards direction selective T5c cell lead to its fast hyperpolarization, whereas impulses on the trailing side of its receptive field lead to a delayed depolarization. Because T5c is OFF-selective, its OFF-impulse responses are inverted, resembling the T4c spatiotemporal receptive field (Extended Data Fig. 7a). This suggests that in our model, T5c implements a similar motion tuning mechanism to OFF-edges as T4c to ON-edges. Again, this mechanism also agrees with experimental findings^55^.

Finally, we show that the model can be used to design optimized stimuli: We used the model to screen for video sequences from the Sintel dataset that elicited the largest responses in the motion selective neurons (Fig. 3g; Methods). One might expect that pure ON- or OFF-stimuli would elicit the largest responses in T4 and T5, respectively. However, we find both ON-and OFF-elements in optimized stimuli, suggesting an interplay between ON- and OFF-pathways. We found that the stimulus that elicits the strongest response in the T4c cell is a central OFF-disc followed by an ON-edge moving upwards, matching the preferred direction of the cell. Similarly, for the T5c cell, the stimulus that elicits the strongest response is a central ON-disc followed by an OFF-edge moving upwards in the preferred direction of the cell (Extended Data Fig. 7b for corresponding full-field naturalistic stimuli, numerically optimized stimuli, and preferred moving edge stimuli). Taken together, this individual model predicts a large number of tuning properties for the T4 and T5 cells and their inputs.

### Neural responses across the model ensemble cluster strongly for many cell types

The predictions in the previous section were derived from a single model. How similar or dissimilar are the predictions of different task-optimized models constrained with the same connectome? To address this question, for each cell type, we simulated the neural activity of a single neuron, in response to naturalistic video sequences from the Sintel dataset. We then used UMAP^102^ to perform nonlinear dimensionality reduction on high-dimensional activity vectors of that neuron across the model ensemble, and clustered the models in the resulting 2D projections (Fig. 4a, see Methods, Supplement). For many cell types, we found that models predict strongly clustered neural responses. For T4c neurons for example, we found three clusters corresponding to qualitatively distinct responses of this cell type for naturalistic inputs: Two clusters contain models with direction selective T4c cells (Fig. 4b and c) with up- and down-selective cardinal tuning, respectively, whereas neurons in the third cluster are not direction tuned. The direction selective cluster with the (correct) upward preference has lowest average task error (circular marker, average task error 5.297) and contains the best task performing model analyzed previously (shown in Fig. 3), followed by the cluster with opposite preference (triangular marker, average task error 5.316). The non-selective cluster has the worst performance (square marker, average task error 5.357), suggesting that models with accurate tuning correlate with lower task error (see also Extended Data Fig. 6).

**Figure 4:**
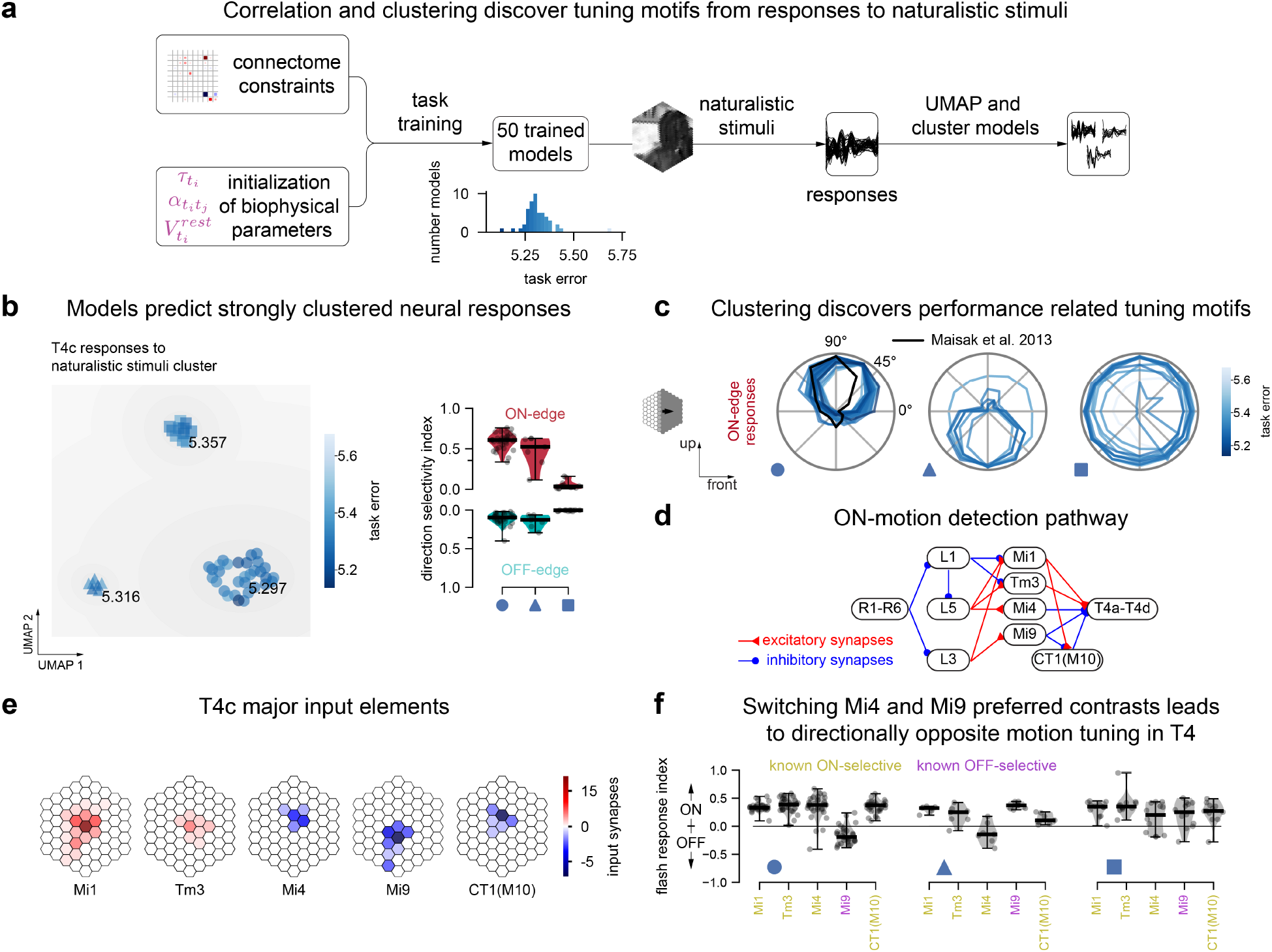
Cluster analysis of DMN ensembles enables hypotheses generation and suggests experimental tests. **(a)** We clustered 50 DMNs after performing nonlinear dimensionality reduction of their predicted responses to naturalistic scenes for each cell type (Inset: Distribution of task performance across models). We compared their tuning to simple stimuli to identify whether clusters correspond to qualitatively different tuning mechanisms. **(b)** Responses of T4c cells to naturalistic scenes reveal three distinct clusters. T4c cells in DMNs in the first and second clusters (circular and triangular marker) are ON-motion direction selective, whereas those in the third cluster (square marker) are not. **(c)** The DMNs reveal three distinct solutions for the T4c cells (which are known to be tuned to upwards ON-motion): (1) upwards tuning (cluster with lowest average task error of 5.297, circular marker in panel b), (2) downwards tuning (5.316 average error, triangular marker), or (3) no motion tuning to ON-edges (5.357 average error, square marker). **(d)** ON-motion detection pathway from Fig 3c. **(e)** Connectivity of major input elements to T4c. Blue and red represent putative hyper- and depolarizing inputs. Saturation represents average number of input synapses for each offset location in the T4c dendrite (see Fig. 1e). **(f)** Tuning properties within each cluster reveal dependencies between T4 tuning and that of Mi4 and Mi9 cells in the ensemble: Switching Mi4 (known ON-contrast selective) and Mi9 (known OFF-contrast selective) contrast preferences results in directionally opposite motion tuning solutions in T4. Cluster 1 (T4c in DMN upwards tuned, circle) indicates ON-selectivity for Mi1, Tm3, Mi4, and CT1(M10), and OFF-selectivity for Mi9. For ON-motion stimuli, in these DMNs T4c receives central depolarizing input from Mi1 and Tm3 and dorsal hyperpolarizing input from Mi4 and CT1(M10). For cluster 2 (T4c in DMN downwards tuned, triangle), Mi1, Tm3, Mi9, and CT1(M10) are ON-selective and Mi4 is OFF-selective. For ON-motion stimuli, in these DMNs, T4c receives central depolarizing input from Mi1 and Tm3, ventral hyperpolarizing input from Mi9 and dorsal hyperpolarizing input from CT1(M10). For cluster 3 (T4c in DMN not tuned, square), all major input elements are ON-selective. For ON-motion stimuli, in these DMNs, T4c receives central depolarizing input from Mi1 and Tm3, dorsal hyperpolarizing input from Mi4 and CT1(M10), and ventral hyperpolarizing input from Mi9.

What are the circuit mechanisms in the ON-motion detection pathway (Fig. 4d) underlying tuning computations in the different clusters? We found that direction selectivity in the two tuned clusters is associated with opposite preferred contrast tuning of Mi4 and Mi9 neurons which provide direct flanking inhibitory input to T4 neurons (Fig. 4e). Models with the correct direction selectivity for T4 neurons also predict the correct contrast selectivity for Mi4 and M9 neurons, and vice versa (Fig. 4f).

This shows how the space of task-optimized and connectome-constrained models can be used to provide hypotheses about different circuit mechanisms which might underlie the tuning properties of individual cells. Conversely, it shows that experimentally measuring the tuning of one neuron automatically translates to constraints on other neurons in the circuit. Here, ‘clamping’ the T4c neurons to their measured tuning properties (by only selecting models from the correct cluster) is sufficient to correctly constrain the tuning of both Mi4 and Mi9 neurons.

### Models predict motion tuning for TmY3

Amongst models with the best task performance, TmY3, TmY4, and TmY18 are often ON-motion selective (Fig. 2c). As these neurons have yet to be experimentally characterized, we analyzed these prediction in our models. Since TmY3 neurons do not receive inputs from other known motion selective neurons, we were intrigued by the possibility that it might directly compute a motion signal and possibly constitute a parallel pathway to the well-known T4 and T5 neurons. In contrast, TmY4 and TmY18 cells receive inputs from T4 cells, potentially inheriting their motion tuning.

In the model ensemble, we found four distinct clusters for TmY3 (Fig. 5a). In the best-performing cluster (circular marker) TmY3 responds to ON-edges from front to back or downwards (Fig. 5b). In contrast, in the second cluster (triangular marker), TmY3 is not direction selective. In the third cluster (square marker) TmY3 is direction selective to ON-edges moving from the back to the front. In the fourth cluster (star marker), TmY3 is, again, not direction selective. Together, the ensemble suggests ON-motion sensitivity for TmY3, but different clusters disagree in their predictions for direction and contrast selectivity.

**Figure 5:**
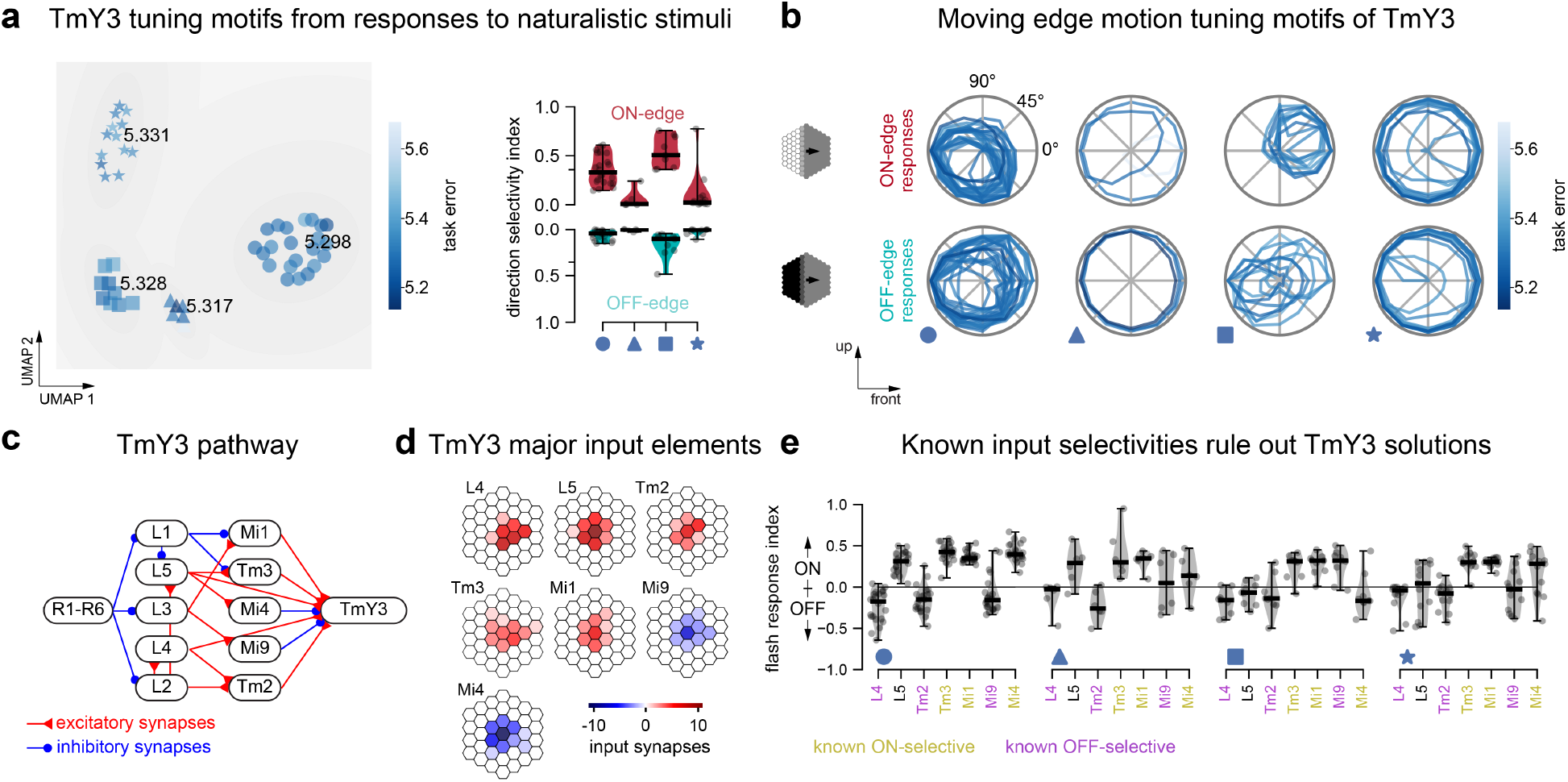
DMNs suggest that TmY3 neurons compute motion independently of T4 and T5 neurons. **(a)** Dimensionality reduction on TmY3 responses to naturalistic stimuli reveals 4 clusters of DMNs with average task errors 5.298 (circle), 5.317 (triangle), 5.328 (square) and 5.331 (star). Across clusters, TmY3 shows different strengths of direction selectivity (evaluated with moving edge stimuli). ON-edge direction selectivity is strong in the first and the third cluster. **(b)** Normalized peak responses of TmY3 to moving edge stimuli in the DMNs of each cluster. **(c)** Major cell types and synaptic connections in the pathway that projects onto TmY3 (simplified). **(d)** The input elements of TmY3 with the highest amount of synapses are L4, L5, Tm2, Tm3, Mi1, Mi9, and Mi4. The asymmetries of their projective fields could allow TmY3 to become motion selective. **(e)** Dependencies between TmY3 tuning and the contrast preference of its input cells. For clusters in which TmY3 is motion selective, cluster 1 (TmY3 tuning to downwards/front-to-back motion, circular marker) indicates ON- selectivity for Tm3, Mi1, and Mi4 cells, and OFF-selectivity for L4, Tm2, and Mi9 cells, in agreement to the known selectivities. In contrast, cluster 3 (TmY3 tuning to upwards/back-to-front motion, square marker) indicates ON- selectivity for Mi9 in contradiction to the known selectivities and hence ruling out the third TmY3 tuning solution.

In our connectome data, the strongest input elements of TmY3 by number of synapses are L4, L5, Tm2, Tm3, Mi1, Mi9, and Mi4 (Fig. 5c and d). While none of these input neurons are motion-selective, the asymmetries in their connectivity to TmY3 might allow it to detect motion. We asked if we could better constrain our predictions by asking which clusters also predicted the correct preferred contrast for these input neurons. We found that the first model cluster (Fig. 5a, circular marker), in which TmY3 is tuned to front-back or downards motion, most accurately captures the known contrast selectivity of all TmY3 input cells (Fig. 5e). In contrast, all three other clusters fail to consistently capture the OFF-selectivity of Mi9. Thus our model proposes TmY3 as a novel candidate motion detector independent of the well-known T4 and T5 motion pathways.

### Sparse synaptic connectivity enables accurate prediction of neural responses from synthetic connectomes

We sought to understand the conditions under which connectomic measurements might strongly constrain models of neural computation. Universal function approximation theorems for artificial neural networks^26,103^ suggest that a single general-purpose connectivity can underlie many possible computations. Empirically, different deep neural network architectures trained to solve the same task have very different selectivity at the level of single units^38,82,104–106^, even when they share population-level representations^30,107,108^. We hypothesized that while general-purpose neural architectures used in machine learning have a many-to-many relationship between the computational task and neural connectivity, biological neural circuits might have a tighter relationship with their computational task due to their sparse structured connectivity, and therefore connectomic measurements and task-constraints in such circuits would strongly constrain mechanistic computational models.

We investigated this counter-factual using synthetic feedforward networks designed to perform a classic handwritten digit recognition network task. We constructed a variety of synthetic networks which perform the same task, but with different degrees of sparsity in their connectivity (Fig. 6). While the synthetic networks all perform the same task with near perfect accuracy, they use dramatically different connectivity, and so use different mechanisms to perform the same task. We then used connectomic measurements from each synthetic network to construct a corresponding connectome-constrained model optimized to perform the same task (Fig. 6a). We then compared the neural responses of hidden neurons in each synthetic network to corresponding neurons in the corresponding task and connectome-constrained model. We considered two settings, one where the connectomic measurements only indicated whether a pair of neurons were connected but not their strength, and one where the measurements indicated both the connectivity and a noisy estimate of the strength of the connection. In our model of the fly visual system, we have connectomic measurements of the synapse count for each connection, which can inform the relative strength of a connection, but not the absolute strength. This constitutes an intermediate regime between known strength and unknown strength.

**Figure 6:**
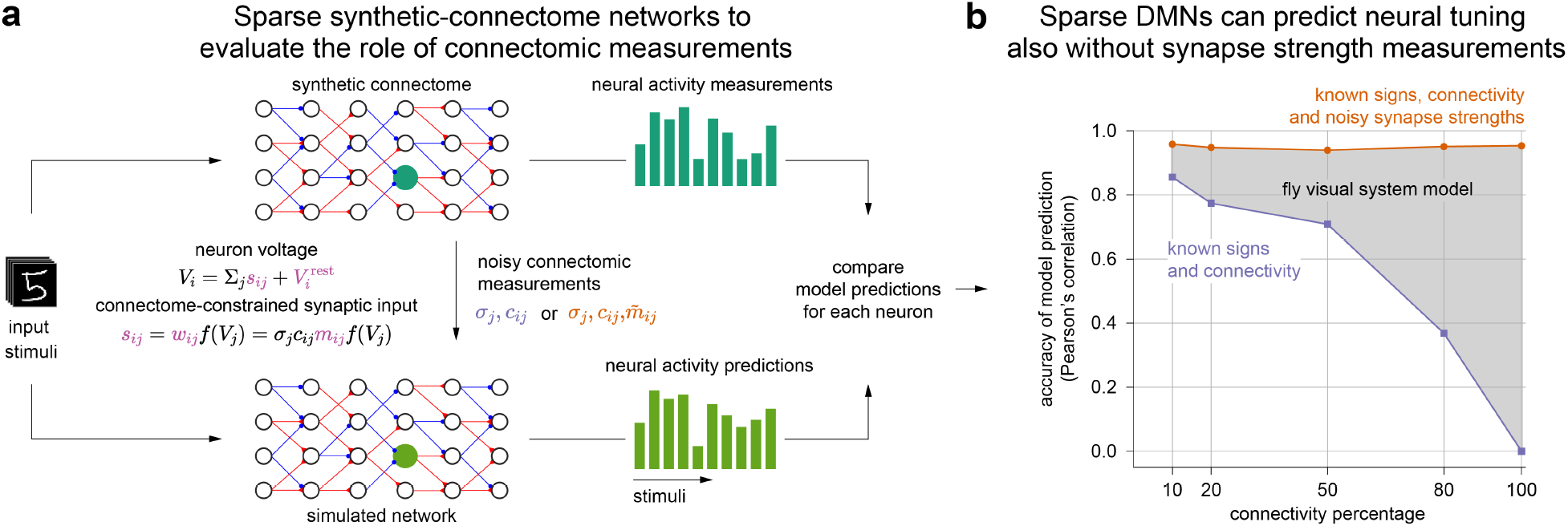
Connectomic measurements can strongly constrain neural networks in circuits with sparse connectivity. **(a)** We constructed synthetic-connectome networks for classifying hand-written digits with varying degrees of sparse connectivity. For each synthetic-connectome network, we simulated connectomic measurements and constructed a connectome-constrained and task-optimized simulated DMN (Methods). We measured the correlation of the neural responses across stimuli of the same neuron in the synthetic-connectome network (dark green) and the fitted neural network (light green). **(b)** Median neural response correlation coefficient from 100 randomly-sampled neuron pairs from each layer and across 25 network pairs. Simulations constrained only with connectivity measurements only correlate well for low connectivities (dark blue), while simulations constrained by measurements of both connectivity and connection strength correlate well across all average connectivity percentages (orange). The fly visual system likely falls in the region between the two curves, since measured synapse counts inform relative connection strengths between pairs of neurons for the same pair of cell types, but not absolute connection strength.

We found that densely connected synthetic networks, where each neuron in a given layer connects to every neuron in the next layer, are essentially uncorrelated with their task- and connectome-constrained models constructed with known connectivity but unknown strength (Fig. 6b, Extended Data Fig. 9a). In contrast, sparsely connected synthetic networks, where each neuron only connects to a small percentage of neurons, are well correlated with their task and connectome-constrained models (median Pearson’s correlation of 0.856 for networks with 10% average connectivity). The correlation between synthetic networks and their simulations drops smoothly as the connection density increases. For connectome-constrained models constructed with known connectivity and a noisy estimate of connection strength, we found that connectivity sparseness is not required for models to correlate with their corresponding synthetic network (Fig. 6b, Extended Data Fig. 9b).

## Discussion

Here we constructed a neural network with neural connectivity measured at the microscopic scale. We also required that at the macroscopic scale, the collective neural activity dynamics across the entire network result in an ethologically relevant computation. The combination of microscopic and macroscopic constraints enabled us to construct a large-scale computational model spanning many tens of cell types and tens of thousands of neurons. We showed that such large-scale mechanistic models could accurately make detailed predictions of the neural responses of individual neurons to dynamic visual stimuli, revealing the mechanisms by which computations are performed. Knowledge of the connectome played a critical role in this success, in part by leading to a massive reduction in the number of free model parameters.

We have taken a reductionist modeling approach to emphasize the important role played by the connectivity of a neural network. We found that for the motion pathways of the fruit fly visual system, this model correctly predicts many aspects of visual selectivity. However our reductionist model cannot, for example, account for the role played in this circuit by electrical synapses^109^, complex chemical synapse dynamics^96^, and neuromodulation^110^. Richer models of neurons, synapses, and extra-synaptic modulation will be essential to correctly predict these effects. Further, we only considered the role of this circuit in detecting motion, which is but one of many computations performed by the visual system^14,111–114^.

Our study provides a direct link between artificial neural networks and biological circuit models. In our task-optimized deep network, every model neuron and synapse corresponds to a real neuron and synapse in the brain. This correspondence enables detailed experimentally testable predictions for each neuron. In contrast, most previous studies using task-optimized deep neural networks have predominantly focused on modeling computations without such detailed connectomic measurements: in fly motion vision path- ways^115,116^ and olfactory system^36^, in fish oculomotor system^117^ in mammalian visual pathways^44,76,106,118^ and prefrontal cortex^45^. More recently, there has been tantalizing early evidence, in small scale models^46^ and larger models^119^ of the fly motion pathways, that our approach to combining connectome and task-constraints in a single model might prove successful.

Our modeling approach provides a discovery tool, aimed at using connectomic measurements to generate detailed, experimentally testable hypotheses for the computational role of individual neurons. Measurements of neural activity are necessarily sparse and involve difficult trade-offs. Activity can frequently only be measured in a limited number of contexts, and for either a limited number of neurons or for a larger number of neurons with poorer temporal resolution. Connectome-constrained DMN models generate meaningful predictions even in the complete absence of neural activity measurements, but can be further constrained by sparse measurements of neural activity as we showed (Fig. 4), or even be directly optimized to match measured neural activity^120^.

Whole nervous system connectome projects are nearing completion for the adult fruit fl^y2,13^, and whole mouse brain connectome projects are now being discussed^121^. Large-scale whole nervous system models^120^ will be of critical importance for integrating connectomic, transcriptomic, neural activity and animal behavior measurements across labs, scales, and the nervous system^2^. Further, with the recent development of detailed biomechanical body models for the fruit fly^122^ and rodent^123^, we can now contemplate constructing whole animal models spanning brain and body.

## Supporting information

Supplemental Cell Profiles

## Author Contributions

Conceptualization, Methodology: JKL, FDT, JHM, SCT. Data curation: JKL, FDT, AN, KS, S-yT. Software and Investigation: JKL, MM, SP, FDT. Analysis: JKL, EG, AN, SCT. Writing: JKL, JHM, SCT. Writing (Review & Editing): EG, AN, KS, MM, SP, FDT. Supervision and funding: SCT, JHM.

## Acknowledgements

We are grateful to Lou Scheffer and Lowell Umayam for assistance with accessing connectomic reconstructions. We thank Axel Borst, James Fitzgerald, Nathan Klapoetke, Gerry Rubin, Michael Reiser, and Karel Svoboda for valuable discussions. We thank James Fitzgerald, David Stern, Nathan Klapoetke, Albert Lee, Richard Gao, Jakob Voigts, Brett Mensh for valuable feedback on the manuscript. We thank Tory Herman for sharing the colorization of the optic lobe figure^65^ (Fig. 1b).

This article is subject to HHMI’s Open Access to Publications policy. HHMI lab heads have previously granted a nonexclusive CC BY 4.0 license to the public and a sublicensable license to HHMI in their research articles. Pursuant to those licenses, the author-accepted manuscript of this article can be made freely available under a CC BY 4.0 license immediately upon publication.

## Methods

### Construction of spatially invariant connectome from local reconstructions

We built a computational model of the fly visual system which is consistent with available connectomic data^48–50,68,124,125^, which has biophysically plausible neural dynamics, and which can be computationally trained to solve an ethiologically relevant behavioural task, namely estimation of optic flow. To achieve this, we developed algorithms to blend annotations from two separate data-sets by transforming, sanitizing, combining and pruning the raw data sets into a coherent connectome spanning all neuropils of the optic lobe (Supplementary Note 1).

The original data stems from focused ion beam scanning EM datasets (FIBSEM) from the FlyEM project at Janelia Research Campus. The FIB-25 dataset volume comprises seven medulla columns and the FIB-19 dataset volume comprises the entire optic lobe and, in particular, detailed connectivity information for inputs to both the T4 and T5 pathways^48,49,68^. The data available to us consisted of 1801 neurons, 702 neurons from FIB-25 and 1099 neurons from FIB-19. For about 830 neurons the visual column was known from hand annotation. These served as reference positions. Of the 830 reference positions, 722 belong to neuron types selected for simulation. None of the T5 cells, whose directional selectivity we aimed to elucidate, were annotated. We therefore built an automated, probabilistic expectation maximization algorithm that takes synaptic connection statistics, projected synapse center-of-mass clusters and existing column annotations into account. We verified the quality of our reconstruction as described in Supplementary Note 1 Only the neurons consistently annotated with both 100% and 90% of reference positions used were counted to estimate the number of synapses between cell types and columns, in order to prune neuron offsets with low confidences.

Synaptic signs for most cell types were predicted based on known expression of neurotransmitter markers (primarily the cell type specific transcriptomics data from Davis et al 2020). For a minority of cell types included in the model, no experimental data on transmitter phenotypes were available. For these neurons, we used guesses of plausible transmitter phenotypes. To derive predicted synaptic signs from transmitter phenotypes, we assigned the output of histaminergic, GABAergic and glutamatergic neurons as hyperpolarizing and the output of cholinergic neurons as depolarizing. In a few cases, we further modified these predictions based on distinct known patterns of neurotransmitter receptor expression (see Davis et al. for details). For example, output from R8 photoreceptor neurons, predicted to release both acetylcholine and histamine, was treated as hyperpolarizing or depolarizing, respectively, depending on whether a target cell type is known to express the histamine receptor ort (a histamine-gated chloride channel).

### Representing the model as a hexagonal convolutional neural network

Our end-to-end differentiable^126^ DMN model of the fly visual system can be interpreted as a continuous-time neural ordinary differential equation (neural ODE)^127^ with a deep convolutional recurrent neural network (convRNN)^128^ architecture that is trained to perform a computer vision task using backpropagation through time (BPTT)^41,79^. Our goal was to optimize a simulation of the fly visual system to perform a complex visual information processing task using optimization methods from deep learning. One hallmark of visual systems that has been widely exploited in such tasks are their convolutional nature^129–132^, i.e. the fact that the same computations are applied to each pixel of the visual input. To model the hexagonal arrangement of photoreceptors in the fly retina, we developed a hexagonal convolutational neural network in the widely used deep learning framework Pytorch^40^, which we used for simulation and optimization of the model.

### Neuronal Dynamics

In detail, we simulated point neurons with voltages *V_i_* of a postsynaptic neuron i, belonging to cell type *t_i_* using threshold-linear dynamics, mathematically equivalent to commonly used formulations of firing-rate models^133^

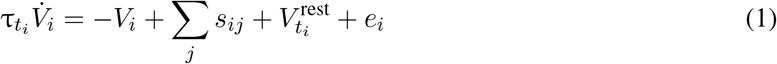

Neurons of the same cell type share time constants, *τ_t_i__*, and resting potentials, 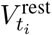. Dynamic visual stimuli were delivered as external input currents *e_i_* to the photoreceptor (R1-R8), for all other cell types, *e_i_* = 0. In our model, instantaneous graded synaptic release from presynaptic neuron *j* to postsynaptic neuron *i* is described by

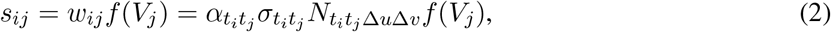

comprising the anatomical filters in terms of the synapse count from EM-reconstruction, 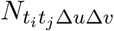, at the offset location Δ*u* = *u_i_* – *u_j_* and Δ*v* = *v_i_* – *v_j_* in the hexagonal lattice between two types of cells, *t_i_* ar *t_j_*, and further characterised by a sign, *σ_t_i_t_j__* ∈ {–1, +1}, and a non-negative scaling factor, *α_t_i_t_j__*.

The synapse model in Equation 2 entails a trainable non-negative scaling factor per filter that is initiaized as

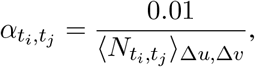

with the denominator describing the average synapse count of the filter. Synapse counts, *N_t_i_t_j__*, and signs, *σ_t_i_t_j__*, from reconstruction and neurotransmitter and receptor profiling were kept fixed. The scaling factor was clamped during training to remain non-negative.

Moreover, at initialization, the resting potentials were sampled from a Gaussian distribution

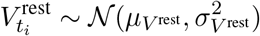

with mean *μ*_*V*_^rest^ = 0.5 (a.u.) and variance 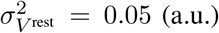. The time constants were initialized at *τ_t_i__* = 50ms. The 50 task-optimized DMNs were initialized with the same parameter values. During training, in Euler integration of the dynamics, we clamped the time constants as *τ_i_* = max(*τ_i_, Δ_t_*), so that they remain above the integration time step *Δ_t_* at all times.

In total, the model comprises 45669 neurons and 1513231 synapses, across two-dimensional hexagonal arrays 31 columns across. Independently of the extent of the two-dimensional hexagonal arrays are the numbers of free parameters: 65 resting potentials, 65 membrane time constants, 604 scaling factors; and connectome determined parameters: 604 signs, and 2355 synapse counts. Thus, the number of free parameters in the visual system model is 734.

In the absence of connectomic measurements, the number of parameters to be estimated is much larger. With *T* = 65 cell types (counting CT1 twice for the compartments in the medulla and lobula) and *C* = 721 cells per type for simplicity, the number of cells in our model would be TC = 46, 865. Assuming an RNN with completely unconstrained connectivity and simple dynamics 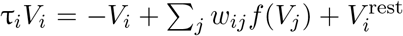 we would have to find (*TC*)^2^ + 2(*TC*) = 2,196, 421, 955 free parameters. Assuming a convolutional RNN with shared filters between cells of the same postsynaptic type, shared time constants, and resting potential, the amount of parameters reduces drastically to *T*^2^*C* + 2*T* = 3, 046, 355. Further assuming the same convolutional RNN but additionally convolutional filters are constrained to *F* = 5 visual columns, i.e. the number of presynaptic input columns in hexagonal lattice is *P* = 3*F*(*F* + 1) + 1, the amount of parameters reduces to *T*^2^*P* + 2*T* = 384, 605. Assuming as in our connectome only *Q* = 604 connections between cell types exist, this reduces the number of parameters further to *Q*P + 2*T* = 55,185. Instead of parametrizing each individual synapse strength, we assume that synapse strength is proportional to synapse count from the connectome times a scalar for each filter, reducing the number of parameters to *Q* + 2*T* = 734 while providing enough capacity for the DMNs to yield realistic tuning to solve the task.

#### Convolutions using scatter and gather operations

For training the network, we compiled the convolutional architecture specified by the connectome and the sign constraints to a graph representation containing (1) a collection of parameter buffers shared across neurons and/or connections, (2) a collection of corresponding index buffers indicating where the parameters relevant to a given neuron or connection can be found in the parameter buffers, and (3) a list of pairs (presynaptic neuron index, postsynaptic neuron index) denoting connectivity. This allowed us to efficiently simulate the network dynamics via Euler integration using a small number of element-wise, scatter, and gather operations at each time step. We found that this is more efficient than using a single convolution operation, or performing a separate convolution for each cell type, since each cell type has its own receptive field - some much larger than others - and the number of cells per type is relatively small.

### Optic flow task

#### Model training

An optic flow field for a video sequence consists of a 2D vector field for each frame. The 2D vector at each pixel represents the magnitude and direction of the apparent local movement of the brightness pattern in an image.

We frame the training objective as a regression task

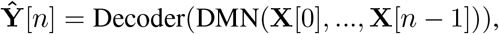

with **Ŷ** the optic flow prediction, and **X** the visual stimulus sequence from the Sintel dataset, both sampled to a regular hexagonal lattice of 721 columns. With the objective to minimize the square error loss between predicted optic flow and target optic flow fields, we jointly optimized the parameters of both the decoder and the visual system network model described above.

In detail, for training the network, we added randomly augmented grey-scaled video sequences from the Sintel dataset sampled to a regular hexagonal lattice of 721 columns to the voltage of the eight photoreceptor cell types (Fig. 1f, Equation 1). We denote a sample from a minibatch of video sequences as **X** ∈ ℝ^*N,C*^ with *N* the number of time steps, and *C* the number of photoreceptor columns. The dynamic range of the input lies between 0 and 1. Input sequences during training entailed 19 consecutive frames drawn randomly from the dataset and resampled to match the integration rate. The original framerate of 24 Hz and 19 frames lead to a simulation of 792ms. We did not find that an integration time step smaller than 20 ms, i.e. a framerate of 50 Hz after resampling, yielded qualitatively superior task performance nor more realistic tuning predictions. We interpolated the target optic flow in time to 50 Hz temporal resolution, instead of resampling it. To increase the amount of training data for better generalization, we augmented input and target sequences as described further below. At the start of each epoch, we computed an initial state of the network’s voltages after 500ms of grey stimuli presentation to initialize the network at a steady state for each minibatch during that epoch. The network integration given input **X** results in simulated sequences of voltages **V** ∈ ∝^*N,T_C_*^ with *T_C_* the total number of cells. The subset of voltages, **V**_out_ ∈ ℝ^*N,D,C*^, of the *D* cell types in the black box in Fig. 1f was passed to a decoding network. For decoding, the voltage was rectified to avoid that the network finds biologically implausible solutions by encoding in negative dynamic ranges. Further, it was mapped to cartesian coordinates to apply Pytorch’s standard spatial convolution layers for decoding and on each timestep independently. In the decoding network, one layer implementing spatial convolution, batch normalization, softplus activation, and dropout, followed by one layer of spatial convolution, transformed the *D* feature maps into the two-dimensional representation of the estimated optic flow, **Ŷ** ∈ ℝ^N,2,C^.

Using stochastic gradient descent with adaptive moment estimation (*β*_1_ = 0.9, *β*_2_ = 0.999, learning rate decreased from 5 × 10^-5^ to 5 × 10^-6^ in ten steps over iterations, batch size of four) and the automatic gradient calculation of the fully differentiable pipeline, we optimized the biophysical parameters through backpropagation through time such that they minimize the L2-norm between the predicted optic flow, **Ŷ**, and the groundtruth optic flow, **Y**:

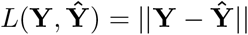

We additionally regularized the shared resting potentials for 150,000 iterations, using stochastic gradient descent without momentum, based on time-averaged responses to naturalistic stimuli of the central column cell of each cell type, *t*_central_, to encourage configurations of resting potentials that lead to nonzero and nonexploding activity in all neurons in the network. We weighted these terms independently with *γ* = 1, encouraging activity greater than *a*, and *δ* = 0.1, encouraging activity less than *a*. We chose *λ_V_* = 0.1 and *a* = 5 in arbitrary units. With *B* being the batch size and *T* the number of all cell types, the regularizer is

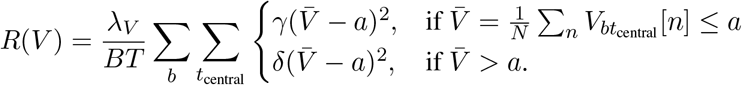

We regularly checkpointed the above error measure *L*(**Y**, **Ŷ**) averaged across a held out validation set of Sintel video clips. Models generalized on optic flow computation after round about 250,000 iterations, yielding functional candidates for our fruit fly visual system models that we analyzed with respect to their tuning properties.

#### Task-optimization dataset

We optimized the network on 23 sequences from the publicly available computer-animated movie Sintel^134^. The sequences have 20-50 frames, at a frame rate of 24 frames per second and a pixel resolution of 1024×436. The dataset provides optical flow in pixel space for each frame after the first of each sequence. Since the integration time steps we use are faster than the actual sampling rate of the sequences, we resample input frames accordingly over time and interpolate the optic flow.

#### Fly-eye rendering

We first transformed the RGB pixel values of the visual stimulus to normalized greyscale between 0 and 1. We translated cartesian frames into receptor activations by placing simulated photoreceptors in a two-dimensional hexagonal array in pixel space, 31 columns across resulting in 721 columns in total, spaced 13 pixels apart. The transduced luminance at each photoreceptor is the greyscale mean value in the 13 × 13-pixel region surrounding it.

#### Augmentation

We used (1) random flips of input and target across one of the three principal axes of the hexagonal lattice, (2) random rotation of input and target around its six-fold rotation axis, (3) adding element-wise Gaussian noise with mean zero and variance *σ_n_* = 0.08 to the input *X* (then clamped at 0) (4) random adjustments of contrasts, log 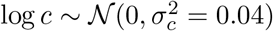, and brightness, 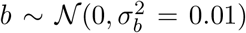, of the input with *X*′ = *c*(*X* – 0.5) + 0.5 + cb.

In addition, we strided fly-eye rendering across the rectangular raw frames in width, subsampling multiple scenes from one. We ensured that such subsamples from the same scene do not distribute across training and validation sets. Input sequences in chunks of 19 consecutive frames were drawn randomly in time from the full sequences.

#### Black-box decoding network

The decoding network is feedforward, convolutional and has no temporal structure. Aspects of the architecture are explained in the paragraph Model training. The spatial convolutions have a filter size of 5 × 5. The first layer transforms the D = 34 feature maps to an eight-channel intermediate representation, that is further translated by an additional convolutional layer to a three-channel intermediate representation of optic flow. The third channel is used as shared normalization of each coordinate of the remaining two-dimensional flow prediction. The decoder uses Pytorch-native implementations for two-dimensional convolutions, batch normalization, softplus activation, and dropout. We initialized its filter weights homogeneously at 0.001.

### Model characterization

#### Task error

To rank models based on their task performance, we computed the standard optic flow metric of average end-to-end point error (EPE)^135^ which calculates the average over all timesteps and pixels (i.e. here columns) of the error

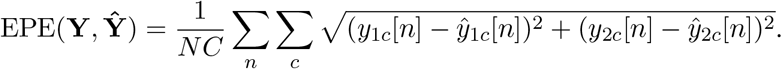

between predicted optic flow and groundtruth optic flow, and averaged across the held out validation set of Sintel sequences.

#### Unconstrained CNN

We trained unconstrained, fully convolutional neural networks on the same dataset and task yielding a lower bound for the task error of the DMN. Optic flow was predicted by the CNN from two consecutive frames

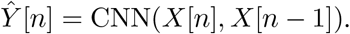

with the original frame rate of the Sintel movie. We chose 5 layers for the CNN with 32, 92, 136, 8, 2 channels respectively and kernel size 5 for all but the first layer which kernel size is 1. Each layer performs a convolution, batch normalization, and ELU activation except the last layer which only performs a convolution. We optimized an ensemble of 5 unconstrained CNNs with 414,666 free parameters each using the same loss function, *L*(*Y, Ŷ*), as for the DMN. We used the same dataset, i.e. hexagonal sequences and augmentations from Sintel, for training and validating the CNN as for training and validating the DMN, allowed by two custom modules mapping from hexagonal lattice to cartesian map and back.

#### Random DMNs

We created 50 DMNs with random parameters (all sampled from Gaussians of different means and standard deviations), and task-optimized only their decoding network, yielding an upper bound for the task error of the task-optimized DMN and a lower bound for the accuracy of tuning predictions without task-optimization.

#### Circular flash stimuli

To evaluate the contrast selectivity of cell types in task-constraint model candidates, we simulated responses of each DMN to circular flashes. The networks were initialized at an approximate steady state after 1s of grey-screen stimulation. Afterwards the flashes were presented for 1s. The flashes with a radius of 6 columns were ON (intensity *I* = 1) or OFF (*I* = 0) on grey (*I* = 0.5) background. We integrated the network dynamics with an integration time step of 5 ms. We recorded the responses of the modeled cells in the central columns to compute the flash response index.

#### Flash response index

To derive the contrast selectivity of a cell type, *t_i_*, we computed the flash response index as

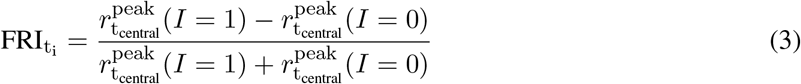

from the non-negative activity

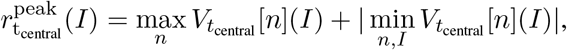

from voltage responses *V_t_central__* [*n*](*I*) to circular flash stimuli of intensities *I* ∈ {0,1} lasting for 1s after 1s of grey stimulus. We note that our index quantifies whether the cell depolarizes to ON- or to OFF-stimuli. However, cells like R1-R8, L1, and L2 can be unrectified, i.e., sensitive to both light increments and light decrements, which is not captured by our index.

For the p-values reported in the results, we performed a binomial test with probability of correct prediction 0.5 (H0) or greater (H1) to both test whether the median FRI from the DMN-ensemble and the best-performing model predict the contrast preferences significantly. Additionally, we found for each individual cell type across 50 DMS that predictions for 29 out of 31 cell types are significant (P < 0.05, binomial).

#### Moving edge stimuli

To predict the motion sensitivity of each cell type in task-constrained DMNs, we simulated the response of each network, initialized at an approximate steady state after 1s of grey-screen stimulation, to custom generated edges moving to 12 different directions, *θ* ∈ [0°, 30°, 60°, 90°, 120°, 150°, 180°, 210°, 240°, 270°, 300°, 330°]. We integrated the network dynamics with an integration time step of 5ms. ON-edges (I = 1) or OFF-edges (*I* = 0) moved on grey (*I* = 0.5) background. Their movement ranged from −13.5° to 13.5° visual angle and we moved them at six different speeds, ranging from 13.92°/s to 145°/s (*S* ∈ [13.92°/s, 27.84°/s, 56.26°/s, 75.4°/s, 110.2°/s, 145.0°/s]).

#### Direction selectivity index

We computed a direction selectivity index of a particular type *t_i_* as

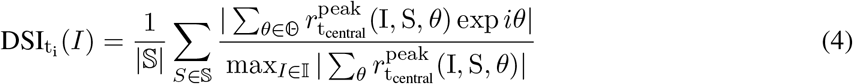

from peak voltages

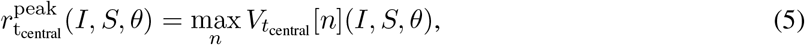

elicited from moving edge stimuli. We parametrized movement angle 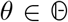, intensities 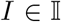, and speeds 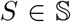 of moving edges. To take the response magnitudes into account for comparing DSI for on- and for off-edges, we normalized by the maximum over both intensities in the denominator. To take different speeds into account, we averaged over 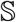.

#### Determining whether a cell type with asymmetric inputs counts as direction selective

We counted a cell type as direction selective if the DSIs from its synthetic measurements were larger than 99% of DSIs from non-motion selective cell types (i.e. those with symmetric filters). We note, however, that estimates of the spatial asymmetry of connectivity from existing connectomic reconstructions can be noisy.

For deriving the 99%-threshold, we first defined a distribution *p*(*d*\*|*t*_sym_) over the direction selectivity index for non-direction selective cells, from peak responses to moving edges of cell types with symmetric inputs, *t*_sym_. We computed that distribution numerically by sampling

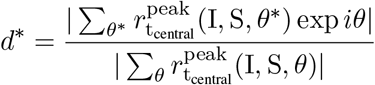

for 100 independent permutations of the angle *θ*\*. We independently computed *d*\* for all stimulus conditions, models, and cell types with symmetric inputs. From *p*(*d*\*|*t*_sym_), we derived the threshold *d*_thresh_ = 0.357 as the 99% quantile of the random variable *d*\*, meaning that the probability that a realization of *d*\* > *d*_thresh_ is less than 1% for cell types with symmetric inputs. To determine whether an asymmetric cell type counts as direction selective, we tested whether synthetically measuring direction selectivity larger than *d*_thresh_ in that cell type is binomial with probability 0.1 (H0) or greater (H1). We identified 12 cell types with asymmetric inputs (T4a, T4b, T4c, T4d, T5a, T5b, T5c, T5d, TmY3, TmY4, TmY5a, TmY18) as direction selective (*P* < 0.05) from our models, and seven cell types with asymmetric inputs to not count as direction selective (T2, T2a, T3, Tm3, Tm4, TmY14, TmY15). See Supplemental Data for reference of cell types with symmetric and asymmetric inputs in our model.

#### UMAP and clustering

We first simulated central column responses to naturalistic scenes (24Hz Sintel video clips from the full augmented dataset) with an integration time step of 10 ms. We clustered models in feature space of concatenated central column responses and sample dimension. Next, we computed a nonlinear dimensionality reduction to 2d (UMAP), and finally fitted Gaussian mixtures of 2 to 5 components to the embedding to label the clusters based on the Guassian mixture model with the number of components that minimize the Bayesian information criterion, using the python libraries umap-learn and scikit-learn^102,136^.

#### Single ommatidium impulse stimuli

To derive spatio-temporal receptive fields, we simulated the response of each network to single ommatidium impulses. Impulses were ON (*I* = 1) on grey (*I* = 0.5) background and presented for 5 ms after 2 s of grey-screen stimulation and followed by 5 s of grey-screen stimulation.

#### Spatio-temporal, spatial and temporal receptive fields

We derived the spatio-temporal receptive field (STRF) of a cell type *t_i_* as the baseline subtracted responses of the central column cell to single ommatidium impulses *J*(*u, v*) at ommatidium locations (*u, v*):

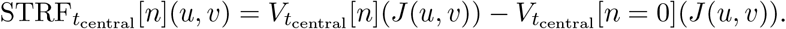

We derived spatial receptive fields, SRFs, from the responses to impulses *J*(*u, v*) at the point in time at which the response to the central ommatidium impulse is at its extremum:

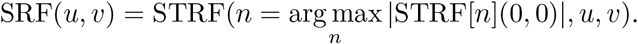

We derive temporal receptive fields, TRFs, from the response to an impulse *J*(0,0) at the central ommatidium: TRF[*n*] = STRF[*n*](0,0).

#### Maximally excitatory naturalistic and artificial stimuli

First, we found the naturalistic maximally excitatory stimulus, **X**^*^, by identifying the Sintel video clip, **X**, from the full dataset with geometric augmentations that elicited the highest possible response in the central column cell of a particular cell type in our models.

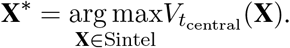

Next, we regularized the naturalistic maximally excitatory stimulus, to yield **X**’, capturing only the stimulus information within the receptive field of the cell, with the objective to minimize

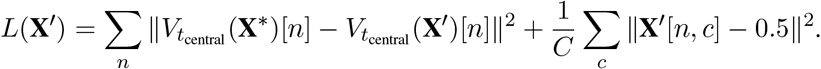

The first summand maintains the exact central response to **X**\*, while the second sets the redundant stimulus information outside of the receptive field to grey (*I* = 0.5).

In addition, we computed artificial maximally excitatory stimuli^137^.

### Training synthetic connectomes

#### Training feedforward synthetic-connectome networks

Sparsified feedforward neural networks with 6 hidden layers (linear transformations sandwiched between rectifications) with equal number of neurons in each hidden layer functioned as synthetic-connectome networks (SCN). The main results describe networks with 128 neurons per hidden layer. We interpret the individual units in the SCN’s as neurons with voltage

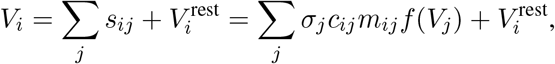

with presynaptic inputs *s_ij_* and resting potentials 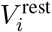. The connectome-constrained synapse strength, *w_ij_*, is characterized by the adjacency matrix *c_ij_*, the signs, *σ_j_*, and the non-negative weight magnitudes *m_ij_*. *c_ij_* = 1 if the connection exists, else *c_ij_* = 0. To respect Dale’s law, the signs were tied to the presynaptic identity, *j*.

We identified the SCN’s parameters *σ_j_, m_ij_*, and 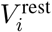 by task-optimization on handwritten digit classification (MNIST)^138^. We determined adjacency matrices, *c_ij_*, for a given connectivity percentage using an iterative local pruning technique, the Lottery Ticket Hypothesis algorithm^139^. The algorithm decreases the connectivity percentage of the SCNs while maintaining high task accuracy.

We optimized SCNs and all simulated networks described below in Pytorch with stochastic gradient descent with adaptive moment estimation (ADAM with AMSGrad), learning rate 0.001, batch size 500, and an exponentially decaying learning rate decay factor of 0.5 per epoch. To constrain the weight magnitudes to stay non-negative, we clamped the values at zero after each optimization step (projected gradient descent). The parameters after convergence minimize the cross-entropy loss between the predicted and the groundtruth classes of the handwritten digits.

#### Simulated networks with known connectivity and unknown strength

Simulated networks inherited connectivity, *c_ij_*, and synapse signs, *σ_j_*, from their respective SCN. In simulated networks, signs and connectivity were held fixed. Weight magnitudes, *m_ij_*, and resting potentials, 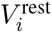, were initialized randomly and task-optimized. Just like SCNs, simulated networks were trained on the MNIST handwritten digit classification task until convergence.

#### Simulated networks with known connectivity and known strength

Alternatively, we imitate measurements of synaptic counts from the SCN’s weight magnitudes:

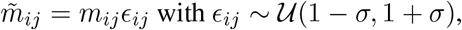

with multiplicative noise to imitate spurious measurements. We used *σ* = 0.5 for the main results. Weight magnitudes were initialized at the measurement, 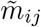, and task-optimized on MNIST with the additional objective to minimize the squared distance between optimized and measured weight magnitudes, 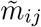 (L2 constraint, Gaussian weight magnitude prior centered around the simulated network’s initialization). We weighted the L2 constraint ten times higher than the cross-entropy objective to keep weight magnitudes of the simulated networks close to the noisy connectomic measurements. Resting potentials, 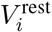, were again initialized randomly and task-optimized.

#### Measuring SCN-simulated network similarity

SCN-simulated network similarity was measured by calculating the median Pearson’s correlation of tuning responses (rectified voltages) of corresponding neurons in the SCN-simulated network pair. In each of the 6 hidden layers, *N* = 100 randomly-sampled neurons were used for comparison. Response tuning was measured over input stimuli from the MNIST test-set (*N* = 10,000 images). Results are medians over all hidden layers and over 25 SCN-simulated network pairs.

## Extended Data Figures

**Extended Data Figure 1:**
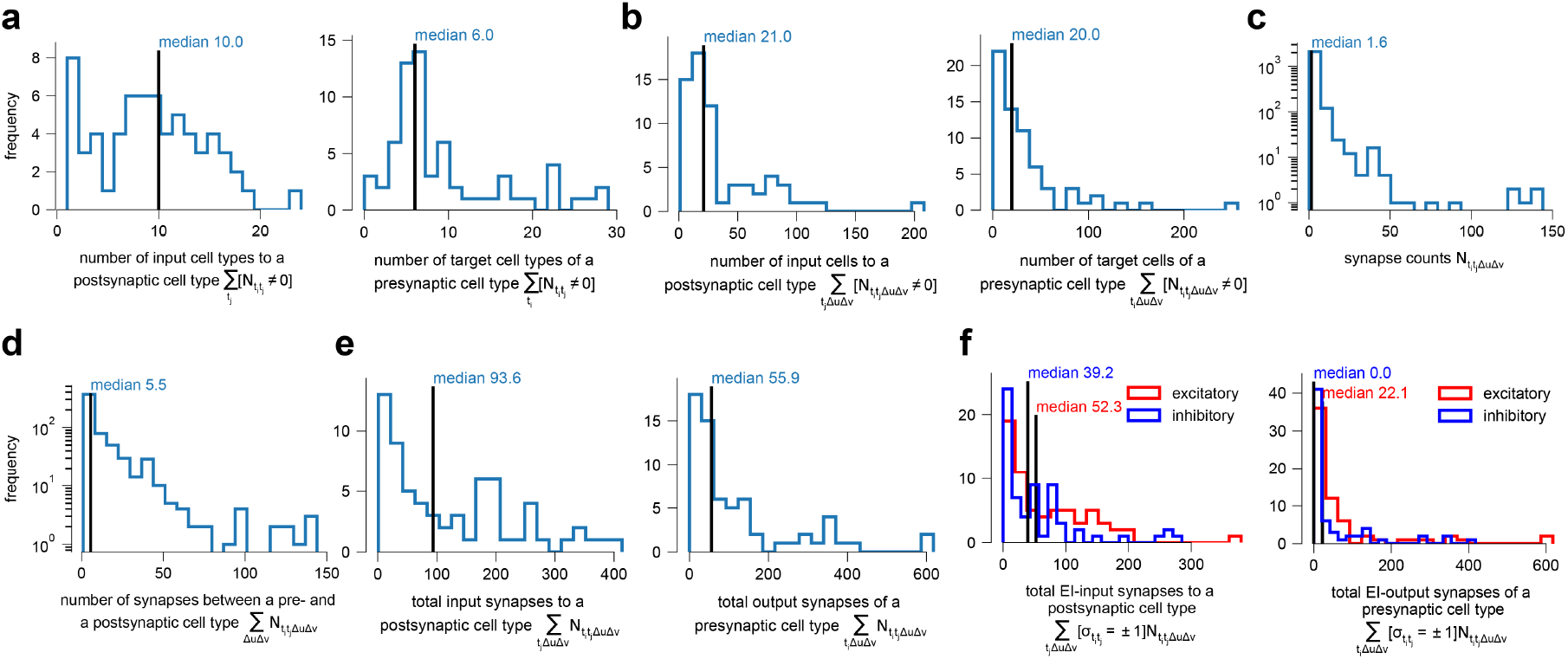
Statistics of derived connectome. **(a)** (left) Half of the 65 cell types receive input from more than ten other cell types, while the other half receives input from less than ten. (right) Half of the 65 cell types project onto more then six other cell types, while the other half projects onto less than six. **(b)** (left) Half of the 65 cell types receive input from 21 up to 200 cells, while the other half receives input from less than 21 cells. (right) Half of the 65 cell types project output onto 20 up to 200 cells, while the other half projects output onto less than 20 cells. **(c)** Half of the connections are characterized by less than 1.6 synapses while the other half are characterized by 1.6 up to hundreds of synapses. **(d)** A pair of presynaptic and postsynaptic cell type is connected by 5.5 synapses in half of the cases and by more than 5.5 up to hundreds in the other half of the cases. **(e)** (left) Half of the 65 cell types receive input from less than 93.6 synapses and the other half between 93.6 to 400 synapses. (right) Half of the 65 cell types project less than 55.9 synapses and the other half projects between 55.9 to 600 synapses. **(f)** Separating (e) into excitatory and inihibitory synapses, (left) we see that half of the 65 cell types receive excitatory inputs from less than 52.3 synapses and the other half from 52.3 to hundreds. Half of the 65 cell types receive inhibitory inputs from less than 39.2 synapses and the other half from 39.2 to hundreds. (right) Half of the 65 cell types project less than 22.1 excitatory synapses and the other half from 22.1 to hundreds. At least half of the 65 cell types project no inhibitory synapses and the rest project between zero to hundreds.

**Extended Data Figure 2:**
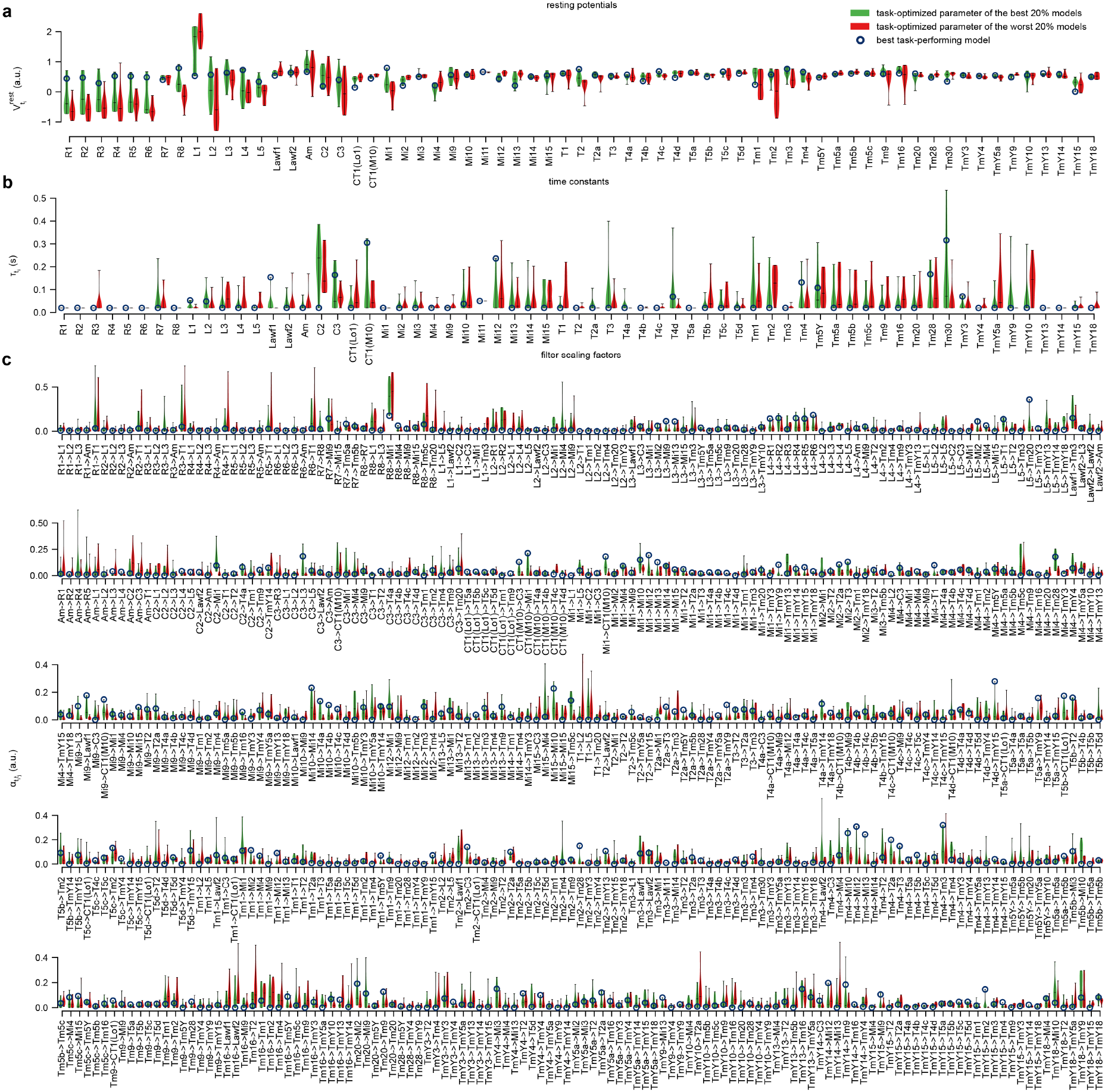
Statistics of learned parameters of best 20% models vs. worst 20% models. **(a)** Task-optimized resting potentials. **(b)** Task-optimized time constants. **(c)** Task-optimized filter scaling factors.

**Extended Data Figure 3:**
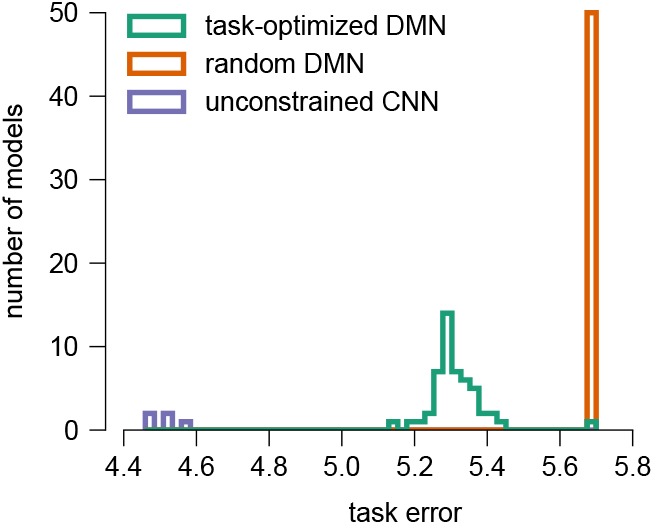
Models with random parameter configurations consistently performed worse on the task than task-optimized models. Task error distributions over ensembles. **(green)** The task error of 50 DMNs distributes between 5.1 and 5.5 (a.u.) after optimization. **(red)** The task error of 50 random DMNs collapses at 5.7 despite optimization of the decoders. **(blue)** An unconstrained CNN with 414,666 parameters reaches a task error between 4.5 and 4.6 (here 5 models).

**Extended Data Figure 4:**
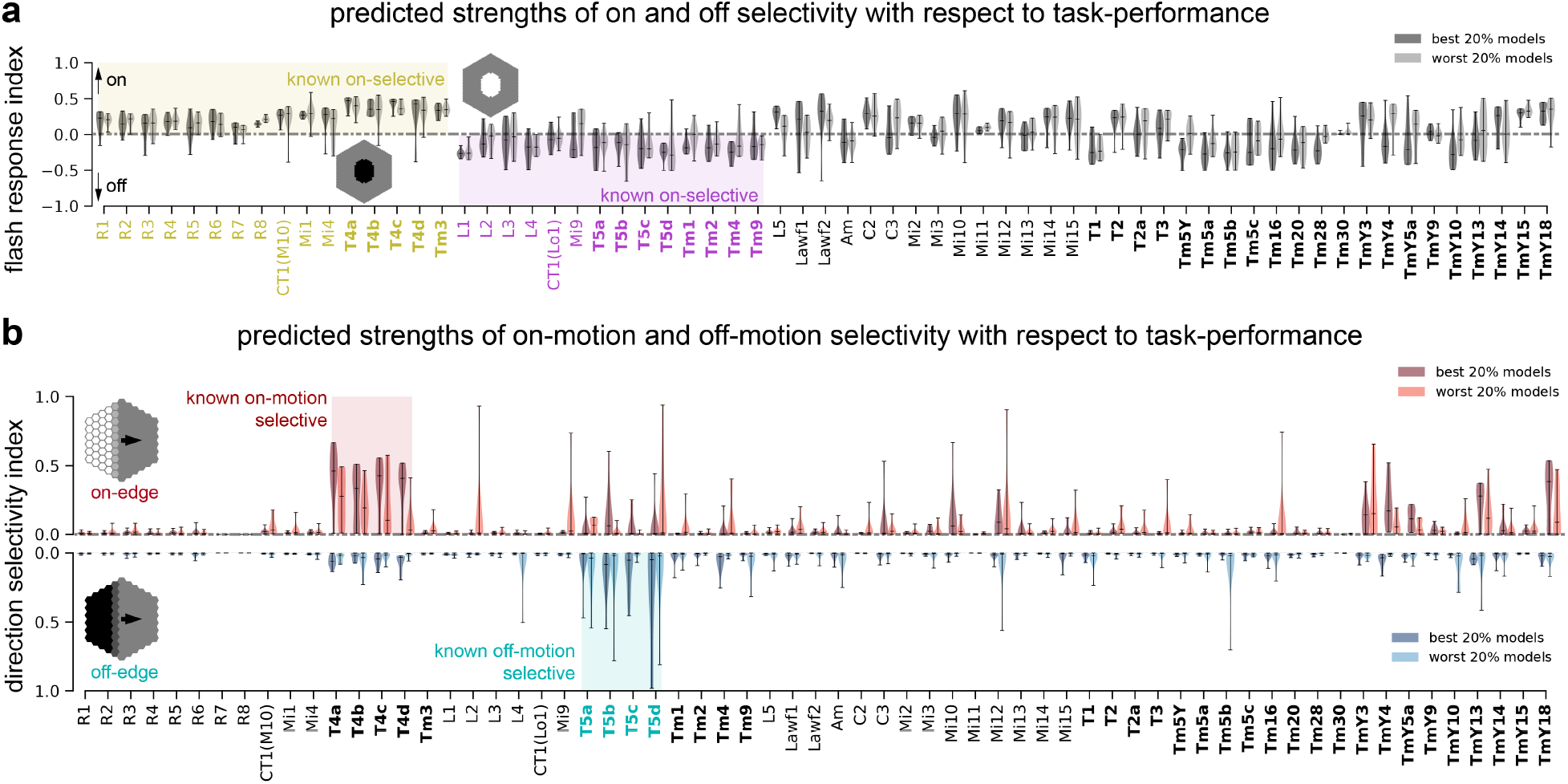
Predicted tuning with respect to task-performance. **(a)** Flash response index computed as the max-abs-scaled peak response to an off flash subtracted from the max-abs-scaled peak response to an on flash - both of approximately 35° radius and presented for 1s after 2 seconds of grey input. Values above 0 indicate on-polarity, values below zero indicate off-polarity. Known on-polar and off-polar cell types are colored in yellow and magenta. **(b)** Single cell type direction selectivity of best 20% task-performing models versus worst 20% task-performing models of an ensemble of 50 models as a result of peak voltage responses in central columns to on-edges and off-edges moving towards all possible directions on grey background (Equation 9). The bolded cell types are those which optic flow is decoded from.

**Extended Data Figure 5:**
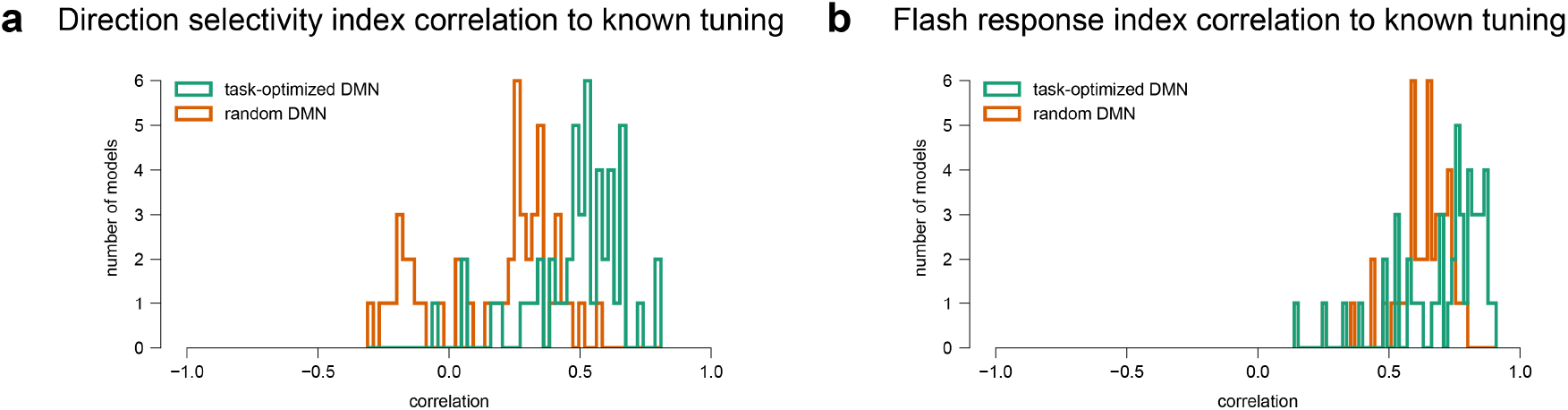
Models with random parameter configurations predict the known direction selectivity indices and flash response indices worse than task-optimized models. We correlate predicted tuning metrics from each model to the known tuning properties to answer if models with random parameter configurations lead to similarly accurate tuning predictions as task-optimized models. **(a)** Task-optimized DMNs (green) predict more accurate direction selectivity indices than randomly parametrized models (red) (P =2 × 10^-11^, Mann-Whitney-U). **(b)** Task-optimized DMNs (green) predict more accurate flash response indices than randomly parametrized models (red) (P = 5.3 × 10^-5^, Mann-Whitney-U).

**Extended Data Figure 6:**
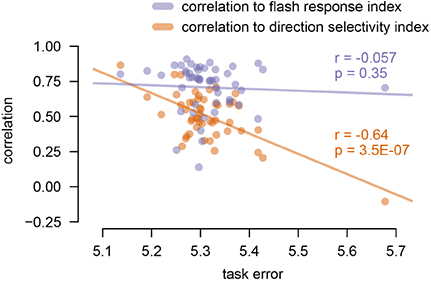
Better task performing models predict motion tuning neurons better. We correlate predicted tuning metrics from each model to the known tuning properties to understand when better performing models give us better tuning predictions. **(orange)** When correlating the direction selectivity index of each model to the binary known properties for T4 and T5 and their input cell types, we find that this correlation is higher for better performing models. **(magenta)** While the models predicted the known contrast preferences generally well, the correlation of flash response index to the binary known contrast preferences of 31 cell types did not significantly increase with better performing models.

**Extended Data Figure 7:**
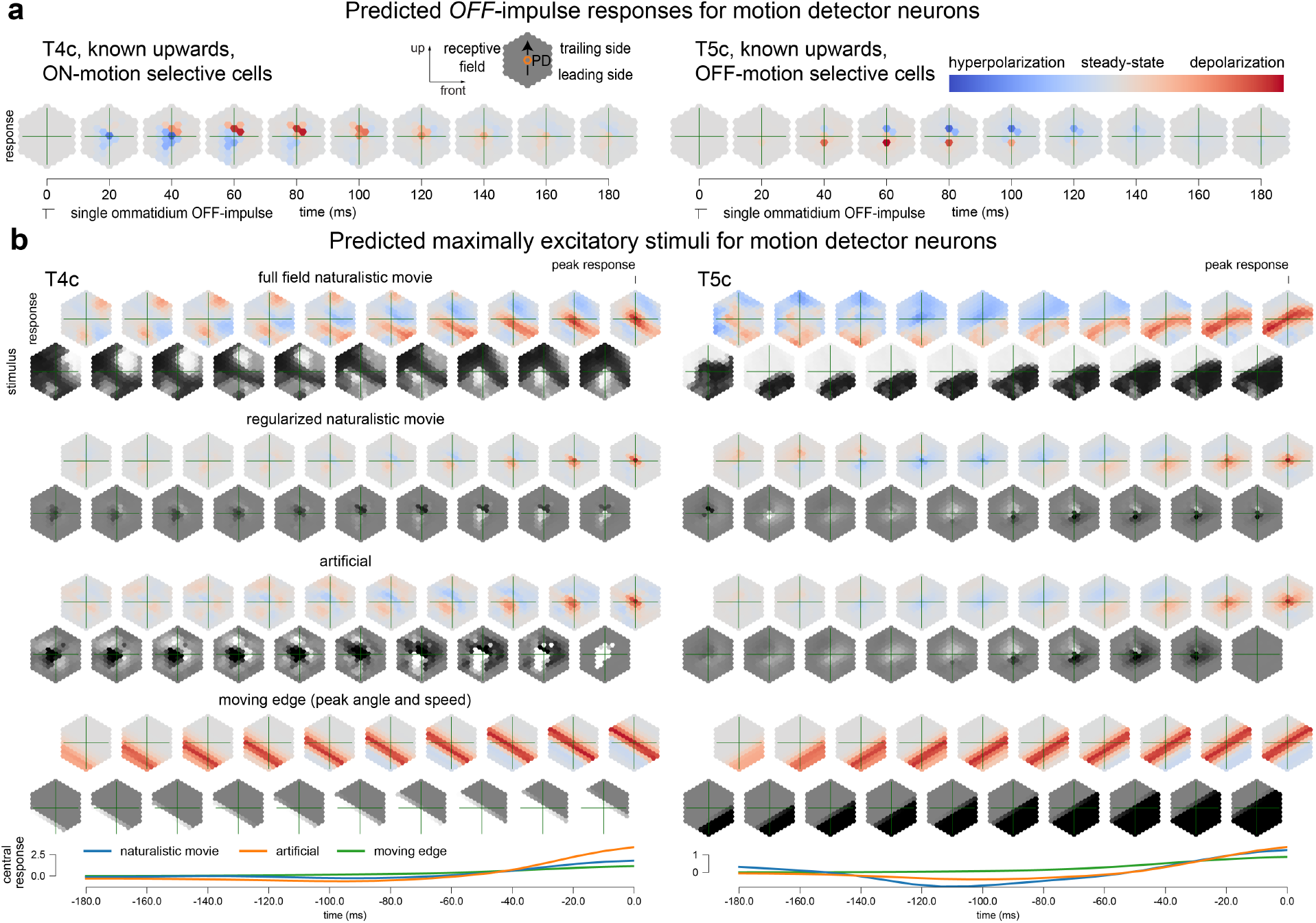
Spatio-temporal receptive fields mapped with OFF-impulses and maximally excitatory stimuli. **(a)** Spatio-temporal receptive field mapping with single ommatidium OFF-impulses. **(b)** Maximally excitatory stimuli and baseline-subtracted responses. Including full-field naturalistic, regularized naturalistic, artificial, and moving edge stimuli and responses. Moving edge angle and speed maximize the central cell peak response. Artificial stimuli are optimized from initial noise to maximize the central cell activity using gradient ascent plus full-field regularization towards grey. The last row shows the baseline-subtracted central cell responses. Peak central cell responses at time point zero.

**Extended Data Figure 8:**
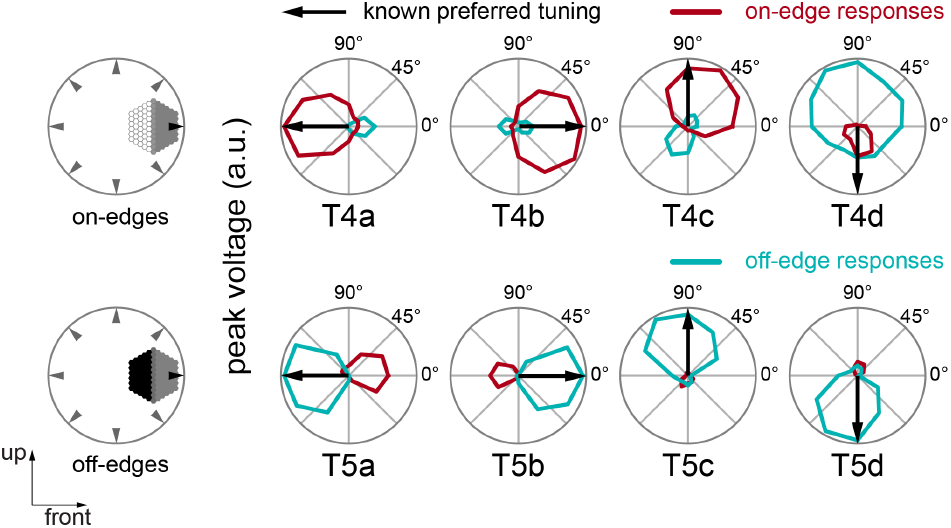
Motion tuning predictions for T4 and T5 subtypes to preferred and null contrast edges in the best-task-performing model.

**Extended Data Figure 9:**
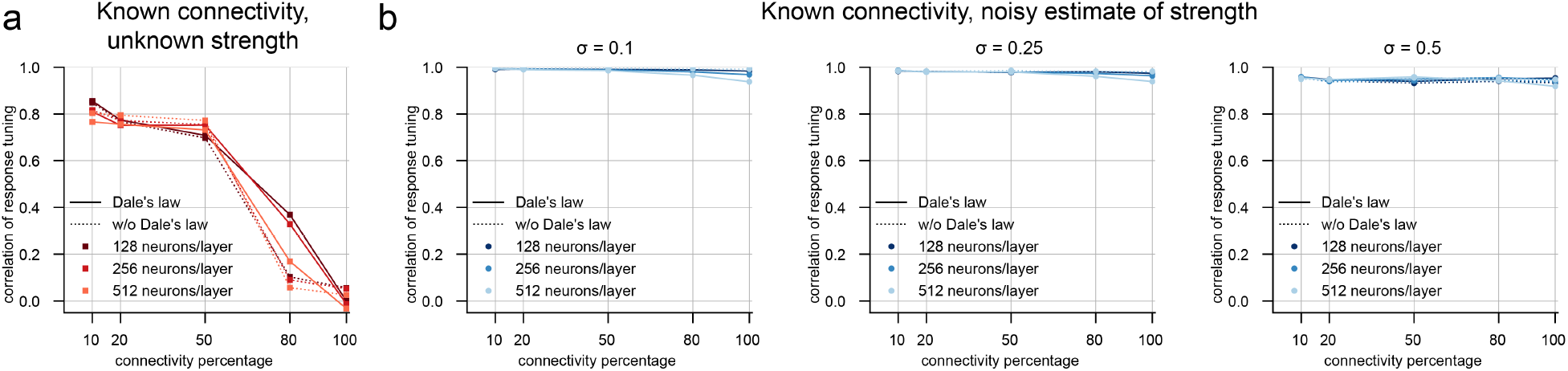
Investigating the role of sparse connectivity with synthetic networks for MNIST handwritten digit recognition. **(a)** Median hidden-layer response correlation as a function of synthetic network connectivity percentage for task and connectome constrained models that had access to only connectivity information but not connection strength. **(b)** Median hidden-layer response correlation as a function of synthetic network connectivity percentage for task and connectome constrained models with access to noisy estimates of connection strength (multiplicative noise levels of *σ* = 0.1, *σ* = 0.25, and *σ* = 0.5, respectively). Connectome constrained models were task optimized with a soft (L2) constraint with the noisy connectomic measurements.

## Supplementary Information

### Supplementary Note 1 Probabilistic model for automatic construction of connectome

The EM-datasets primarily contain the lamina projections, medulla (FIB-25), lobula and lobula plate (FIB- 19) cells, and the important cell types of the primary motion detection circuit (T4, T5). In total, they contain 1801 neurons (702 from FIB-25 and 1099 from FIB-19), with hand-annotated positions available for 830 of these neurons (SI Figure 1). To accurately localize the remaining neurons and synapses and to derive cell-type connectivity (Fig. 1b), we build a probabilistic expectation maximization algorithm that takes synaptic connection statistics, projected synapse center-of-mass clusters and existing column annotations into account. We verified the quality of our reconstruction, concluding that even in the absence of 90% of the hand-annotations available to us, we could accurately position the majority of the neurons in our circuit reconstruction (SI Table 1, SI Figure 2). In the absence of ground-truth annotations, we verified the quality of our reconstruction by the *recovery* and *consistency* rates (Table 1). The *recovery* rate is defined as the ratio of reference positions successfully recovered by our algorithm after removing a random proportion of reference positions from the data. For each 10% of the reference positions removed, on average, only 2.5% are not correctly recovered. The *consistency* rate is defined as the fraction of neurons obtaining the same position between evaluations of the algorithm starting with a different fraction of reference positions. For each 10% of the reference positions removed, on average, an additional 3.9% of neurons are not consistently estimated. We found that even just 10% (83 positions) of the available ground truth was sufficient to robustly position the majority of the neurons (64.3%, 534 positions) into the correct columns, and annotate 48% (865 neurons) perfectly consistent.

**Table 1:**
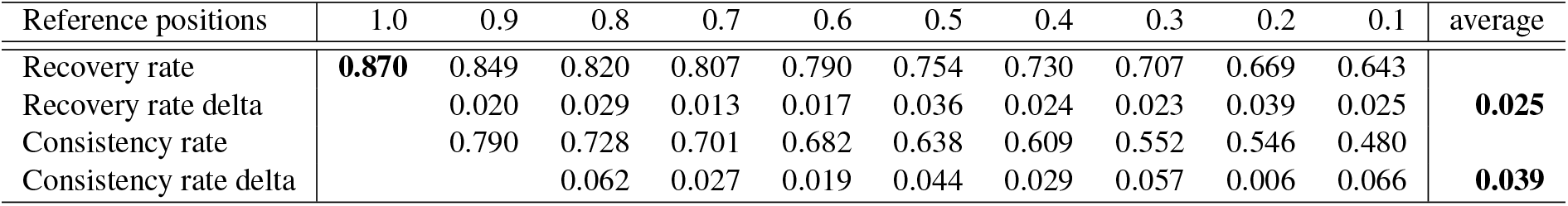
Recovery and consistency of columnar cell position estimation.

#### Probabilistic expectation maximization for unassigned neurons

Each neuron is either annotated in the dataset (in 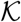), assigned to a position (in 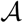) by our algorithm, or still under evaluation (in set 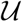). Iteratively, the EM-expectation step updates the normal distribution 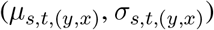 of expected synapse counts between neurons, while the EM-maximization step updates the positions (*y, x*) of all neurons not yet assigned to a column (set 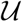).

#### Synapse center-of-mass as a neuron column position proxy

For offset assignment, we take the center-of-mass of all synapses belonging (pre- or postsynaptic) to a neuron into account. These are generally a more useful hint than the physical location of the cell body, as the cell bodies are mostly positioned on the side of the neuropiles, and not near the column to which the neuron belongs to, but most cells have a majority of synapses in close proximity to their own column. Since many neurons span more than one layer in a neuropile, or even multiple neuropiles, we first group all synapses per neuron into clusters, and then assign the center-of-mass of these clusters to one of up to *N* = 5 super-clusters (approximately matching the medulla, lobula and lobula plate). Clustering is done via k-means with the ideal number of clusters determined by silhouette scores. The super-clusters allow to project 3D synapse coordinates onto a retinotopic 2D hexagonal lattice with a simple projection and affine transformation.

#### Hybrid cost-model for neuron-position likelihood estimates

Prior knowledge about the normal distribution 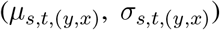 of expected synapse counts between neurons from existing annotations is required to express the probability of any cell specimen *c* to be located at position (*y, x*). This metric correlates a neuron to all pre- and postsynaptic neurons it is connected to, of which some already have a fixed, known position. Thereby, the neighbouring neurons with known position (in 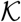 and 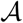) contribute to stabilize the probabilities of unassigned neurons (in set 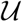). For columnar, spatially repeated neurons, we can also assume that only one neuron per position is present. This prior rapidly discounts the number of possible positions an unassigned neuron can have, each time another neuron becomes assigned (moves from 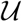 to 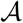).

#### Fusion of sepratately evaluated datasets

Since FIB-19 and FIB-25 are evaluated separately, we have to combine the estimated parameters of both models to a single, coherent model. The datasets overlap partially, in terms of the neurpoiles and cell types covered, and our method therefore fuses the model by always taking the larger estimated parameter.

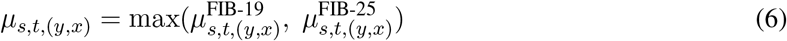

This method of model fusion only underestimates the number of synapses between two neurons if they have connections in two different neuropiles, and if each neuropile is exclusively covered by only one dataset.

#### Pruning spurious synapses

Some automatic annotations, which are not proof read in the FlyEM DVID data, contain a large number of autapses per neuron on most neuron types, arising from wrongly detected synapses in the cell bodies themselves. Additionally, there are many statistically insignificant single synapses left from the assignment algorithm. We imposed the following additional filter on our estimated model parameters, to remove both autapses and spurious connections with less than one synapse on average.

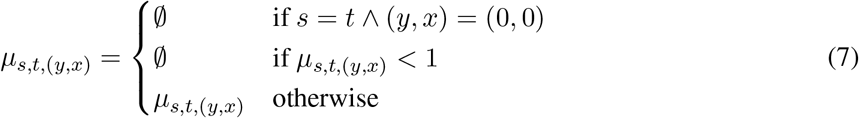

Finally, neuron types without connections and synapses with either missing target or source can be removed. The resulting mean synapse counts *μ*_*s,t*,(*y,x*)_ form the convolutional filters for our simulation.

**Table 2:** Results of the probabilistic model construction.

| Pos. removed [%] | 0 | 10 | 20 | 30 | 40 | 50 | 60 | 70 | 80 | 90 | 100 |
| --- | --- | --- | --- | --- | --- | --- | --- | --- | --- | --- | --- |
| Reference positions | 830 | 747 | 664 | 581 | 498 | 415 | 332 | 249 | 166 | 83 | 0 |
| Combined | 1801 | 1459 | 1305 | 1346 | 1248 | 1152 | 1229 | 1081 | 972 | 944 | 0 |
| FIB-25 | 702 | 592 | 573 | 566 | 538 | 509 | 491 | 464 | 445 | 399 | 0 |
| FIB-19 | 1099 | 867 | 732 | 780 | 710 | 643 | 738 | 617 | 527 | 545 | 0 |
| Combined (annotated) | 721 | 708 | 694 | 684 | 658 | 643 | 626 | 576 | 566 | 499 | 0 |
| FIB-25 (annotated) | 393 | 387 | 379 | 375 | 360 | 353 | 339 | 307 | 304 | 272 | 0 |
| FIB-19 (annotated) | 328 | 321 | 315 | 309 | 298 | 290 | 287 | 269 | 262 | 227 | 0 |

**Supplementary Figure 1:**
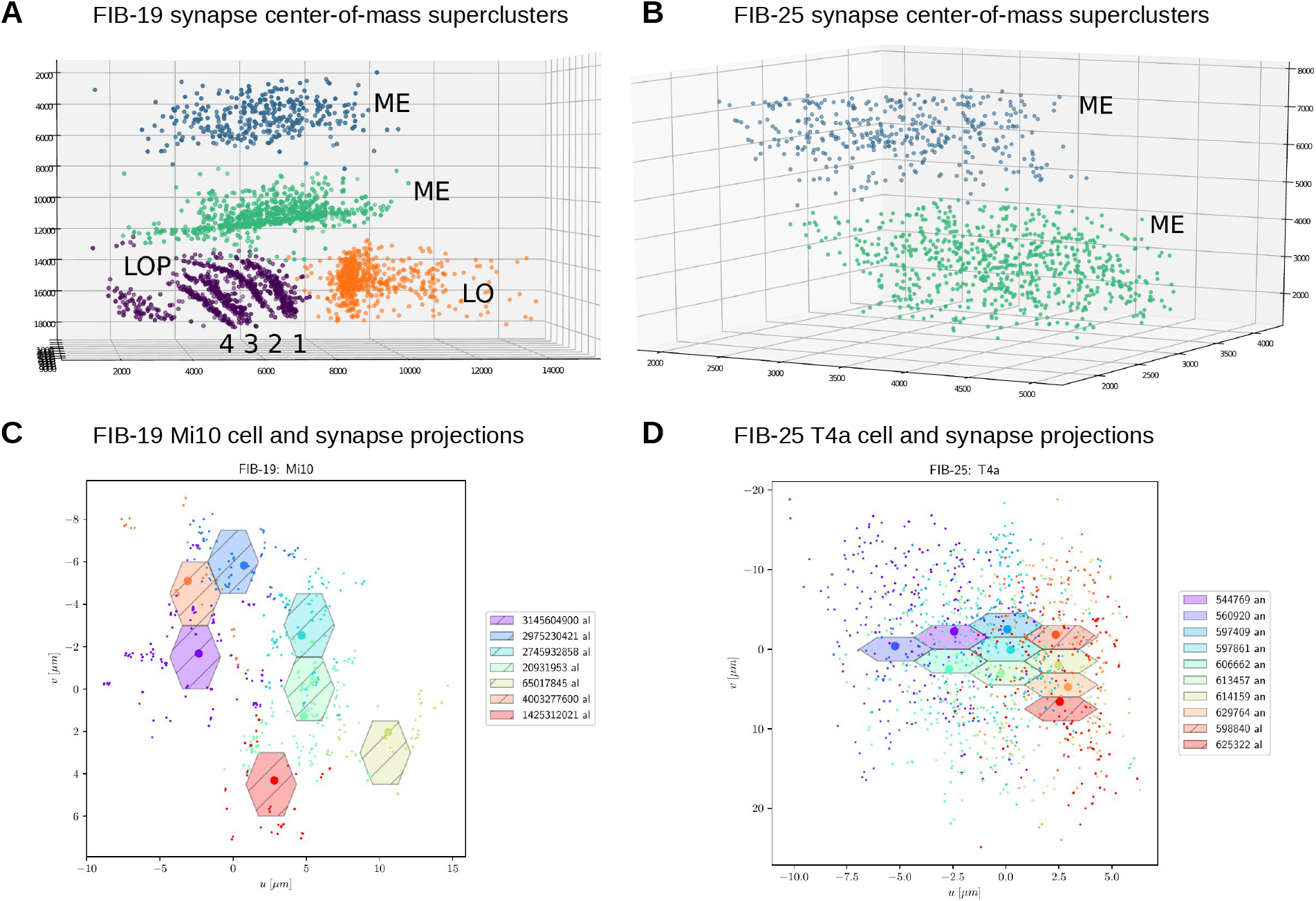
**(A)** FIB-19 synapse center-of-mass superclusters. The clusters form two strata in the medulla (ME), and in the lobula (LO) and lobula plate layers (LOP, 1-4) additionally. Each dot corresponds to the center-of-mass of all synapses belonging to the super-cluster. Typically, each diverging arborization of a cell becomes a distinct location, which helps our probabilistic model to project 3D positions of synapses into retinotopic 2D planes, despite the lobula having a different spatial orientation (perpendicular) than the medulla and lobula (Fig. 1a and c). **(B)** FIB- 25 synapse center-of-mass superclusters. The clusters form two strata (ME) in the medulla. **(C)** Mi10 cell type in FIB-19 with no pre-annotated lattice positions. The seven cell specimen (hexagons) are recovered by our probabilistic algorithm. Individual synapses and synapse center-of-mass projections are superimposed. **(D)** T4a cell type in FIB-25 with eight pre-annotated (an) and two recovered (al) lattice positions. The projected synapse positions show directional displacement consistent with the direction selectivity of T4a cells (Fig. 3a).

**Supplementary Figure 2:**
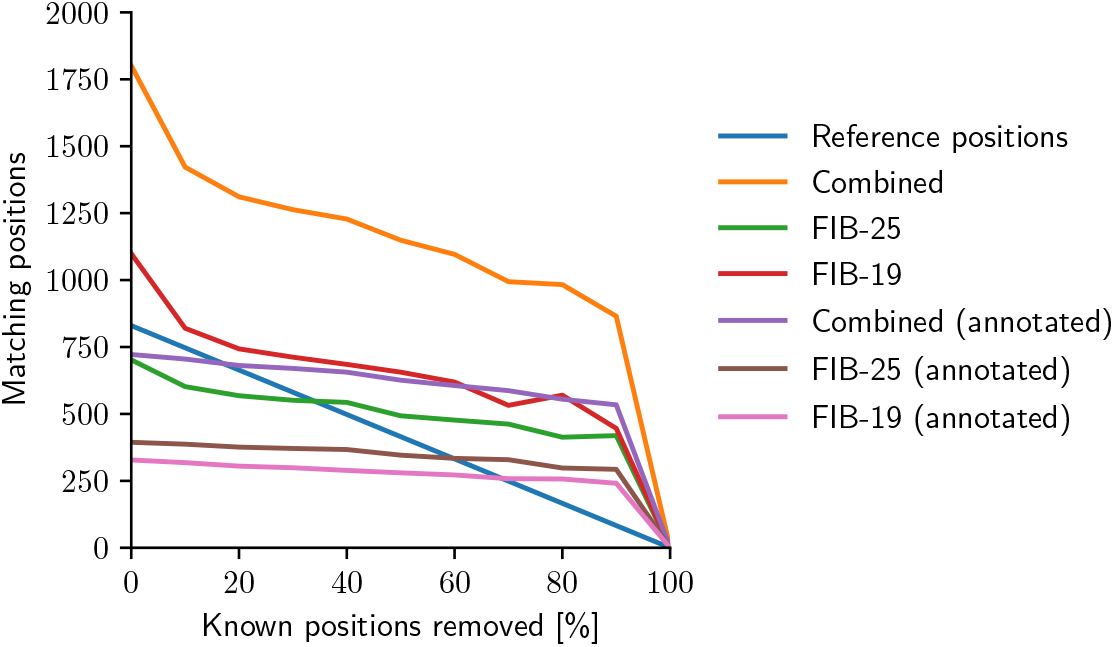
Results of the probabilistic model construction.

### Supplementary Note 2 Manually constructed connectome

#### Lamina and ommatidia model

Since neither FIB-19 nor FIB-25 contain the connections of the ommatidia or first neuropile, the lamina^48,68^, we reused and refined the existing hand-crafted model from our previous work^119^, which is based on data from Rivera-Alba *et al*.^50^ and Tuthill *et al*.^124,125^.

#### Non-columnar single CT1 cell model

While we did in general not model any neurons with large tangential branches, such as Mt, Mt, Pm, Dm, which span many columns and are therefore insufficiently segmented in FIB-19 and FIB-25, we did model the single CT1 cell present in the lobula CT1(Lo1) and medulla CT1(M10)^49^. Because a multi-compartment model with bidirectional electrical synapses resulted often in oscillatory dynamics in earlier modeling attempts and because CT1 terminals were found to act as functionally independent units^70^ we modelled CT1 as two anatomically separate cell types CT1(Lo1) and CT1(M10).

#### Non-columnar periodic cells

Because lamina-wide-field neurons, Lawf1 and Lawf2, do not occur in each individual column but more sparsely, we modeled them with an inferred spatial stride to occur more sparsely resulting in 123 cell of each type in our model (there are approx. 140 Lawf2 neurons per optic lobe, and each of approx. 700 columns is innervated by approx. 5 Lawf2 cells^124^).

#### Hexagonal lattice rendering of connectome

For compilation into the hexagonal grid, the convex hull of the filters is filled with ones to remove spatial discontinuities. Although these are considered mostly false positives from the connectome reconstruction^12^, this allowed for weak autapses in our hexagonal model that did not affect the tuning predictions.

#### Additional proofreading

We manually proofread filters on the hexagonal lattice and compared them to the reported filters in the literature to ensure overall correspondence. We found that the reconstruction did not fully capture the asymmetry reported in^48^ of the T5 anatomical receptive field of Tm9, which we then substituted by a Gaussian at the reported offset column scaled by the reported number of input synapses. For few T4 and T5 inputs the number of input synapses reported in the literature^48^ slightly deviated from our reconstruction. To get a better initialization of our filter scale we scaled them to closely match the number of input synapses reported^48^.

### Supplementary Note 3 Investigating the role of sparse connectivity with synthetic networks for MNIST digit recognition

#### Training feedforward synthetic networks

The weight matrix for each layer in Dale’s-law-based synthetic networks (DLTrues) is decomposed into three components: binary adjacency matrix, non-negative weight magnitudes, and a sign vector.

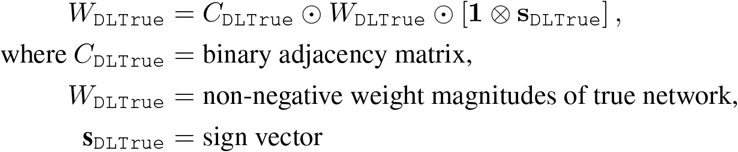

By means of projected gradient descent, *W*_DLTrue_ is enforced to be non-negative and is initialized from the absolute value of the He initialization distribution^140^. Although sign vector **s**_DLTrue_ is randomly initialized with equal probability to be either −1 or +1 to represent inhibitory and excitatory synapses respectively, its elements are allowed to assume values in R over the course of training.

#### Inducing sparsity

Binary adjacency matrix *C*_DLTrue_ is initialized to be a unit matrix and is later updated according to the desired true network connectivity level. Following the LTH algorithm, a portion of the lowest-magnitude weights, designated to be pruned, were identified from 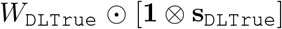. For pruning synapses, the corresponding entries in the adjacency matrix *C*_DLTrue_ were then set to zero. After each pruning iteration, weight magnitudes were reset back to their original initialization, followed by a final training run post-pruning.

#### Training with/without sign constraints

In addition to a Dale’s-law-based sign constraint, we also experimented with networks trained without any sign constraint. No restrictions were imposed on the nature of outgoing synapses i.e., a neuron can have both excitatory and inhibitory outgoing synapses. For true network variants trained without a sign constraint, **1**⊗**s**_DLTrue_ was simply replaced by a sign matrix *S*_nonDLTrue_ initialized in a similar fashion; that is, all entries were initialized to be in {-1, +1} with equal probability. We will refer to true networks trained without a sign constraint as `nonDLTrues`.

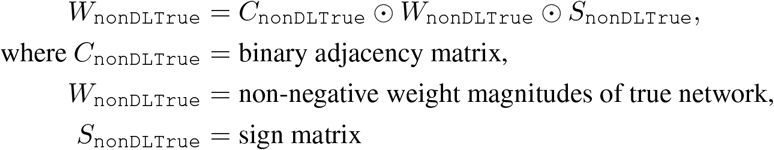

#### Training simulated networks

As elements in a true network’s **s**_DLTrue_ or *S*_nonDLTrue_ are allowed to assume values in ℝ while training, only the signs of these elements are inherited by the true networks’s respective simulated network.

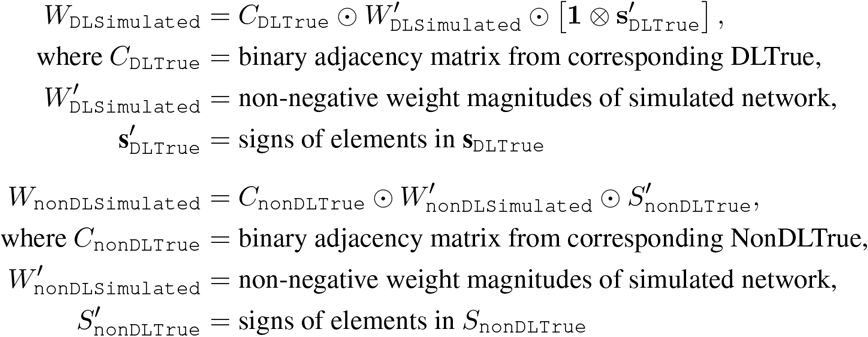

Extended Data Fig. 9 A shows median hidden-layer tuning correlations for networks trained with and without Dale’s law sign constraint for three different architectures.

#### Training simulated networks that had access to weight magnitudes

Three levels of multiplicative noise *σ* = 0.1, 0.25, 0.5 were explored, inducing low-noise, medium-noise, and high-noise weight estimates, respectively. Each noise level represents the maximum percentage by which a weight magnitude could be perturbed.

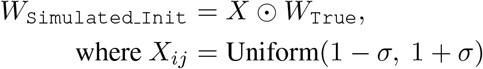

Simulated networks were trained with a Gaussian prior on the weights centered around the noisy initialization. In effect, this additional loss term penalizes trainable weights for deviating from their noisy initialization.

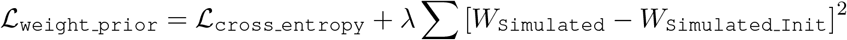

Fig. 9B shows median hidden-layer tuning correlations for networks with low-, medium-, and high-noise weight perturbations.

#### Sparse networks have larger local minima

We inspected the size of the simulated networks’ local minima by analyzing how well they converged when initialized from a perturbed local optimum. Response tuning correlation in this context was used to quantify efficacy of convergence. After training the connec- tome simulations on the same handwritten digit classification task, we found that sparser networks were able to recover function even from highly-perturbed network initializations. By virtue of weight pruning, as the number of free parameters in the true network (and hence simulated network) are reduced, we believe that the size of the simulations’ local minima increase, allowing sparser simulated networks to converge even when initialized far from optimum.

Simulated networks were initialized with a noisy version of their respective true network’s weights. Three varying levels of multiplicative noise (*σ* = 0.1, 0.25, 0.5) were used to perturb the simulations’ local minima.

Extended Data Fig ?? shows median hidden-layer tuning correlations at initialization distances of *σ* = 0.1, 0.25, 0.5 for networks with 128, 256, and 512 neurons per hidden layer, respectively.

## Footnotes

1 https://github.com/TuragaLab/flyvis

