## Supplemental Cell Profiles for "Connectome-constrained deep mechanistic networks predict neural responses across the fly visual system at single-neuron resolution"

### Cell type

1 R1 3

2 R2 10

3 R3 16

4 R4 23

5 R5 30

6 R6 37

7 R7 43

8 R8 49

9 L1 55

10 L2 62

11 L3 69

12 L4 76

13 L5 83

14 Lawf1 90

15 Lawf2 96

16 Am 102

17 C2 109

18 C3 116

19 CT1(Lo1) 123

20 CT1(M10) 130

21 Mi1 136

22 Mi2 143

23 Mi3 150

24 Mi4 156

25 Mi9 163

26 Mi10 170

27 Mi11 177

28 Mi12 183

29 Mi13 190

30 Mi14 197

31 Mi15 204

32 T1 210

| | | |
|---|---|---|
| 33 | T2 | 216 |
| 34 | T2a | 223 |
| 35 | T3 | 230 |
| 36 | T4a | 236 |
| 37 | T4b | 243 |
| 38 | T4c | 250 |
| 39 | T4d | 257 |
| 40 | T5a | 264 |
| 41 | T5b | 270 |
| 42 | T5c | 276 |
| 43 | T5d | 283 |
| 44 | Tm1 | 290 |
| 45 | Tm2 | 297 |
| 46 | Tm3 | 304 |
| 47 | Tm4 | 311 |
| 48 | Tm5Y | 318 |
| 49 | Tm5a | 324 |
| 50 | Tm5b | 330 |
| 51 | Tm5c | 336 |
| 52 | Tm9 | 343 |
| 53 | Tm16 | 350 |
| 54 | Tm20 | 357 |
| 55 | Tm28 | 363 |
| 56 | Tm30 | 370 |
| 57 | TmY3 | 376 |
| 58 | TmY4 | 383 |
| 59 | TmY5a | 390 |
| 60 | TmY9 | 397 |
| 61 | TmY10 | 403 |
| 62 | TmY13 | 409 |
| 63 | TmY14 | 416 |
| 64 | TmY15 | 423 |
| 65 | TmY18 | 430 |

# 1 R1

← Cell types

#### Figures

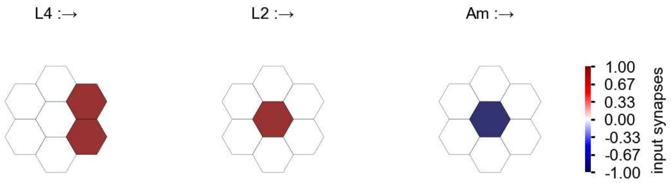

R1 - Figure 1: **Anatomical receptive fields.** Each colored hexagon is a projection, with the projection strength characterized by the average number of synapses that we count from the EM reconstruction. Red indicates excitatory synapses, blue indicates inhibitory synapses from inferred signs. Filters in the order of their total number of synapses.

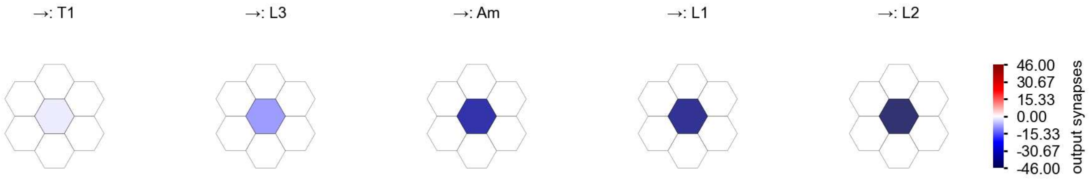

R1 - Figure 2: **Anatomical projective fields.** Each colored hexagon is a projection, with the projection strength characterized by the average number of synapses that we count from the EM reconstruction. Red indicates excitatory synapses, blue indicates inhibitory synapses from inferred signs. Filters in the order of their total number of synapses.

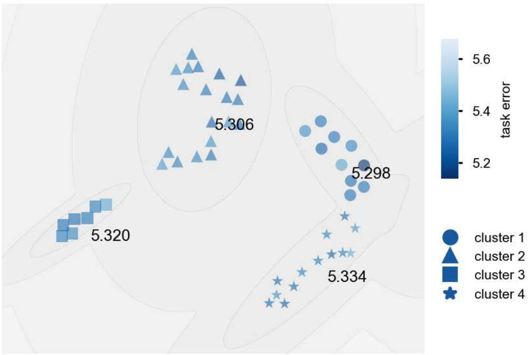

R1 - Figure 3: **Clustering of the responses to naturalistic stimuli.** From responses to naturalistic scenes from Sintel. Scatterpoints represent individual models colored by their task error.

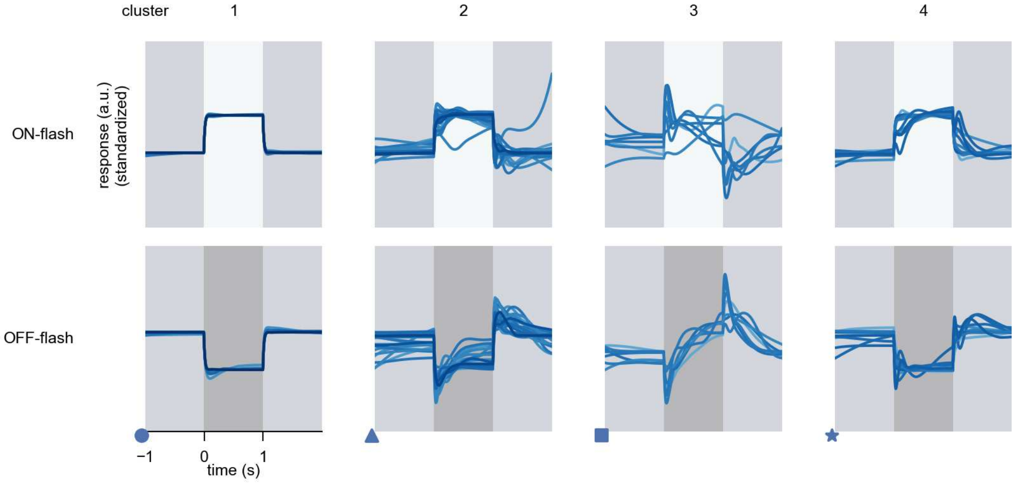

R1 - Figure 4: **Responses to flashes.** The top row shows responses to ON-flashes, the bottom row shows responses to OFF-flashes. The responses from the different models in the different clusters (columns) overlay and are colored in the respective task error of the model. The circular flashes occur at second zero, cover 6 ommatidia in radius, and last for one second. Before and after, a grey-stimulus leads to a stationary state of the network. The traces are standardized.

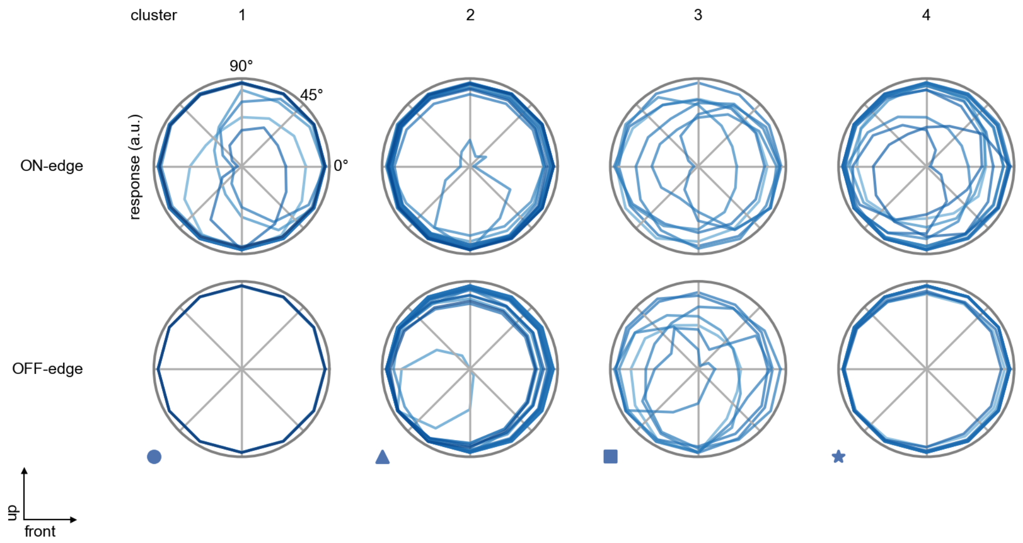

R1 - Figure 5: **Peak responses to moving edges.** The top row shows normalized peak responses to moving ON-edges, the bottom row shows normalized peak responses to moving OFF-edges. The edge-stimuli move in different directions from 0 to 360 degrees. Before, a grey-stimulus led the network to a stationary state. The responses from the different models in the different clusters (columns) overlay and are colored in the respective task error of the model. The peak responses are averaged over all assessed edge-speeds.

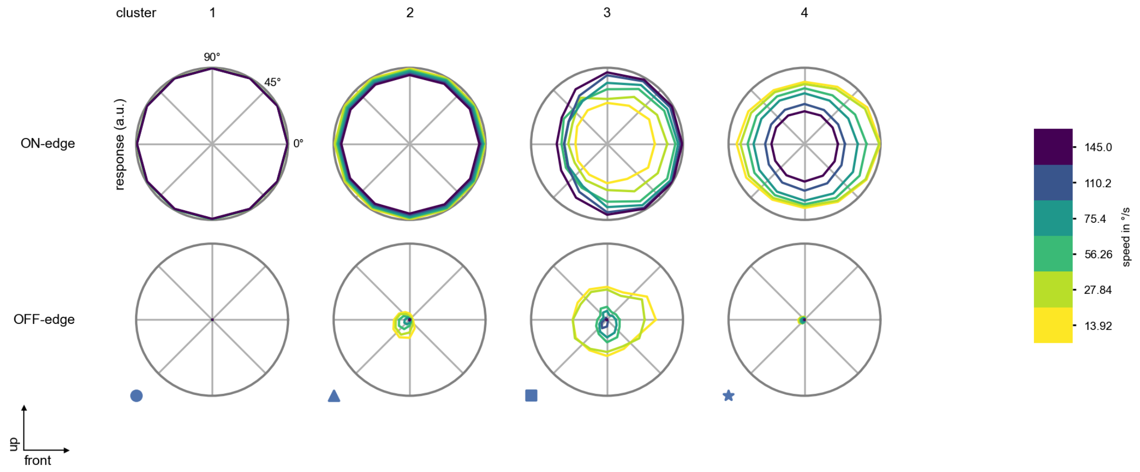

R1 - Figure 6: **Peak responses to moving edges from best-performing models.** The top row shows peak responses to moving ON-edges, the bottom row shows peak responses to moving OFF-edges. The edge-stimuli move in different directions from 0 to 360 degrees and at different speeds. Before, a grey-stimulus led the network to a stationary state. The columns show responses from the best-task-performing model in the respective cluster.

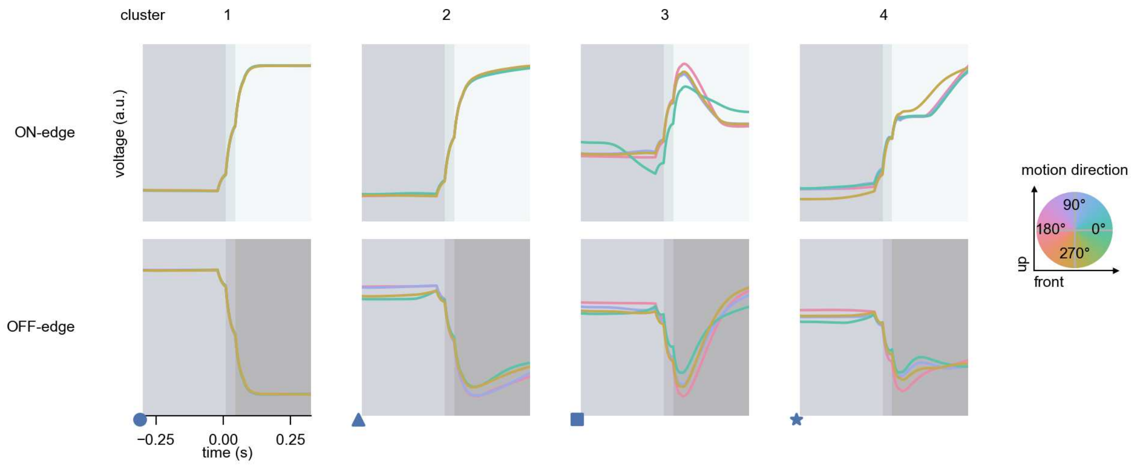

R1 - Figure 7: **Responses to moving edges from best-performing models.** The top row shows responses to moving ON-edges, the bottom row shows peak responses to moving OFF-edges. The edge-stimuli move in different directions from 0 to 360 degrees and at different speeds. Before, a grey-stimulus led the network to a stationary state. The columns show responses from the best-task-performing model in the respective cluster. Example responses to edges moving at 75.4°/s in all cardinal directions (green 0°, blue 90°, red 180°, yellow 270°) from -22.5 to 22.5° visual angle. Background contour indicates the edge, passing by the central column at second zero.

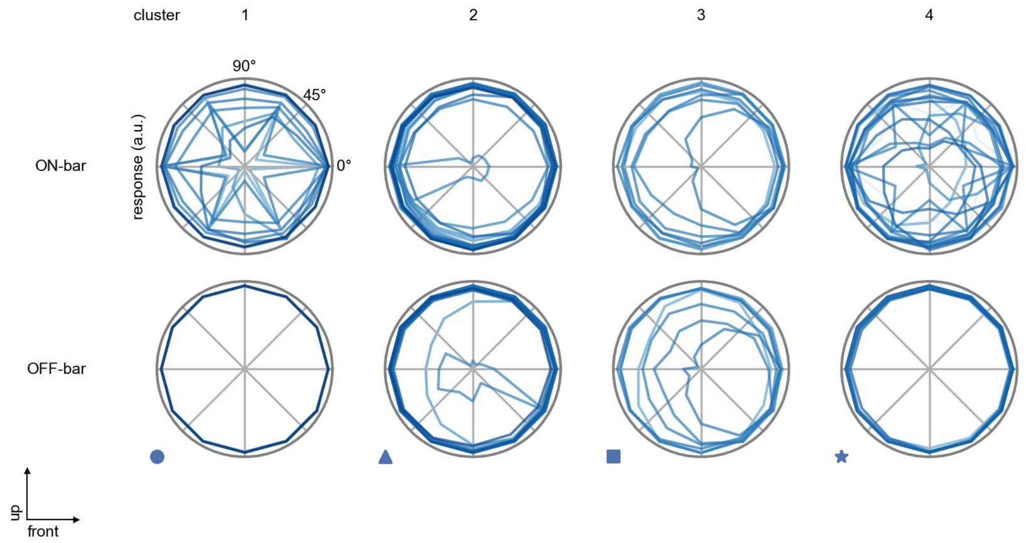

R1 - Figure 8: **Peak responses to moving bars.** The top row shows normalized peak responses to moving ON-bars, the bottom row shows normalized peak responses to moving OFF-bars. The bar-stimuli move in different directions from 0 to 360 degrees. Before, a grey-stimulus led the network to a stationary state. The responses from the different models in the different clusters (columns) overlay and are colored in the respective task error of the model. The peak responses are averaged over all assessed bar-speeds.

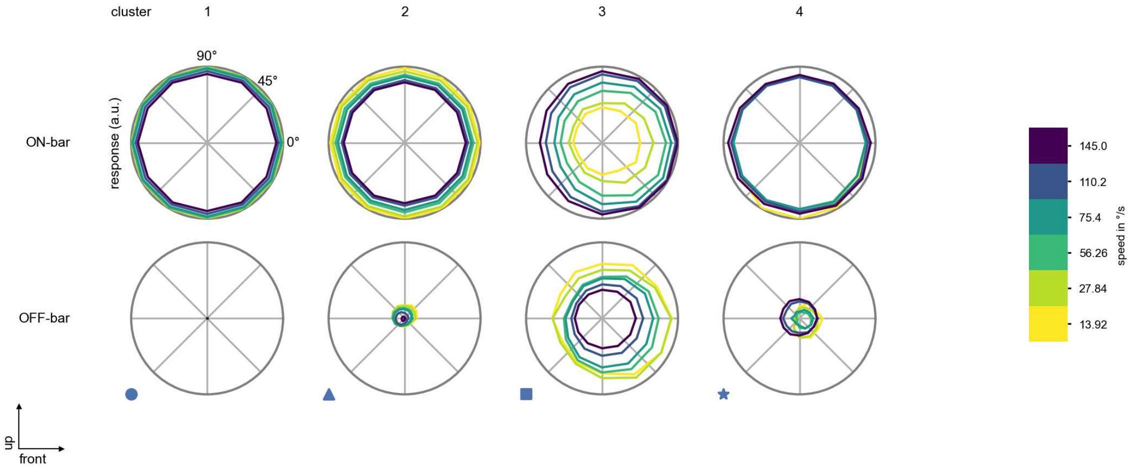

R1 - Figure 9: **Peak responses to moving bars from best-performing models.** The top row shows peak responses to moving ON-bars, the bottom row shows peak responses to moving OFF-bars. The bar-stimuli move in different directions from 0 to 360 degrees and at different speeds. Before, a grey-stimulus led the network to a stationary state. The columns show responses from the best-task-performing model in the respective cluster.

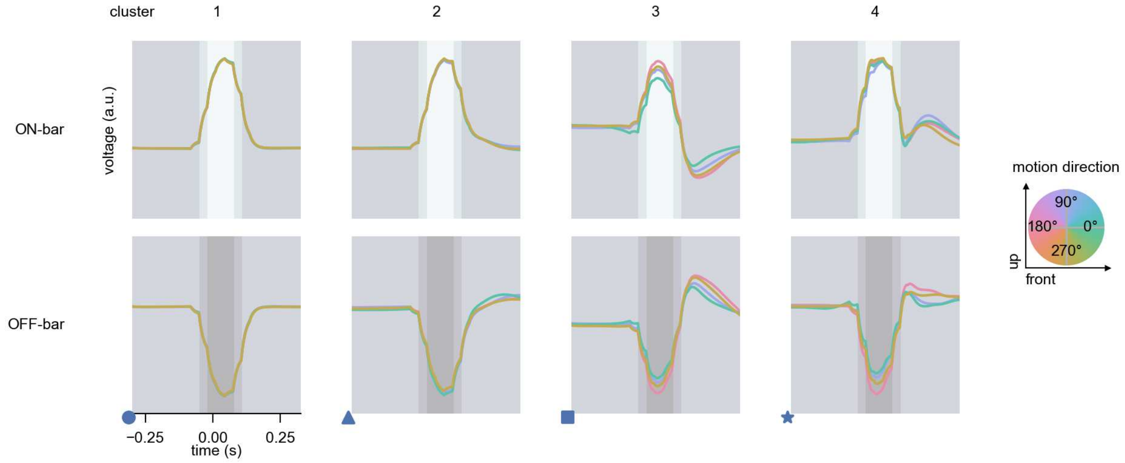

R1 - Figure 10: **Responses to moving bars from best-performing models.** The top row shows responses to moving ON-bars, the bottom row shows peak responses to moving OFF-bars. The bar-stimuli move in different directions from 0 to 360 degrees and at different speeds. Before, a grey-stimulus led the network to a stationary state. The columns show responses from the best-task-performing model in the respective cluster. Example responses to bars moving at 75.4°/s in all cardinal directions (green 0°, blue 90°, red 180°, yellow 270°) from -22.5 to 22.5° visual angle. Background contour indicates the bar, passing by the central column at second zero.

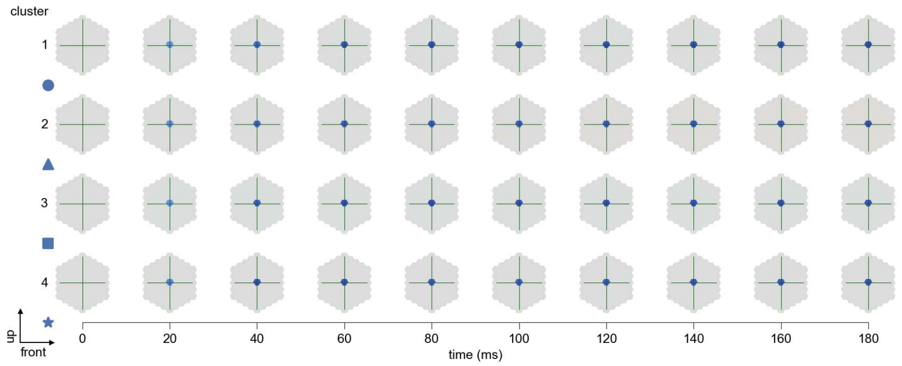

R1 - Figure 11: **Spatio-temporal receptive field.** Responses of the central cell to ON-impulses at single ommatidium impulse locations. The impulse occurs at second zero. Before, a grey-stimulus led the network to a stationary state. Responses are baseline-subtracted. Each row illustrates responses from the best-performing model of the respective cluster. Red indicated depolarization, blue indicates hyperpolarization.

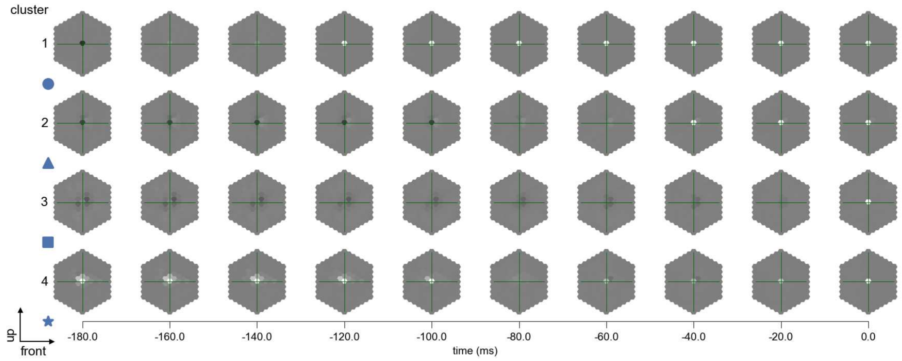

R1 - Figure 12: **Maximally excitatory stimuli.** Each row presents the regularized naturalistic-stimulus from the Sintel dataset that maximizes the cell type's central column response at second zero in the best-performing model of the respective cluster.

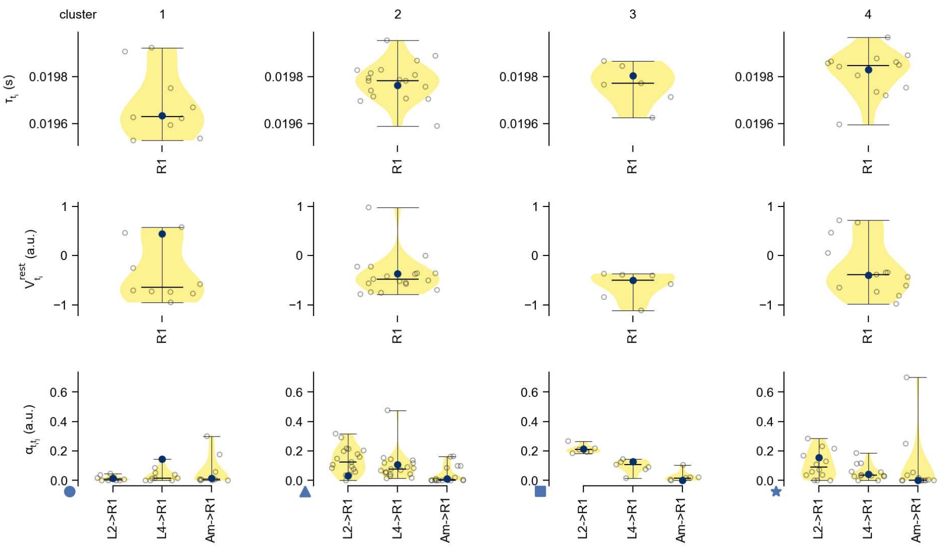

R1 - Figure 13: **Task-constrained parameters.** Each column shows the parameters inferred within the respective cluster. First row: learned time constants of the cell type. Second row: resting potentials of the cell type. Third row: scaling factors for the convolutional filters. The blue scatter represents the parameters from the best-performing model within the cluster.

# 2 R2

← Cell types

#### Figures

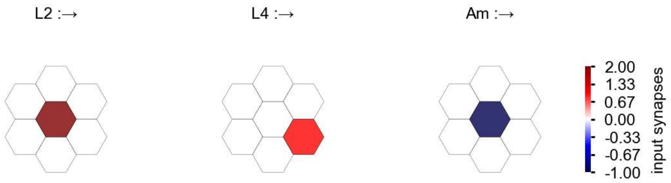

R2 - Figure 1: **Anatomical receptive fields.** Each colored hexagon is a projection, with the projection strength characterized by the average number of synapses that we count from the EM reconstruction. Red indicates excitatory synapses, blue indicates inhibitory synapses from inferred signs. Filters in the order of their total number of synapses.

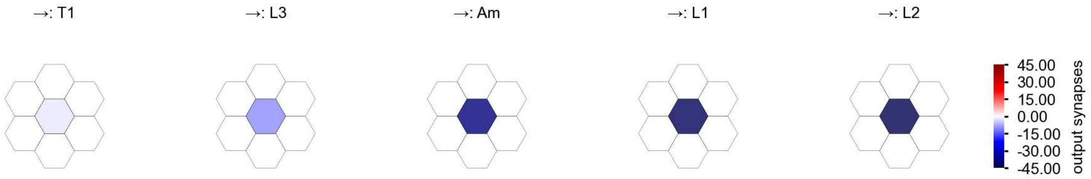

R2 - Figure 2: **Anatomical projective fields.** Each colored hexagon is a projection, with the projection strength characterized by the average number of synapses that we count from the EM reconstruction. Red indicates excitatory synapses, blue indicates inhibitory synapses from inferred signs. Filters in the order of their total number of synapses.

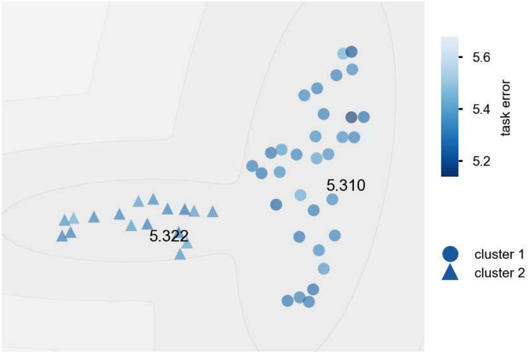

R2 - Figure 3: **Clustering of the responses to naturalistic stimuli.** From responses to naturalistic scenes from Sintel. Scatterpoints represent individual models colored by their task error.

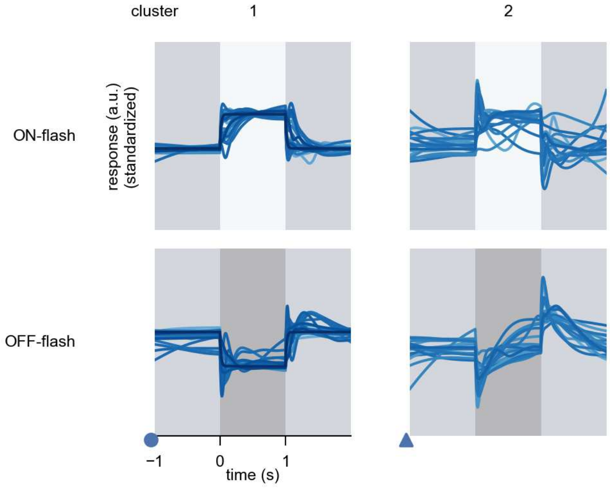

R2 - Figure 4: **Responses to flashes.** The top row shows responses to ON-flashes, the bottom row shows responses to OFF-flashes. The responses from the different models in the different clusters (columns) overlay and are colored in the respective task error of the model. The circular flashes occur at second zero, cover 6 ommatidia in radius, and last for one second. Before and after, a grey-stimulus leads to a stationary state of the network. The traces are standardized.

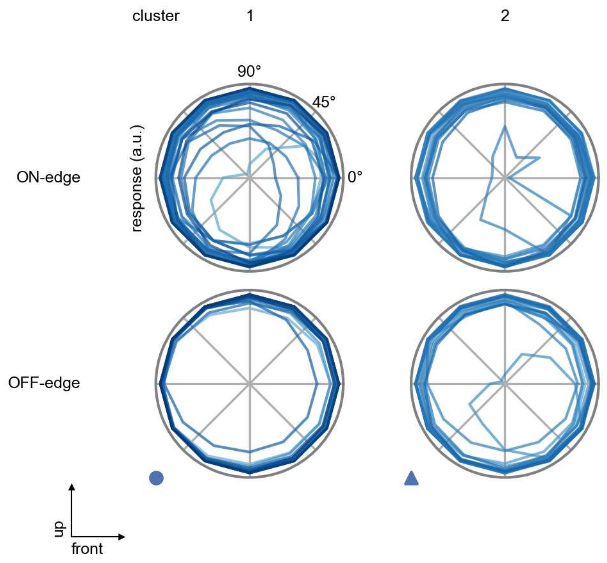

R2 - Figure 5: **Peak responses to moving edges.** The top row shows normalized peak responses to moving ON-edges, the bottom row shows normalized peak responses to moving OFF-edges. The edge-stimuli move in different directions from 0 to 360 degrees. Before, a grey-stimulus led the network to a stationary state. The responses from the different models in the different clusters (columns) overlay and are colored in the respective task error of the model. The peak responses are averaged over all assessed edge-speeds.

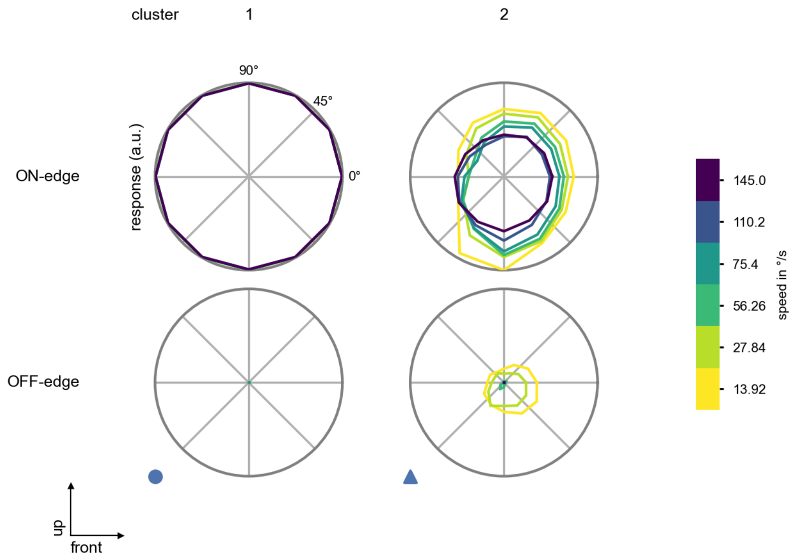

R2 - Figure 6: **Peak responses to moving edges from best-performing models.** The top row shows peak responses to moving ON-edges, the bottom row shows peak responses to moving OFF-edges. The edge-stimuli move in different directions from 0 to 360 degrees and at different speeds. Before, a grey-stimulus led the network to a stationary state. The columns show responses from the best-task-performing model in the respective cluster.

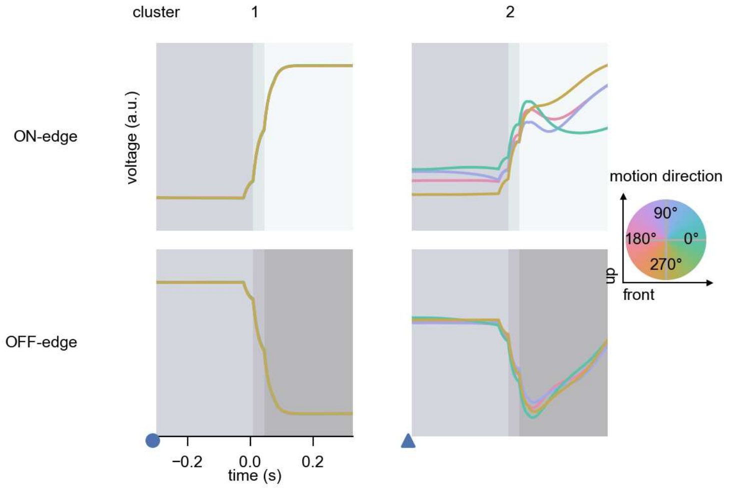

R2 - Figure 7: **Responses to moving edges from best-performing models.** The top row shows responses to moving ON-edges, the bottom row shows peak responses to moving OFF-edges. The edge-stimuli move in different directions from 0 to 360 degrees and at different speeds. Before, a grey-stimulus led the network to a stationary state. The columns show responses from the best-task-performing model in the respective cluster. Example responses to edges moving at 75.4°/s in all cardinal directions (green 0°, blue 90°, red 180°, yellow 270°) from -22.5 to 22.5° visual angle. Background contour indicates the edge, passing by the central column at second zero.

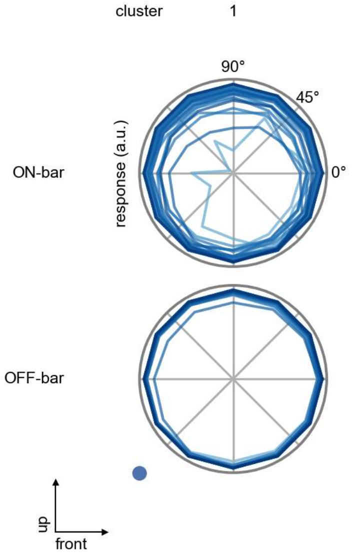

R2 - Figure 8: **Peak responses to moving bars.** The top row shows normalized peak responses to moving ON-bars, the bottom row shows normalized peak responses to moving OFF-bars. The bar-stimuli move in different directions from 0 to 360 degrees. Before, a grey-stimulus led the network to a stationary state. The responses from the different models in the different clusters (columns) overlay and are colored in the respective task error of the model. The peak responses are averaged over all assessed bar-speeds.

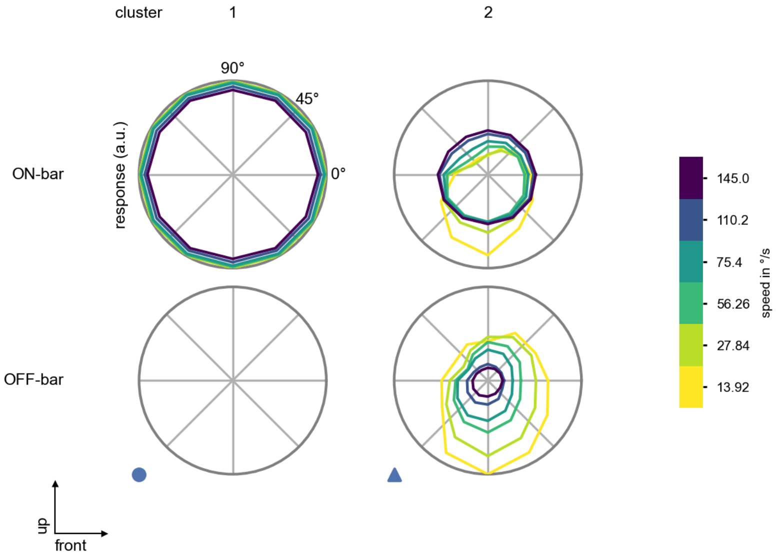

R2 - Figure 9: **Peak responses to moving bars from best-performing models.** The top row shows peak responses to moving ON-bars, the bottom row shows peak responses to moving OFF-bars. The bar-stimuli move in different directions from 0 to 360 degrees and at different speeds. Before, a grey-stimulus led the network to a stationary state. The columns show responses from the best-task-performing model in the respective cluster.

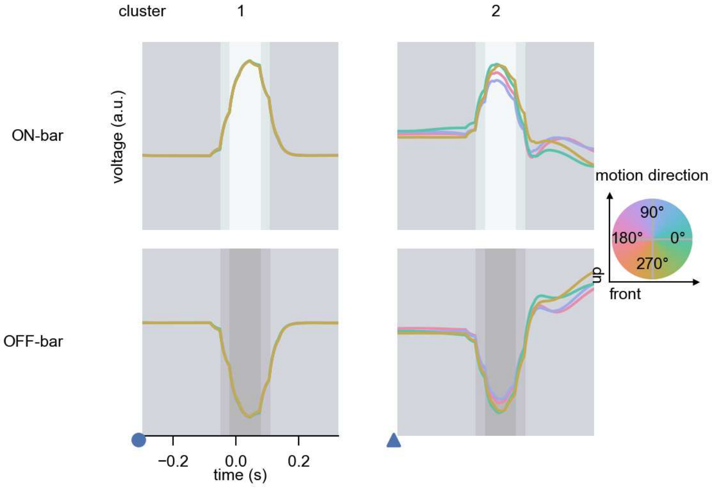

R2 - Figure 10: **Responses to moving bars from best-performing models.** The top row shows responses to moving ON-bars, the bottom row shows peak responses to moving OFF-bars. The bar-stimuli move in different directions from 0 to 360 degrees and at different speeds. Before, a grey-stimulus led the network to a stationary state. The columns show responses from the best-task-performing model in the respective cluster. Example responses to bars moving at 75.4°/s in all cardinal directions (green 0°, blue 90°, red 180°, yellow 270°) from -22.5 to 22.5° visual angle. Background contour indicates the bar, passing by the central column at second zero.

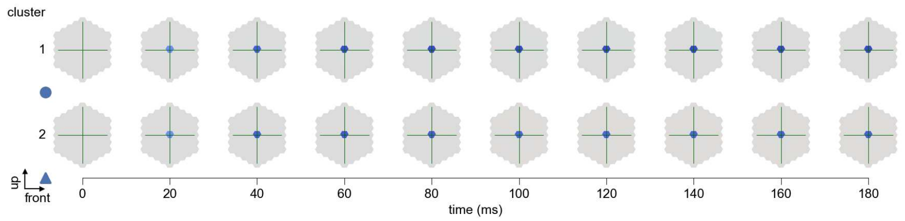

R2 - Figure 11: **Spatio-temporal receptive field.** Responses of the central cell to ON-impulses at single ommatidium impulse locations. The impulse occurs at second zero. Before, a grey-stimulus led the network to a stationary state. Responses are baseline-subtracted. Each row illustrates responses from the best-performing model of the respective cluster. Red indicated depolarization, blue indicates hyperpolarization.

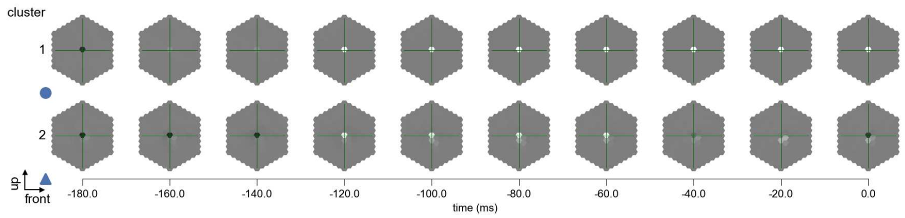

R2 - Figure 12: **Maximally excitatory stimuli.** Each row presents the regularized naturalistic-stimulus from the Sintel dataset that maximizes the cell type's central column response at second zero in the best-performing model of the respective cluster.

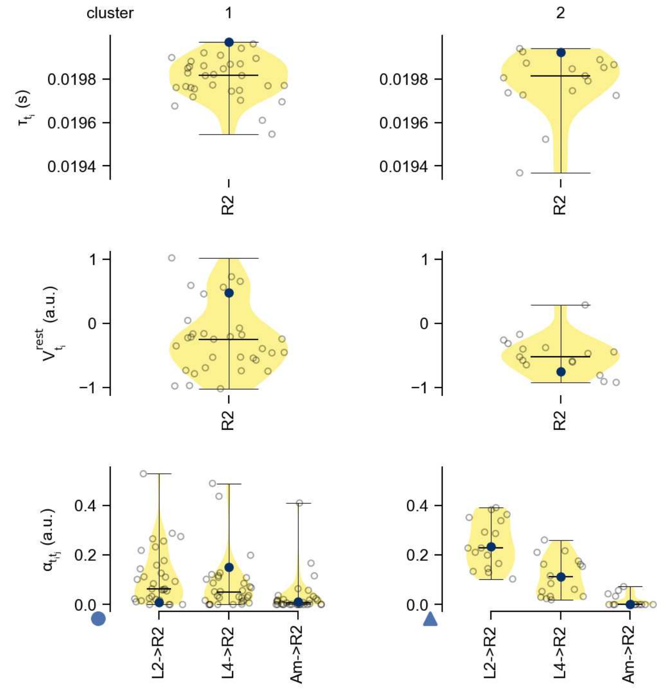

R2 - Figure 13: **Task-constrained parameters.** Each column shows the parameters inferred within the respective cluster. First row: learned time constants of the cell type. Second row: resting potentials of the cell type. Third row: scaling factors for the convolutional filters. The blue scatter represents the parameters from the best-performing model within the cluster.

## 3 R3

← Cell types

##### Figures

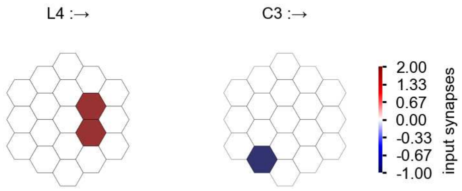

R3 - Figure 1: **Anatomical receptive fields.** Each colored hexagon is a projection, with the projection strength characterized by the average number of synapses that we count from the EM reconstruction. Red indicates excitatory synapses, blue indicates inhibitory synapses from inferred signs. Filters in the order of their total number of synapses.

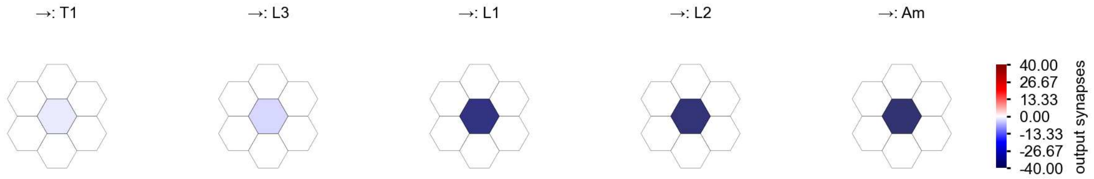

R3 - Figure 2: **Anatomical projective fields.** Each colored hexagon is a projection, with the projection strength characterized by the average number of synapses that we count from the EM reconstruction. Red indicates excitatory synapses, blue indicates inhibitory synapses from inferred signs. Filters in the order of their total number of synapses.

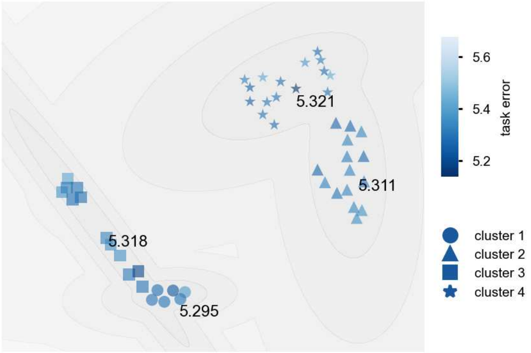

R3 - Figure 3: **Clustering of the responses to naturalistic stimuli.** From responses to naturalistic scenes from Sintel. Scatterpoints represent individual models colored by their task error.

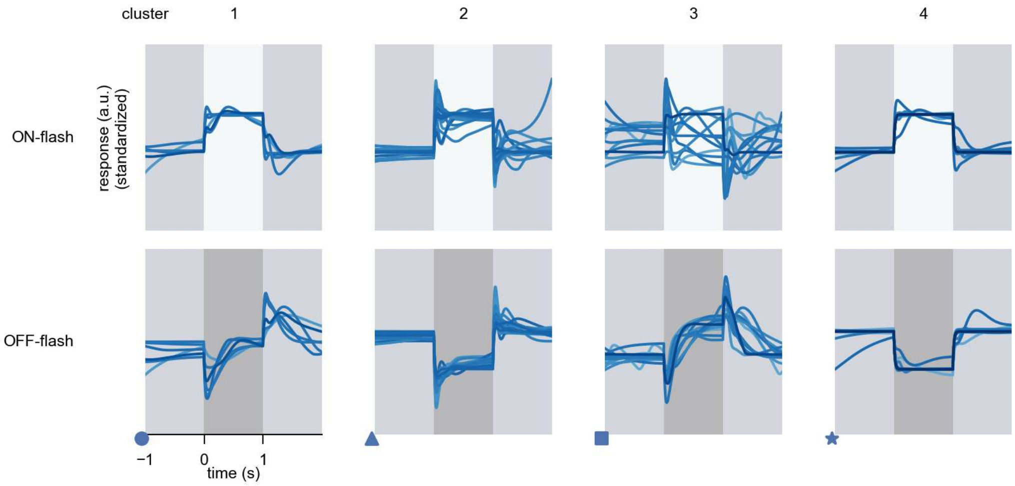

R3 - Figure 4: **Responses to flashes.** The top row shows responses to ON-flashes, the bottom row shows responses to OFF-flashes. The responses from the different models in the different clusters (columns) overlay and are colored in the respective task error of the model. The circular flashes occur at second zero, cover 6 ommatidia in radius, and last for one second. Before and after, a grey-stimulus leads to a stationary state of the network. The traces are standardized.

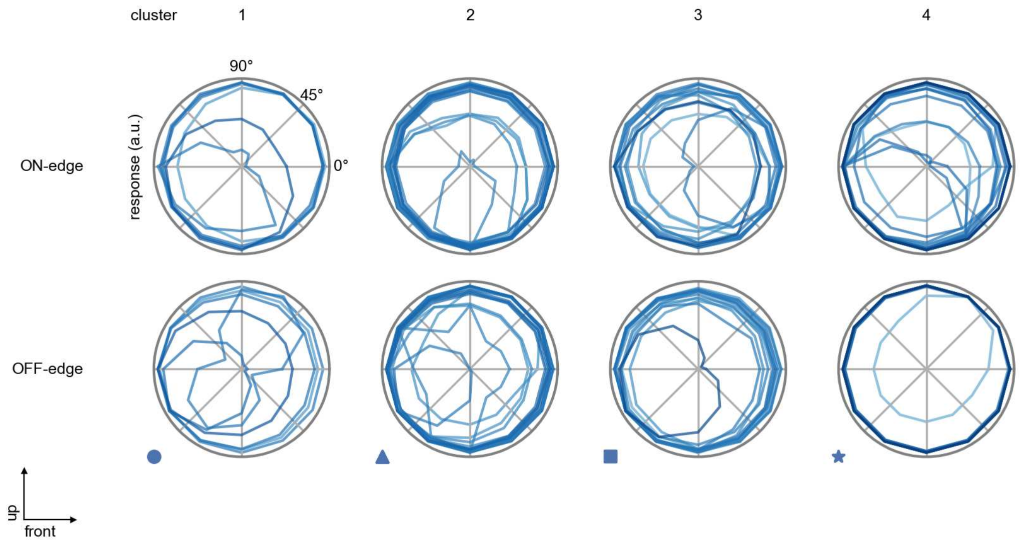

R3 - Figure 5: **Peak responses to moving edges.** The top row shows normalized peak responses to moving ON-edges, the bottom row shows normalized peak responses to moving OFF-edges. The edge-stimuli move in different directions from 0 to 360 degrees. Before, a grey-stimulus led the network to a stationary state. The responses from the different models in the different clusters (columns) overlay and are colored in the respective task error of the model. The peak responses are averaged over all assessed edge-speeds.

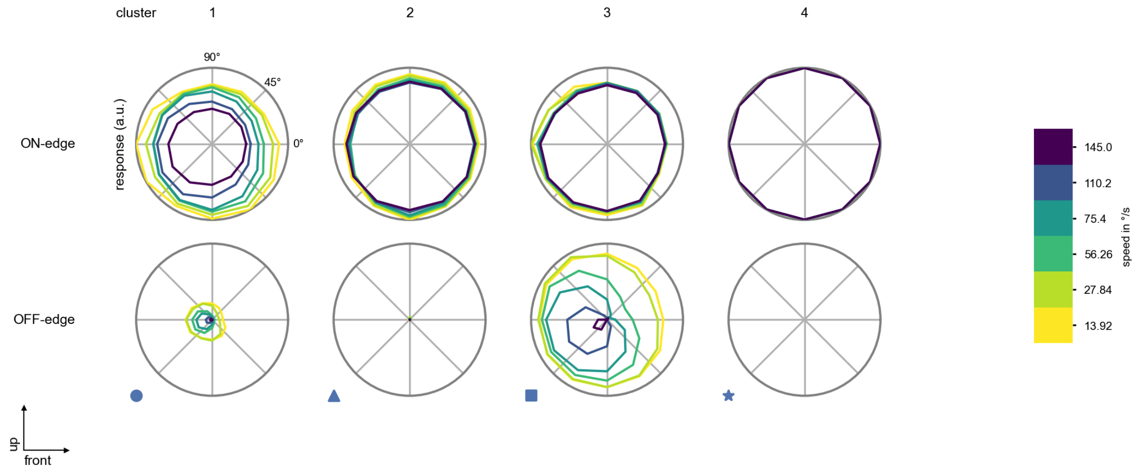

R3 - Figure 6: **Peak responses to moving edges from best-performing models.** The top row shows peak responses to moving ON-edges, the bottom row shows peak responses to moving OFF-edges. The edge-stimuli move in different directions from 0 to 360 degrees and at different speeds. Before, a grey-stimulus led the network to a stationary state. The columns show responses from the best-task-performing model in the respective cluster.

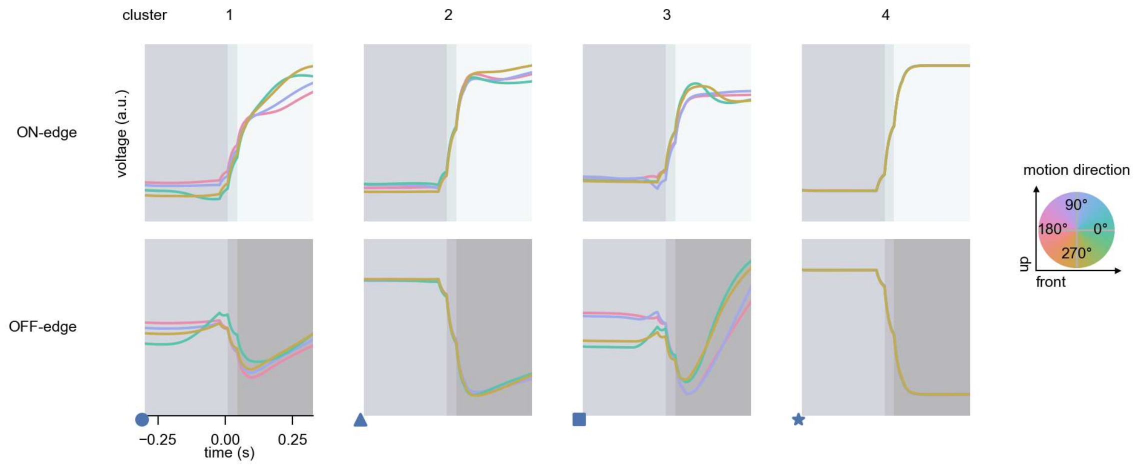

R3 - Figure 7: **Responses to moving edges from best-performing models.** The top row shows responses to moving ON-edges, the bottom row shows peak responses to moving OFF-edges. The edge-stimuli move in different directions from 0 to 360 degrees and at different speeds. Before, a grey-stimulus led the network to a stationary state. The columns show responses from the best-task-performing model in the respective cluster. Example responses to edges moving at 75.4°/s in all cardinal directions (green 0°, blue 90°, red 180°, yellow 270°) from -22.5 to 22.5° visual angle. Background contour indicates the edge, passing by the central column at second zero.

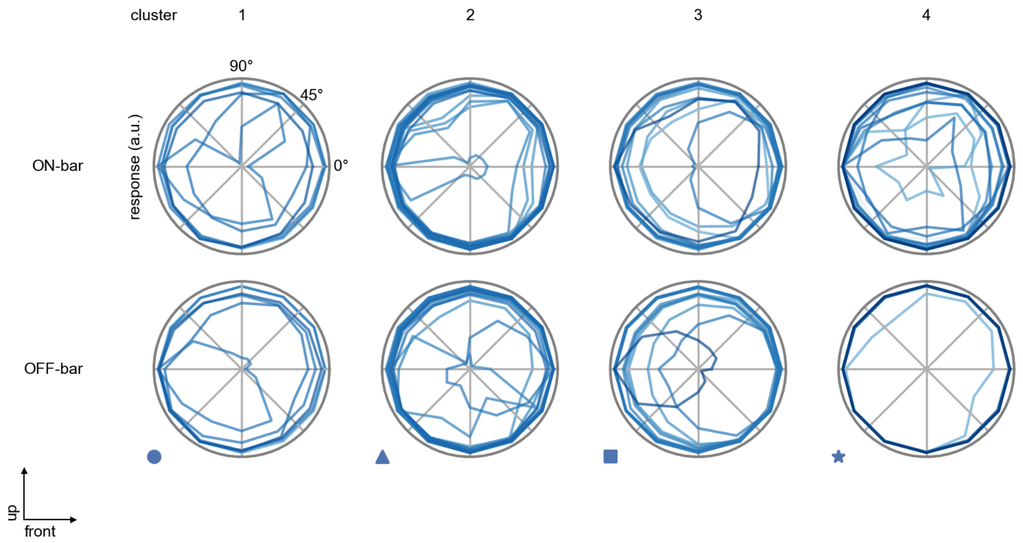

R3 - Figure 8: **Peak responses to moving bars.** The top row shows normalized peak responses to moving ON-bars, the bottom row shows normalized peak responses to moving OFF-bars. The bar-stimuli move in different directions from 0 to 360 degrees. Before, a grey-stimulus led the network to a stationary state. The responses from the different models in the different clusters (columns) overlay and are colored in the respective task error of the model. The peak responses are averaged over all assessed bar-speeds.

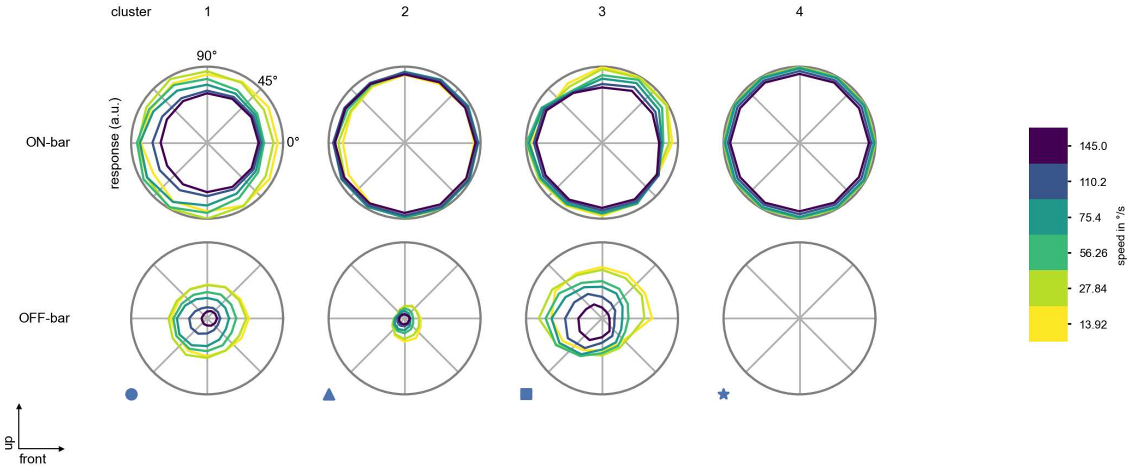

R3 - Figure 9: **Peak responses to moving bars from best-performing models.** The top row shows peak responses to moving ON-bars, the bottom row shows peak responses to moving OFF-bars. The bar-stimuli move in different directions from 0 to 360 degrees and at different speeds. Before, a grey-stimulus led the network to a stationary state. The columns show responses from the best-task-performing model in the respective cluster.

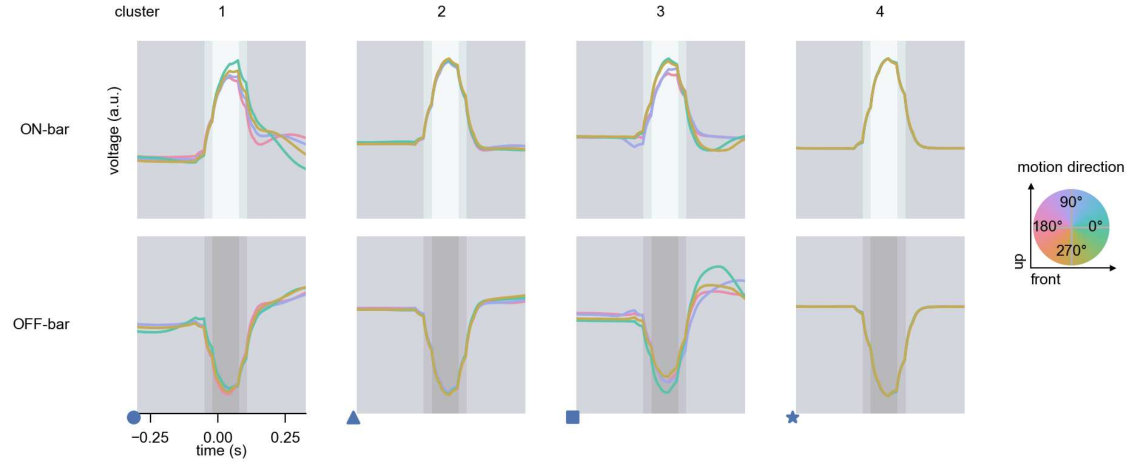

R3 - Figure 10: **Responses to moving bars from best-performing models.** The top row shows responses to moving ON-bars, the bottom row shows peak responses to moving OFF-bars. The bar-stimuli move in different directions from 0 to 360 degrees and at different speeds. Before, a grey-stimulus led the network to a stationary state. The columns show responses from the best-task-performing model in the respective cluster. Example responses to bars moving at 75.4°/s in all cardinal directions (green 0°, blue 90°, red 180°, yellow 270°) from -22.5 to 22.5° visual angle. Background contour indicates the bar, passing by the central column at second zero.

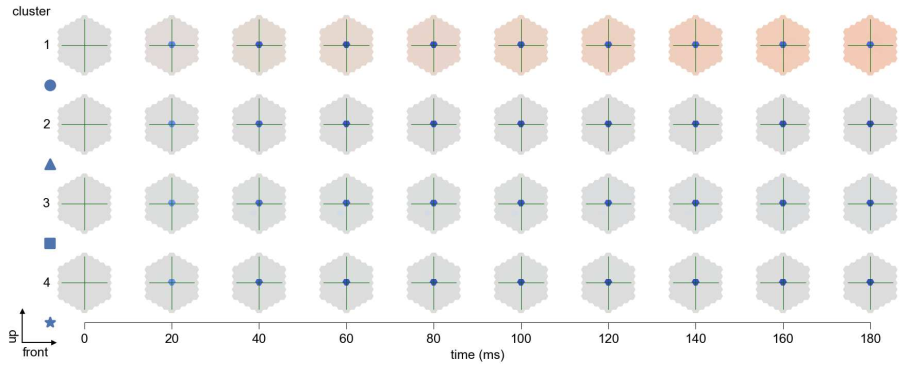

R3 - Figure 11: **Spatio-temporal receptive field.** Responses of the central cell to ON-impulses at single ommatidium impulse locations. The impulse occurs at second zero. Before, a grey-stimulus led the network to a stationary state. Responses are baseline-subtracted. Each row illustrates responses from the best-performing model of the respective cluster. Red indicated depolarization, blue indicates hyperpolarization.

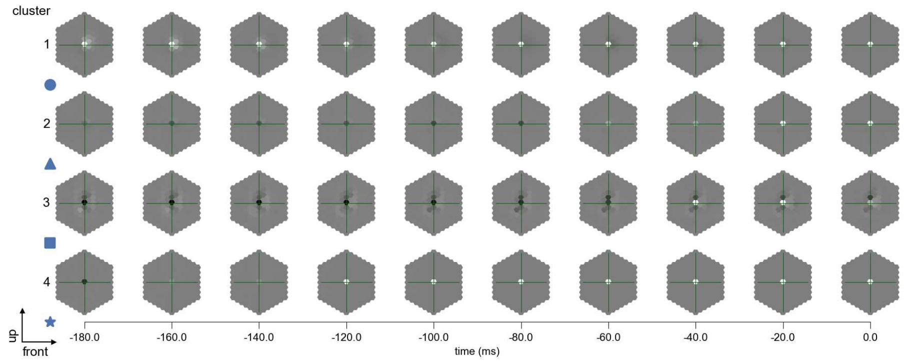

R3 - Figure 12: **Maximally excitatory stimuli.** Each row presents the regularized naturalistic-stimulus from the Sintel dataset that maximizes the cell type's central column response at second zero in the best-performing model of the respective cluster.

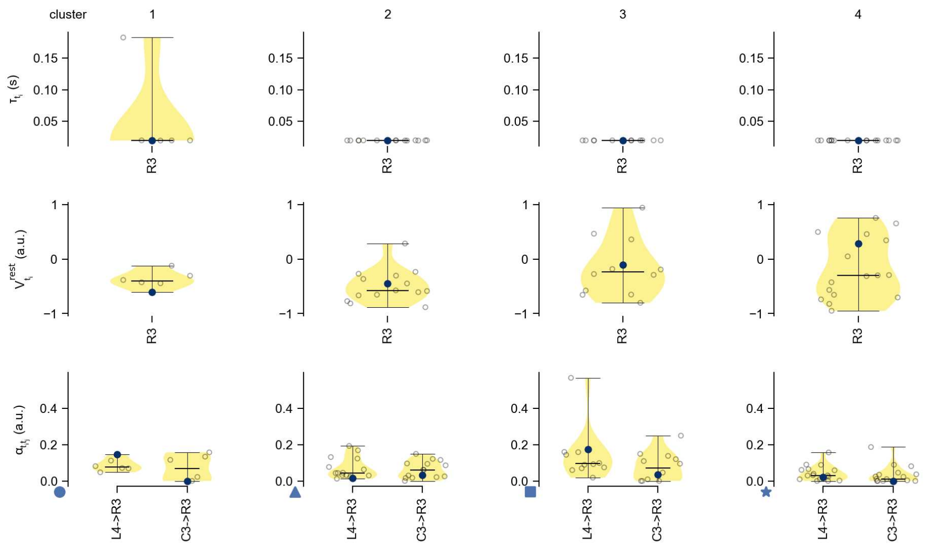

R3 - Figure 13: **Task-constrained parameters.** Each column shows the parameters inferred within the respective cluster. First row: learned time constants of the cell type. Second row: resting potentials of the cell type. Third row: scaling factors for the convolutional filters. The blue scatter represents the parameters from the best-performing model within the cluster.

# 4 R4

← Cell types

#### Figures

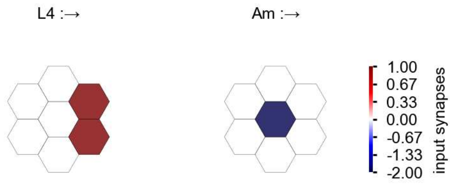

R4 - Figure 1: **Anatomical receptive fields.** Each colored hexagon is a projection, with the projection strength characterized by the average number of synapses that we count from the EM reconstruction. Red indicates excitatory synapses, blue indicates inhibitory synapses from inferred signs. Filters in the order of their total number of synapses.

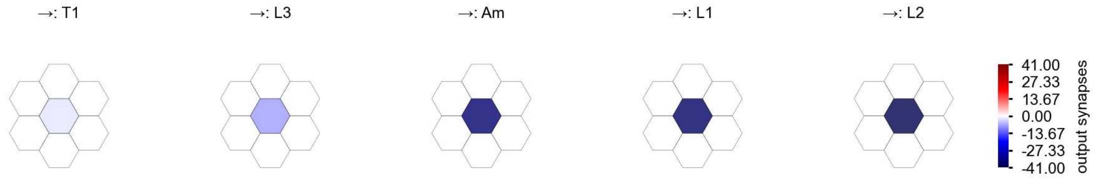

R4 - Figure 2: **Anatomical projective fields.** Each colored hexagon is a projection, with the projection strength characterized by the average number of synapses that we count from the EM reconstruction. Red indicates excitatory synapses, blue indicates inhibitory synapses from inferred signs. Filters in the order of their total number of synapses.

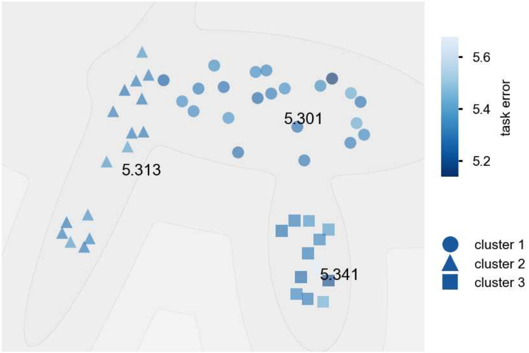

R4 - Figure 3: **Clustering of the responses to naturalistic stimuli.** From responses to naturalistic scenes from Sintel. Scatterpoints represent individual models colored by their task error.

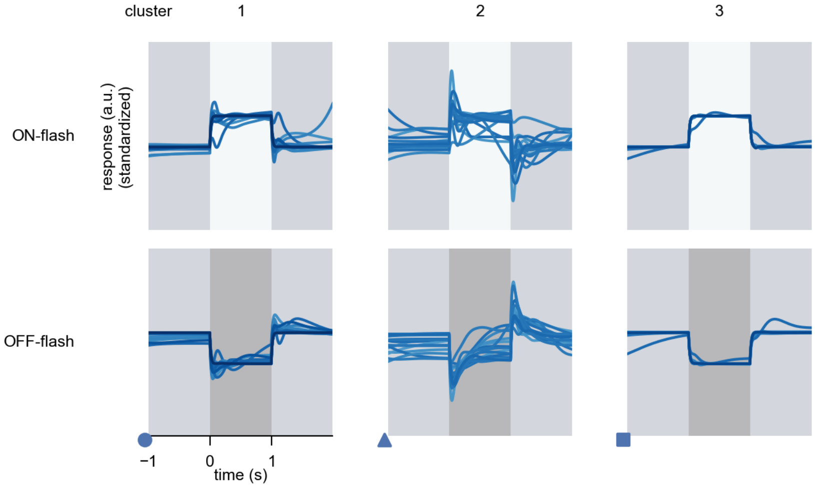

R4 - Figure 4: **Responses to flashes.** The top row shows responses to ON-flashes, the bottom row shows responses to OFF-flashes. The responses from the different models in the different clusters (columns) overlay and are colored in the respective task error of the model. The circular flashes occur at second zero, cover 6 ommatidia in radius, and last for one second. Before and after, a grey-stimulus leads to a stationary state of the network. The traces are standardized.

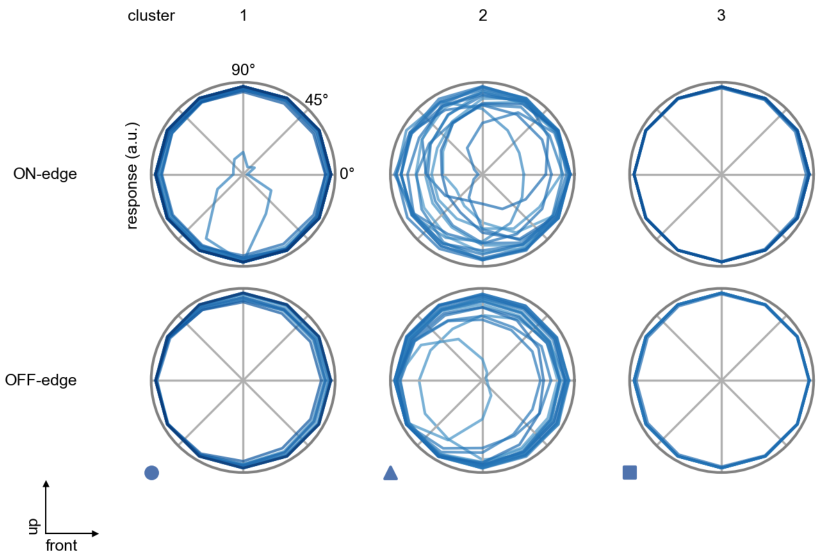

R4 - Figure 5: **Peak responses to moving edges.** The top row shows normalized peak responses to moving ON-edges, the bottom row shows normalized peak responses to moving OFF-edges. The edge-stimuli move in different directions from 0 to 360 degrees. Before, a grey-stimulus led the network to a stationary state. The responses from the different models in the different clusters (columns) overlay and are colored in the respective task error of the model. The peak responses are averaged over all assessed edge-speeds.

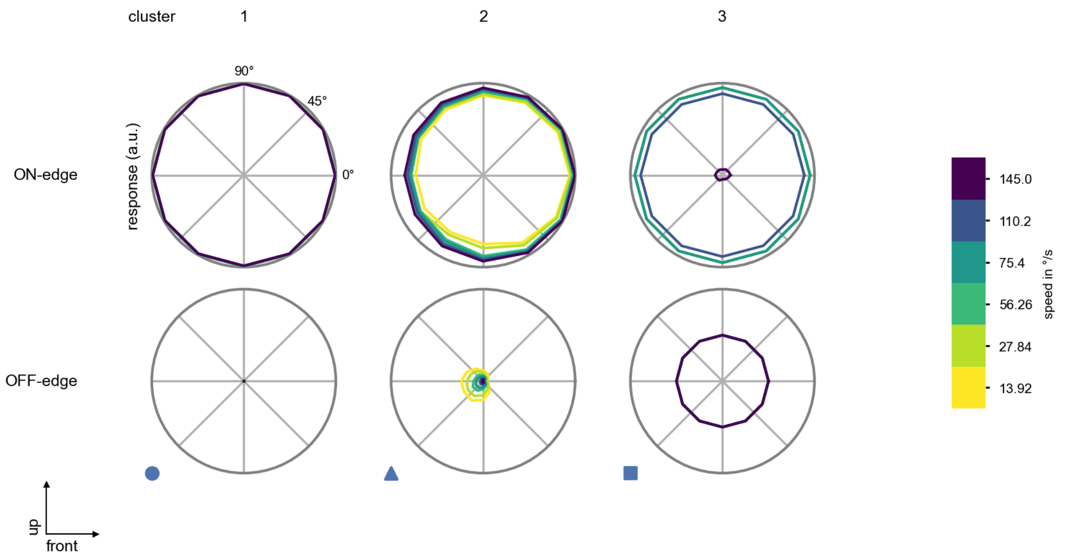

R4 - Figure 6: **Peak responses to moving edges from best-performing models.** The top row shows peak responses to moving ON-edges, the bottom row shows peak responses to moving OFF-edges. The edge-stimuli move in different directions from 0 to 360 degrees and at different speeds. Before, a grey-stimulus led the network to a stationary state. The columns show responses from the best-task-performing model in the respective cluster.

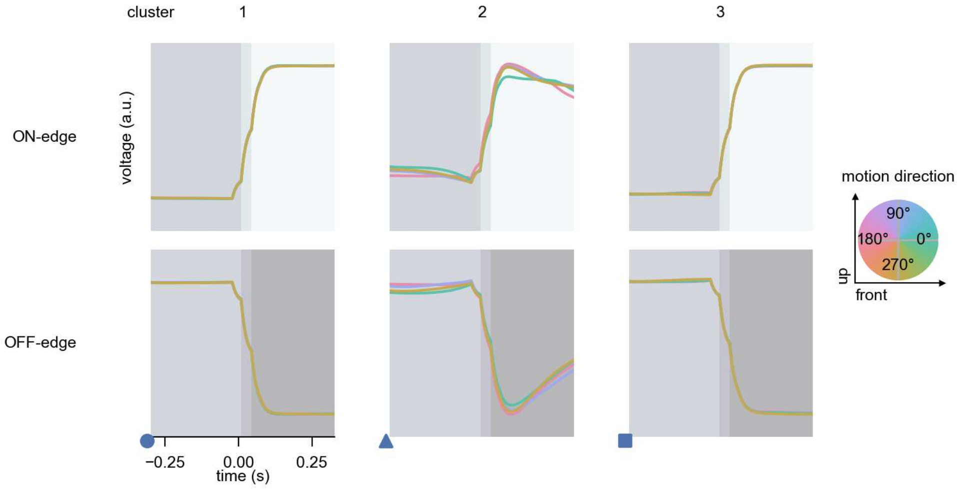

R4 - Figure 7: **Responses to moving edges from best-performing models.** The top row shows responses to moving ON-edges, the bottom row shows peak responses to moving OFF-edges. The edge-stimuli move in different directions from 0 to 360 degrees and at different speeds. Before, a grey-stimulus led the network to a stationary state. The columns show responses from the best-task-performing model in the respective cluster. Example responses to edges moving at 75.4°/s in all cardinal directions (green 0°, blue 90°, red 180°, yellow 270°) from -22.5 to 22.5° visual angle. Background contour indicates the edge, passing by the central column at second zero.

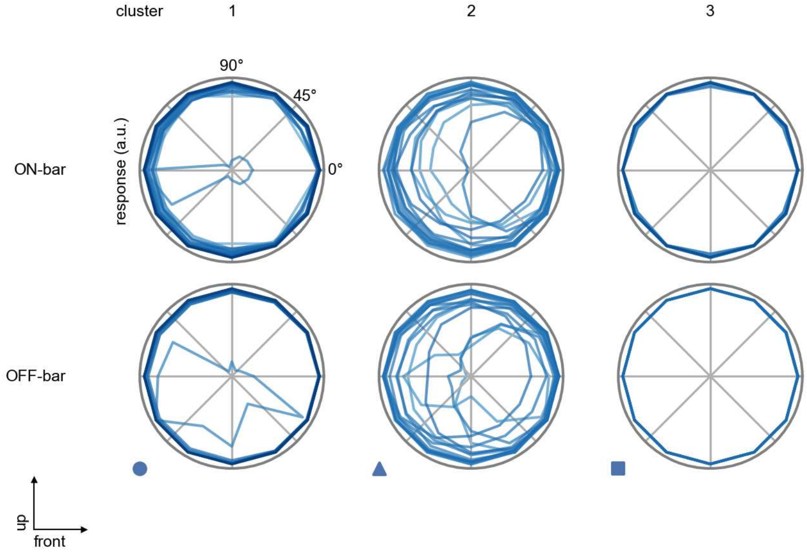

R4 - Figure 8: **Peak responses to moving bars.** The top row shows normalized peak responses to moving ON-bars, the bottom row shows normalized peak responses to moving OFF-bars. The bar-stimuli move in different directions from 0 to 360 degrees. Before, a grey-stimulus led the network to a stationary state. The responses from the different models in the different clusters (columns) overlay and are colored in the respective task error of the model. The peak responses are averaged over all assessed bar-speeds.

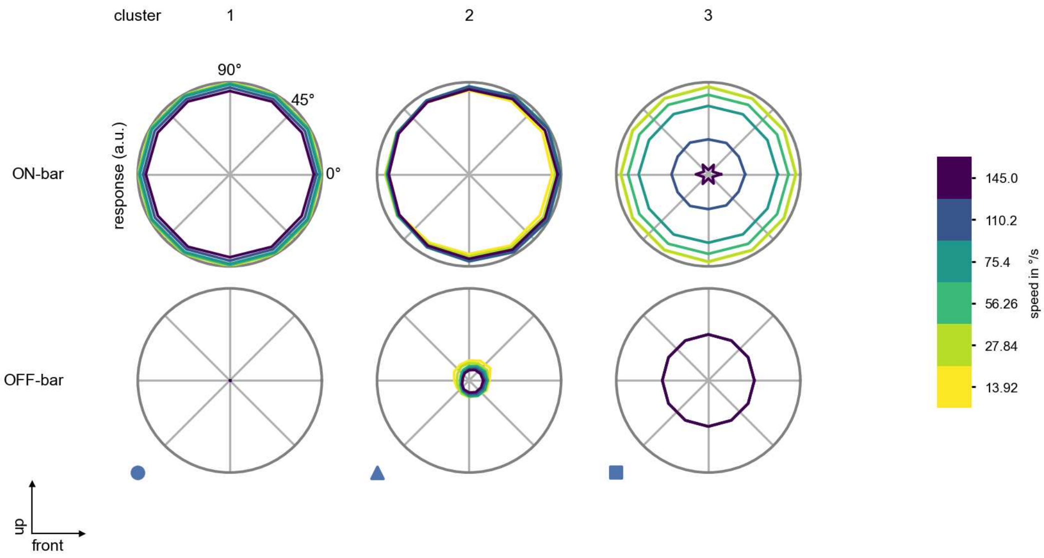

R4 - Figure 9: **Peak responses to moving bars from best-performing models.** The top row shows peak responses to moving ON-bars, the bottom row shows peak responses to moving OFF-bars. The bar-stimuli move in different directions from 0 to 360 degrees and at different speeds. Before, a grey-stimulus led the network to a stationary state. The columns show responses from the best-task-performing model in the respective cluster.

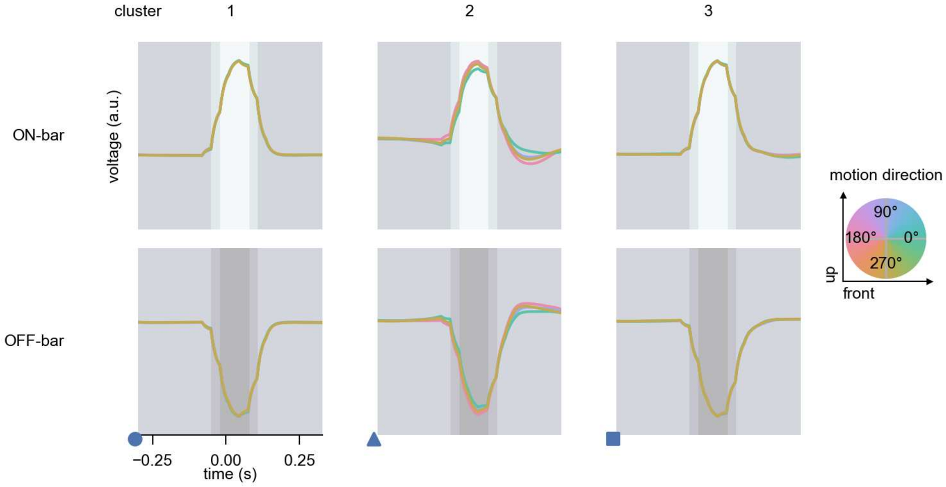

R4 - Figure 10: **Responses to moving bars from best-performing models.** The top row shows responses to moving ON-bars, the bottom row shows peak responses to moving OFF-bars. The bar-stimuli move in different directions from 0 to 360 degrees and at different speeds. Before, a grey-stimulus led the network to a stationary state. The columns show responses from the best-task-performing model in the respective cluster. Example responses to bars moving at 75.4°/s in all cardinal directions (green 0°, blue 90°, red 180°, yellow 270°) from -22.5 to 22.5° visual angle. Background contour indicates the bar, passing by the central column at second zero.

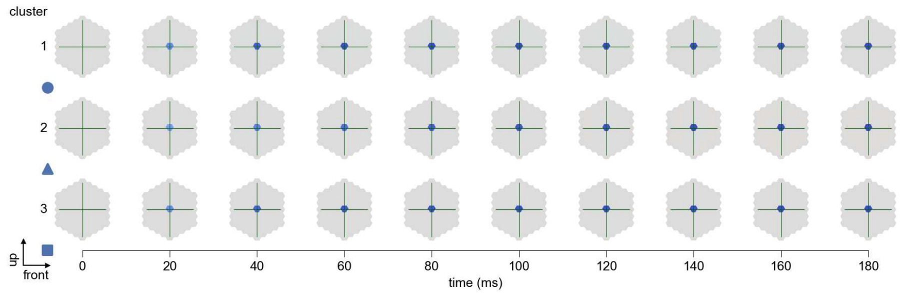

R4 - Figure 11: **Spatio-temporal receptive field.** Responses of the central cell to ON-impulses at single ommatidium impulse locations. The impulse occurs at second zero. Before, a grey-stimulus led the network to a stationary state. Responses are baseline-subtracted. Each row illustrates responses from the best-performing model of the respective cluster. Red indicated depolarization, blue indicates hyperpolarization.

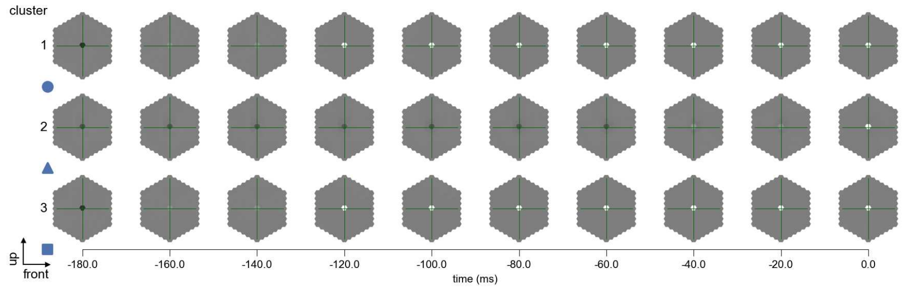

R4 - Figure 12: **Maximally excitatory stimuli.** Each row presents the regularized naturalistic-stimulus from the Sintel dataset that maximizes the cell type's central column response at second zero in the best-performing model of the respective cluster.

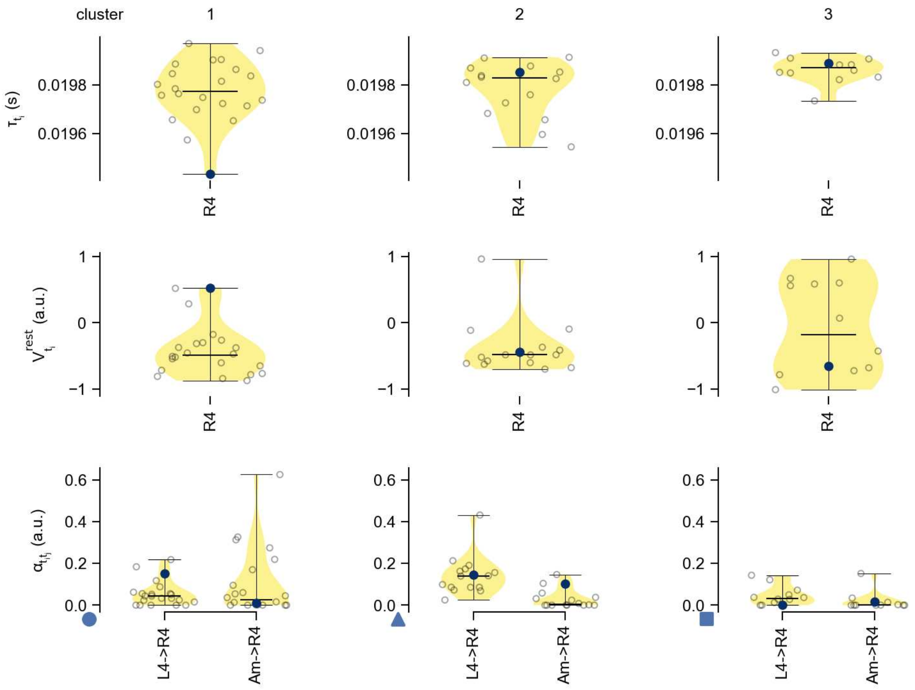

R4 - Figure 13: **Task-constrained parameters.** Each column shows the parameters inferred within the respective cluster. First row: learned time constants of the cell type. Second row: resting potentials of the cell type. Third row: scaling factors for the convolutional filters. The blue scatter represents the parameters from the best-performing model within the cluster.

# 5 R5

← Cell types

#### Figures

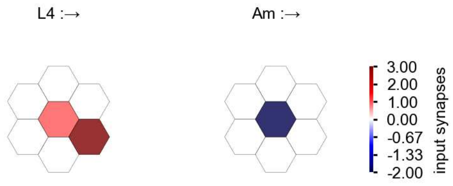

R5 - Figure 1: **Anatomical receptive fields.** Each colored hexagon is a projection, with the projection strength characterized by the average number of synapses that we count from the EM reconstruction. Red indicates excitatory synapses, blue indicates inhibitory synapses from inferred signs. Filters in the order of their total number of synapses.

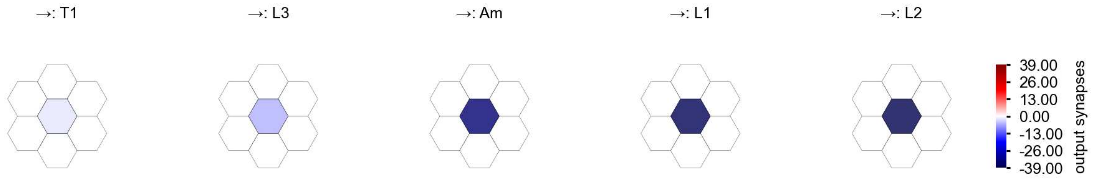

R5 - Figure 2: **Anatomical projective fields.** Each colored hexagon is a projection, with the projection strength characterized by the average number of synapses that we count from the EM reconstruction. Red indicates excitatory synapses, blue indicates inhibitory synapses from inferred signs. Filters in the order of their total number of synapses.

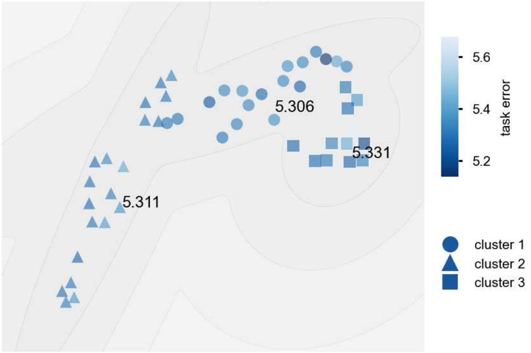

R5 - Figure 3: **Clustering of the responses to naturalistic stimuli.** From responses to naturalistic scenes from Sintel. Scatterpoints represent individual models colored by their task error.

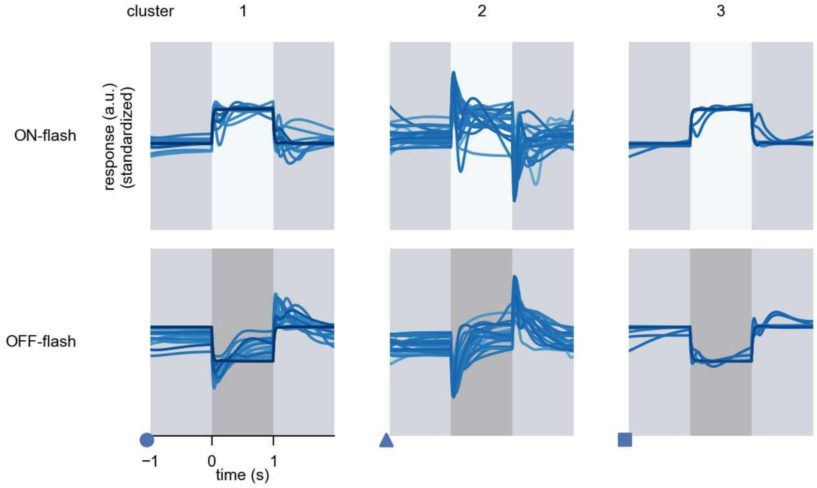

R5 - Figure 4: **Responses to flashes.** The top row shows responses to ON-flashes, the bottom row shows responses to OFF-flashes. The responses from the different models in the different clusters (columns) overlay and are colored in the respective task error of the model. The circular flashes occur at second zero, cover 6 ommatidia in radius, and last for one second. Before and after, a grey-stimulus leads to a stationary state of the network. The traces are standardized.

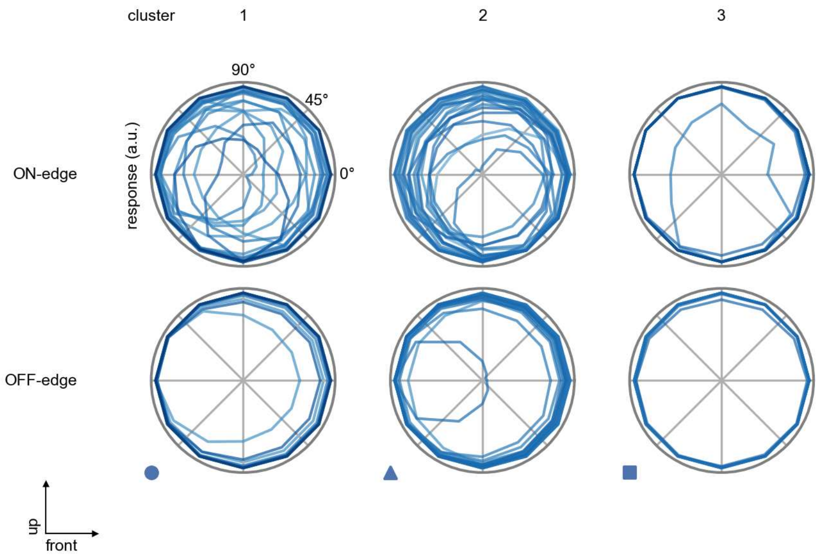

R5 - Figure 5: **Peak responses to moving edges.** The top row shows normalized peak responses to moving ON-edges, the bottom row shows normalized peak responses to moving OFF-edges. The edge-stimuli move in different directions from 0 to 360 degrees. Before, a grey-stimulus led the network to a stationary state. The responses from the different models in the different clusters (columns) overlay and are colored in the respective task error of the model. The peak responses are averaged over all assessed edge-speeds.

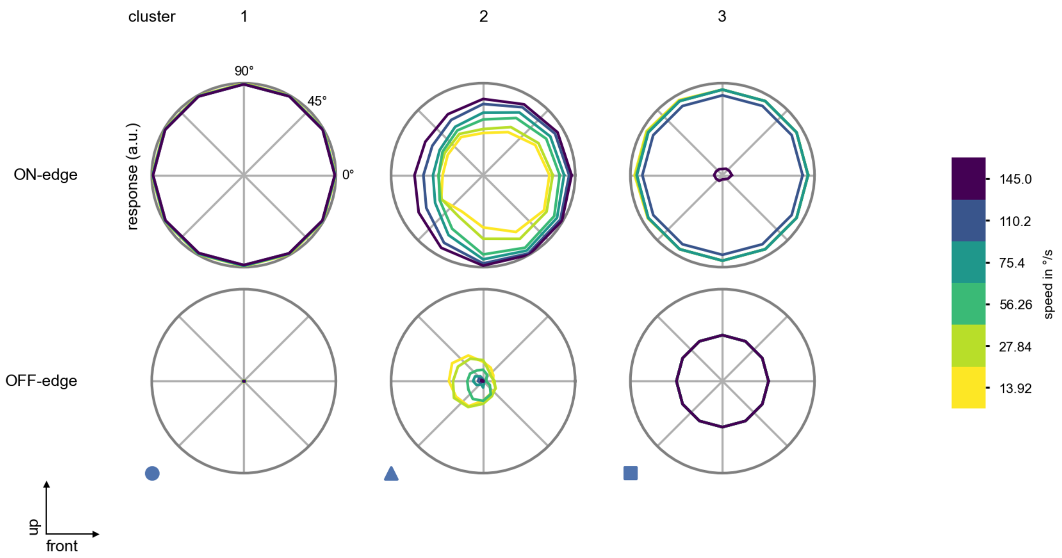

R5 - Figure 6: **Peak responses to moving edges from best-performing models.** The top row shows peak responses to moving ON-edges, the bottom row shows peak responses to moving OFF-edges. The edge-stimuli move in different directions from 0 to 360 degrees and at different speeds. Before, a grey-stimulus led the network to a stationary state. The columns show responses from the best-task-performing model in the respective cluster.

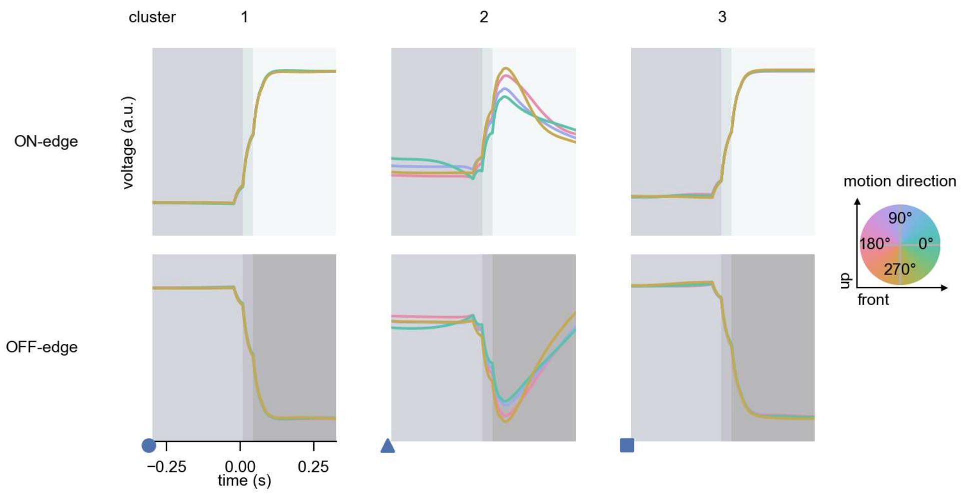

R5 - Figure 7: **Responses to moving edges from best-performing models.** The top row shows responses to moving ON-edges, the bottom row shows peak responses to moving OFF-edges. The edge-stimuli move in different directions from 0 to 360 degrees and at different speeds. Before, a grey-stimulus led the network to a stationary state. The columns show responses from the best-task-performing model in the respective cluster. Example responses to edges moving at 75.4°/s in all cardinal directions (green 0°, blue 90°, red 180°, yellow 270°) from -22.5 to 22.5° visual angle. Background contour indicates the edge, passing by the central column at second zero.

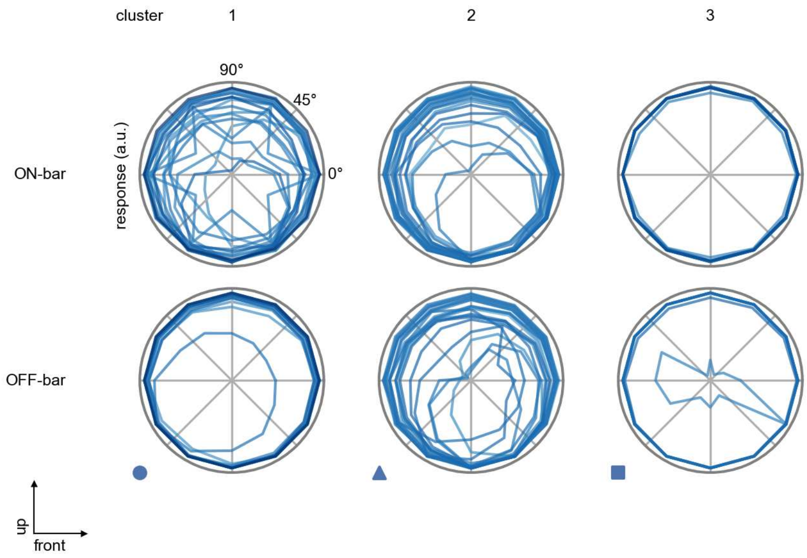

R5 - Figure 8: **Peak responses to moving bars.** The top row shows normalized peak responses to moving ON-bars, the bottom row shows normalized peak responses to moving OFF-bars. The bar-stimuli move in different directions from 0 to 360 degrees. Before, a grey-stimulus led the network to a stationary state. The responses from the different models in the different clusters (columns) overlay and are colored in the respective task error of the model. The peak responses are averaged over all assessed bar-speeds.

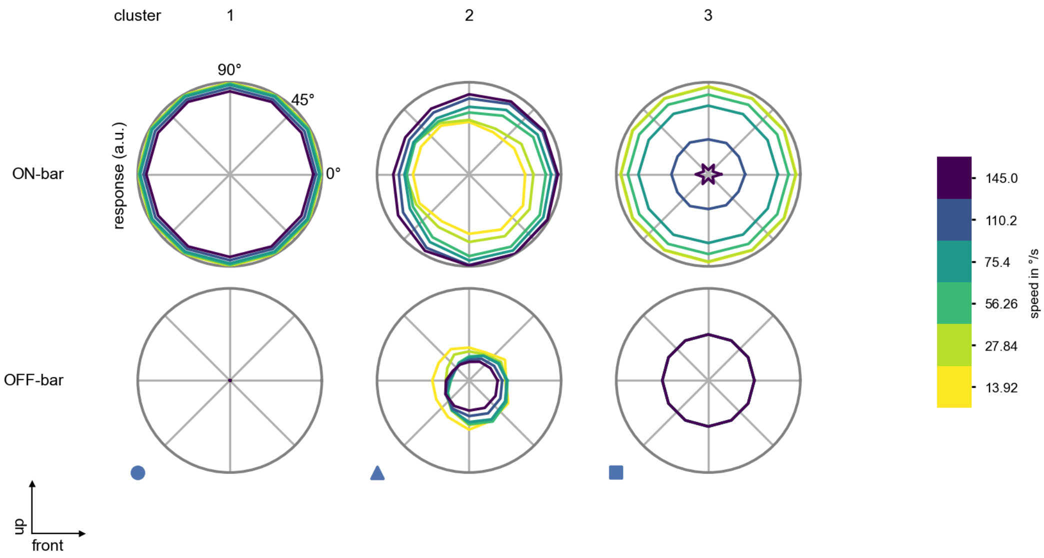

R5 - Figure 9: **Peak responses to moving bars from best-performing models.** The top row shows peak responses to moving ON-bars, the bottom row shows peak responses to moving OFF-bars. The bar-stimuli move in different directions from 0 to 360 degrees and at different speeds. Before, a grey-stimulus led the network to a stationary state. The columns show responses from the best-task-performing model in the respective cluster.

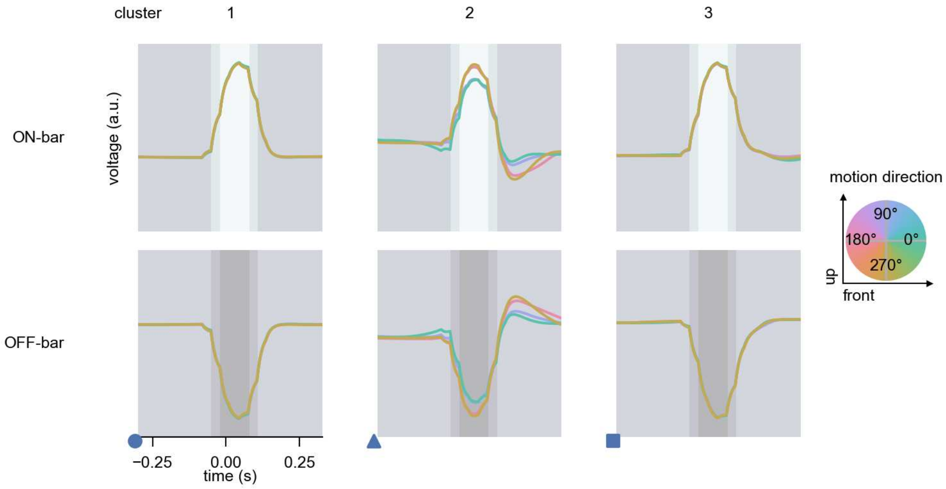

R5 - Figure 10: **Responses to moving bars from best-performing models.** The top row shows responses to moving ON-bars, the bottom row shows peak responses to moving OFF-bars. The bar-stimuli move in different directions from 0 to 360 degrees and at different speeds. Before, a grey-stimulus led the network to a stationary state. The columns show responses from the best-task-performing model in the respective cluster. Example responses to bars moving at 75.4°/s in all cardinal directions (green 0°, blue 90°, red 180°, yellow 270°) from -22.5 to 22.5° visual angle. Background contour indicates the bar, passing by the central column at second zero.

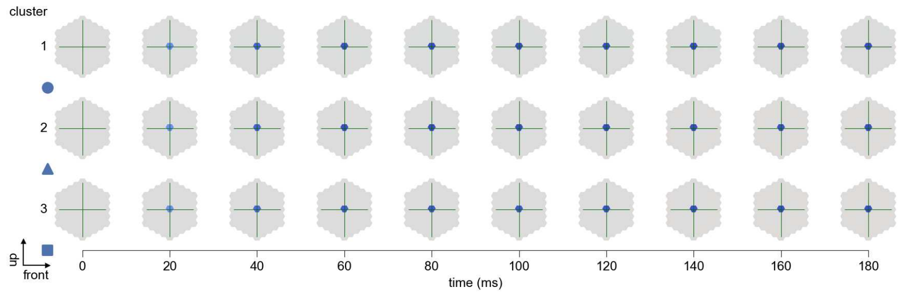

R5 - Figure 11: **Spatio-temporal receptive field.** Responses of the central cell to ON-impulses at single ommatidium impulse locations. The impulse occurs at second zero. Before, a grey-stimulus led the network to a stationary state. Responses are baseline-subtracted. Each row illustrates responses from the best-performing model of the respective cluster. Red indicated depolarization, blue indicates hyperpolarization.

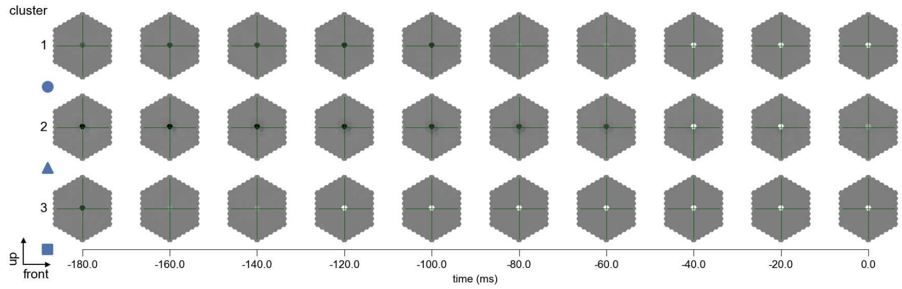

R5 - Figure 12: **Maximally excitatory stimuli.** Each row presents the regularized naturalistic-stimulus from the Sintel dataset that maximizes the cell type's central column response at second zero in the best-performing model of the respective cluster.

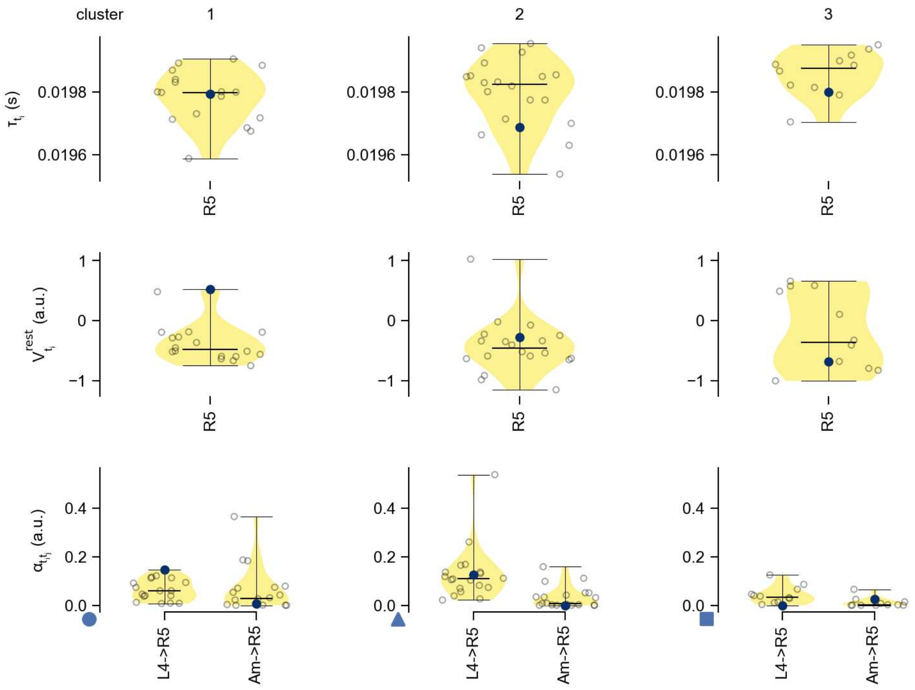

R5 - Figure 13: **Task-constrained parameters.** Each column shows the parameters inferred within the respective cluster. First row: learned time constants of the cell type. Second row: resting potentials of the cell type. Third row: scaling factors for the convolutional filters. The blue scatter represents the parameters from the best-performing model within the cluster.

# 6 R6

← Cell types

#### Figures

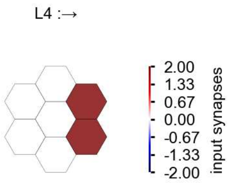

R6 - Figure 1: **Anatomical receptive fields.** Each colored hexagon is a projection, with the projection strength characterized by the average number of synapses that we count from the EM reconstruction. Red indicates excitatory synapses, blue indicates inhibitory synapses from inferred signs. Filters in the order of their total number of synapses.

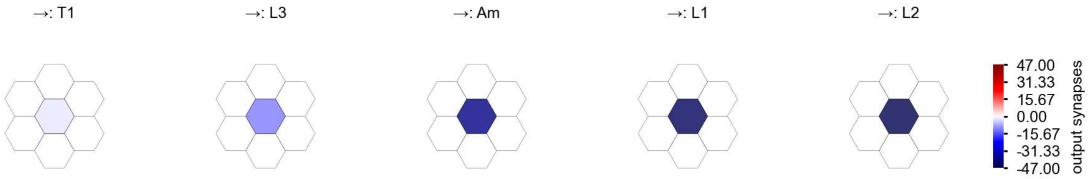

R6 - Figure 2: **Anatomical projective fields.** Each colored hexagon is a projection, with the projection strength characterized by the average number of synapses that we count from the EM reconstruction. Red indicates excitatory synapses, blue indicates inhibitory synapses from inferred signs. Filters in the order of their total number of synapses.

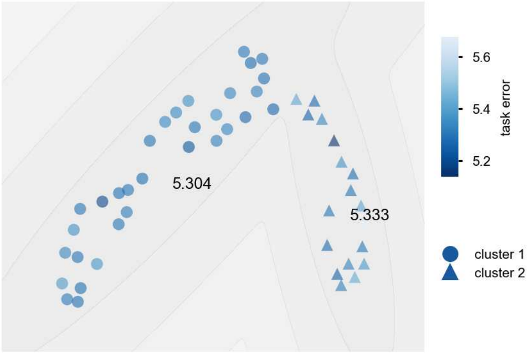

R6 - Figure 3: **Clustering of the responses to naturalistic stimuli.** From responses to naturalistic scenes from Sintel. Scatterpoints represent individual models colored by their task error.

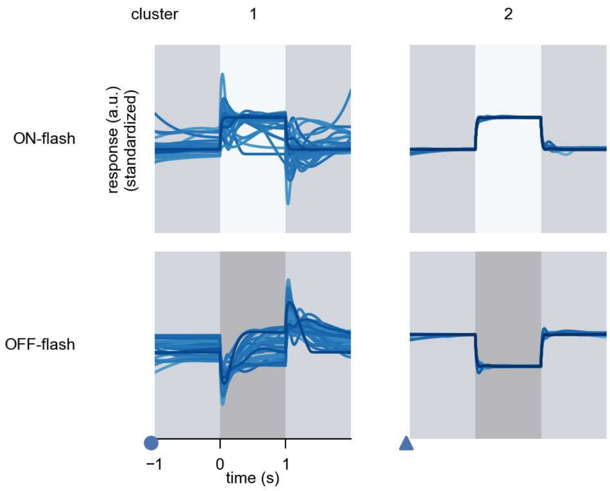

R6 - Figure 4: **Responses to flashes.** The top row shows responses to ON-flashes, the bottom row shows responses to OFF-flashes. The responses from the different models in the different clusters (columns) overlay and are colored in the respective task error of the model. The circular flashes occur at second zero, cover 6 ommatidia in radius, and last for one second. Before and after, a grey-stimulus leads to a stationary state of the network. The traces are standardized.

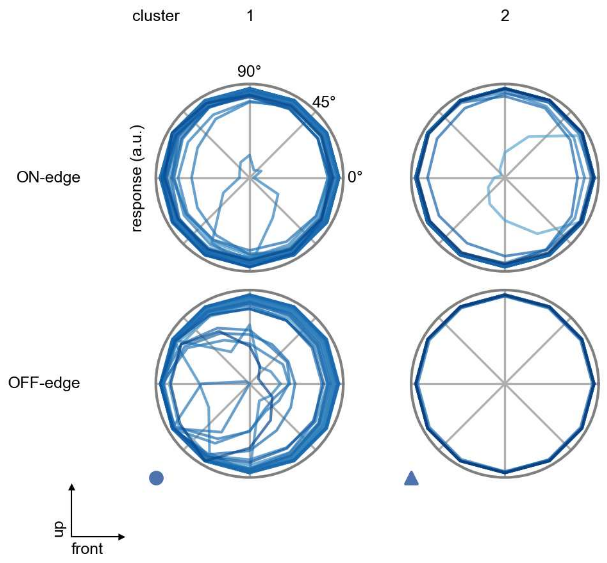

R6 - Figure 5: **Peak responses to moving edges.** The top row shows normalized peak responses to moving ON-edges, the bottom row shows normalized peak responses to moving OFF-edges. The edge-stimuli move in different directions from 0 to 360 degrees. Before, a grey-stimulus led the network to a stationary state. The responses from the different models in the different clusters (columns) overlay and are colored in the respective task error of the model. The peak responses are averaged over all assessed edge-speeds.

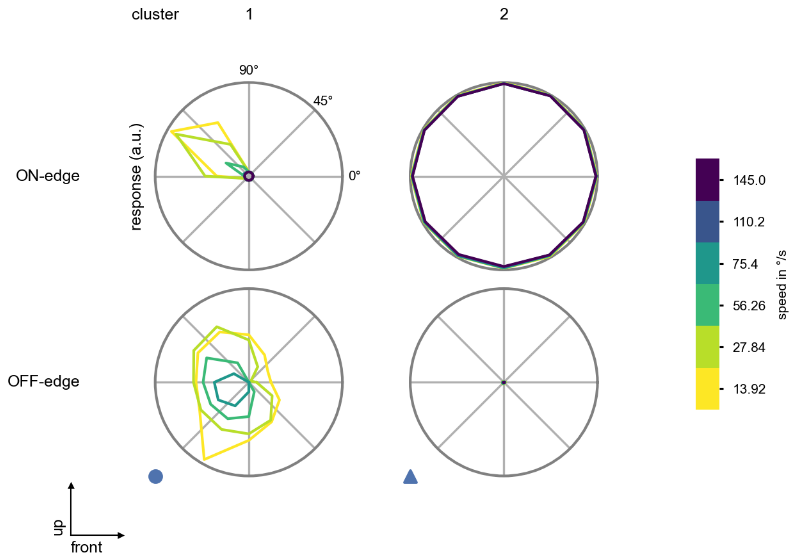

R6 - Figure 6: **Peak responses to moving edges from best-performing models.** The top row shows peak responses to moving ON-edges, the bottom row shows peak responses to moving OFF-edges. The edge-stimuli move in different directions from 0 to 360 degrees and at different speeds. Before, a grey-stimulus led the network to a stationary state. The columns show responses from the best-task-performing model in the respective cluster.

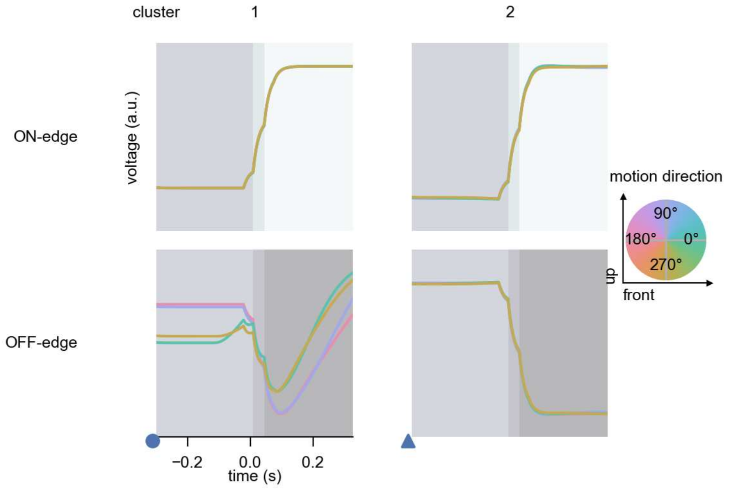

R6 - Figure 7: **Responses to moving edges from best-performing models.** The top row shows responses to moving ON-edges, the bottom row shows peak responses to moving OFF-edges. The edge-stimuli move in different directions from 0 to 360 degrees and at different speeds. Before, a grey-stimulus led the network to a stationary state. The columns show responses from the best-task-performing model in the respective cluster. Example responses to edges moving at 75.4°/s in all cardinal directions (green 0°, blue 90°, red 180°, yellow 270°) from -22.5 to 22.5° visual angle. Background contour indicates the edge, passing by the central column at second zero.

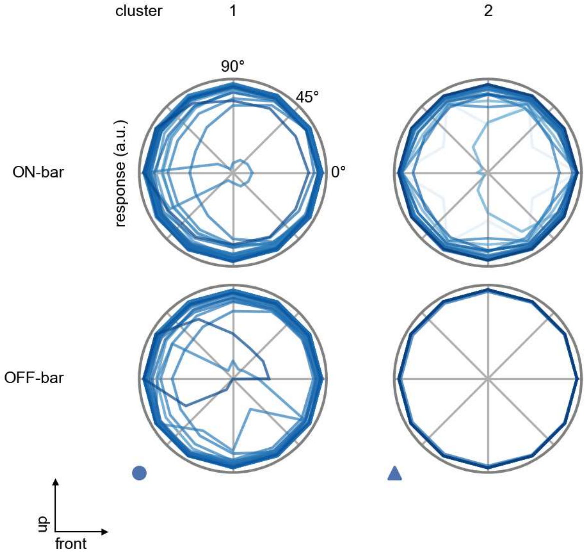

R6 - Figure 8: **Peak responses to moving bars.** The top row shows normalized peak responses to moving ON-bars, the bottom row shows normalized peak responses to moving OFF-bars. The bar-stimuli move in different directions from 0 to 360 degrees. Before, a grey-stimulus led the network to a stationary state. The responses from the different models in the different clusters (columns) overlay and are colored in the respective task error of the model. The peak responses are averaged over all assessed bar-speeds.

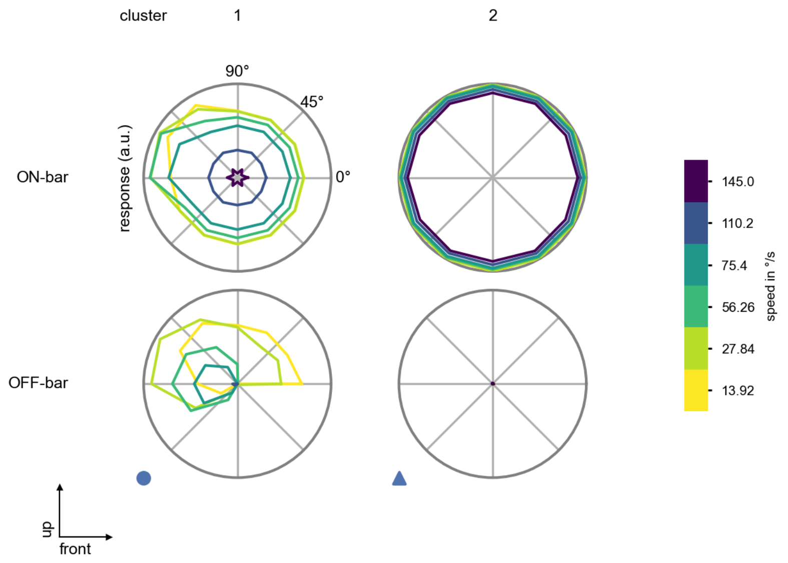

R6 - Figure 9: **Peak responses to moving bars from best-performing models.** The top row shows peak responses to moving ON-bars, the bottom row shows peak responses to moving OFF-bars. The bar-stimuli move in different directions from 0 to 360 degrees and at different speeds. Before, a grey-stimulus led the network to a stationary state. The columns show responses from the best-task-performing model in the respective cluster.

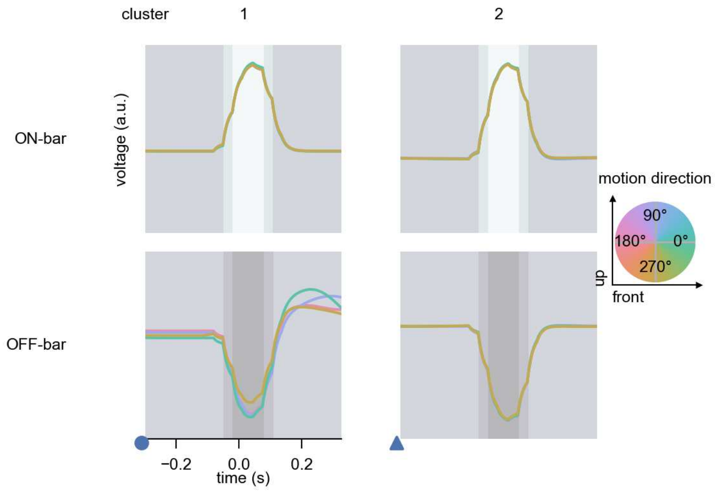

R6 - Figure 10: **Responses to moving bars from best-performing models.** The top row shows responses to moving ON-bars, the bottom row shows peak responses to moving OFF-bars. The bar-stimuli move in different directions from 0 to 360 degrees and at different speeds. Before, a grey-stimulus led the network to a stationary state. The columns show responses from the best-task-performing model in the respective cluster. Example responses to bars moving at 75.4°/s in all cardinal directions (green 0°, blue 90°, red 180°, yellow 270°) from -22.5 to 22.5° visual angle. Background contour indicates the bar, passing by the central column at second zero.

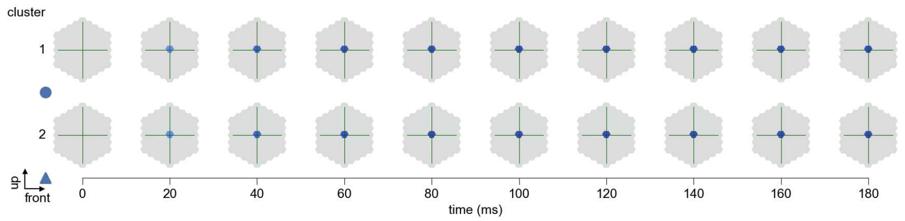

R6 - Figure 11: **Spatio-temporal receptive field.** Responses of the central cell to ON-impulses at single ommatidium impulse locations. The impulse occurs at second zero. Before, a grey-stimulus led the network to a stationary state. Responses are baseline-subtracted. Each row illustrates responses from the best-performing model of the respective cluster. Red indicated depolarization, blue indicates hyperpolarization.

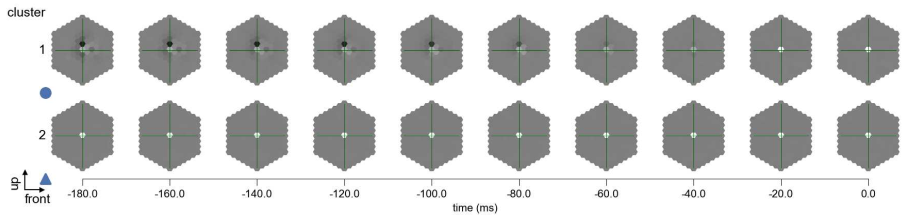

R6 - Figure 12: **Maximally excitatory stimuli.** Each row presents the regularized naturalistic-stimulus from the Sintel dataset that maximizes the cell type's central column response at second zero in the best-performing model of the respective cluster.

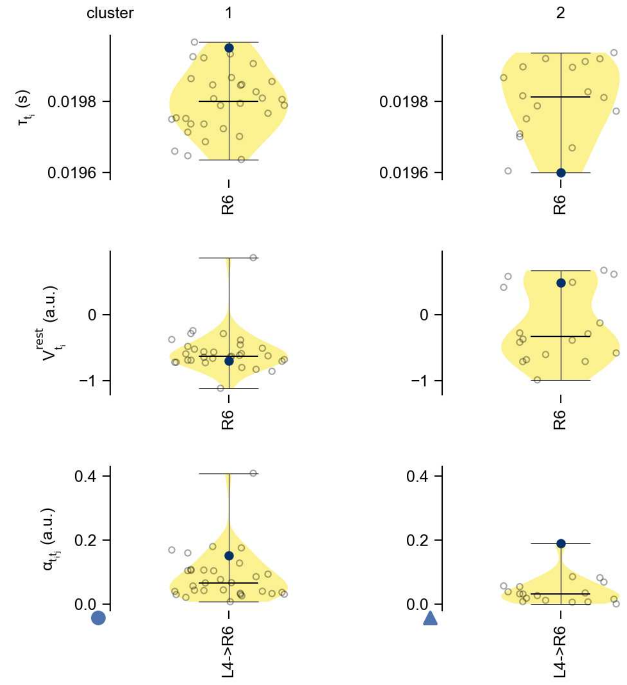

R6 - Figure 13: **Task-constrained parameters.** Each column shows the parameters inferred within the respective cluster. First row: learned time constants of the cell type. Second row: resting potentials of the cell type. Third row: scaling factors for the convolutional filters. The blue scatter represents the parameters from the best-performing model within the cluster.

# 7 R7

← Cell types

#### Figures

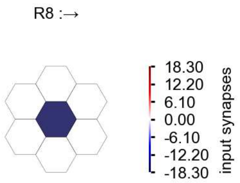

R7 - Figure 1: **Anatomical receptive fields.** Each colored hexagon is a projection, with the projection strength characterized by the average number of synapses that we count from the EM reconstruction. Red indicates excitatory synapses, blue indicates inhibitory synapses from inferred signs. Filters in the order of their total number of synapses.

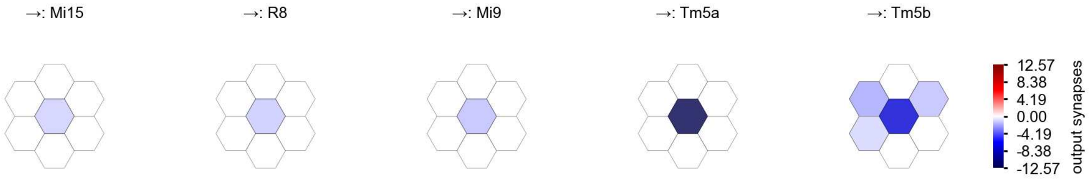

R7 - Figure 2: **Anatomical projective fields.** Each colored hexagon is a projection, with the projection strength characterized by the average number of synapses that we count from the EM reconstruction. Red indicates excitatory synapses, blue indicates inhibitory synapses from inferred signs. Filters in the order of their total number of synapses.

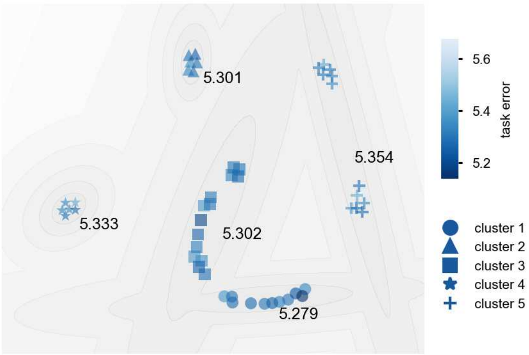

R7 - Figure 3: **Clustering of the responses to naturalistic stimuli.** From responses to naturalistic scenes from Sintel. Scatterpoints represent individual models colored by their task error.

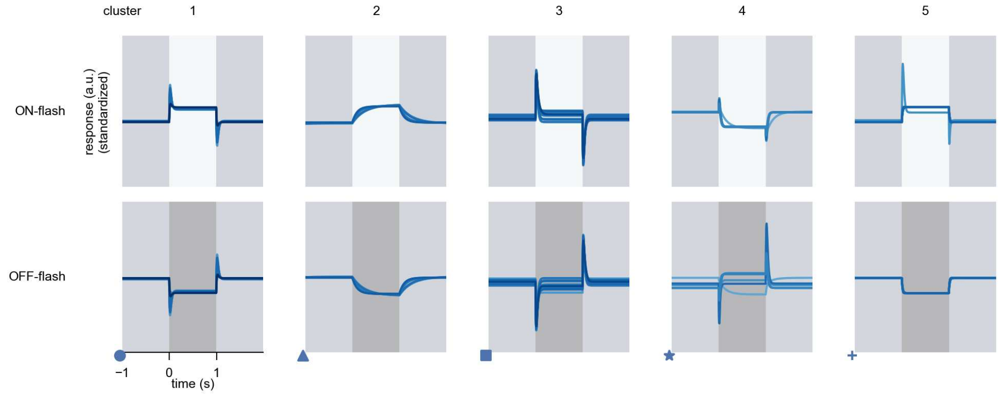

R7 - Figure 4: **Responses to flashes.** The top row shows responses to ON-flashes, the bottom row shows responses to OFF-flashes. The responses from the different models in the different clusters (columns) overlay and are colored in the respective task error of the model. The circular flashes occur at second zero, cover 6 ommatidia in radius, and last for one second. Before and after, a grey-stimulus leads to a stationary state of the network. The traces are standardized.

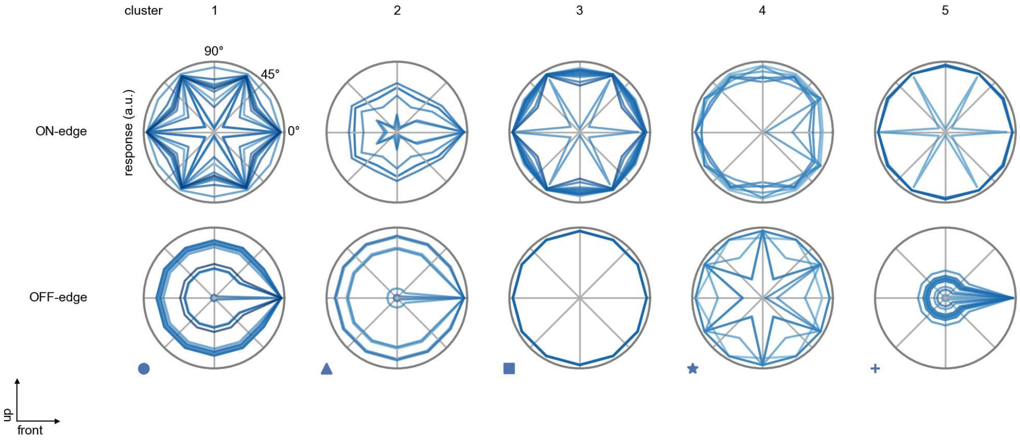

R7 - Figure 5: **Peak responses to moving edges.** The top row shows normalized peak responses to moving ON-edges, the bottom row shows normalized peak responses to moving OFF-edges. The edge-stimuli move in different directions from 0 to 360 degrees. Before, a grey-stimulus led the network to a stationary state. The responses from the different models in the different clusters (columns) overlay and are colored in the respective task error of the model. The peak responses are averaged over all assessed edge-speeds.

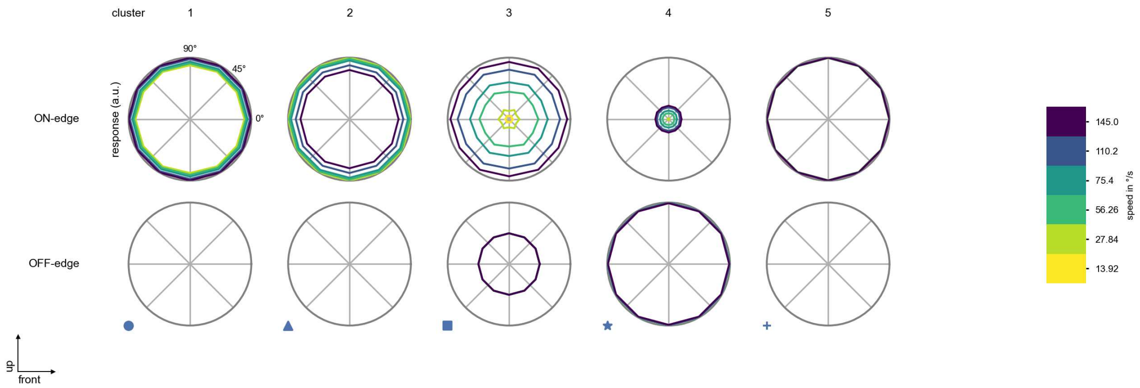

R7 - Figure 6: **Peak responses to moving edges from best-performing models.** The top row shows peak responses to moving ON-edges, the bottom row shows peak responses to moving OFF-edges. The edge-stimuli move in different directions from 0 to 360 degrees and at different speeds. Before, a grey-stimulus led the network to a stationary state. The columns show responses from the best-task-performing model in the respective cluster.

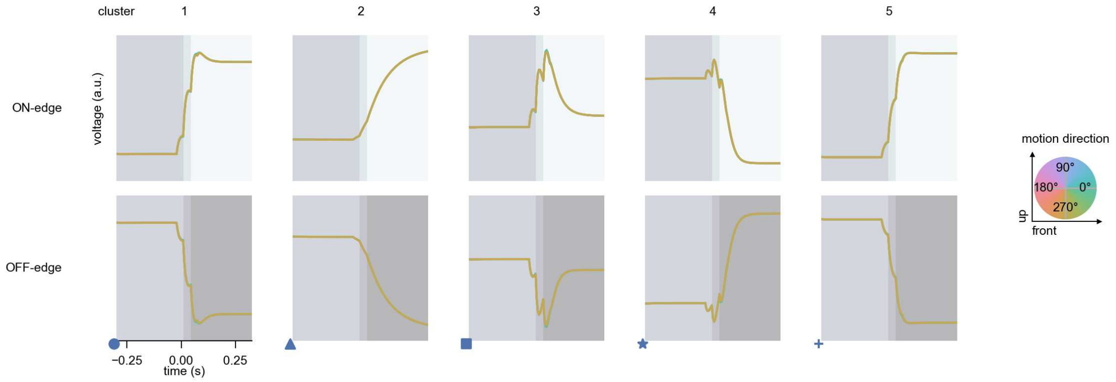

R7 - Figure 7: **Responses to moving edges from best-performing models.** The top row shows responses to moving ON-edges, the bottom row shows peak responses to moving OFF-edges. The edge-stimuli move in different directions from 0 to 360 degrees and at different speeds. Before, a grey-stimulus led the network to a stationary state. The columns show responses from the best-task-performing model in the respective cluster. Example responses to edges moving at 75.4°/s in all cardinal directions (green 0°, blue 90°, red 180°, yellow 270°) from -22.5 to 22.5° visual angle. Background contour indicates the edge, passing by the central column at second zero.

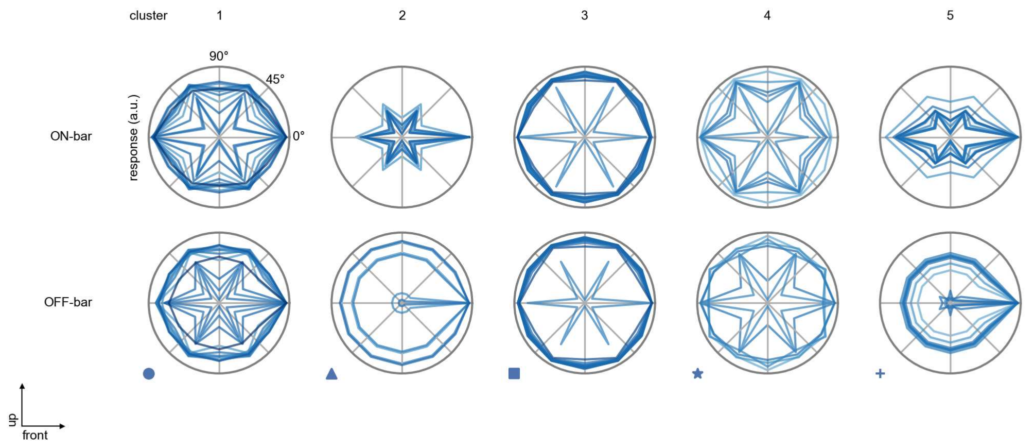

R7 - Figure 8: **Peak responses to moving bars.** The top row shows normalized peak responses to moving ON-bars, the bottom row shows normalized peak responses to moving OFF-bars. The bar-stimuli move in different directions from 0 to 360 degrees. Before, a grey-stimulus led the network to a stationary state. The responses from the different models in the different clusters (columns) overlay and are colored in the respective task error of the model. The peak responses are averaged over all assessed bar-speeds.

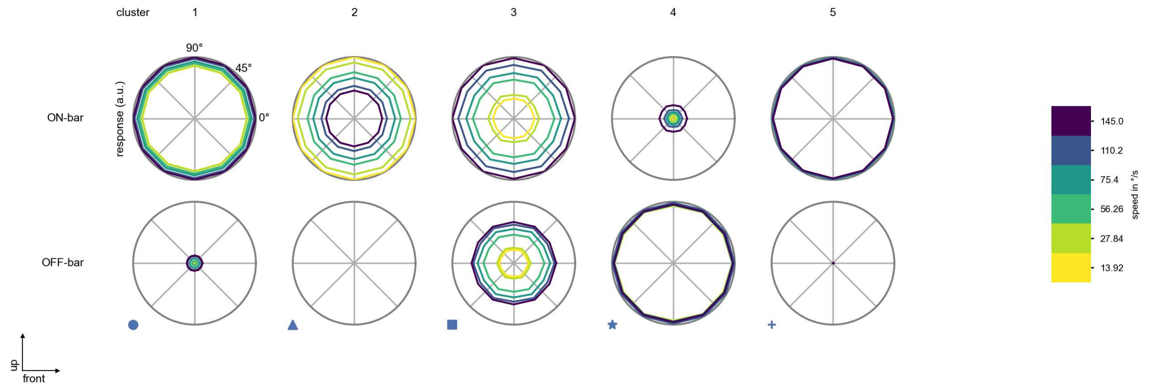

R7 - Figure 9: **Peak responses to moving bars from best-performing models.** The top row shows peak responses to moving ON-bars, the bottom row shows peak responses to moving OFF-bars. The bar-stimuli move in different directions from 0 to 360 degrees and at different speeds. Before, a grey-stimulus led the network to a stationary state. The columns show responses from the best-task-performing model in the respective cluster.

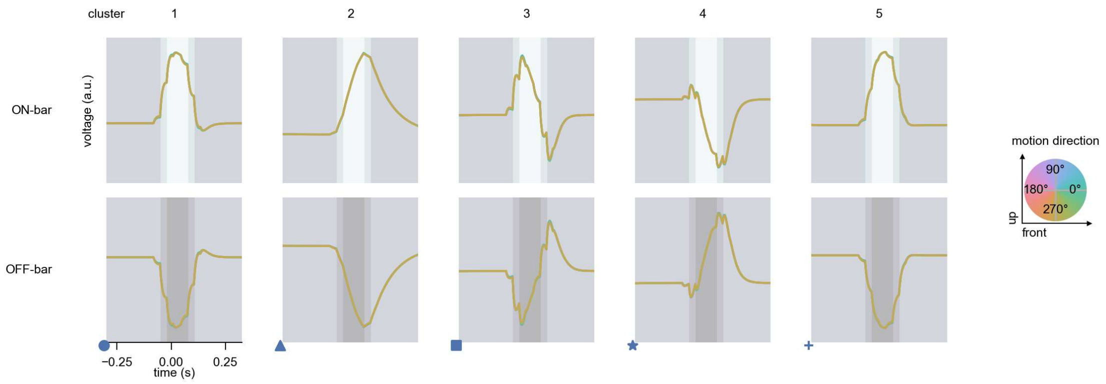

R7 - Figure 10: **Responses to moving bars from best-performing models.** The top row shows responses to moving ON-bars, the bottom row shows peak responses to moving OFF-bars. The bar-stimuli move in different directions from 0 to 360 degrees and at different speeds. Before, a grey-stimulus led the network to a stationary state. The columns show responses from the best-task-performing model in the respective cluster. Example responses to bars moving at 75.4°/s in all cardinal directions (green 0°, blue 90°, red 180°, yellow 270°) from -22.5 to 22.5° visual angle. Background contour indicates the bar, passing by the central column at second zero.

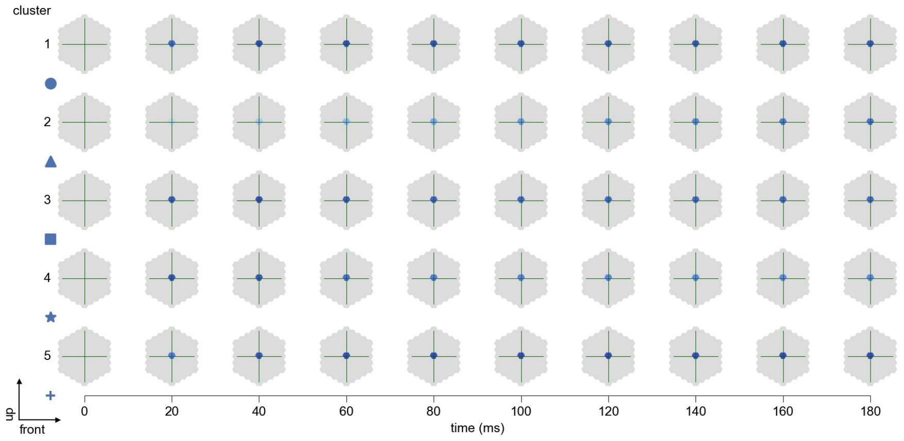

R7 - Figure 11: **Spatio-temporal receptive field.** Responses of the central cell to ON-impulses at single ommatidium impulse locations. The impulse occurs at second zero. Before, a grey-stimulus led the network to a stationary state. Responses are baseline-subtracted. Each row illustrates responses from the best-performing model of the respective cluster. Red indicated depolarization, blue indicates hyperpolarization.

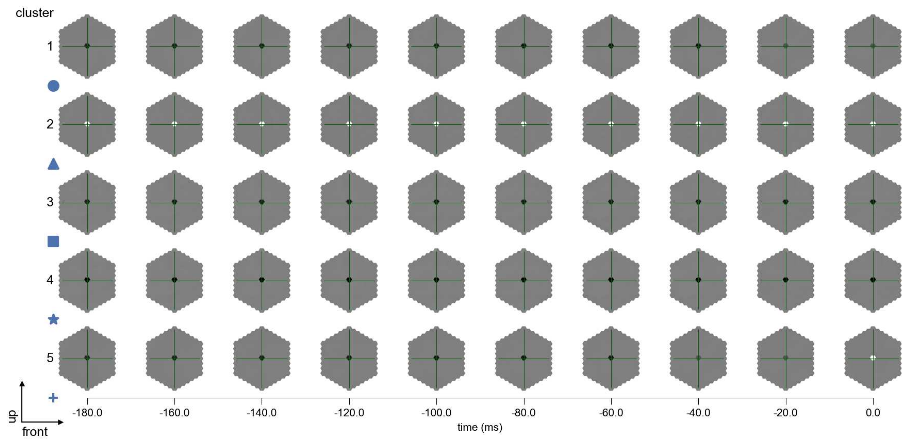

R7 - Figure 12: **Maximally excitatory stimuli.** Each row presents the regularized naturalistic-stimulus from the Sintel dataset that maximizes the cell type's central column response at second zero in the best-performing model of the respective cluster.

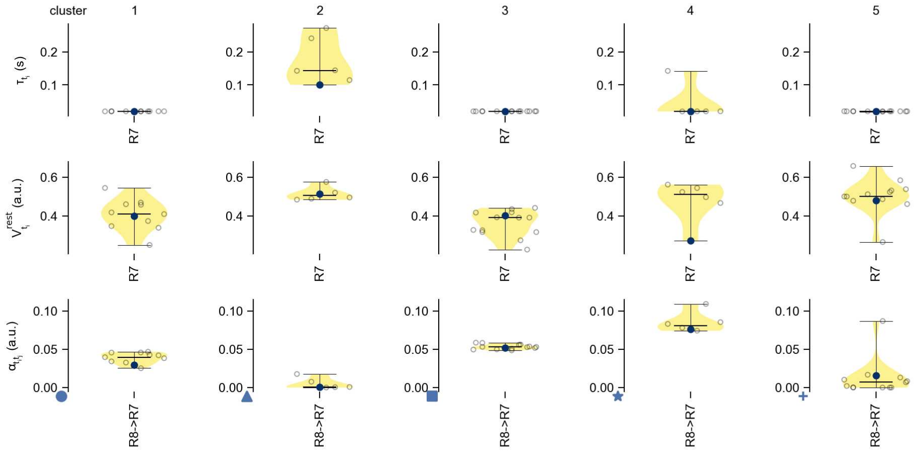

R7 - Figure 13: **Task-constrained parameters.** Each column shows the parameters inferred within the respective cluster. First row: learned time constants of the cell type. Second row: resting potentials of the cell type. Third row: scaling factors for the convolutional filters. The blue scatter represents the parameters from the best-performing model within the cluster.

# 8 R8

← Cell types

#### Figures

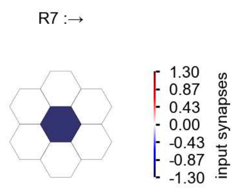

R8 - Figure 1: **Anatomical receptive fields.** Each colored hexagon is a projection, with the projection strength characterized by the average number of synapses that we count from the EM reconstruction. Red indicates excitatory synapses, blue indicates inhibitory synapses from inferred signs. Filters in the order of their total number of synapses.

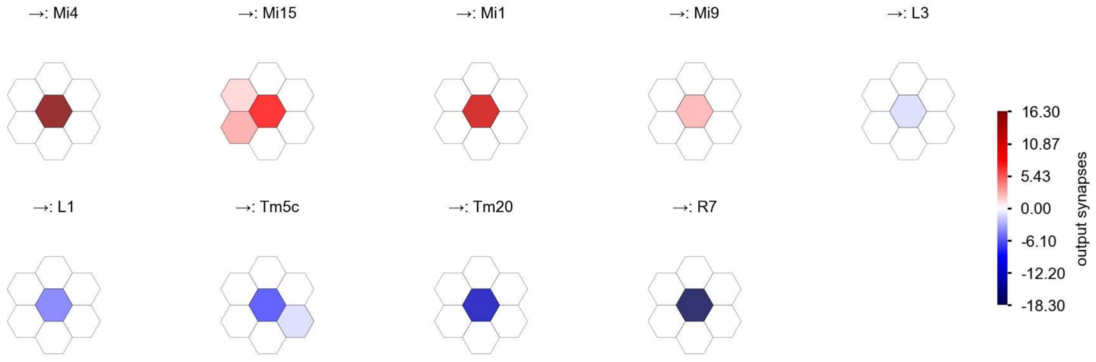

R8 - Figure 2: **Anatomical projective fields.** Each colored hexagon is a projection, with the projection strength characterized by the average number of synapses that we count from the EM reconstruction. Red indicates excitatory synapses, blue indicates inhibitory synapses from inferred signs. Filters in the order of their total number of synapses.

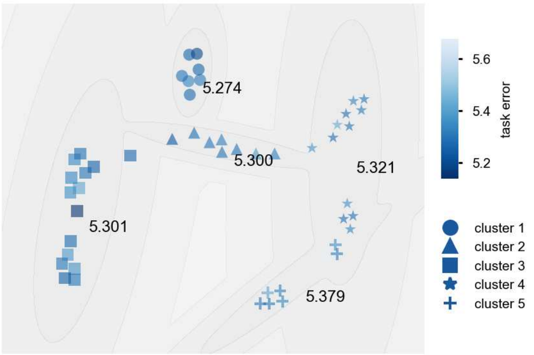

R8 - Figure 3: **Clustering of the responses to naturalistic stimuli.** From responses to naturalistic scenes from Sintel. Scatterpoints represent individual models colored by their task error.

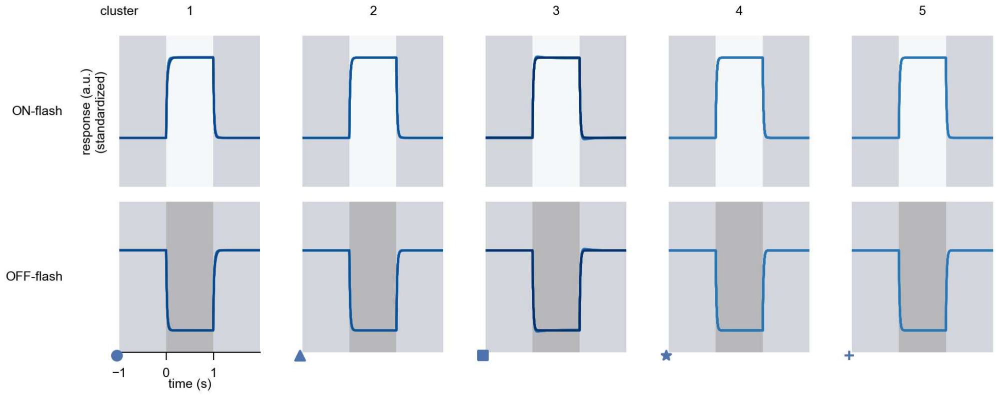

R8 - Figure 4: **Responses to flashes.** The top row shows responses to ON-flashes, the bottom row shows responses to OFF-flashes. The responses from the different models in the different clusters (columns) overlay and are colored in the respective task error of the model. The circular flashes occur at second zero, cover 6 ommatidia in radius, and last for one second. Before and after, a grey-stimulus leads to a stationary state of the network. The traces are standardized.

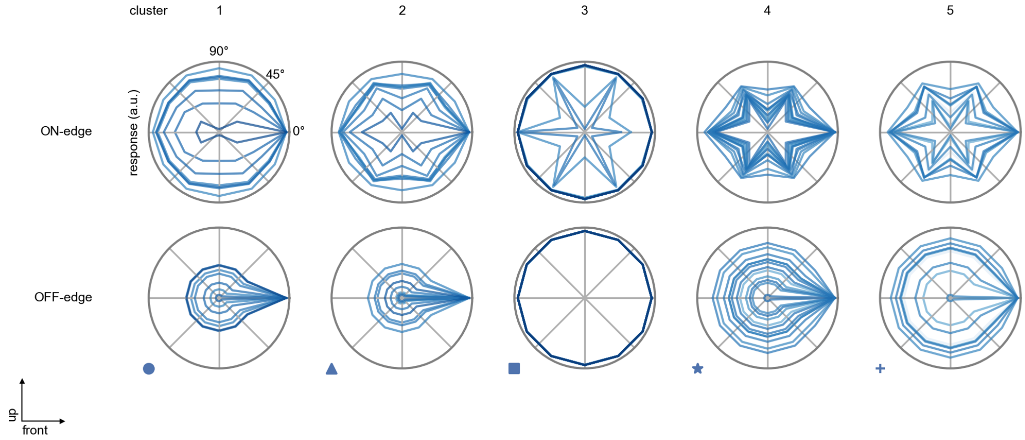

R8 - Figure 5: **Peak responses to moving edges.** The top row shows normalized peak responses to moving ON-edges, the bottom row shows normalized peak responses to moving OFF-edges. The edge-stimuli move in different directions from 0 to 360 degrees. Before, a grey-stimulus led the network to a stationary state. The responses from the different models in the different clusters (columns) overlay and are colored in the respective task error of the model. The peak responses are averaged over all assessed edge-speeds.

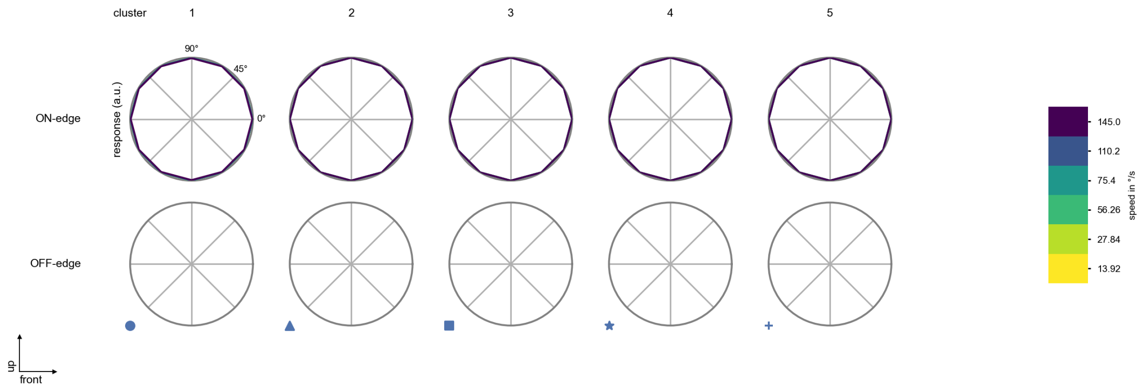

R8 - Figure 6: **Peak responses to moving edges from best-performing models.** The top row shows peak responses to moving ON-edges, the bottom row shows peak responses to moving OFF-edges. The edge-stimuli move in different directions from 0 to 360 degrees and at different speeds. Before, a grey-stimulus led the network to a stationary state. The columns show responses from the best-task-performing model in the respective cluster.

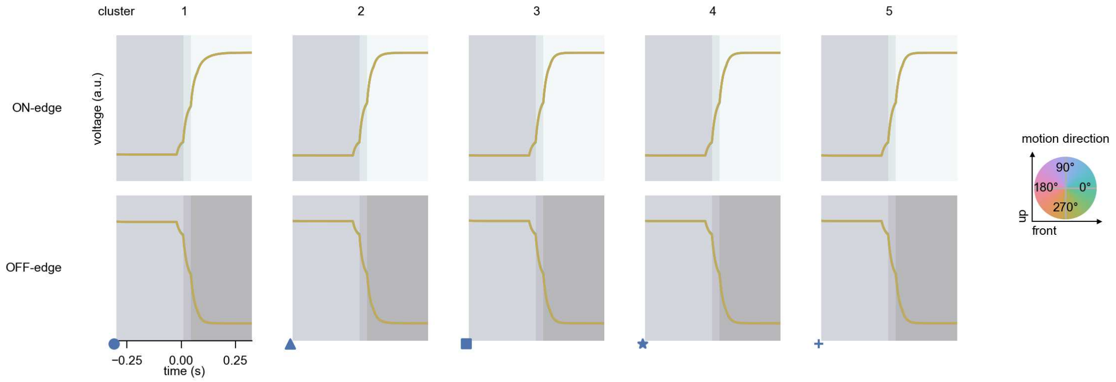

R8 - Figure 7: **Responses to moving edges from best-performing models.** The top row shows responses to moving ON-edges, the bottom row shows peak responses to moving OFF-edges. The edge-stimuli move in different directions from 0 to 360 degrees and at different speeds. Before, a grey-stimulus led the network to a stationary state. The columns show responses from the best-task-performing model in the respective cluster. Example responses to edges moving at 75.4°/s in all cardinal directions (green 0°, blue 90°, red 180°, yellow 270°) from -22.5 to 22.5° visual angle. Background contour indicates the edge, passing by the central column at second zero.

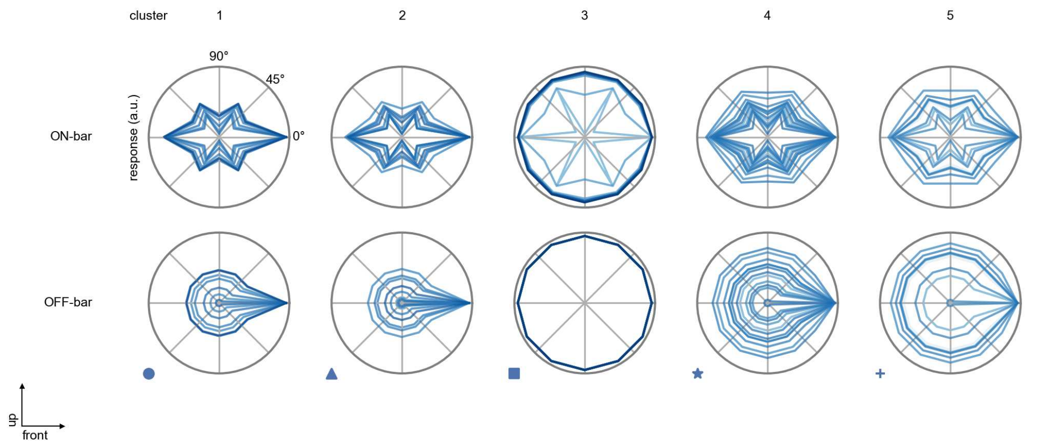

R8 - Figure 8: **Peak responses to moving bars.** The top row shows normalized peak responses to moving ON-bars, the bottom row shows normalized peak responses to moving OFF-bars. The bar-stimuli move in different directions from 0 to 360 degrees. Before, a grey-stimulus led the network to a stationary state. The responses from the different models in the different clusters (columns) overlay and are colored in the respective task error of the model. The peak responses are averaged over all assessed bar-speeds.

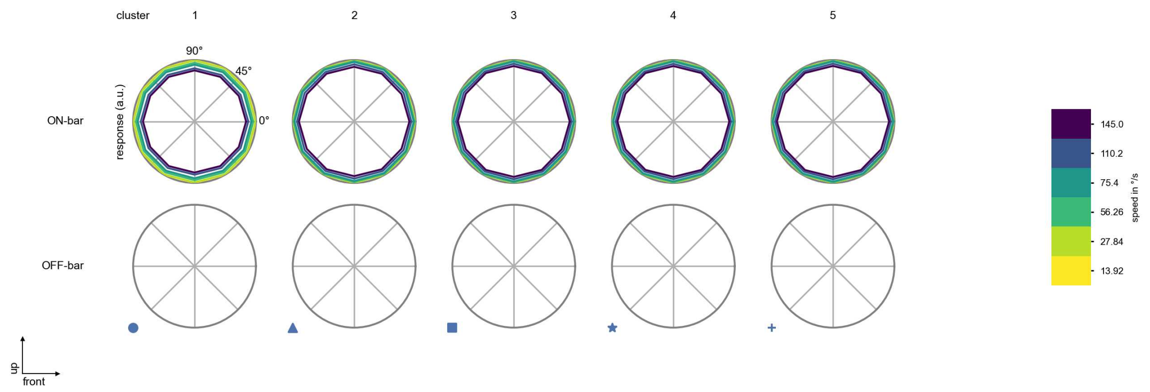

R8 - Figure 9: **Peak responses to moving bars from best-performing models.** The top row shows peak responses to moving ON-bars, the bottom row shows peak responses to moving OFF-bars. The bar-stimuli move in different directions from 0 to 360 degrees and at different speeds. Before, a grey-stimulus led the network to a stationary state. The columns show responses from the best-task-performing model in the respective cluster.

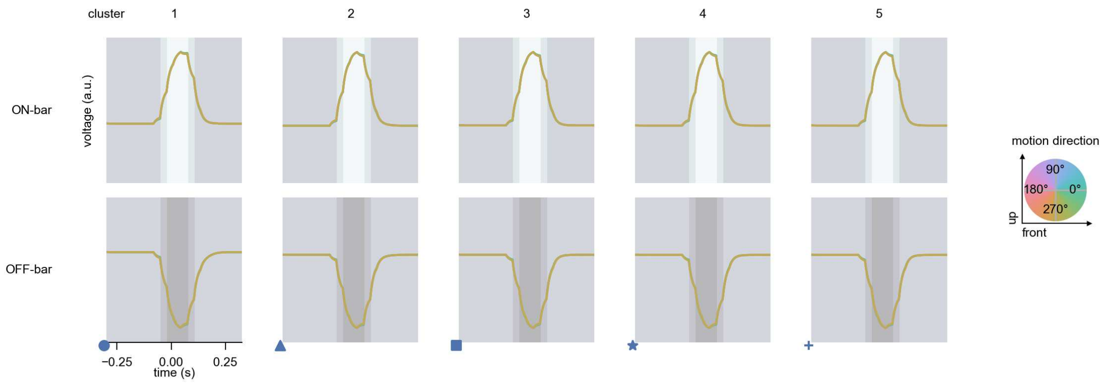

R8 - Figure 10: **Responses to moving bars from best-performing models.** The top row shows responses to moving ON-bars, the bottom row shows peak responses to moving OFF-bars. The bar-stimuli move in different directions from 0 to 360 degrees and at different speeds. Before, a grey-stimulus led the network to a stationary state. The columns show responses from the best-task-performing model in the respective cluster. Example responses to bars moving at 75.4°/s in all cardinal directions (green 0°, blue 90°, red 180°, yellow 270°) from -22.5 to 22.5° visual angle. Background contour indicates the bar, passing by the central column at second zero.

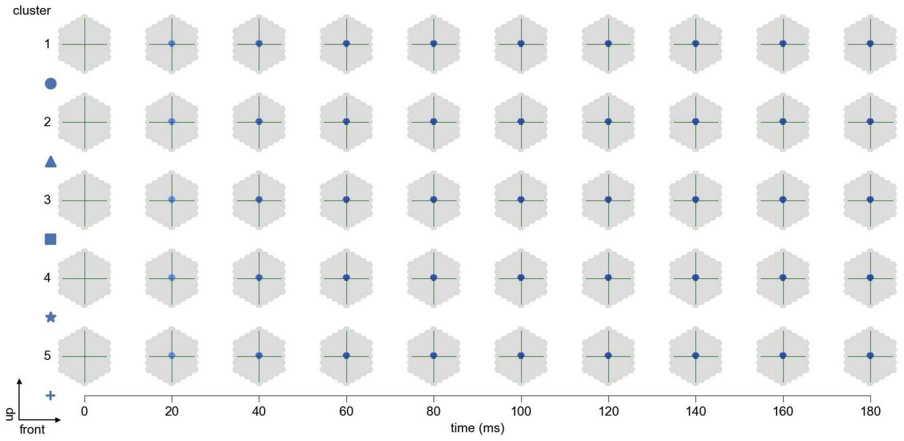

R8 - Figure 11: **Spatio-temporal receptive field.** Responses of the central cell to ON-impulses at single ommatidium impulse locations. The impulse occurs at second zero. Before, a grey-stimulus led the network to a stationary state. Responses are baseline-subtracted. Each row illustrates responses from the best-performing model of the respective cluster. Red indicated depolarization, blue indicates hyperpolarization.

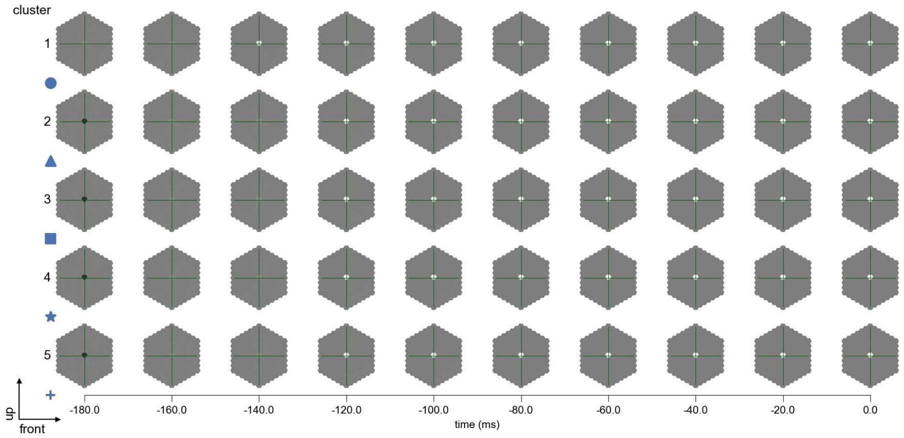

R8 - Figure 12: **Maximally excitatory stimuli.** Each row presents the regularized naturalistic-stimulus from the Sintel dataset that maximizes the cell type's central column response at second zero in the best-performing model of the respective cluster.

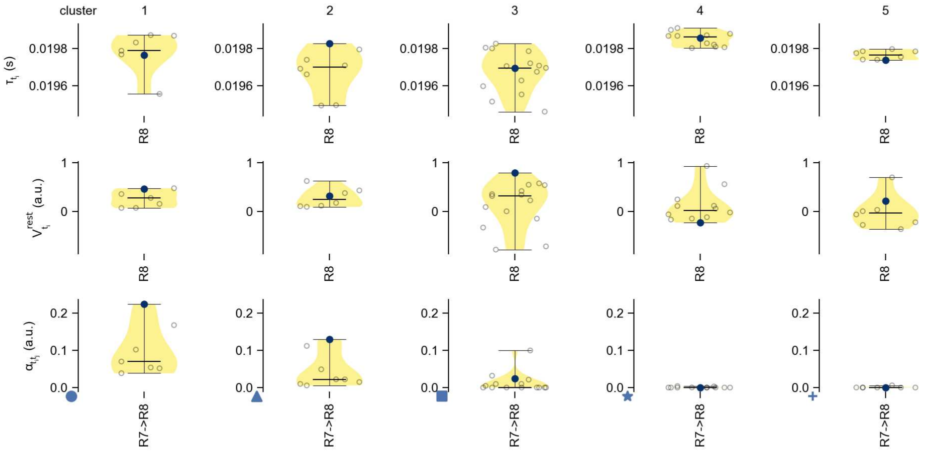

R8 - Figure 13: **Task-constrained parameters.** Each column shows the parameters inferred within the respective cluster. First row: learned time constants of the cell type. Second row: resting potentials of the cell type. Third row: scaling factors for the convolutional filters. The blue scatter represents the parameters from the best-performing model within the cluster.

# 9 L1

← Cell types

#### Figures

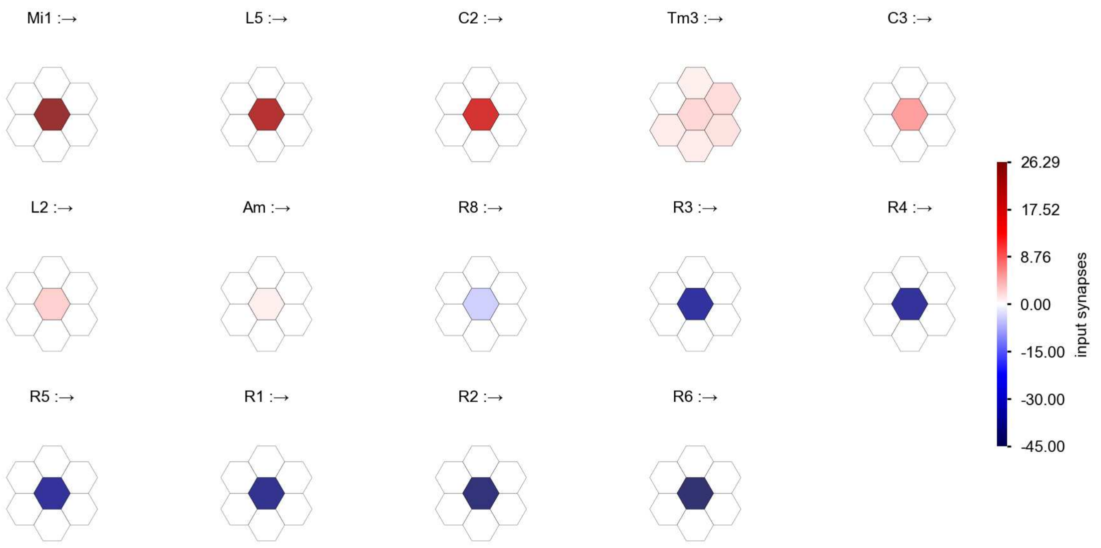

L1 - Figure 1: **Anatomical receptive fields.** Each colored hexagon is a projection, with the projection strength characterized by the average number of synapses that we count from the EM reconstruction. Red indicates excitatory synapses, blue indicates inhibitory synapses from inferred signs. Filters in the order of their total number of synapses.

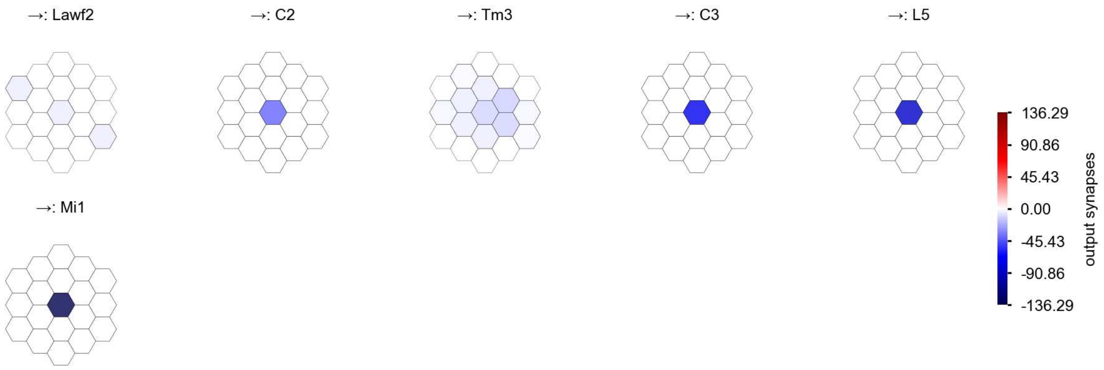

L1 - Figure 2: **Anatomical projective fields.** Each colored hexagon is a projection, with the projection strength characterized by the average number of synapses that we count from the EM reconstruction. Red indicates excitatory synapses, blue indicates inhibitory synapses from inferred signs. Filters in the order of their total number of synapses.

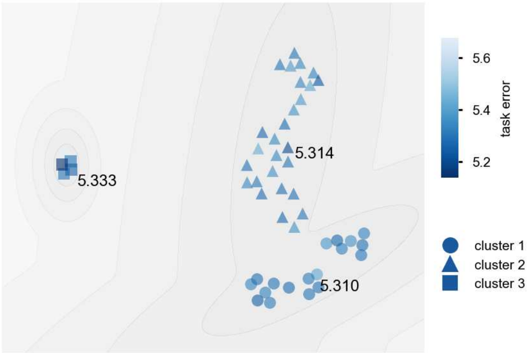

L1 - Figure 3: **Clustering of the responses to naturalistic stimuli.** From responses to naturalistic scenes from Sintel. Scatterpoints represent individual models colored by their task error.

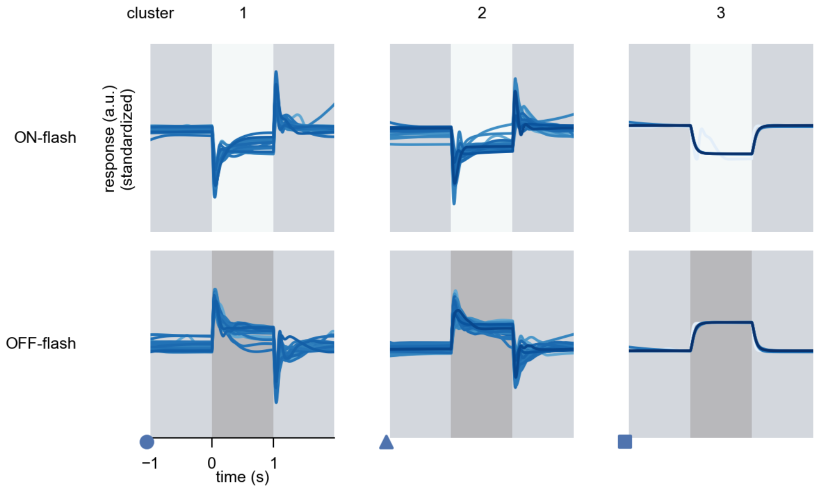

L1 - Figure 4: **Responses to flashes.** The top row shows responses to ON-flashes, the bottom row shows responses to OFF-flashes. The responses from the different models in the different clusters (columns) overlay and are colored in the respective task error of the model. The circular flashes occur at second zero, cover 6 ommatidia in radius, and last for one second. Before and after, a grey-stimulus leads to a stationary state of the network. The traces are standardized.

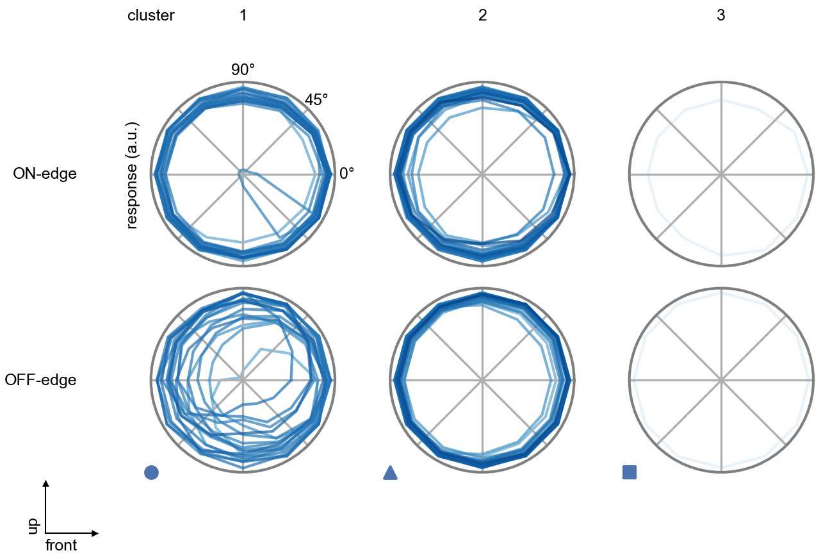

L1 - Figure 5: **Peak responses to moving edges.** The top row shows normalized peak responses to moving ON-edges, the bottom row shows normalized peak responses to moving OFF-edges. The edge-stimuli move in different directions from 0 to 360 degrees. Before, a grey-stimulus led the network to a stationary state. The responses from the different models in the different clusters (columns) overlay and are colored in the respective task error of the model. The peak responses are averaged over all assessed edge-speeds.

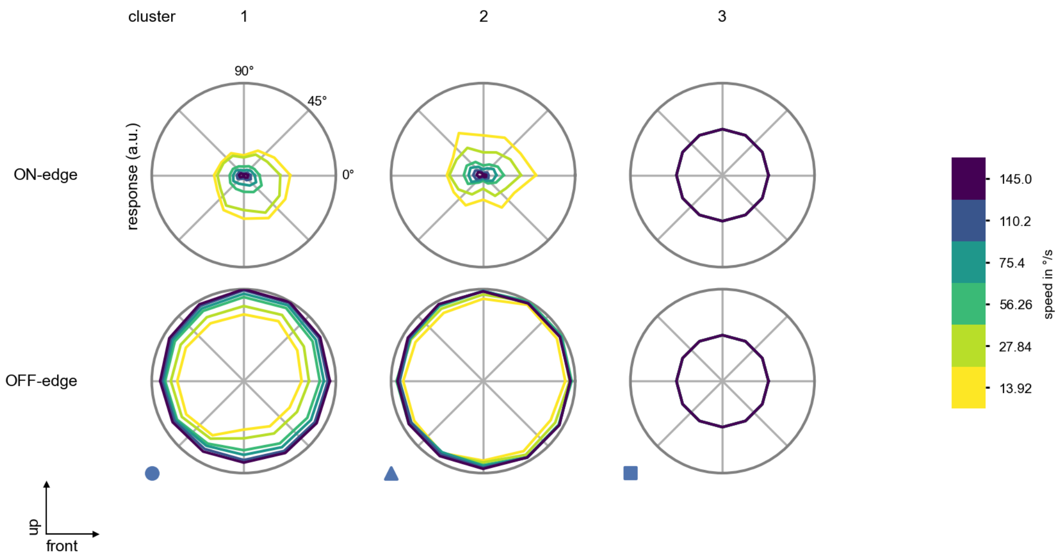

L1 - Figure 6: **Peak responses to moving edges from best-performing models.** The top row shows peak responses to moving ON-edges, the bottom row shows peak responses to moving OFF-edges. The edge-stimuli move in different directions from 0 to 360 degrees and at different speeds. Before, a grey-stimulus led the network to a stationary state. The columns show responses from the best-task-performing model in the respective cluster.

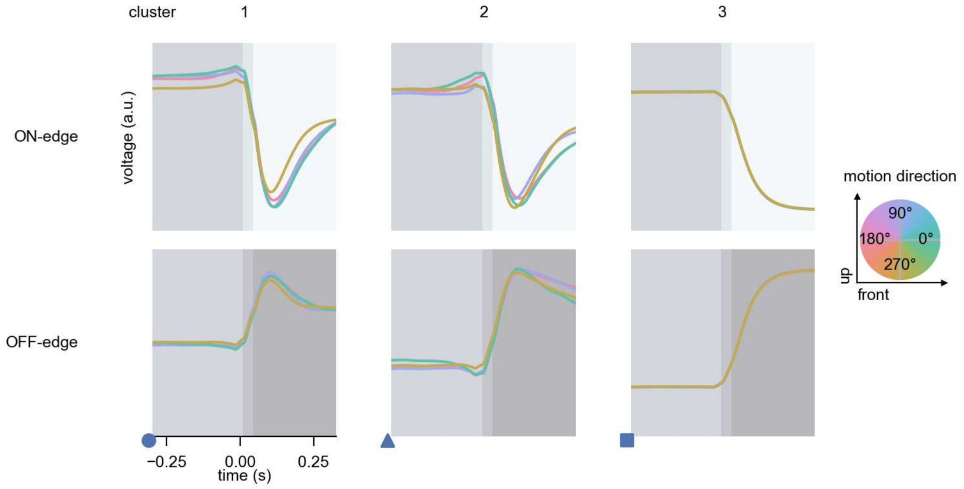

L1 - Figure 7: **Responses to moving edges from best-performing models.** The top row shows responses to moving ON-edges, the bottom row shows peak responses to moving OFF-edges. The edge-stimuli move in different directions from 0 to 360 degrees and at different speeds. Before, a grey-stimulus led the network to a stationary state. The columns show responses from the best-task-performing model in the respective cluster. Example responses to edges moving at 75.4°/s in all cardinal directions (green 0°, blue 90°, red 180°, yellow 270°) from -22.5 to 22.5° visual angle. Background contour indicates the edge, passing by the central column at second zero.

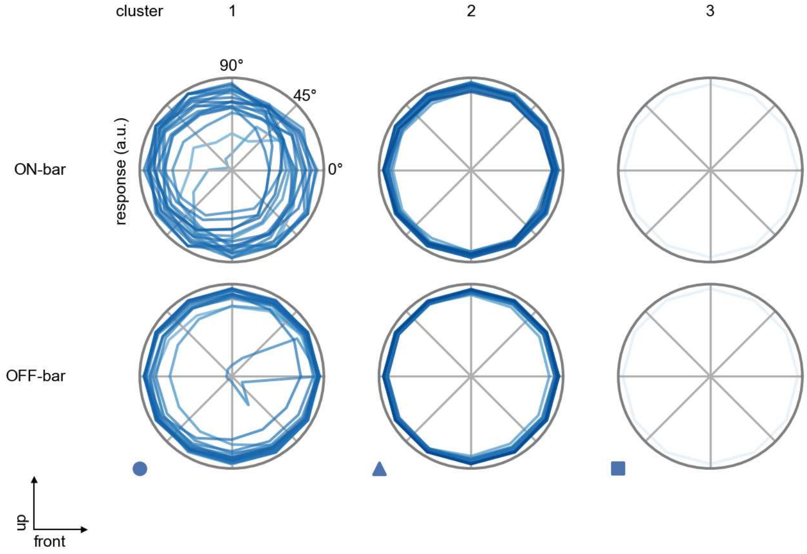

L1 - Figure 8: **Peak responses to moving bars.** The top row shows normalized peak responses to moving ON-bars, the bottom row shows normalized peak responses to moving OFF-bars. The bar-stimuli move in different directions from 0 to 360 degrees. Before, a grey-stimulus led the network to a stationary state. The responses from the different models in the different clusters (columns) overlay and are colored in the respective task error of the model. The peak responses are averaged over all assessed bar-speeds.

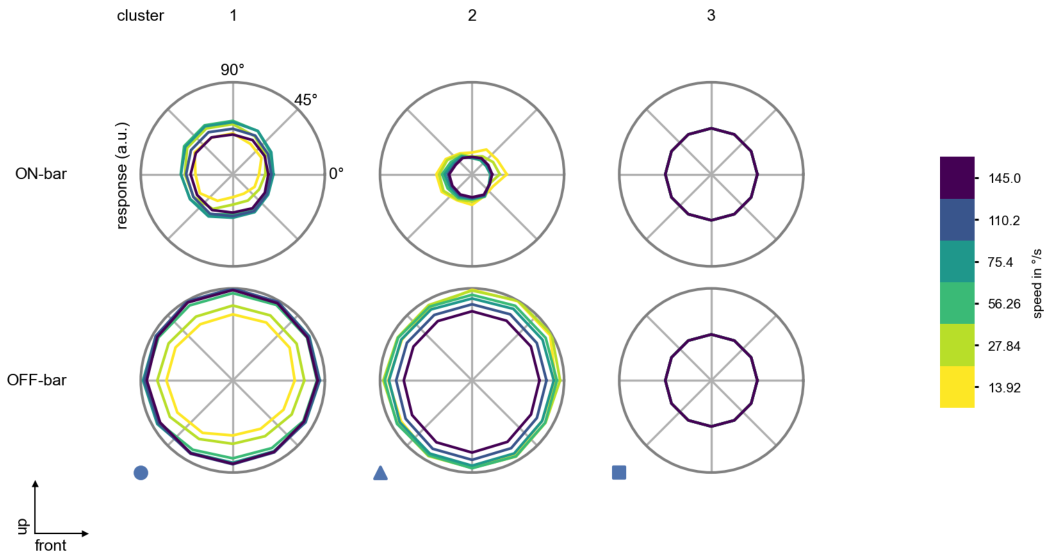

L1 - Figure 9: **Peak responses to moving bars from best-performing models.** The top row shows peak responses to moving ON-bars, the bottom row shows peak responses to moving OFF-bars. The bar-stimuli move in different directions from 0 to 360 degrees and at different speeds. Before, a grey-stimulus led the network to a stationary state. The columns show responses from the best-task-performing model in the respective cluster.

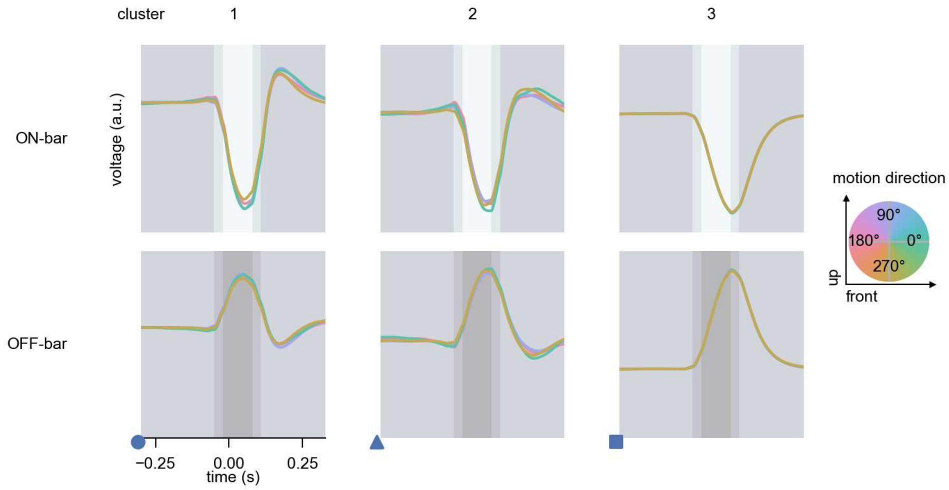

L1 - Figure 10: **Responses to moving bars from best-performing models.** The top row shows responses to moving ON-bars, the bottom row shows peak responses to moving OFF-bars. The bar-stimuli move in different directions from 0 to 360 degrees and at different speeds. Before, a grey-stimulus led the network to a stationary state. The columns show responses from the best-task-performing model in the respective cluster. Example responses to bars moving at 75.4°/s in all cardinal directions (green 0°, blue 90°, red 180°, yellow 270°) from -22.5 to 22.5° visual angle. Background contour indicates the bar, passing by the central column at second zero.

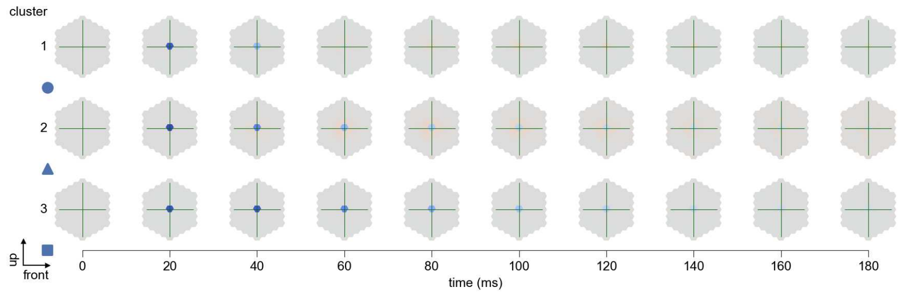

L1 - Figure 11: **Spatio-temporal receptive field.** Responses of the central cell to ON-impulses at single ommatidium impulse locations. The impulse occurs at second zero. Before, a grey-stimulus led the network to a stationary state. Responses are baseline-subtracted. Each row illustrates responses from the best-performing model of the respective cluster. Red indicated depolarization, blue indicates hyperpolarization.

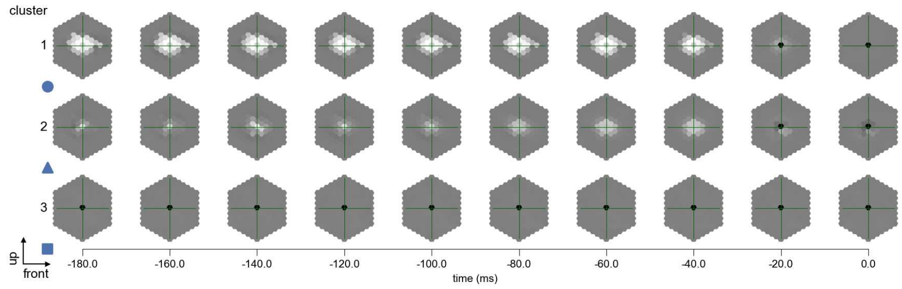

L1 - Figure 12: **Maximally excitatory stimuli.** Each row presents the regularized naturalistic-stimulus from the Sintel dataset that maximizes the cell type's central column response at second zero in the best-performing model of the respective cluster.

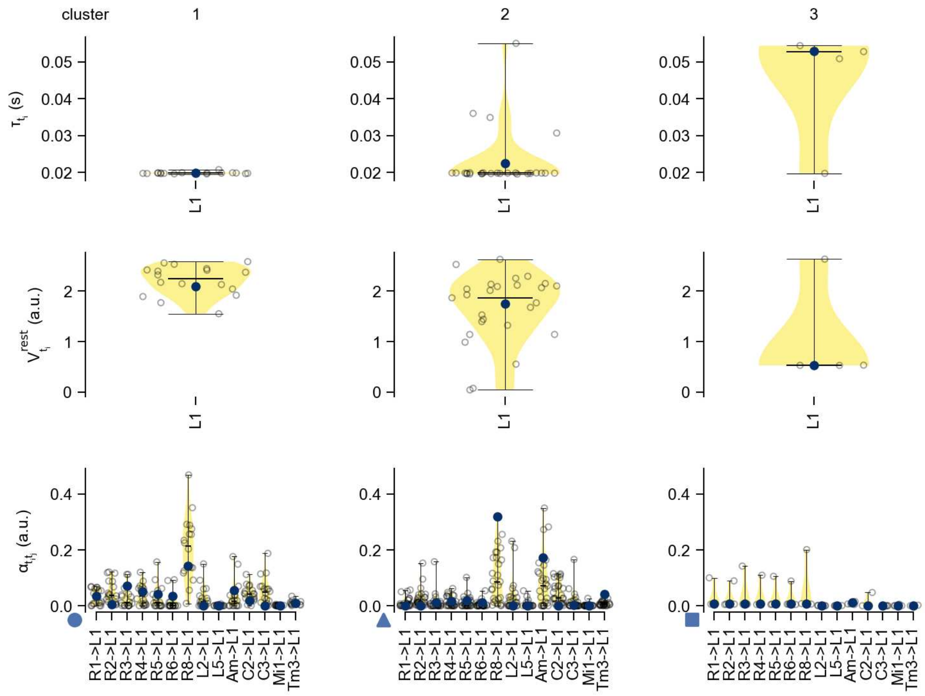

L1 - Figure 13: **Task-constrained parameters.** Each column shows the parameters inferred within the respective cluster. First row: learned time constants of the cell type. Second row: resting potentials of the cell type. Third row: scaling factors for the convolutional filters. The blue scatter represents the parameters from the best-performing model within the cluster.

# 10 L2

← Cell types

#### Figures

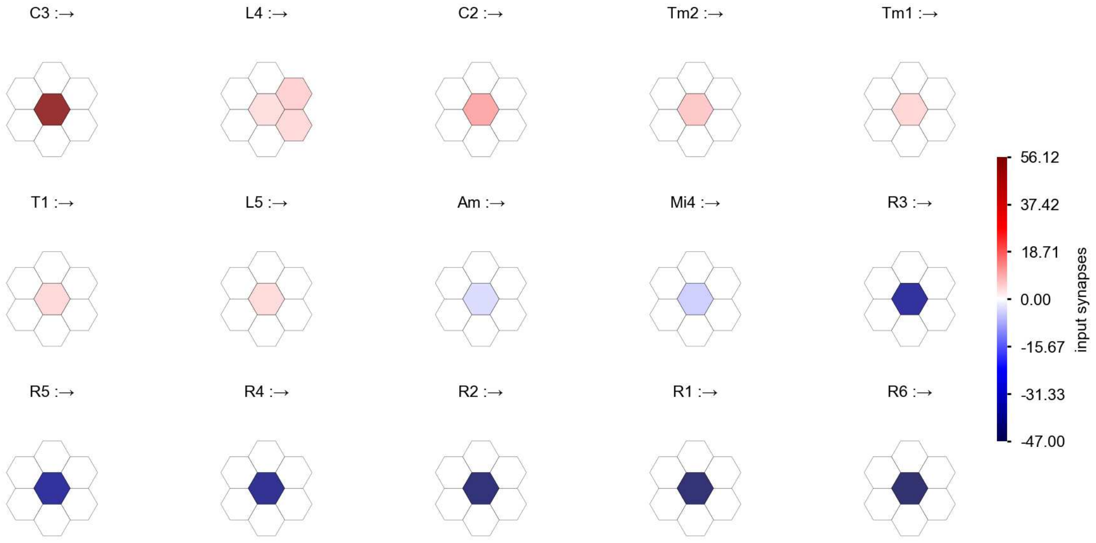

L2 - Figure 1: **Anatomical receptive fields.** Each colored hexagon is a projection, with the projection strength characterized by the average number of synapses that we count from the EM reconstruction. Red indicates excitatory synapses, blue indicates inhibitory synapses from inferred signs. Filters in the order of their total number of synapses.

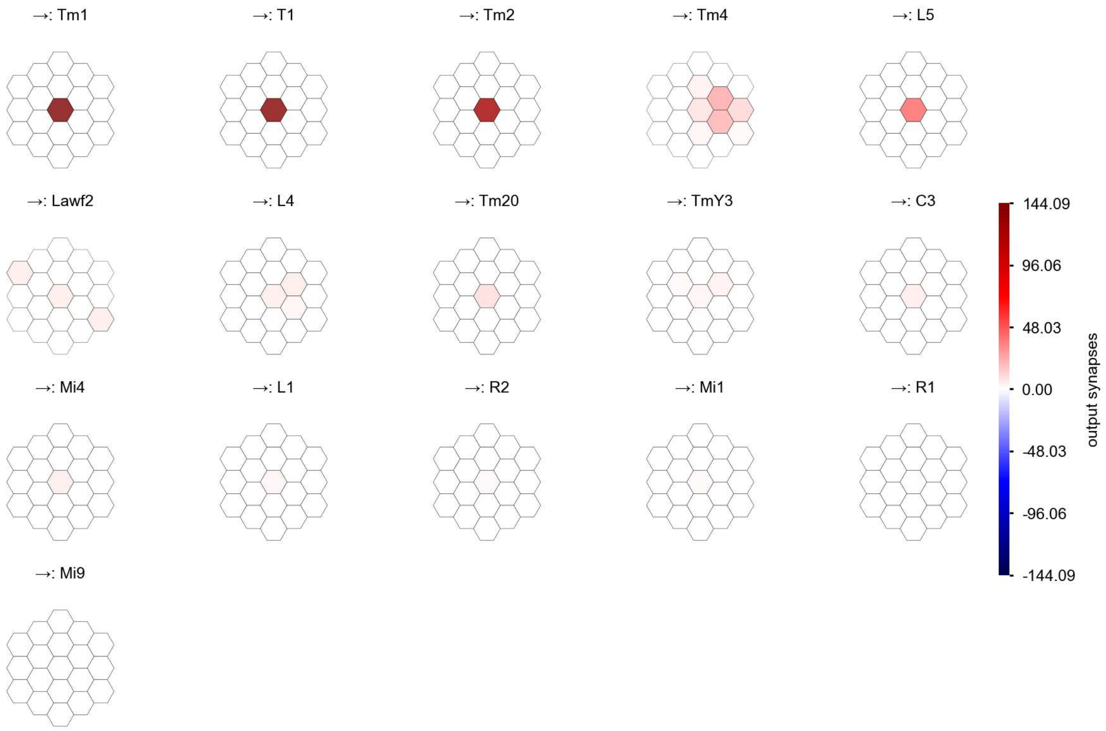

L2 - Figure 2: **Anatomical projective fields.** Each colored hexagon is a projection, with the projection strength characterized by the average number of synapses that we count from the EM reconstruction. Red indicates excitatory synapses, blue indicates inhibitory synapses from inferred signs. Filters in the order of their total number of synapses.

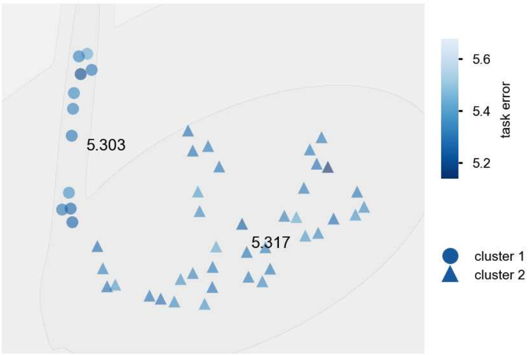

L2 - Figure 3: **Clustering of the responses to naturalistic stimuli.** From responses to naturalistic scenes from Sintel. Scatterpoints represent individual models colored by their task error.

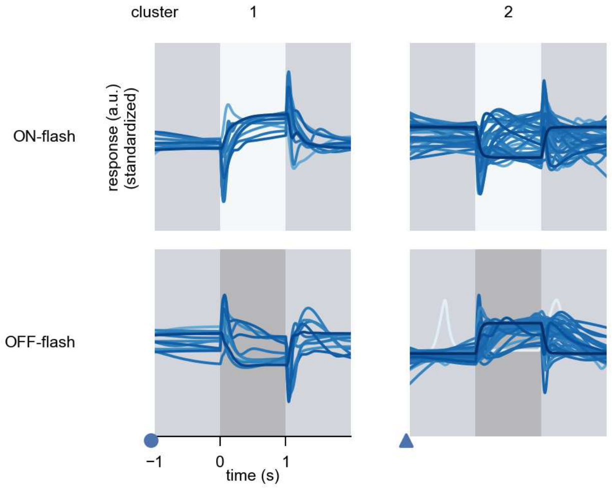

L2 - Figure 4: **Responses to flashes.** The top row shows responses to ON-flashes, the bottom row shows responses to OFF-flashes. The responses from the different models in the different clusters (columns) overlay and are colored in the respective task error of the model. The circular flashes occur at second zero, cover 6 ommatidia in radius, and last for one second. Before and after, a grey-stimulus leads to a stationary state of the network. The traces are standardized.

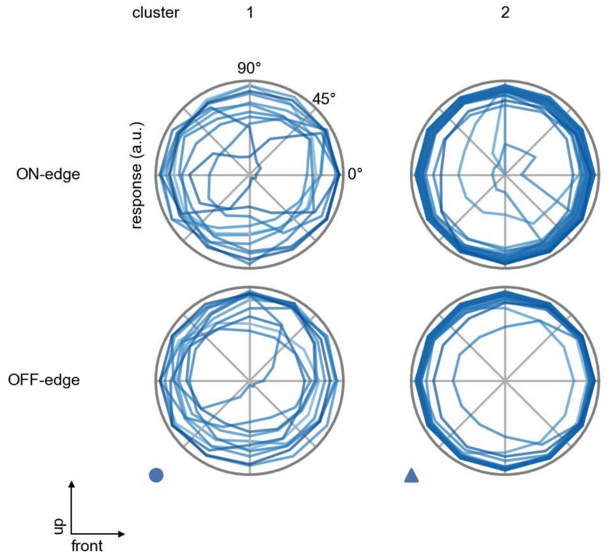

L2 - Figure 5: **Peak responses to moving edges.** The top row shows normalized peak responses to moving ON-edges, the bottom row shows normalized peak responses to moving OFF-edges. The edge-stimuli move in different directions from 0 to 360 degrees. Before, a grey-stimulus led the network to a stationary state. The responses from the different models in the different clusters (columns) overlay and are colored in the respective task error of the model. The peak responses are averaged over all assessed edge-speeds.

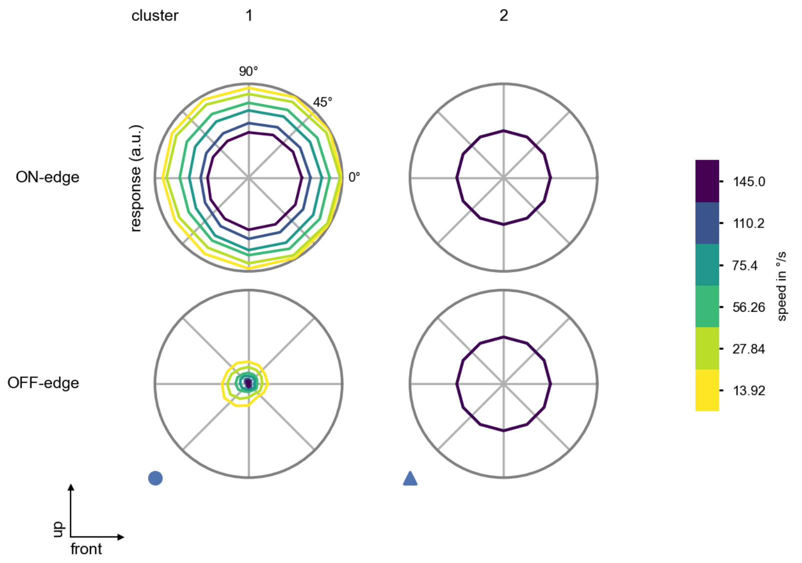

L2 - Figure 6: **Peak responses to moving edges from best-performing models.** The top row shows peak responses to moving ON-edges, the bottom row shows peak responses to moving OFF-edges. The edge-stimuli move in different directions from 0 to 360 degrees and at different speeds. Before, a grey-stimulus led the network to a stationary state. The columns show responses from the best-task-performing model in the respective cluster.

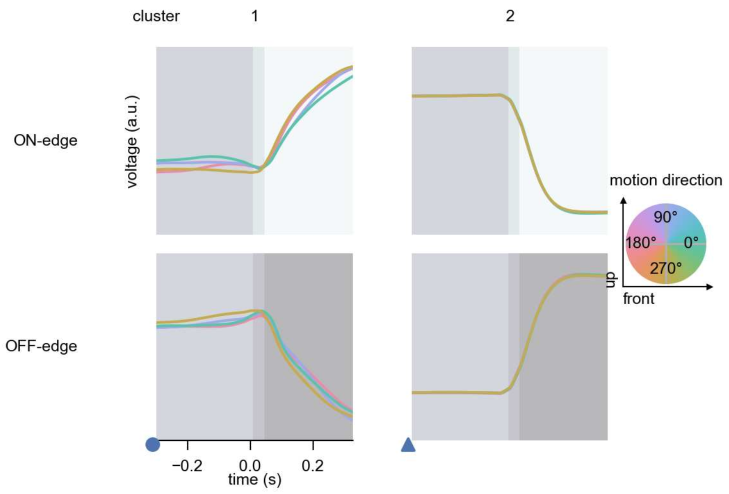

L2 - Figure 7: **Responses to moving edges from best-performing models.** The top row shows responses to moving ON-edges, the bottom row shows peak responses to moving OFF-edges. The edge-stimuli move in different directions from 0 to 360 degrees and at different speeds. Before, a grey-stimulus led the network to a stationary state. The columns show responses from the best-task-performing model in the respective cluster. Example responses to edges moving at 75.4°/s in all cardinal directions (green 0°, blue 90°, red 180°, yellow 270°) from -22.5 to 22.5° visual angle. Background contour indicates the edge, passing by the central column at second zero.

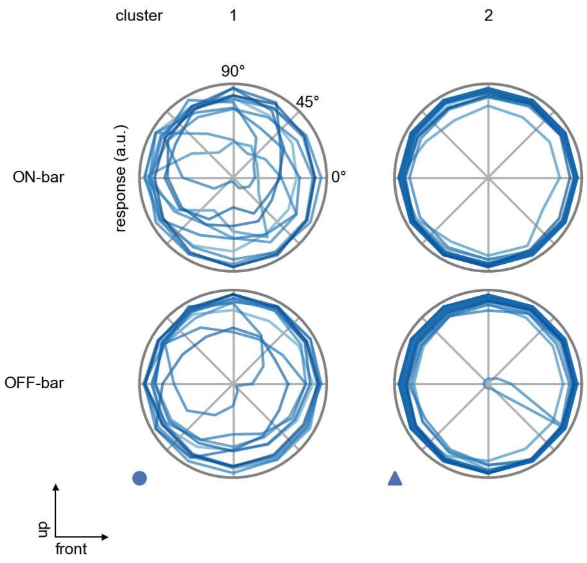

L2 - Figure 8: **Peak responses to moving bars.** The top row shows normalized peak responses to moving ON-bars, the bottom row shows normalized peak responses to moving OFF-bars. The bar-stimuli move in different directions from 0 to 360 degrees. Before, a grey-stimulus led the network to a stationary state. The responses from the different models in the different clusters (columns) overlay and are colored in the respective task error of the model. The peak responses are averaged over all assessed bar-speeds.

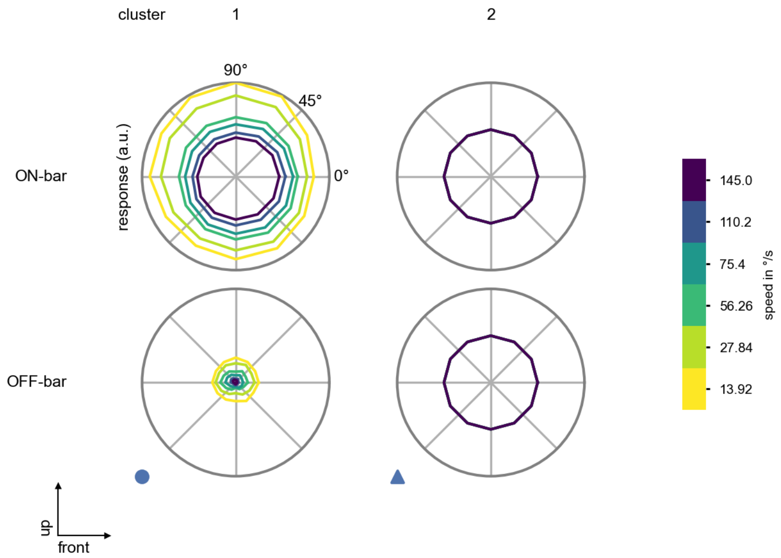

L2 - Figure 9: **Peak responses to moving bars from best-performing models.** The top row shows peak responses to moving ON-bars, the bottom row shows peak responses to moving OFF-bars. The bar-stimuli move in different directions from 0 to 360 degrees and at different speeds. Before, a grey-stimulus led the network to a stationary state. The columns show responses from the best-task-performing model in the respective cluster.

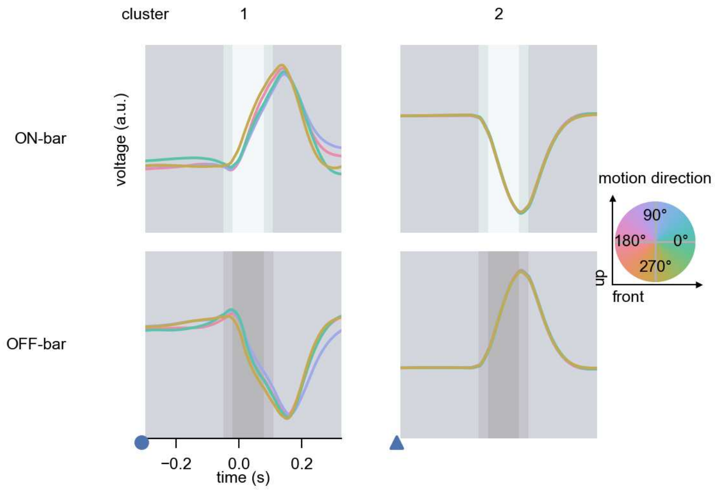

L2 - Figure 10: **Responses to moving bars from best-performing models.** The top row shows responses to moving ON-bars, the bottom row shows peak responses to moving OFF-bars. The bar-stimuli move in different directions from 0 to 360 degrees and at different speeds. Before, a grey-stimulus led the network to a stationary state. The columns show responses from the best-task-performing model in the respective cluster. Example responses to bars moving at 75.4°/s in all cardinal directions (green 0°, blue 90°, red 180°, yellow 270°) from -22.5 to 22.5° visual angle. Background contour indicates the bar, passing by the central column at second zero.

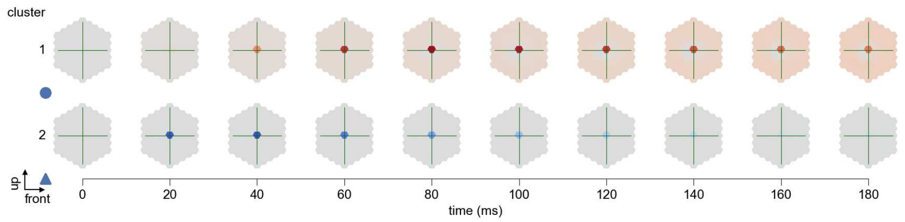

L2 - Figure 11: **Spatio-temporal receptive field.** Responses of the central cell to ON-impulses at single ommatidium impulse locations. The impulse occurs at second zero. Before, a grey-stimulus led the network to a stationary state. Responses are baseline-subtracted. Each row illustrates responses from the best-performing model of the respective cluster. Red indicated depolarization, blue indicates hyperpolarization.

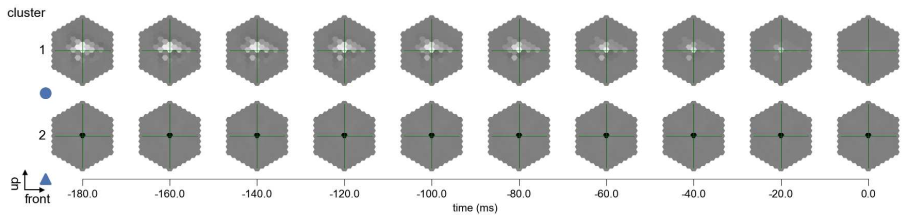

L2 - Figure 12: **Maximally excitatory stimuli.** Each row presents the regularized naturalistic-stimulus from the Sintel dataset that maximizes the cell type's central column response at second zero in the best-performing model of the respective cluster.

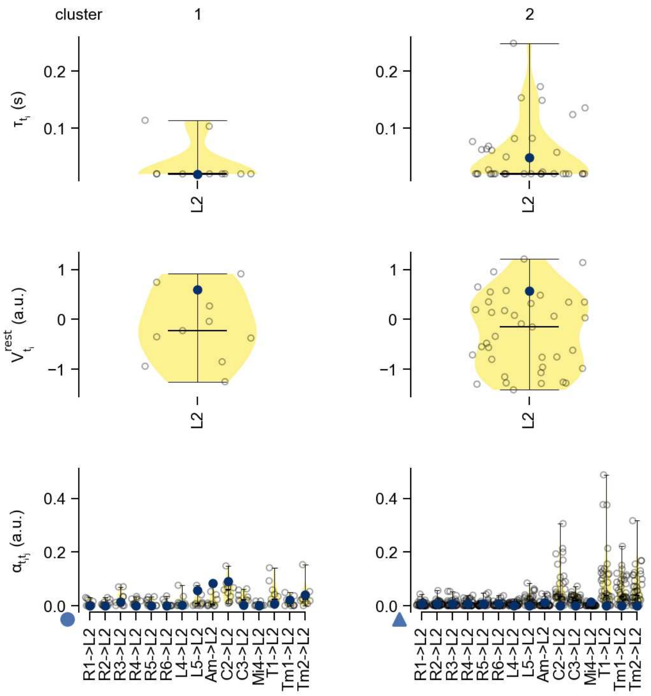

L2 - Figure 13: **Task-constrained parameters.** Each column shows the parameters inferred within the respective cluster. First row: learned time constants of the cell type. Second row: resting potentials of the cell type. Third row: scaling factors for the convolutional filters. The blue scatter represents the parameters from the best-performing model within the cluster.

# 11 L3

← Cell types

#### Figures

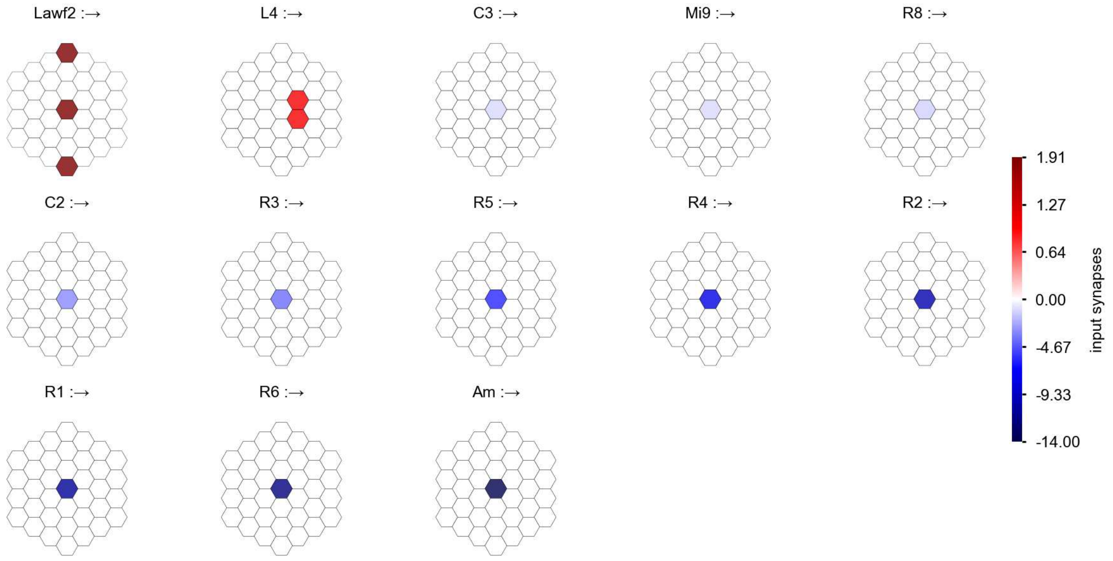

L3 - Figure 1: **Anatomical receptive fields.** Each colored hexagon is a projection, with the projection strength characterized by the average number of synapses that we count from the EM reconstruction. Red indicates excitatory synapses, blue indicates inhibitory synapses from inferred signs. Filters in the order of their total number of synapses.

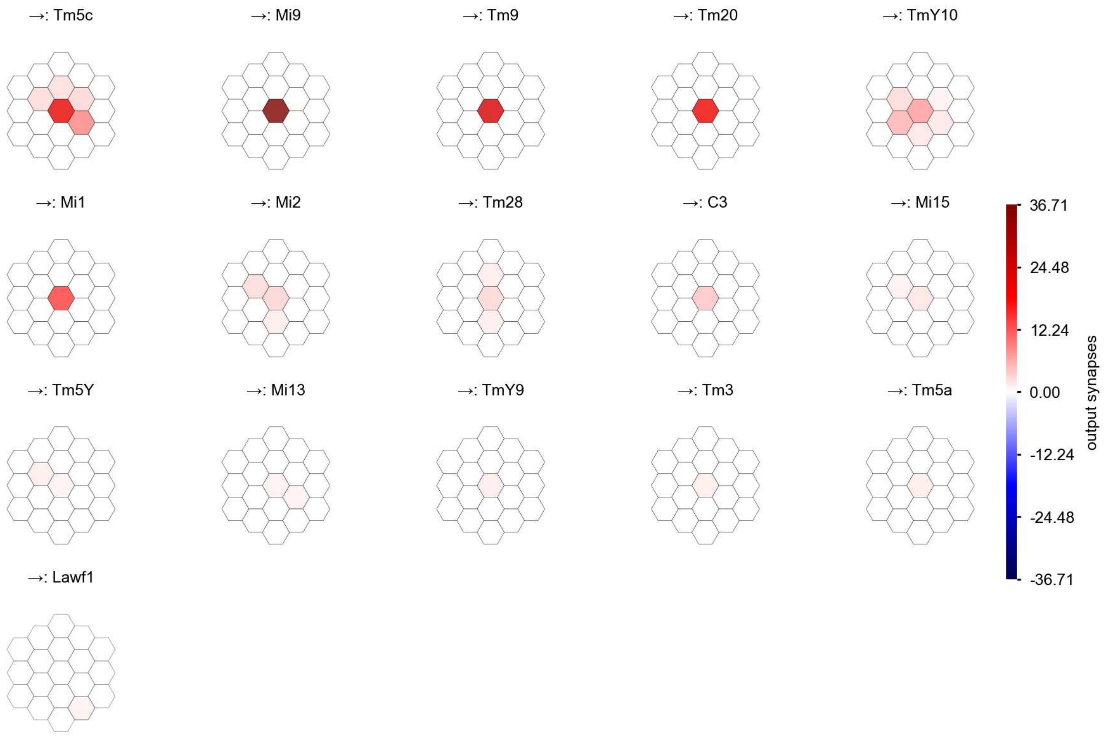

L3 - Figure 2: **Anatomical projective fields.** Each colored hexagon is a projection, with the projection strength characterized by the average number of synapses that we count from the EM reconstruction. Red indicates excitatory synapses, blue indicates inhibitory synapses from inferred signs. Filters in the order of their total number of synapses.

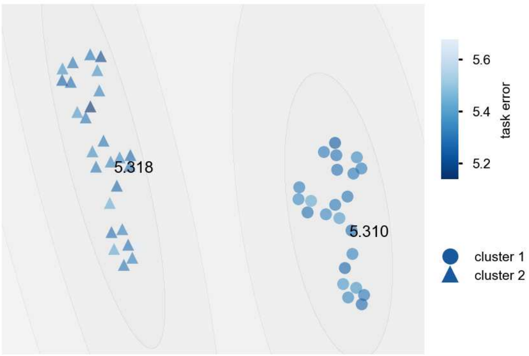

L3 - Figure 3: **Clustering of the responses to naturalistic stimuli.** From responses to naturalistic scenes from Sintel. Scatterpoints represent individual models colored by their task error.

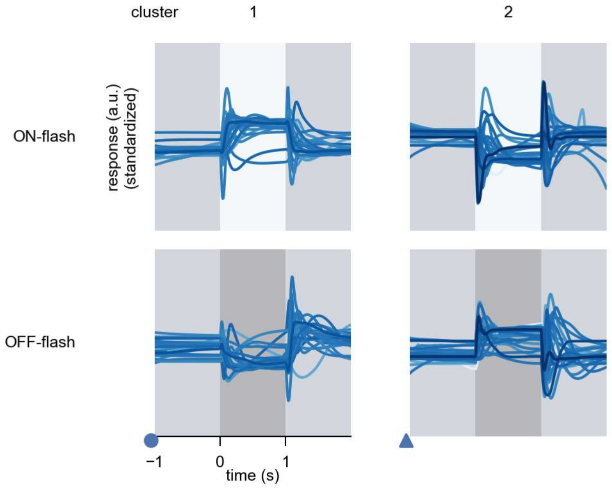

L3 - Figure 4: **Responses to flashes.** The top row shows responses to ON-flashes, the bottom row shows responses to OFF-flashes. The responses from the different models in the different clusters (columns) overlay and are colored in the respective task error of the model. The circular flashes occur at second zero, cover 6 ommatidia in radius, and last for one second. Before and after, a grey-stimulus leads to a stationary state of the network. The traces are standardized.

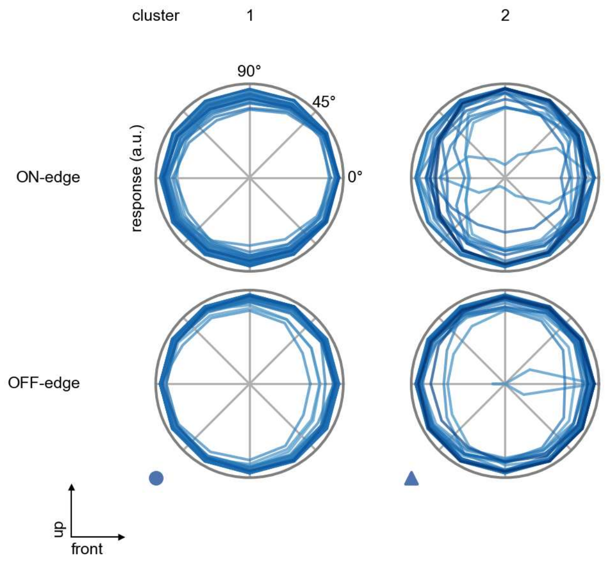

L3 - Figure 5: **Peak responses to moving edges.** The top row shows normalized peak responses to moving ON-edges, the bottom row shows normalized peak responses to moving OFF-edges. The edge-stimuli move in different directions from 0 to 360 degrees. Before, a grey-stimulus led the network to a stationary state. The responses from the different models in the different clusters (columns) overlay and are colored in the respective task error of the model. The peak responses are averaged over all assessed edge-speeds.

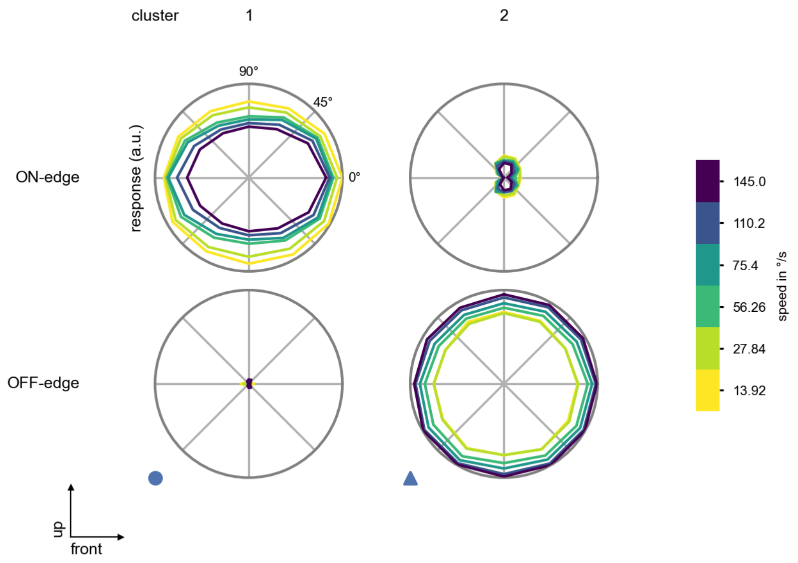

L3 - Figure 6: **Peak responses to moving edges from best-performing models.** The top row shows peak responses to moving ON-edges, the bottom row shows peak responses to moving OFF-edges. The edge-stimuli move in different directions from 0 to 360 degrees and at different speeds. Before, a grey-stimulus led the network to a stationary state. The columns show responses from the best-task-performing model in the respective cluster.

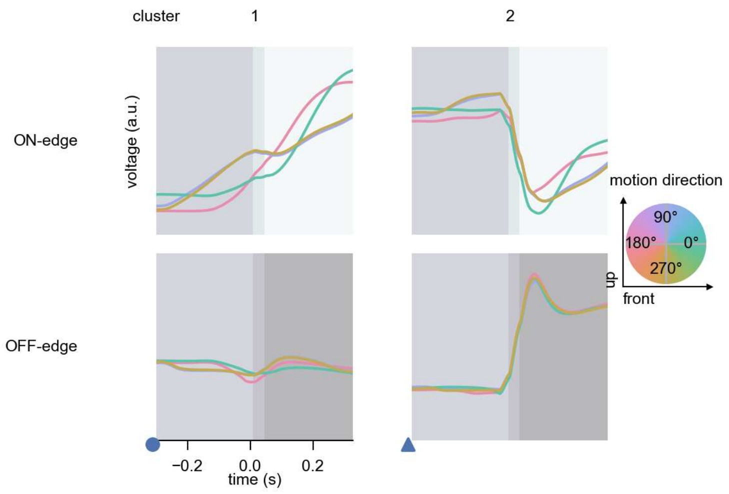

L3 - Figure 7: **Responses to moving edges from best-performing models.** The top row shows responses to moving ON-edges, the bottom row shows peak responses to moving OFF-edges. The edge-stimuli move in different directions from 0 to 360 degrees and at different speeds. Before, a grey-stimulus led the network to a stationary state. The columns show responses from the best-task-performing model in the respective cluster. Example responses to edges moving at 75.4°/s in all cardinal directions (green 0°, blue 90°, red 180°, yellow 270°) from -22.5 to 22.5° visual angle. Background contour indicates the edge, passing by the central column at second zero.

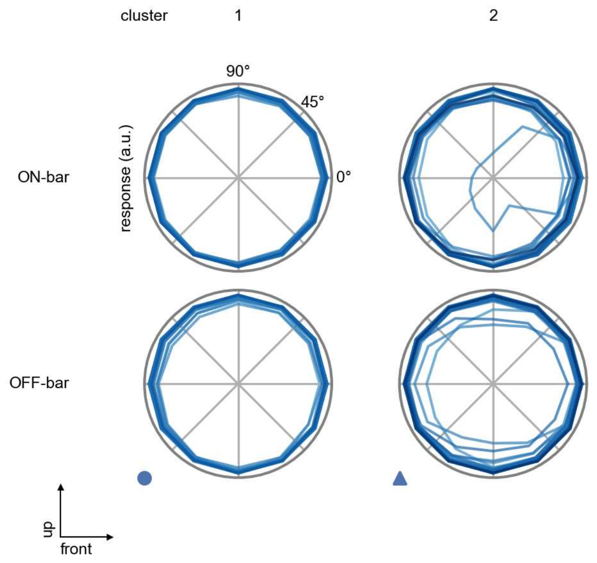

L3 - Figure 8: **Peak responses to moving bars.** The top row shows normalized peak responses to moving ON-bars, the bottom row shows normalized peak responses to moving OFF-bars. The bar-stimuli move in different directions from 0 to 360 degrees. Before, a grey-stimulus led the network to a stationary state. The responses from the different models in the different clusters (columns) overlay and are colored in the respective task error of the model. The peak responses are averaged over all assessed bar-speeds.

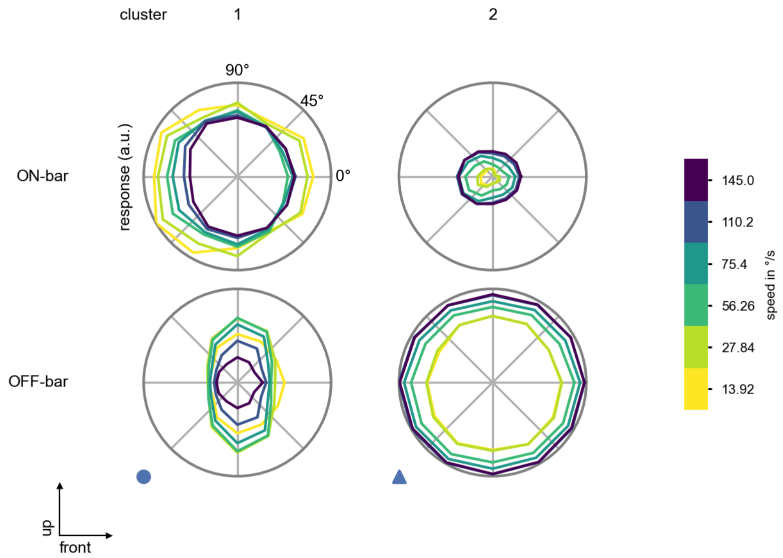

L3 - Figure 9: **Peak responses to moving bars from best-performing models.** The top row shows peak responses to moving ON-bars, the bottom row shows peak responses to moving OFF-bars. The bar-stimuli move in different directions from 0 to 360 degrees and at different speeds. Before, a grey-stimulus led the network to a stationary state. The columns show responses from the best-task-performing model in the respective cluster.

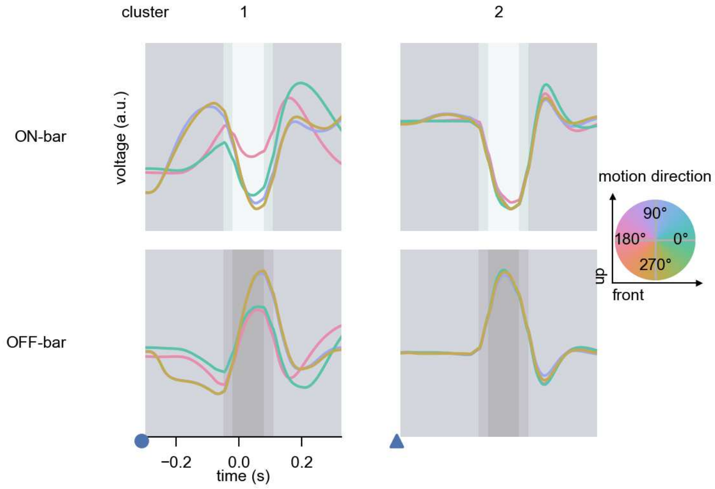

L3 - Figure 10: **Responses to moving bars from best-performing models.** The top row shows responses to moving ON-bars, the bottom row shows peak responses to moving OFF-bars. The bar-stimuli move in different directions from 0 to 360 degrees and at different speeds. Before, a grey-stimulus led the network to a stationary state. The columns show responses from the best-task-performing model in the respective cluster. Example responses to bars moving at 75.4°/s in all cardinal directions (green 0°, blue 90°, red 180°, yellow 270°) from -22.5 to 22.5° visual angle. Background contour indicates the bar, passing by the central column at second zero.

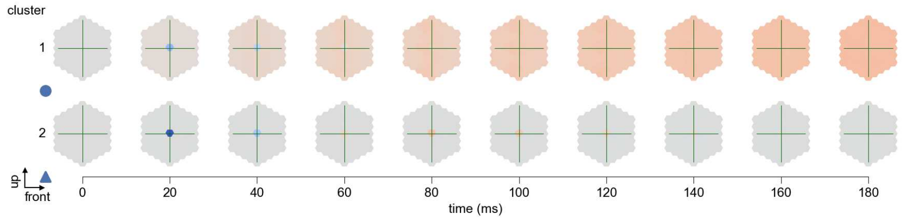

L3 - Figure 11: **Spatio-temporal receptive field.** Responses of the central cell to ON-impulses at single ommatidium impulse locations. The impulse occurs at second zero. Before, a grey-stimulus led the network to a stationary state. Responses are baseline-subtracted. Each row illustrates responses from the best-performing model of the respective cluster. Red indicated depolarization, blue indicates hyperpolarization.

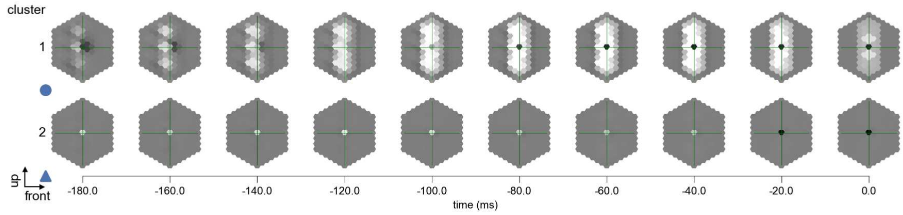

L3 - Figure 12: **Maximally excitatory stimuli.** Each row presents the regularized naturalistic-stimulus from the Sintel dataset that maximizes the cell type's central column response at second zero in the best-performing model of the respective cluster.

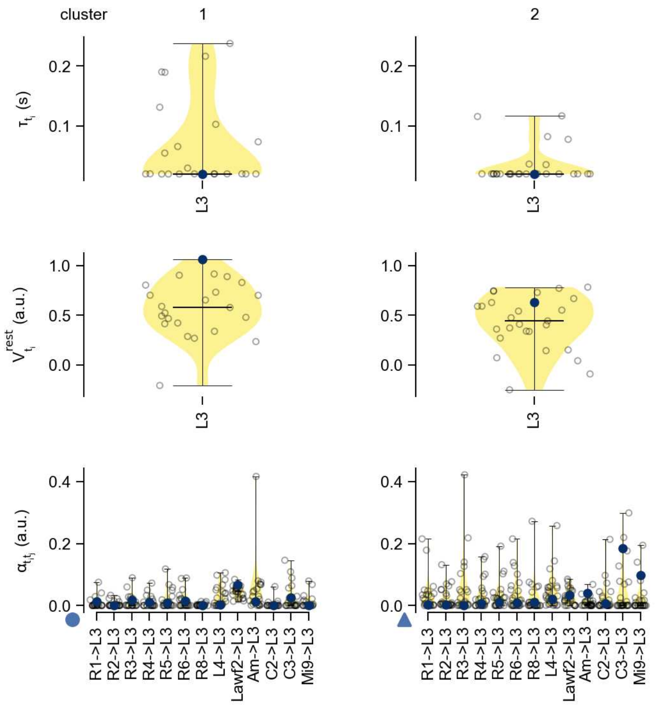

L3 - Figure 13: **Task-constrained parameters.** Each column shows the parameters inferred within the respective cluster. First row: learned time constants of the cell type. Second row: resting potentials of the cell type. Third row: scaling factors for the convolutional filters. The blue scatter represents the parameters from the best-performing model within the cluster.

# 12 L4

← Cell types

#### Figures

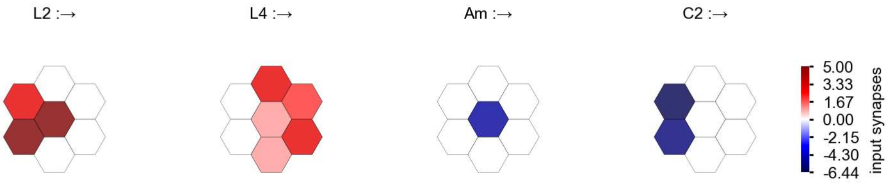

L4 - Figure 1: **Anatomical receptive fields.** Each colored hexagon is a projection, with the projection strength characterized by the average number of synapses that we count from the EM reconstruction. Red indicates excitatory synapses, blue indicates inhibitory synapses from inferred signs. Filters in the order of their total number of synapses.

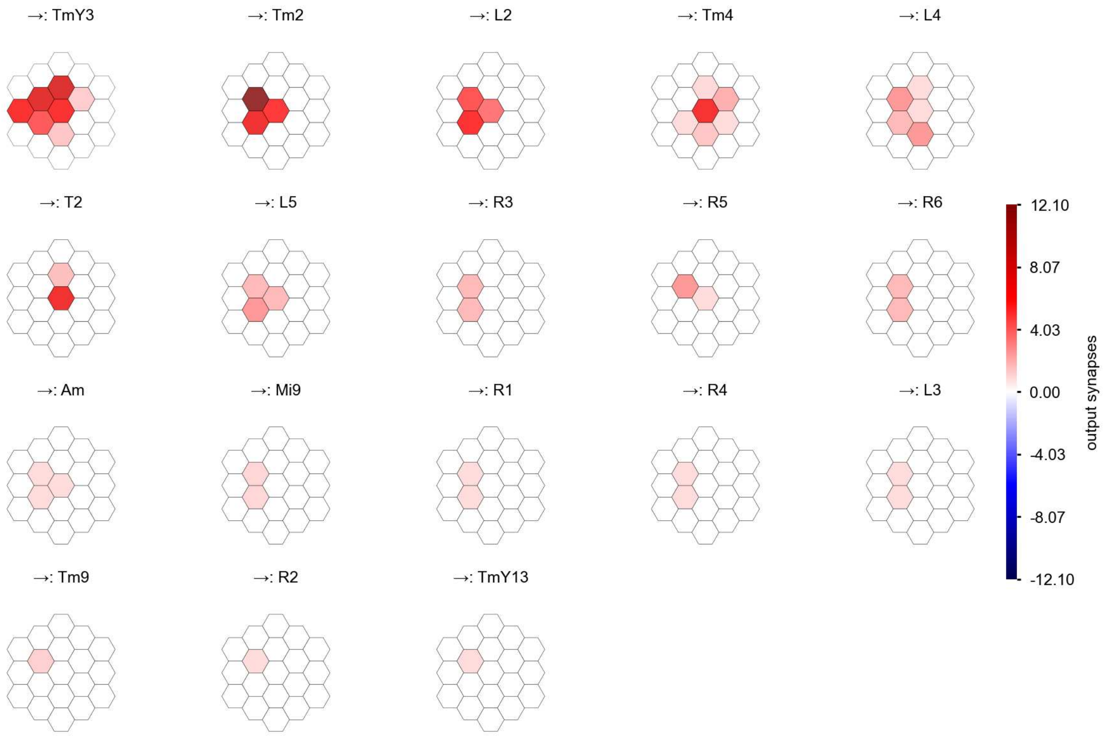

L4 - Figure 2: **Anatomical projective fields.** Each colored hexagon is a projection, with the projection strength characterized by the average number of synapses that we count from the EM reconstruction. Red indicates excitatory synapses, blue indicates inhibitory synapses from inferred signs. Filters in the order of their total number of synapses.

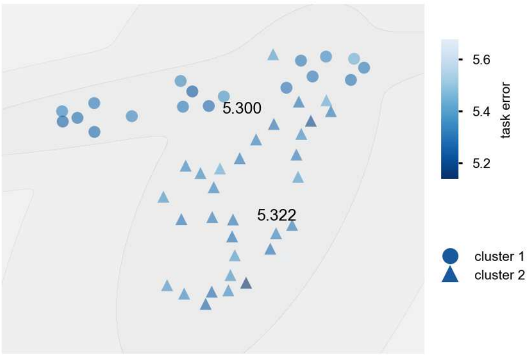

L4 - Figure 3: **Clustering of the responses to naturalistic stimuli.** From responses to naturalistic scenes from Sintel. Scatterpoints represent individual models colored by their task error.

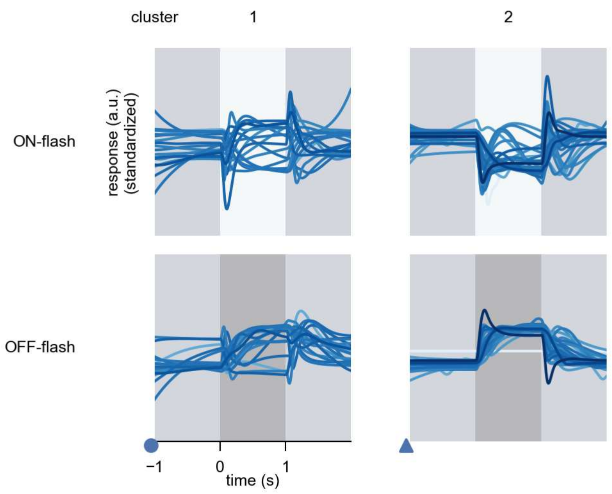

L4 - Figure 4: **Responses to flashes.** The top row shows responses to ON-flashes, the bottom row shows responses to OFF-flashes. The responses from the different models in the different clusters (columns) overlay and are colored in the respective task error of the model. The circular flashes occur at second zero, cover 6 ommatidia in radius, and last for one second. Before and after, a grey-stimulus leads to a stationary state of the network. The traces are standardized.

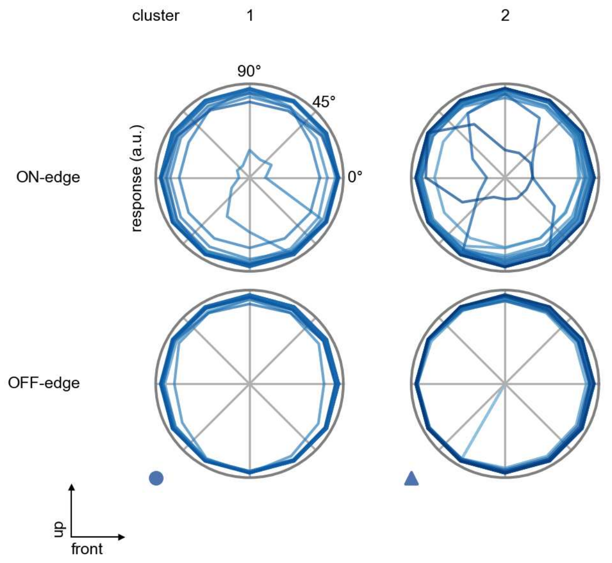

L4 - Figure 5: **Peak responses to moving edges.** The top row shows normalized peak responses to moving ON-edges, the bottom row shows normalized peak responses to moving OFF-edges. The edge-stimuli move in different directions from 0 to 360 degrees. Before, a grey-stimulus led the network to a stationary state. The responses from the different models in the different clusters (columns) overlay and are colored in the respective task error of the model. The peak responses are averaged over all assessed edge-speeds.

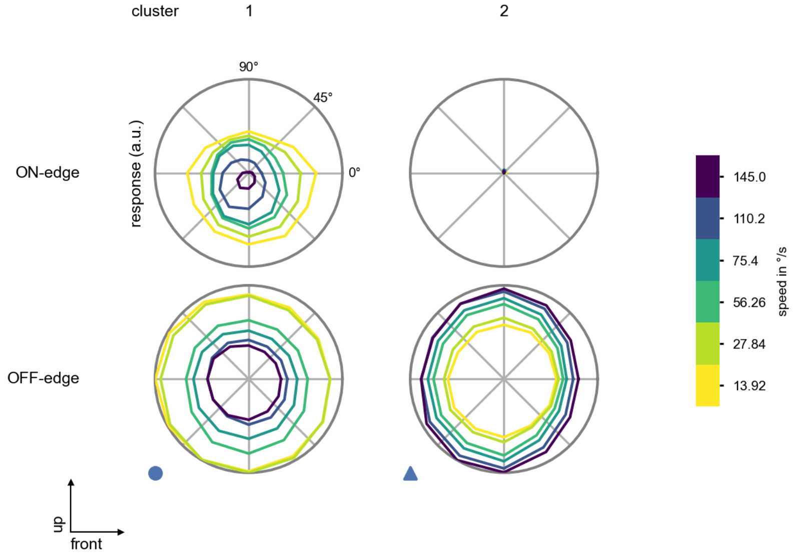

L4 - Figure 6: **Peak responses to moving edges from best-performing models.** The top row shows peak responses to moving ON-edges, the bottom row shows peak responses to moving OFF-edges. The edge-stimuli move in different directions from 0 to 360 degrees and at different speeds. Before, a grey-stimulus led the network to a stationary state. The columns show responses from the best-task-performing model in the respective cluster.

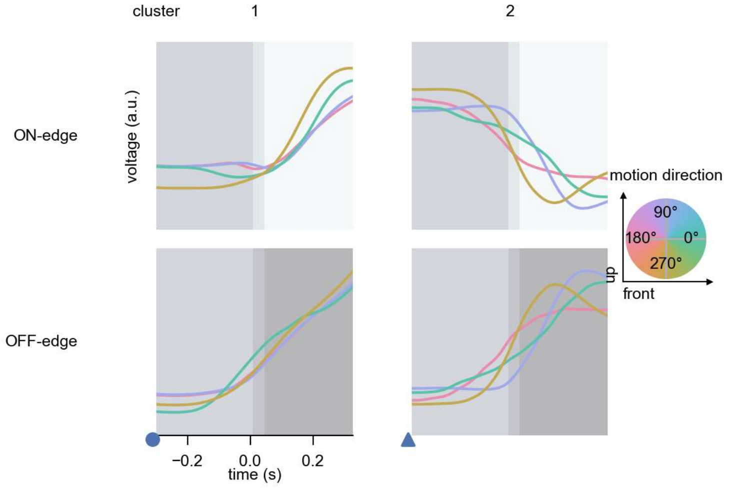

L4 - Figure 7: **Responses to moving edges from best-performing models.** The top row shows responses to moving ON-edges, the bottom row shows peak responses to moving OFF-edges. The edge-stimuli move in different directions from 0 to 360 degrees and at different speeds. Before, a grey-stimulus led the network to a stationary state. The columns show responses from the best-task-performing model in the respective cluster. Example responses to edges moving at 75.4°/s in all cardinal directions (green 0°, blue 90°, red 180°, yellow 270°) from -22.5 to 22.5° visual angle. Background contour indicates the edge, passing by the central column at second zero.

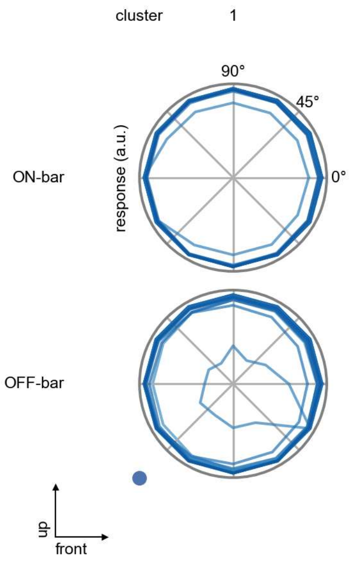

L4 - Figure 8: **Peak responses to moving bars.** The top row shows normalized peak responses to moving ON-bars, the bottom row shows normalized peak responses to moving OFF-bars. The bar-stimuli move in different directions from 0 to 360 degrees. Before, a grey-stimulus led the network to a stationary state. The responses from the different models in the different clusters (columns) overlay and are colored in the respective task error of the model. The peak responses are averaged over all assessed bar-speeds.

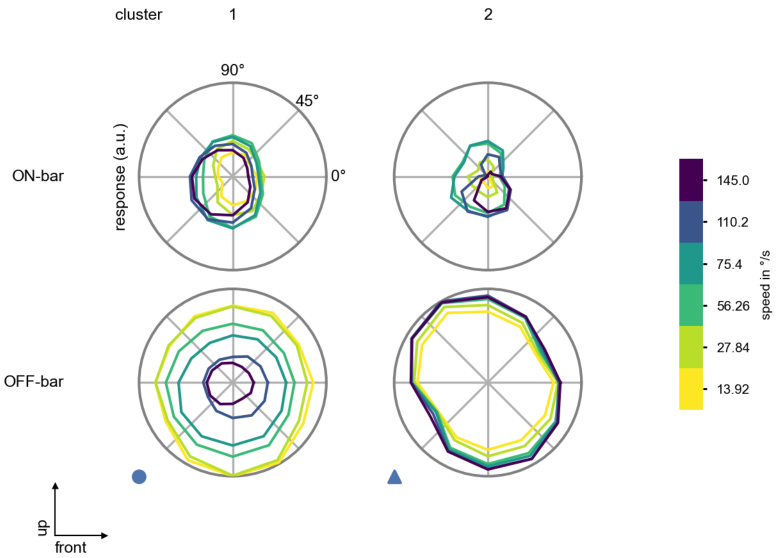

L4 - Figure 9: **Peak responses to moving bars from best-performing models.** The top row shows peak responses to moving ON-bars, the bottom row shows peak responses to moving OFF-bars. The bar-stimuli move in different directions from 0 to 360 degrees and at different speeds. Before, a grey-stimulus led the network to a stationary state. The columns show responses from the best-task-performing model in the respective cluster.

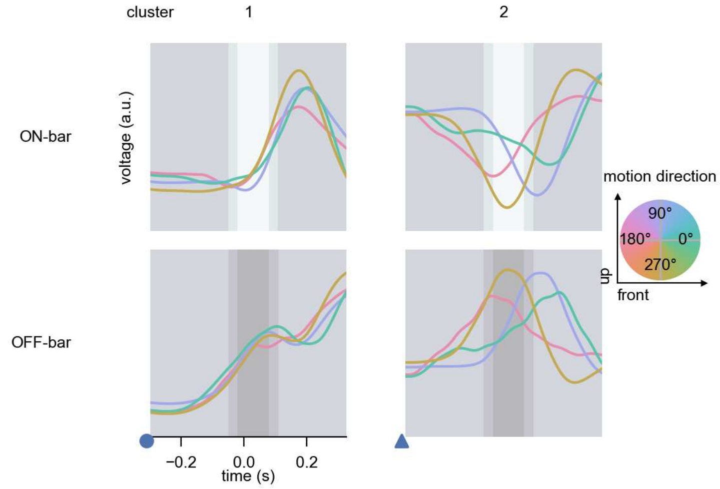

L4 - Figure 10: **Responses to moving bars from best-performing models.** The top row shows responses to moving ON-bars, the bottom row shows peak responses to moving OFF-bars. The bar-stimuli move in different directions from 0 to 360 degrees and at different speeds. Before, a grey-stimulus led the network to a stationary state. The columns show responses from the best-task-performing model in the respective cluster. Example responses to bars moving at 75.4°/s in all cardinal directions (green 0°, blue 90°, red 180°, yellow 270°) from -22.5 to 22.5° visual angle. Background contour indicates the bar, passing by the central column at second zero.

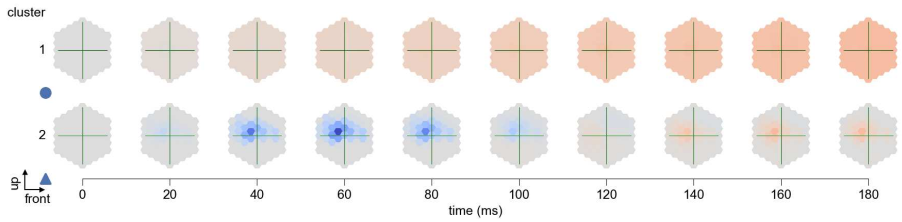

L4 - Figure 11: **Spatio-temporal receptive field.** Responses of the central cell to ON-impulses at single ommatidium impulse locations. The impulse occurs at second zero. Before, a grey-stimulus led the network to a stationary state. Responses are baseline-subtracted. Each row illustrates responses from the best-performing model of the respective cluster. Red indicated depolarization, blue indicates hyperpolarization.

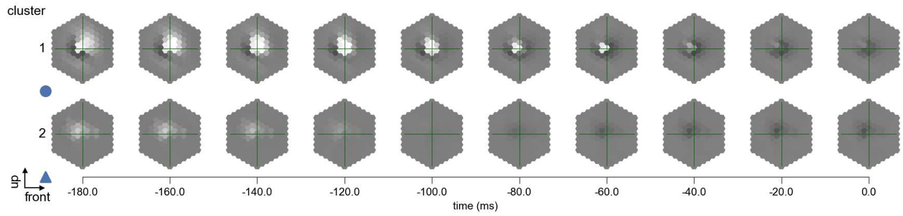

L4 - Figure 12: **Maximally excitatory stimuli.** Each row presents the regularized naturalistic-stimulus from the Sintel dataset that maximizes the cell type's central column response at second zero in the best-performing model of the respective cluster.

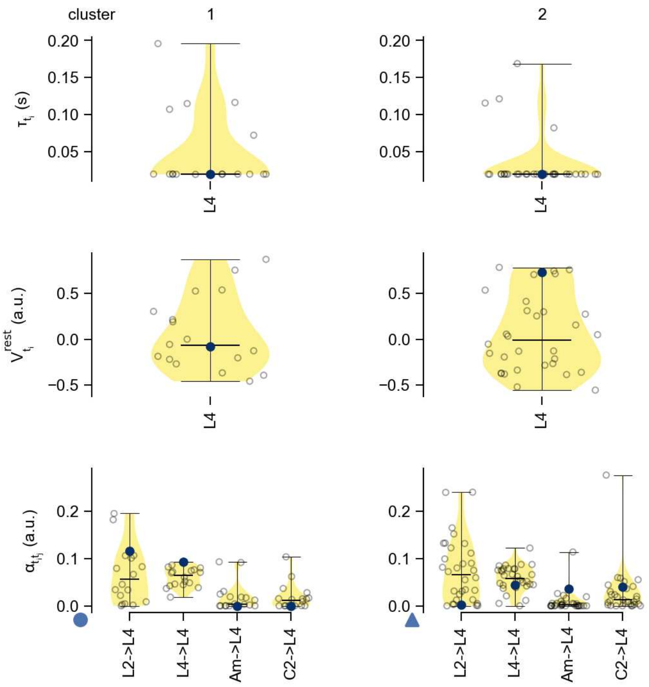

L4 - Figure 13: **Task-constrained parameters.** Each column shows the parameters inferred within the respective cluster. First row: learned time constants of the cell type. Second row: resting potentials of the cell type. Third row: scaling factors for the convolutional filters. The blue scatter represents the parameters from the best-performing model within the cluster.

# 13 L5

← Cell types

#### Figures

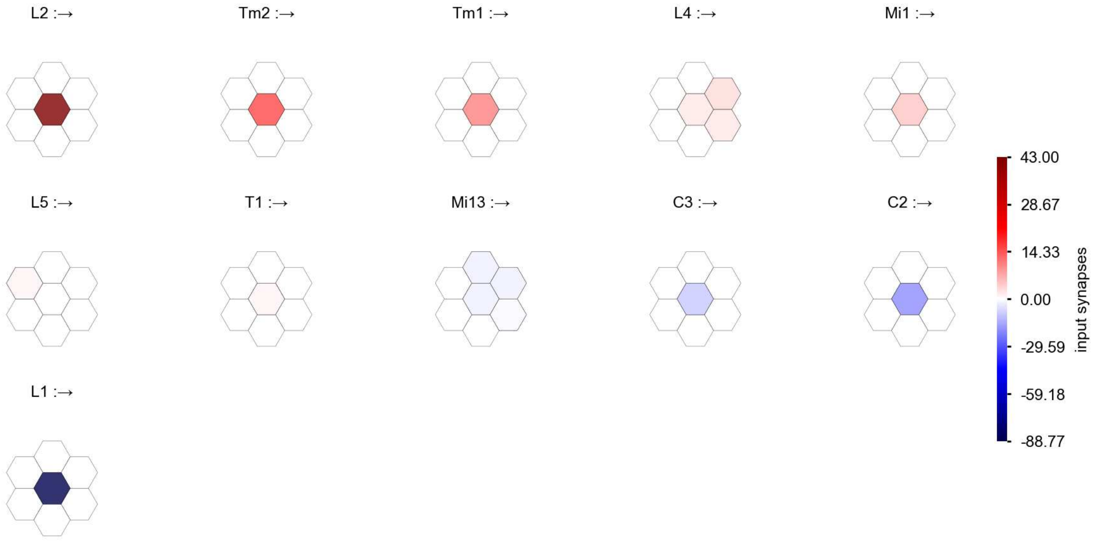

L5 - Figure 1: **Anatomical receptive fields.** Each colored hexagon is a projection, with the projection strength characterized by the average number of synapses that we count from the EM reconstruction. Red indicates excitatory synapses, blue indicates inhibitory synapses from inferred signs. Filters in the order of their total number of synapses.

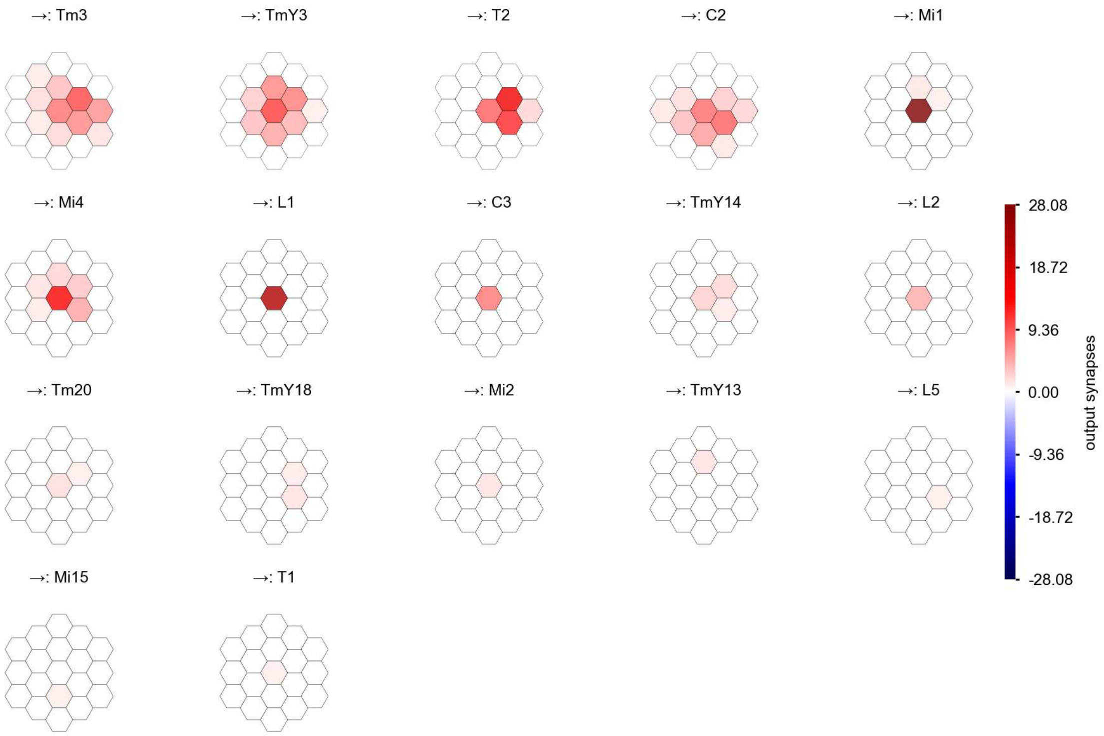

L5 - Figure 2: **Anatomical projective fields.** Each colored hexagon is a projection, with the projection strength characterized by the average number of synapses that we count from the EM reconstruction. Red indicates excitatory synapses, blue indicates inhibitory synapses from inferred signs. Filters in the order of their total number of synapses.

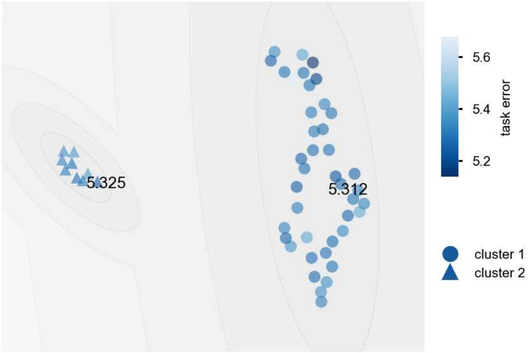

L5 - Figure 3: **Clustering of the responses to naturalistic stimuli.** From responses to naturalistic scenes from Sintel. Scatterpoints represent individual models colored by their task error.

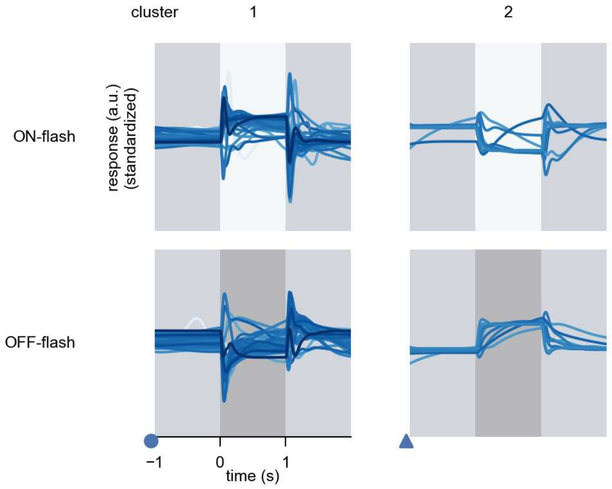

L5 - Figure 4: **Responses to flashes.** The top row shows responses to ON-flashes, the bottom row shows responses to OFF-flashes. The responses from the different models in the different clusters (columns) overlay and are colored in the respective task error of the model. The circular flashes occur at second zero, cover 6 ommatidia in radius, and last for one second. Before and after, a grey-stimulus leads to a stationary state of the network. The traces are standardized.

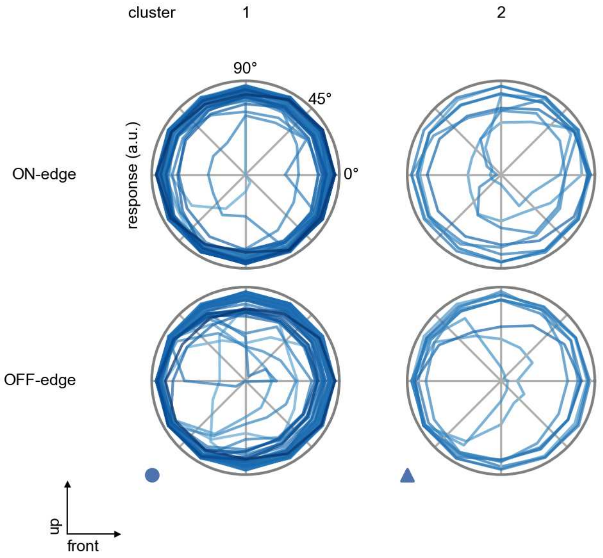

L5 - Figure 5: **Peak responses to moving edges.** The top row shows normalized peak responses to moving ON-edges, the bottom row shows normalized peak responses to moving OFF-edges. The edge-stimuli move in different directions from 0 to 360 degrees. Before, a grey-stimulus led the network to a stationary state. The responses from the different models in the different clusters (columns) overlay and are colored in the respective task error of the model. The peak responses are averaged over all assessed edge-speeds.

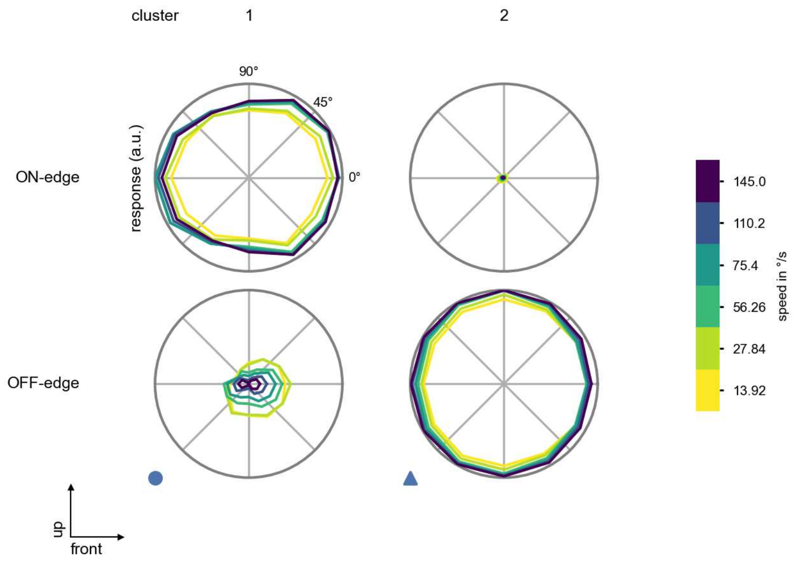

L5 - Figure 6: **Peak responses to moving edges from best-performing models.** The top row shows peak responses to moving ON-edges, the bottom row shows peak responses to moving OFF-edges. The edge-stimuli move in different directions from 0 to 360 degrees and at different speeds. Before, a grey-stimulus led the network to a stationary state. The columns show responses from the best-task-performing model in the respective cluster.

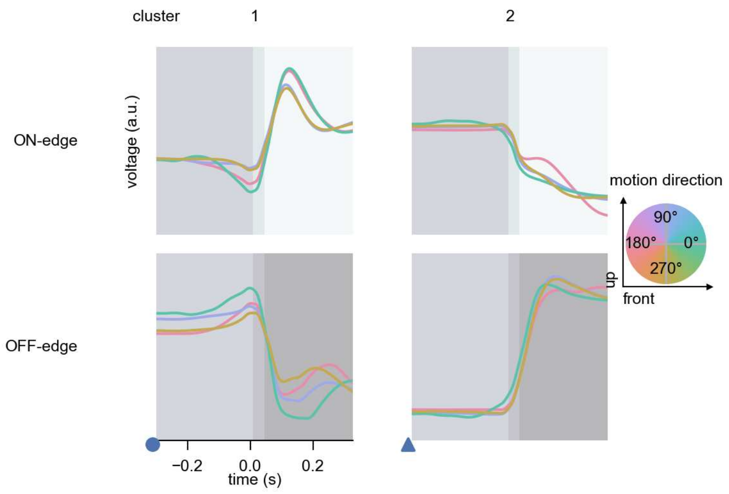

L5 - Figure 7: **Responses to moving edges from best-performing models.** The top row shows responses to moving ON-edges, the bottom row shows peak responses to moving OFF-edges. The edge-stimuli move in different directions from 0 to 360 degrees and at different speeds. Before, a grey-stimulus led the network to a stationary state. The columns show responses from the best-task-performing model in the respective cluster. Example responses to edges moving at 75.4°/s in all cardinal directions (green 0°, blue 90°, red 180°, yellow 270°) from -22.5 to 22.5° visual angle. Background contour indicates the edge, passing by the central column at second zero.

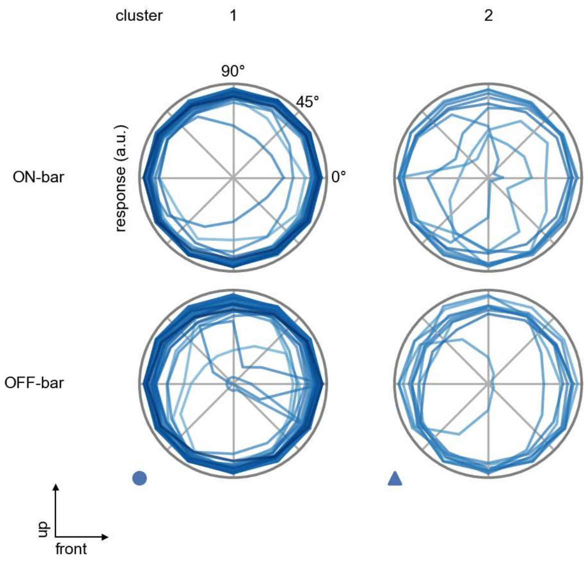

L5 - Figure 8: **Peak responses to moving bars.** The top row shows normalized peak responses to moving ON-bars, the bottom row shows normalized peak responses to moving OFF-bars. The bar-stimuli move in different directions from 0 to 360 degrees. Before, a grey-stimulus led the network to a stationary state. The responses from the different models in the different clusters (columns) overlay and are colored in the respective task error of the model. The peak responses are averaged over all assessed bar-speeds.

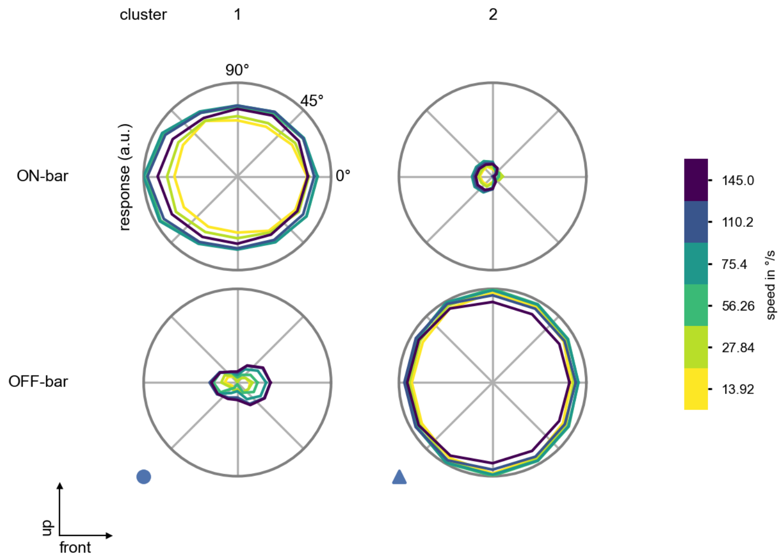

L5 - Figure 9: **Peak responses to moving bars from best-performing models.** The top row shows peak responses to moving ON-bars, the bottom row shows peak responses to moving OFF-bars. The bar-stimuli move in different directions from 0 to 360 degrees and at different speeds. Before, a grey-stimulus led the network to a stationary state. The columns show responses from the best-task-performing model in the respective cluster.

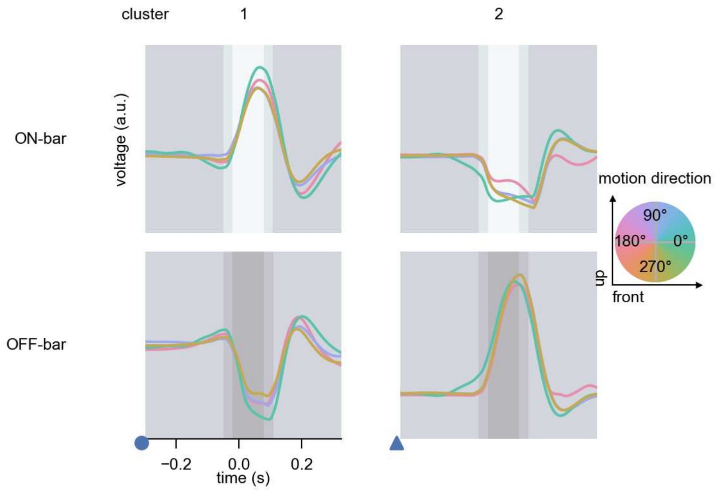

L5 - Figure 10: **Responses to moving bars from best-performing models.** The top row shows responses to moving ON-bars, the bottom row shows peak responses to moving OFF-bars. The bar-stimuli move in different directions from 0 to 360 degrees and at different speeds. Before, a grey-stimulus led the network to a stationary state. The columns show responses from the best-task-performing model in the respective cluster. Example responses to bars moving at 75.4°/s in all cardinal directions (green 0°, blue 90°, red 180°, yellow 270°) from -22.5 to 22.5° visual angle. Background contour indicates the bar, passing by the central column at second zero.

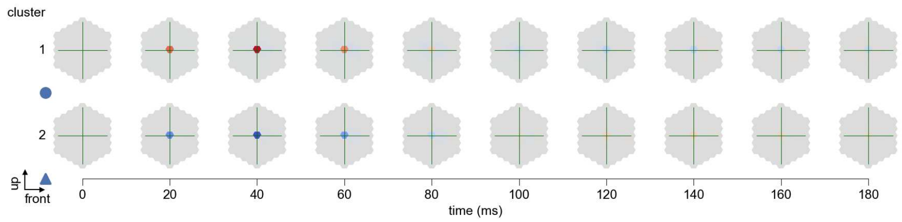

L5 - Figure 11: **Spatio-temporal receptive field.** Responses of the central cell to ON-impulses at single ommatidium impulse locations. The impulse occurs at second zero. Before, a grey-stimulus led the network to a stationary state. Responses are baseline-subtracted. Each row illustrates responses from the best-performing model of the respective cluster. Red indicated depolarization, blue indicates hyperpolarization.

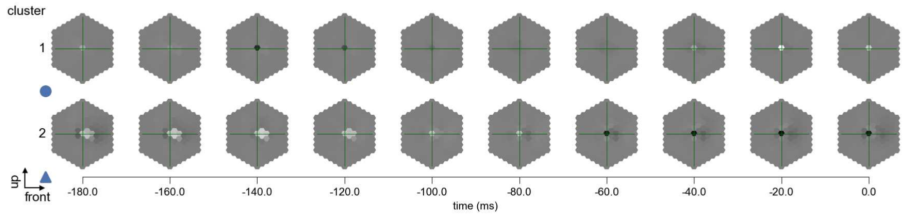

L5 - Figure 12: **Maximally excitatory stimuli.** Each row presents the regularized naturalistic-stimulus from the Sintel dataset that maximizes the cell type's central column response at second zero in the best-performing model of the respective cluster.

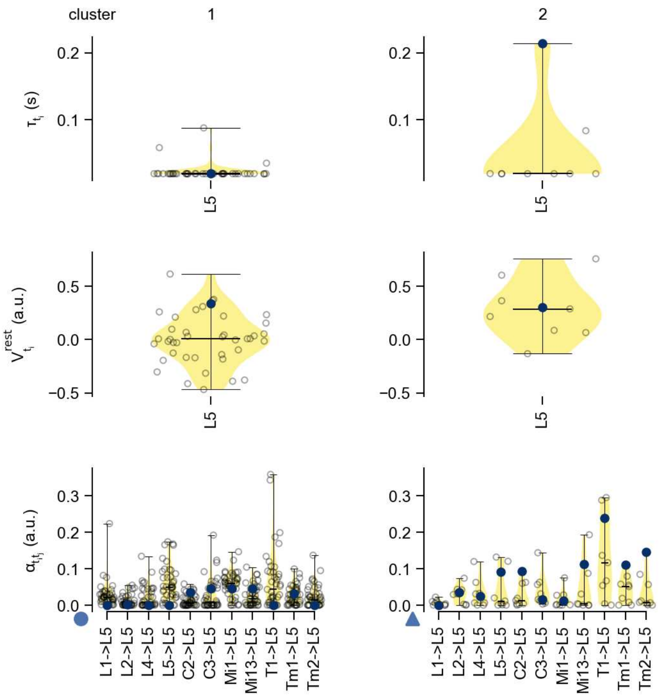

L5 - Figure 13: **Task-constrained parameters.** Each column shows the parameters inferred within the respective cluster. First row: learned time constants of the cell type. Second row: resting potentials of the cell type. Third row: scaling factors for the convolutional filters. The blue scatter represents the parameters from the best-performing model within the cluster.

### 14 Lawf1

← Cell types

#### Figures

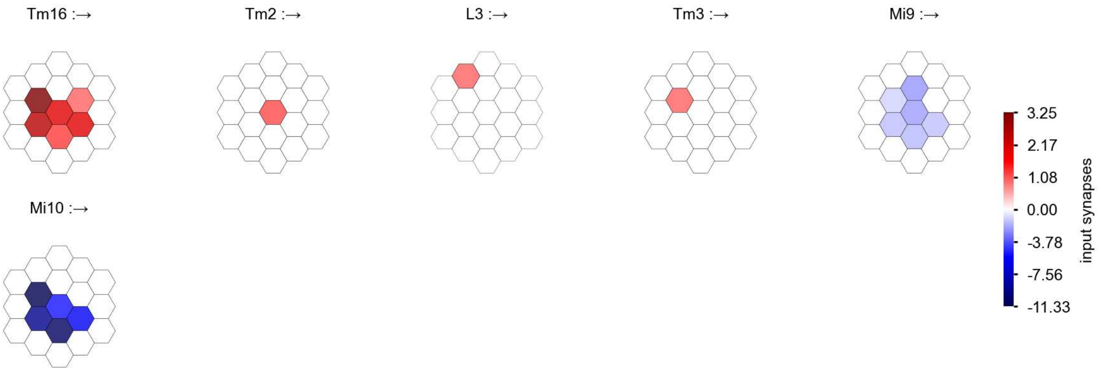

Lawf1 - Figure 1: **Anatomical receptive fields.** Each colored hexagon is a projection, with the projection strength characterized by the average number of synapses that we count from the EM reconstruction. Red indicates excitatory synapses, blue indicates inhibitory synapses from inferred signs. Filters in the order of their total number of synapses.

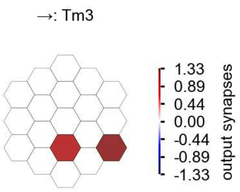

Lawf1 - Figure 2: **Anatomical projective fields.** Each colored hexagon is a projection, with the projection strength characterized by the average number of synapses that we count from the EM reconstruction. Red indicates excitatory synapses, blue indicates inhibitory synapses from inferred signs. Filters in the order of their total number of synapses.

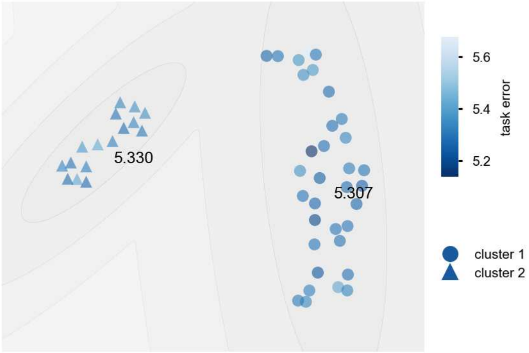

Lawf1 - Figure 3: **Clustering of the responses to naturalistic stimuli.** From responses to naturalistic scenes from Sintel. Scatterpoints represent individual models colored by their task error.

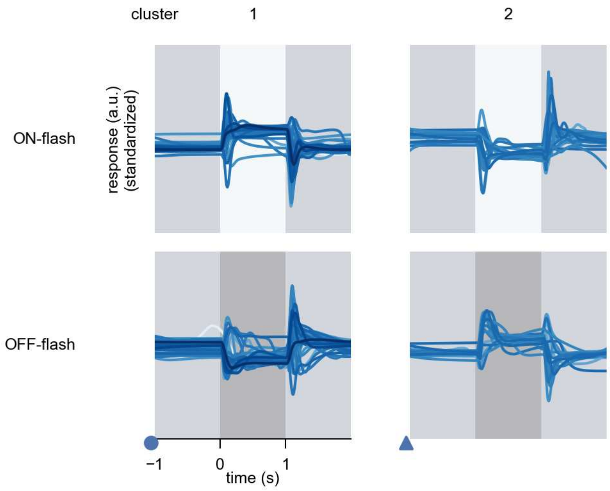

Lawf1 - Figure 4: **Responses to flashes.** The top row shows responses to ON-flashes, the bottom row shows responses to OFF-flashes. The responses from the different models in the different clusters (columns) overlay and are colored in the respective task error of the model. The circular flashes occur at second zero, cover 6 ommatidia in radius, and last for one second. Before and after, a grey-stimulus leads to a stationary state of the network. The traces are standardized.

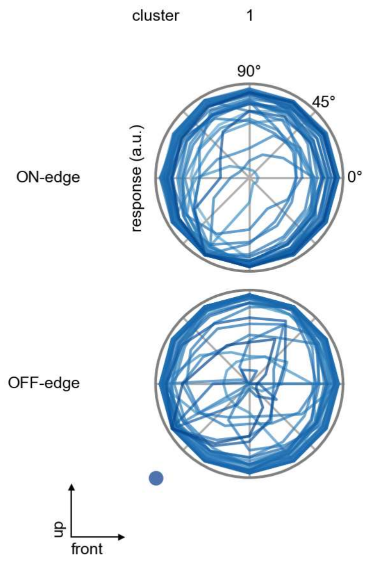

Lawf1 - Figure 5: **Peak responses to moving edges.** The top row shows normalized peak responses to moving ON-edges, the bottom row shows normalized peak responses to moving OFF-edges. The edge-stimuli move in different directions from 0 to 360 degrees. Before, a grey-stimulus led the network to a stationary state. The responses from the different models in the different clusters (columns) overlay and are colored in the respective task error of the model. The peak responses are averaged over all assessed edge-speeds.

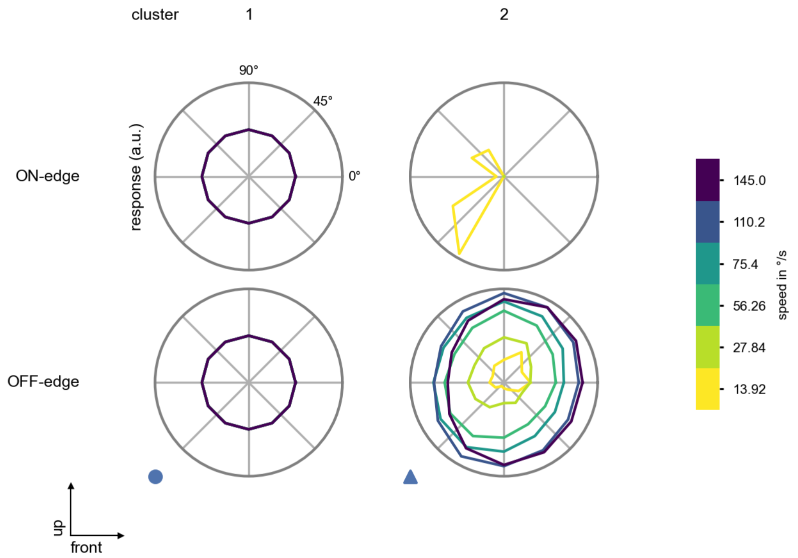

Lawf1 - Figure 6: **Peak responses to moving edges from best-performing models.** The top row shows peak responses to moving ON-edges, the bottom row shows peak responses to moving OFF-edges. The edge-stimuli move in different directions from 0 to 360 degrees and at different speeds. Before, a grey-stimulus led the network to a stationary state. The columns show responses from the best-task-performing model in the respective cluster.

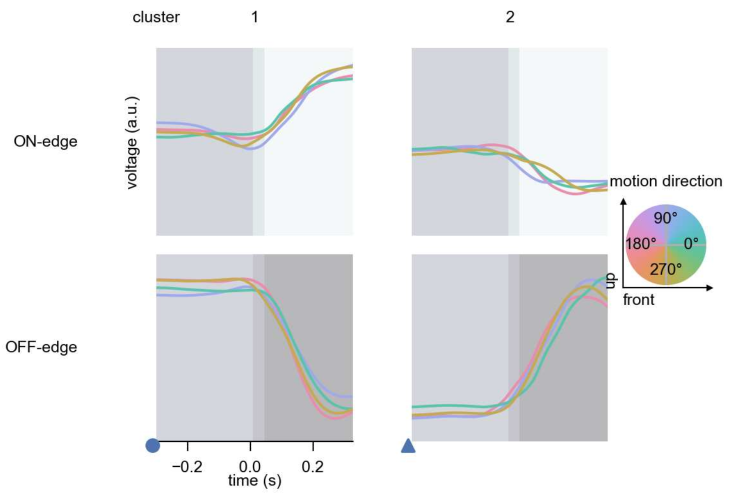

Lawf1 - Figure 7: **Responses to moving edges from best-performing models.** The top row shows responses to moving ON-edges, the bottom row shows peak responses to moving OFF-edges. The edge-stimuli move in different directions from 0 to 360 degrees and at different speeds. Before, a grey-stimulus led the network to a stationary state. The columns show responses from the best-task-performing model in the respective cluster. Example responses to edges moving at 75.4°/s in all cardinal directions (green 0°, blue 90°, red 180°, yellow 270°) from -22.5 to 22.5° visual angle. Background contour indicates the edge, passing by the central column at second zero.

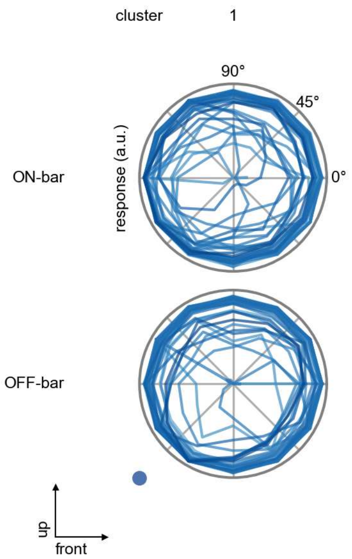

Lawf1 - Figure 8: **Peak responses to moving bars.** The top row shows normalized peak responses to moving ON-bars, the bottom row shows normalized peak responses to moving OFF-bars. The bar-stimuli move in different directions from 0 to 360 degrees. Before, a grey-stimulus led the network to a stationary state. The responses from the different models in the different clusters (columns) overlay and are colored in the respective task error of the model. The peak responses are averaged over all assessed bar-speeds.

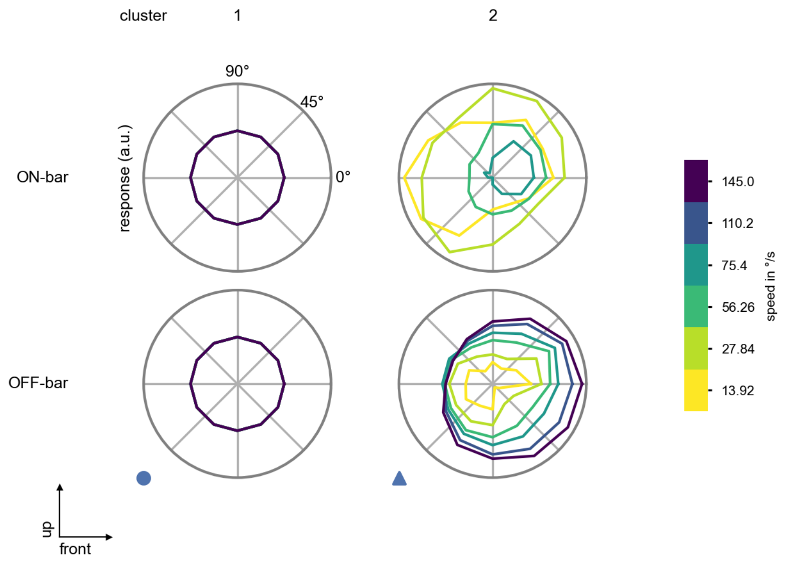

Lawf1 - Figure 9: **Peak responses to moving bars from best-performing models.** The top row shows peak responses to moving ON-bars, the bottom row shows peak responses to moving OFF-bars. The bar-stimuli move in different directions from 0 to 360 degrees and at different speeds. Before, a grey-stimulus led the network to a stationary state. The columns show responses from the best-task-performing model in the respective cluster.

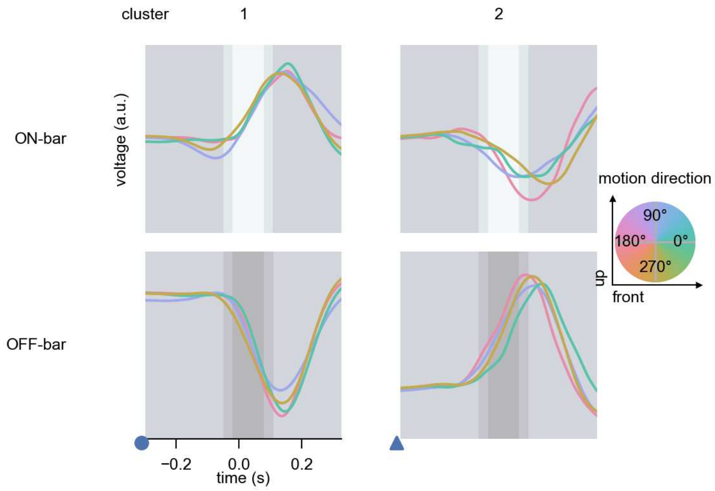

Lawf1 - Figure 10: **Responses to moving bars from best-performing models.** The top row shows responses to moving ON-bars, the bottom row shows peak responses to moving OFF-bars. The bar-stimuli move in different directions from 0 to 360 degrees and at different speeds. Before, a grey-stimulus led the network to a stationary state. The columns show responses from the best-task-performing model in the respective cluster. Example responses to bars moving at 75.4°/s in all cardinal directions (green 0°, blue 90°, red 180°, yellow 270°) from -22.5 to 22.5° visual angle. Background contour indicates the bar, passing by the central column at second zero.

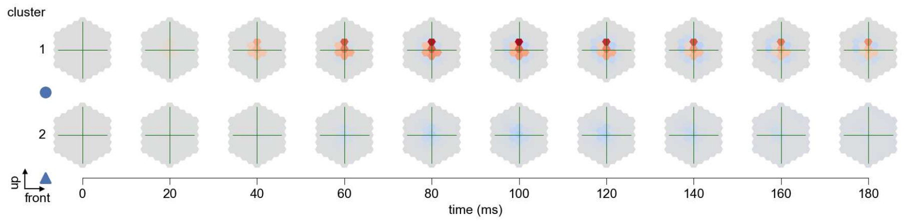

Lawf1 - Figure 11: **Spatio-temporal receptive field.** Responses of the central cell to ON-impulses at single ommatidium impulse locations. The impulse occurs at second zero. Before, a grey-stimulus led the network to a stationary state. Responses are baseline-subtracted. Each row illustrates responses from the best-performing model of the respective cluster. Red indicated depolarization, blue indicates hyperpolarization.

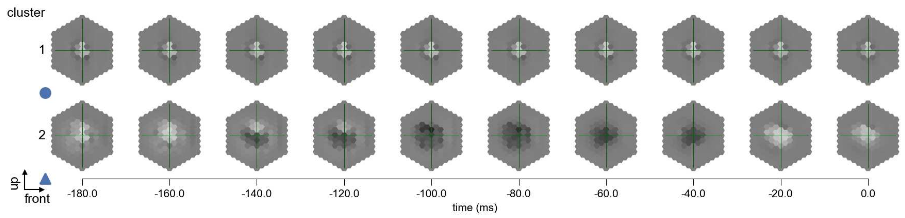

Lawf1 - Figure 12: **Maximally excitatory stimuli.** Each row presents the regularized naturalistic-stimulus from the Sintel dataset that maximizes the cell type's central column response at second zero in the best-performing model of the respective cluster.

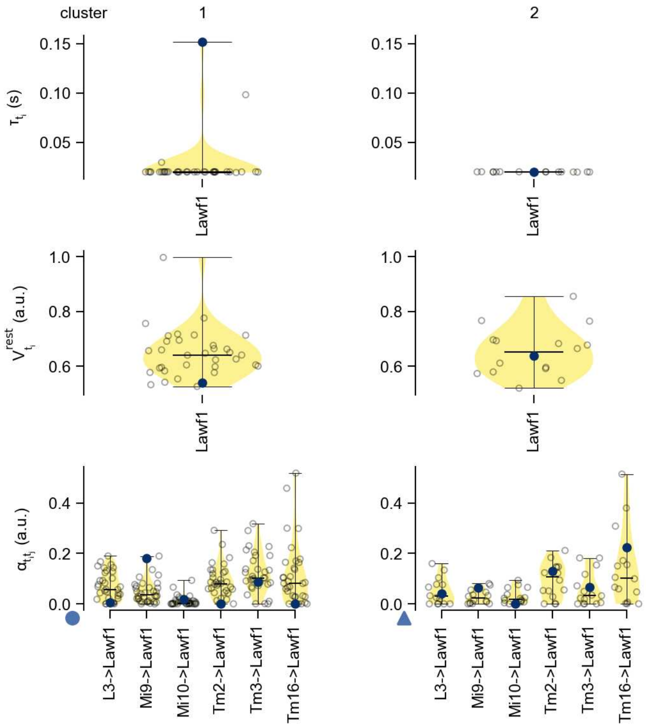

Lawf1 - Figure 13: **Task-constrained parameters.** Each column shows the parameters inferred within the respective cluster. First row: learned time constants of the cell type. Second row: resting potentials of the cell type. Third row: scaling factors for the convolutional filters. The blue scatter represents the parameters from the best-performing model within the cluster.

### 15 Lawf2

← Cell types

#### Figures

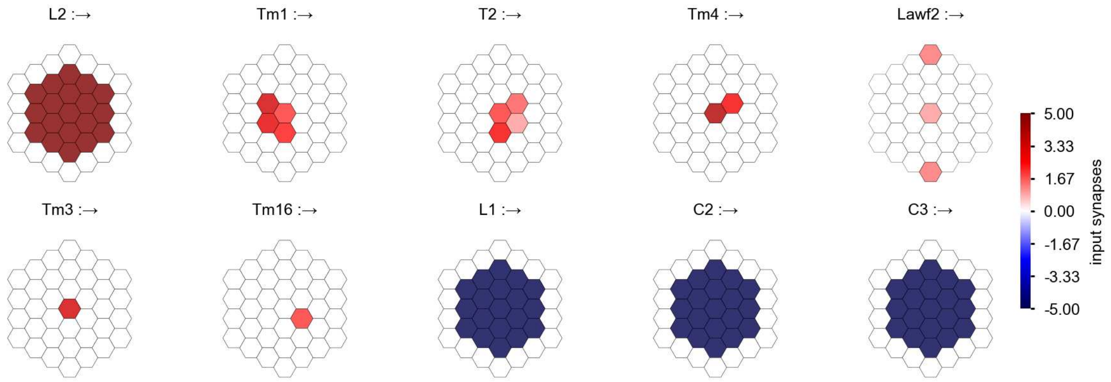

Lawf2 - Figure 1: **Anatomical receptive fields.** Each colored hexagon is a projection, with the projection strength characterized by the average number of synapses that we count from the EM reconstruction. Red indicates excitatory synapses, blue indicates inhibitory synapses from inferred signs. Filters in the order of their total number of synapses.

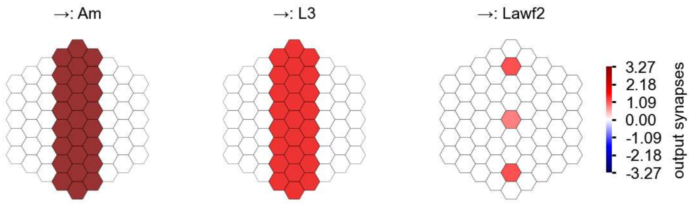

Lawf2 - Figure 2: **Anatomical projective fields.** Each colored hexagon is a projection, with the projection strength characterized by the average number of synapses that we count from the EM reconstruction. Red indicates excitatory synapses, blue indicates inhibitory synapses from inferred signs. Filters in the order of their total number of synapses.

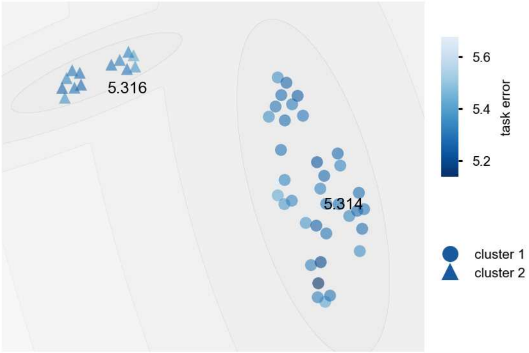

Lawf2 - Figure 3: **Clustering of the responses to naturalistic stimuli.** From responses to naturalistic scenes from Sintel. Scatterpoints represent individual models colored by their task error.

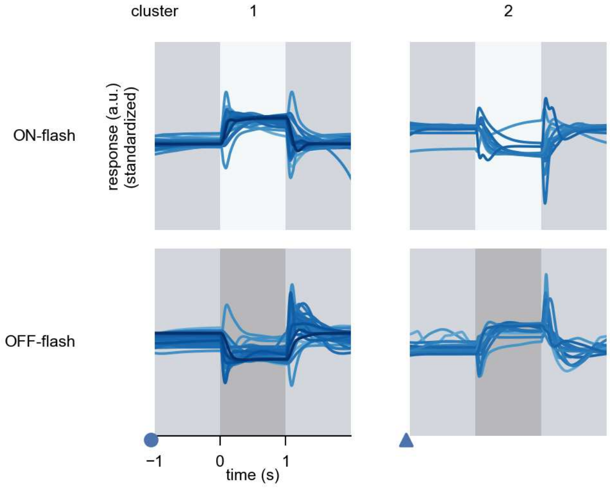

Lawf2 - Figure 4: **Responses to flashes.** The top row shows responses to ON-flashes, the bottom row shows responses to OFF-flashes. The responses from the different models in the different clusters (columns) overlay and are colored in the respective task error of the model. The circular flashes occur at second zero, cover 6 ommatidia in radius, and last for one second. Before and after, a grey-stimulus leads to a stationary state of the network. The traces are standardized.

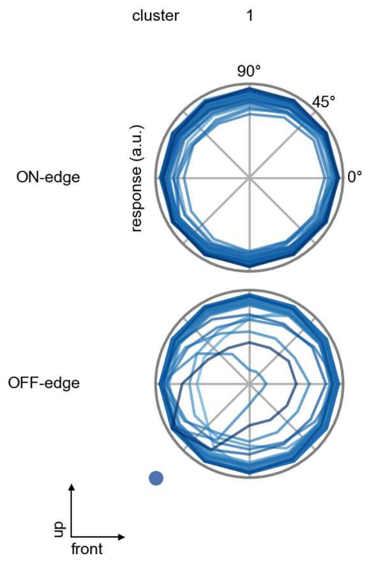

Lawf2 - Figure 5: **Peak responses to moving edges.** The top row shows normalized peak responses to moving ON-edges, the bottom row shows normalized peak responses to moving OFF-edges. The edge-stimuli move in different directions from 0 to 360 degrees. Before, a grey-stimulus led the network to a stationary state. The responses from the different models in the different clusters (columns) overlay and are colored in the respective task error of the model. The peak responses are averaged over all assessed edge-speeds.

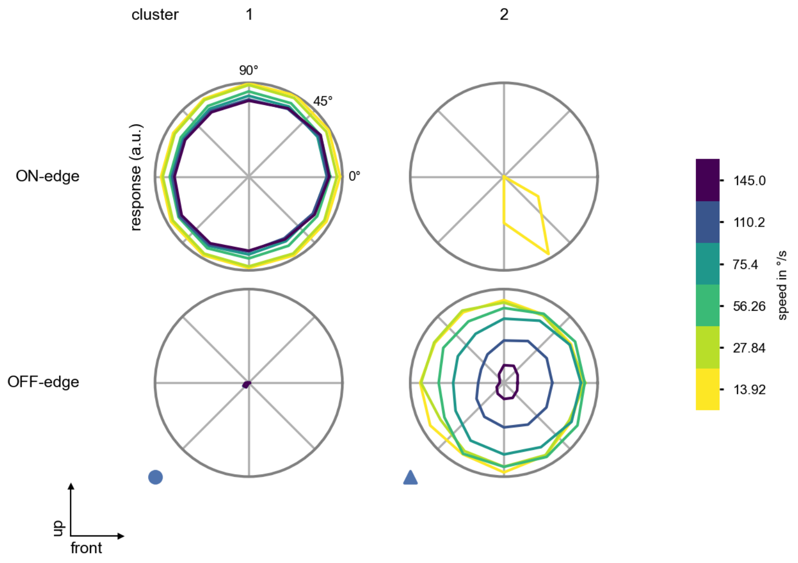

Lawf2 - Figure 6: **Peak responses to moving edges from best-performing models.** The top row shows peak responses to moving ON-edges, the bottom row shows peak responses to moving OFF-edges. The edge-stimuli move in different directions from 0 to 360 degrees and at different speeds. Before, a grey-stimulus led the network to a stationary state. The columns show responses from the best-task-performing model in the respective cluster.

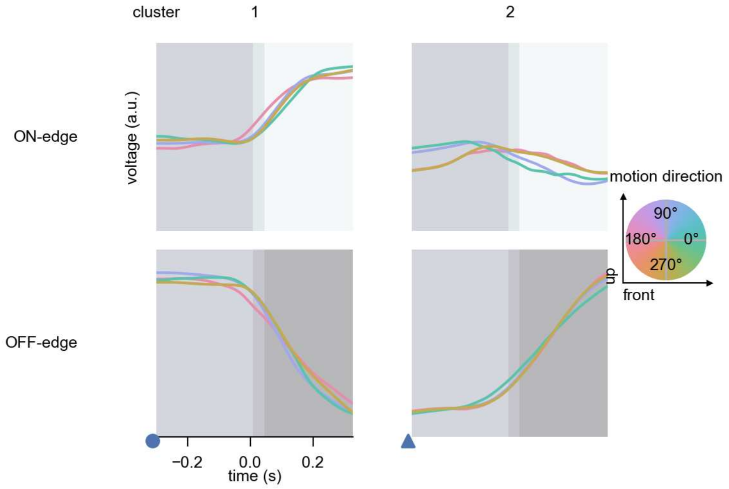

Lawf2 - Figure 7: **Responses to moving edges from best-performing models.** The top row shows responses to moving ON-edges, the bottom row shows peak responses to moving OFF-edges. The edge-stimuli move in different directions from 0 to 360 degrees and at different speeds. Before, a grey-stimulus led the network to a stationary state. The columns show responses from the best-task-performing model in the respective cluster. Example responses to edges moving at 75.4°/s in all cardinal directions (green 0°, blue 90°, red 180°, yellow 270°) from -22.5 to 22.5° visual angle. Background contour indicates the edge, passing by the central column at second zero.

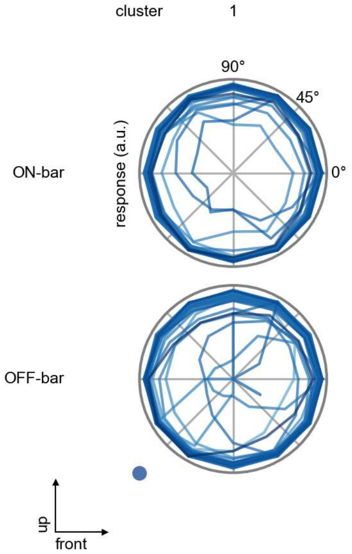

Lawf2 - Figure 8: **Peak responses to moving bars.** The top row shows normalized peak responses to moving ON-bars, the bottom row shows normalized peak responses to moving OFF-bars. The bar-stimuli move in different directions from 0 to 360 degrees. Before, a grey-stimulus led the network to a stationary state. The responses from the different models in the different clusters (columns) overlay and are colored in the respective task error of the model. The peak responses are averaged over all assessed bar-speeds.

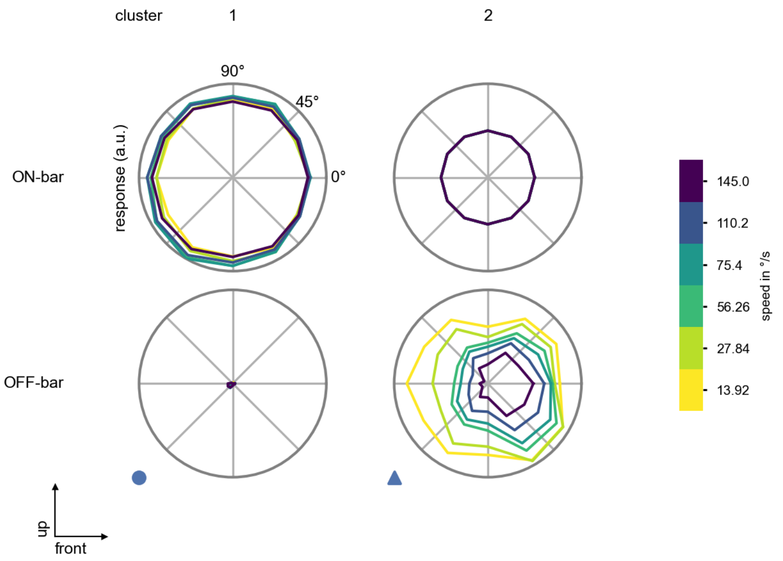

Lawf2 - Figure 9: **Peak responses to moving bars from best-performing models.** The top row shows peak responses to moving ON-bars, the bottom row shows peak responses to moving OFF-bars. The bar-stimuli move in different directions from 0 to 360 degrees and at different speeds. Before, a grey-stimulus led the network to a stationary state. The columns show responses from the best-task-performing model in the respective cluster.

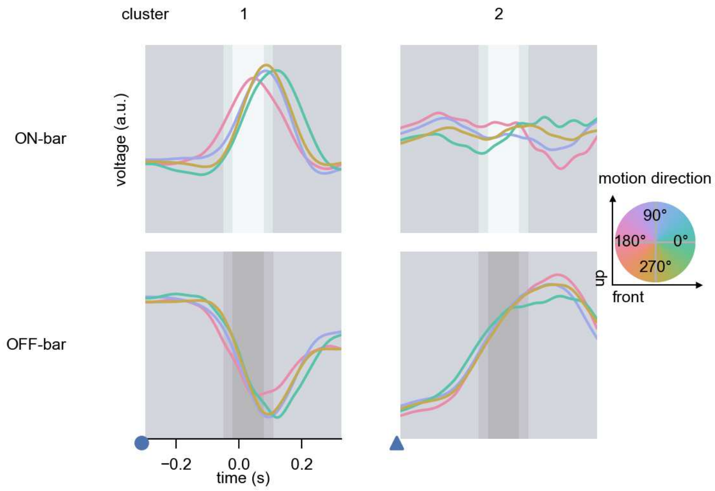

Lawf2 - Figure 10: **Responses to moving bars from best-performing models.** The top row shows responses to moving ON-bars, the bottom row shows peak responses to moving OFF-bars. The bar-stimuli move in different directions from 0 to 360 degrees and at different speeds. Before, a grey-stimulus led the network to a stationary state. The columns show responses from the best-task-performing model in the respective cluster. Example responses to bars moving at 75.4°/s in all cardinal directions (green 0°, blue 90°, red 180°, yellow 270°) from -22.5 to 22.5° visual angle. Background contour indicates the bar, passing by the central column at second zero.

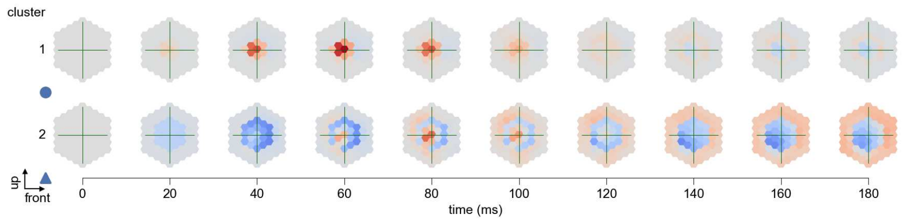

Lawf2 - Figure 11: **Spatio-temporal receptive field.** Responses of the central cell to ON-impulses at single ommatidium impulse locations. The impulse occurs at second zero. Before, a grey-stimulus led the network to a stationary state. Responses are baseline-subtracted. Each row illustrates responses from the best-performing model of the respective cluster. Red indicated depolarization, blue indicates hyperpolarization.

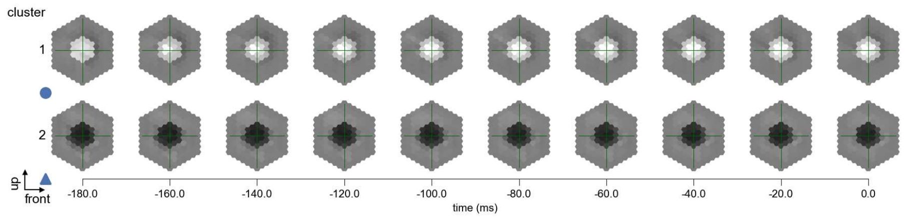

Lawf2 - Figure 12: **Maximally excitatory stimuli.** Each row presents the regularized naturalistic-stimulus from the Sintel dataset that maximizes the cell type's central column response at second zero in the best-performing model of the respective cluster.

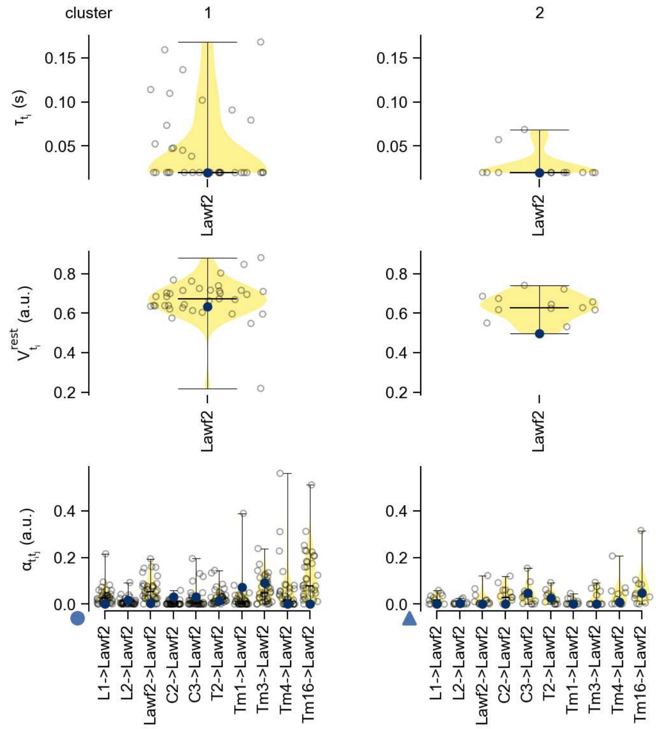

Lawf2 - Figure 13: **Task-constrained parameters.** Each column shows the parameters inferred within the respective cluster. First row: learned time constants of the cell type. Second row: resting potentials of the cell type. Third row: scaling factors for the convolutional filters. The blue scatter represents the parameters from the best-performing model within the cluster.

# 16 Am

← Cell types

#### Figures

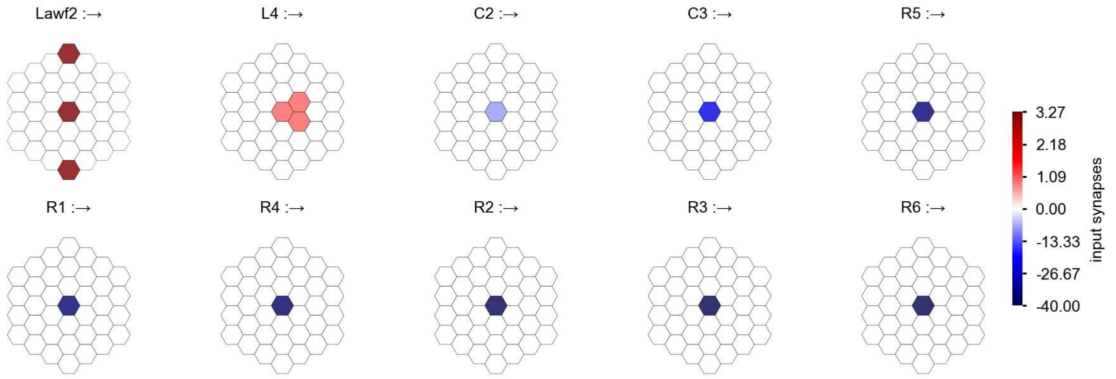

Am - Figure 1: **Anatomical receptive fields.** Each colored hexagon is a projection, with the projection strength characterized by the average number of synapses that we count from the EM reconstruction. Red indicates excitatory synapses, blue indicates inhibitory synapses from inferred signs. Filters in the order of their total number of synapses.

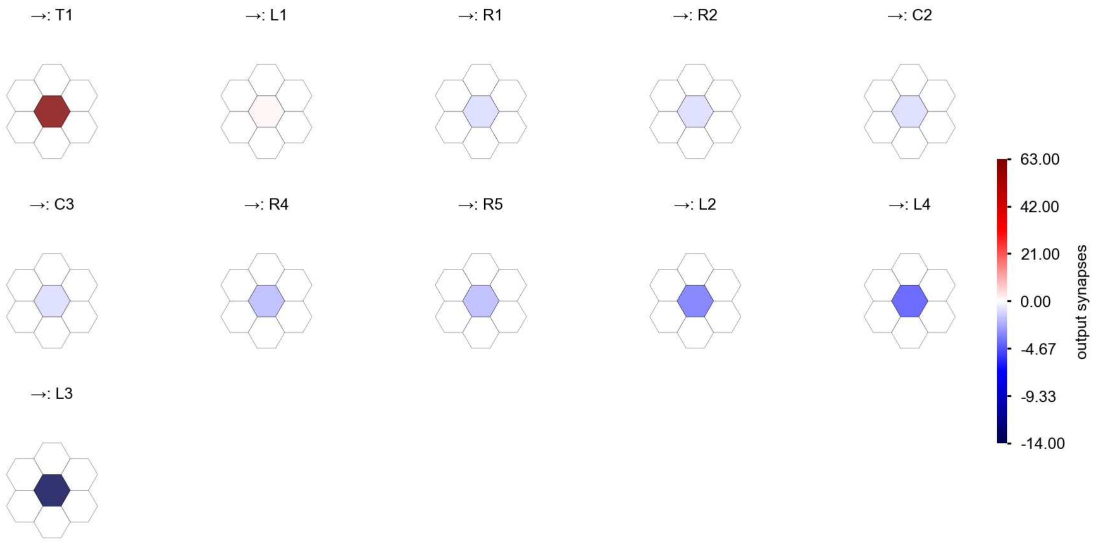

Am - Figure 2: **Anatomical projective fields.** Each colored hexagon is a projection, with the projection strength characterized by the average number of synapses that we count from the EM reconstruction. Red indicates excitatory synapses, blue indicates inhibitory synapses from inferred signs. Filters in the order of their total number of synapses.

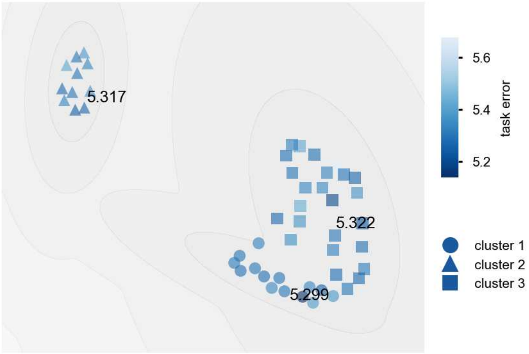

Am - Figure 3: **Clustering of the responses to naturalistic stimuli.** From responses to naturalistic scenes from Sintel. Scatterpoints represent individual models colored by their task error.

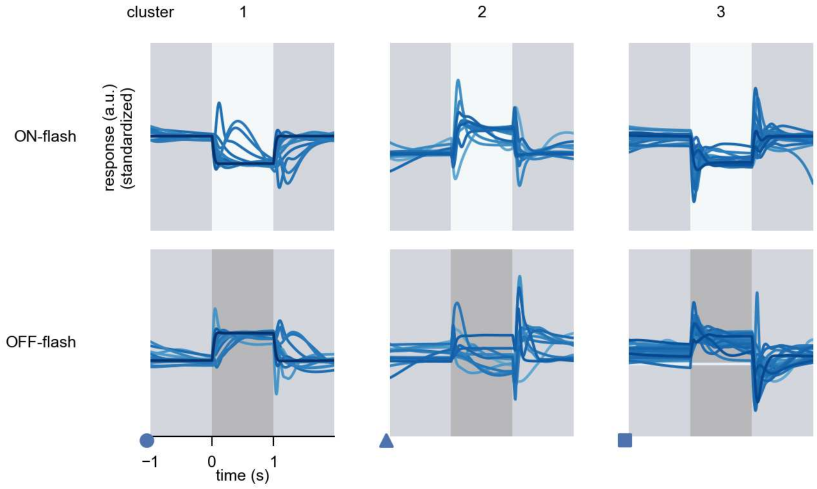

Am - Figure 4: **Responses to flashes.** The top row shows responses to ON-flashes, the bottom row shows responses to OFF-flashes. The responses from the different models in the different clusters (columns) overlay and are colored in the respective task error of the model. The circular flashes occur at second zero, cover 6 ommatidia in radius, and last for one second. Before and after, a grey-stimulus leads to a stationary state of the network. The traces are standardized.

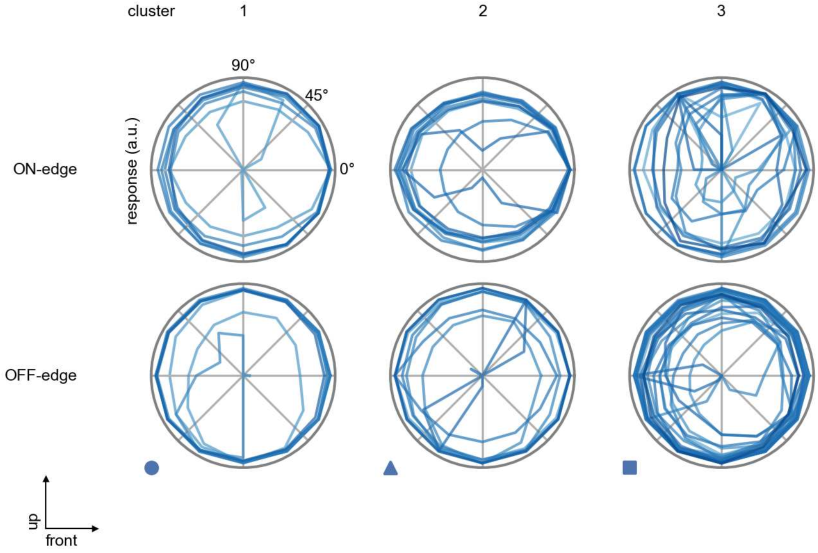

Am - Figure 5: **Peak responses to moving edges.** The top row shows normalized peak responses to moving ON-edges, the bottom row shows normalized peak responses to moving OFF-edges. The edge-stimuli move in different directions from 0 to 360 degrees. Before, a grey-stimulus led the network to a stationary state. The responses from the different models in the different clusters (columns) overlay and are colored in the respective task error of the model. The peak responses are averaged over all assessed edge-speeds.

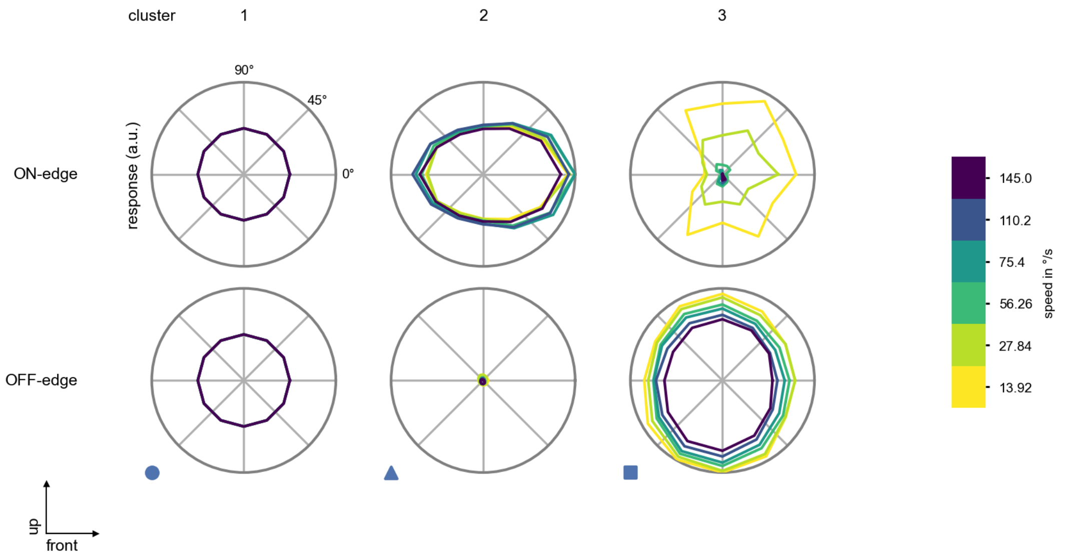

Am - Figure 6: **Peak responses to moving edges from best-performing models.** The top row shows peak responses to moving ON-edges, the bottom row shows peak responses to moving OFF-edges. The edge-stimuli move in different directions from 0 to 360 degrees and at different speeds. Before, a grey-stimulus led the network to a stationary state. The columns show responses from the best-task-performing model in the respective cluster.

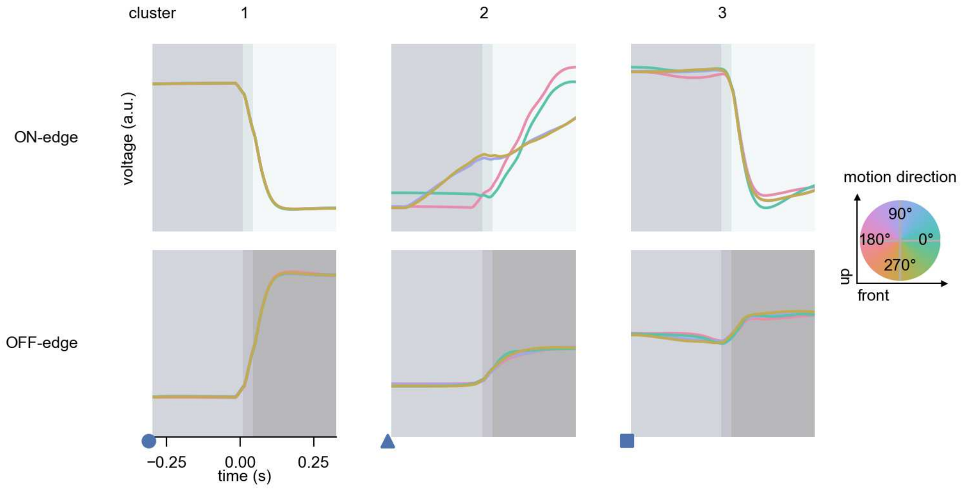

Am - Figure 7: **Responses to moving edges from best-performing models.** The top row shows responses to moving ON-edges, the bottom row shows peak responses to moving OFF-edges. The edge-stimuli move in different directions from 0 to 360 degrees and at different speeds. Before, a grey-stimulus led the network to a stationary state. The columns show responses from the best-task-performing model in the respective cluster. Example responses to edges moving at 75.4°/s in all cardinal directions (green 0°, blue 90°, red 180°, yellow 270°) from -22.5 to 22.5° visual angle. Background contour indicates the edge, passing by the central column at second zero.

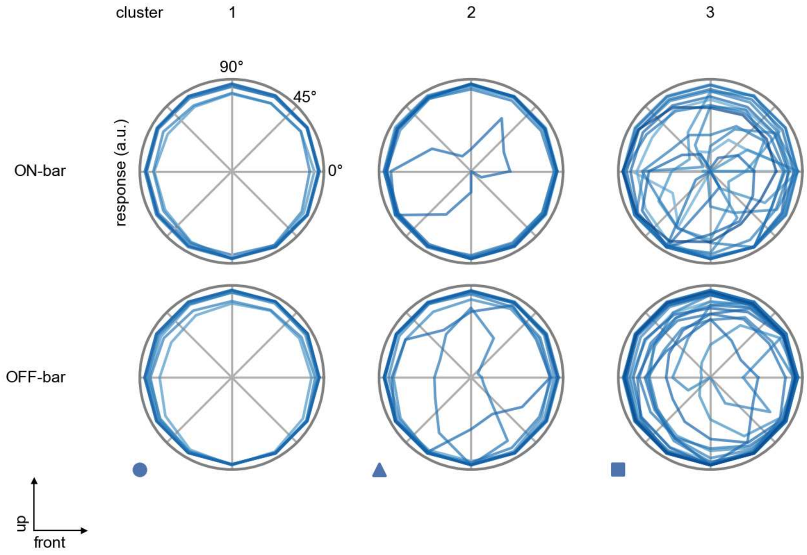

Am - Figure 8: **Peak responses to moving bars.** The top row shows normalized peak responses to moving ON-bars, the bottom row shows normalized peak responses to moving OFF-bars. The bar-stimuli move in different directions from 0 to 360 degrees. Before, a grey-stimulus led the network to a stationary state. The responses from the different models in the different clusters (columns) overlay and are colored in the respective task error of the model. The peak responses are averaged over all assessed bar-speeds.

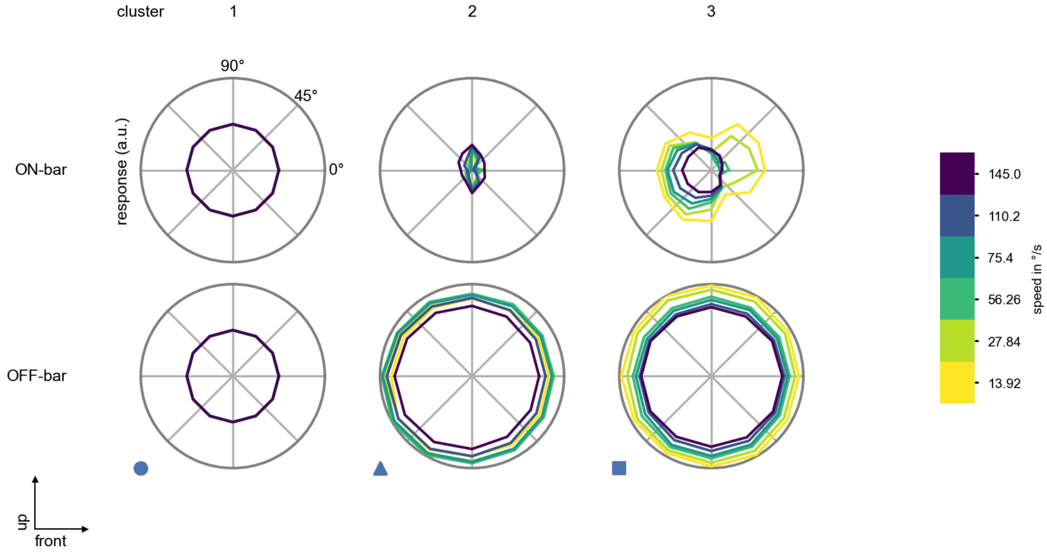

Am - Figure 9: **Peak responses to moving bars from best-performing models.** The top row shows peak responses to moving ON-bars, the bottom row shows peak responses to moving OFF-bars. The bar-stimuli move in different directions from 0 to 360 degrees and at different speeds. Before, a grey-stimulus led the network to a stationary state. The columns show responses from the best-task-performing model in the respective cluster.

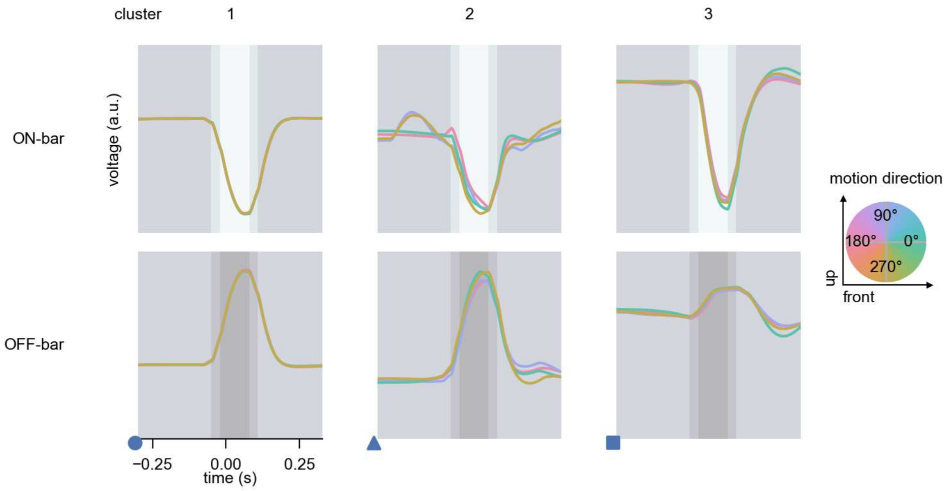

Am - Figure 10: **Responses to moving bars from best-performing models.** The top row shows responses to moving ON-bars, the bottom row shows peak responses to moving OFF-bars. The bar-stimuli move in different directions from 0 to 360 degrees and at different speeds. Before, a grey-stimulus led the network to a stationary state. The columns show responses from the best-task-performing model in the respective cluster. Example responses to bars moving at 75.4°/s in all cardinal directions (green 0°, blue 90°, red 180°, yellow 270°) from -22.5 to 22.5° visual angle. Background contour indicates the bar, passing by the central column at second zero.

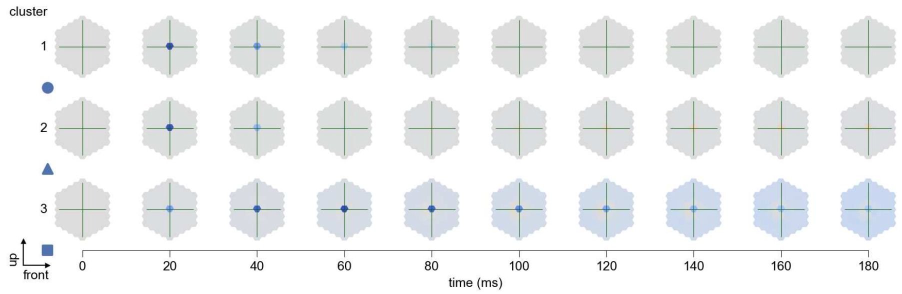

Am - Figure 11: **Spatio-temporal receptive field.** Responses of the central cell to ON-impulses at single ommatidium impulse locations. The impulse occurs at second zero. Before, a grey-stimulus led the network to a stationary state. Responses are baseline-subtracted. Each row illustrates responses from the best-performing model of the respective cluster. Red indicated depolarization, blue indicates hyperpolarization.

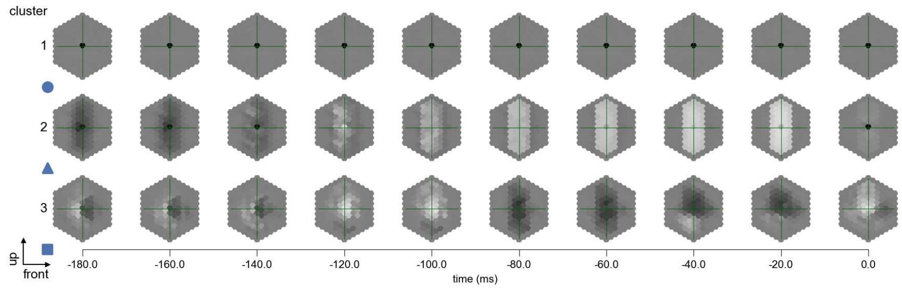

Am - Figure 12: **Maximally excitatory stimuli.** Each row presents the regularized naturalistic-stimulus from the Sintel dataset that maximizes the cell type's central column response at second zero in the best-performing model of the respective cluster.

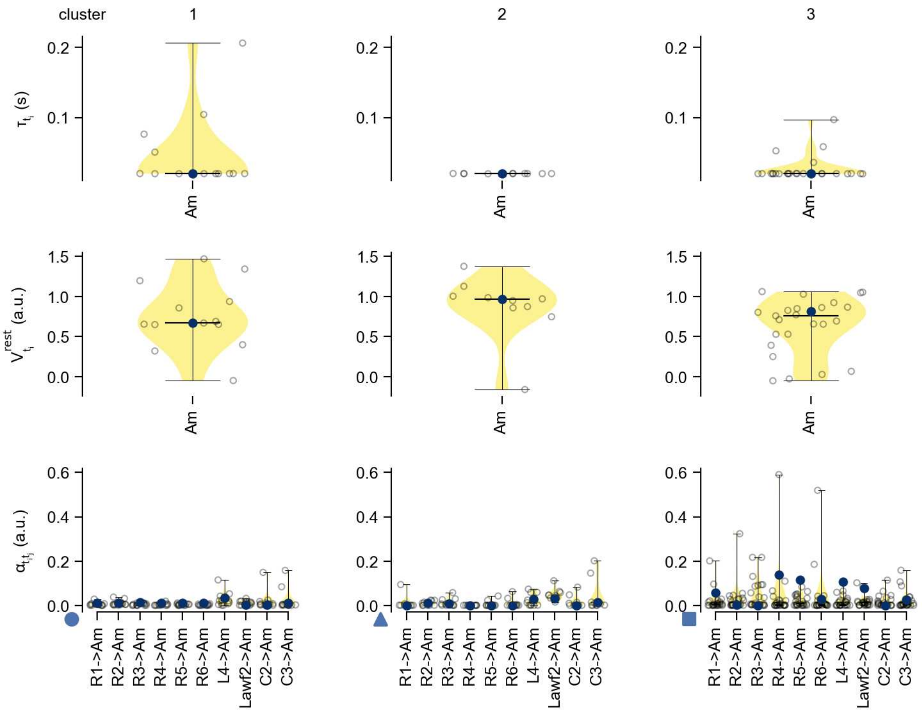

Am - Figure 13: **Task-constrained parameters.** Each column shows the parameters inferred within the respective cluster. First row: learned time constants of the cell type. Second row: resting potentials of the cell type. Third row: scaling factors for the convolutional filters. The blue scatter represents the parameters from the best-performing model within the cluster.

# 17 C2

← Cell types

#### Figures

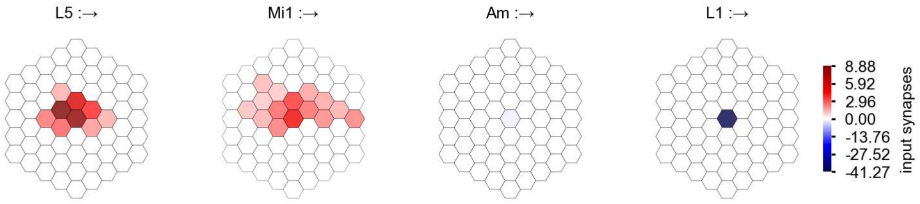

C2 - Figure 1: **Anatomical receptive fields.** Each colored hexagon is a projection, with the projection strength characterized by the average number of synapses that we count from the EM reconstruction. Red indicates excitatory synapses, blue indicates inhibitory synapses from inferred signs. Filters in the order of their total number of synapses.

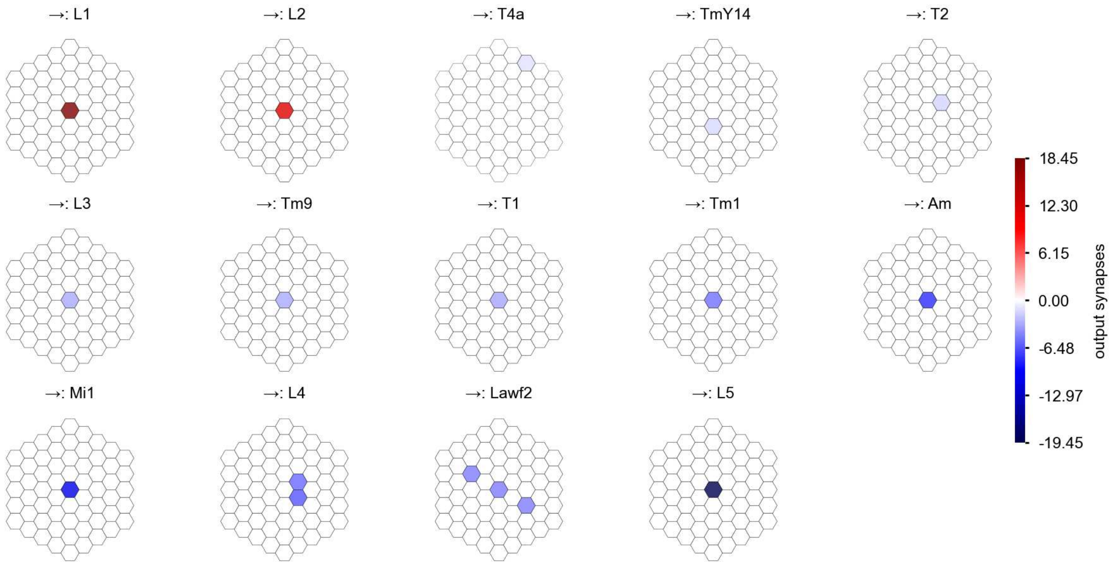

C2 - Figure 2: **Anatomical projective fields.** Each colored hexagon is a projection, with the projection strength characterized by the average number of synapses that we count from the EM reconstruction. Red indicates excitatory synapses, blue indicates inhibitory synapses from inferred signs. Filters in the order of their total number of synapses.

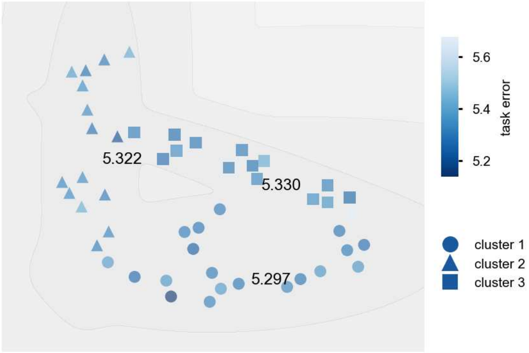

C2 - Figure 3: **Clustering of the responses to naturalistic stimuli.** From responses to naturalistic scenes from Sintel. Scatterpoints represent individual models colored by their task error.

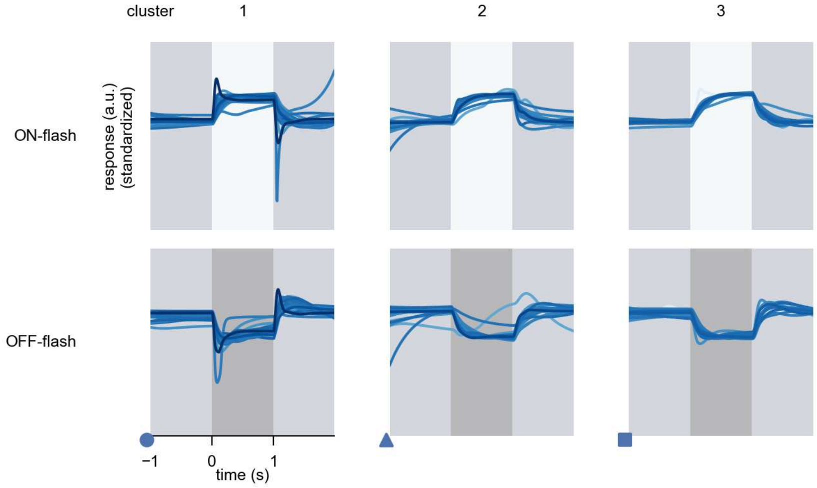

C2 - Figure 4: **Responses to flashes.** The top row shows responses to ON-flashes, the bottom row shows responses to OFF-flashes. The responses from the different models in the different clusters (columns) overlay and are colored in the respective task error of the model. The circular flashes occur at second zero, cover 6 ommatidia in radius, and last for one second. Before and after, a grey-stimulus leads to a stationary state of the network. The traces are standardized.

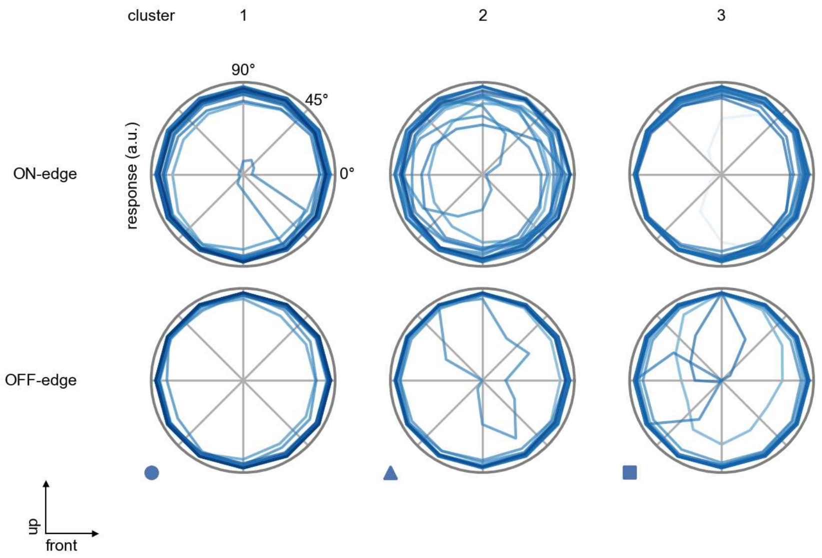

C2 - Figure 5: **Peak responses to moving edges.** The top row shows normalized peak responses to moving ON-edges, the bottom row shows normalized peak responses to moving OFF-edges. The edge-stimuli move in different directions from 0 to 360 degrees. Before, a grey-stimulus led the network to a stationary state. The responses from the different models in the different clusters (columns) overlay and are colored in the respective task error of the model. The peak responses are averaged over all assessed edge-speeds.

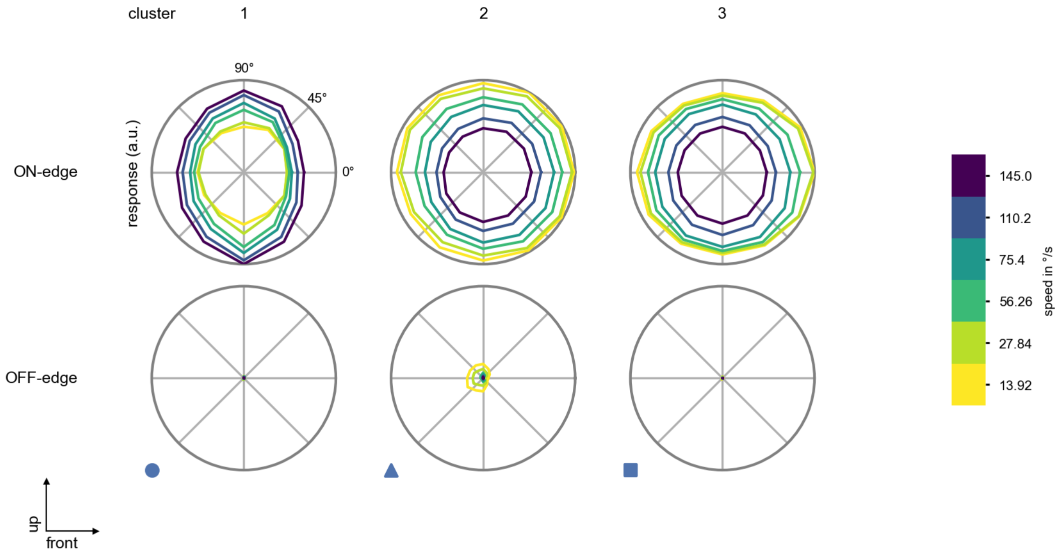

C2 - Figure 6: **Peak responses to moving edges from best-performing models.** The top row shows peak responses to moving ON-edges, the bottom row shows peak responses to moving OFF-edges. The edge-stimuli move in different directions from 0 to 360 degrees and at different speeds. Before, a grey-stimulus led the network to a stationary state. The columns show responses from the best-task-performing model in the respective cluster.

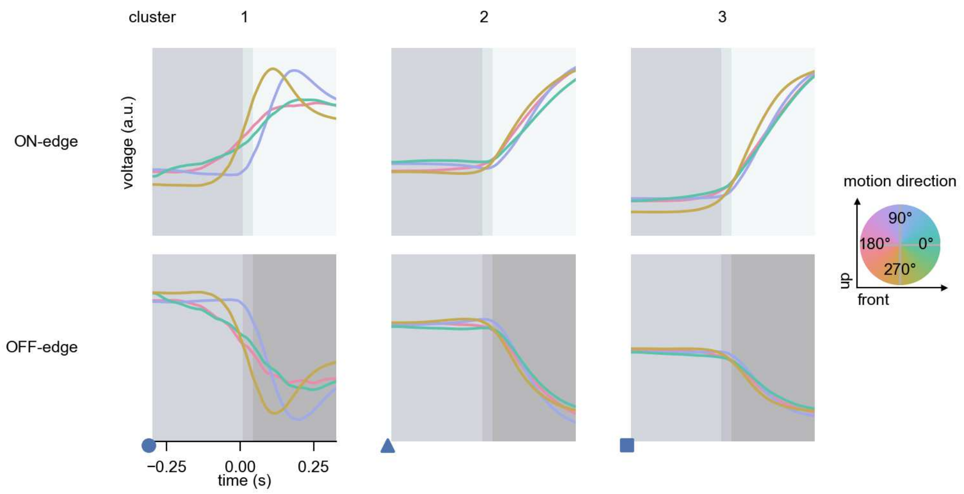

C2 - Figure 7: **Responses to moving edges from best-performing models.** The top row shows responses to moving ON-edges, the bottom row shows peak responses to moving OFF-edges. The edge-stimuli move in different directions from 0 to 360 degrees and at different speeds. Before, a grey-stimulus led the network to a stationary state. The columns show responses from the best-task-performing model in the respective cluster. Example responses to edges moving at 75.4°/s in all cardinal directions (green 0°, blue 90°, red 180°, yellow 270°) from -22.5 to 22.5° visual angle. Background contour indicates the edge, passing by the central column at second zero.

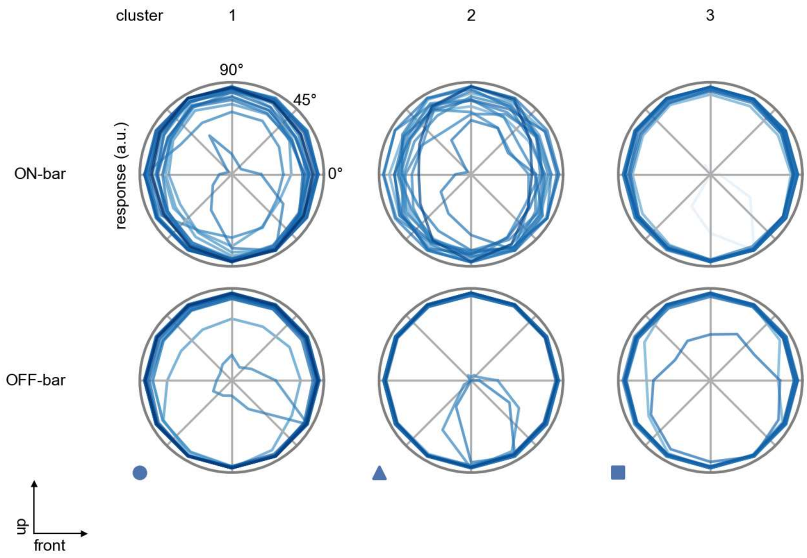

C2 - Figure 8: **Peak responses to moving bars.** The top row shows normalized peak responses to moving ON-bars, the bottom row shows normalized peak responses to moving OFF-bars. The bar-stimuli move in different directions from 0 to 360 degrees. Before, a grey-stimulus led the network to a stationary state. The responses from the different models in the different clusters (columns) overlay and are colored in the respective task error of the model. The peak responses are averaged over all assessed bar-speeds.

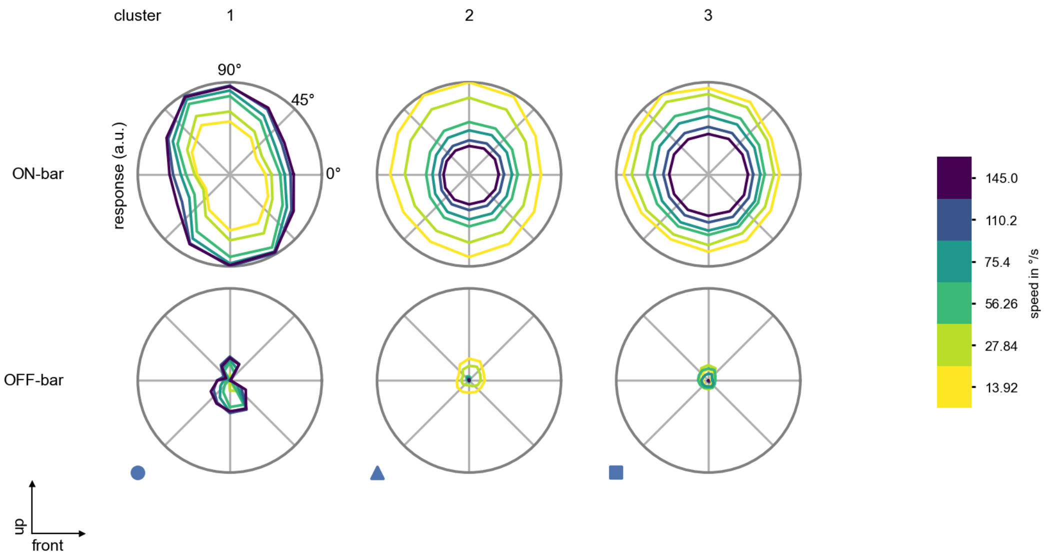

C2 - Figure 9: **Peak responses to moving bars from best-performing models.** The top row shows peak responses to moving ON-bars, the bottom row shows peak responses to moving OFF-bars. The bar-stimuli move in different directions from 0 to 360 degrees and at different speeds. Before, a grey-stimulus led the network to a stationary state. The columns show responses from the best-task-performing model in the respective cluster.

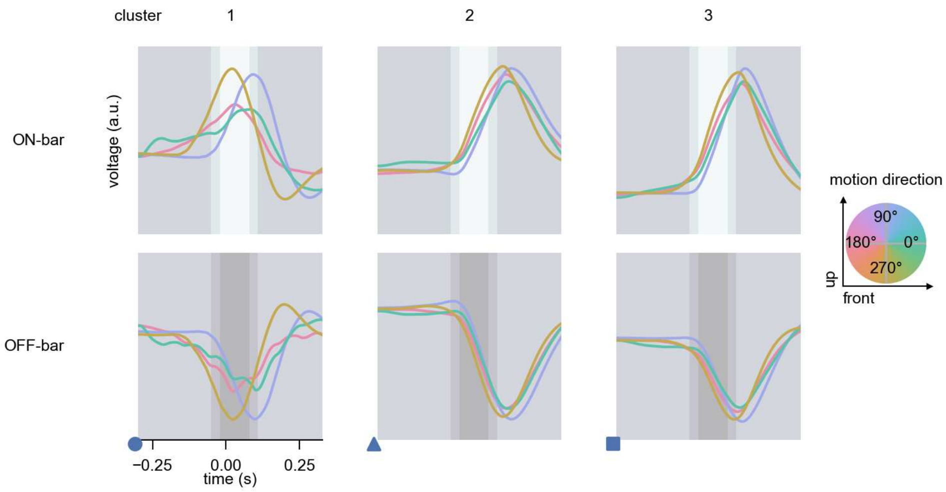

C2 - Figure 10: **Responses to moving bars from best-performing models.** The top row shows responses to moving ON-bars, the bottom row shows peak responses to moving OFF-bars. The bar-stimuli move in different directions from 0 to 360 degrees and at different speeds. Before, a grey-stimulus led the network to a stationary state. The columns show responses from the best-task-performing model in the respective cluster. Example responses to bars moving at 75.4°/s in all cardinal directions (green 0°, blue 90°, red 180°, yellow 270°) from -22.5 to 22.5° visual angle. Background contour indicates the bar, passing by the central column at second zero.

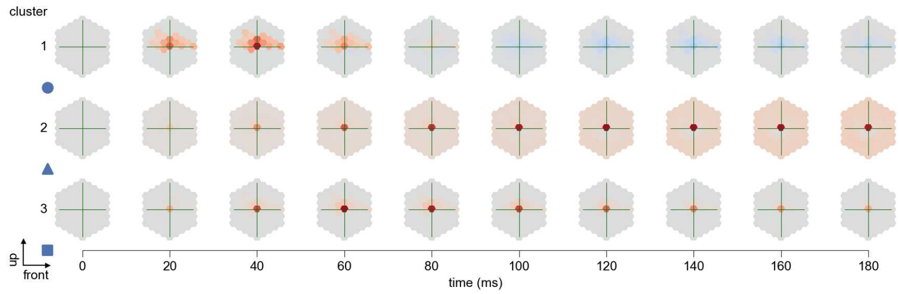

C2 - Figure 11: **Spatio-temporal receptive field.** Responses of the central cell to ON-impulses at single ommatidium impulse locations. The impulse occurs at second zero. Before, a grey-stimulus led the network to a stationary state. Responses are baseline-subtracted. Each row illustrates responses from the best-performing model of the respective cluster. Red indicated depolarization, blue indicates hyperpolarization.

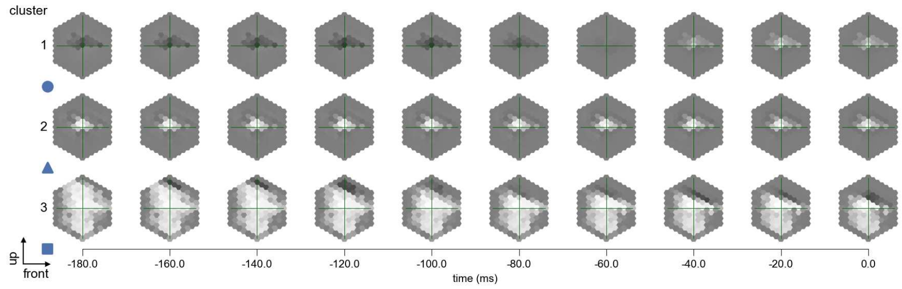

C2 - Figure 12: **Maximally excitatory stimuli.** Each row presents the regularized naturalistic-stimulus from the Sintel dataset that maximizes the cell type's central column response at second zero in the best-performing model of the respective cluster.

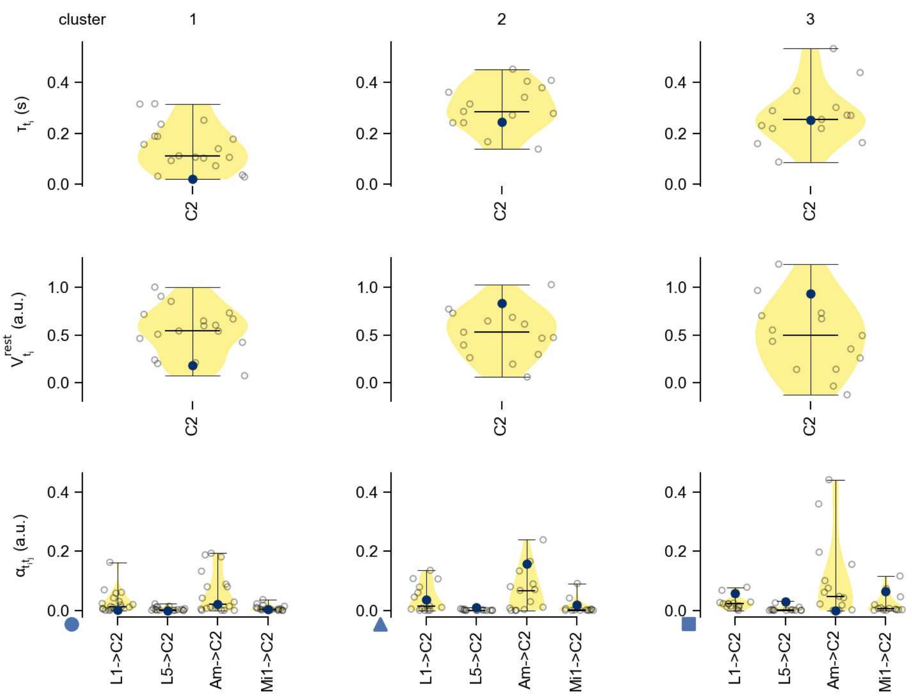

C2 - Figure 13: **Task-constrained parameters.** Each column shows the parameters inferred within the respective cluster. First row: learned time constants of the cell type. Second row: resting potentials of the cell type. Third row: scaling factors for the convolutional filters. The blue scatter represents the parameters from the best-performing model within the cluster.

# 18 C3

← Cell types

#### Figures

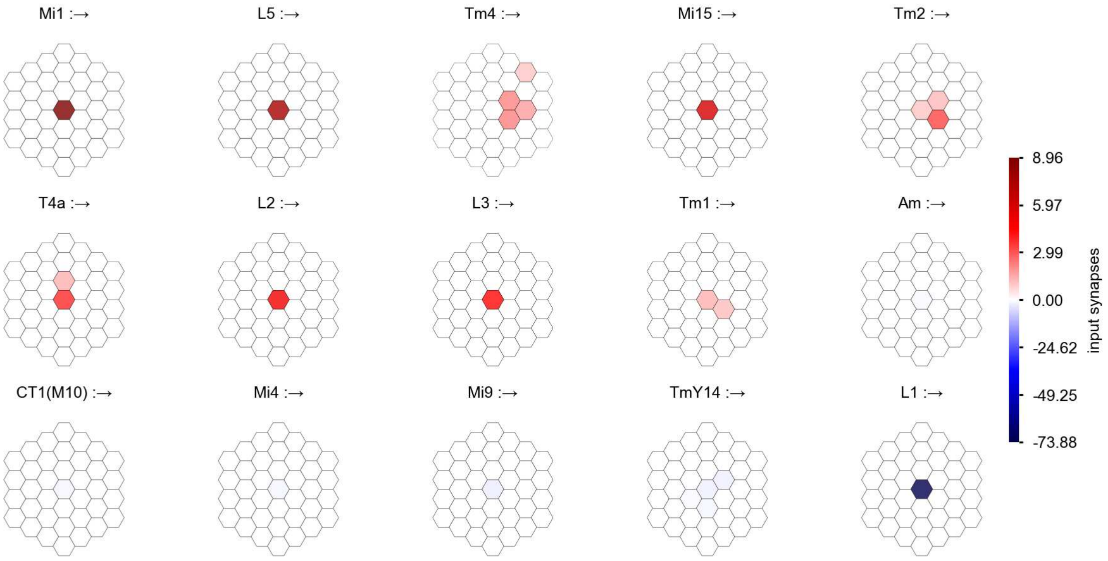

C3 - Figure 1: **Anatomical receptive fields.** Each colored hexagon is a projection, with the projection strength characterized by the average number of synapses that we count from the EM reconstruction. Red indicates excitatory synapses, blue indicates inhibitory synapses from inferred signs. Filters in the order of their total number of synapses.

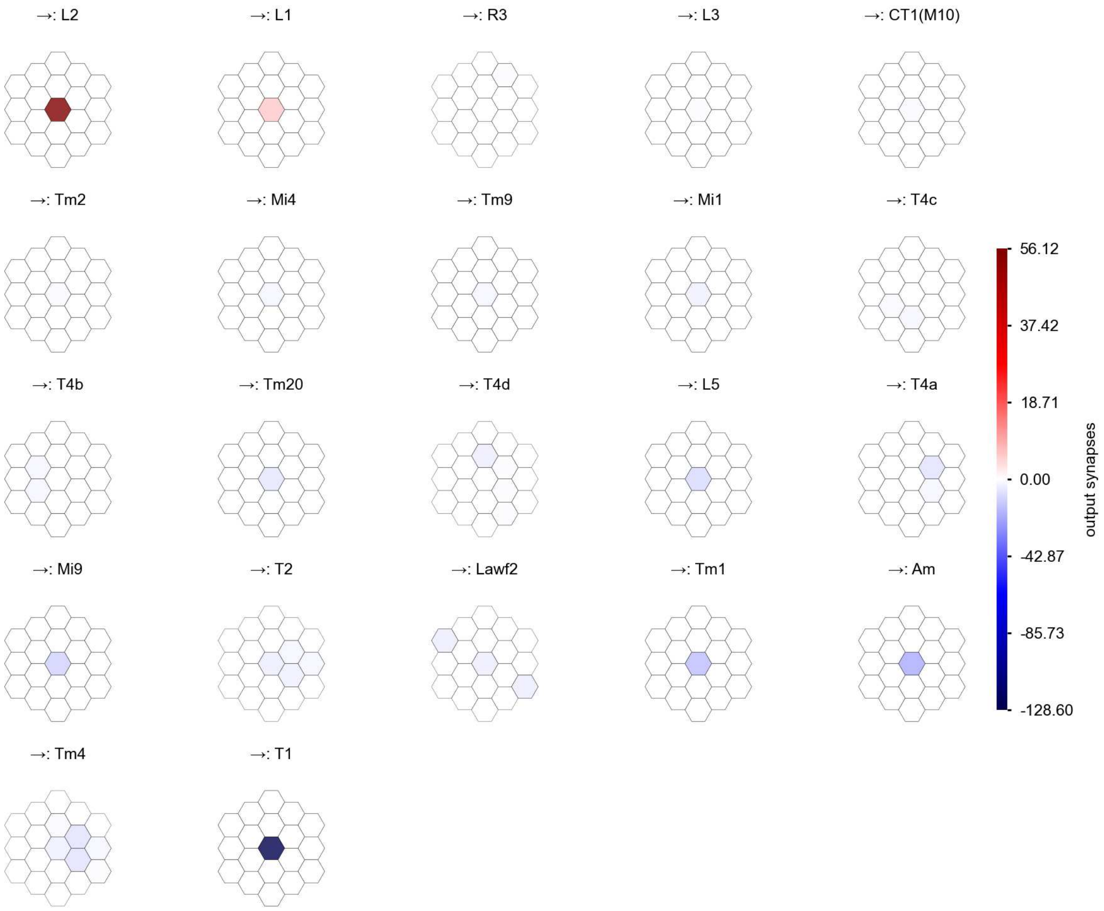

C3 - Figure 2: **Anatomical projective fields.** Each colored hexagon is a projection, with the projection strength characterized by the average number of synapses that we count from the EM reconstruction. Red indicates excitatory synapses, blue indicates inhibitory synapses from inferred signs. Filters in the order of their total number of synapses.

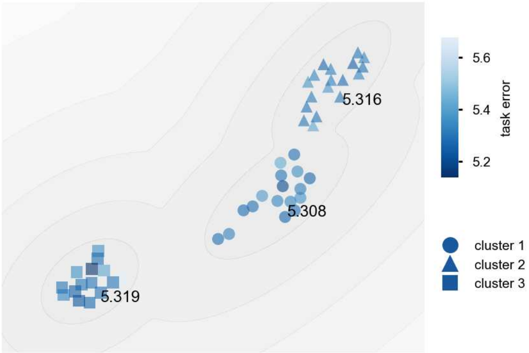

C3 - Figure 3: **Clustering of the responses to naturalistic stimuli.** From responses to naturalistic scenes from Sintel. Scatterpoints represent individual models colored by their task error.

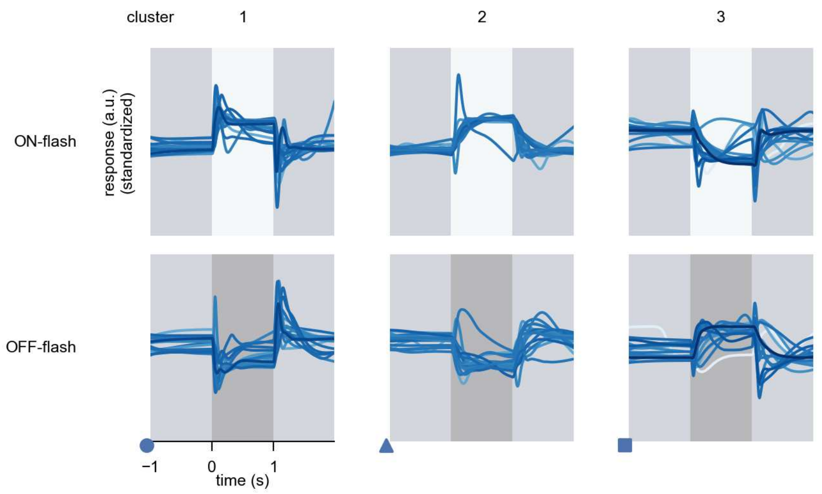

C3 - Figure 4: **Responses to flashes.** The top row shows responses to ON-flashes, the bottom row shows responses to OFF-flashes. The responses from the different models in the different clusters (columns) overlay and are colored in the respective task error of the model. The circular flashes occur at second zero, cover 6 ommatidia in radius, and last for one second. Before and after, a grey-stimulus leads to a stationary state of the network. The traces are standardized.

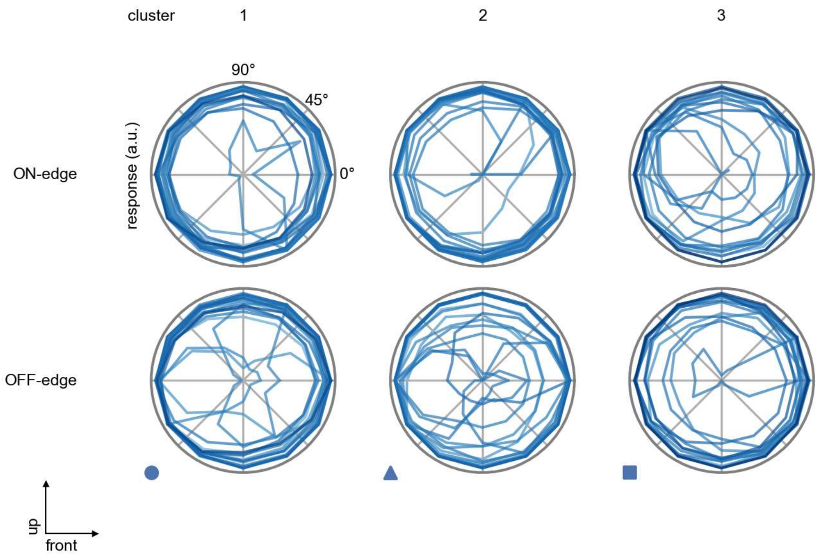

C3 - Figure 5: **Peak responses to moving edges.** The top row shows normalized peak responses to moving ON-edges, the bottom row shows normalized peak responses to moving OFF-edges. The edge-stimuli move in different directions from 0 to 360 degrees. Before, a grey-stimulus led the network to a stationary state. The responses from the different models in the different clusters (columns) overlay and are colored in the respective task error of the model. The peak responses are averaged over all assessed edge-speeds.

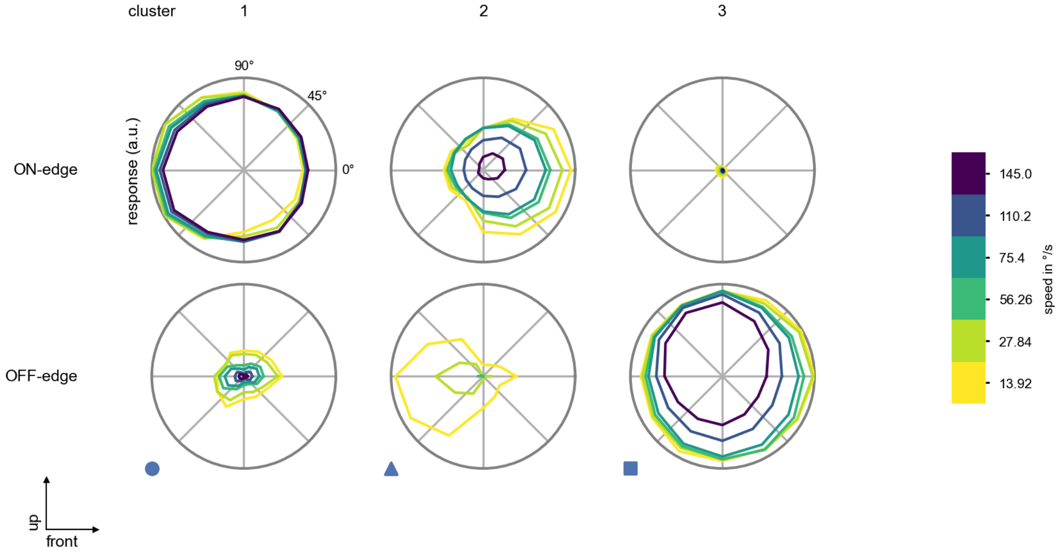

C3 - Figure 6: **Peak responses to moving edges from best-performing models.** The top row shows peak responses to moving ON-edges, the bottom row shows peak responses to moving OFF-edges. The edge-stimuli move in different directions from 0 to 360 degrees and at different speeds. Before, a grey-stimulus led the network to a stationary state. The columns show responses from the best-task-performing model in the respective cluster.

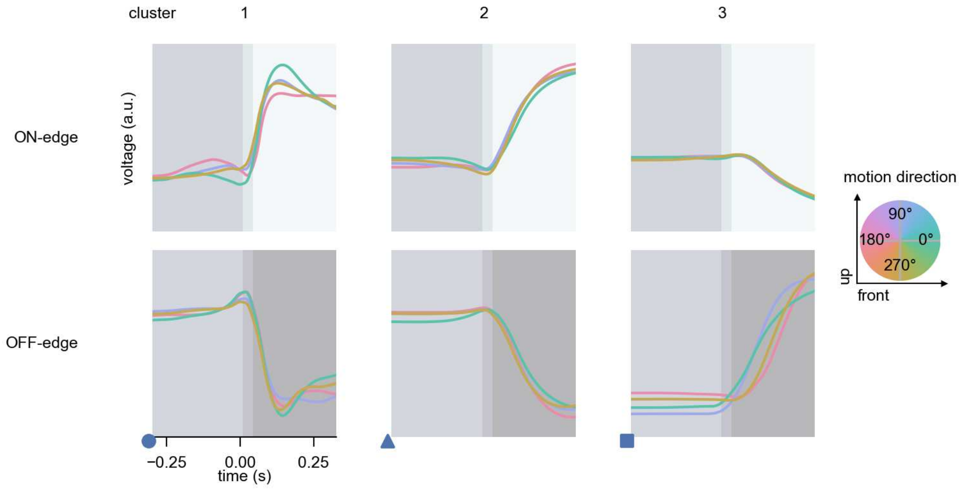

C3 - Figure 7: **Responses to moving edges from best-performing models.** The top row shows responses to moving ON-edges, the bottom row shows peak responses to moving OFF-edges. The edge-stimuli move in different directions from 0 to 360 degrees and at different speeds. Before, a grey-stimulus led the network to a stationary state. The columns show responses from the best-task-performing model in the respective cluster. Example responses to edges moving at 75.4°/s in all cardinal directions (green 0°, blue 90°, red 180°, yellow 270°) from -22.5 to 22.5° visual angle. Background contour indicates the edge, passing by the central column at second zero.

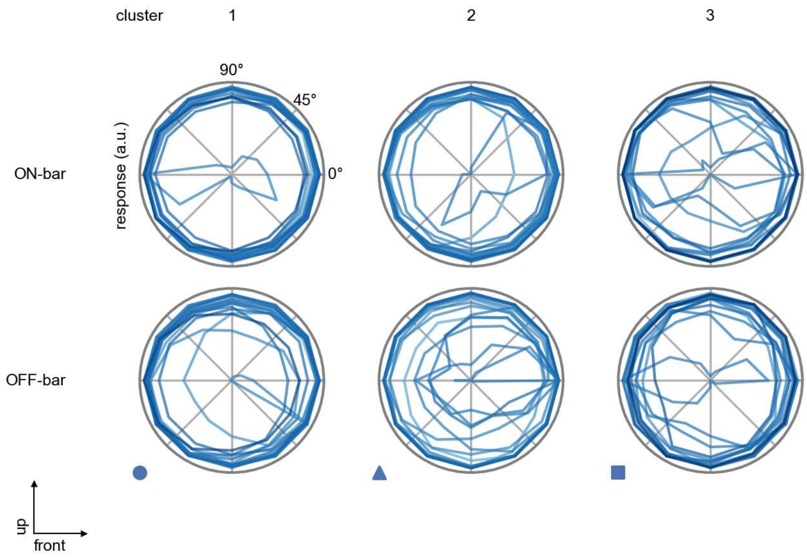

C3 - Figure 8: **Peak responses to moving bars.** The top row shows normalized peak responses to moving ON-bars, the bottom row shows normalized peak responses to moving OFF-bars. The bar-stimuli move in different directions from 0 to 360 degrees. Before, a grey-stimulus led the network to a stationary state. The responses from the different models in the different clusters (columns) overlay and are colored in the respective task error of the model. The peak responses are averaged over all assessed bar-speeds.

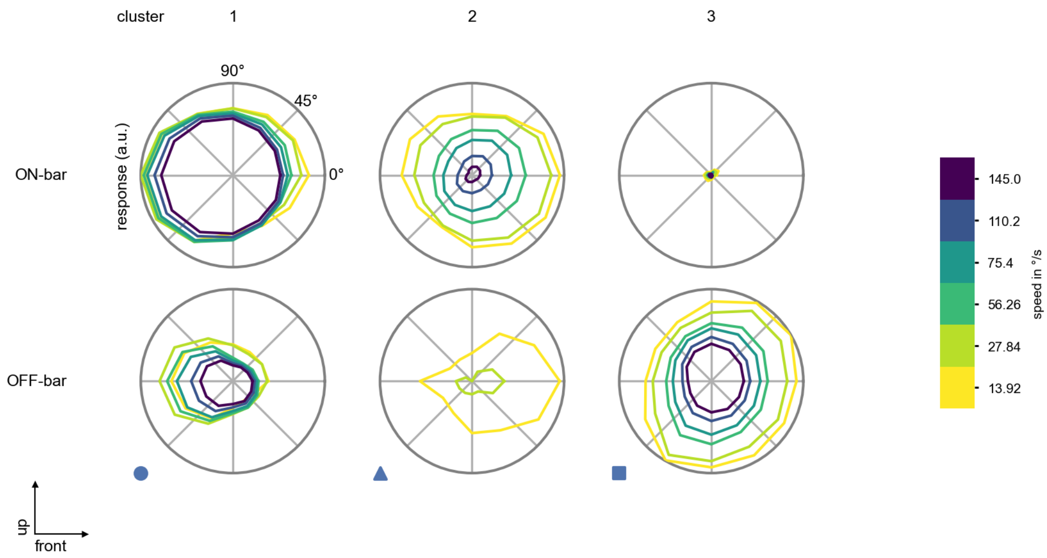

C3 - Figure 9: **Peak responses to moving bars from best-performing models.** The top row shows peak responses to moving ON-bars, the bottom row shows peak responses to moving OFF-bars. The bar-stimuli move in different directions from 0 to 360 degrees and at different speeds. Before, a grey-stimulus led the network to a stationary state. The columns show responses from the best-task-performing model in the respective cluster.

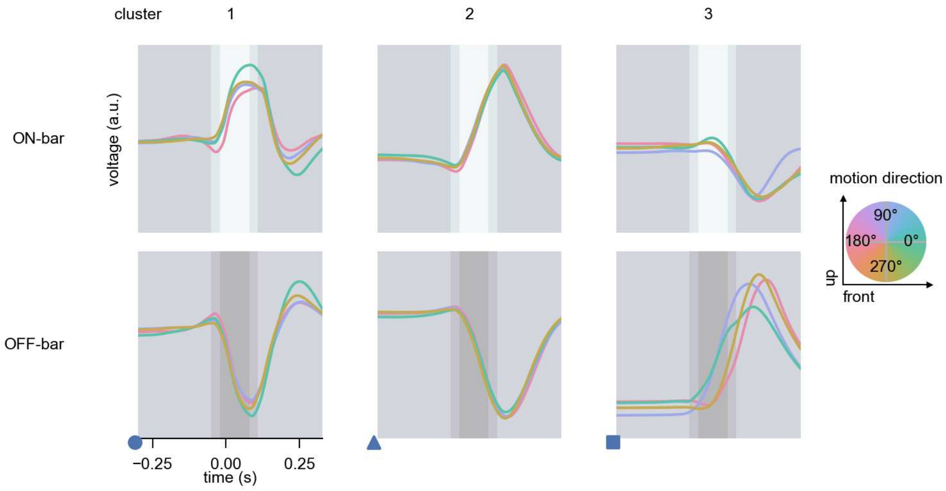

C3 - Figure 10: **Responses to moving bars from best-performing models.** The top row shows responses to moving ON-bars, the bottom row shows peak responses to moving OFF-bars. The bar-stimuli move in different directions from 0 to 360 degrees and at different speeds. Before, a grey-stimulus led the network to a stationary state. The columns show responses from the best-task-performing model in the respective cluster. Example responses to bars moving at 75.4°/s in all cardinal directions (green 0°, blue 90°, red 180°, yellow 270°) from -22.5 to 22.5° visual angle. Background contour indicates the bar, passing by the central column at second zero.

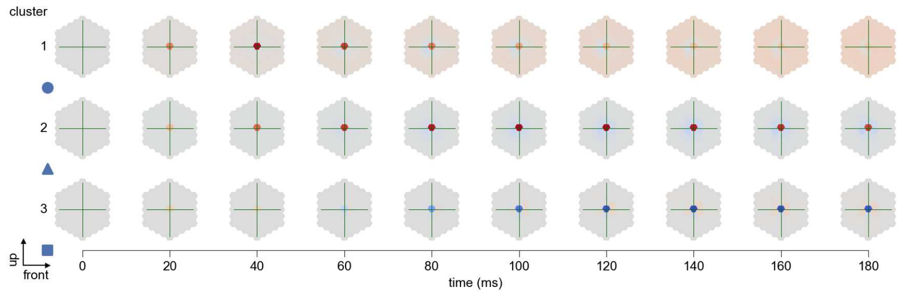

C3 - Figure 11: **Spatio-temporal receptive field.** Responses of the central cell to ON-impulses at single ommatidium impulse locations. The impulse occurs at second zero. Before, a grey-stimulus led the network to a stationary state. Responses are baseline-subtracted. Each row illustrates responses from the best-performing model of the respective cluster. Red indicated depolarization, blue indicates hyperpolarization.

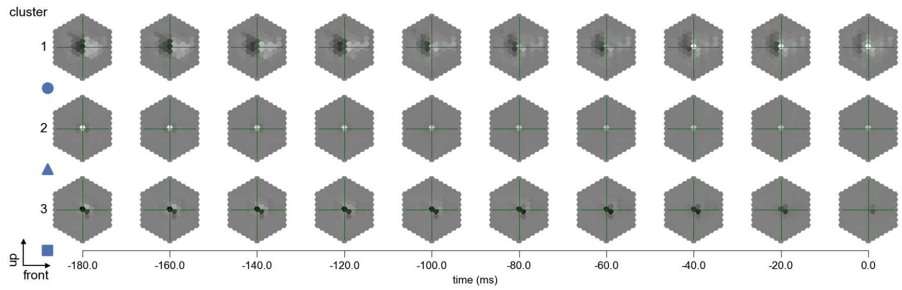

C3 - Figure 12: **Maximally excitatory stimuli.** Each row presents the regularized naturalistic-stimulus from the Sintel dataset that maximizes the cell type's central column response at second zero in the best-performing model of the respective cluster.

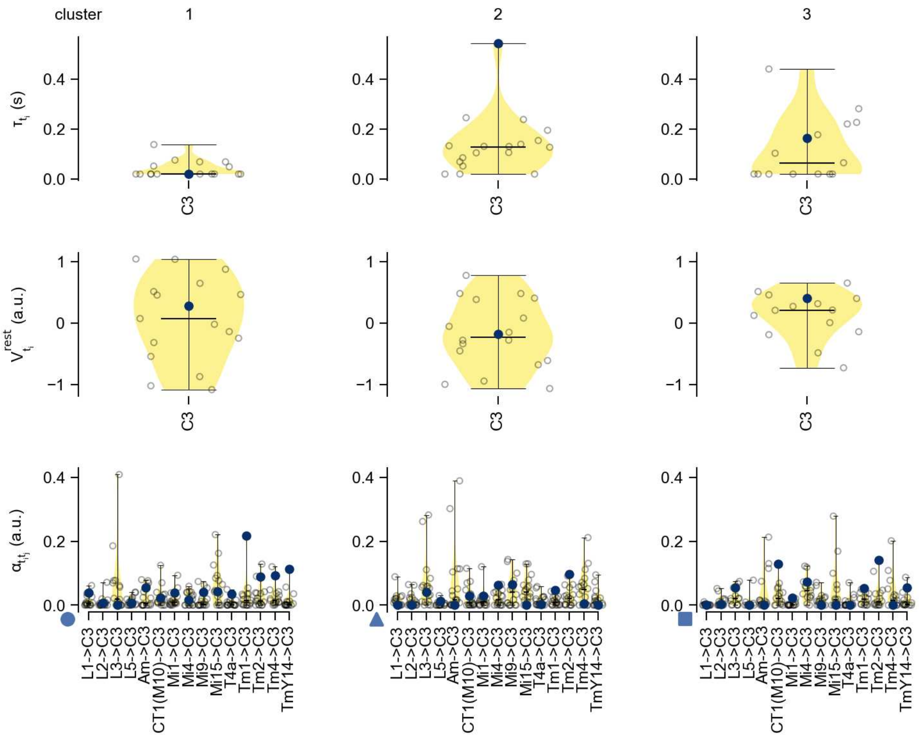

C3 - Figure 13: **Task-constrained parameters.** Each column shows the parameters inferred within the respective cluster. First row: learned time constants of the cell type. Second row: resting potentials of the cell type. Third row: scaling factors for the convolutional filters. The blue scatter represents the parameters from the best-performing model within the cluster.

# 19 CT1(Lo1)

← Cell types

#### Figures

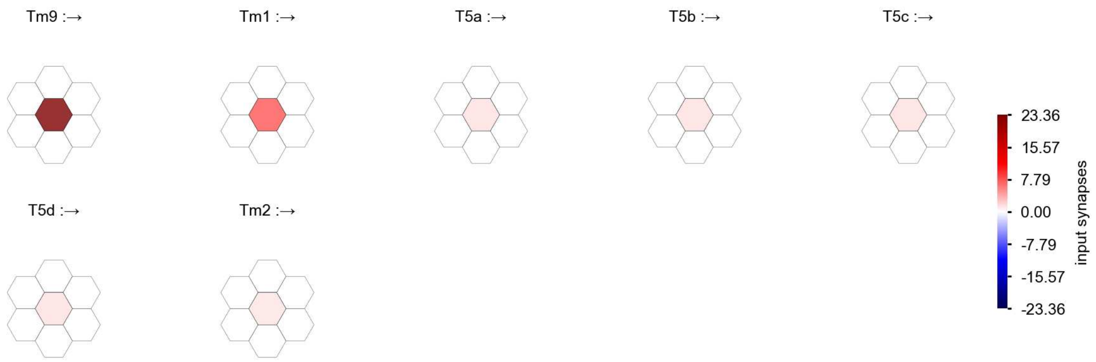

CT1(Lo1) - Figure 1: **Anatomical receptive fields.** Each colored hexagon is a projection, with the projection strength characterized by the average number of synapses that we count from the EM reconstruction. Red indicates excitatory synapses, blue indicates inhibitory synapses from inferred signs. Filters in the order of their total number of synapses.

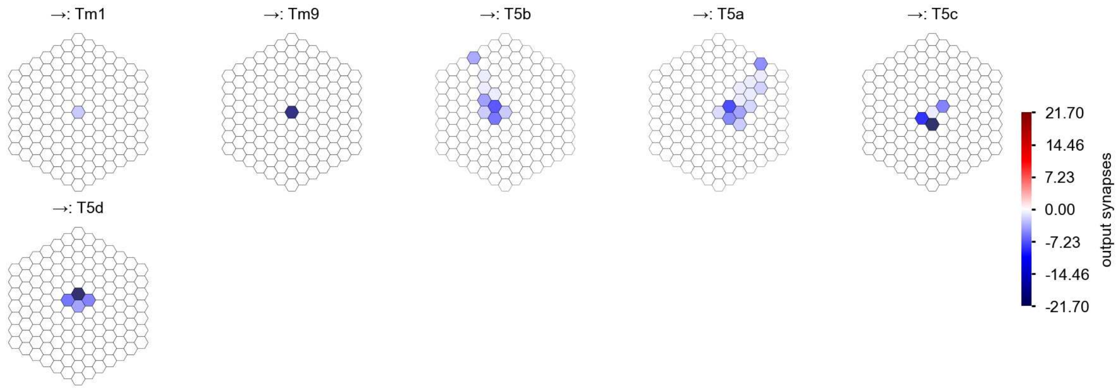

CT1(Lo1) - Figure 2: **Anatomical projective fields.** Each colored hexagon is a projection, with the projection strength characterized by the average number of synapses that we count from the EM reconstruction. Red indicates excitatory synapses, blue indicates inhibitory synapses from inferred signs. Filters in the order of their total number of synapses.

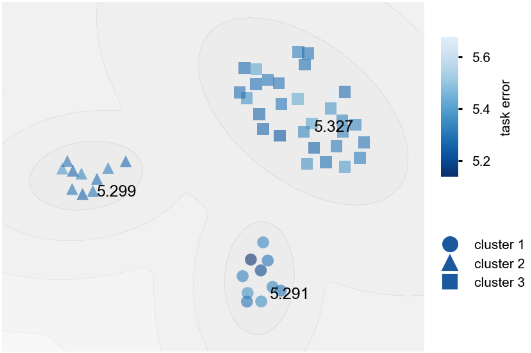

CT1(Lo1) - Figure 3: **Clustering of the responses to naturalistic stimuli.** From responses to naturalistic scenes from Sintel. Scatterpoints represent individual models colored by their task error.

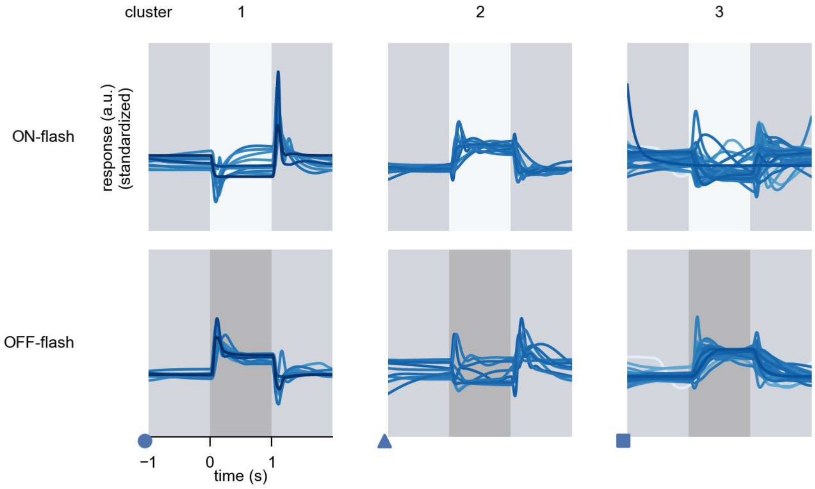

CT1(Lo1) - Figure 4: **Responses to flashes.** The top row shows responses to ON-flashes, the bottom row shows responses to OFF-flashes. The responses from the different models in the different clusters (columns) overlay and are colored in the respective task error of the model. The circular flashes occur at second zero, cover 6 ommatidia in radius, and last for one second. Before and after, a grey-stimulus leads to a stationary state of the network. The traces are standardized.

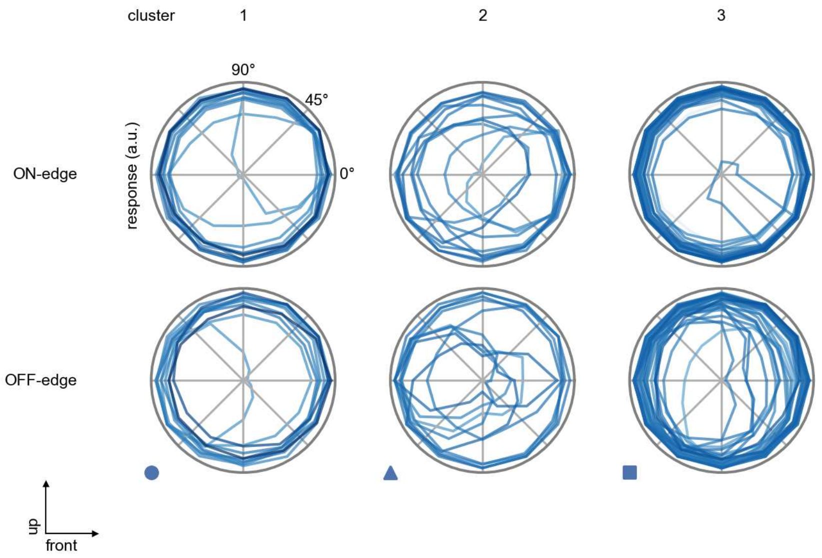

CT1(Lo1) - Figure 5: **Peak responses to moving edges.** The top row shows normalized peak responses to moving ON-edges, the bottom row shows normalized peak responses to moving OFF-edges. The edge-stimuli move in different directions from 0 to 360 degrees. Before, a grey-stimulus led the network to a stationary state. The responses from the different models in the different clusters (columns) overlay and are colored in the respective task error of the model. The peak responses are averaged over all assessed edge-speeds.

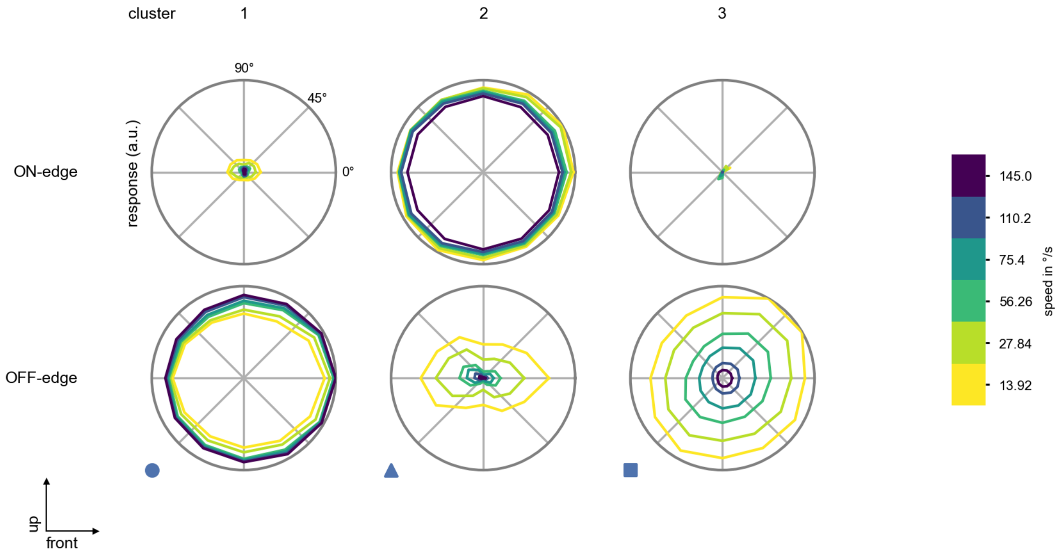

CT1(Lo1) - Figure 6: **Peak responses to moving edges from best-performing models.** The top row shows peak responses to moving ON-edges, the bottom row shows peak responses to moving OFF-edges. The edge-stimuli move in different directions from 0 to 360 degrees and at different speeds. Before, a grey-stimulus led the network to a stationary state. The columns show responses from the best-task-performing model in the respective cluster.

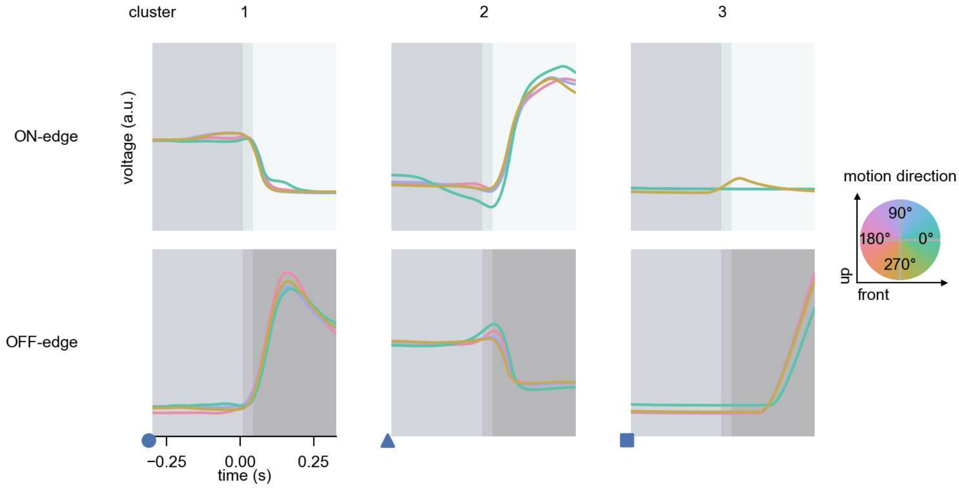

CT1(Lo1) - Figure 7: **Responses to moving edges from best-performing models.** The top row shows responses to moving ON-edges, the bottom row shows peak responses to moving OFF-edges. The edge-stimuli move in different directions from 0 to 360 degrees and at different speeds. Before, a grey-stimulus led the network to a stationary state. The columns show responses from the best-task-performing model in the respective cluster. Example responses to edges moving at 75.4°/s in all cardinal directions (green 0°, blue 90°, red 180°, yellow 270°) from -22.5 to 22.5° visual angle. Background contour indicates the edge, passing by the central column at second zero.

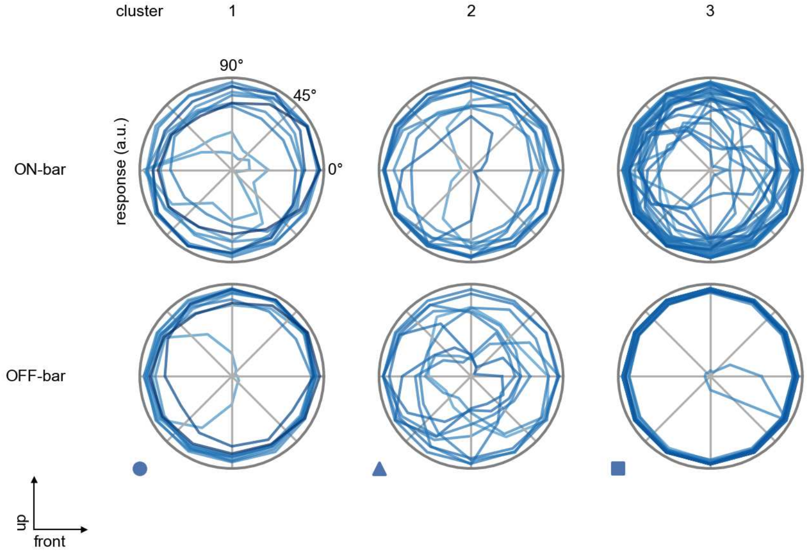

CT1(Lo1) - Figure 8: **Peak responses to moving bars.** The top row shows normalized peak responses to moving ON-bars, the bottom row shows normalized peak responses to moving OFF-bars. The bar-stimuli move in different directions from 0 to 360 degrees. Before, a grey-stimulus led the network to a stationary state. The responses from the different models in the different clusters (columns) overlay and are colored in the respective task error of the model. The peak responses are averaged over all assessed bar-speeds.

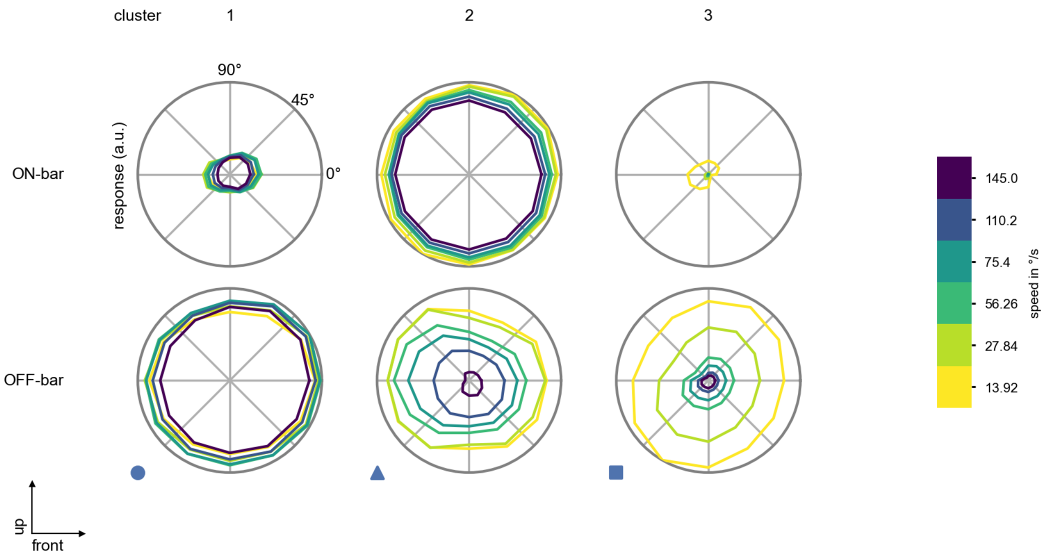

CT1(Lo1) - Figure 9: **Peak responses to moving bars from best-performing models.** The top row shows peak responses to moving ON-bars, the bottom row shows peak responses to moving OFF-bars. The bar-stimuli move in different directions from 0 to 360 degrees and at different speeds. Before, a grey-stimulus led the network to a stationary state. The columns show responses from the best-task-performing model in the respective cluster.

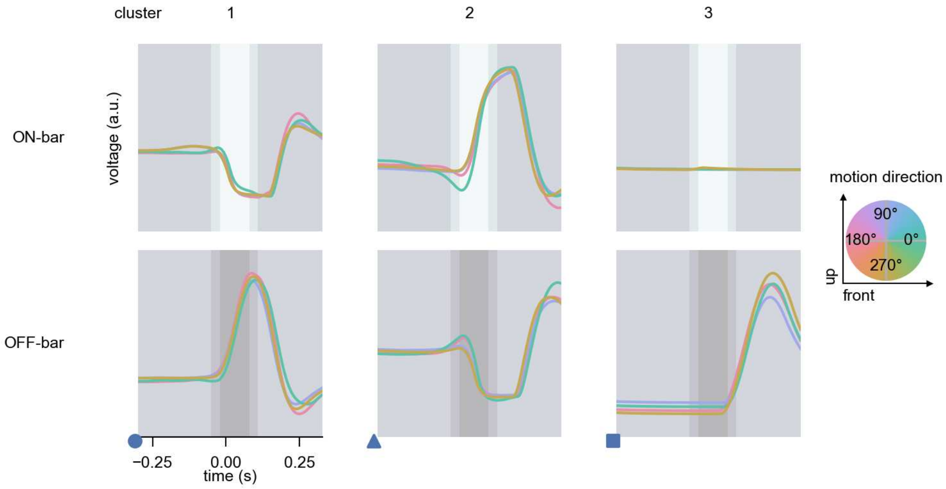

CT1(Lo1) - Figure 10: **Responses to moving bars from best-performing models.** The top row shows responses to moving ON-bars, the bottom row shows peak responses to moving OFF-bars. The bar-stimuli move in different directions from 0 to 360 degrees and at different speeds. Before, a grey-stimulus led the network to a stationary state. The columns show responses from the best-task-performing model in the respective cluster. Example responses to bars moving at 75.4°/s in all cardinal directions (green 0°, blue 90°, red 180°, yellow 270°) from -22.5 to 22.5° visual angle. Background contour indicates the bar, passing by the central column at second zero.

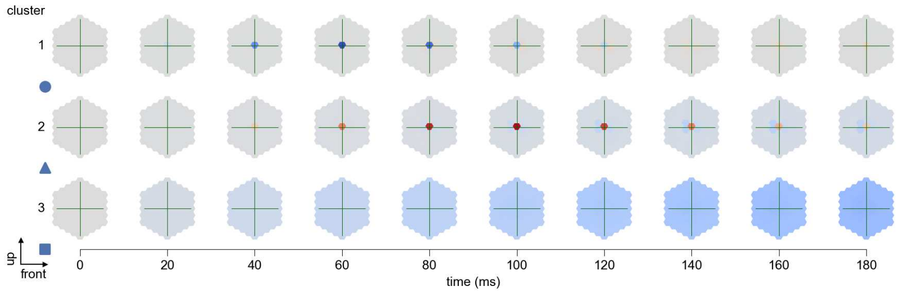

CT1(Lo1) - Figure 11: **Spatio-temporal receptive field.** Responses of the central cell to ON-impulses at single ommatidium impulse locations. The impulse occurs at second zero. Before, a grey-stimulus led the network to a stationary state. Responses are baseline-subtracted. Each row illustrates responses from the best-performing model of the respective cluster. Red indicated depolarization, blue indicates hyperpolarization.

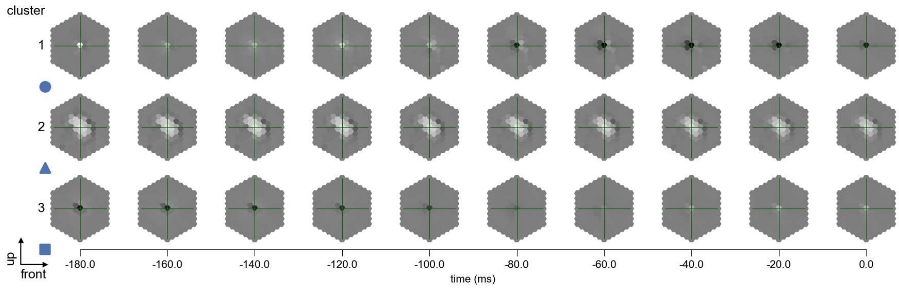

CT1(Lo1) - Figure 12: **Maximally excitatory stimuli.** Each row presents the regularized naturalistic-stimulus from the Sintel dataset that maximizes the cell type's central column response at second zero in the best-performing model of the respective cluster.

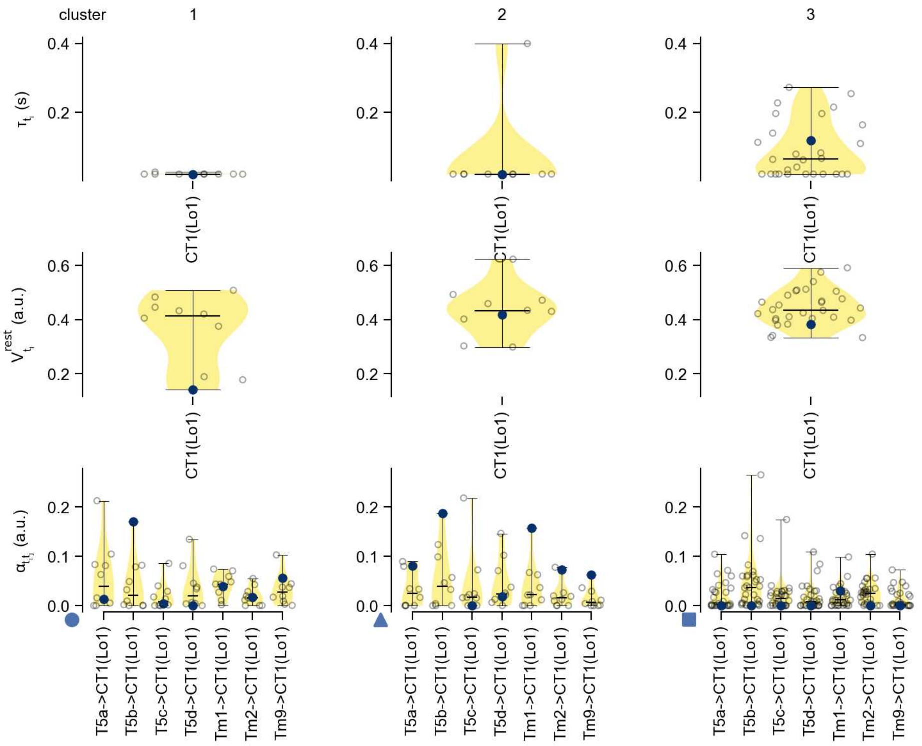

CT1(Lo1) - Figure 13: **Task-constrained parameters.** Each column shows the parameters inferred within the respective cluster. First row: learned time constants of the cell type. Second row: resting potentials of the cell type. Third row: scaling factors for the convolutional filters. The blue scatter represents the parameters from the best-performing model within the cluster.

# 20 CT1(M10)

← Cell types

#### Figures

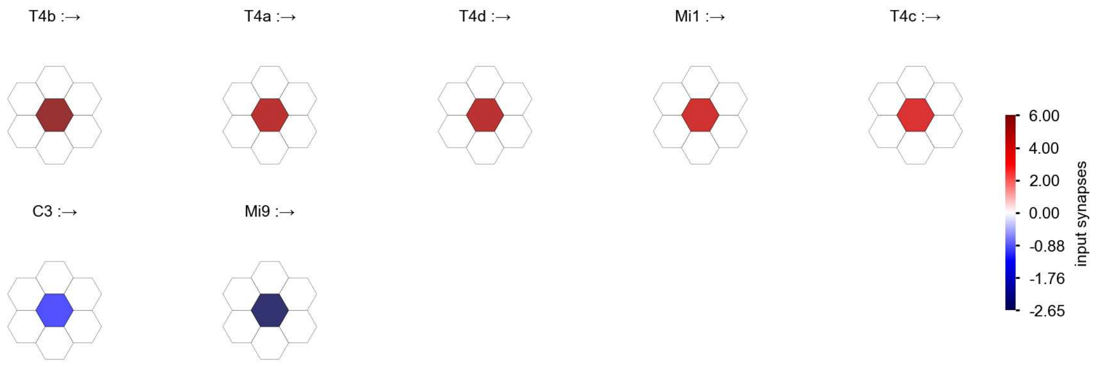

CT1(M10) - Figure 1: **Anatomical receptive fields.** Each colored hexagon is a projection, with the projection strength characterized by the average number of synapses that we count from the EM reconstruction. Red indicates excitatory synapses, blue indicates inhibitory synapses from inferred signs. Filters in the order of their total number of synapses.

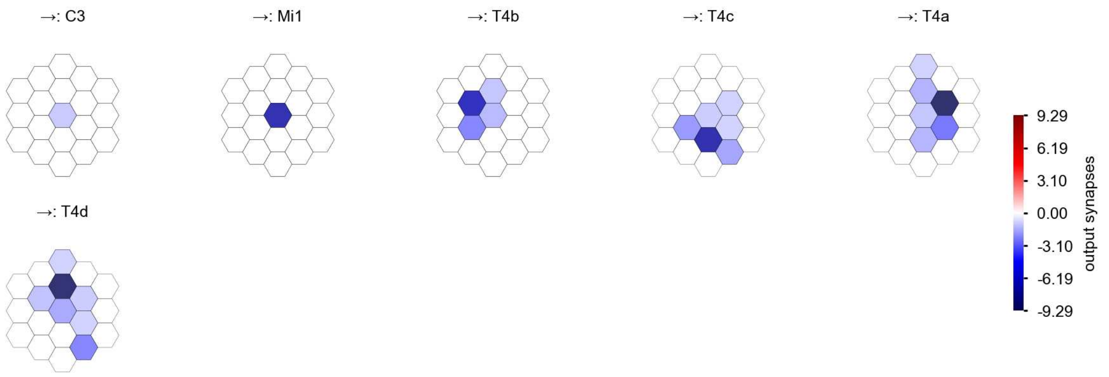

CT1(M10) - Figure 2: **Anatomical projective fields.** Each colored hexagon is a projection, with the projection strength characterized by the average number of synapses that we count from the EM reconstruction. Red indicates excitatory synapses, blue indicates inhibitory synapses from inferred signs. Filters in the order of their total number of synapses.

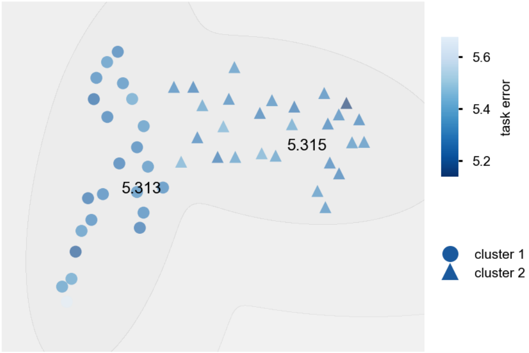

CT1(M10) - Figure 3: **Clustering of the responses to naturalistic stimuli.** From responses to naturalistic scenes from Sintel. Scatterpoints represent individual models colored by their task error.

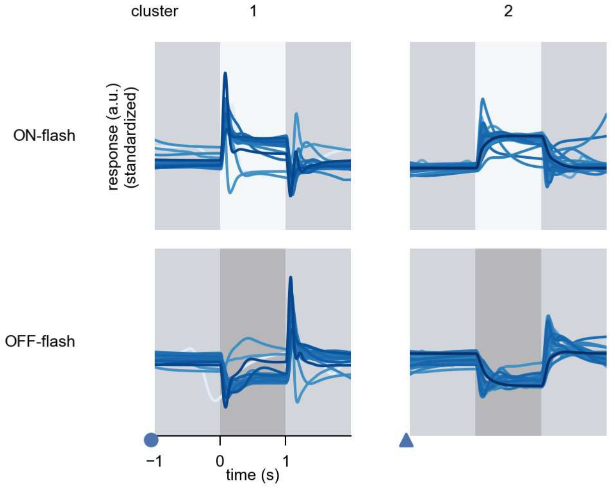

CT1(M10) - Figure 4: **Responses to flashes.** The top row shows responses to ON-flashes, the bottom row shows responses to OFF-flashes. The responses from the different models in the different clusters (columns) overlay and are colored in the respective task error of the model. The circular flashes occur at second zero, cover 6 ommatidia in radius, and last for one second. Before and after, a grey-stimulus leads to a stationary state of the network. The traces are standardized.

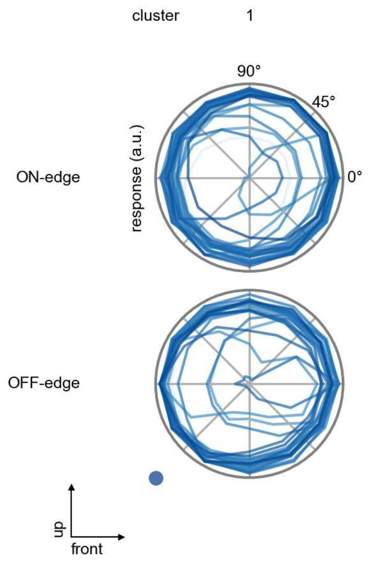

CT1(M10) - Figure 5: **Peak responses to moving edges.** The top row shows normalized peak responses to moving ON-edges, the bottom row shows normalized peak responses to moving OFF-edges. The edge-stimuli move in different directions from 0 to 360 degrees. Before, a grey-stimulus led the network to a stationary state. The responses from the different models in the different clusters (columns) overlay and are colored in the respective task error of the model. The peak responses are averaged over all assessed edge-speeds.

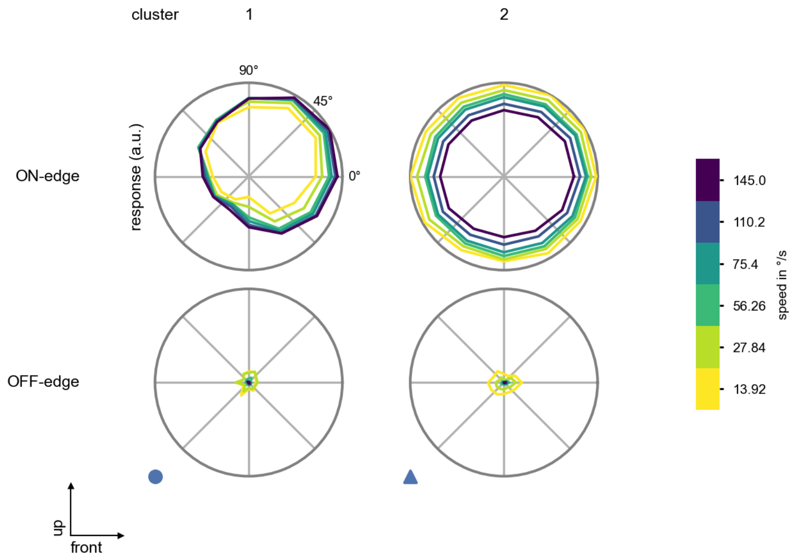

CT1(M10) - Figure 6: **Peak responses to moving edges from best-performing models.** The top row shows peak responses to moving ON-edges, the bottom row shows peak responses to moving OFF-edges. The edge-stimuli move in different directions from 0 to 360 degrees and at different speeds. Before, a grey-stimulus led the network to a stationary state. The columns show responses from the best-task-performing model in the respective cluster.

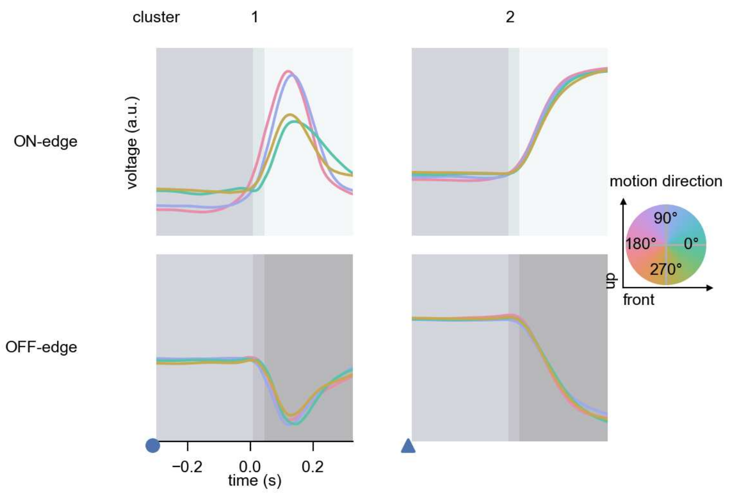

CT1(M10) - Figure 7: **Responses to moving edges from best-performing models.** The top row shows responses to moving ON-edges, the bottom row shows peak responses to moving OFF-edges. The edge-stimuli move in different directions from 0 to 360 degrees and at different speeds. Before, a grey-stimulus led the network to a stationary state. The columns show responses from the best-task-performing model in the respective cluster. Example responses to edges moving at 75.4°/s in all cardinal directions (green 0°, blue 90°, red 180°, yellow 270°) from -22.5 to 22.5° visual angle. Background contour indicates the edge, passing by the central column at second zero.

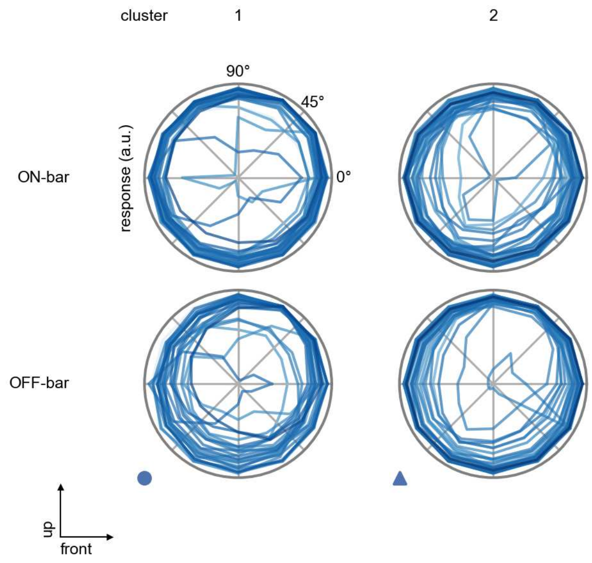

CT1(M10) - Figure 8: **Peak responses to moving bars.** The top row shows normalized peak responses to moving ON-bars, the bottom row shows normalized peak responses to moving OFF-bars. The bar-stimuli move in different directions from 0 to 360 degrees. Before, a grey-stimulus led the network to a stationary state. The responses from the different models in the different clusters (columns) overlay and are colored in the respective task error of the model. The peak responses are averaged over all assessed bar-speeds.

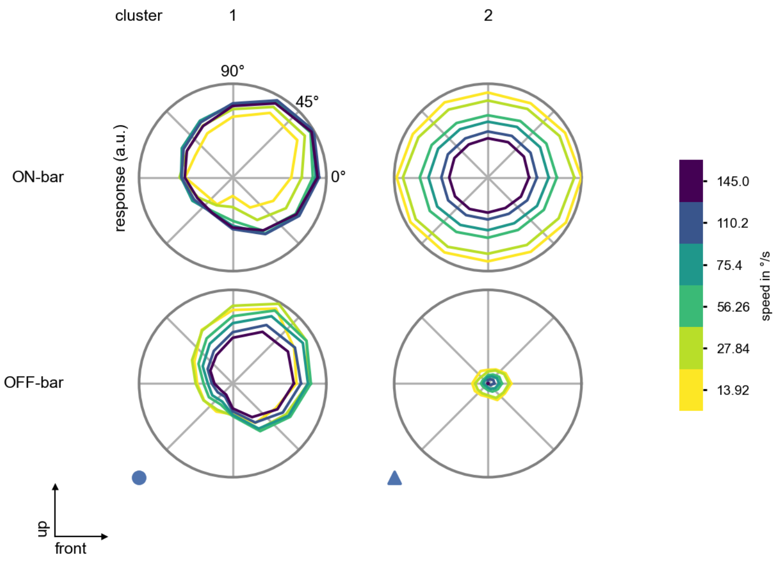

CT1(M10) - Figure 9: **Peak responses to moving bars from best-performing models.** The top row shows peak responses to moving ON-bars, the bottom row shows peak responses to moving OFF-bars. The bar-stimuli move in different directions from 0 to 360 degrees and at different speeds. Before, a grey-stimulus led the network to a stationary state. The columns show responses from the best-task-performing model in the respective cluster.

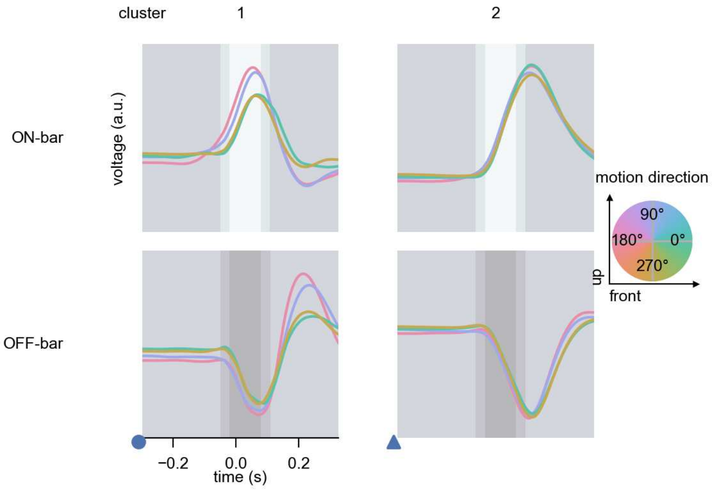

CT1(M10) - Figure 10: **Responses to moving bars from best-performing models.** The top row shows responses to moving ON-bars, the bottom row shows peak responses to moving OFF-bars. The bar-stimuli move in different directions from 0 to 360 degrees and at different speeds. Before, a grey-stimulus led the network to a stationary state. The columns show responses from the best-task-performing model in the respective cluster. Example responses to bars moving at 75.4°/s in all cardinal directions (green 0°, blue 90°, red 180°, yellow 270°) from -22.5 to 22.5° visual angle. Background contour indicates the bar, passing by the central column at second zero.

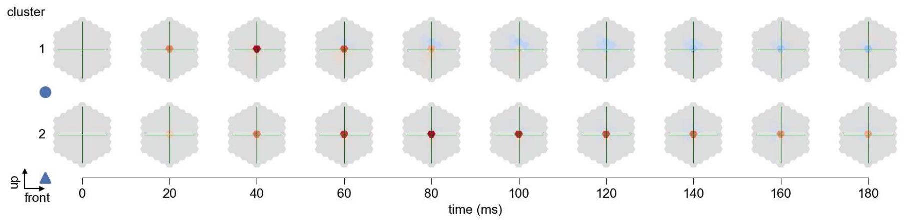

CT1(M10) - Figure 11: **Spatio-temporal receptive field.** Responses of the central cell to ON-impulses at single ommatidium impulse locations. The impulse occurs at second zero. Before, a grey-stimulus led the network to a stationary state. Responses are baseline-subtracted. Each row illustrates responses from the best-performing model of the respective cluster. Red indicated depolarization, blue indicates hyperpolarization.

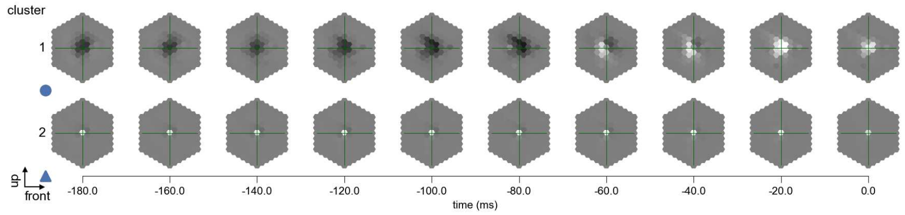

CT1(M10) - Figure 12: **Maximally excitatory stimuli.** Each row presents the regularized naturalistic-stimulus from the Sintel dataset that maximizes the cell type's central column response at second zero in the best-performing model of the respective cluster.

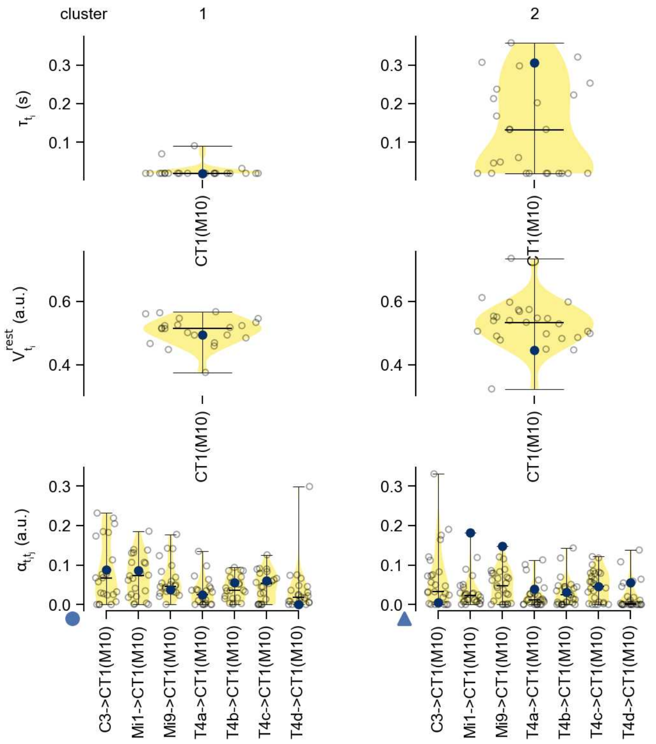

CT1(M10) - Figure 13: **Task-constrained parameters.** Each column shows the parameters inferred within the respective cluster. First row: learned time constants of the cell type. Second row: resting potentials of the cell type. Third row: scaling factors for the convolutional filters. The blue scatter represents the parameters from the best-performing model within the cluster.

# 21 Mi1

← Cell types

#### Figures

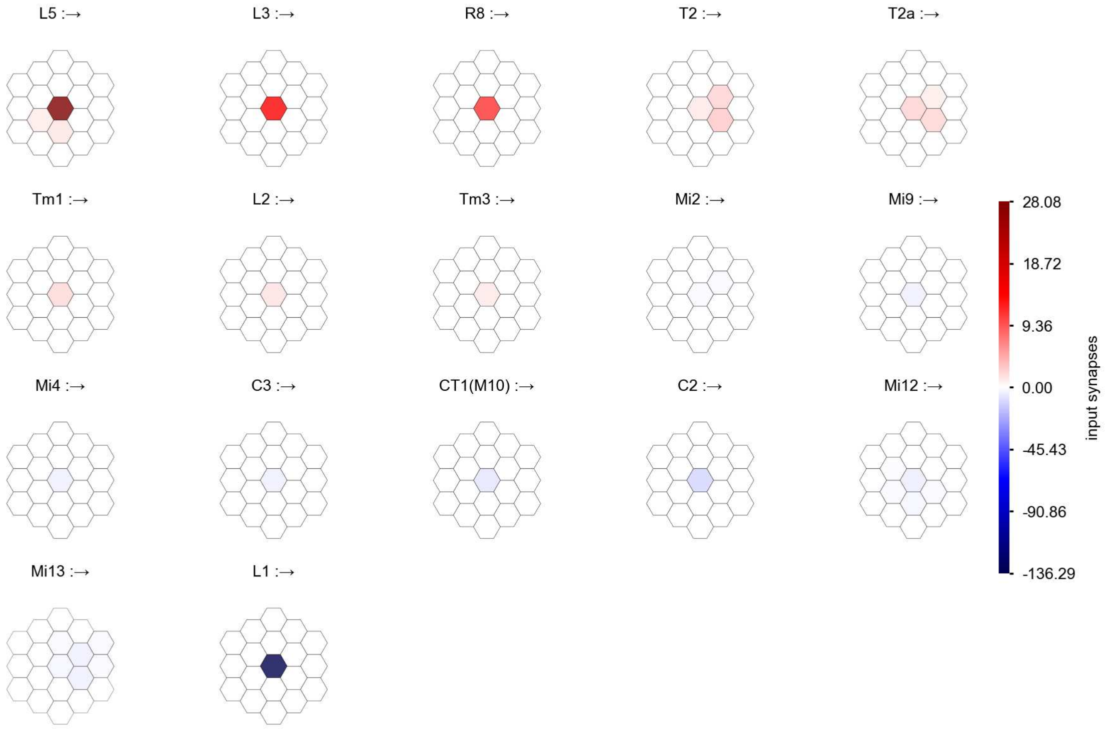

Mi1 - Figure 1: **Anatomical receptive fields.** Each colored hexagon is a projection, with the projection strength characterized by the average number of synapses that we count from the EM reconstruction. Red indicates excitatory synapses, blue indicates inhibitory synapses from inferred signs. Filters in the order of their total number of synapses.

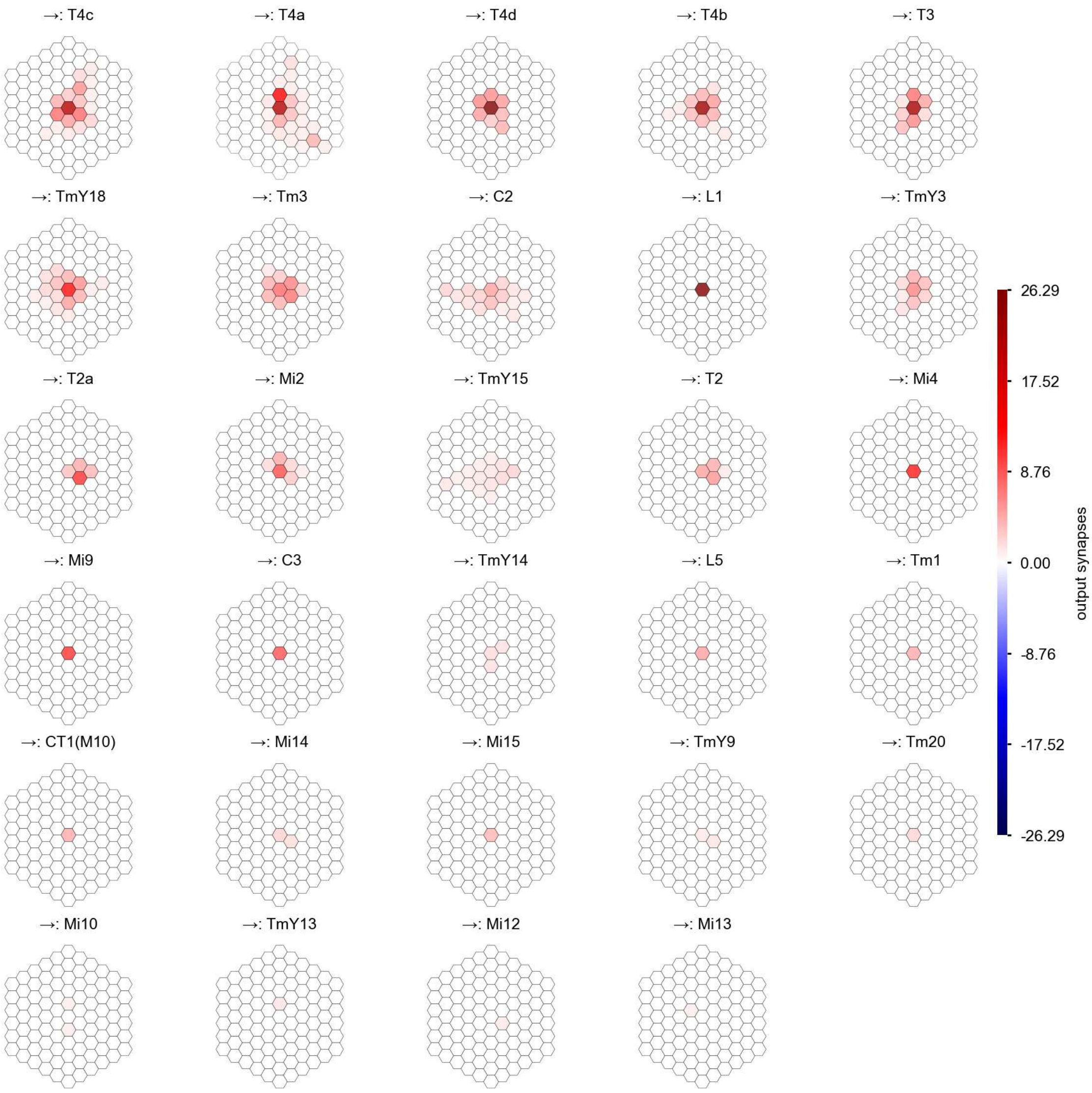

Mi1 - Figure 2: **Anatomical projective fields.** Each colored hexagon is a projection, with the projection strength characterized by the average number of synapses that we count from the EM reconstruction. Red indicates excitatory synapses, blue indicates inhibitory synapses from inferred signs. Filters in the order of their total number of synapses.

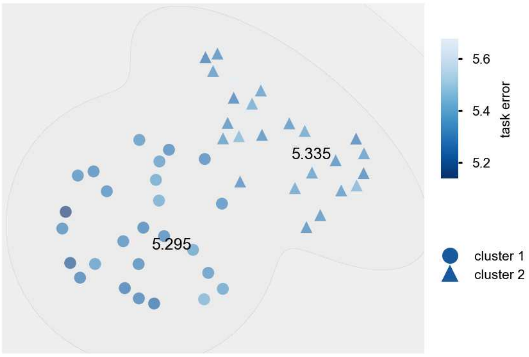

Mi1 - Figure 3: **Clustering of the responses to naturalistic stimuli.** From responses to naturalistic scenes from Sintel. Scatterpoints represent individual models colored by their task error.

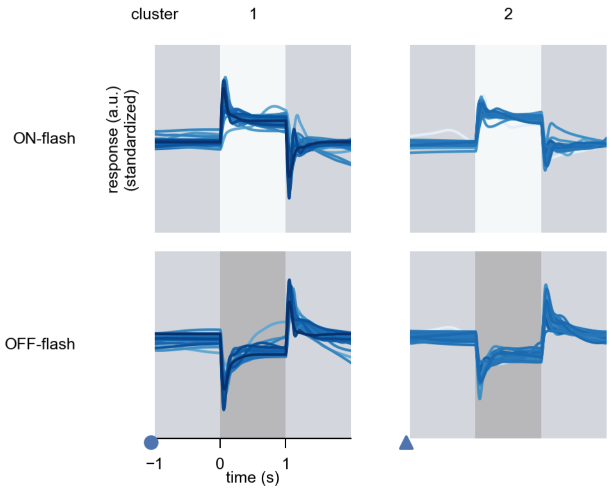

Mi1 - Figure 4: **Responses to flashes.** The top row shows responses to ON-flashes, the bottom row shows responses to OFF-flashes. The responses from the different models in the different clusters (columns) overlay and are colored in the respective task error of the model. The circular flashes occur at second zero, cover 6 ommatidia in radius, and last for one second. Before and after, a grey-stimulus leads to a stationary state of the network. The traces are standardized.

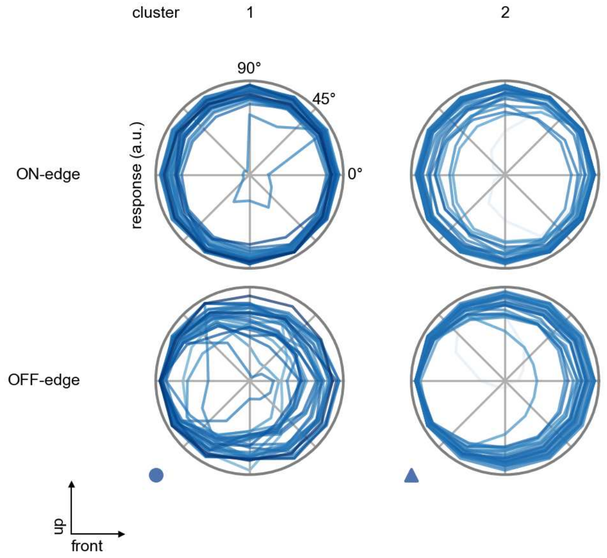

Mi1 - Figure 5: **Peak responses to moving edges.** The top row shows normalized peak responses to moving ON-edges, the bottom row shows normalized peak responses to moving OFF-edges. The edge-stimuli move in different directions from 0 to 360 degrees. Before, a grey-stimulus led the network to a stationary state. The responses from the different models in the different clusters (columns) overlay and are colored in the respective task error of the model. The peak responses are averaged over all assessed edge-speeds.

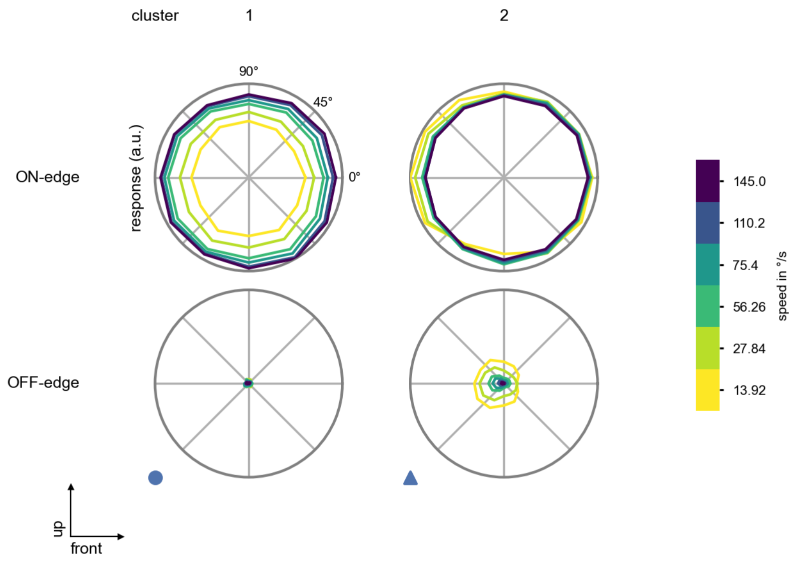

Mi1 - Figure 6: **Peak responses to moving edges from best-performing models.** The top row shows peak responses to moving ON-edges, the bottom row shows peak responses to moving OFF-edges. The edge-stimuli move in different directions from 0 to 360 degrees and at different speeds. Before, a grey-stimulus led the network to a stationary state. The columns show responses from the best-task-performing model in the respective cluster.

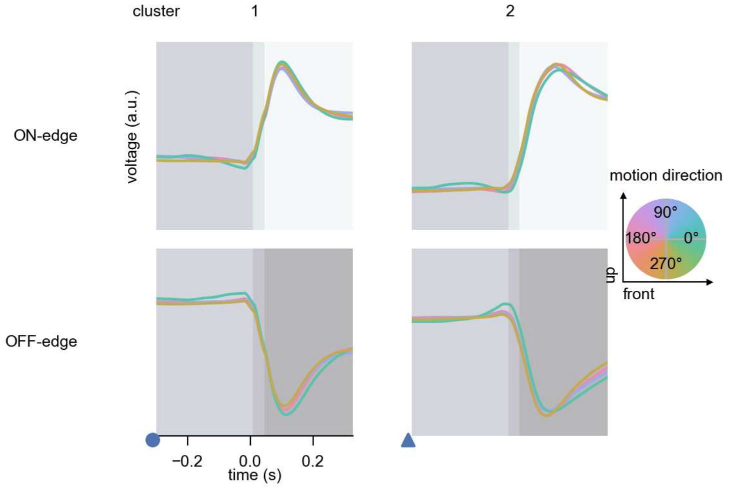

Mi1 - Figure 7: **Responses to moving edges from best-performing models.** The top row shows responses to moving ON-edges, the bottom row shows peak responses to moving OFF-edges. The edge-stimuli move in different directions from 0 to 360 degrees and at different speeds. Before, a grey-stimulus led the network to a stationary state. The columns show responses from the best-task-performing model in the respective cluster. Example responses to edges moving at 75.4°/s in all cardinal directions (green 0°, blue 90°, red 180°, yellow 270°) from -22.5 to 22.5° visual angle. Background contour indicates the edge, passing by the central column at second zero.

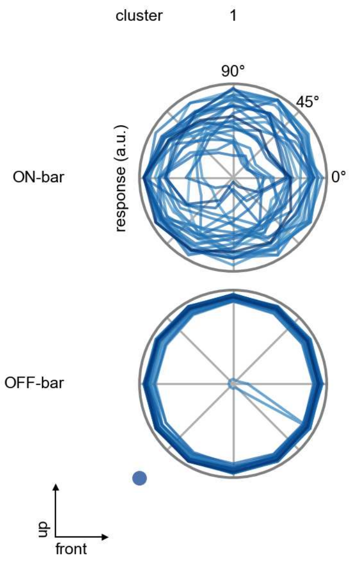

Mi1 - Figure 8: **Peak responses to moving bars.** The top row shows normalized peak responses to moving ON-bars, the bottom row shows normalized peak responses to moving OFF-bars. The bar-stimuli move in different directions from 0 to 360 degrees. Before, a grey-stimulus led the network to a stationary state. The responses from the different models in the different clusters (columns) overlay and are colored in the respective task error of the model. The peak responses are averaged over all assessed bar-speeds.

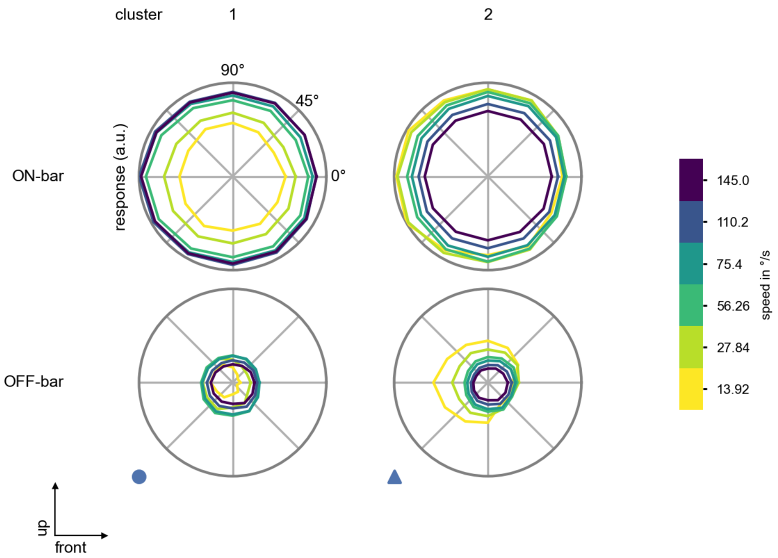

Mi1 - Figure 9: **Peak responses to moving bars from best-performing models.** The top row shows peak responses to moving ON-bars, the bottom row shows peak responses to moving OFF-bars. The bar-stimuli move in different directions from 0 to 360 degrees and at different speeds. Before, a grey-stimulus led the network to a stationary state. The columns show responses from the best-task-performing model in the respective cluster.

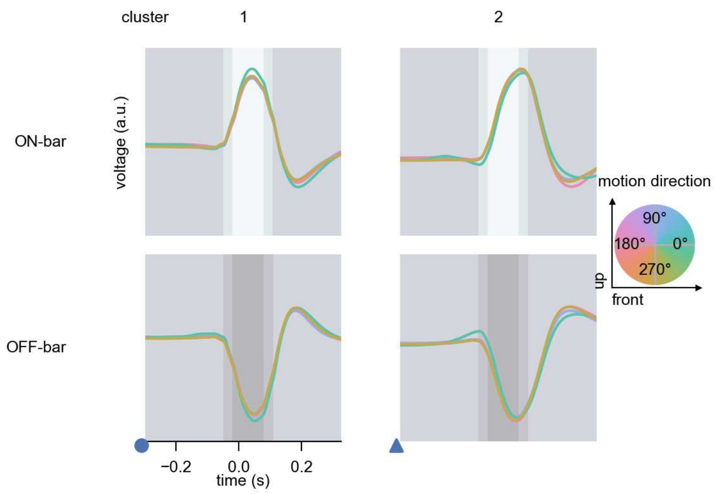

Mi1 - Figure 10: **Responses to moving bars from best-performing models.** The top row shows responses to moving ON-bars, the bottom row shows peak responses to moving OFF-bars. The bar-stimuli move in different directions from 0 to 360 degrees and at different speeds. Before, a grey-stimulus led the network to a stationary state. The columns show responses from the best-task-performing model in the respective cluster. Example responses to bars moving at 75.4°/s in all cardinal directions (green 0°, blue 90°, red 180°, yellow 270°) from -22.5 to 22.5° visual angle. Background contour indicates the bar, passing by the central column at second zero.

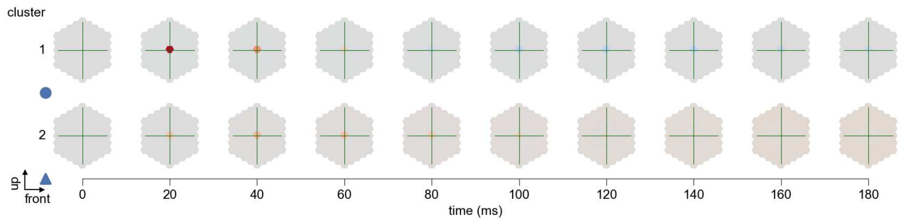

Mi1 - Figure 11: **Spatio-temporal receptive field.** Responses of the central cell to ON-impulses at single ommatidium impulse locations. The impulse occurs at second zero. Before, a grey-stimulus led the network to a stationary state. Responses are baseline-subtracted. Each row illustrates responses from the best-performing model of the respective cluster. Red indicated depolarization, blue indicates hyperpolarization.

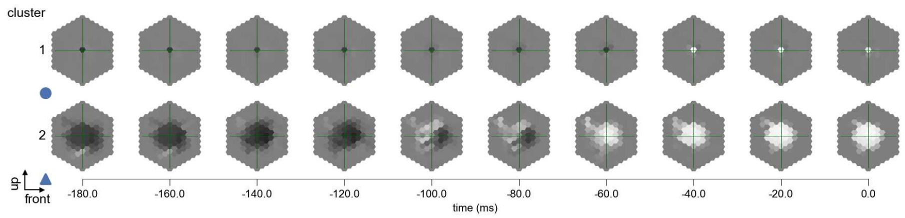

Mi1 - Figure 12: **Maximally excitatory stimuli.** Each row presents the regularized naturalistic-stimulus from the Sintel dataset that maximizes the cell type's central column response at second zero in the best-performing model of the respective cluster.

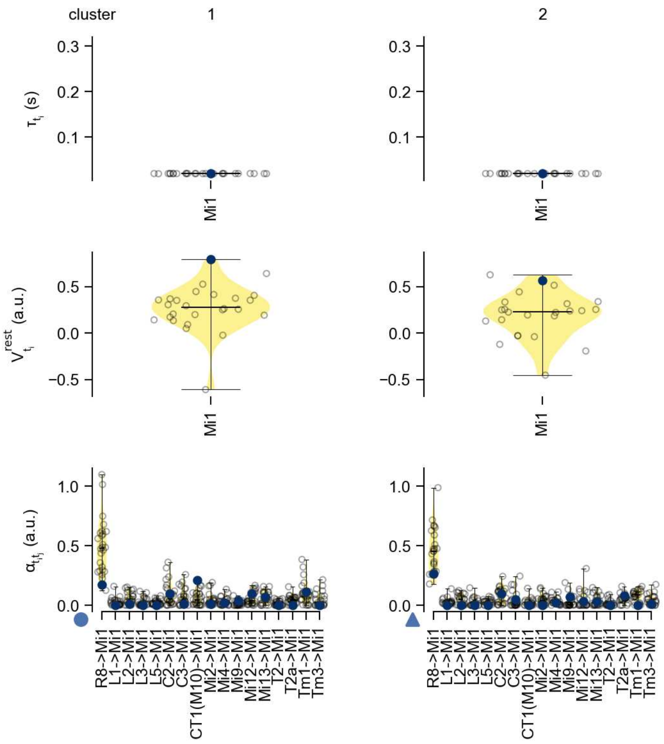

Mi1 - Figure 13: **Task-constrained parameters.** Each column shows the parameters inferred within the respective cluster. First row: learned time constants of the cell type. Second row: resting potentials of the cell type. Third row: scaling factors for the convolutional filters. The blue scatter represents the parameters from the best-performing model within the cluster.

# 22 Mi2

← Cell types

#### Figures

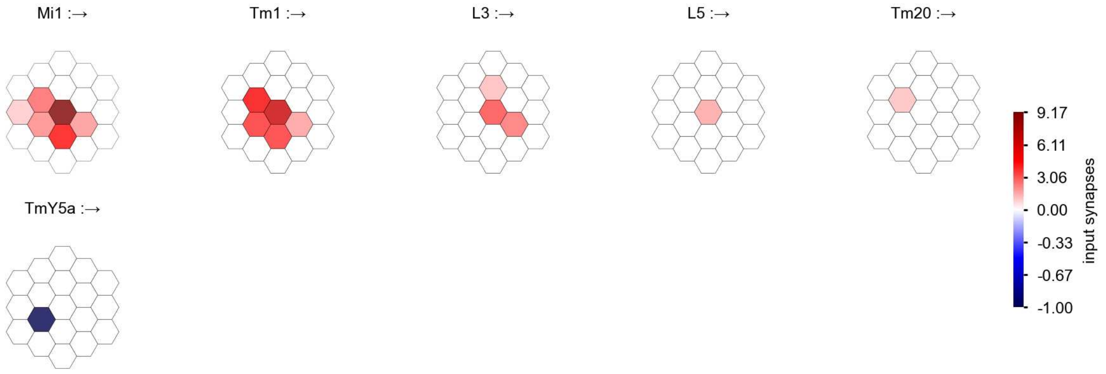

Mi2 - Figure 1: **Anatomical receptive fields.** Each colored hexagon is a projection, with the projection strength characterized by the average number of synapses that we count from the EM reconstruction. Red indicates excitatory synapses, blue indicates inhibitory synapses from inferred signs. Filters in the order of their total number of synapses.

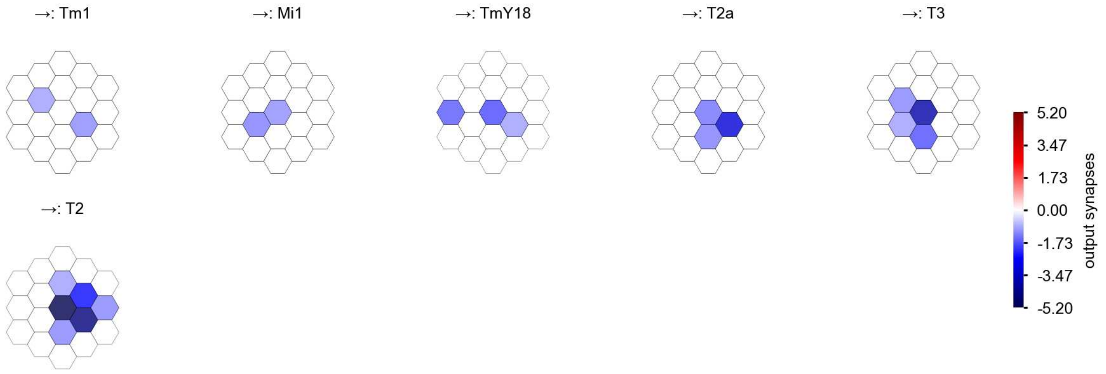

Mi2 - Figure 2: **Anatomical projective fields.** Each colored hexagon is a projection, with the projection strength characterized by the average number of synapses that we count from the EM reconstruction. Red indicates excitatory synapses, blue indicates inhibitory synapses from inferred signs. Filters in the order of their total number of synapses.

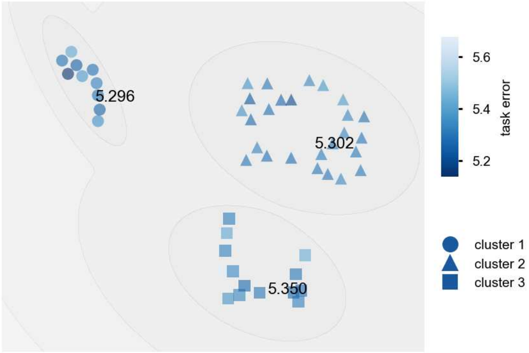

Mi2 - Figure 3: **Clustering of the responses to naturalistic stimuli.** From responses to naturalistic scenes from Sintel. Scatterpoints represent individual models colored by their task error.

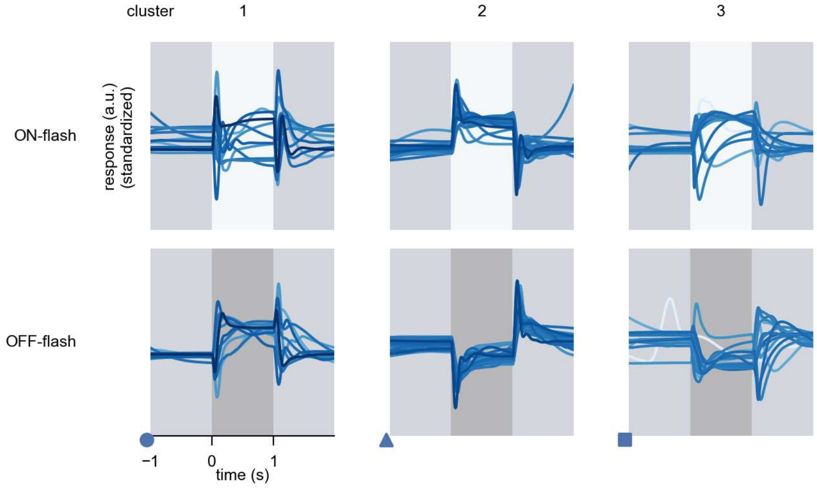

Mi2 - Figure 4: **Responses to flashes.** The top row shows responses to ON-flashes, the bottom row shows responses to OFF-flashes. The responses from the different models in the different clusters (columns) overlay and are colored in the respective task error of the model. The circular flashes occur at second zero, cover 6 ommatidia in radius, and last for one second. Before and after, a grey-stimulus leads to a stationary state of the network. The traces are standardized.

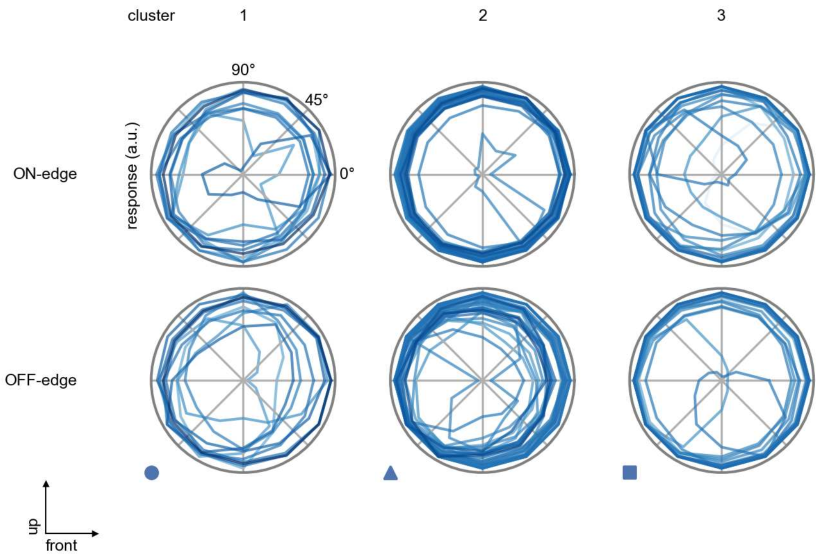

Mi2 - Figure 5: **Peak responses to moving edges.** The top row shows normalized peak responses to moving ON-edges, the bottom row shows normalized peak responses to moving OFF-edges. The edge-stimuli move in different directions from 0 to 360 degrees. Before, a grey-stimulus led the network to a stationary state. The responses from the different models in the different clusters (columns) overlay and are colored in the respective task error of the model. The peak responses are averaged over all assessed edge-speeds.

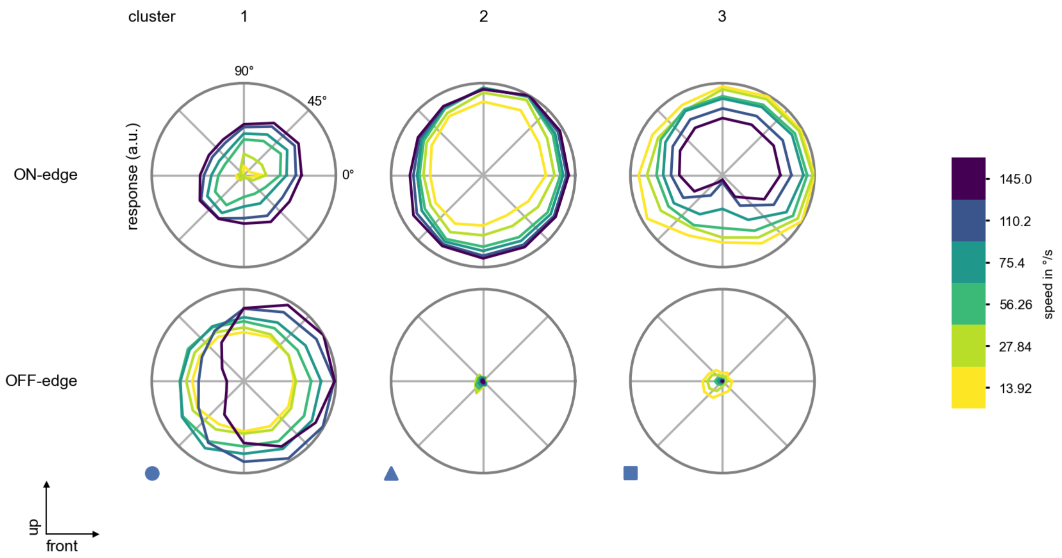

Mi2 - Figure 6: **Peak responses to moving edges from best-performing models.** The top row shows peak responses to moving ON-edges, the bottom row shows peak responses to moving OFF-edges. The edge-stimuli move in different directions from 0 to 360 degrees and at different speeds. Before, a grey-stimulus led the network to a stationary state. The columns show responses from the best-task-performing model in the respective cluster.

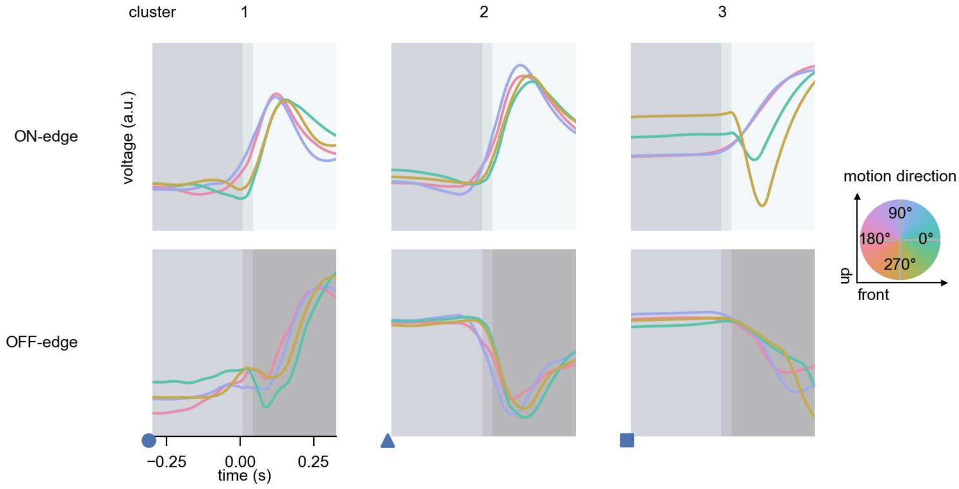

Mi2 - Figure 7: **Responses to moving edges from best-performing models.** The top row shows responses to moving ON-edges, the bottom row shows peak responses to moving OFF-edges. The edge-stimuli move in different directions from 0 to 360 degrees and at different speeds. Before, a grey-stimulus led the network to a stationary state. The columns show responses from the best-task-performing model in the respective cluster. Example responses to edges moving at 75.4°/s in all cardinal directions (green 0°, blue 90°, red 180°, yellow 270°) from -22.5 to 22.5° visual angle. Background contour indicates the edge, passing by the central column at second zero.

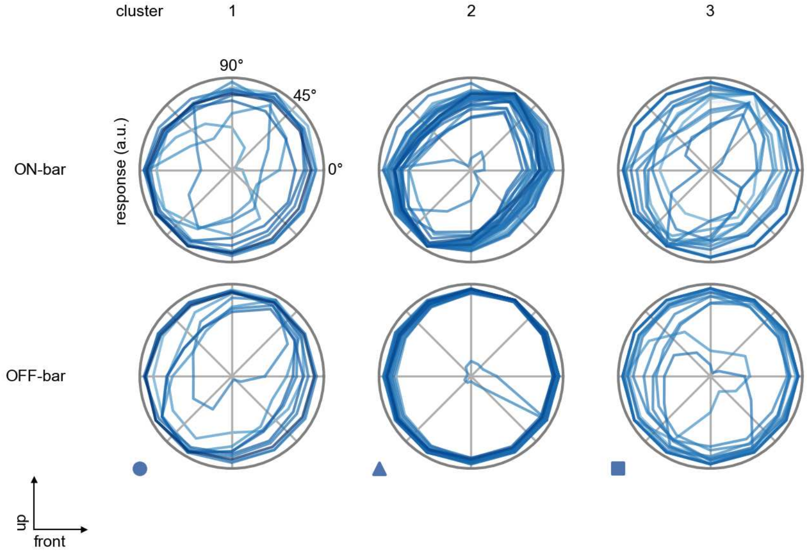

Mi2 - Figure 8: **Peak responses to moving bars.** The top row shows normalized peak responses to moving ON-bars, the bottom row shows normalized peak responses to moving OFF-bars. The bar-stimuli move in different directions from 0 to 360 degrees. Before, a grey-stimulus led the network to a stationary state. The responses from the different models in the different clusters (columns) overlay and are colored in the respective task error of the model. The peak responses are averaged over all assessed bar-speeds.

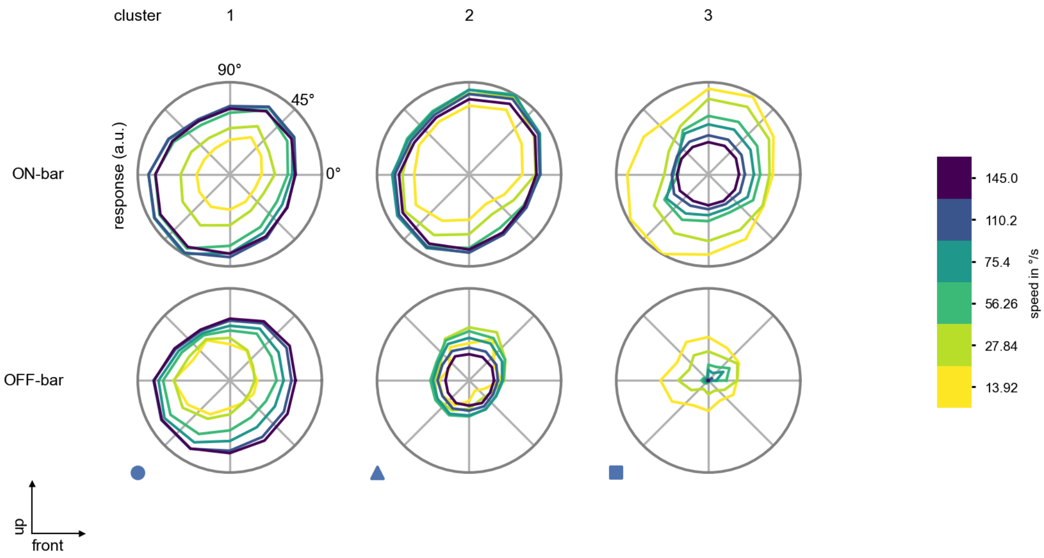

Mi2 - Figure 9: **Peak responses to moving bars from best-performing models.** The top row shows peak responses to moving ON-bars, the bottom row shows peak responses to moving OFF-bars. The bar-stimuli move in different directions from 0 to 360 degrees and at different speeds. Before, a grey-stimulus led the network to a stationary state. The columns show responses from the best-task-performing model in the respective cluster.

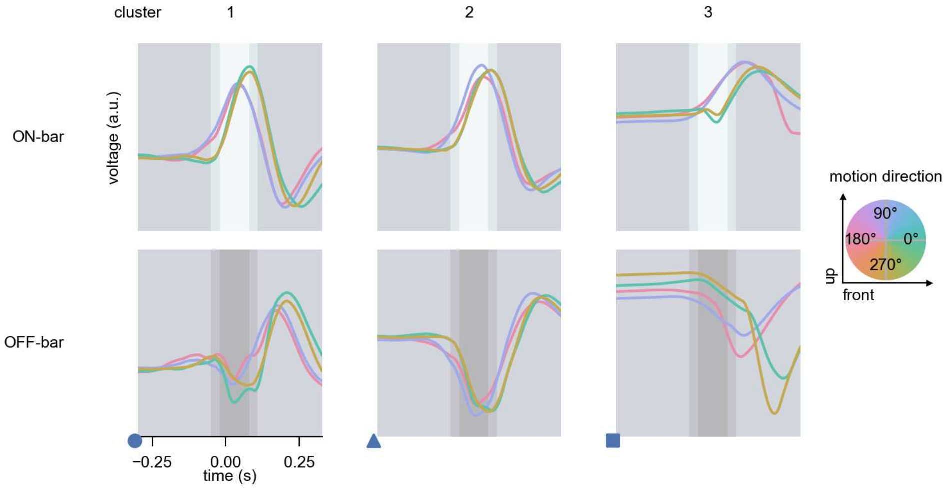

Mi2 - Figure 10: **Responses to moving bars from best-performing models.** The top row shows responses to moving ON-bars, the bottom row shows peak responses to moving OFF-bars. The bar-stimuli move in different directions from 0 to 360 degrees and at different speeds. Before, a grey-stimulus led the network to a stationary state. The columns show responses from the best-task-performing model in the respective cluster. Example responses to bars moving at 75.4°/s in all cardinal directions (green 0°, blue 90°, red 180°, yellow 270°) from -22.5 to 22.5° visual angle. Background contour indicates the bar, passing by the central column at second zero.

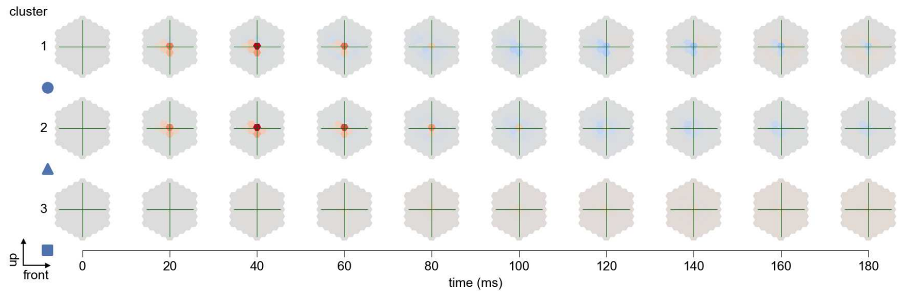

Mi2 - Figure 11: **Spatio-temporal receptive field.** Responses of the central cell to ON-impulses at single ommatidium impulse locations. The impulse occurs at second zero. Before, a grey-stimulus led the network to a stationary state. Responses are baseline-subtracted. Each row illustrates responses from the best-performing model of the respective cluster. Red indicated depolarization, blue indicates hyperpolarization.

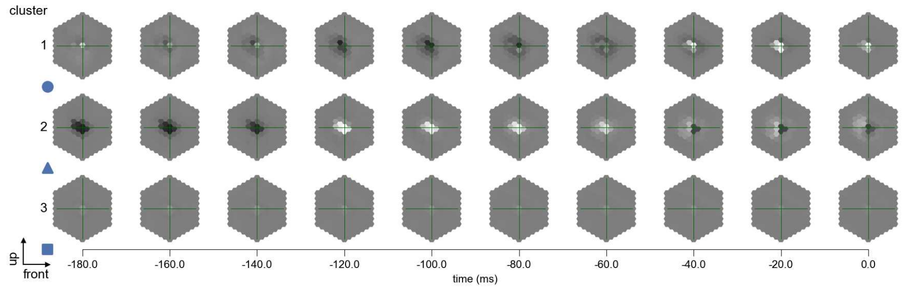

Mi2 - Figure 12: **Maximally excitatory stimuli.** Each row presents the regularized naturalistic-stimulus from the Sintel dataset that maximizes the cell type's central column response at second zero in the best-performing model of the respective cluster.

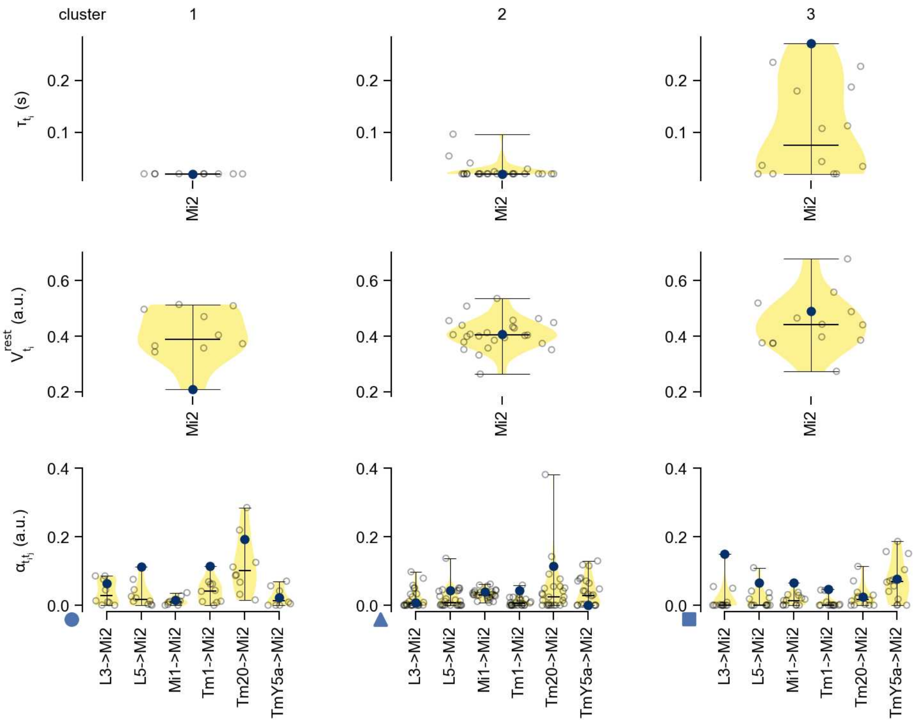

Mi2 - Figure 13: **Task-constrained parameters.** Each column shows the parameters inferred within the respective cluster. First row: learned time constants of the cell type. Second row: resting potentials of the cell type. Third row: scaling factors for the convolutional filters. The blue scatter represents the parameters from the best-performing model within the cluster.

# 23 Mi3

← Cell types

#### Figures

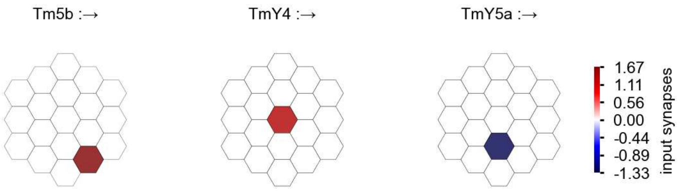

Mi3 - Figure 1: **Anatomical receptive fields.** Each colored hexagon is a projection, with the projection strength characterized by the average number of synapses that we count from the EM reconstruction. Red indicates excitatory synapses, blue indicates inhibitory synapses from inferred signs. Filters in the order of their total number of synapses.

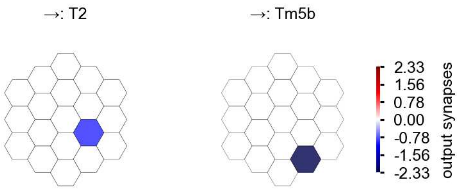

Mi3 - Figure 2: **Anatomical projective fields.** Each colored hexagon is a projection, with the projection strength characterized by the average number of synapses that we count from the EM reconstruction. Red indicates excitatory synapses, blue indicates inhibitory synapses from inferred signs. Filters in the order of their total number of synapses.

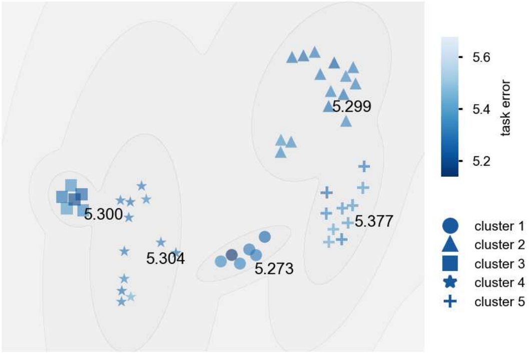

Mi3 - Figure 3: **Clustering of the responses to naturalistic stimuli.** From responses to naturalistic scenes from Sintel. Scatterpoints represent individual models colored by their task error.

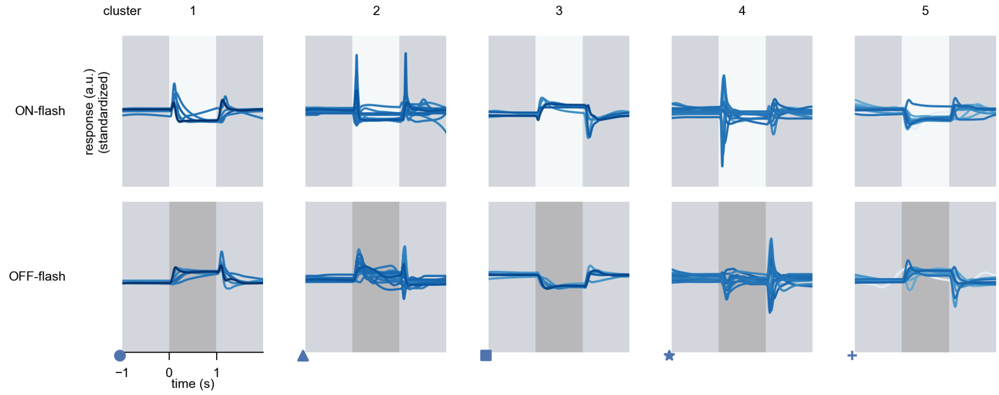

Mi3 - Figure 4: **Responses to flashes.** The top row shows responses to ON-flashes, the bottom row shows responses to OFF-flashes. The responses from the different models in the different clusters (columns) overlay and are colored in the respective task error of the model. The circular flashes occur at second zero, cover 6 ommatidia in radius, and last for one second. Before and after, a grey-stimulus leads to a stationary state of the network. The traces are standardized.

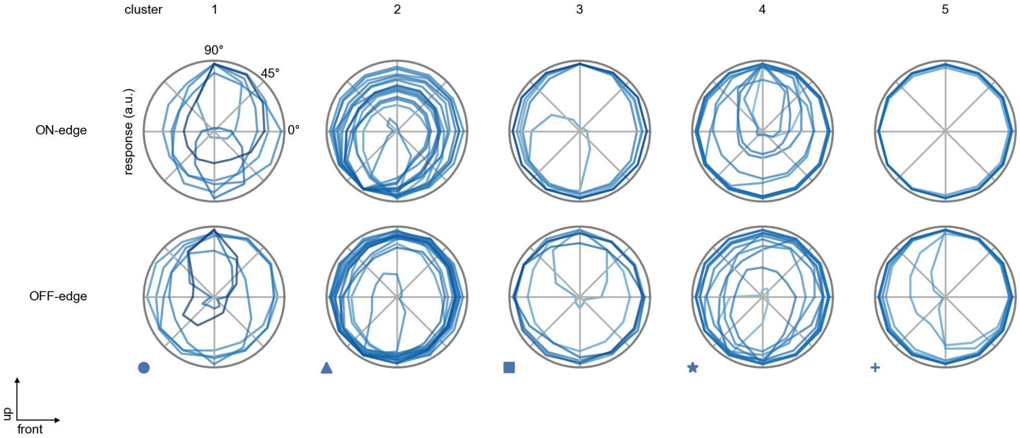

Mi3 - Figure 5: **Peak responses to moving edges.** The top row shows normalized peak responses to moving ON-edges, the bottom row shows normalized peak responses to moving OFF-edges. The edge-stimuli move in different directions from 0 to 360 degrees. Before, a grey-stimulus led the network to a stationary state. The responses from the different models in the different clusters (columns) overlay and are colored in the respective task error of the model. The peak responses are averaged over all assessed edge-speeds.

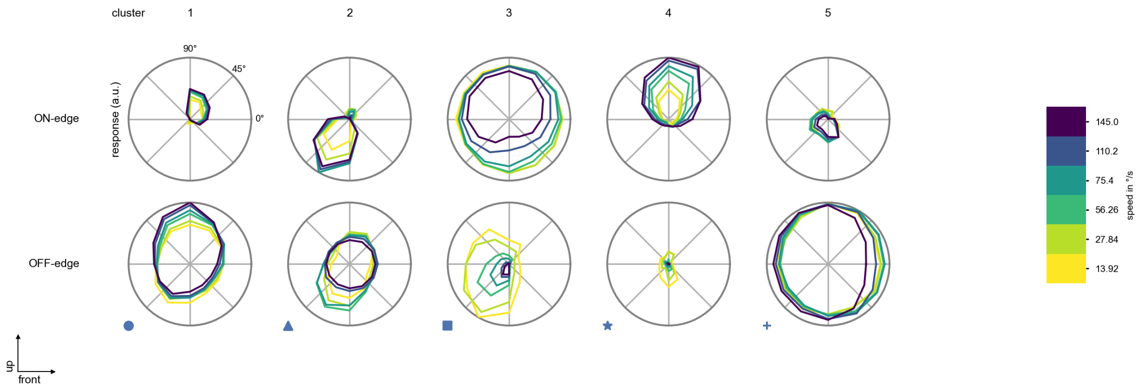

Mi3 - Figure 6: **Peak responses to moving edges from best-performing models.** The top row shows peak responses to moving ON-edges, the bottom row shows peak responses to moving OFF-edges. The edge-stimuli move in different directions from 0 to 360 degrees and at different speeds. Before, a grey-stimulus led the network to a stationary state. The columns show responses from the best-task-performing model in the respective cluster.

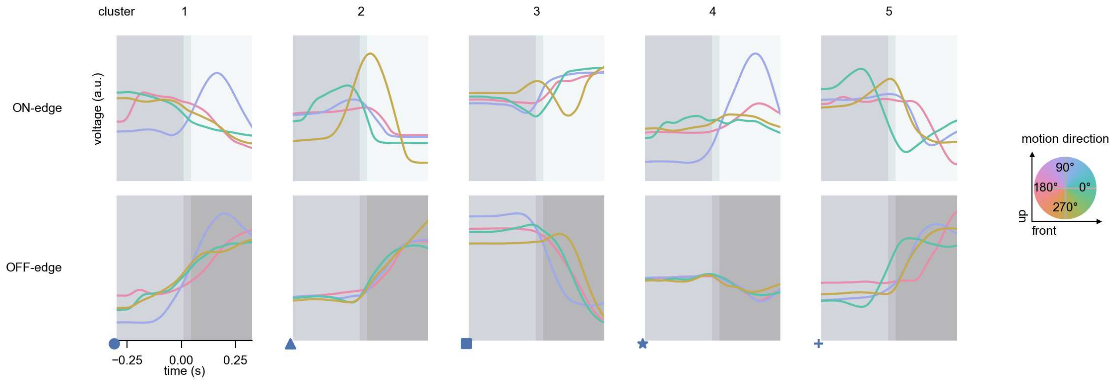

Mi3 - Figure 7: **Responses to moving edges from best-performing models.** The top row shows responses to moving ON-edges, the bottom row shows peak responses to moving OFF-edges. The edge-stimuli move in different directions from 0 to 360 degrees and at different speeds. Before, a grey-stimulus led the network to a stationary state. The columns show responses from the best-task-performing model in the respective cluster. Example responses to edges moving at 75.4°/s in all cardinal directions (green 0°, blue 90°, red 180°, yellow 270°) from -22.5 to 22.5° visual angle. Background contour indicates the edge, passing by the central column at second zero.

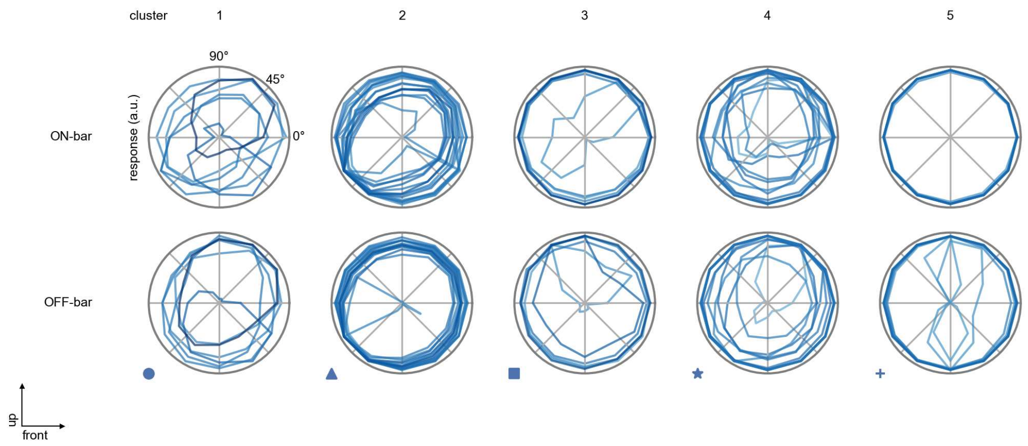

Mi3 - Figure 8: **Peak responses to moving bars.** The top row shows normalized peak responses to moving ON-bars, the bottom row shows normalized peak responses to moving OFF-bars. The bar-stimuli move in different directions from 0 to 360 degrees. Before, a grey-stimulus led the network to a stationary state. The responses from the different models in the different clusters (columns) overlay and are colored in the respective task error of the model. The peak responses are averaged over all assessed bar-speeds.

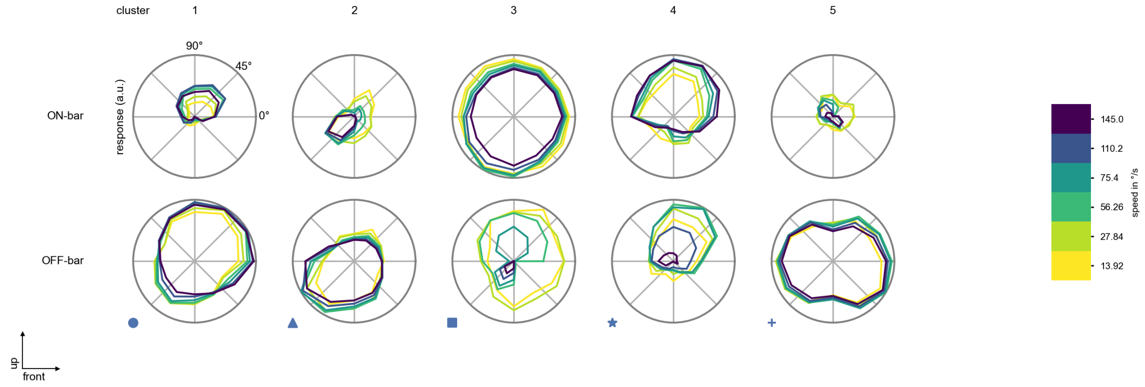

Mi3 - Figure 9: **Peak responses to moving bars from best-performing models.** The top row shows peak responses to moving ON-bars, the bottom row shows peak responses to moving OFF-bars. The bar-stimuli move in different directions from 0 to 360 degrees and at different speeds. Before, a grey-stimulus led the network to a stationary state. The columns show responses from the best-task-performing model in the respective cluster.

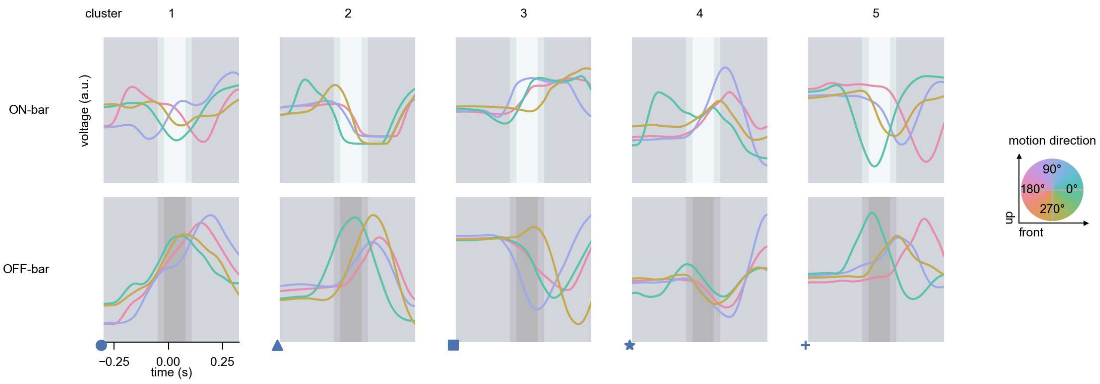

Mi3 - Figure 10: **Responses to moving bars from best-performing models.** The top row shows responses to moving ON-bars, the bottom row shows peak responses to moving OFF-bars. The bar-stimuli move in different directions from 0 to 360 degrees and at different speeds. Before, a grey-stimulus led the network to a stationary state. The columns show responses from the best-task-performing model in the respective cluster. Example responses to bars moving at 75.4°/s in all cardinal directions (green 0°, blue 90°, red 180°, yellow 270°) from -22.5 to 22.5° visual angle. Background contour indicates the bar, passing by the central column at second zero.

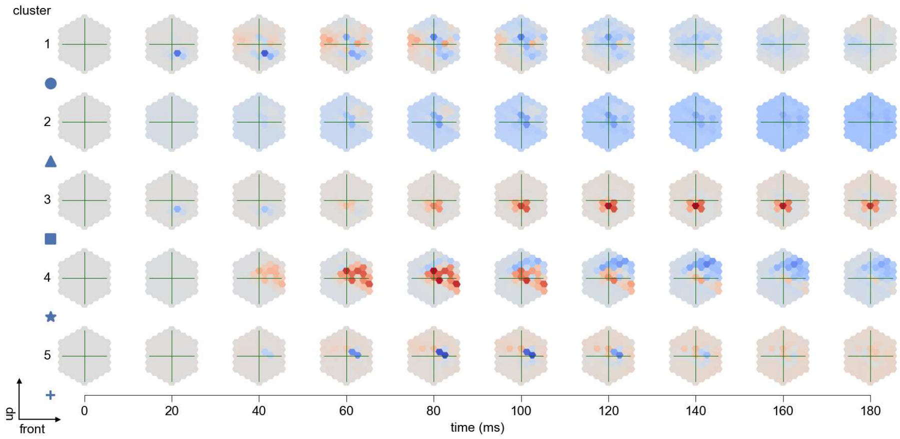

Mi3 - Figure 11: **Spatio-temporal receptive field.** Responses of the central cell to ON-impulses at single ommatidium impulse locations. The impulse occurs at second zero. Before, a grey-stimulus led the network to a stationary state. Responses are baseline-subtracted. Each row illustrates responses from the best-performing model of the respective cluster. Red indicated depolarization, blue indicates hyperpolarization.

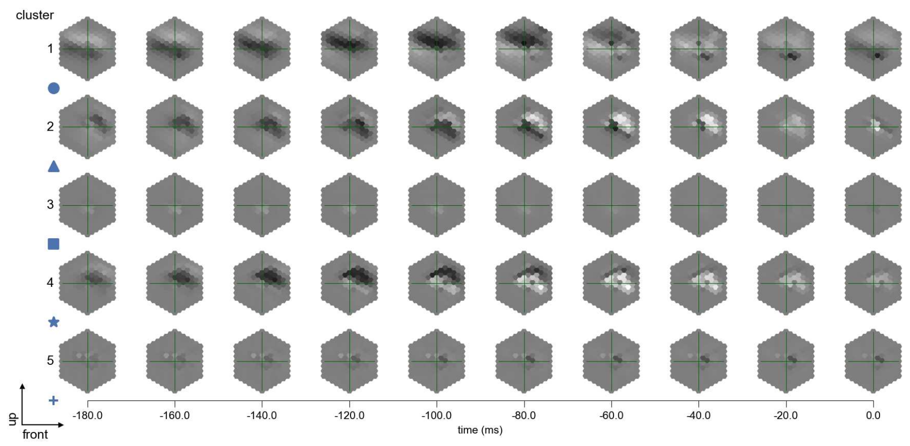

Mi3 - Figure 12: **Maximally excitatory stimuli.** Each row presents the regularized naturalistic-stimulus from the Sintel dataset that maximizes the cell type's central column response at second zero in the best-performing model of the respective cluster.

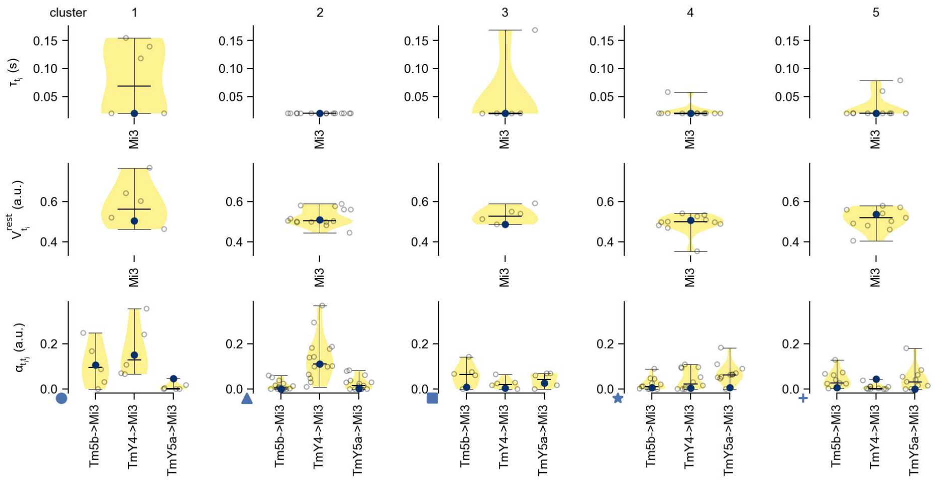

Mi3 - Figure 13: **Task-constrained parameters.** Each column shows the parameters inferred within the respective cluster. First row: learned time constants of the cell type. Second row: resting potentials of the cell type. Third row: scaling factors for the convolutional filters. The blue scatter represents the parameters from the best-performing model within the cluster.

# 24 Mi4

← Cell types

#### Figures

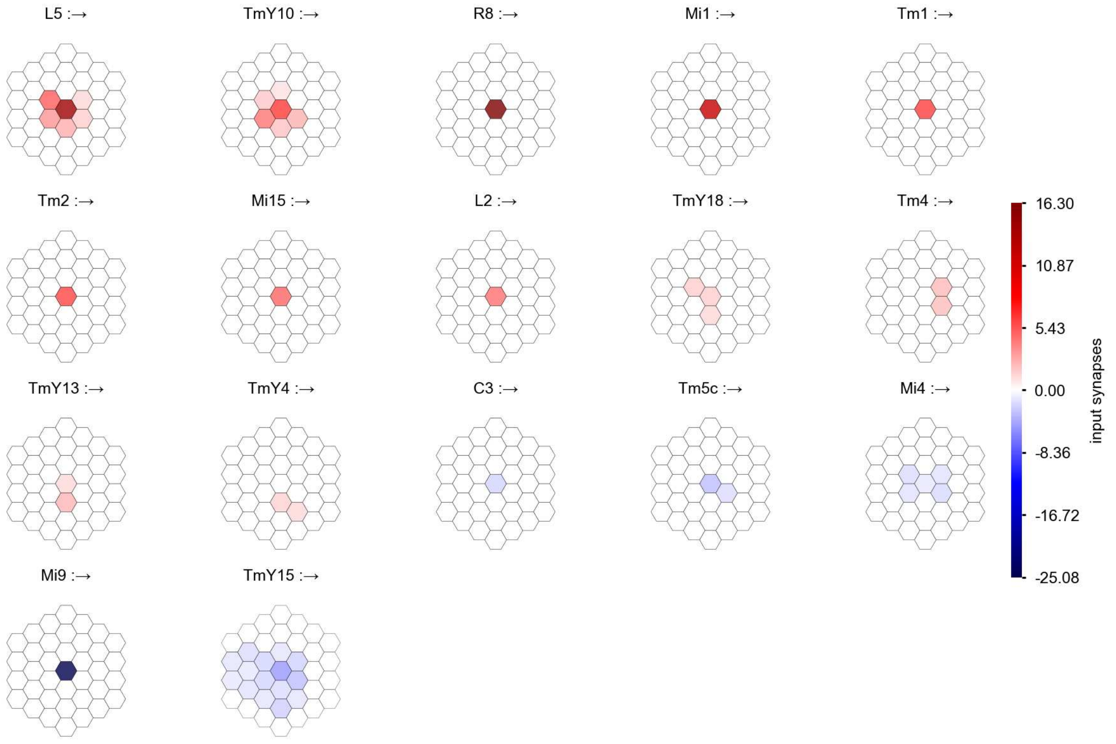

Mi4 - Figure 1: **Anatomical receptive fields.** Each colored hexagon is a projection, with the projection strength characterized by the average number of synapses that we count from the EM reconstruction. Red indicates excitatory synapses, blue indicates inhibitory synapses from inferred signs. Filters in the order of their total number of synapses.

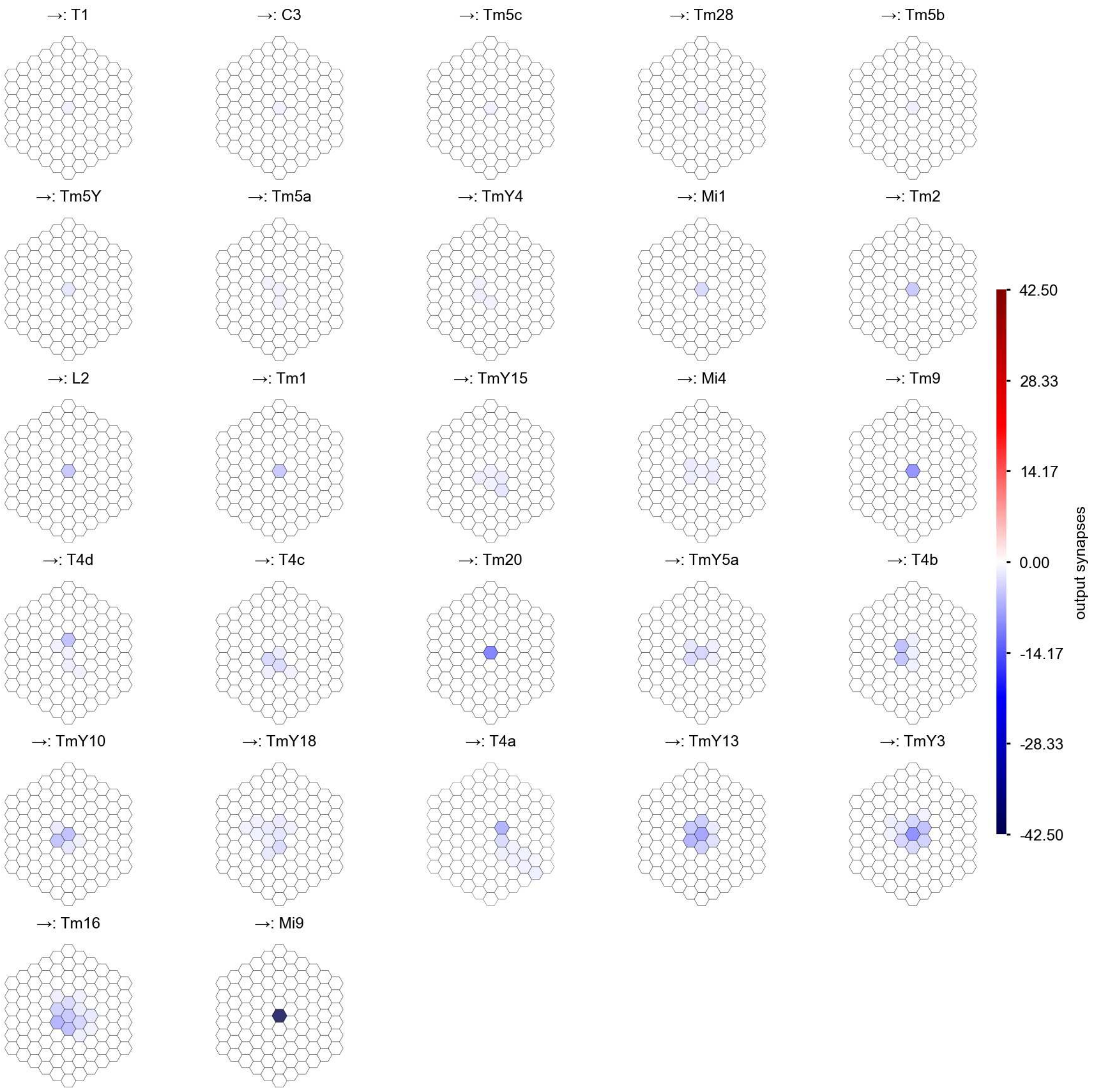

Mi4 - Figure 2: **Anatomical projective fields.** Each colored hexagon is a projection, with the projection strength characterized by the average number of synapses that we count from the EM reconstruction. Red indicates excitatory synapses, blue indicates inhibitory synapses from inferred signs. Filters in the order of their total number of synapses.

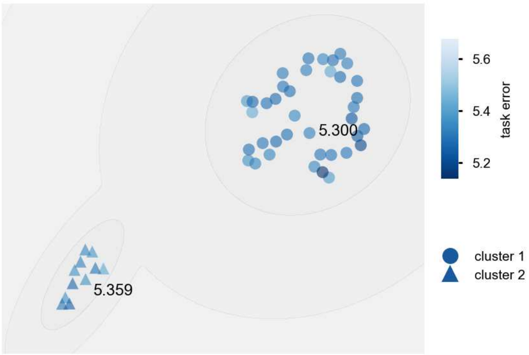

Mi4 - Figure 3: **Clustering of the responses to naturalistic stimuli.** From responses to naturalistic scenes from Sintel. Scatterpoints represent individual models colored by their task error.

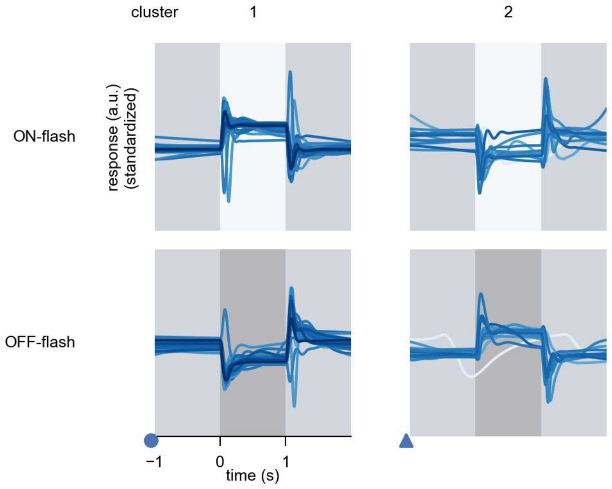

Mi4 - Figure 4: **Responses to flashes.** The top row shows responses to ON-flashes, the bottom row shows responses to OFF-flashes. The responses from the different models in the different clusters (columns) overlay and are colored in the respective task error of the model. The circular flashes occur at second zero, cover 6 ommatidia in radius, and last for one second. Before and after, a grey-stimulus leads to a stationary state of the network. The traces are standardized.

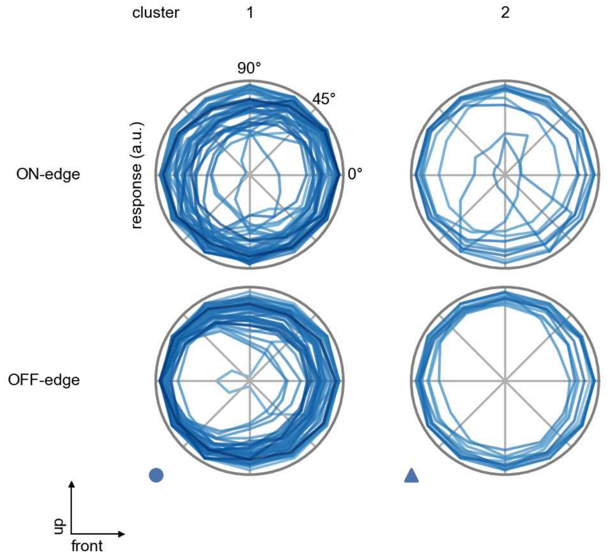

Mi4 - Figure 5: **Peak responses to moving edges.** The top row shows normalized peak responses to moving ON-edges, the bottom row shows normalized peak responses to moving OFF-edges. The edge-stimuli move in different directions from 0 to 360 degrees. Before, a grey-stimulus led the network to a stationary state. The responses from the different models in the different clusters (columns) overlay and are colored in the respective task error of the model. The peak responses are averaged over all assessed edge-speeds.

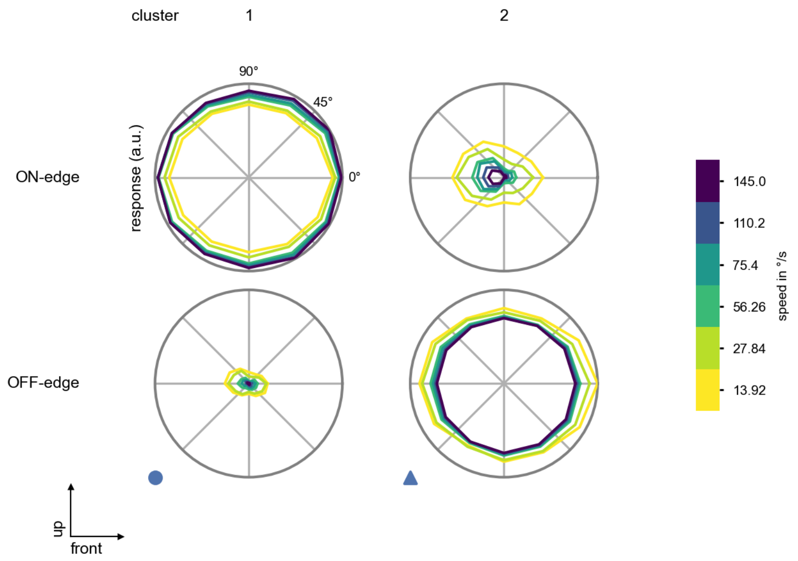

Mi4 - Figure 6: **Peak responses to moving edges from best-performing models.** The top row shows peak responses to moving ON-edges, the bottom row shows peak responses to moving OFF-edges. The edge-stimuli move in different directions from 0 to 360 degrees and at different speeds. Before, a grey-stimulus led the network to a stationary state. The columns show responses from the best-task-performing model in the respective cluster.

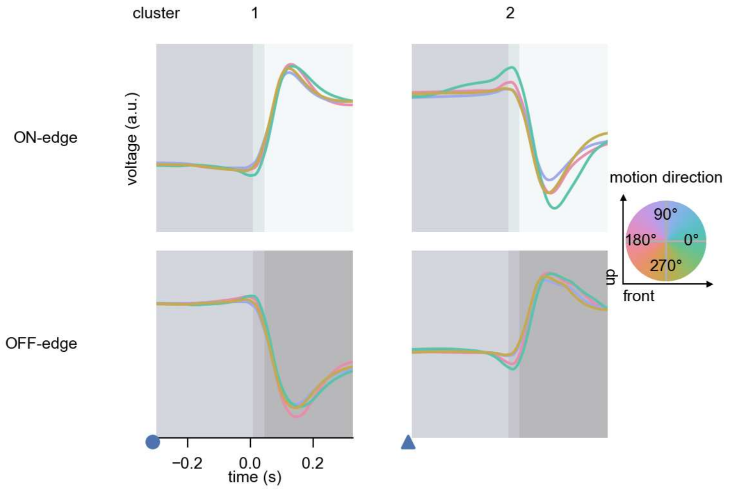

Mi4 - Figure 7: **Responses to moving edges from best-performing models.** The top row shows responses to moving ON-edges, the bottom row shows peak responses to moving OFF-edges. The edge-stimuli move in different directions from 0 to 360 degrees and at different speeds. Before, a grey-stimulus led the network to a stationary state. The columns show responses from the best-task-performing model in the respective cluster. Example responses to edges moving at 75.4°/s in all cardinal directions (green 0°, blue 90°, red 180°, yellow 270°) from -22.5 to 22.5° visual angle. Background contour indicates the edge, passing by the central column at second zero.

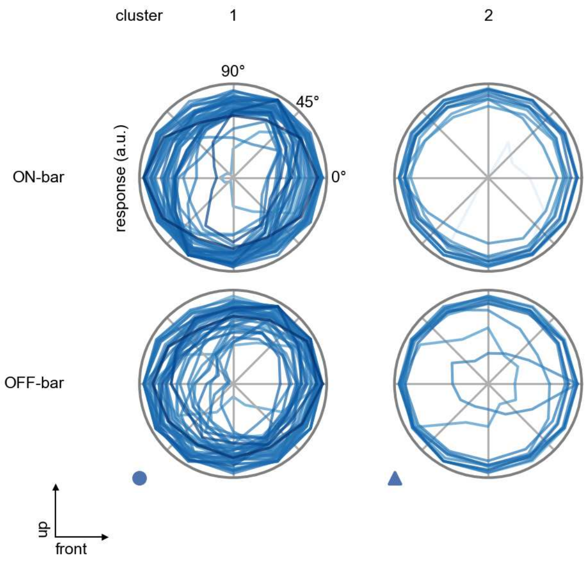

Mi4 - Figure 8: **Peak responses to moving bars.** The top row shows normalized peak responses to moving ON-bars, the bottom row shows normalized peak responses to moving OFF-bars. The bar-stimuli move in different directions from 0 to 360 degrees. Before, a grey-stimulus led the network to a stationary state. The responses from the different models in the different clusters (columns) overlay and are colored in the respective task error of the model. The peak responses are averaged over all assessed bar-speeds.

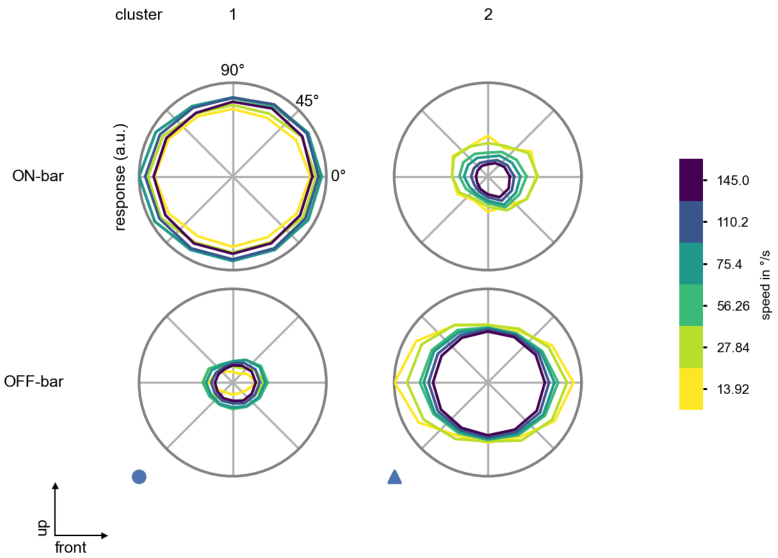

Mi4 - Figure 9: **Peak responses to moving bars from best-performing models.** The top row shows peak responses to moving ON-bars, the bottom row shows peak responses to moving OFF-bars. The bar-stimuli move in different directions from 0 to 360 degrees and at different speeds. Before, a grey-stimulus led the network to a stationary state. The columns show responses from the best-task-performing model in the respective cluster.

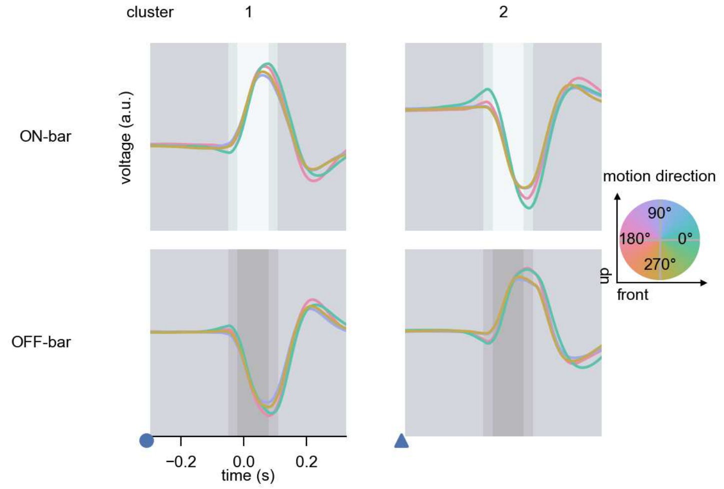

Mi4 - Figure 10: **Responses to moving bars from best-performing models.** The top row shows responses to moving ON-bars, the bottom row shows peak responses to moving OFF-bars. The bar-stimuli move in different directions from 0 to 360 degrees and at different speeds. Before, a grey-stimulus led the network to a stationary state. The columns show responses from the best-task-performing model in the respective cluster. Example responses to bars moving at 75.4°/s in all cardinal directions (green 0°, blue 90°, red 180°, yellow 270°) from -22.5 to 22.5° visual angle. Background contour indicates the bar, passing by the central column at second zero.

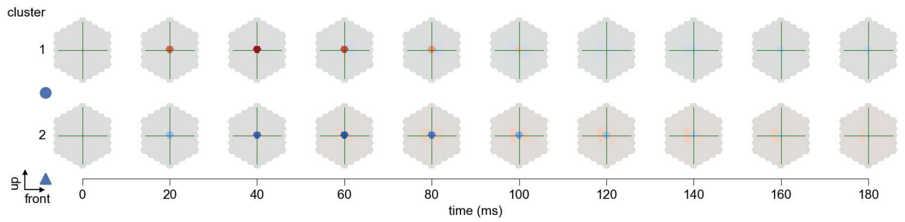

Mi4 - Figure 11: **Spatio-temporal receptive field.** Responses of the central cell to ON-impulses at single ommatidium impulse locations. The impulse occurs at second zero. Before, a grey-stimulus led the network to a stationary state. Responses are baseline-subtracted. Each row illustrates responses from the best-performing model of the respective cluster. Red indicated depolarization, blue indicates hyperpolarization.

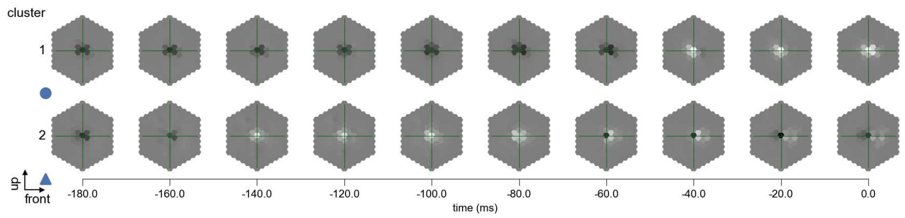

Mi4 - Figure 12: **Maximally excitatory stimuli.** Each row presents the regularized naturalistic-stimulus from the Sintel dataset that maximizes the cell type's central column response at second zero in the best-performing model of the respective cluster.

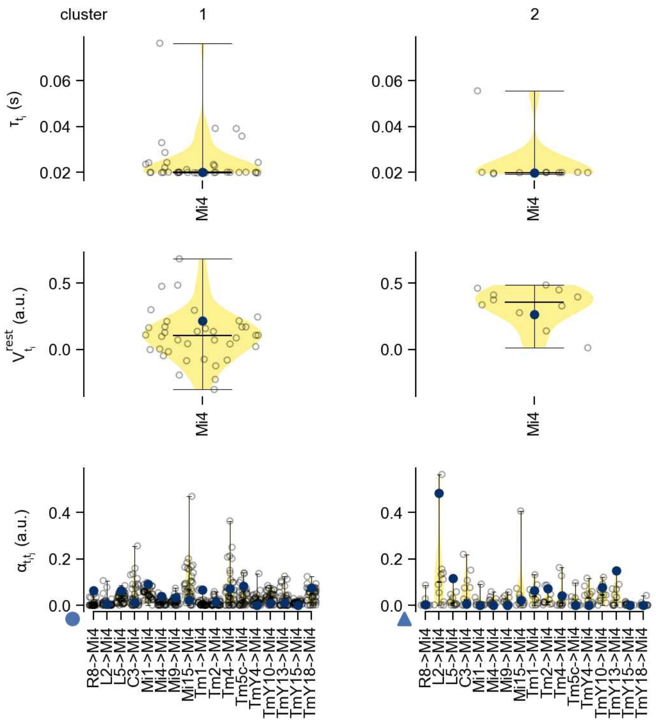

Mi4 - Figure 13: **Task-constrained parameters.** Each column shows the parameters inferred within the respective cluster. First row: learned time constants of the cell type. Second row: resting potentials of the cell type. Third row: scaling factors for the convolutional filters. The blue scatter represents the parameters from the best-performing model within the cluster.

# 25 Mi9

← Cell types

#### Figures

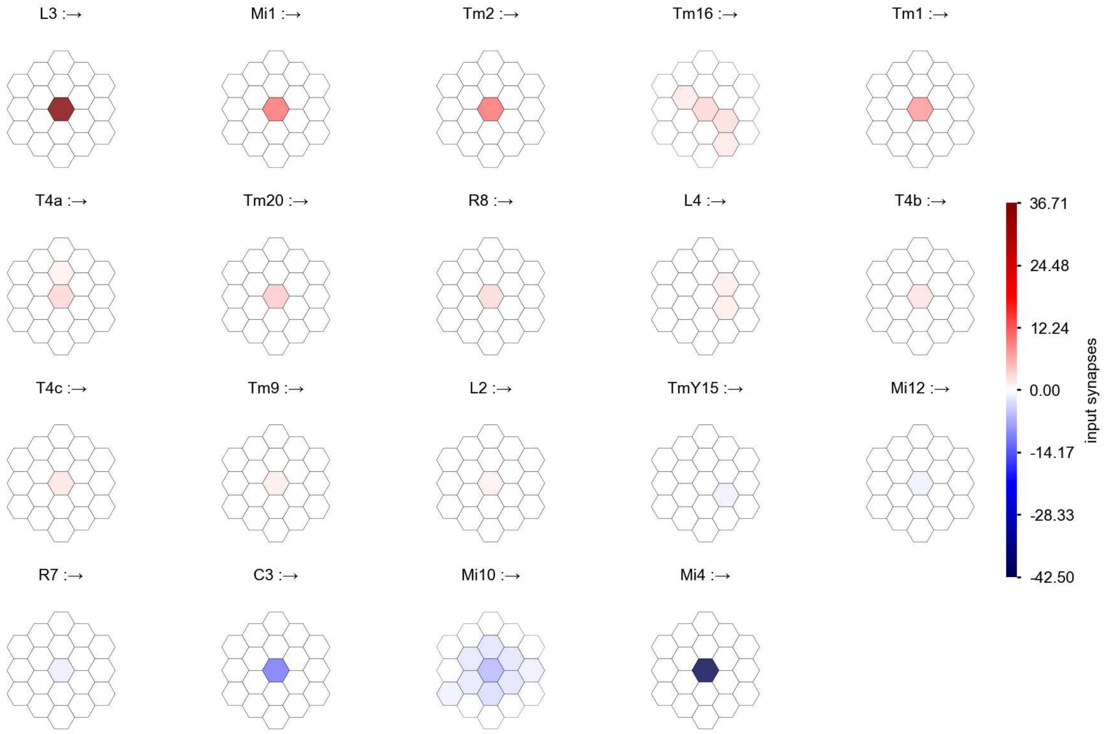

Mi9 - Figure 1: **Anatomical receptive fields.** Each colored hexagon is a projection, with the projection strength characterized by the average number of synapses that we count from the EM reconstruction. Red indicates excitatory synapses, blue indicates inhibitory synapses from inferred signs. Filters in the order of their total number of synapses.

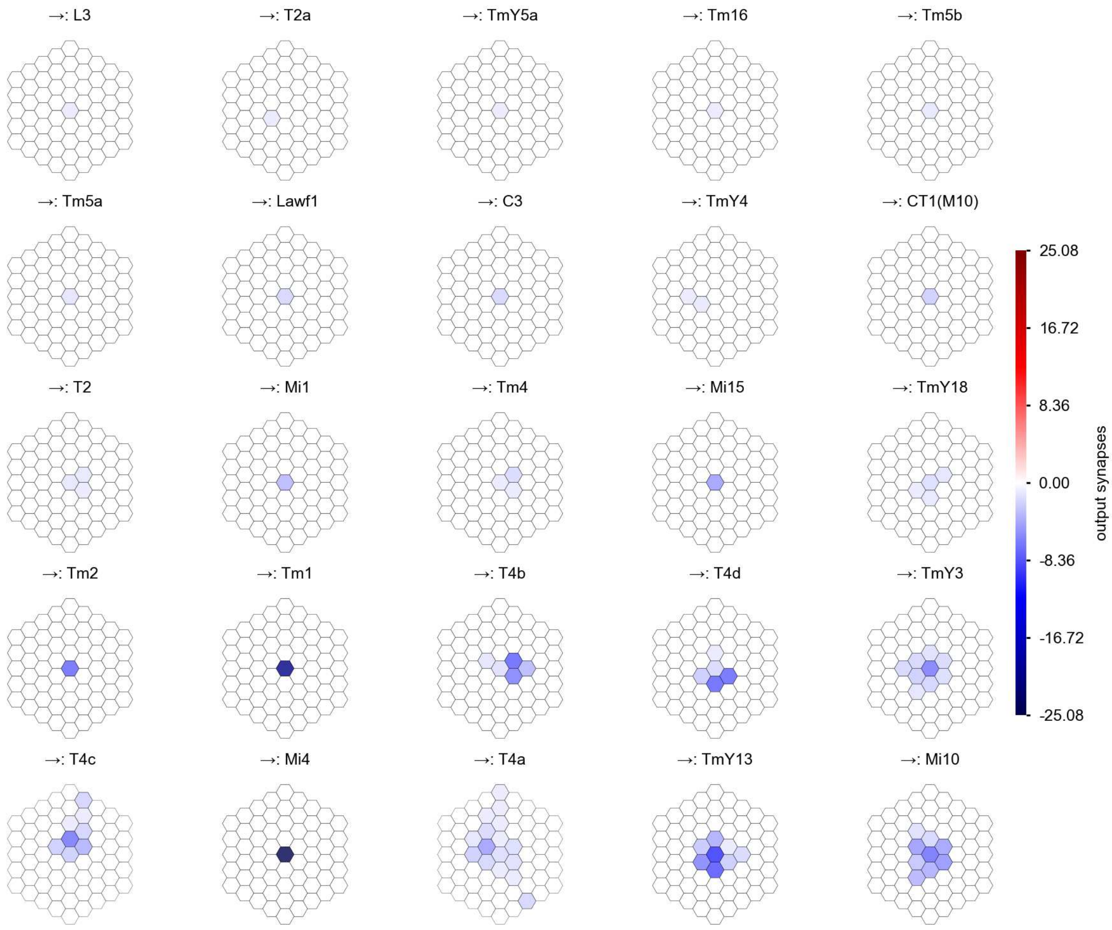

Mi9 - Figure 2: **Anatomical projective fields.** Each colored hexagon is a projection, with the projection strength characterized by the average number of synapses that we count from the EM reconstruction. Red indicates excitatory synapses, blue indicates inhibitory synapses from inferred signs. Filters in the order of their total number of synapses.

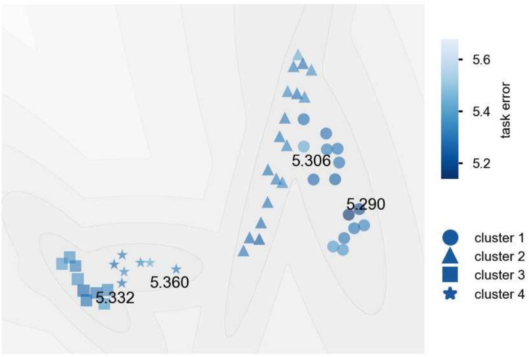

Mi9 - Figure 3: **Clustering of the responses to naturalistic stimuli.** From responses to naturalistic scenes from Sintel. Scatterpoints represent individual models colored by their task error.

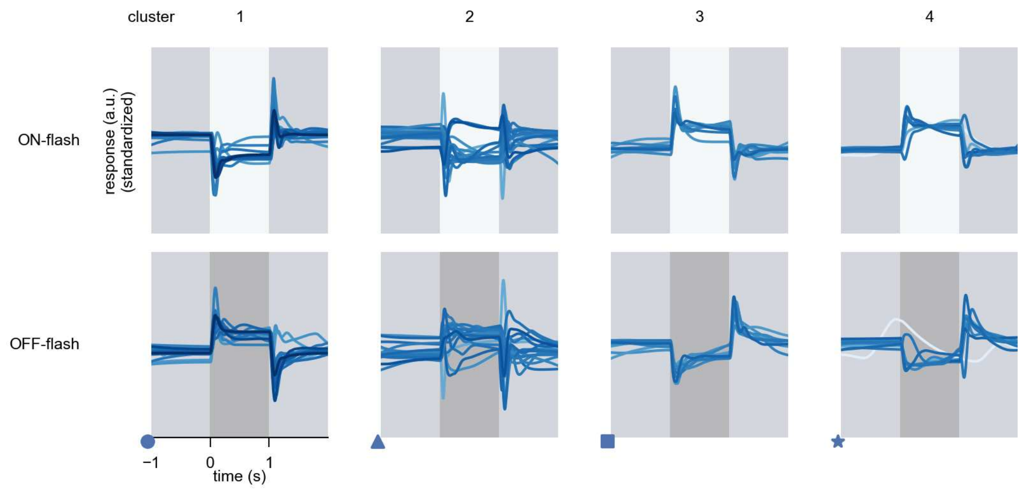

Mi9 - Figure 4: **Responses to flashes.** The top row shows responses to ON-flashes, the bottom row shows responses to OFF-flashes. The responses from the different models in the different clusters (columns) overlay and are colored in the respective task error of the model. The circular flashes occur at second zero, cover 6 ommatidia in radius, and last for one second. Before and after, a grey-stimulus leads to a stationary state of the network. The traces are standardized.

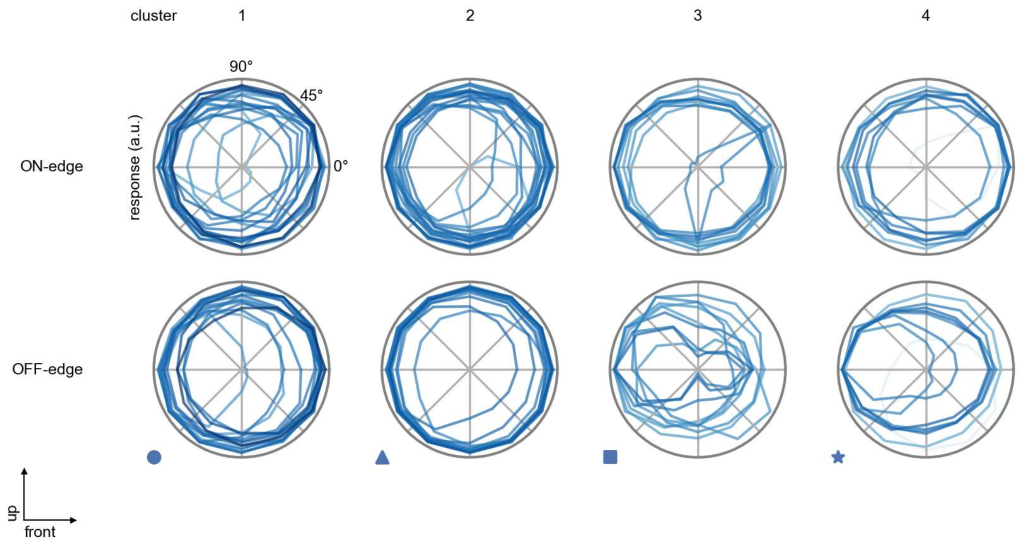

Mi9 - Figure 5: **Peak responses to moving edges.** The top row shows normalized peak responses to moving ON-edges, the bottom row shows normalized peak responses to moving OFF-edges. The edge-stimuli move in different directions from 0 to 360 degrees. Before, a grey-stimulus led the network to a stationary state. The responses from the different models in the different clusters (columns) overlay and are colored in the respective task error of the model. The peak responses are averaged over all assessed edge-speeds.

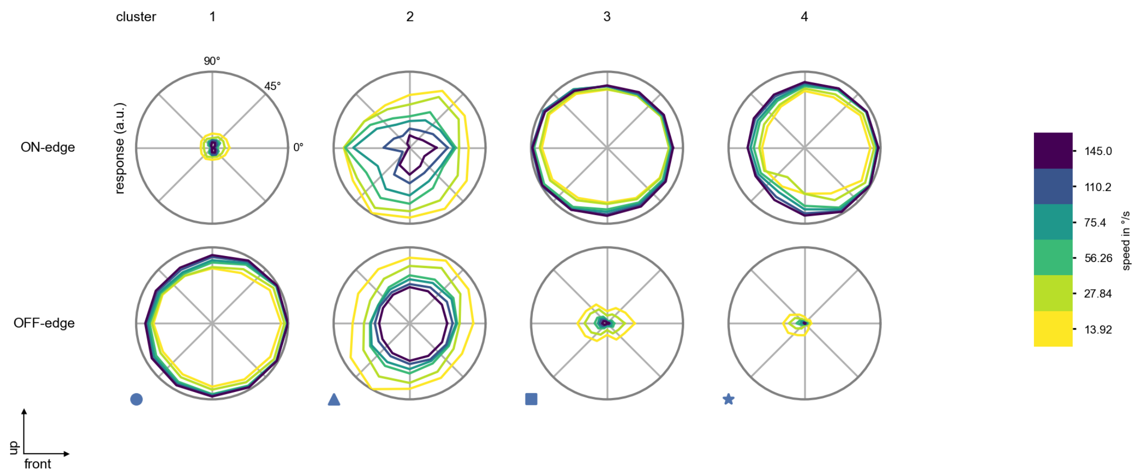

Mi9 - Figure 6: **Peak responses to moving edges from best-performing models.** The top row shows peak responses to moving ON-edges, the bottom row shows peak responses to moving OFF-edges. The edge-stimuli move in different directions from 0 to 360 degrees and at different speeds. Before, a grey-stimulus led the network to a stationary state. The columns show responses from the best-task-performing model in the respective cluster.

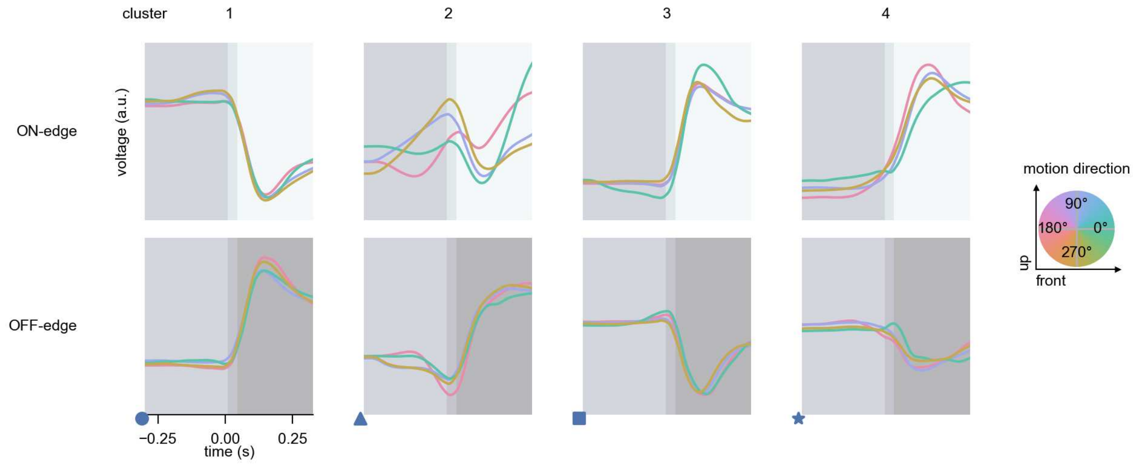

Mi9 - Figure 7: **Responses to moving edges from best-performing models.** The top row shows responses to moving ON-edges, the bottom row shows peak responses to moving OFF-edges. The edge-stimuli move in different directions from 0 to 360 degrees and at different speeds. Before, a grey-stimulus led the network to a stationary state. The columns show responses from the best-task-performing model in the respective cluster. Example responses to edges moving at 75.4°/s in all cardinal directions (green 0°, blue 90°, red 180°, yellow 270°) from -22.5 to 22.5° visual angle. Background contour indicates the edge, passing by the central column at second zero.

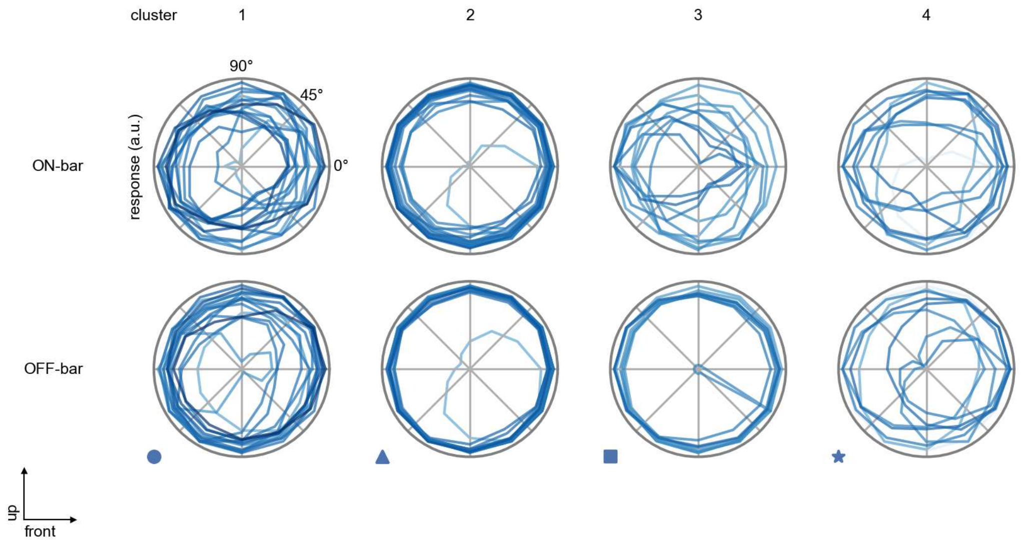

Mi9 - Figure 8: **Peak responses to moving bars.** The top row shows normalized peak responses to moving ON-bars, the bottom row shows normalized peak responses to moving OFF-bars. The bar-stimuli move in different directions from 0 to 360 degrees. Before, a grey-stimulus led the network to a stationary state. The responses from the different models in the different clusters (columns) overlay and are colored in the respective task error of the model. The peak responses are averaged over all assessed bar-speeds.

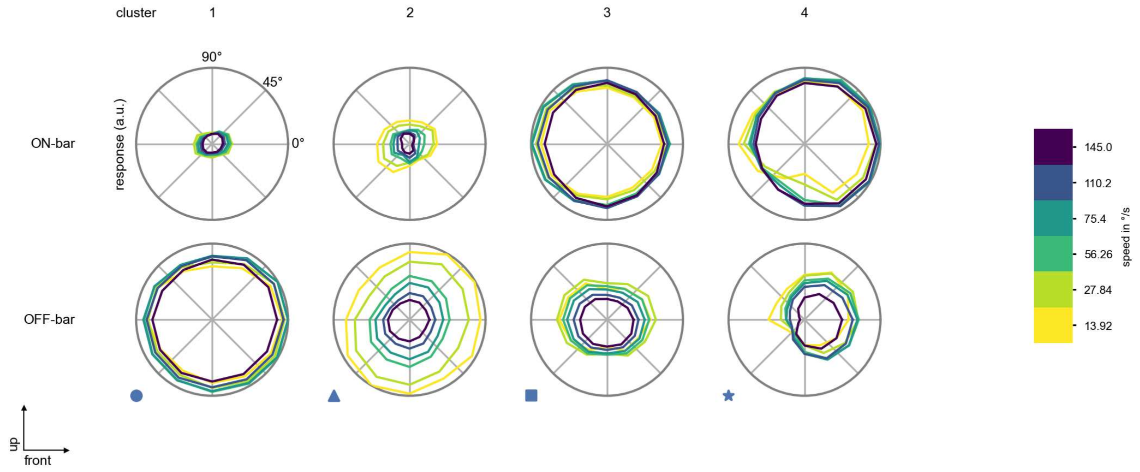

Mi9 - Figure 9: **Peak responses to moving bars from best-performing models.** The top row shows peak responses to moving ON-bars, the bottom row shows peak responses to moving OFF-bars. The bar-stimuli move in different directions from 0 to 360 degrees and at different speeds. Before, a grey-stimulus led the network to a stationary state. The columns show responses from the best-task-performing model in the respective cluster.

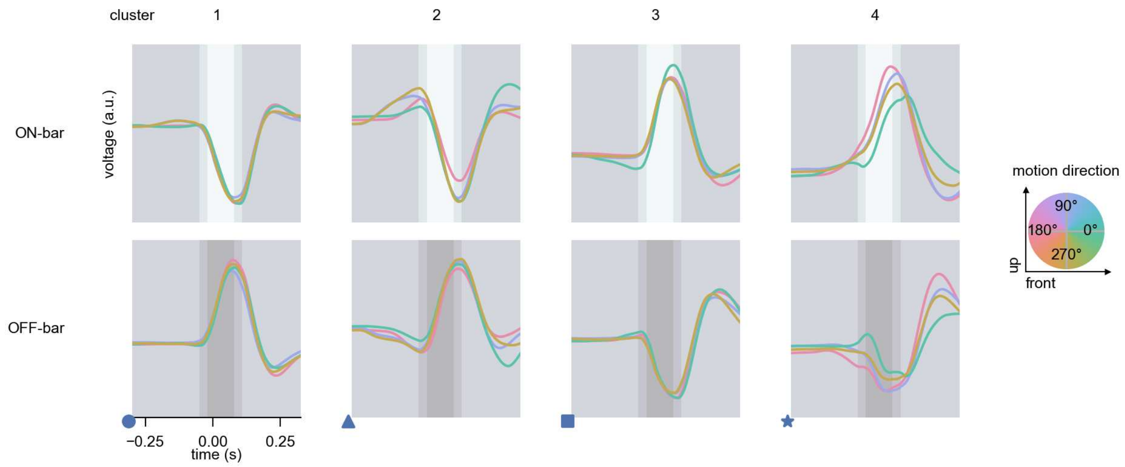

Mi9 - Figure 10: **Responses to moving bars from best-performing models.** The top row shows responses to moving ON-bars, the bottom row shows peak responses to moving OFF-bars. The bar-stimuli move in different directions from 0 to 360 degrees and at different speeds. Before, a grey-stimulus led the network to a stationary state. The columns show responses from the best-task-performing model in the respective cluster. Example responses to bars moving at 75.4°/s in all cardinal directions (green 0°, blue 90°, red 180°, yellow 270°) from -22.5 to 22.5° visual angle. Background contour indicates the bar, passing by the central column at second zero.

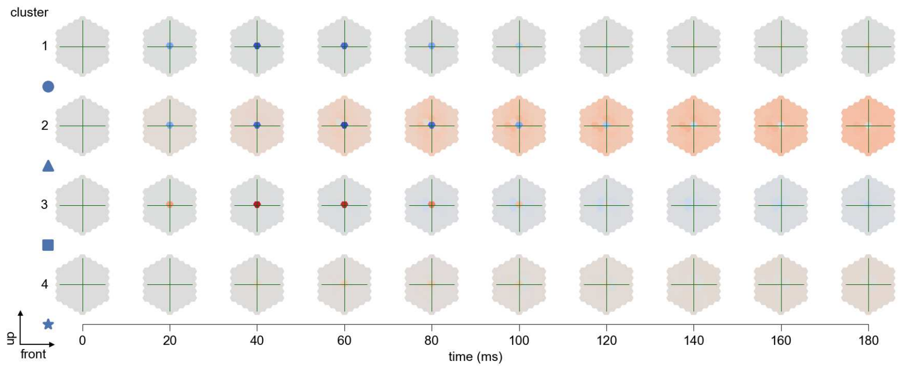

Mi9 - Figure 11: **Spatio-temporal receptive field.** Responses of the central cell to ON-impulses at single ommatidium impulse locations. The impulse occurs at second zero. Before, a grey-stimulus led the network to a stationary state. Responses are baseline-subtracted. Each row illustrates responses from the best-performing model of the respective cluster. Red indicated depolarization, blue indicates hyperpolarization.

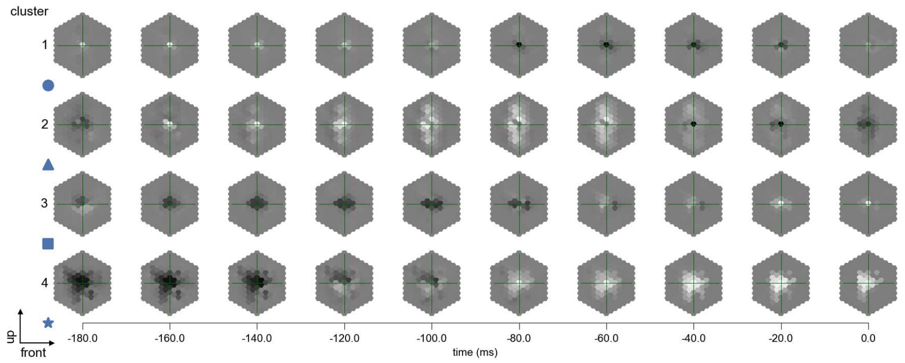

Mi9 - Figure 12: **Maximally excitatory stimuli.** Each row presents the regularized naturalistic-stimulus from the Sintel dataset that maximizes the cell type's central column response at second zero in the best-performing model of the respective cluster.

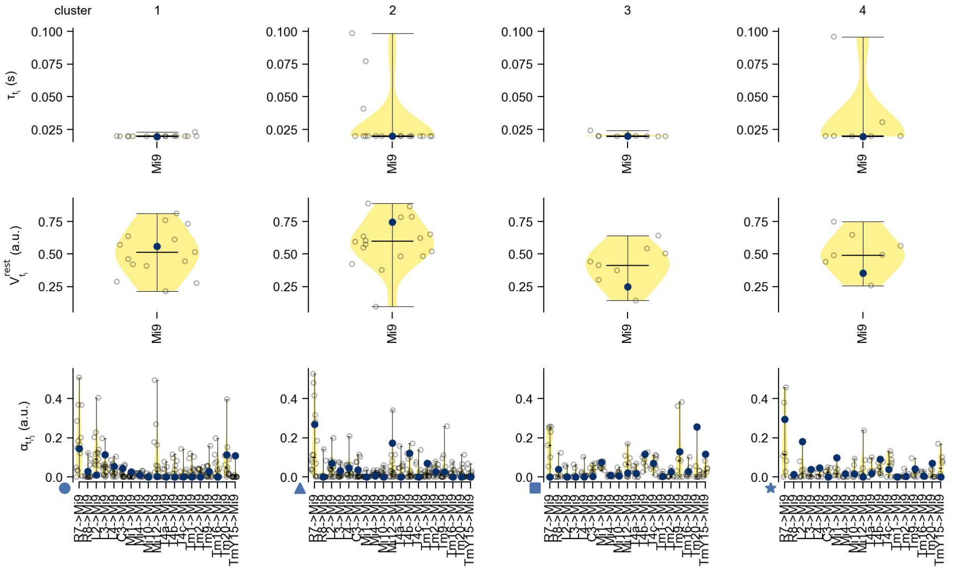

Mi9 - Figure 13: **Task-constrained parameters.** Each column shows the parameters inferred within the respective cluster. First row: learned time constants of the cell type. Second row: resting potentials of the cell type. Third row: scaling factors for the convolutional filters. The blue scatter represents the parameters from the best-performing model within the cluster.

# 26 Mi10

← Cell types

#### Figures

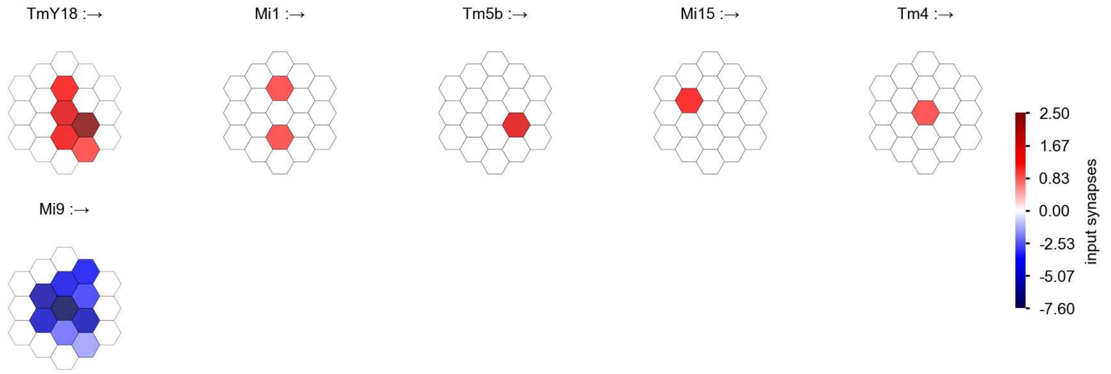

Mi10 - Figure 1: **Anatomical receptive fields.** Each colored hexagon is a projection, with the projection strength characterized by the average number of synapses that we count from the EM reconstruction. Red indicates excitatory synapses, blue indicates inhibitory synapses from inferred signs. Filters in the order of their total number of synapses.

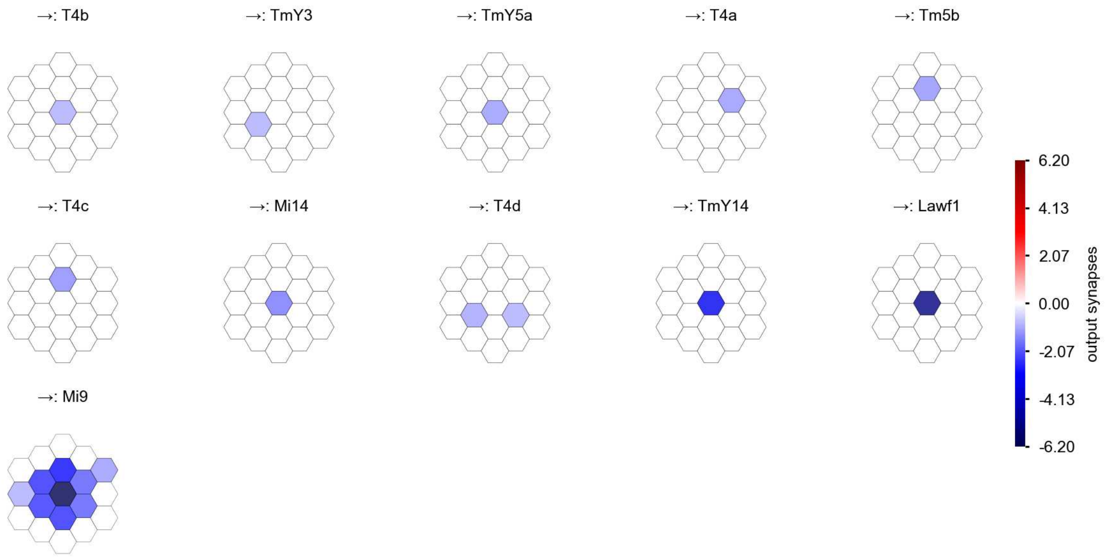

Mi10 - Figure 2: **Anatomical projective fields.** Each colored hexagon is a projection, with the projection strength characterized by the average number of synapses that we count from the EM reconstruction. Red indicates excitatory synapses, blue indicates inhibitory synapses from inferred signs. Filters in the order of their total number of synapses.

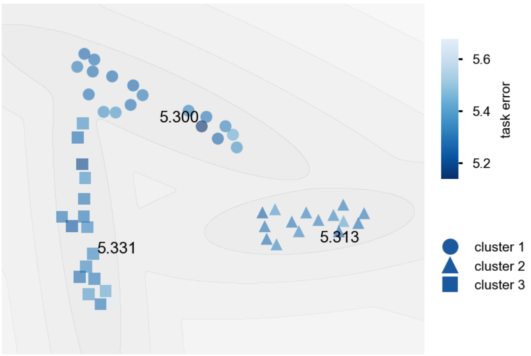

Mi10 - Figure 3: **Clustering of the responses to naturalistic stimuli.** From responses to naturalistic scenes from Sintel. Scatterpoints represent individual models colored by their task error.

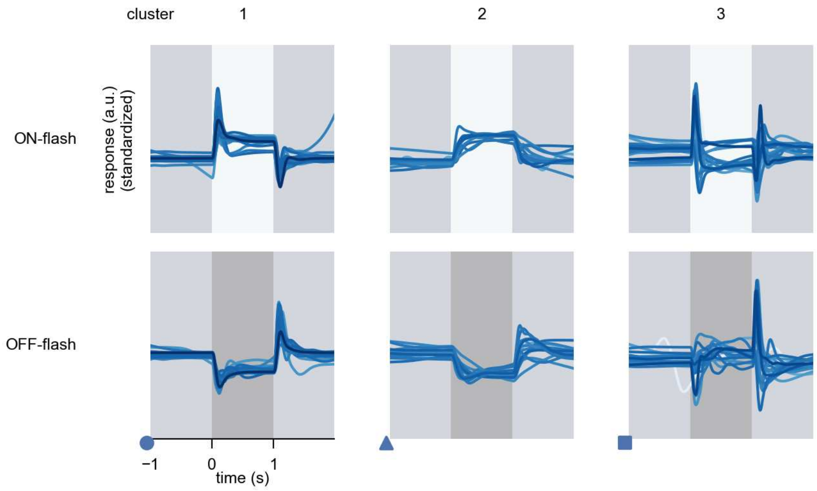

Mi10 - Figure 4: **Responses to flashes.** The top row shows responses to ON-flashes, the bottom row shows responses to OFF-flashes. The responses from the different models in the different clusters (columns) overlay and are colored in the respective task error of the model. The circular flashes occur at second zero, cover 6 ommatidia in radius, and last for one second. Before and after, a grey-stimulus leads to a stationary state of the network. The traces are standardized.

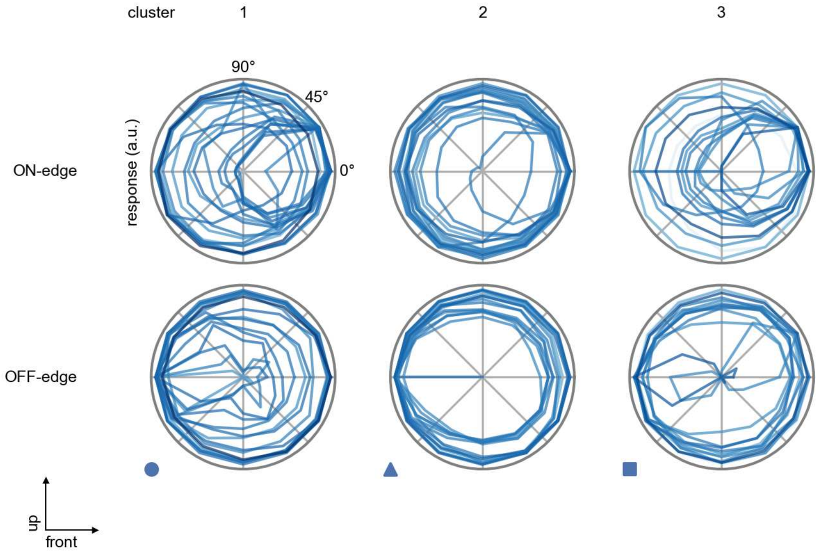

Mi10 - Figure 5: **Peak responses to moving edges.** The top row shows normalized peak responses to moving ON-edges, the bottom row shows normalized peak responses to moving OFF-edges. The edge-stimuli move in different directions from 0 to 360 degrees. Before, a grey-stimulus led the network to a stationary state. The responses from the different models in the different clusters (columns) overlay and are colored in the respective task error of the model. The peak responses are averaged over all assessed edge-speeds.

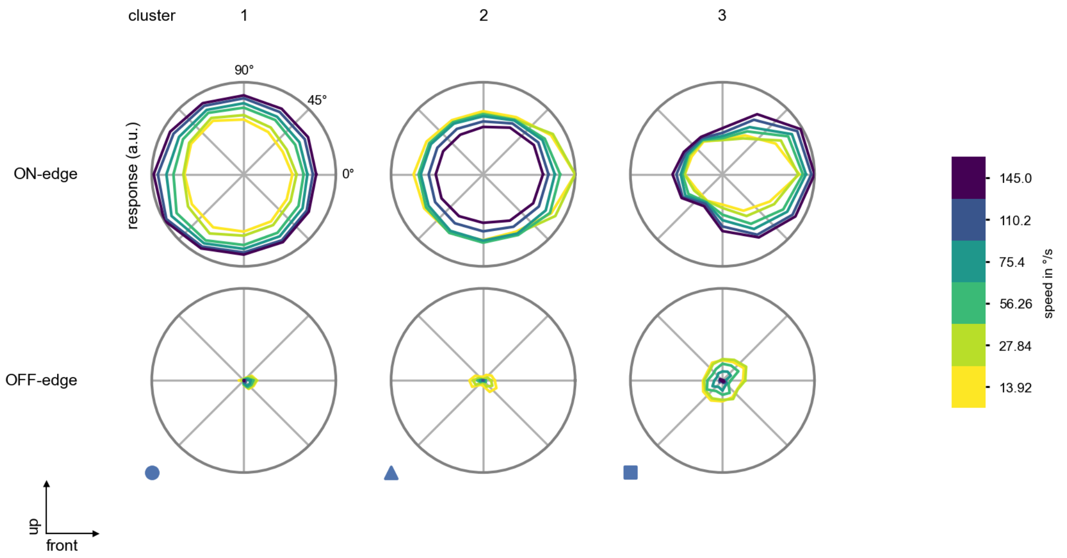

Mi10 - Figure 6: **Peak responses to moving edges from best-performing models.** The top row shows peak responses to moving ON-edges, the bottom row shows peak responses to moving OFF-edges. The edge-stimuli move in different directions from 0 to 360 degrees and at different speeds. Before, a grey-stimulus led the network to a stationary state. The columns show responses from the best-task-performing model in the respective cluster.

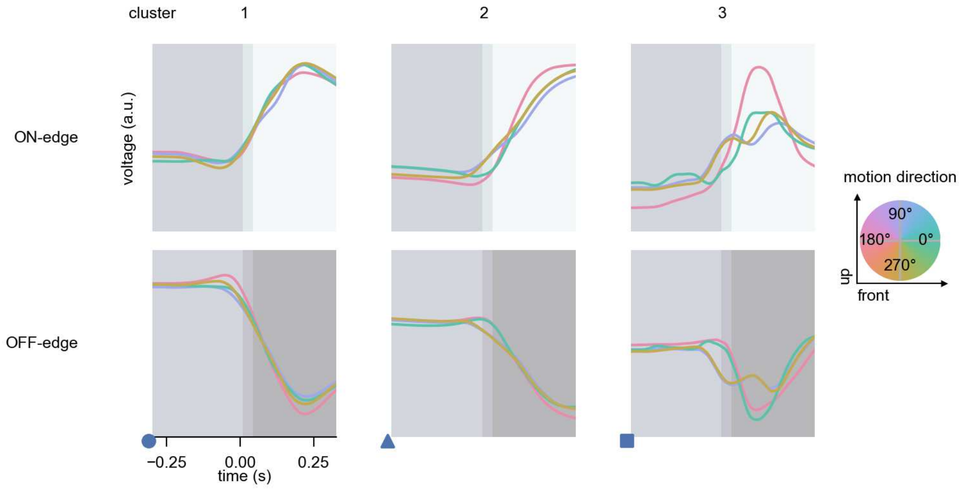

Mi10 - Figure 7: **Responses to moving edges from best-performing models.** The top row shows responses to moving ON-edges, the bottom row shows peak responses to moving OFF-edges. The edge-stimuli move in different directions from 0 to 360 degrees and at different speeds. Before, a grey-stimulus led the network to a stationary state. The columns show responses from the best-task-performing model in the respective cluster. Example responses to edges moving at 75.4°/s in all cardinal directions (green 0°, blue 90°, red 180°, yellow 270°) from -22.5 to 22.5° visual angle. Background contour indicates the edge, passing by the central column at second zero.

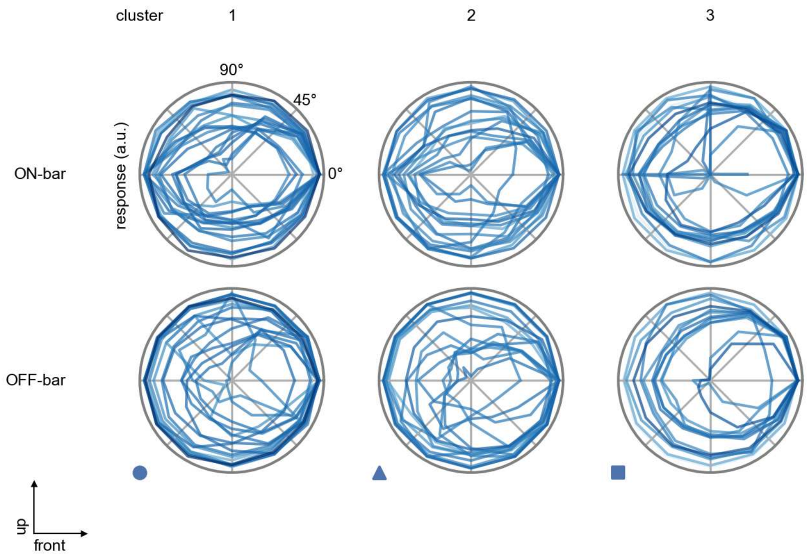

Mi10 - Figure 8: **Peak responses to moving bars.** The top row shows normalized peak responses to moving ON-bars, the bottom row shows normalized peak responses to moving OFF-bars. The bar-stimuli move in different directions from 0 to 360 degrees. Before, a grey-stimulus led the network to a stationary state. The responses from the different models in the different clusters (columns) overlay and are colored in the respective task error of the model. The peak responses are averaged over all assessed bar-speeds.

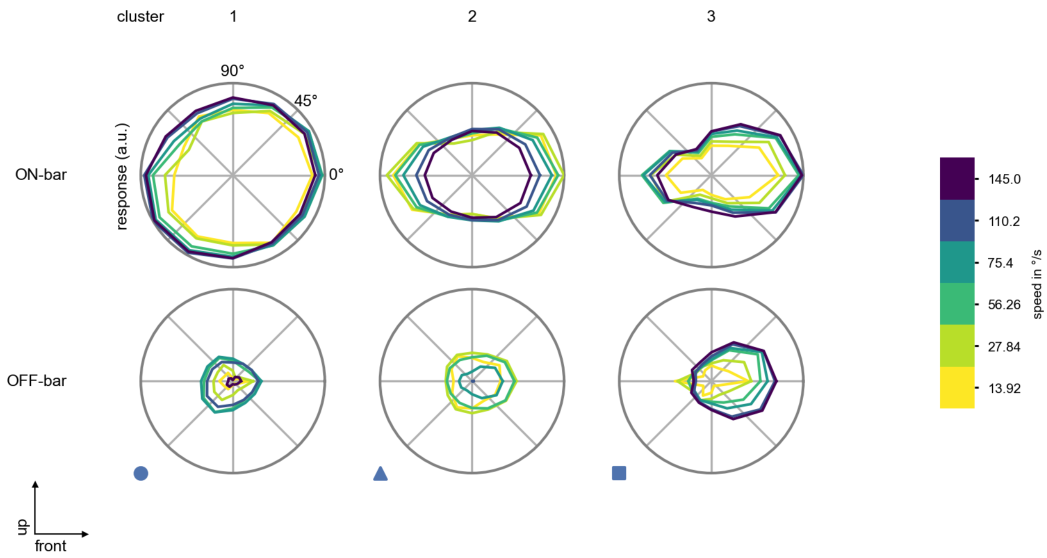

Mi10 - Figure 9: **Peak responses to moving bars from best-performing models.** The top row shows peak responses to moving ON-bars, the bottom row shows peak responses to moving OFF-bars. The bar-stimuli move in different directions from 0 to 360 degrees and at different speeds. Before, a grey-stimulus led the network to a stationary state. The columns show responses from the best-task-performing model in the respective cluster.

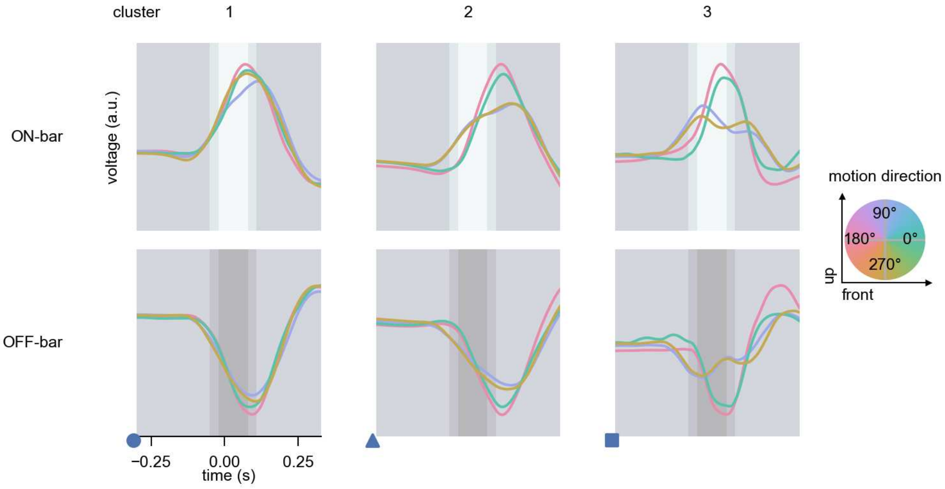

Mi10 - Figure 10: **Responses to moving bars from best-performing models.** The top row shows responses to moving ON-bars, the bottom row shows peak responses to moving OFF-bars. The bar-stimuli move in different directions from 0 to 360 degrees and at different speeds. Before, a grey-stimulus led the network to a stationary state. The columns show responses from the best-task-performing model in the respective cluster. Example responses to bars moving at 75.4°/s in all cardinal directions (green 0°, blue 90°, red 180°, yellow 270°) from -22.5 to 22.5° visual angle. Background contour indicates the bar, passing by the central column at second zero.

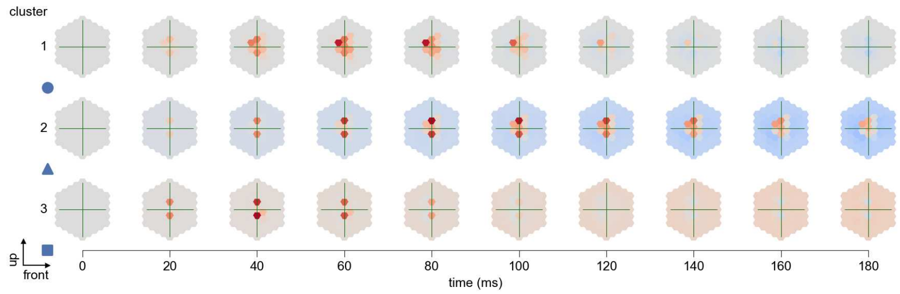

Mi10 - Figure 11: **Spatio-temporal receptive field.** Responses of the central cell to ON-impulses at single ommatidium impulse locations. The impulse occurs at second zero. Before, a grey-stimulus led the network to a stationary state. Responses are baseline-subtracted. Each row illustrates responses from the best-performing model of the respective cluster. Red indicated depolarization, blue indicates hyperpolarization.

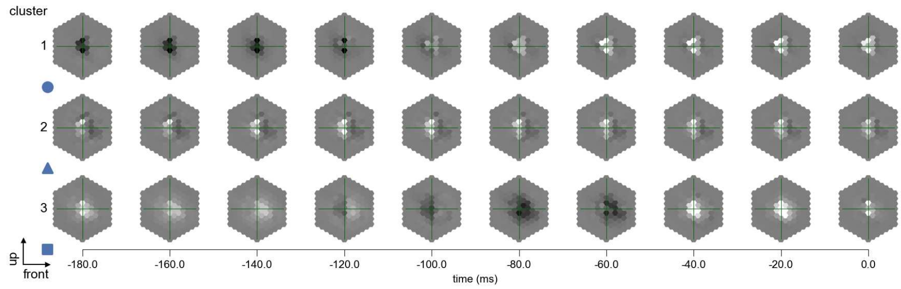

Mi10 - Figure 12: **Maximally excitatory stimuli.** Each row presents the regularized naturalistic-stimulus from the Sintel dataset that maximizes the cell type's central column response at second zero in the best-performing model of the respective cluster.

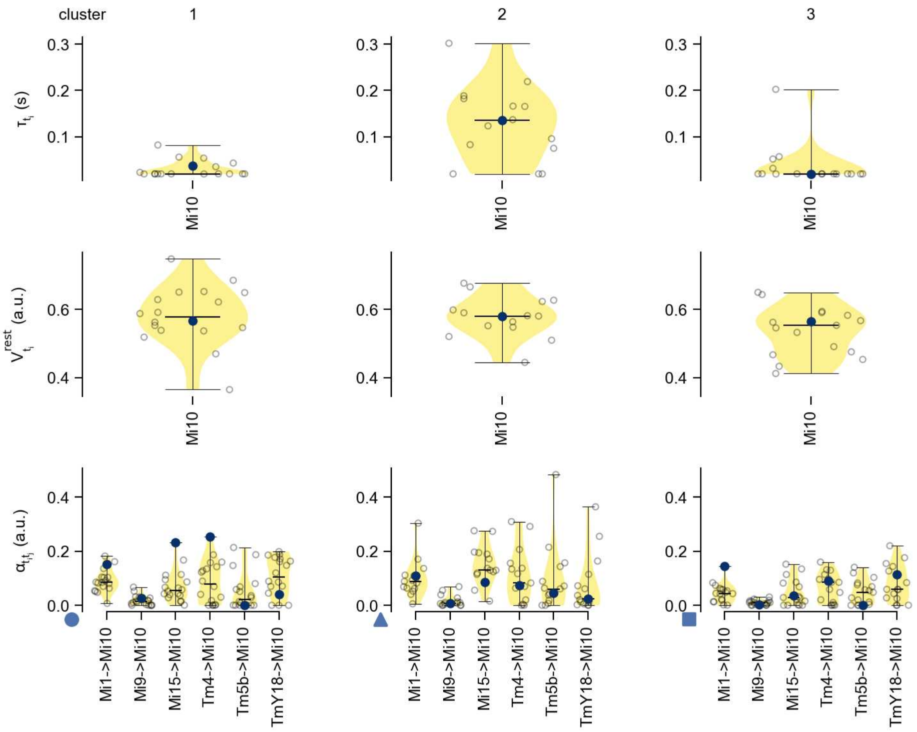

Mi10 - Figure 13: **Task-constrained parameters.** Each column shows the parameters inferred within the respective cluster. First row: learned time constants of the cell type. Second row: resting potentials of the cell type. Third row: scaling factors for the convolutional filters. The blue scatter represents the parameters from the best-performing model within the cluster.

# 27 Mi11

← Cell types

#### Figures

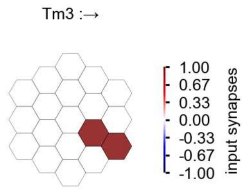

Mi11 - Figure 1: **Anatomical receptive fields.** Each colored hexagon is a projection, with the projection strength characterized by the average number of synapses that we count from the EM reconstruction. Red indicates excitatory synapses, blue indicates inhibitory synapses from inferred signs. Filters in the order of their total number of synapses.

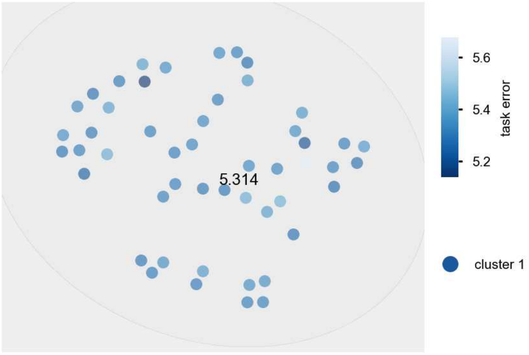

Mi11 - Figure 2: **Clustering of the responses to naturalistic stimuli.** From responses to naturalistic scenes from Sintel. Scatterpoints represent individual models colored by their task error.

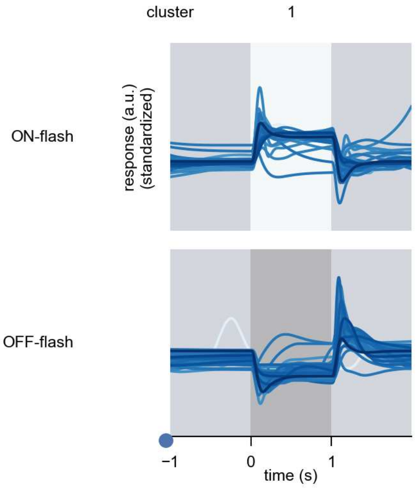

Mi11 - Figure 3: **Responses to flashes.** The top row shows responses to ON-flashes, the bottom row shows responses to OFF-flashes. The responses from the different models in the different clusters (columns) overlay and are colored in the respective task error of the model. The circular flashes occur at second zero, cover 6 ommatidia in radius, and last for one second. Before and after, a grey-stimulus leads to a stationary state of the network. The traces are standardized.

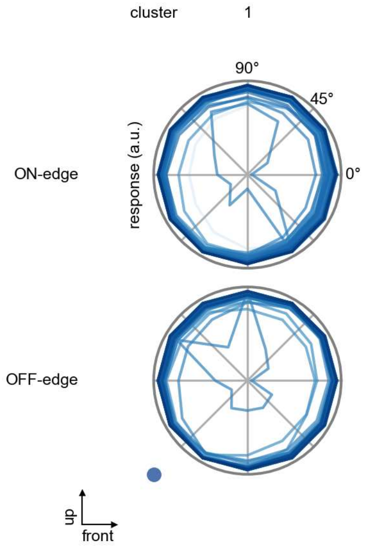

Mi11 - Figure 4: **Peak responses to moving edges.** The top row shows normalized peak responses to moving ON-edges, the bottom row shows normalized peak responses to moving OFF-edges. The edge-stimuli move in different directions from 0 to 360 degrees. Before, a grey-stimulus led the network to a stationary state. The responses from the different models in the different clusters (columns) overlay and are colored in the respective task error of the model. The peak responses are averaged over all assessed edge-speeds.

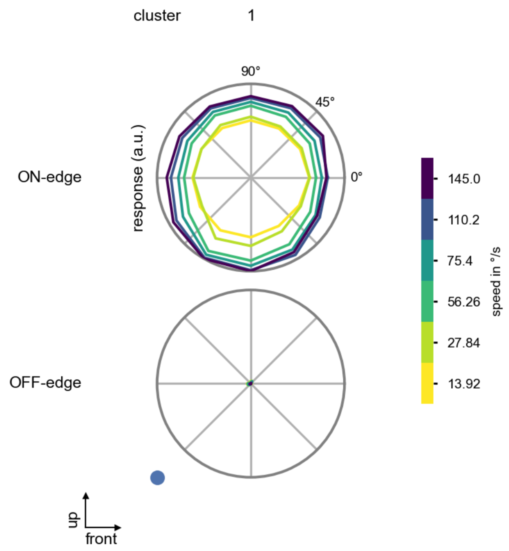

Mi11 - Figure 5: **Peak responses to moving edges from best-performing models.** The top row shows peak responses to moving ON-edges, the bottom row shows peak responses to moving OFF-edges. The edge-stimuli move in different directions from 0 to 360 degrees and at different speeds. Before, a grey-stimulus led the network to a stationary state. The columns show responses from the best-task-performing model in the respective cluster.

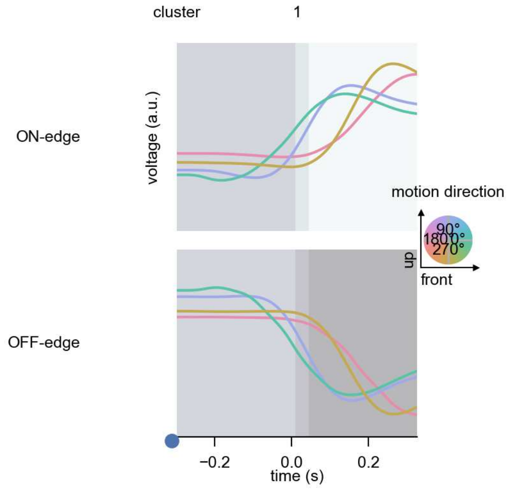

Mi11 - Figure 6: **Responses to moving edges from best-performing models.** The top row shows responses to moving ON-edges, the bottom row shows peak responses to moving OFF-edges. The edge-stimuli move in different directions from 0 to 360 degrees and at different speeds. Before, a grey-stimulus led the network to a stationary state. The columns show responses from the best-task-performing model in the respective cluster. Example responses to edges moving at 75.4°/s in all cardinal directions (green 0°, blue 90°, red 180°, yellow 270°) from -22.5 to 22.5° visual angle. Background contour indicates the edge, passing by the central column at second zero.

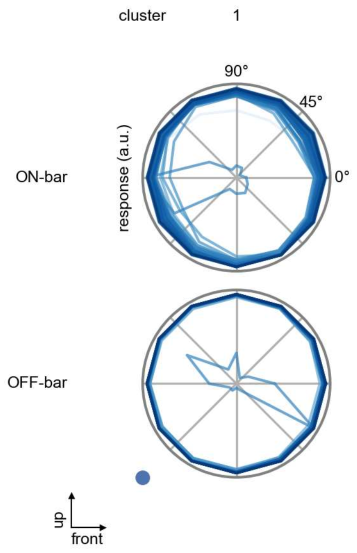

Mi11 - Figure 7: **Peak responses to moving bars.** The top row shows normalized peak responses to moving ON-bars, the bottom row shows normalized peak responses to moving OFF-bars. The bar-stimuli move in different directions from 0 to 360 degrees. Before, a grey-stimulus led the network to a stationary state. The responses from the different models in the different clusters (columns) overlay and are colored in the respective task error of the model. The peak responses are averaged over all assessed bar-speeds.

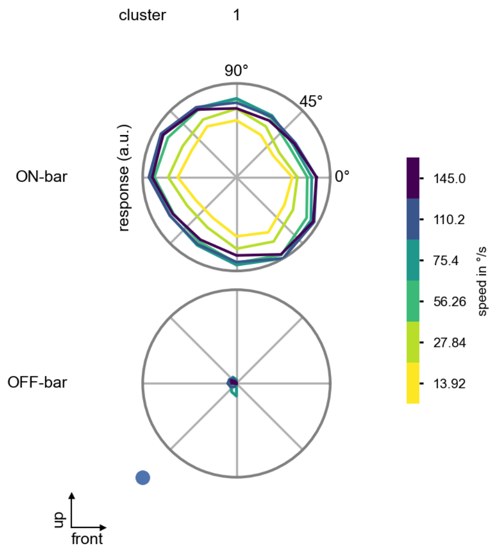

Mi11 - Figure 8: **Peak responses to moving bars from best-performing models.** The top row shows peak responses to moving ON-bars, the bottom row shows peak responses to moving OFF-bars. The bar-stimuli move in different directions from 0 to 360 degrees and at different speeds. Before, a grey-stimulus led the network to a stationary state. The columns show responses from the best-task-performing model in the respective cluster.

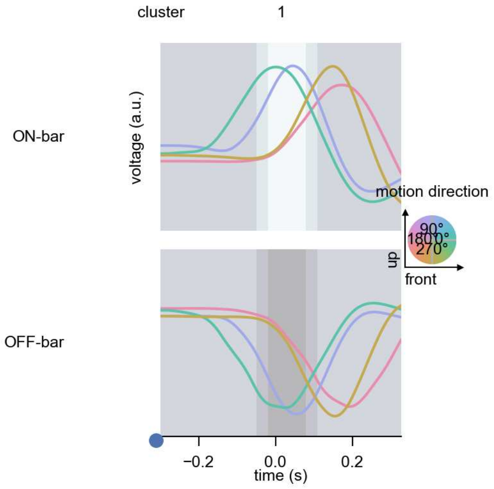

Mi11 - Figure 9: **Responses to moving bars from best-performing models.** The top row shows responses to moving ON-bars, the bottom row shows peak responses to moving OFF-bars. The bar-stimuli move in different directions from 0 to 360 degrees and at different speeds. Before, a grey-stimulus led the network to a stationary state. The columns show responses from the best-task-performing model in the respective cluster. Example responses to bars moving at 75.4°/s in all cardinal directions (green 0°, blue 90°, red 180°, yellow 270°) from -22.5 to 22.5° visual angle. Background contour indicates the bar, passing by the central column at second zero.

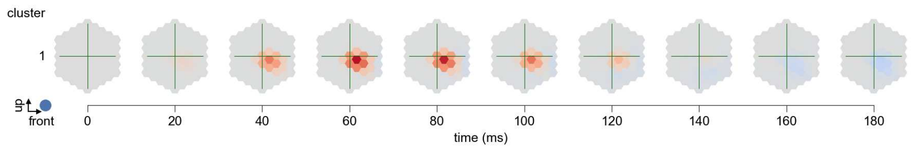

Mi11 - Figure 10: **Spatio-temporal receptive field.** Responses of the central cell to ON-impulses at single ommatidium impulse locations. The impulse occurs at second zero. Before, a grey-stimulus led the network to a stationary state. Responses are baseline-subtracted. Each row illustrates responses from the best-performing model of the respective cluster. Red indicated depolarization, blue indicates hyperpolarization.

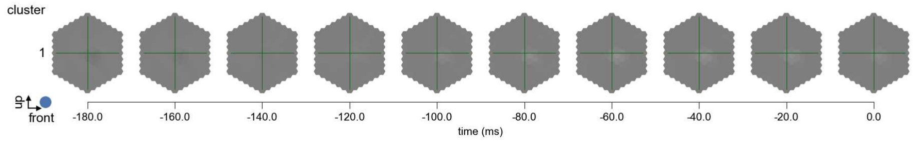

Mi11 - Figure 11: **Maximally excitatory stimuli.** Each row presents the regularized naturalistic-stimulus from the Sintel dataset that maximizes the cell type's central column response at second zero in the best-performing model of the respective cluster.

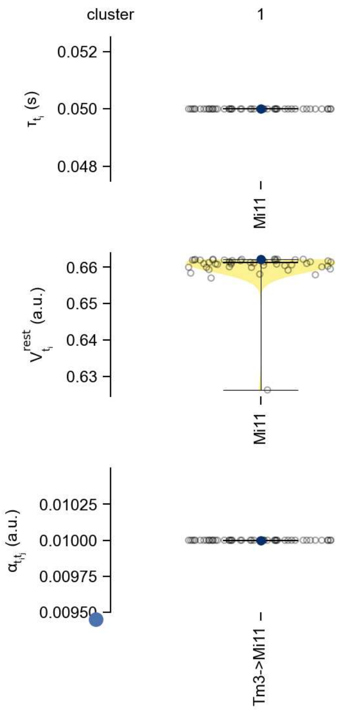

Mi11 - Figure 12: **Task-constrained parameters.** Each column shows the parameters inferred within the respective cluster. First row: learned time constants of the cell type. Second row: resting potentials of the cell type. Third row: scaling factors for the convolutional filters. The blue scatter represents the parameters from the best-performing model within the cluster.

# 28 Mi12

← Cell types

#### Figures

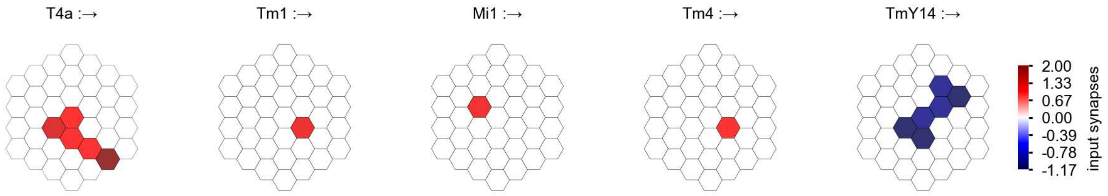

Mi12 - Figure 1: **Anatomical receptive fields.** Each colored hexagon is a projection, with the projection strength characterized by the average number of synapses that we count from the EM reconstruction. Red indicates excitatory synapses, blue indicates inhibitory synapses from inferred signs. Filters in the order of their total number of synapses.

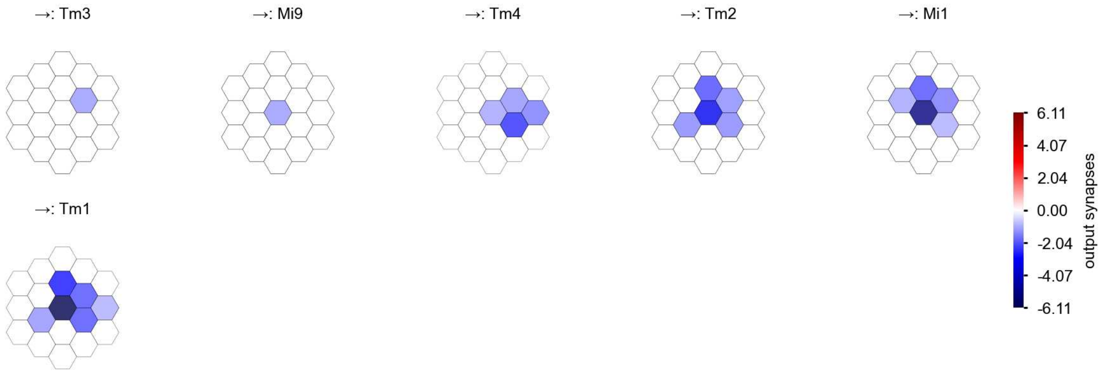

Mi12 - Figure 2: **Anatomical projective fields.** Each colored hexagon is a projection, with the projection strength characterized by the average number of synapses that we count from the EM reconstruction. Red indicates excitatory synapses, blue indicates inhibitory synapses from inferred signs. Filters in the order of their total number of synapses.

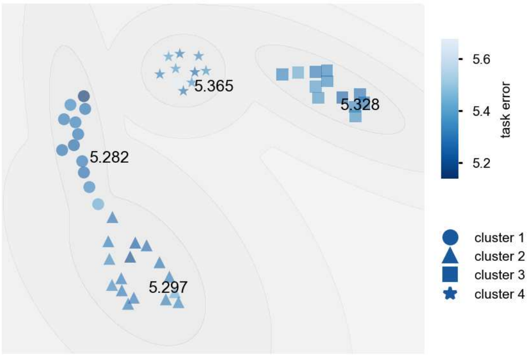

Mi12 - Figure 3: **Clustering of the responses to naturalistic stimuli.** From responses to naturalistic scenes from Sintel. Scatterpoints represent individual models colored by their task error.

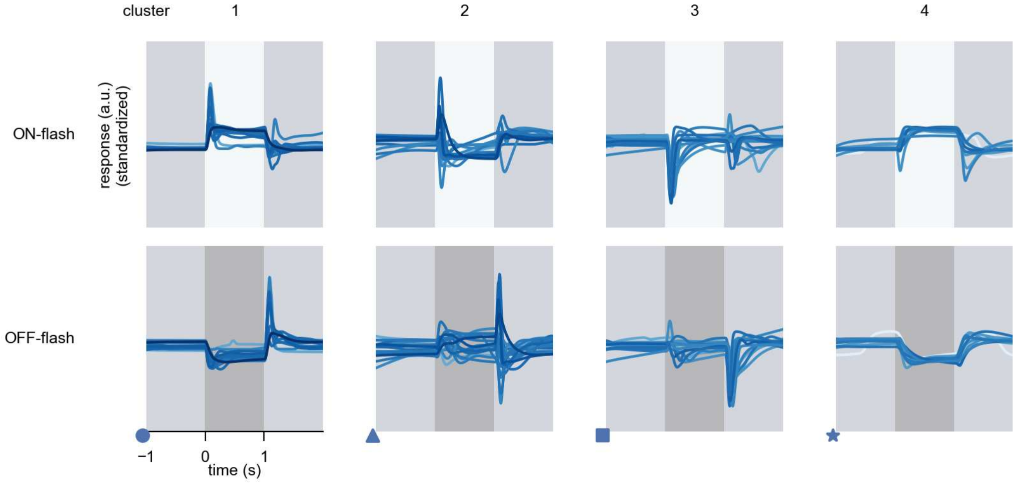

Mi12 - Figure 4: **Responses to flashes.** The top row shows responses to ON-flashes, the bottom row shows responses to OFF-flashes. The responses from the different models in the different clusters (columns) overlay and are colored in the respective task error of the model. The circular flashes occur at second zero, cover 6 ommatidia in radius, and last for one second. Before and after, a grey-stimulus leads to a stationary state of the network. The traces are standardized.

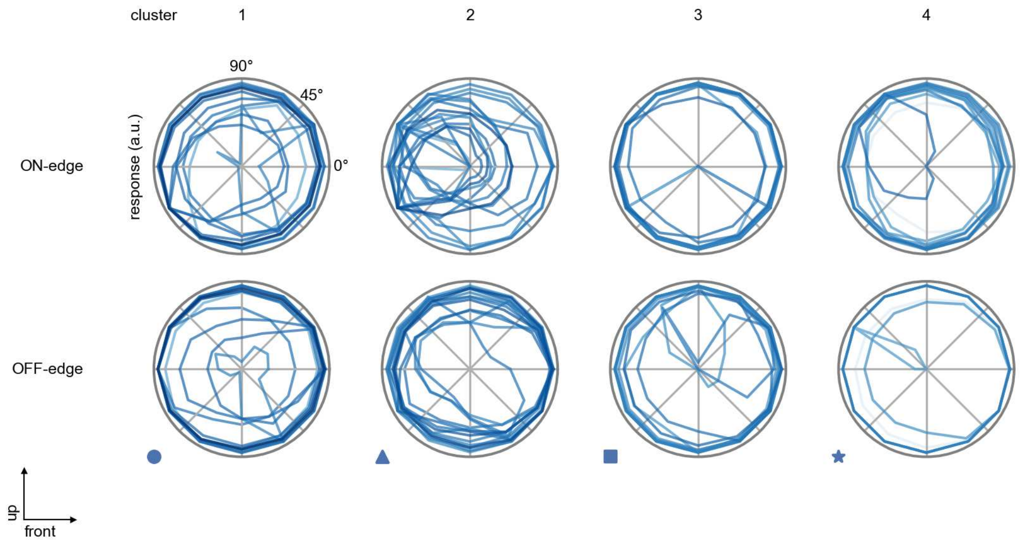

Mi12 - Figure 5: **Peak responses to moving edges.** The top row shows normalized peak responses to moving ON-edges, the bottom row shows normalized peak responses to moving OFF-edges. The edge-stimuli move in different directions from 0 to 360 degrees. Before, a grey-stimulus led the network to a stationary state. The responses from the different models in the different clusters (columns) overlay and are colored in the respective task error of the model. The peak responses are averaged over all assessed edge-speeds.

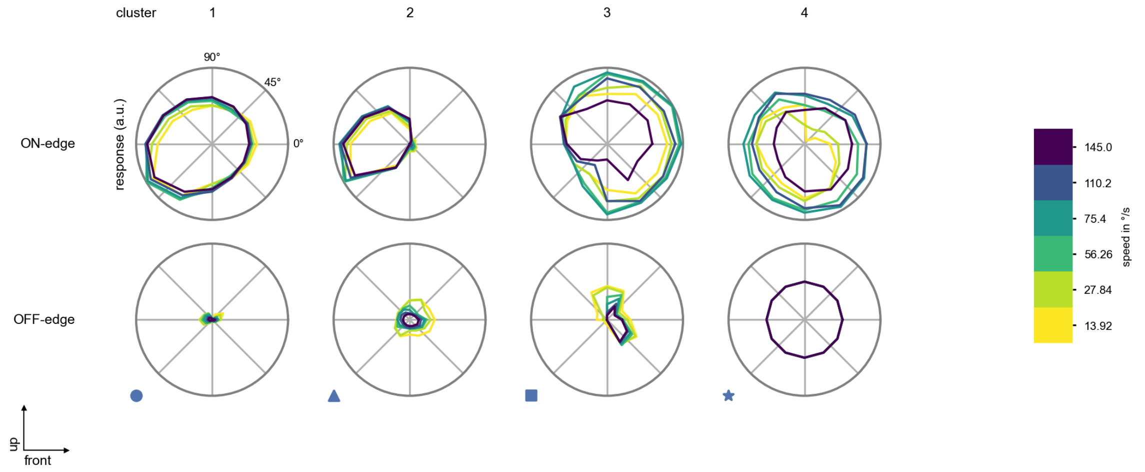

Mi12 - Figure 6: **Peak responses to moving edges from best-performing models.** The top row shows peak responses to moving ON-edges, the bottom row shows peak responses to moving OFF-edges. The edge-stimuli move in different directions from 0 to 360 degrees and at different speeds. Before, a grey-stimulus led the network to a stationary state. The columns show responses from the best-task-performing model in the respective cluster.

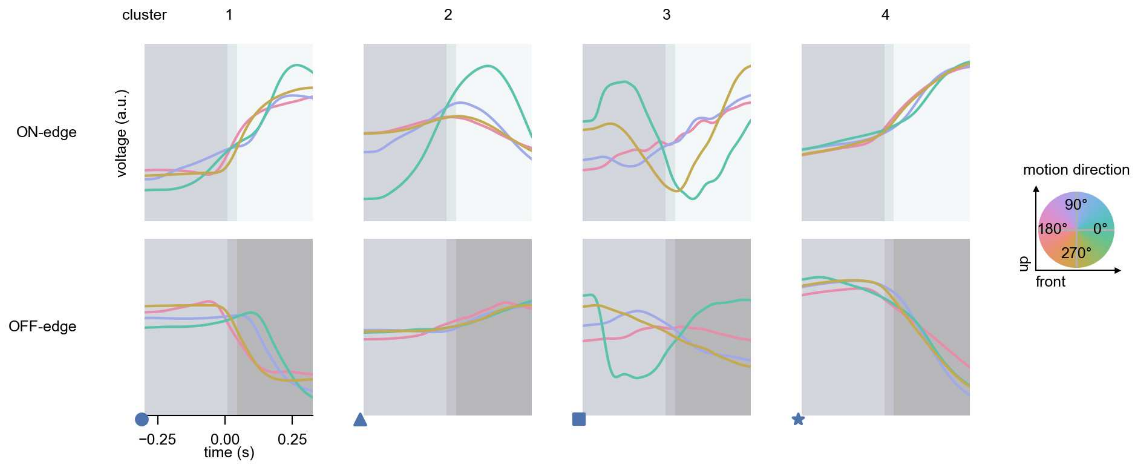

Mi12 - Figure 7: **Responses to moving edges from best-performing models.** The top row shows responses to moving ON-edges, the bottom row shows peak responses to moving OFF-edges. The edge-stimuli move in different directions from 0 to 360 degrees and at different speeds. Before, a grey-stimulus led the network to a stationary state. The columns show responses from the best-task-performing model in the respective cluster. Example responses to edges moving at 75.4°/s in all cardinal directions (green 0°, blue 90°, red 180°, yellow 270°) from -22.5 to 22.5° visual angle. Background contour indicates the edge, passing by the central column at second zero.

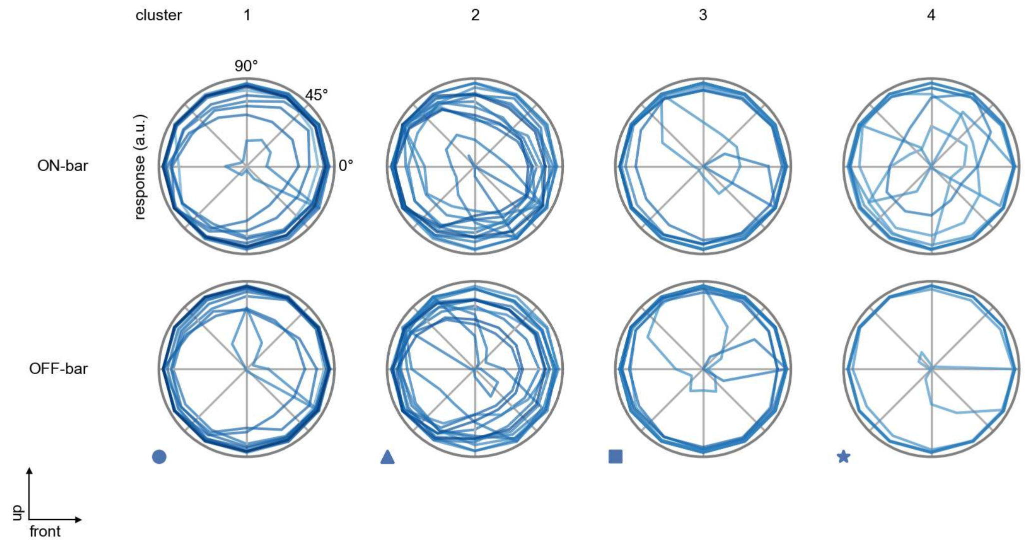

Mi12 - Figure 8: **Peak responses to moving bars.** The top row shows normalized peak responses to moving ON-bars, the bottom row shows normalized peak responses to moving OFF-bars. The bar-stimuli move in different directions from 0 to 360 degrees. Before, a grey-stimulus led the network to a stationary state. The responses from the different models in the different clusters (columns) overlay and are colored in the respective task error of the model. The peak responses are averaged over all assessed bar-speeds.

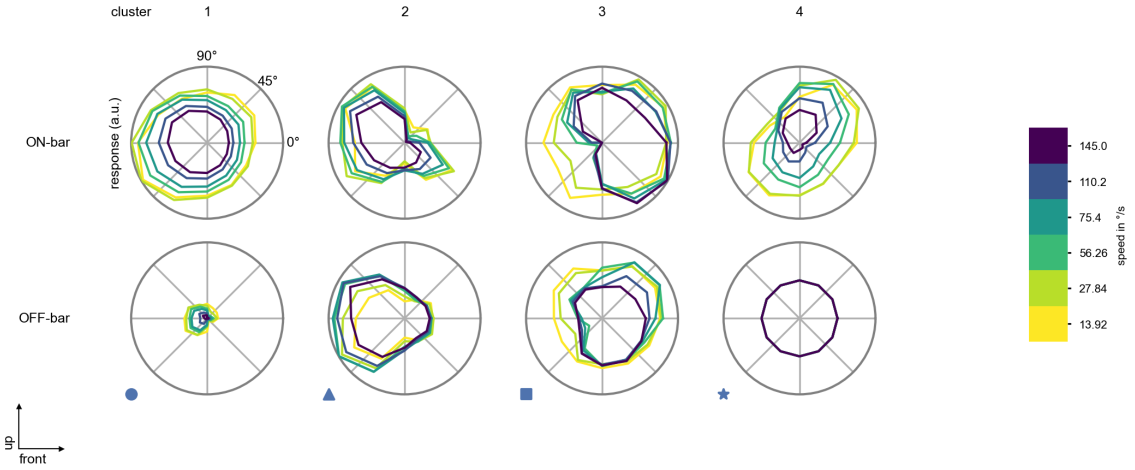

Mi12 - Figure 9: **Peak responses to moving bars from best-performing models.** The top row shows peak responses to moving ON-bars, the bottom row shows peak responses to moving OFF-bars. The bar-stimuli move in different directions from 0 to 360 degrees and at different speeds. Before, a grey-stimulus led the network to a stationary state. The columns show responses from the best-task-performing model in the respective cluster.

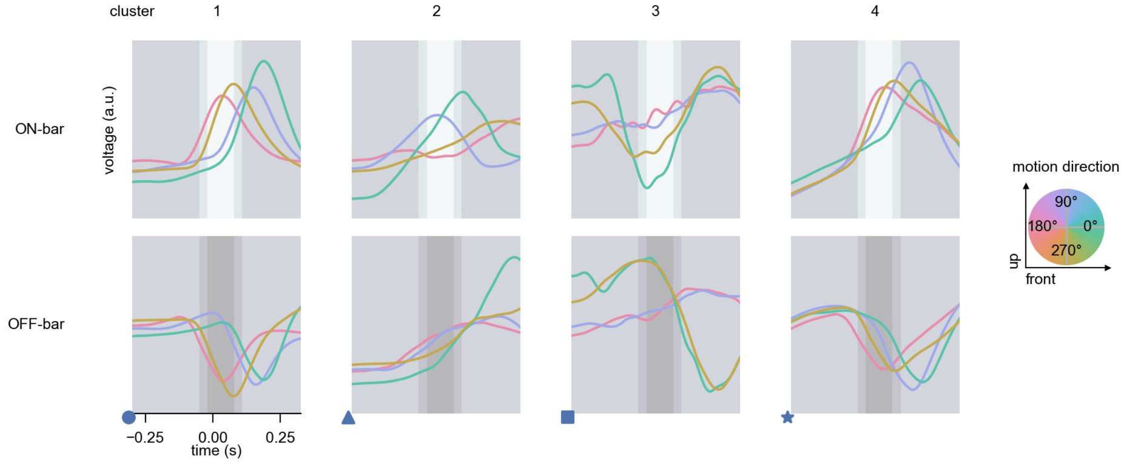

Mi12 - Figure 10: **Responses to moving bars from best-performing models.** The top row shows responses to moving ON-bars, the bottom row shows peak responses to moving OFF-bars. The bar-stimuli move in different directions from 0 to 360 degrees and at different speeds. Before, a grey-stimulus led the network to a stationary state. The columns show responses from the best-task-performing model in the respective cluster. Example responses to bars moving at 75.4°/s in all cardinal directions (green 0°, blue 90°, red 180°, yellow 270°) from -22.5 to 22.5° visual angle. Background contour indicates the bar, passing by the central column at second zero.

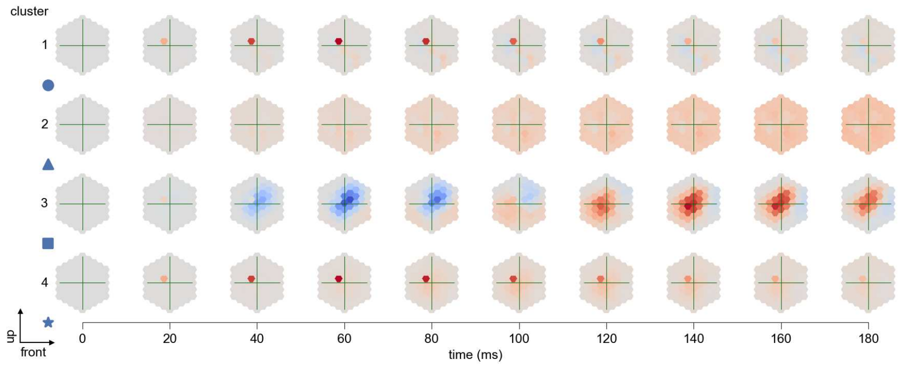

Mi12 - Figure 11: **Spatio-temporal receptive field.** Responses of the central cell to ON-impulses at single ommatidium impulse locations. The impulse occurs at second zero. Before, a grey-stimulus led the network to a stationary state. Responses are baseline-subtracted. Each row illustrates responses from the best-performing model of the respective cluster. Red indicated depolarization, blue indicates hyperpolarization.

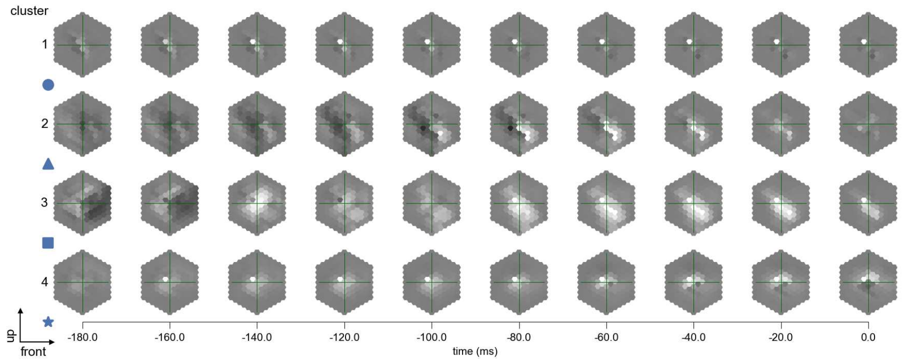

Mi12 - Figure 12: **Maximally excitatory stimuli.** Each row presents the regularized naturalistic-stimulus from the Sintel dataset that maximizes the cell type's central column response at second zero in the best-performing model of the respective cluster.

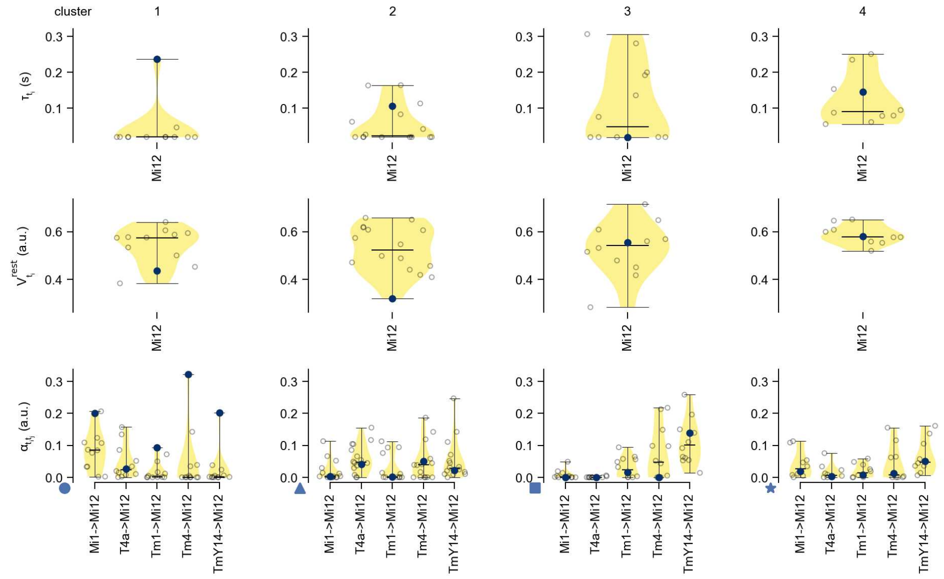

Mi12 - Figure 13: **Task-constrained parameters.** Each column shows the parameters inferred within the respective cluster. First row: learned time constants of the cell type. Second row: resting potentials of the cell type. Third row: scaling factors for the convolutional filters. The blue scatter represents the parameters from the best-performing model within the cluster.

# 29 Mi13

← Cell types

#### Figures

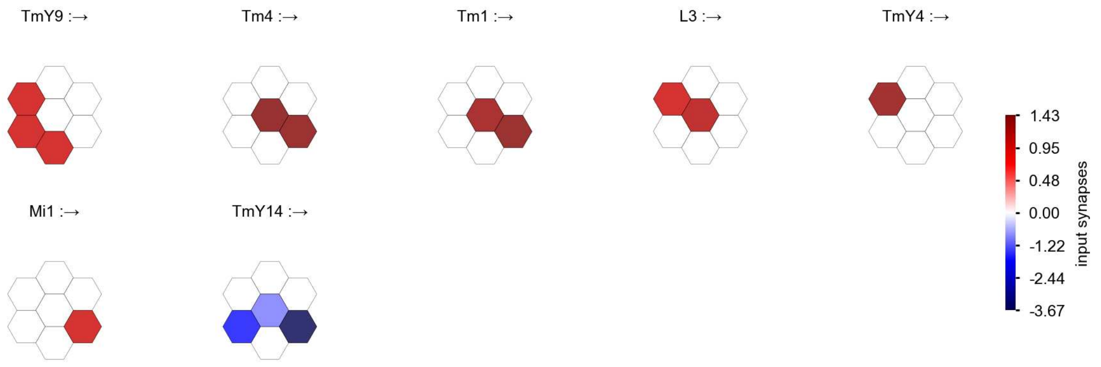

Mi13 - Figure 1: **Anatomical receptive fields.** Each colored hexagon is a projection, with the projection strength characterized by the average number of synapses that we count from the EM reconstruction. Red indicates excitatory synapses, blue indicates inhibitory synapses from inferred signs. Filters in the order of their total number of synapses.

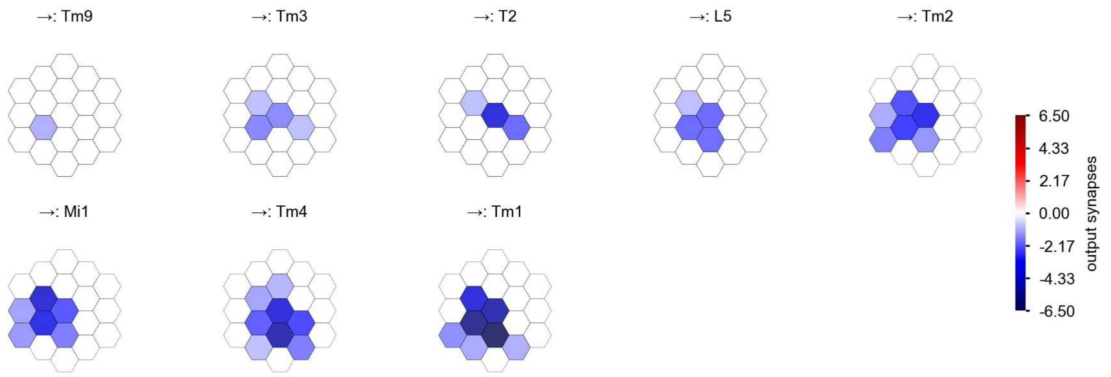

Mi13 - Figure 2: **Anatomical projective fields.** Each colored hexagon is a projection, with the projection strength characterized by the average number of synapses that we count from the EM reconstruction. Red indicates excitatory synapses, blue indicates inhibitory synapses from inferred signs. Filters in the order of their total number of synapses.

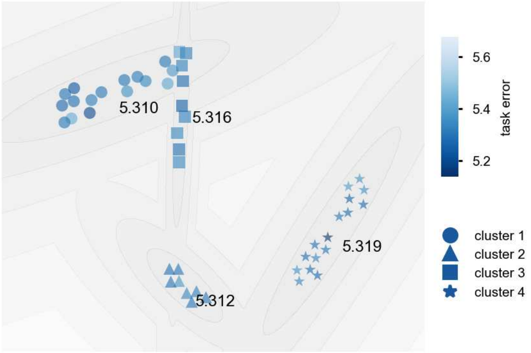

Mi13 - Figure 3: **Clustering of the responses to naturalistic stimuli.** From responses to naturalistic scenes from Sintel. Scatterpoints represent individual models colored by their task error.

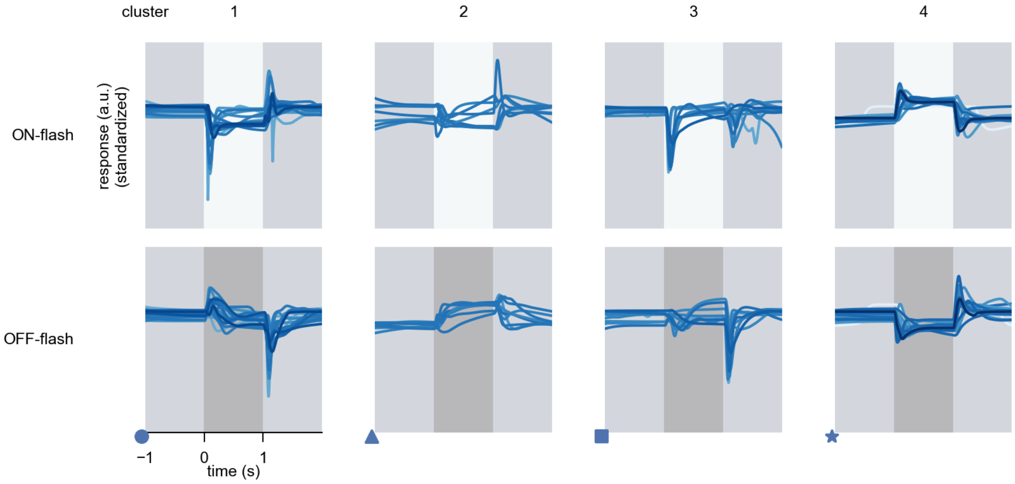

Mi13 - Figure 4: **Responses to flashes.** The top row shows responses to ON-flashes, the bottom row shows responses to OFF-flashes. The responses from the different models in the different clusters (columns) overlay and are colored in the respective task error of the model. The circular flashes occur at second zero, cover 6 ommatidia in radius, and last for one second. Before and after, a grey-stimulus leads to a stationary state of the network. The traces are standardized.

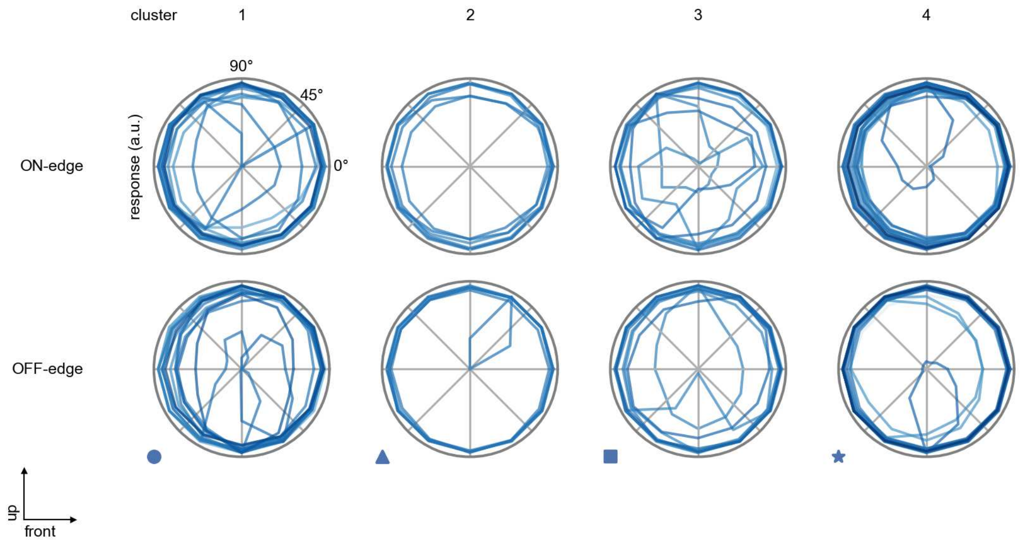

Mi13 - Figure 5: **Peak responses to moving edges.** The top row shows normalized peak responses to moving ON-edges, the bottom row shows normalized peak responses to moving OFF-edges. The edge-stimuli move in different directions from 0 to 360 degrees. Before, a grey-stimulus led the network to a stationary state. The responses from the different models in the different clusters (columns) overlay and are colored in the respective task error of the model. The peak responses are averaged over all assessed edge-speeds.

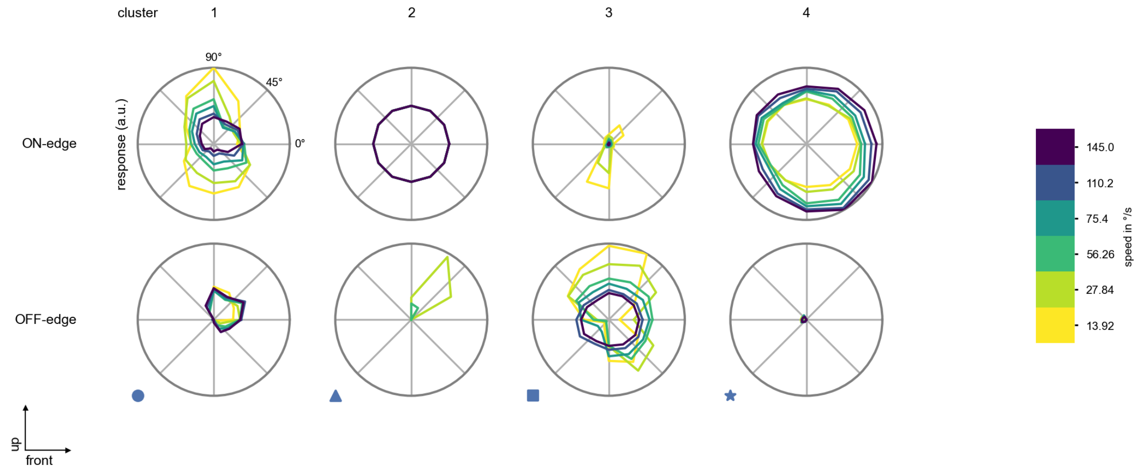

Mi13 - Figure 6: **Peak responses to moving edges from best-performing models.** The top row shows peak responses to moving ON-edges, the bottom row shows peak responses to moving OFF-edges. The edge-stimuli move in different directions from 0 to 360 degrees and at different speeds. Before, a grey-stimulus led the network to a stationary state. The columns show responses from the best-task-performing model in the respective cluster.

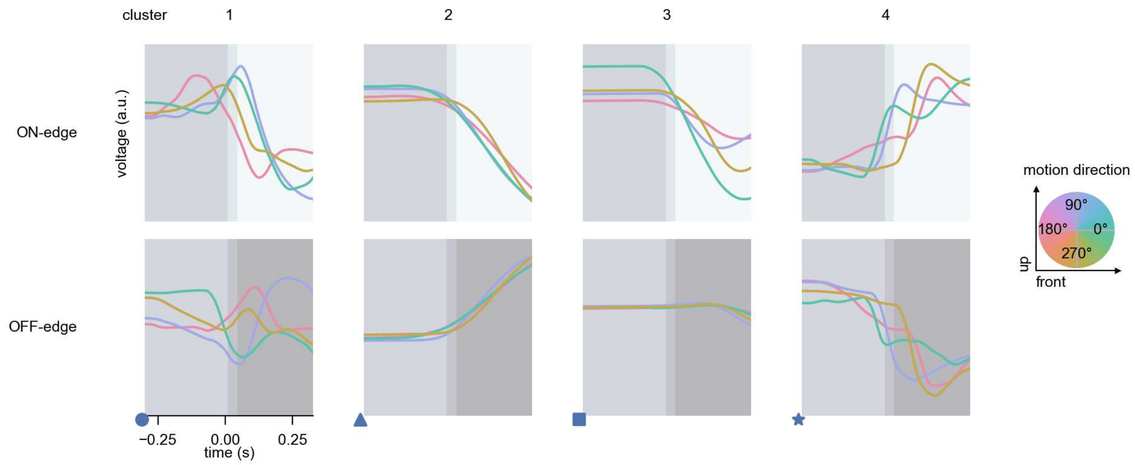

Mi13 - Figure 7: **Responses to moving edges from best-performing models.** The top row shows responses to moving ON-edges, the bottom row shows peak responses to moving OFF-edges. The edge-stimuli move in different directions from 0 to 360 degrees and at different speeds. Before, a grey-stimulus led the network to a stationary state. The columns show responses from the best-task-performing model in the respective cluster. Example responses to edges moving at 75.4°/s in all cardinal directions (green 0°, blue 90°, red 180°, yellow 270°) from -22.5 to 22.5° visual angle. Background contour indicates the edge, passing by the central column at second zero.

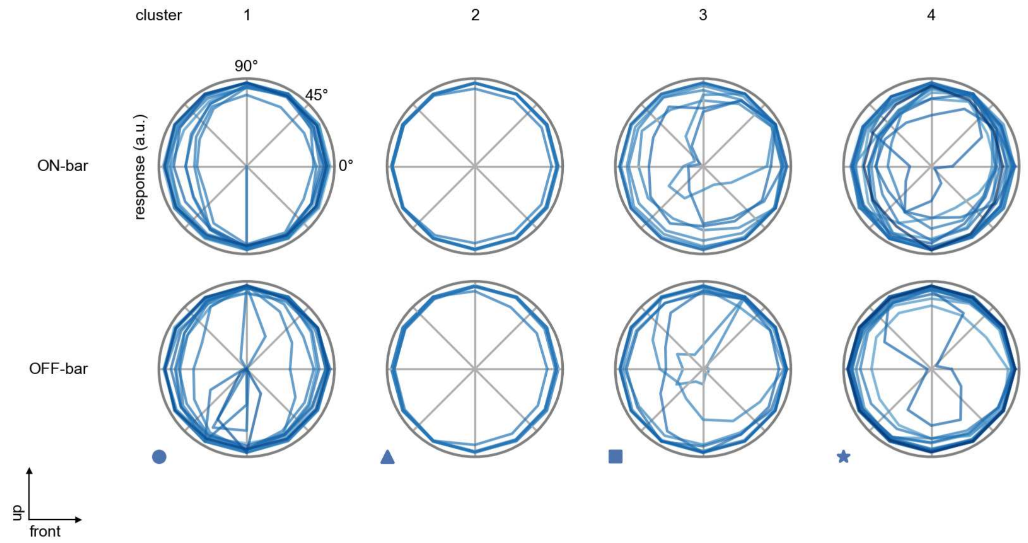

Mi13 - Figure 8: **Peak responses to moving bars.** The top row shows normalized peak responses to moving ON-bars, the bottom row shows normalized peak responses to moving OFF-bars. The bar-stimuli move in different directions from 0 to 360 degrees. Before, a grey-stimulus led the network to a stationary state. The responses from the different models in the different clusters (columns) overlay and are colored in the respective task error of the model. The peak responses are averaged over all assessed bar-speeds.

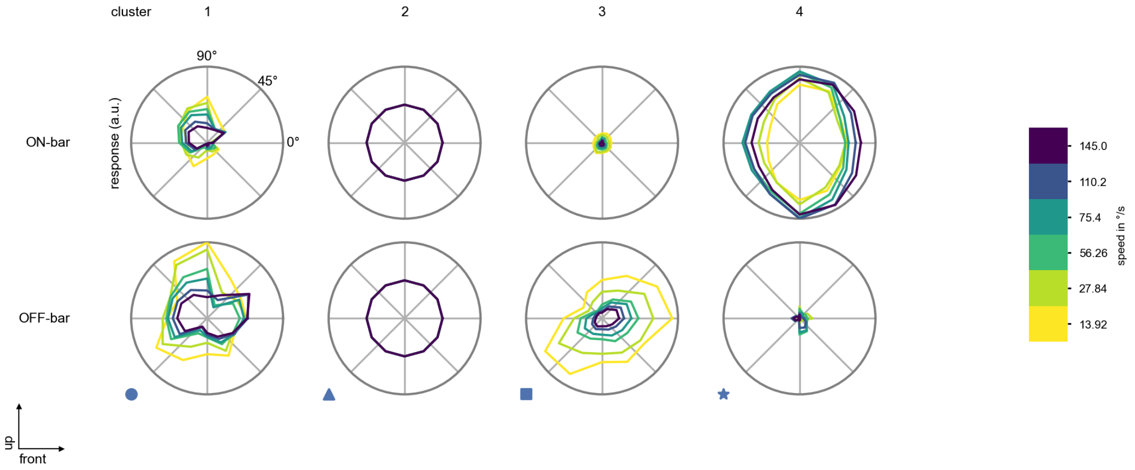

Mi13 - Figure 9: **Peak responses to moving bars from best-performing models.** The top row shows peak responses to moving ON-bars, the bottom row shows peak responses to moving OFF-bars. The bar-stimuli move in different directions from 0 to 360 degrees and at different speeds. Before, a grey-stimulus led the network to a stationary state. The columns show responses from the best-task-performing model in the respective cluster.

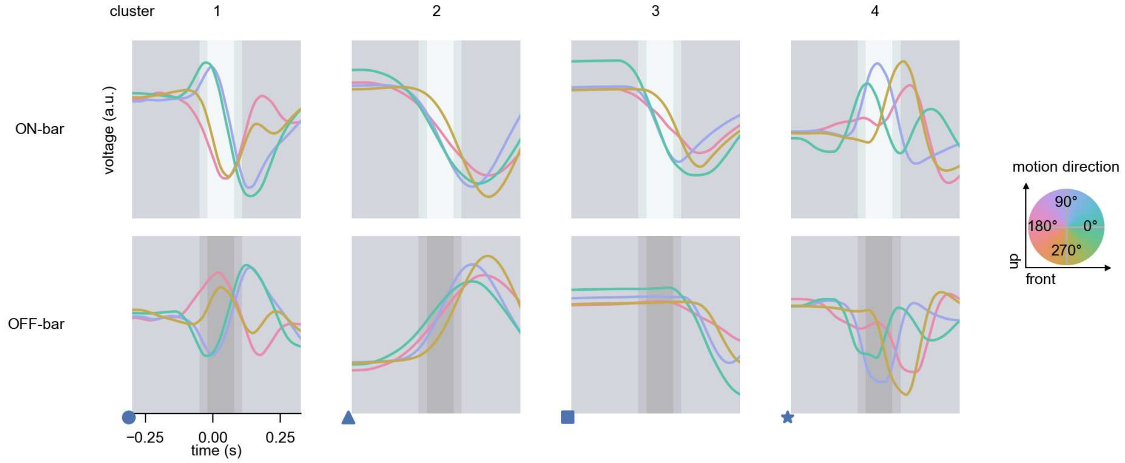

Mi13 - Figure 10: **Responses to moving bars from best-performing models.** The top row shows responses to moving ON-bars, the bottom row shows peak responses to moving OFF-bars. The bar-stimuli move in different directions from 0 to 360 degrees and at different speeds. Before, a grey-stimulus led the network to a stationary state. The columns show responses from the best-task-performing model in the respective cluster. Example responses to bars moving at 75.4°/s in all cardinal directions (green 0°, blue 90°, red 180°, yellow 270°) from -22.5 to 22.5° visual angle. Background contour indicates the bar, passing by the central column at second zero.

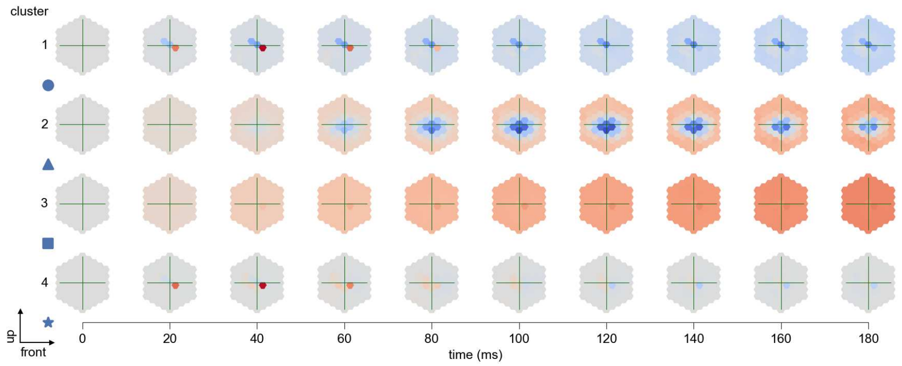

Mi13 - Figure 11: **Spatio-temporal receptive field.** Responses of the central cell to ON-impulses at single ommatidium impulse locations. The impulse occurs at second zero. Before, a grey-stimulus led the network to a stationary state. Responses are baseline-subtracted. Each row illustrates responses from the best-performing model of the respective cluster. Red indicated depolarization, blue indicates hyperpolarization.

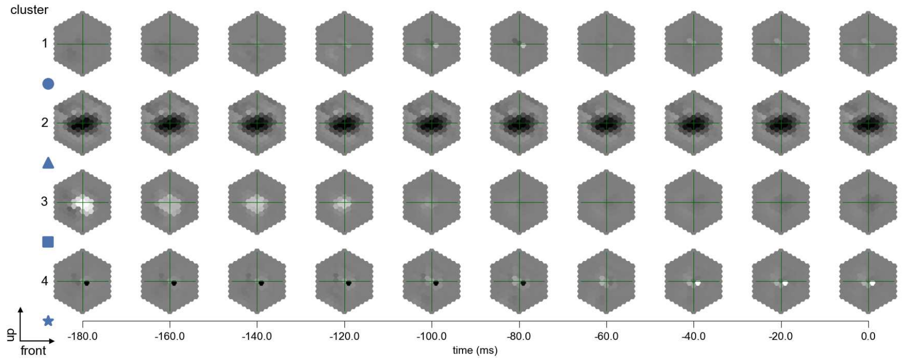

Mi13 - Figure 12: **Maximally excitatory stimuli.** Each row presents the regularized naturalistic-stimulus from the Sintel dataset that maximizes the cell type's central column response at second zero in the best-performing model of the respective cluster.

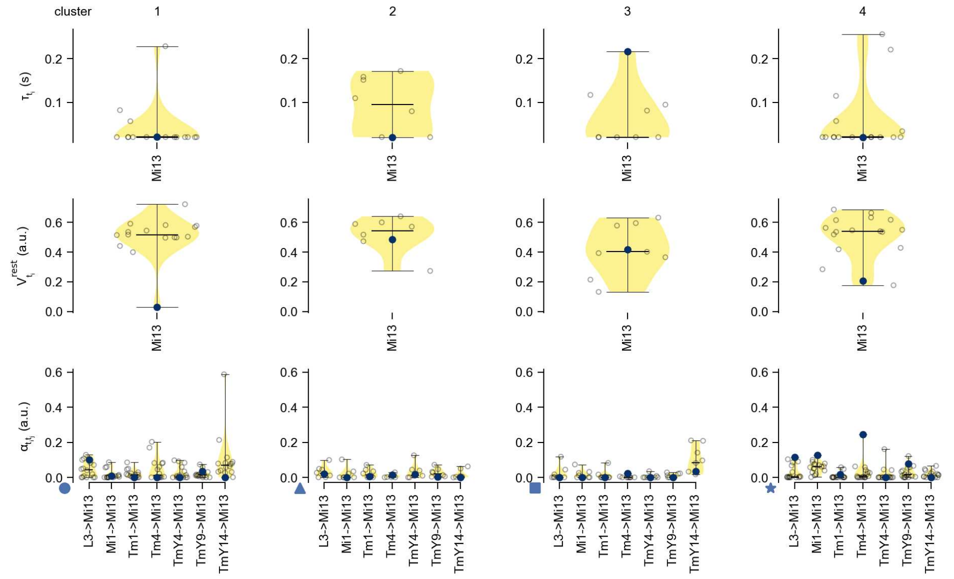

Mi13 - Figure 13: **Task-constrained parameters.** Each column shows the parameters inferred within the respective cluster. First row: learned time constants of the cell type. Second row: resting potentials of the cell type. Third row: scaling factors for the convolutional filters. The blue scatter represents the parameters from the best-performing model within the cluster.

# 30 Mi14

← Cell types

#### Figures

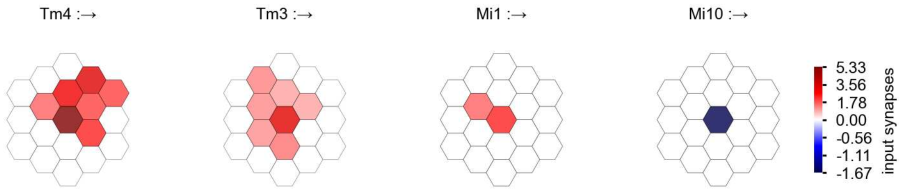

Mi14 - Figure 1: **Anatomical receptive fields.** Each colored hexagon is a projection, with the projection strength characterized by the average number of synapses that we count from the EM reconstruction. Red indicates excitatory synapses, blue indicates inhibitory synapses from inferred signs. Filters in the order of their total number of synapses.

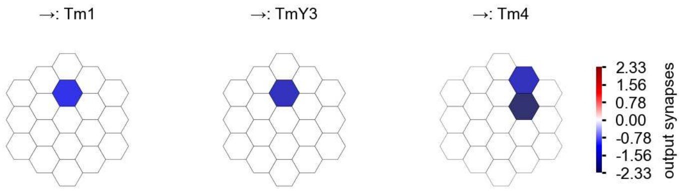

Mi14 - Figure 2: **Anatomical projective fields.** Each colored hexagon is a projection, with the projection strength characterized by the average number of synapses that we count from the EM reconstruction. Red indicates excitatory synapses, blue indicates inhibitory synapses from inferred signs. Filters in the order of their total number of synapses.

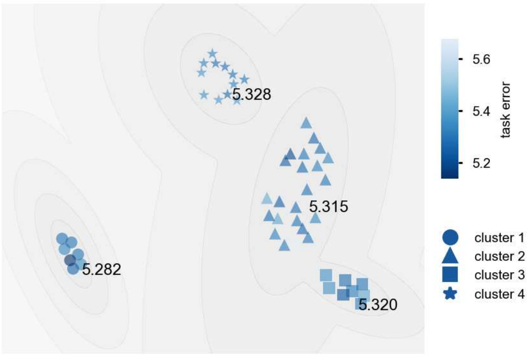

Mi14 - Figure 3: **Clustering of the responses to naturalistic stimuli.** From responses to naturalistic scenes from Sintel. Scatterpoints represent individual models colored by their task error.

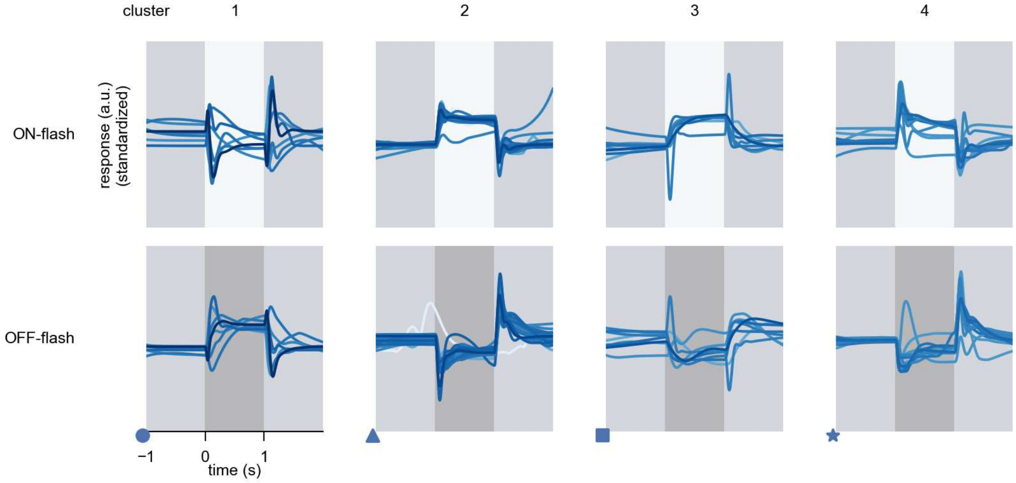

Mi14 - Figure 4: **Responses to flashes.** The top row shows responses to ON-flashes, the bottom row shows responses to OFF-flashes. The responses from the different models in the different clusters (columns) overlay and are colored in the respective task error of the model. The circular flashes occur at second zero, cover 6 ommatidia in radius, and last for one second. Before and after, a grey-stimulus leads to a stationary state of the network. The traces are standardized.

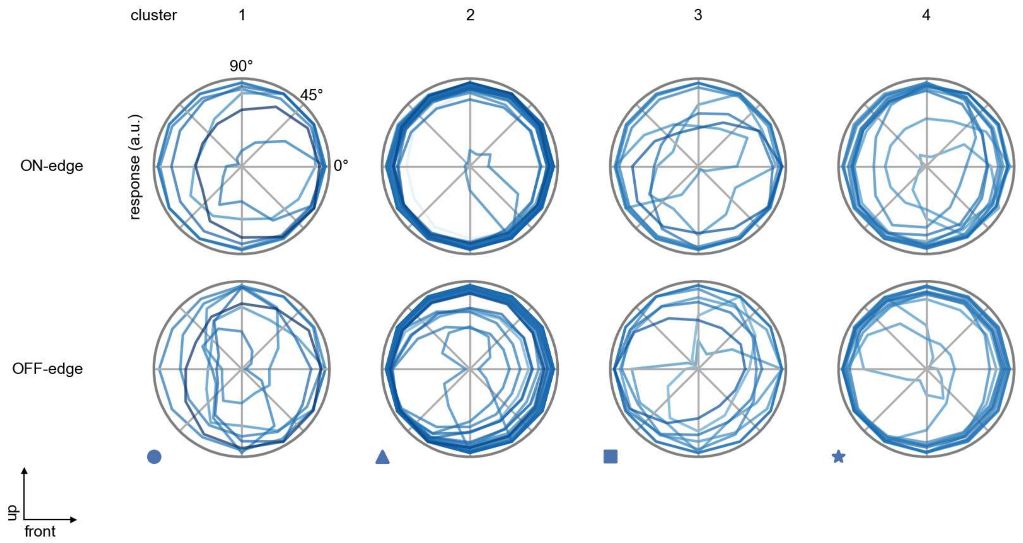

Mi14 - Figure 5: **Peak responses to moving edges.** The top row shows normalized peak responses to moving ON-edges, the bottom row shows normalized peak responses to moving OFF-edges. The edge-stimuli move in different directions from 0 to 360 degrees. Before, a grey-stimulus led the network to a stationary state. The responses from the different models in the different clusters (columns) overlay and are colored in the respective task error of the model. The peak responses are averaged over all assessed edge-speeds.

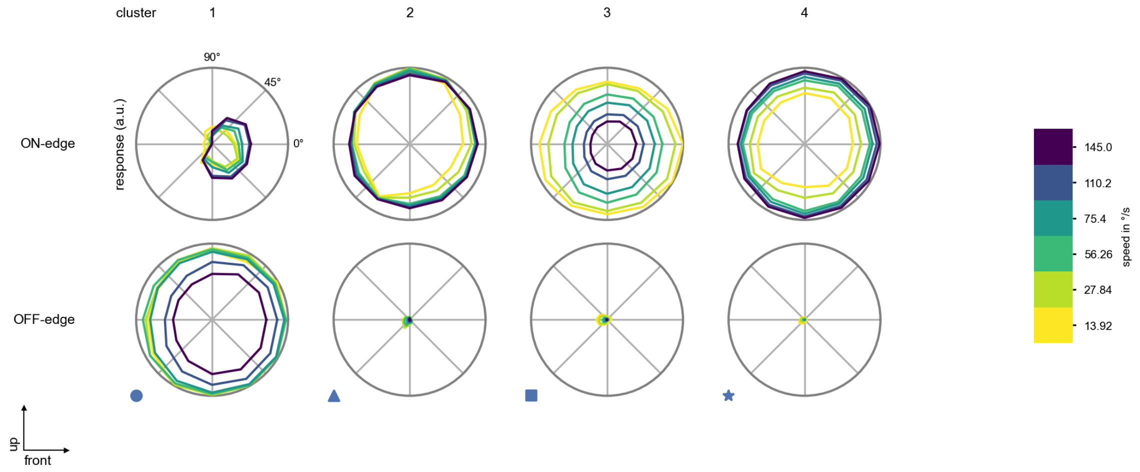

Mi14 - Figure 6: **Peak responses to moving edges from best-performing models.** The top row shows peak responses to moving ON-edges, the bottom row shows peak responses to moving OFF-edges. The edge-stimuli move in different directions from 0 to 360 degrees and at different speeds. Before, a grey-stimulus led the network to a stationary state. The columns show responses from the best-task-performing model in the respective cluster.

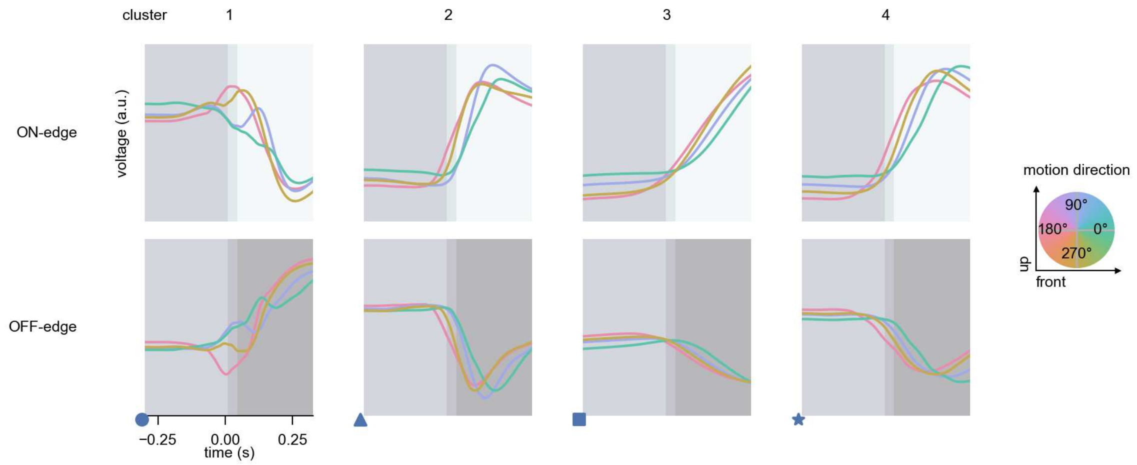

Mi14 - Figure 7: **Responses to moving edges from best-performing models.** The top row shows responses to moving ON-edges, the bottom row shows peak responses to moving OFF-edges. The edge-stimuli move in different directions from 0 to 360 degrees and at different speeds. Before, a grey-stimulus led the network to a stationary state. The columns show responses from the best-task-performing model in the respective cluster. Example responses to edges moving at 75.4°/s in all cardinal directions (green 0°, blue 90°, red 180°, yellow 270°) from -22.5 to 22.5° visual angle. Background contour indicates the edge, passing by the central column at second zero.

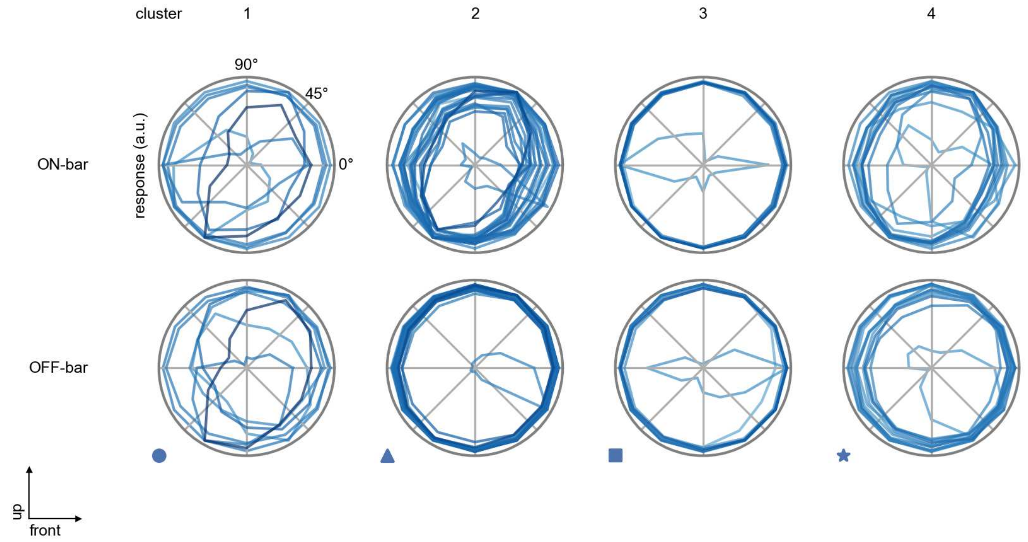

Mi14 - Figure 8: **Peak responses to moving bars.** The top row shows normalized peak responses to moving ON-bars, the bottom row shows normalized peak responses to moving OFF-bars. The bar-stimuli move in different directions from 0 to 360 degrees. Before, a grey-stimulus led the network to a stationary state. The responses from the different models in the different clusters (columns) overlay and are colored in the respective task error of the model. The peak responses are averaged over all assessed bar-speeds.

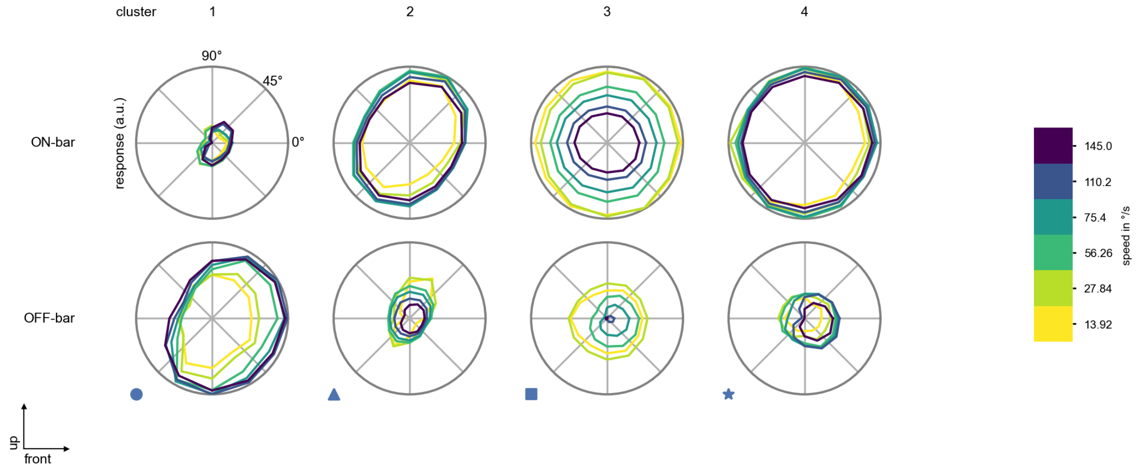

Mi14 - Figure 9: **Peak responses to moving bars from best-performing models.** The top row shows peak responses to moving ON-bars, the bottom row shows peak responses to moving OFF-bars. The bar-stimuli move in different directions from 0 to 360 degrees and at different speeds. Before, a grey-stimulus led the network to a stationary state. The columns show responses from the best-task-performing model in the respective cluster.

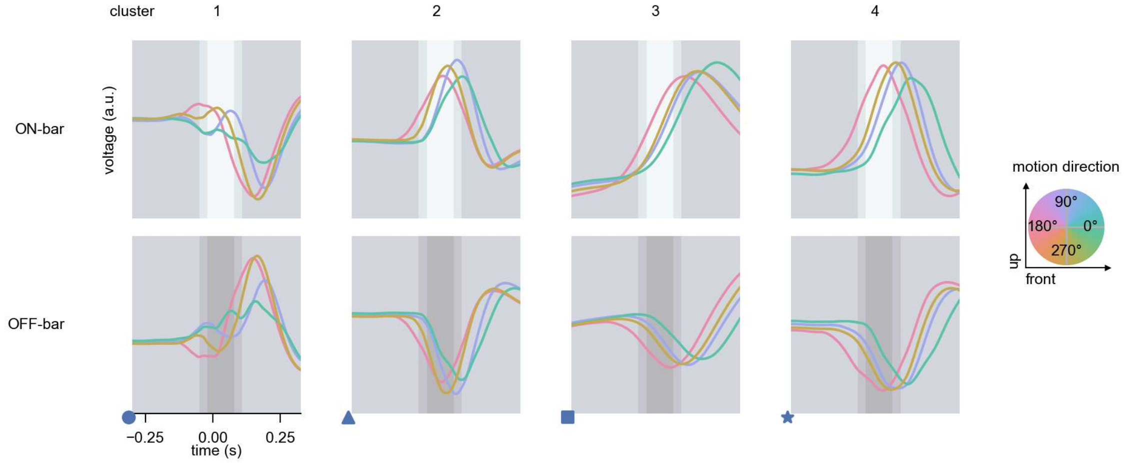

Mi14 - Figure 10: **Responses to moving bars from best-performing models.** The top row shows responses to moving ON-bars, the bottom row shows peak responses to moving OFF-bars. The bar-stimuli move in different directions from 0 to 360 degrees and at different speeds. Before, a grey-stimulus led the network to a stationary state. The columns show responses from the best-task-performing model in the respective cluster. Example responses to bars moving at 75.4°/s in all cardinal directions (green 0°, blue 90°, red 180°, yellow 270°) from -22.5 to 22.5° visual angle. Background contour indicates the bar, passing by the central column at second zero.

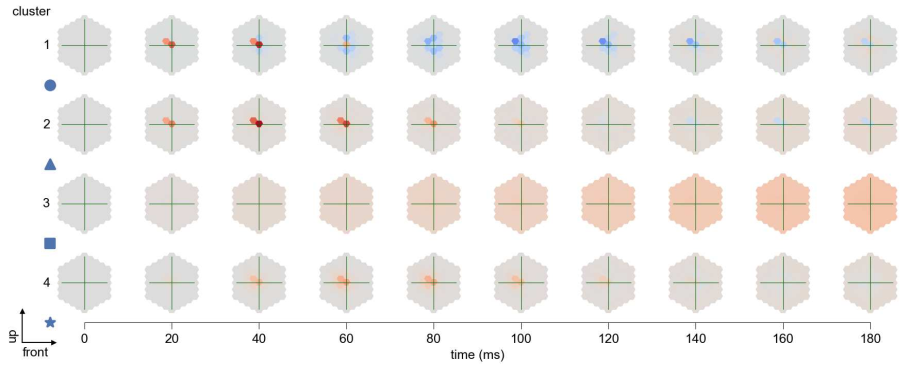

Mi14 - Figure 11: **Spatio-temporal receptive field.** Responses of the central cell to ON-impulses at single ommatidium impulse locations. The impulse occurs at second zero. Before, a grey-stimulus led the network to a stationary state. Responses are baseline-subtracted. Each row illustrates responses from the best-performing model of the respective cluster. Red indicated depolarization, blue indicates hyperpolarization.

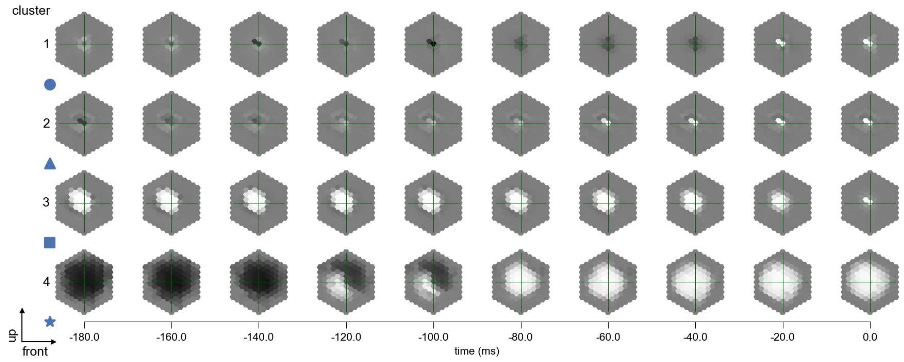

Mi14 - Figure 12: **Maximally excitatory stimuli.** Each row presents the regularized naturalistic-stimulus from the Sintel dataset that maximizes the cell type's central column response at second zero in the best-performing model of the respective cluster.

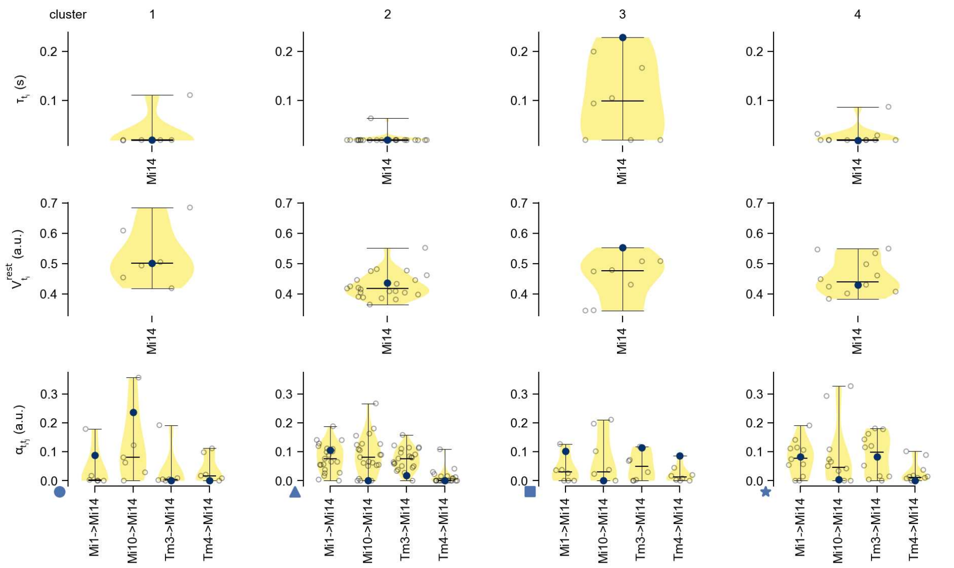

Mi14 - Figure 13: **Task-constrained parameters.** Each column shows the parameters inferred within the respective cluster. First row: learned time constants of the cell type. Second row: resting potentials of the cell type. Third row: scaling factors for the convolutional filters. The blue scatter represents the parameters from the best-performing model within the cluster.

# 31 Mi15

← Cell types

#### Figures

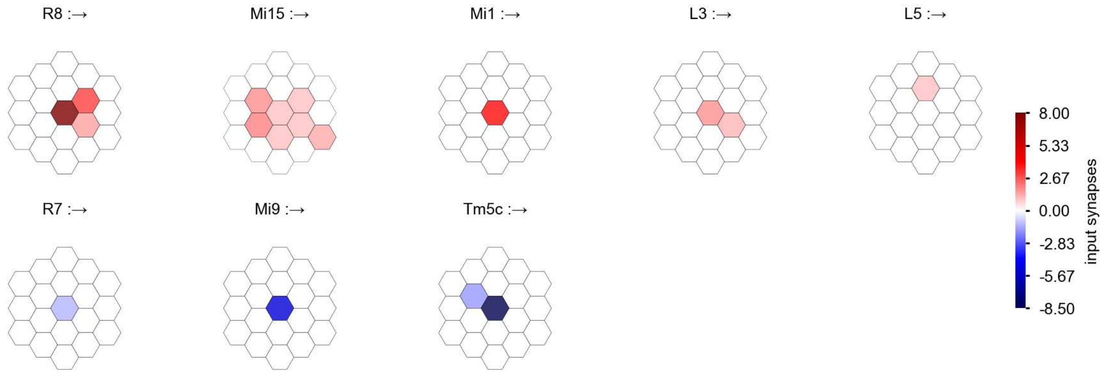

Mi15 - Figure 1: **Anatomical receptive fields.** Each colored hexagon is a projection, with the projection strength characterized by the average number of synapses that we count from the EM reconstruction. Red indicates excitatory synapses, blue indicates inhibitory synapses from inferred signs. Filters in the order of their total number of synapses.

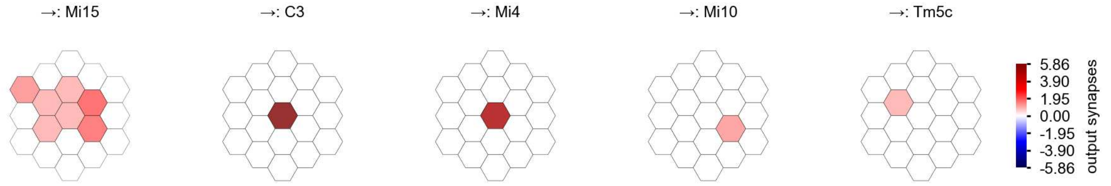

Mi15 - Figure 2: **Anatomical projective fields.** Each colored hexagon is a projection, with the projection strength characterized by the average number of synapses that we count from the EM reconstruction. Red indicates excitatory synapses, blue indicates inhibitory synapses from inferred signs. Filters in the order of their total number of synapses.

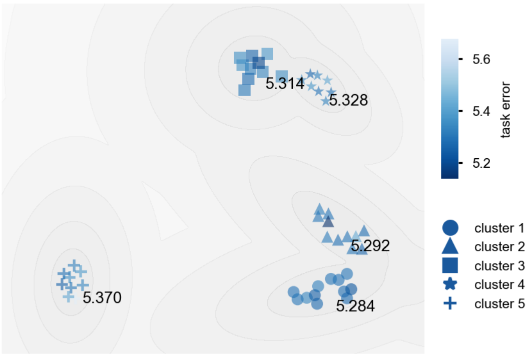

Mi15 - Figure 3: **Clustering of the responses to naturalistic stimuli.** From responses to naturalistic scenes from Sintel. Scatterpoints represent individual models colored by their task error.

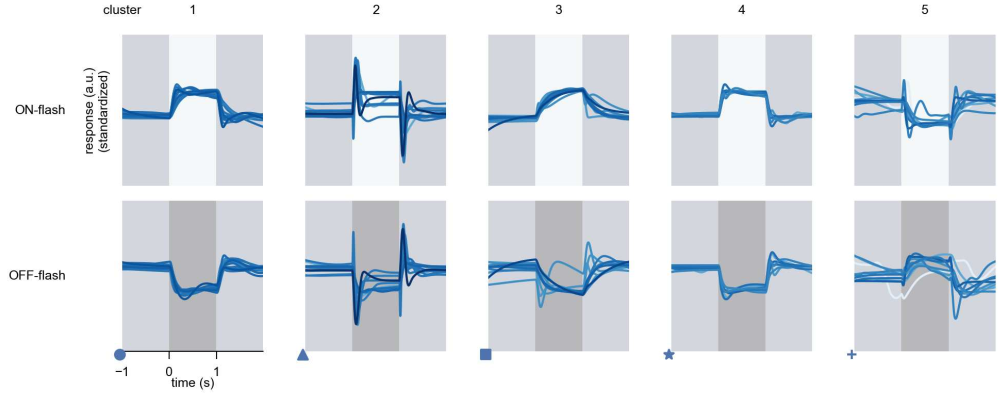

Mi15 - Figure 4: **Responses to flashes.** The top row shows responses to ON-flashes, the bottom row shows responses to OFF-flashes. The responses from the different models in the different clusters (columns) overlay and are colored in the respective task error of the model. The circular flashes occur at second zero, cover 6 ommatidia in radius, and last for one second. Before and after, a grey-stimulus leads to a stationary state of the network. The traces are standardized.

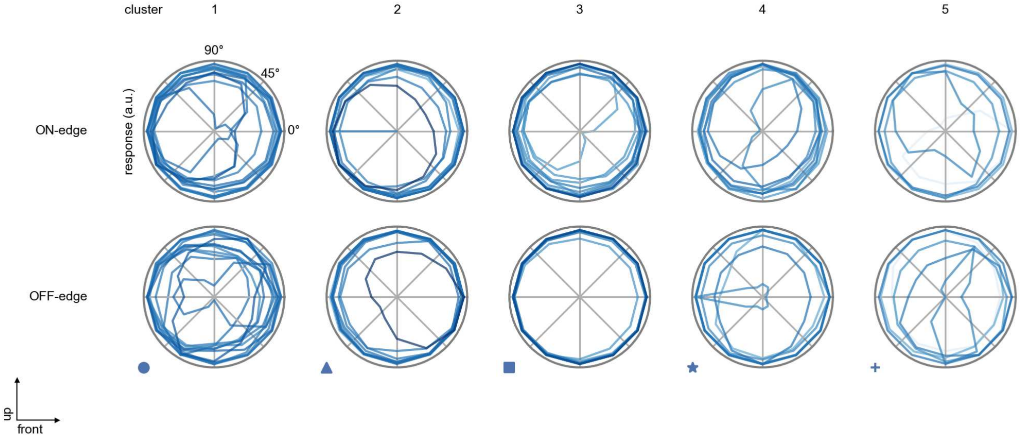

Mi15 - Figure 5: **Peak responses to moving edges.** The top row shows normalized peak responses to moving ON-edges, the bottom row shows normalized peak responses to moving OFF-edges. The edge-stimuli move in different directions from 0 to 360 degrees. Before, a grey-stimulus led the network to a stationary state. The responses from the different models in the different clusters (columns) overlay and are colored in the respective task error of the model. The peak responses are averaged over all assessed edge-speeds.

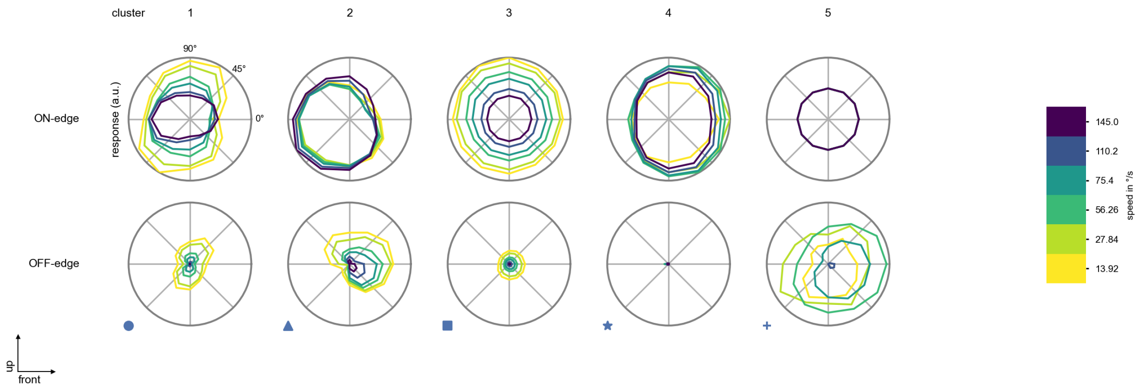

Mi15 - Figure 6: **Peak responses to moving edges from best-performing models.** The top row shows peak responses to moving ON-edges, the bottom row shows peak responses to moving OFF-edges. The edge-stimuli move in different directions from 0 to 360 degrees and at different speeds. Before, a grey-stimulus led the network to a stationary state. The columns show responses from the best-task-performing model in the respective cluster.

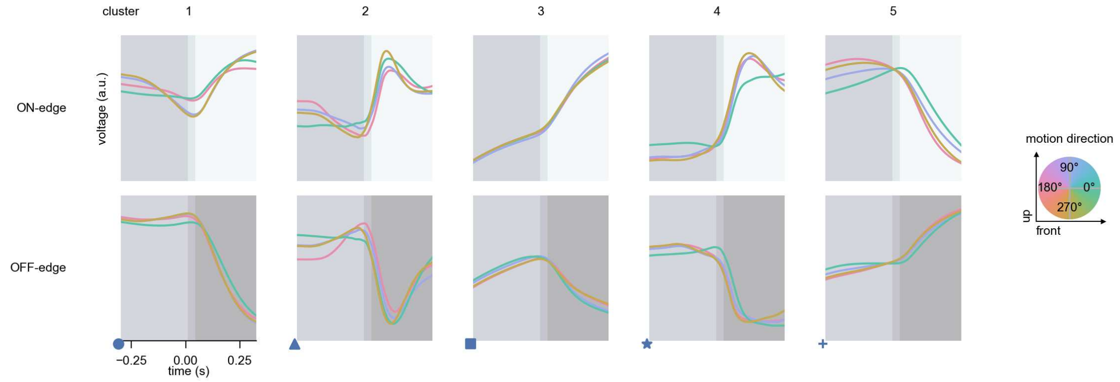

Mi15 - Figure 7: **Responses to moving edges from best-performing models.** The top row shows responses to moving ON-edges, the bottom row shows peak responses to moving OFF-edges. The edge-stimuli move in different directions from 0 to 360 degrees and at different speeds. Before, a grey-stimulus led the network to a stationary state. The columns show responses from the best-task-performing model in the respective cluster. Example responses to edges moving at 75.4°/s in all cardinal directions (green 0°, blue 90°, red 180°, yellow 270°) from -22.5 to 22.5° visual angle. Background contour indicates the edge, passing by the central column at second zero.

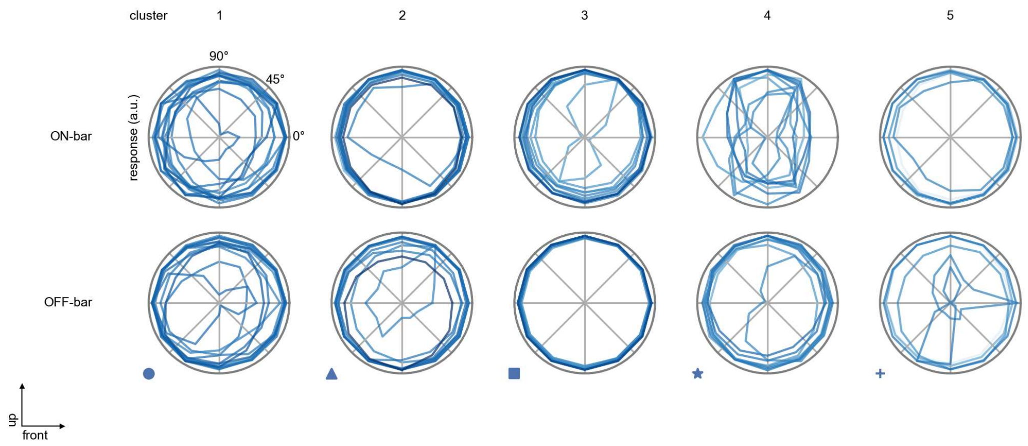

Mi15 - Figure 8: **Peak responses to moving bars.** The top row shows normalized peak responses to moving ON-bars, the bottom row shows normalized peak responses to moving OFF-bars. The bar-stimuli move in different directions from 0 to 360 degrees. Before, a grey-stimulus led the network to a stationary state. The responses from the different models in the different clusters (columns) overlay and are colored in the respective task error of the model. The peak responses are averaged over all assessed bar-speeds.

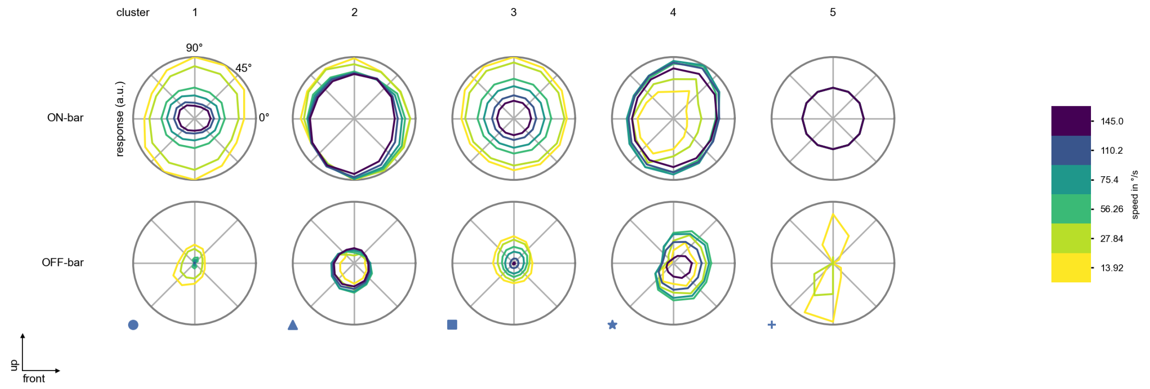

Mi15 - Figure 9: **Peak responses to moving bars from best-performing models.** The top row shows peak responses to moving ON-bars, the bottom row shows peak responses to moving OFF-bars. The bar-stimuli move in different directions from 0 to 360 degrees and at different speeds. Before, a grey-stimulus led the network to a stationary state. The columns show responses from the best-task-performing model in the respective cluster.

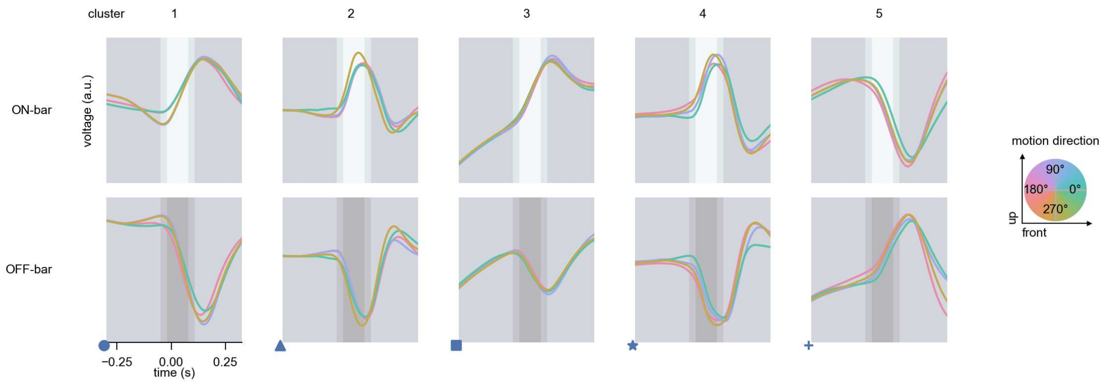

Mi15 - Figure 10: **Responses to moving bars from best-performing models.** The top row shows responses to moving ON-bars, the bottom row shows peak responses to moving OFF-bars. The bar-stimuli move in different directions from 0 to 360 degrees and at different speeds. Before, a grey-stimulus led the network to a stationary state. The columns show responses from the best-task-performing model in the respective cluster. Example responses to bars moving at 75.4°/s in all cardinal directions (green 0°, blue 90°, red 180°, yellow 270°) from -22.5 to 22.5° visual angle. Background contour indicates the bar, passing by the central column at second zero.

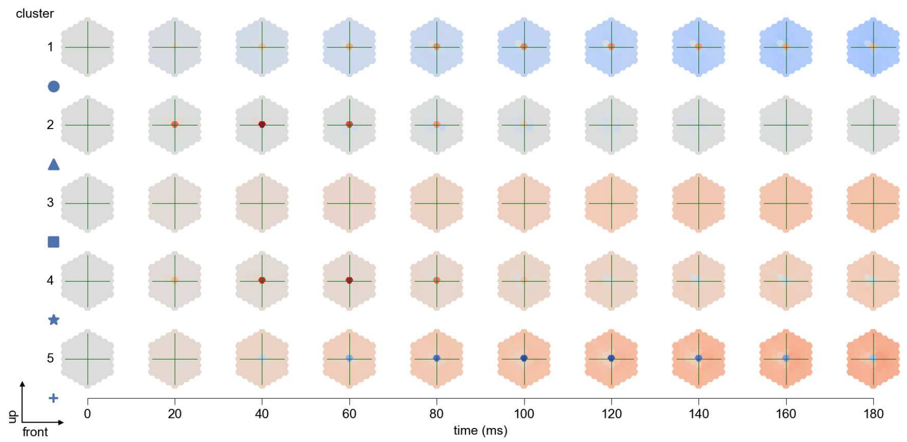

Mi15 - Figure 11: **Spatio-temporal receptive field.** Responses of the central cell to ON-impulses at single ommatidium impulse locations. The impulse occurs at second zero. Before, a grey-stimulus led the network to a stationary state. Responses are baseline-subtracted. Each row illustrates responses from the best-performing model of the respective cluster. Red indicated depolarization, blue indicates hyperpolarization.

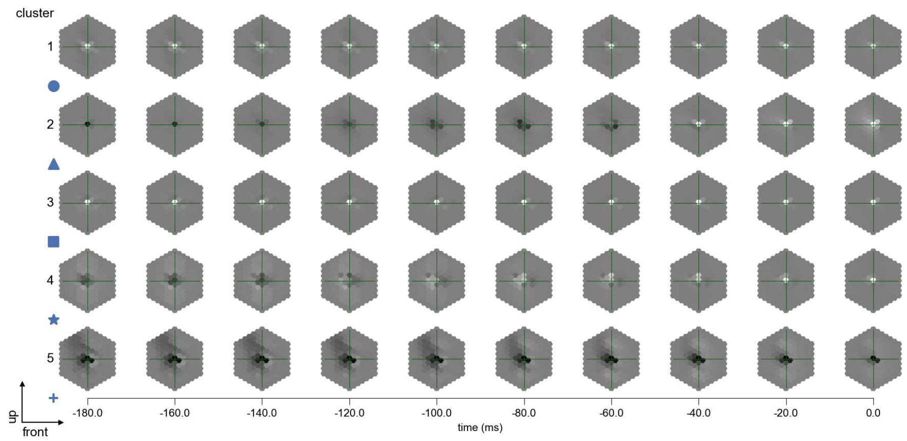

Mi15 - Figure 12: **Maximally excitatory stimuli.** Each row presents the regularized naturalistic-stimulus from the Sintel dataset that maximizes the cell type's central column response at second zero in the best-performing model of the respective cluster.

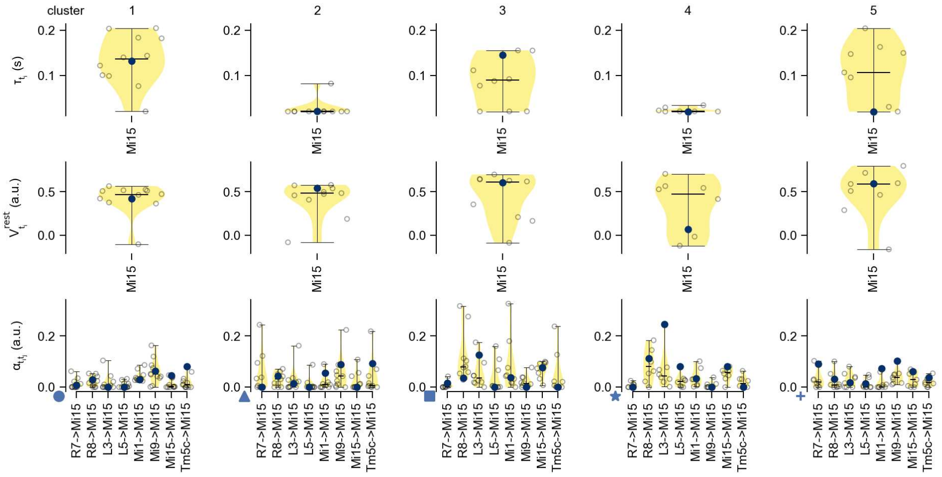

Mi15 - Figure 13: **Task-constrained parameters.** Each column shows the parameters inferred within the respective cluster. First row: learned time constants of the cell type. Second row: resting potentials of the cell type. Third row: scaling factors for the convolutional filters. The blue scatter represents the parameters from the best-performing model within the cluster.

# 32 T1

← Cell types

#### Figures

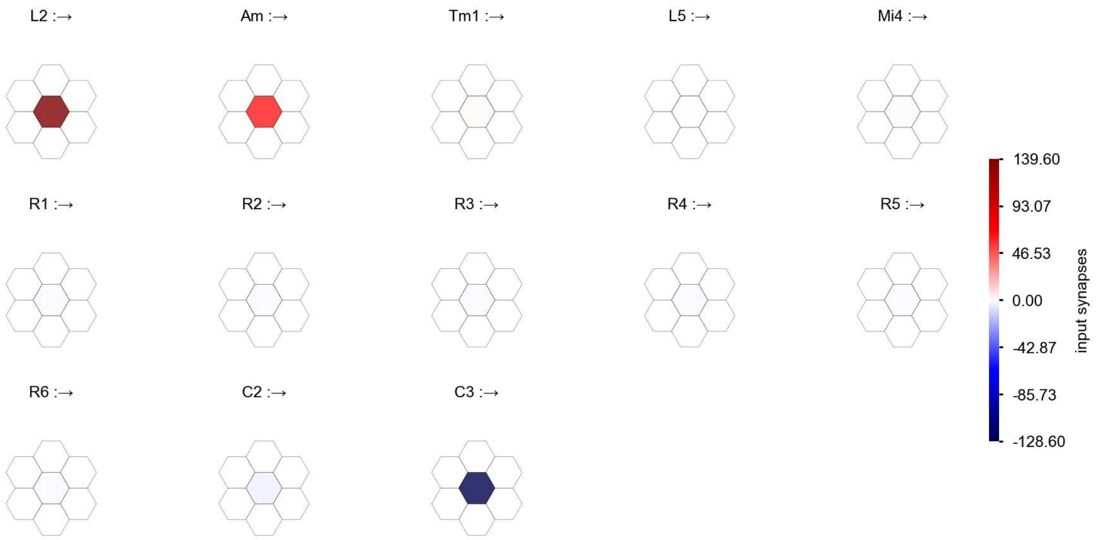

T1 - Figure 1: **Anatomical receptive fields.** Each colored hexagon is a projection, with the projection strength characterized by the average number of synapses that we count from the EM reconstruction. Red indicates excitatory synapses, blue indicates inhibitory synapses from inferred signs. Filters in the order of their total number of synapses.

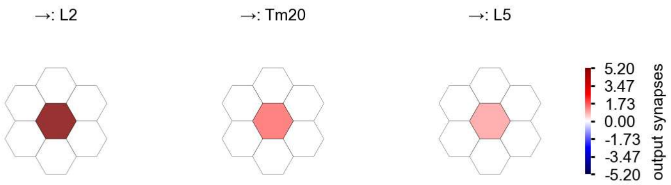

T1 - Figure 2: **Anatomical projective fields.** Each colored hexagon is a projection, with the projection strength characterized by the average number of synapses that we count from the EM reconstruction. Red indicates excitatory synapses, blue indicates inhibitory synapses from inferred signs. Filters in the order of their total number of synapses.

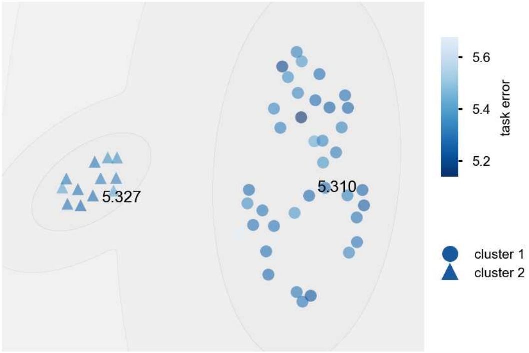

T1 - Figure 3: **Clustering of the responses to naturalistic stimuli.** From responses to naturalistic scenes from Sintel. Scatterpoints represent individual models colored by their task error.

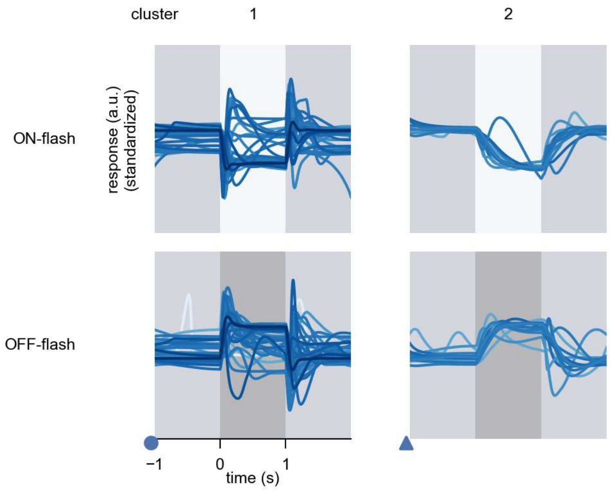

T1 - Figure 4: **Responses to flashes.** The top row shows responses to ON-flashes, the bottom row shows responses to OFF-flashes. The responses from the different models in the different clusters (columns) overlay and are colored in the respective task error of the model. The circular flashes occur at second zero, cover 6 ommatidia in radius, and last for one second. Before and after, a grey-stimulus leads to a stationary state of the network. The traces are standardized.

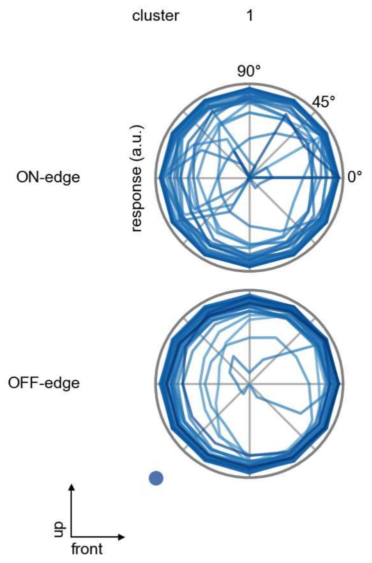

T1 - Figure 5: **Peak responses to moving edges.** The top row shows normalized peak responses to moving ON-edges, the bottom row shows normalized peak responses to moving OFF-edges. The edge-stimuli move in different directions from 0 to 360 degrees. Before, a grey-stimulus led the network to a stationary state. The responses from the different models in the different clusters (columns) overlay and are colored in the respective task error of the model. The peak responses are averaged over all assessed edge-speeds.

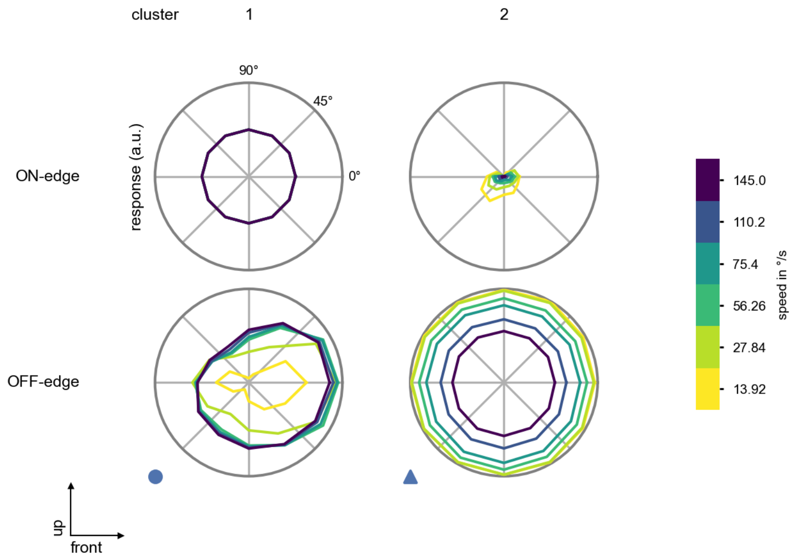

T1 - Figure 6: **Peak responses to moving edges from best-performing models.** The top row shows peak responses to moving ON-edges, the bottom row shows peak responses to moving OFF-edges. The edge-stimuli move in different directions from 0 to 360 degrees and at different speeds. Before, a grey-stimulus led the network to a stationary state. The columns show responses from the best-task-performing model in the respective cluster.

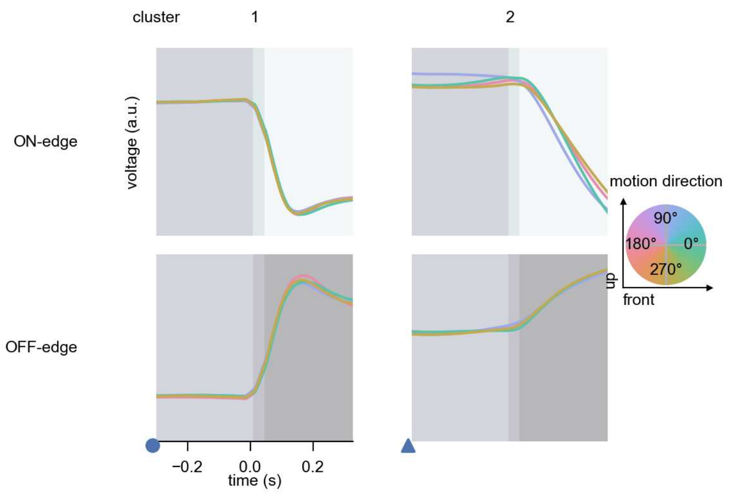

T1 - Figure 7: **Responses to moving edges from best-performing models.** The top row shows responses to moving ON-edges, the bottom row shows peak responses to moving OFF-edges. The edge-stimuli move in different directions from 0 to 360 degrees and at different speeds. Before, a grey-stimulus led the network to a stationary state. The columns show responses from the best-task-performing model in the respective cluster. Example responses to edges moving at 75.4°/s in all cardinal directions (green 0°, blue 90°, red 180°, yellow 270°) from -22.5 to 22.5° visual angle. Background contour indicates the edge, passing by the central column at second zero.

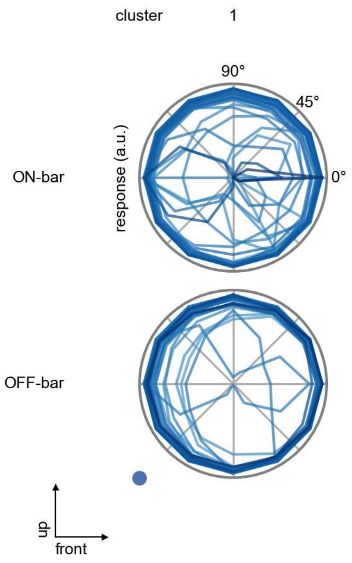

T1 - Figure 8: **Peak responses to moving bars.** The top row shows normalized peak responses to moving ON-bars, the bottom row shows normalized peak responses to moving OFF-bars. The bar-stimuli move in different directions from 0 to 360 degrees. Before, a grey-stimulus led the network to a stationary state. The responses from the different models in the different clusters (columns) overlay and are colored in the respective task error of the model. The peak responses are averaged over all assessed bar-speeds.

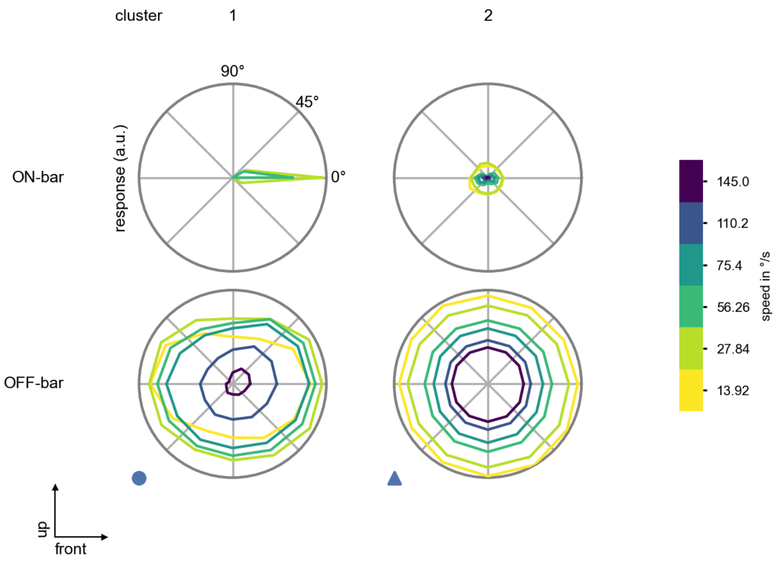

T1 - Figure 9: **Peak responses to moving bars from best-performing models.** The top row shows peak responses to moving ON-bars, the bottom row shows peak responses to moving OFF-bars. The bar-stimuli move in different directions from 0 to 360 degrees and at different speeds. Before, a grey-stimulus led the network to a stationary state. The columns show responses from the best-task-performing model in the respective cluster.

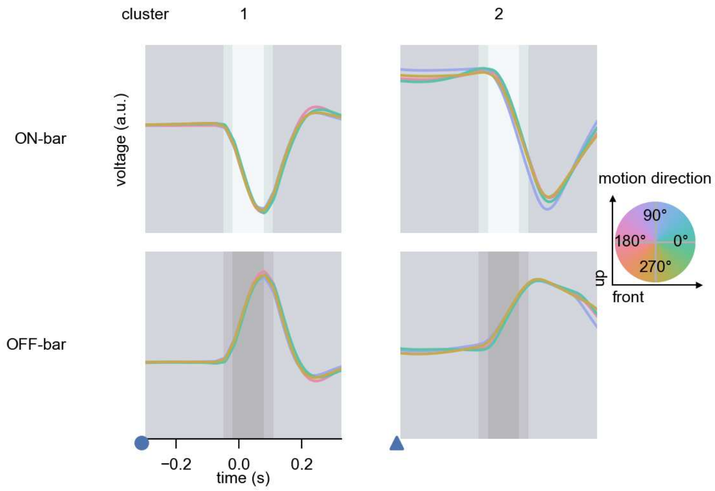

T1 - Figure 10: **Responses to moving bars from best-performing models.** The top row shows responses to moving ON-bars, the bottom row shows peak responses to moving OFF-bars. The bar-stimuli move in different directions from 0 to 360 degrees and at different speeds. Before, a grey-stimulus led the network to a stationary state. The columns show responses from the best-task-performing model in the respective cluster. Example responses to bars moving at 75.4°/s in all cardinal directions (green 0°, blue 90°, red 180°, yellow 270°) from -22.5 to 22.5° visual angle. Background contour indicates the bar, passing by the central column at second zero.

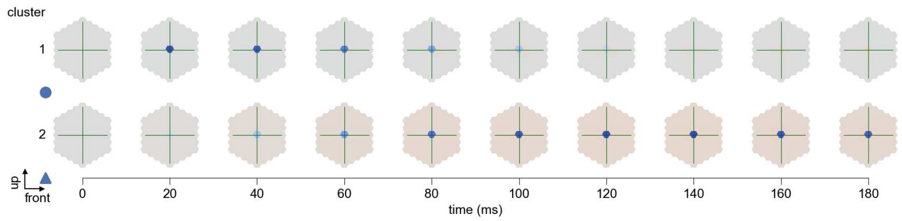

T1 - Figure 11: **Spatio-temporal receptive field.** Responses of the central cell to ON-impulses at single ommatidium impulse locations. The impulse occurs at second zero. Before, a grey-stimulus led the network to a stationary state. Responses are baseline-subtracted. Each row illustrates responses from the best-performing model of the respective cluster. Red indicated depolarization, blue indicates hyperpolarization.

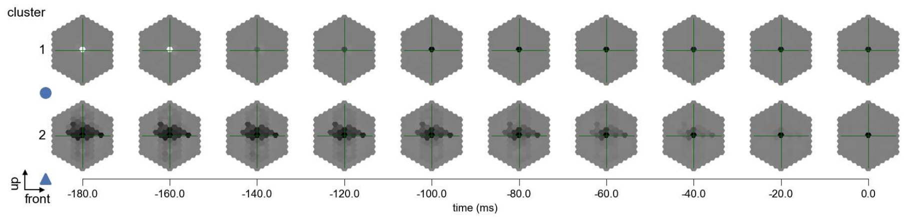

T1 - Figure 12: **Maximally excitatory stimuli.** Each row presents the regularized naturalistic-stimulus from the Sintel dataset that maximizes the cell type's central column response at second zero in the best-performing model of the respective cluster.

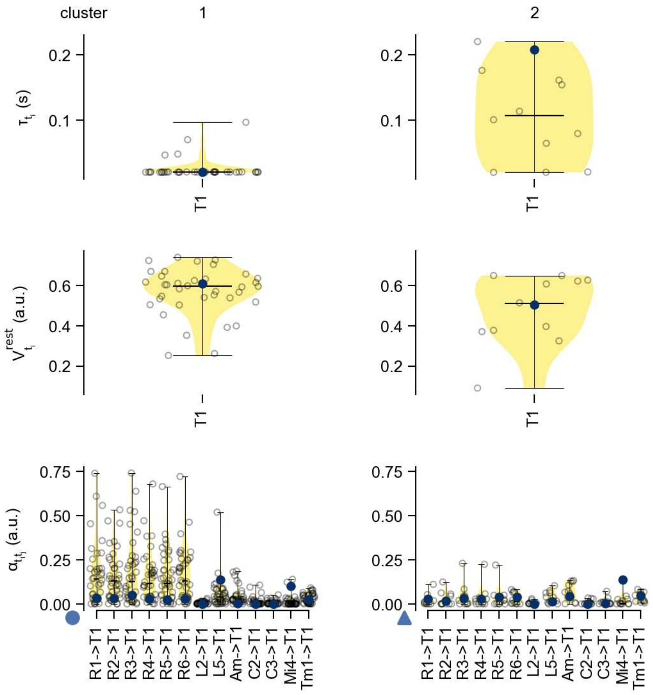

T1 - Figure 13: **Task-constrained parameters.** Each column shows the parameters inferred within the respective cluster. First row: learned time constants of the cell type. Second row: resting potentials of the cell type. Third row: scaling factors for the convolutional filters. The blue scatter represents the parameters from the best-performing model within the cluster.

# 33 T2

← Cell types

#### Figures

1 Anatomical receptive fields. 216
2 Anatomical projective fields. 217
3 Clustering of the responses to naturalistic stimuli. 217
4 Responses to flashes. 217
5 Peak responses to moving edges. 218
6 Peak responses to moving edges from best-performing models. 218
7 Responses to moving edges from best-performing models. 219
8 Peak responses to moving bars. 219
9 Peak responses to moving bars from best-performing models. 220
10 Responses to moving bars from best-performing models. 220
11 Spatio-temporal receptive field. 221
12 Maximally excitatory stimuli. 221
13 Task-constrained parameters. 222

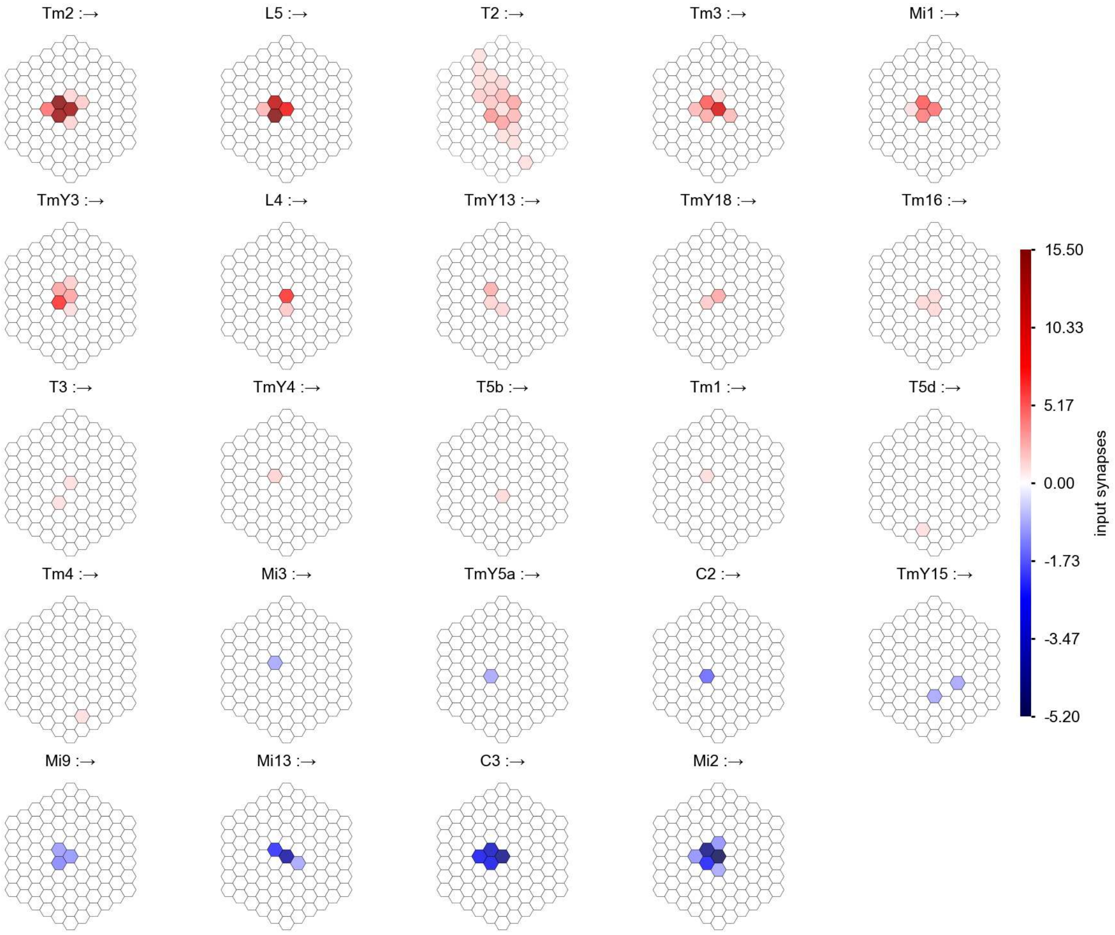

T2 - Figure 1: **Anatomical receptive fields.** Each colored hexagon is a projection, with the projection strength characterized by the average number of synapses that we count from the EM reconstruction. Red indicates excitatory synapses, blue indicates inhibitory synapses from inferred signs. Filters in the order of their total number of synapses.

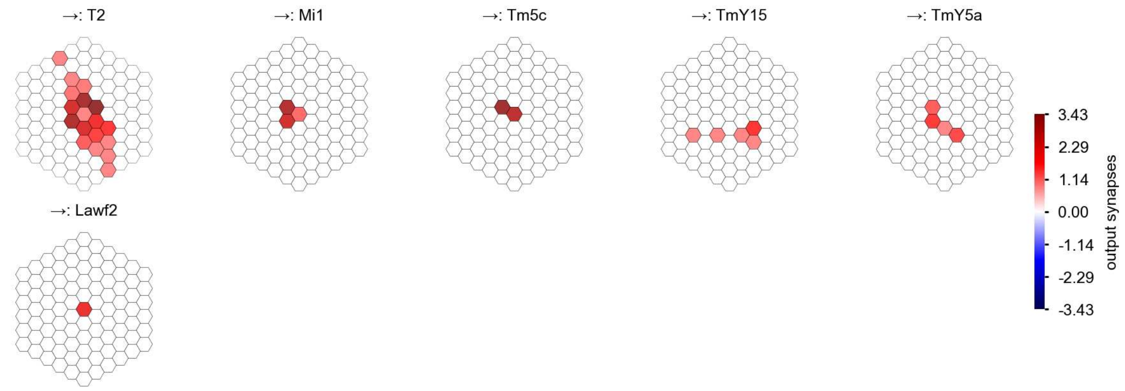

T2 - Figure 2: **Anatomical projective fields.** Each colored hexagon is a projection, with the projection strength characterized by the average number of synapses that we count from the EM reconstruction. Red indicates excitatory synapses, blue indicates inhibitory synapses from inferred signs. Filters in the order of their total number of synapses.

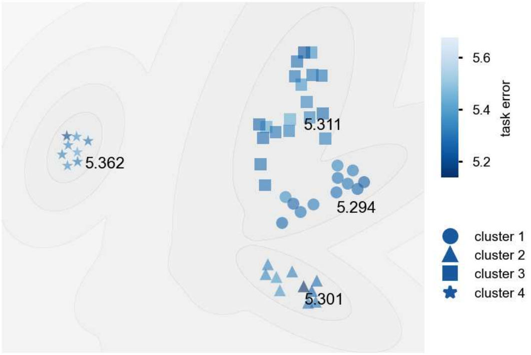

T2 - Figure 3: **Clustering of the responses to naturalistic stimuli.** From responses to naturalistic scenes from Sintel. Scatterpoints represent individual models colored by their task error.

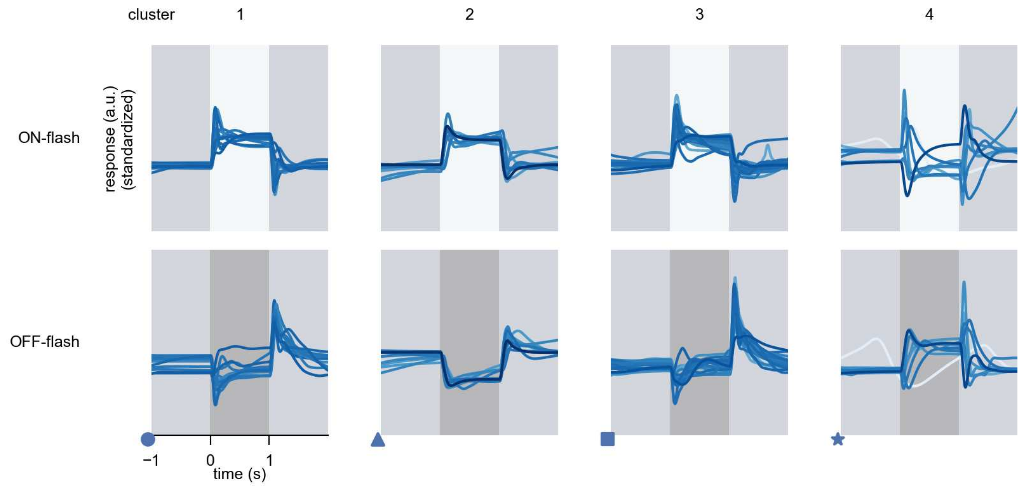

T2 - Figure 4: **Responses to flashes.** The top row shows responses to ON-flashes, the bottom row shows responses to OFF-flashes. The responses from the different models in the different clusters (columns) overlay and are colored in the respective task error of the model. The circular flashes occur at second zero, cover 6 ommatidia in radius, and last for one second. Before and after, a grey-stimulus leads to a stationary state of the network. The traces are standardized.

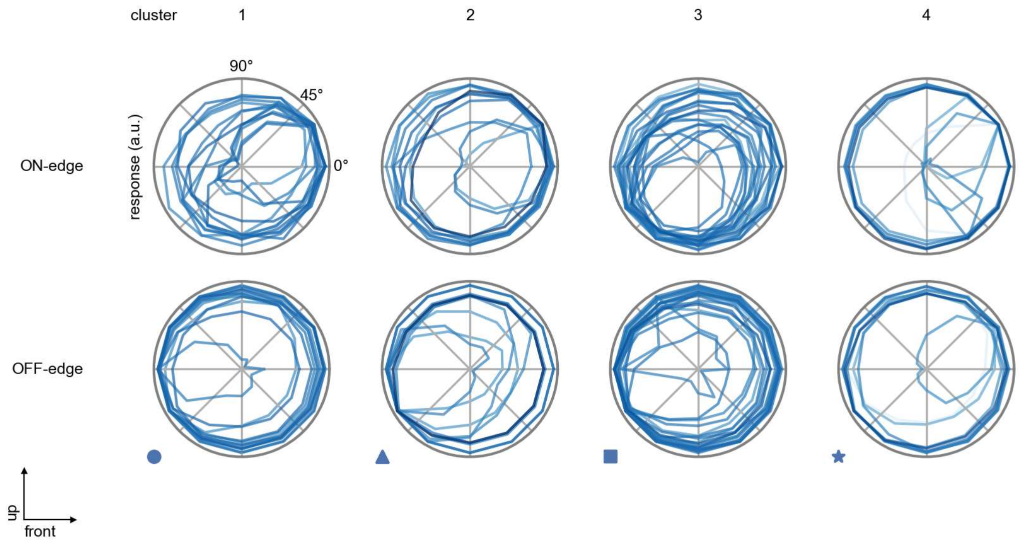

T2 - Figure 5: **Peak responses to moving edges.** The top row shows normalized peak responses to moving ON-edges, the bottom row shows normalized peak responses to moving OFF-edges. The edge-stimuli move in different directions from 0 to 360 degrees. Before, a grey-stimulus led the network to a stationary state. The responses from the different models in the different clusters (columns) overlay and are colored in the respective task error of the model. The peak responses are averaged over all assessed edge-speeds.

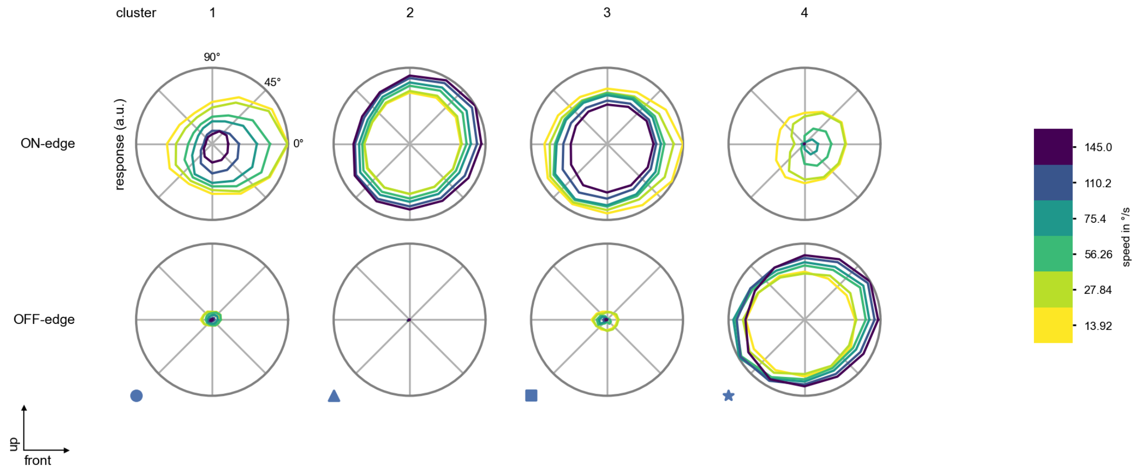

T2 - Figure 6: **Peak responses to moving edges from best-performing models.** The top row shows peak responses to moving ON-edges, the bottom row shows peak responses to moving OFF-edges. The edge-stimuli move in different directions from 0 to 360 degrees and at different speeds. Before, a grey-stimulus led the network to a stationary state. The columns show responses from the best-task-performing model in the respective cluster.

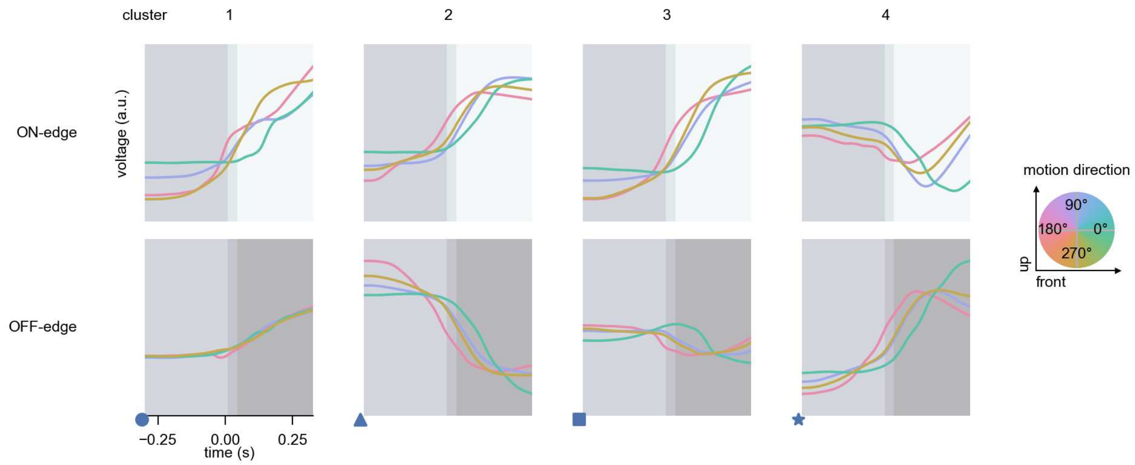

T2 - Figure 7: **Responses to moving edges from best-performing models.** The top row shows responses to moving ON-edges, the bottom row shows peak responses to moving OFF-edges. The edge-stimuli move in different directions from 0 to 360 degrees and at different speeds. Before, a grey-stimulus led the network to a stationary state. The columns show responses from the best-task-performing model in the respective cluster. Example responses to edges moving at 75.4°/s in all cardinal directions (green 0°, blue 90°, red 180°, yellow 270°) from -22.5 to 22.5° visual angle. Background contour indicates the edge, passing by the central column at second zero.

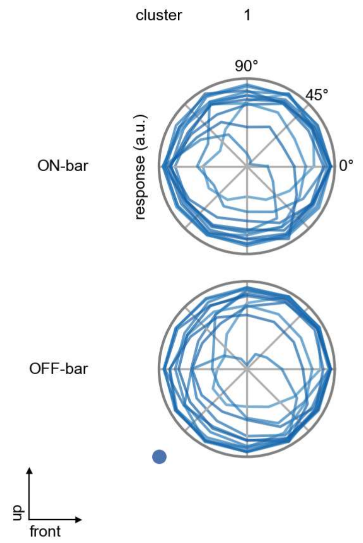

T2 - Figure 8: **Peak responses to moving bars.** The top row shows normalized peak responses to moving ON-bars, the bottom row shows normalized peak responses to moving OFF-bars. The bar-stimuli move in different directions from 0 to 360 degrees. Before, a grey-stimulus led the network to a stationary state. The responses from the different models in the different clusters (columns) overlay and are colored in the respective task error of the model. The peak responses are averaged over all assessed bar-speeds.

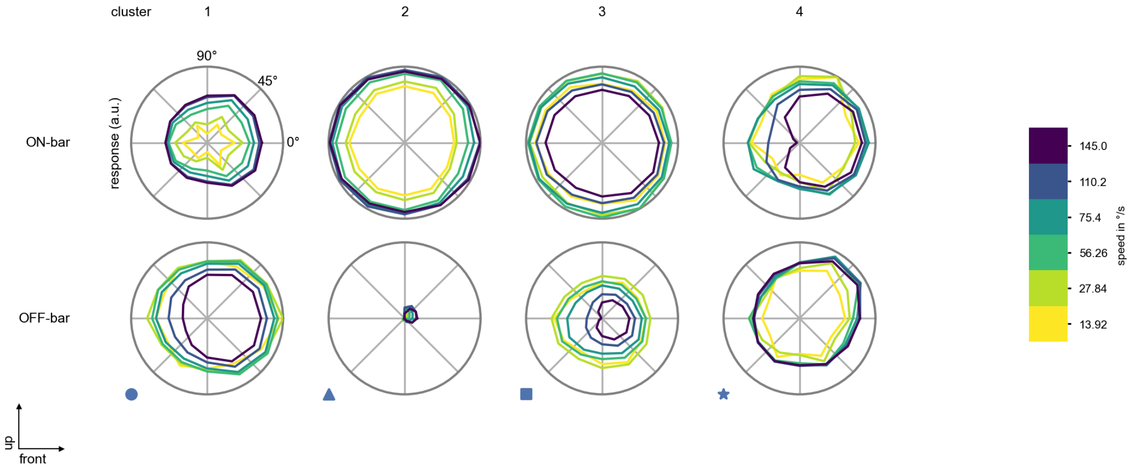

T2 - Figure 9: **Peak responses to moving bars from best-performing models.** The top row shows peak responses to moving ON-bars, the bottom row shows peak responses to moving OFF-bars. The bar-stimuli move in different directions from 0 to 360 degrees and at different speeds. Before, a grey-stimulus led the network to a stationary state. The columns show responses from the best-task-performing model in the respective cluster.

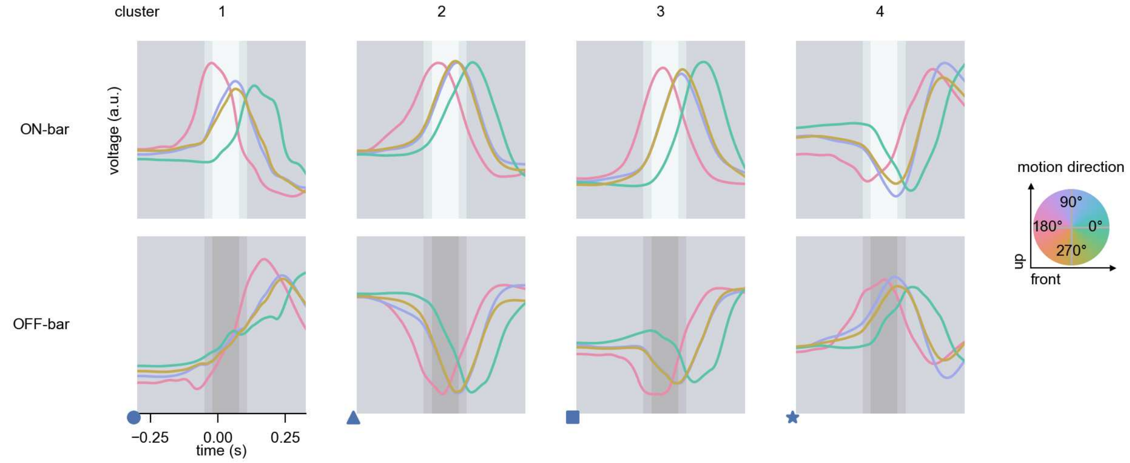

T2 - Figure 10: **Responses to moving bars from best-performing models.** The top row shows responses to moving ON-bars, the bottom row shows peak responses to moving OFF-bars. The bar-stimuli move in different directions from 0 to 360 degrees and at different speeds. Before, a grey-stimulus led the network to a stationary state. The columns show responses from the best-task-performing model in the respective cluster. Example responses to bars moving at 75.4°/s in all cardinal directions (green 0°, blue 90°, red 180°, yellow 270°) from -22.5 to 22.5° visual angle. Background contour indicates the bar, passing by the central column at second zero.

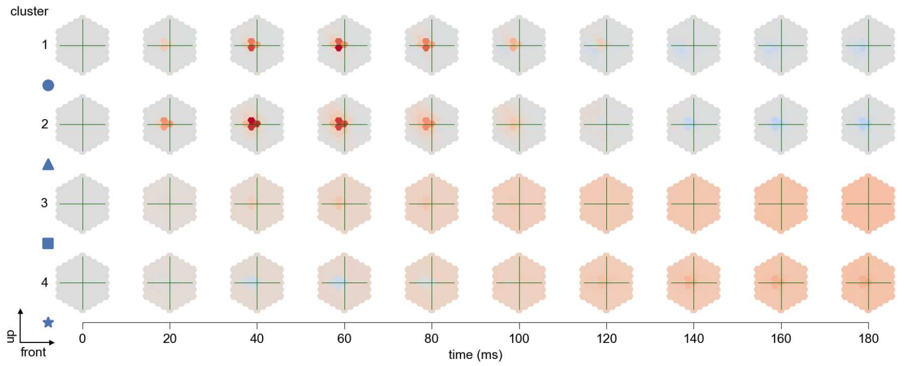

T2 - Figure 11: **Spatio-temporal receptive field.** Responses of the central cell to ON-impulses at single ommatidium impulse locations. The impulse occurs at second zero. Before, a grey-stimulus led the network to a stationary state. Responses are baseline-subtracted. Each row illustrates responses from the best-performing model of the respective cluster. Red indicated depolarization, blue indicates hyperpolarization.

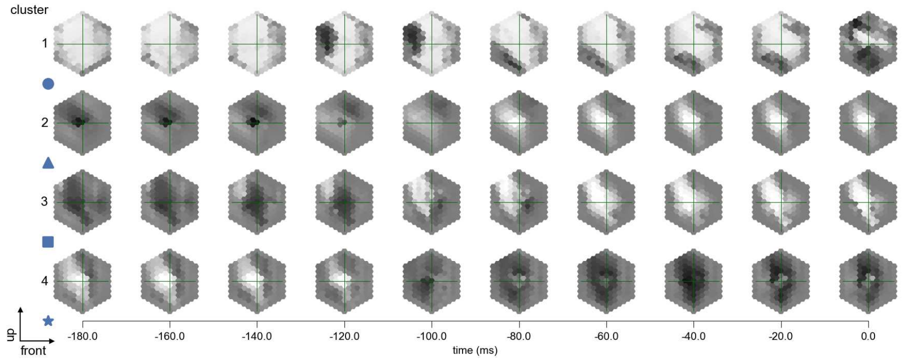

T2 - Figure 12: **Maximally excitatory stimuli.** Each row presents the regularized naturalistic-stimulus from the Sintel dataset that maximizes the cell type's central column response at second zero in the best-performing model of the respective cluster.

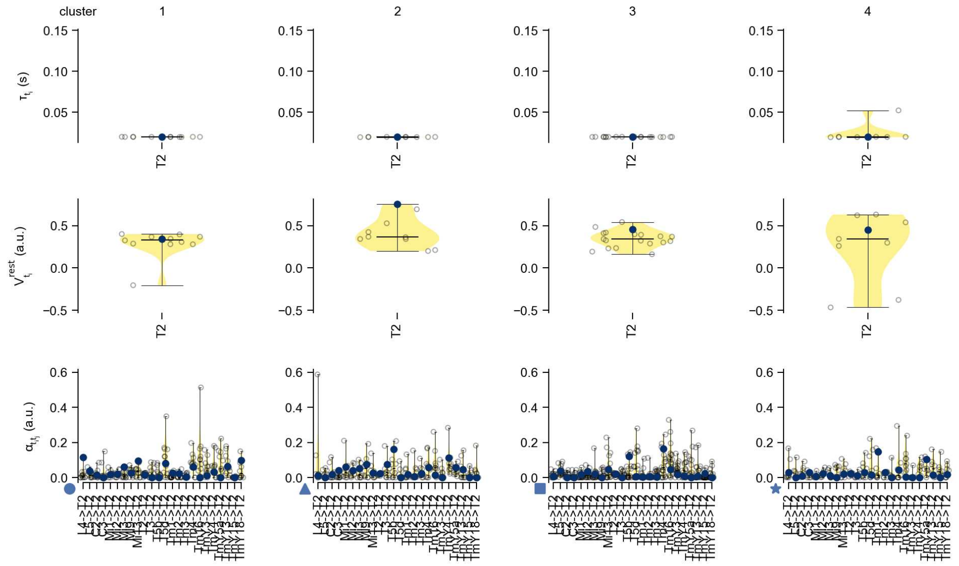

T2 - Figure 13: **Task-constrained parameters.** Each column shows the parameters inferred within the respective cluster. First row: learned time constants of the cell type. Second row: resting potentials of the cell type. Third row: scaling factors for the convolutional filters. The blue scatter represents the parameters from the best-performing model within the cluster.

# 34 T2a

← Cell types

#### Figures

1 Anatomical receptive fields. . . . 223
2 Anatomical projective fields. . . . 224
3 Clustering of the responses to naturalistic stimuli. . . . 224
4 Responses to flashes. . . . 224
5 Peak responses to moving edges. . . . 225
6 Peak responses to moving edges from best-performing models. . . . 225
7 Responses to moving edges from best-performing models. . . . 226
8 Peak responses to moving bars. . . . 226
9 Peak responses to moving bars from best-performing models. . . . 227
10 Responses to moving bars from best-performing models. . . . 227
11 Spatio-temporal receptive field. . . . 228
12 Maximally excitatory stimuli. . . . 228
13 Task-constrained parameters. . . . 229

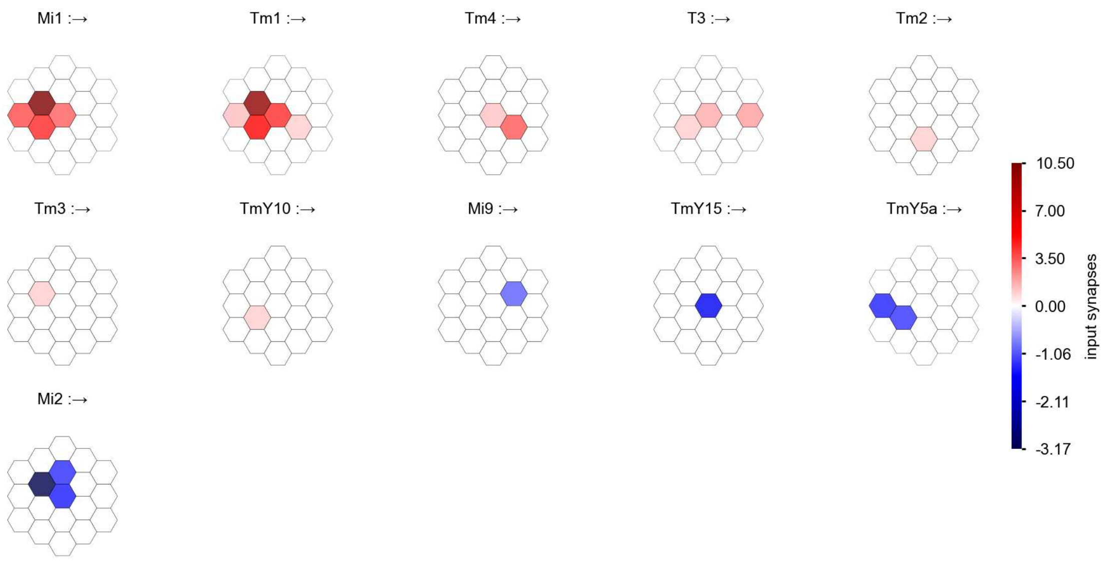

T2a - Figure 1: **Anatomical receptive fields.** Each colored hexagon is a projection, with the projection strength characterized by the average number of synapses that we count from the EM reconstruction. Red indicates excitatory synapses, blue indicates inhibitory synapses from inferred signs. Filters in the order of their total number of synapses.

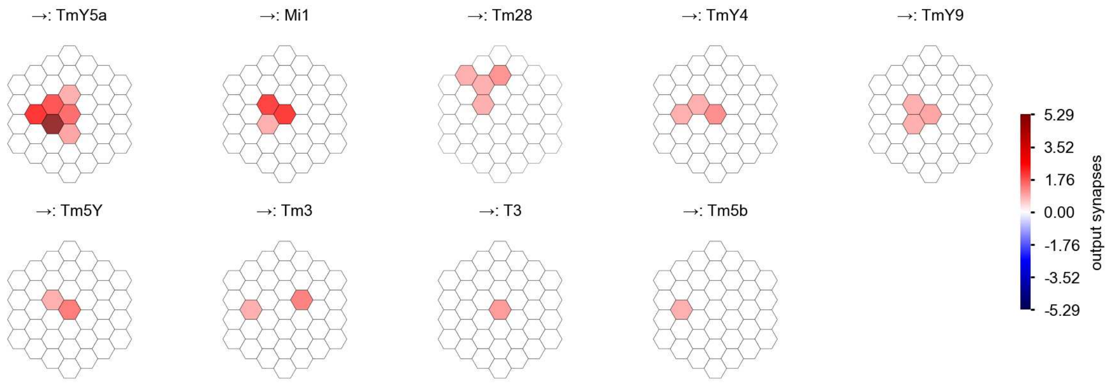

T2a - Figure 2: **Anatomical projective fields.** Each colored hexagon is a projection, with the projection strength characterized by the average number of synapses that we count from the EM reconstruction. Red indicates excitatory synapses, blue indicates inhibitory synapses from inferred signs. Filters in the order of their total number of synapses.

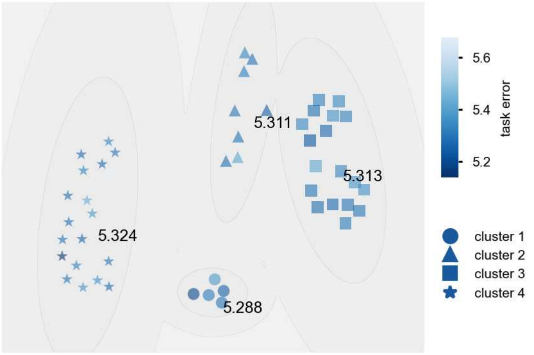

T2a - Figure 3: **Clustering of the responses to naturalistic stimuli.** From responses to naturalistic scenes from Sintel. Scatterpoints represent individual models colored by their task error.

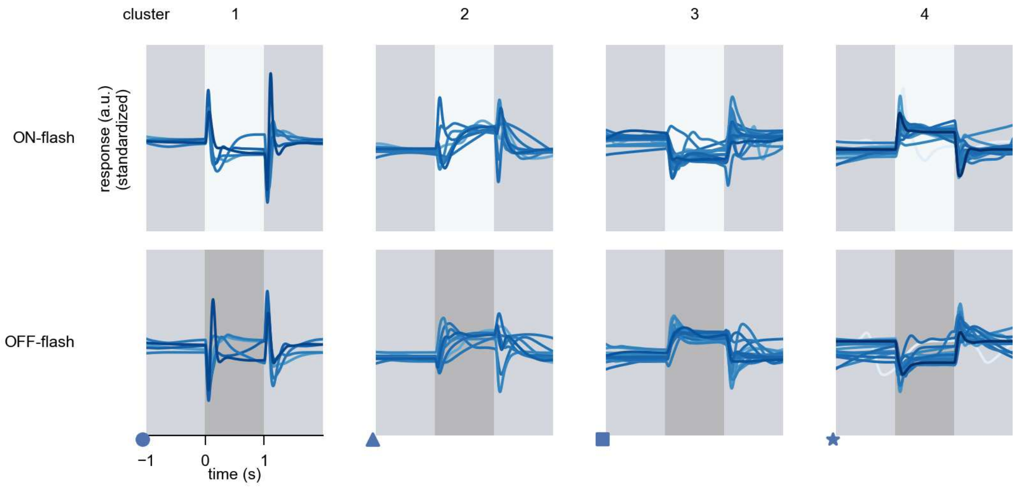

T2a - Figure 4: **Responses to flashes.** The top row shows responses to ON-flashes, the bottom row shows responses to OFF-flashes. The responses from the different models in the different clusters (columns) overlay and are colored in the respective task error of the model. The circular flashes occur at second zero, cover 6 ommatidia in radius, and last for one second. Before and after, a grey-stimulus leads to a stationary state of the network. The traces are standardized.

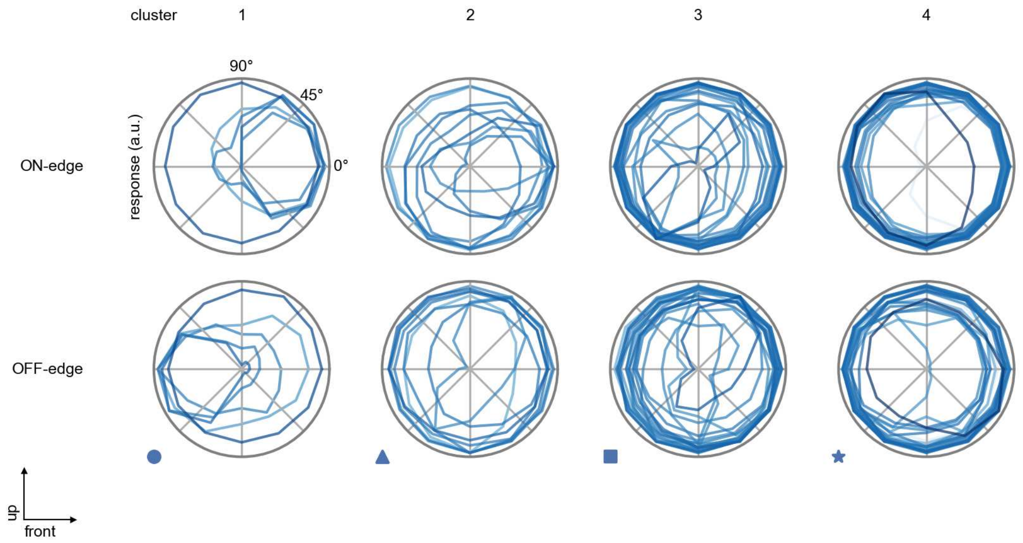

T2a - Figure 5: **Peak responses to moving edges.** The top row shows normalized peak responses to moving ON-edges, the bottom row shows normalized peak responses to moving OFF-edges. The edge-stimuli move in different directions from 0 to 360 degrees. Before, a grey-stimulus led the network to a stationary state. The responses from the different models in the different clusters (columns) overlay and are colored in the respective task error of the model. The peak responses are averaged over all assessed edge-speeds.

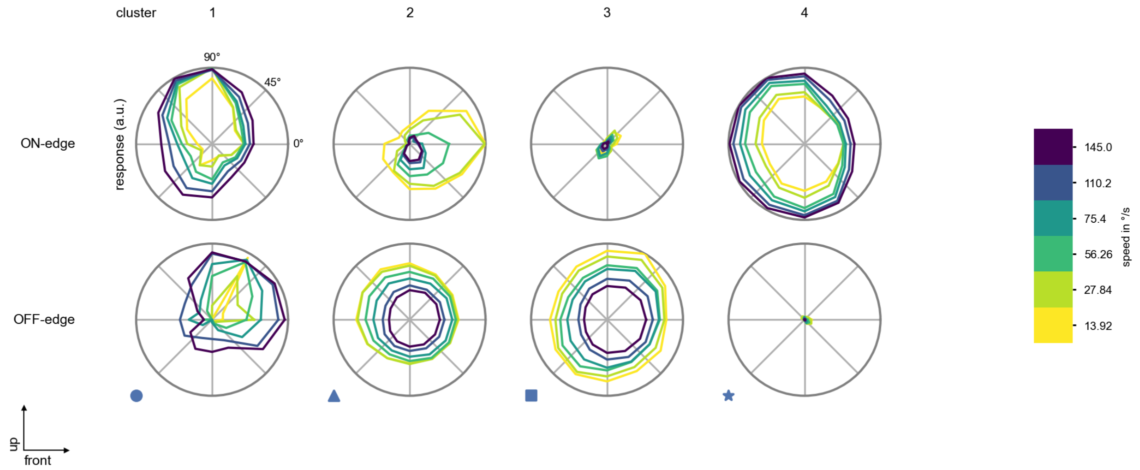

T2a - Figure 6: **Peak responses to moving edges from best-performing models.** The top row shows peak responses to moving ON-edges, the bottom row shows peak responses to moving OFF-edges. The edge-stimuli move in different directions from 0 to 360 degrees and at different speeds. Before, a grey-stimulus led the network to a stationary state. The columns show responses from the best-task-performing model in the respective cluster.

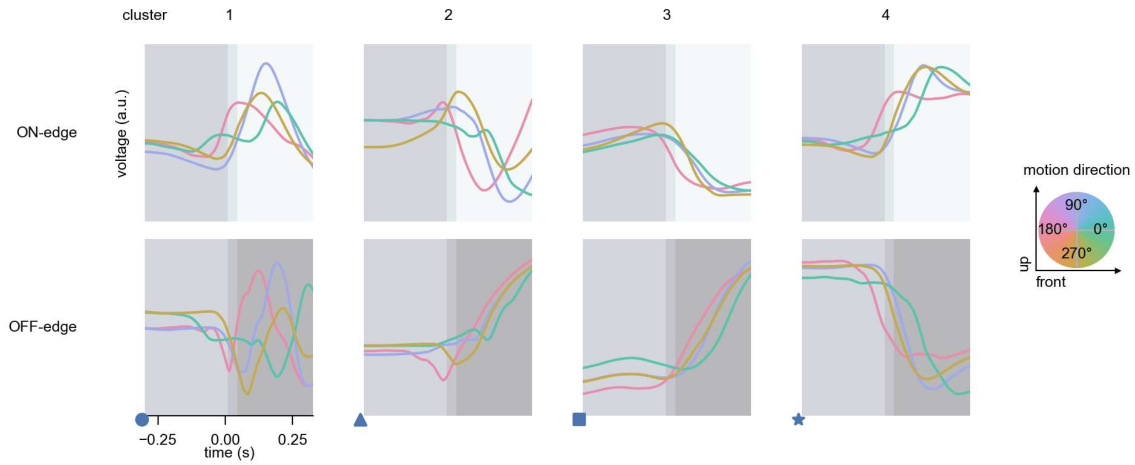

T2a - Figure 7: **Responses to moving edges from best-performing models.** The top row shows responses to moving ON-edges, the bottom row shows peak responses to moving OFF-edges. The edge-stimuli move in different directions from 0 to 360 degrees and at different speeds. Before, a grey-stimulus led the network to a stationary state. The columns show responses from the best-task-performing model in the respective cluster. Example responses to edges moving at 75.4°/s in all cardinal directions (green 0°, blue 90°, red 180°, yellow 270°) from -22.5 to 22.5° visual angle. Background contour indicates the edge, passing by the central column at second zero.

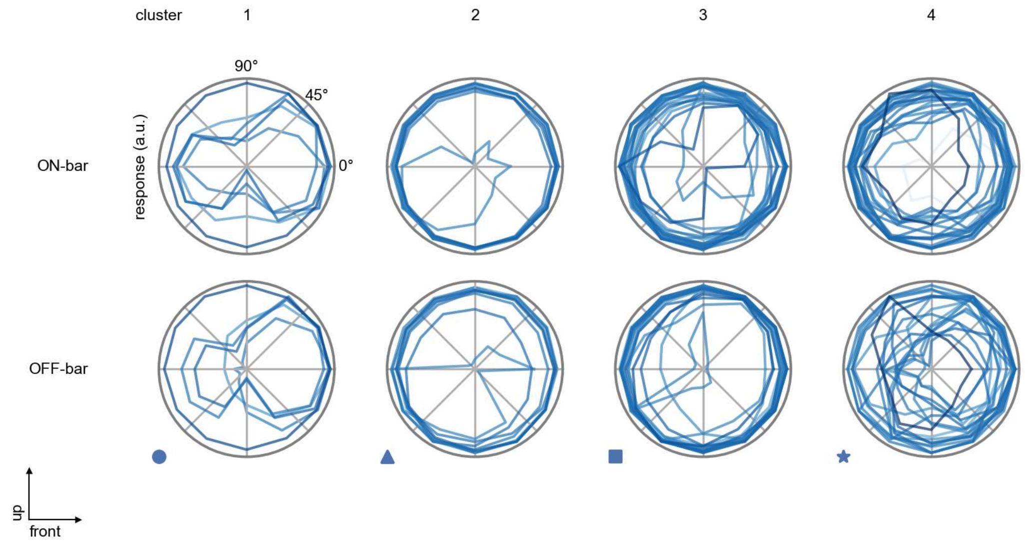

T2a - Figure 8: **Peak responses to moving bars.** The top row shows normalized peak responses to moving ON-bars, the bottom row shows normalized peak responses to moving OFF-bars. The bar-stimuli move in different directions from 0 to 360 degrees. Before, a grey-stimulus led the network to a stationary state. The responses from the different models in the different clusters (columns) overlay and are colored in the respective task error of the model. The peak responses are averaged over all assessed bar-speeds.

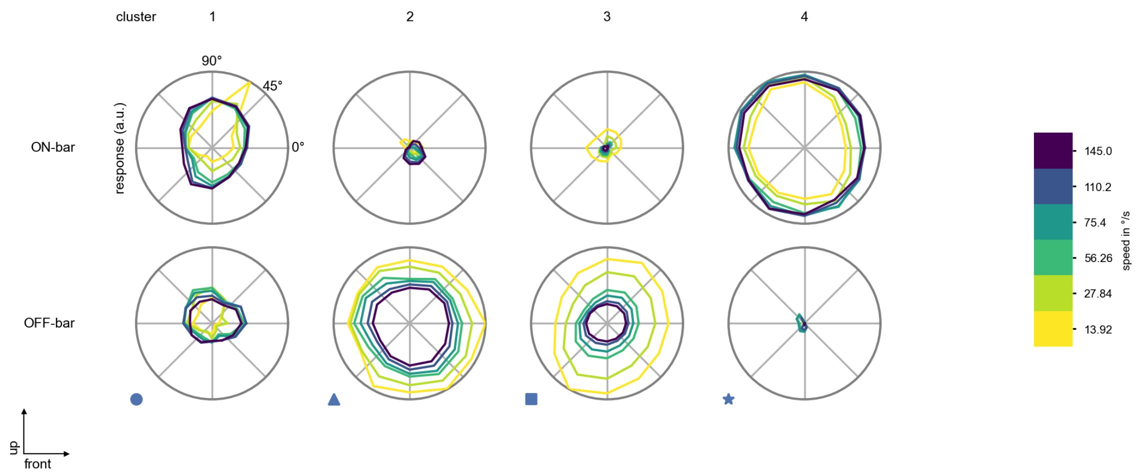

T2a - Figure 9: **Peak responses to moving bars from best-performing models.** The top row shows peak responses to moving ON-bars, the bottom row shows peak responses to moving OFF-bars. The bar-stimuli move in different directions from 0 to 360 degrees and at different speeds. Before, a grey-stimulus led the network to a stationary state. The columns show responses from the best-task-performing model in the respective cluster.

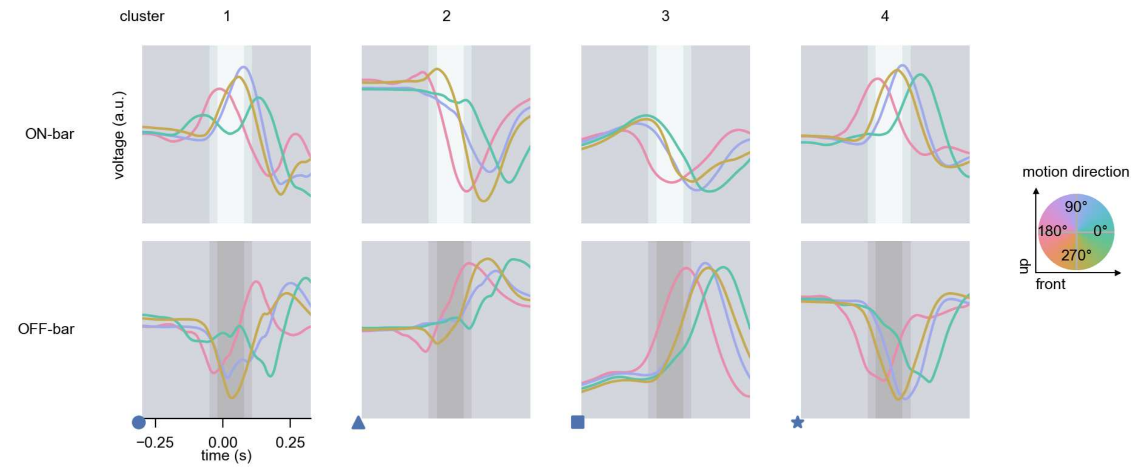

T2a - Figure 10: **Responses to moving bars from best-performing models.** The top row shows responses to moving ON-bars, the bottom row shows peak responses to moving OFF-bars. The bar-stimuli move in different directions from 0 to 360 degrees and at different speeds. Before, a grey-stimulus led the network to a stationary state. The columns show responses from the best-task-performing model in the respective cluster. Example responses to bars moving at 75.4°/s in all cardinal directions (green 0°, blue 90°, red 180°, yellow 270°) from -22.5 to 22.5° visual angle. Background contour indicates the bar, passing by the central column at second zero.

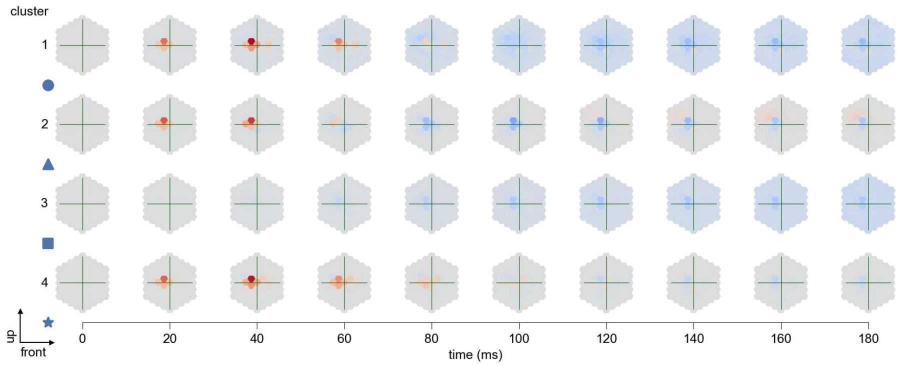

T2a - Figure 11: **Spatio-temporal receptive field.** Responses of the central cell to ON-impulses at single ommatidium impulse locations. The impulse occurs at second zero. Before, a grey-stimulus led the network to a stationary state. Responses are baseline-subtracted. Each row illustrates responses from the best-performing model of the respective cluster. Red indicated depolarization, blue indicates hyperpolarization.

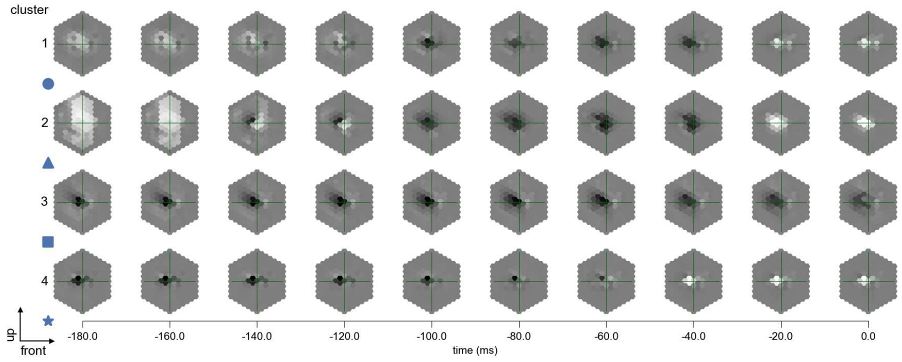

T2a - Figure 12: **Maximally excitatory stimuli.** Each row presents the regularized naturalistic-stimulus from the Sintel dataset that maximizes the cell type's central column response at second zero in the best-performing model of the respective cluster.

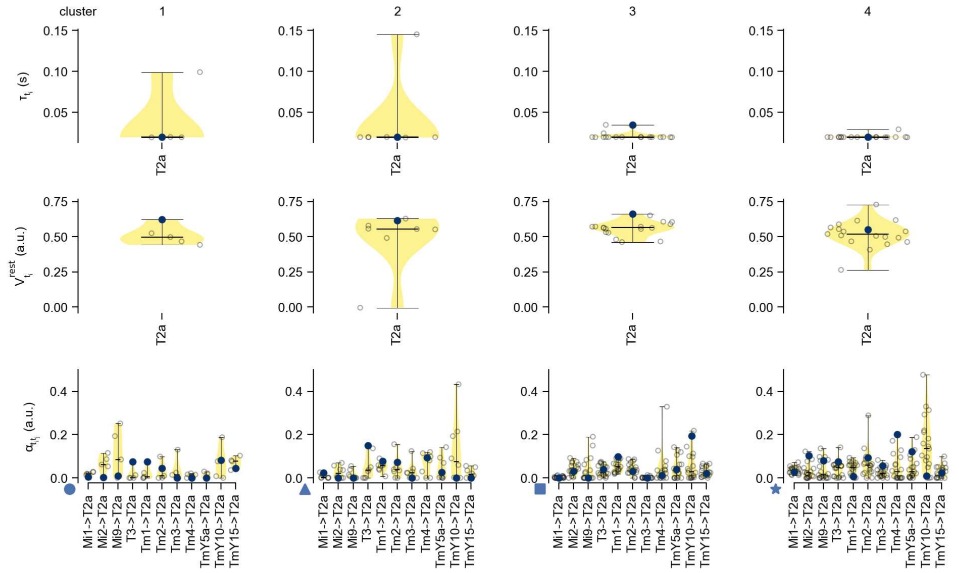

T2a - Figure 13: **Task-constrained parameters.** Each column shows the parameters inferred within the respective cluster. First row: learned time constants of the cell type. Second row: resting potentials of the cell type. Third row: scaling factors for the convolutional filters. The blue scatter represents the parameters from the best-performing model within the cluster.

# 35 T3

← Cell types

#### Figures

1 Anatomical receptive fields. . . . . 230
2 Anatomical projective fields. . . . . 230
3 Clustering of the responses to naturalistic stimuli. . . . . 231
4 Responses to flashes. . . . . 231
5 Peak responses to moving edges. . . . . 232
6 Peak responses to moving edges from best-performing models. . . . . 232
7 Responses to moving edges from best-performing models. . . . . 233
8 Peak responses to moving bars. . . . . 233
9 Peak responses to moving bars from best-performing models. . . . . 234
10 Responses to moving bars from best-performing models. . . . . 234
11 Spatio-temporal receptive field. . . . . 235
12 Maximally excitatory stimuli. . . . . 235
13 Task-constrained parameters. . . . . 235

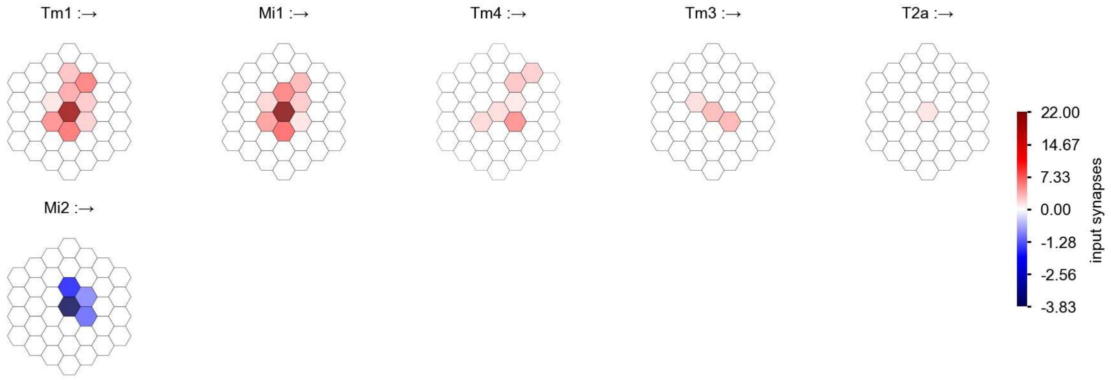

T3 - Figure 1: **Anatomical receptive fields.** Each colored hexagon is a projection, with the projection strength characterized by the average number of synapses that we count from the EM reconstruction. Red indicates excitatory synapses, blue indicates inhibitory synapses from inferred signs. Filters in the order of their total number of synapses.

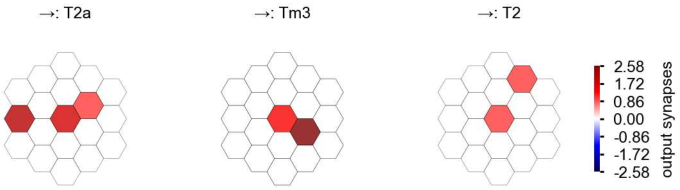

T3 - Figure 2: **Anatomical projective fields.** Each colored hexagon is a projection, with the projection strength characterized by the average number of synapses that we count from the EM reconstruction. Red indicates excitatory synapses, blue indicates inhibitory synapses from inferred signs. Filters in the order of their total number of synapses.

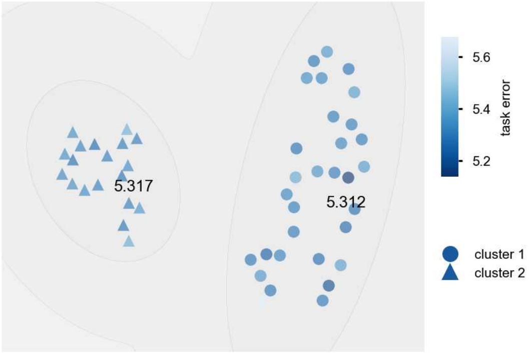

T3 - Figure 3: **Clustering of the responses to naturalistic stimuli.** From responses to naturalistic scenes from Sintel. Scatterpoints represent individual models colored by their task error.

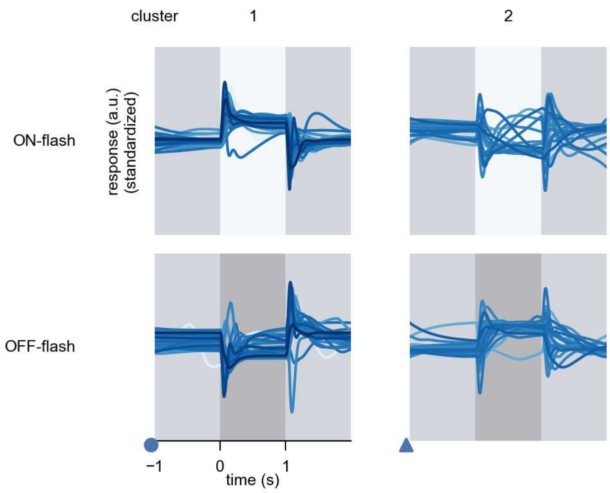

T3 - Figure 4: **Responses to flashes.** The top row shows responses to ON-flashes, the bottom row shows responses to OFF-flashes. The responses from the different models in the different clusters (columns) overlay and are colored in the respective task error of the model. The circular flashes occur at second zero, cover 6 ommatidia in radius, and last for one second. Before and after, a grey-stimulus leads to a stationary state of the network. The traces are standardized.

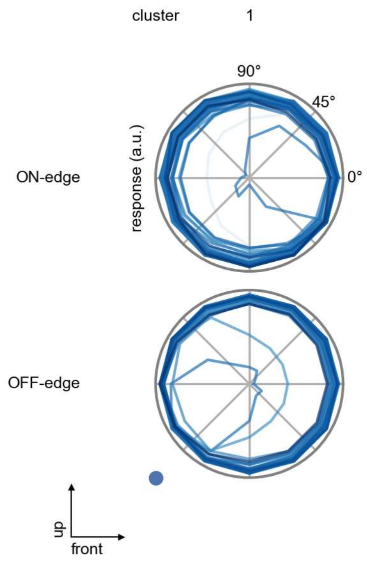

T3 - Figure 5: **Peak responses to moving edges.** The top row shows normalized peak responses to moving ON-edges, the bottom row shows normalized peak responses to moving OFF-edges. The edge-stimuli move in different directions from 0 to 360 degrees. Before, a grey-stimulus led the network to a stationary state. The responses from the different models in the different clusters (columns) overlay and are colored in the respective task error of the model. The peak responses are averaged over all assessed edge-speeds.

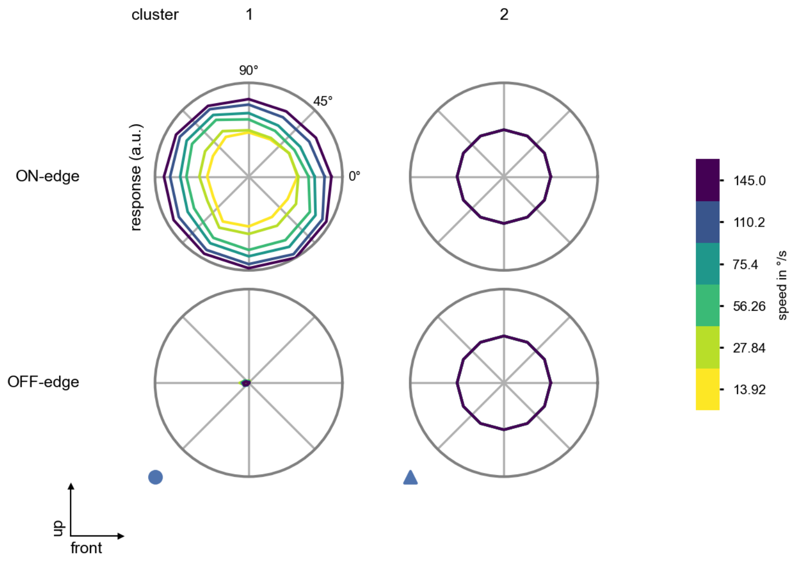

T3 - Figure 6: **Peak responses to moving edges from best-performing models.** The top row shows peak responses to moving ON-edges, the bottom row shows peak responses to moving OFF-edges. The edge-stimuli move in different directions from 0 to 360 degrees and at different speeds. Before, a grey-stimulus led the network to a stationary state. The columns show responses from the best-task-performing model in the respective cluster.

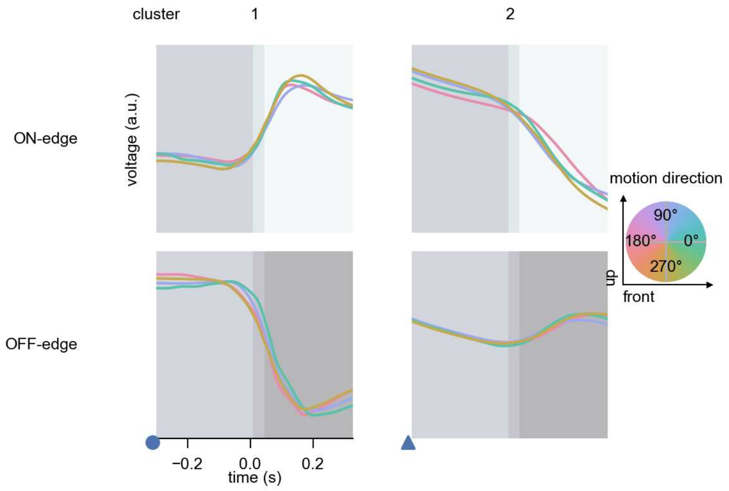

T3 - Figure 7: **Responses to moving edges from best-performing models.** The top row shows responses to moving ON-edges, the bottom row shows peak responses to moving OFF-edges. The edge-stimuli move in different directions from 0 to 360 degrees and at different speeds. Before, a grey-stimulus led the network to a stationary state. The columns show responses from the best-task-performing model in the respective cluster. Example responses to edges moving at 75.4°/s in all cardinal directions (green 0°, blue 90°, red 180°, yellow 270°) from -22.5 to 22.5° visual angle. Background contour indicates the edge, passing by the central column at second zero.

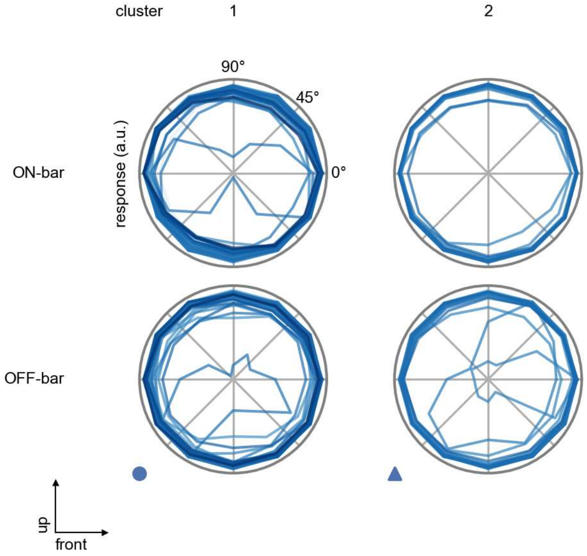

T3 - Figure 8: **Peak responses to moving bars.** The top row shows normalized peak responses to moving ON-bars, the bottom row shows normalized peak responses to moving OFF-bars. The bar-stimuli move in different directions from 0 to 360 degrees. Before, a grey-stimulus led the network to a stationary state. The responses from the different models in the different clusters (columns) overlay and are colored in the respective task error of the model. The peak responses are averaged over all assessed bar-speeds.

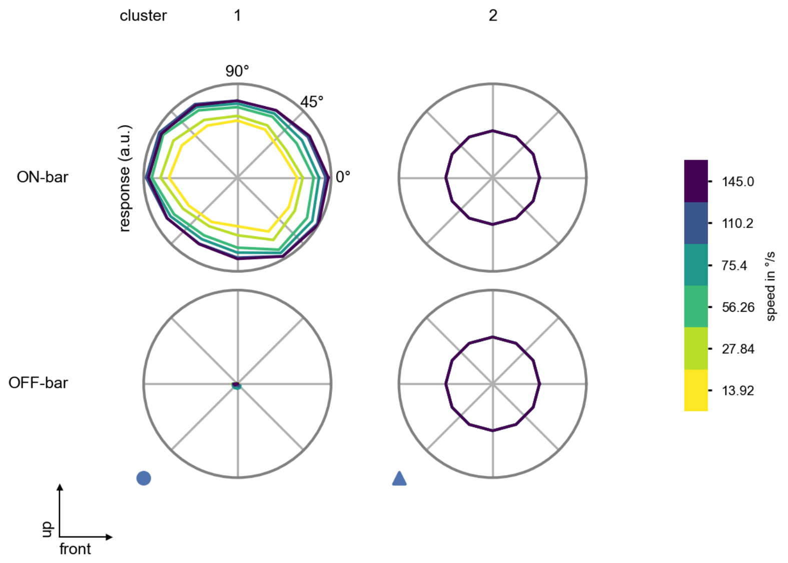

T3 - Figure 9: **Peak responses to moving bars from best-performing models.** The top row shows peak responses to moving ON-bars, the bottom row shows peak responses to moving OFF-bars. The bar-stimuli move in different directions from 0 to 360 degrees and at different speeds. Before, a grey-stimulus led the network to a stationary state. The columns show responses from the best-task-performing model in the respective cluster.

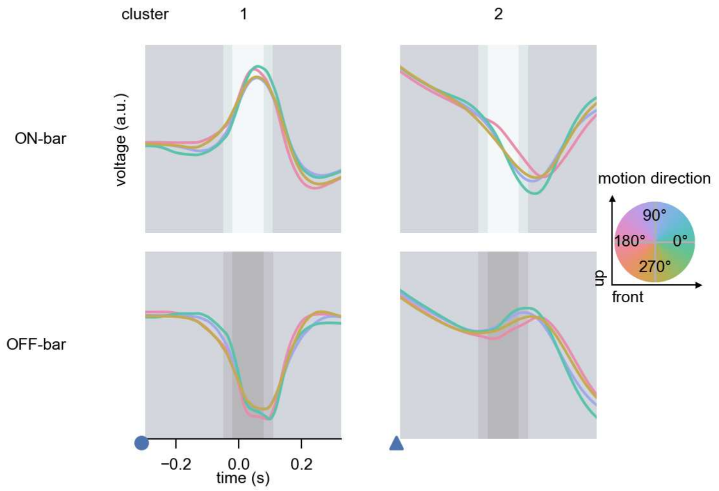

T3 - Figure 10: **Responses to moving bars from best-performing models.** The top row shows responses to moving ON-bars, the bottom row shows peak responses to moving OFF-bars. The bar-stimuli move in different directions from 0 to 360 degrees and at different speeds. Before, a grey-stimulus led the network to a stationary state. The columns show responses from the best-task-performing model in the respective cluster. Example responses to bars moving at 75.4°/s in all cardinal directions (green 0°, blue 90°, red 180°, yellow 270°) from -22.5 to 22.5° visual angle. Background contour indicates the bar, passing by the central column at second zero.

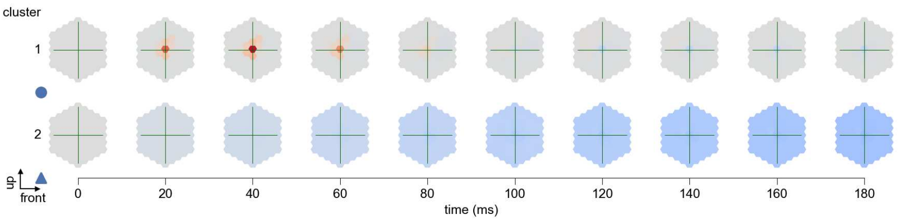

T3 - Figure 11: **Spatio-temporal receptive field.** Responses of the central cell to ON-impulses at single ommatidium impulse locations. The impulse occurs at second zero. Before, a grey-stimulus led the network to a stationary state. Responses are baseline-subtracted. Each row illustrates responses from the best-performing model of the respective cluster. Red indicated depolarization, blue indicates hyperpolarization.

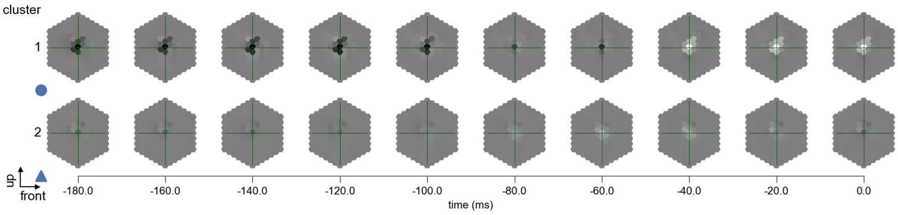

T3 - Figure 12: **Maximally excitatory stimuli.** Each row presents the regularized naturalistic-stimulus from the Sintel dataset that maximizes the cell type's central column response at second zero in the best-performing model of the respective cluster.

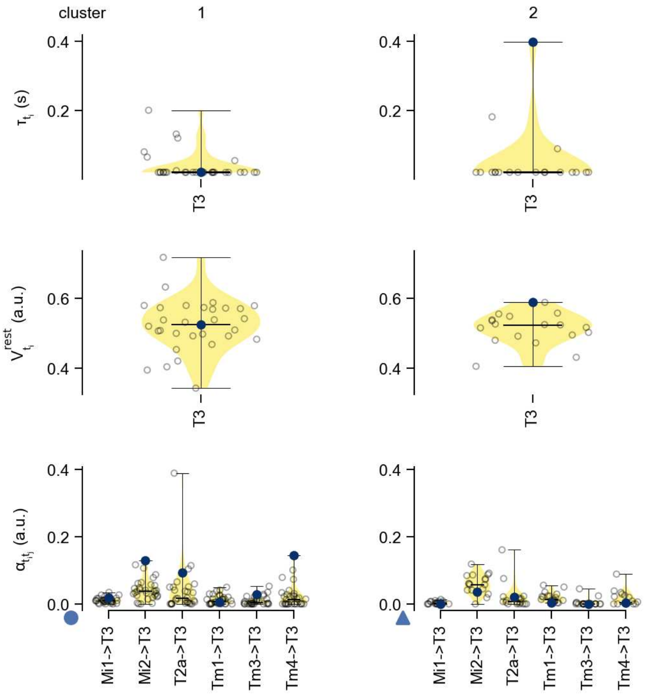

T3 - Figure 13: **Task-constrained parameters.** Each column shows the parameters inferred within the respective cluster. First row: learned time constants of the cell type. Second row: resting potentials of the cell type. Third row: scaling factors for the convolutional filters. The blue scatter represents the parameters from the best-performing model within the cluster.

# 36 T4a

← Cell types

#### Figures

1 Anatomical receptive fields. . . . 236
2 Anatomical projective fields. . . . 237
3 Clustering of the responses to naturalistic stimuli. . . . 237
4 Responses to flashes. . . . 237
5 Peak responses to moving edges. . . . 238
6 Peak responses to moving edges from best-performing models. . . . 238
7 Responses to moving edges from best-performing models. . . . 239
8 Peak responses to moving bars. . . . 239
9 Peak responses to moving bars from best-performing models. . . . 240
10 Responses to moving bars from best-performing models. . . . 240
11 Spatio-temporal receptive field. . . . 241
12 Maximally excitatory stimuli. . . . 241
13 Task-constrained parameters. . . . 242

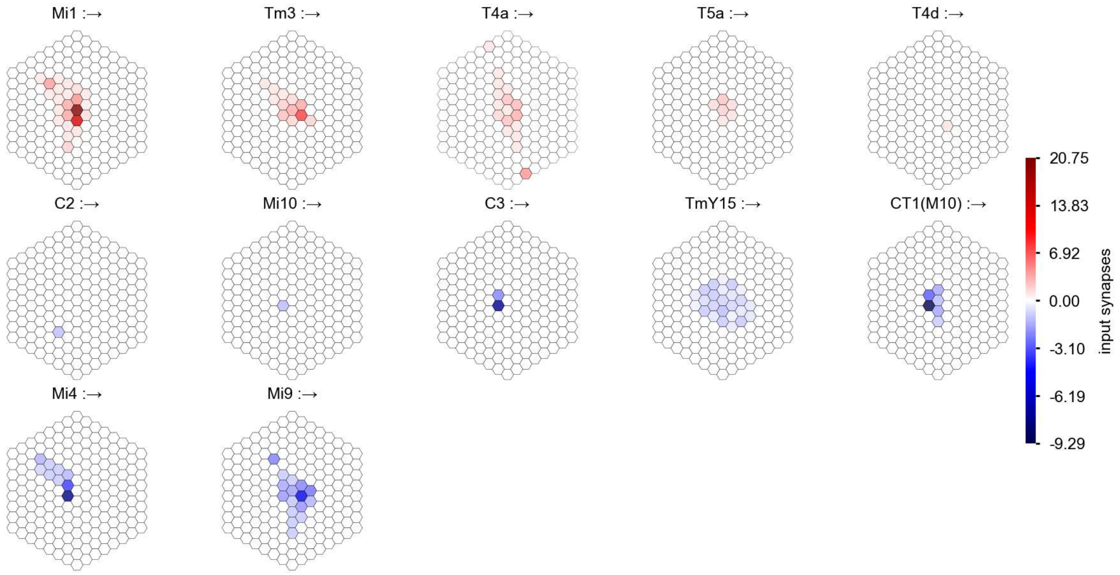

T4a - Figure 1: **Anatomical receptive fields.** Each colored hexagon is a projection, with the projection strength characterized by the average number of synapses that we count from the EM reconstruction. Red indicates excitatory synapses, blue indicates inhibitory synapses from inferred signs. Filters in the order of their total number of synapses.

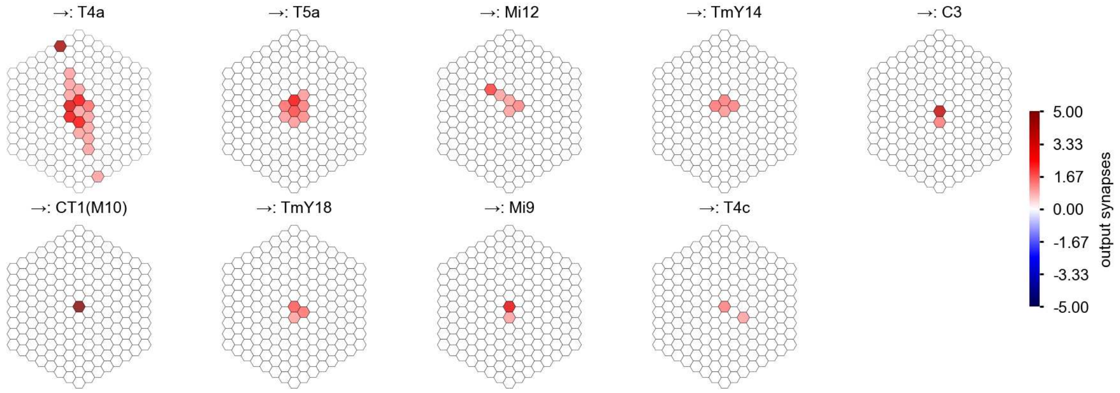

T4a - Figure 2: **Anatomical projective fields.** Each colored hexagon is a projection, with the projection strength characterized by the average number of synapses that we count from the EM reconstruction. Red indicates excitatory synapses, blue indicates inhibitory synapses from inferred signs. Filters in the order of their total number of synapses.

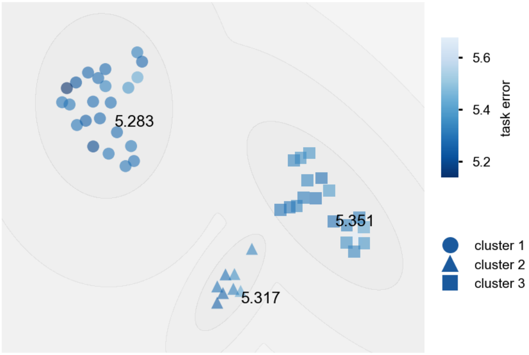

T4a - Figure 3: **Clustering of the responses to naturalistic stimuli.** From responses to naturalistic scenes from Sintel. Scatterpoints represent individual models colored by their task error.

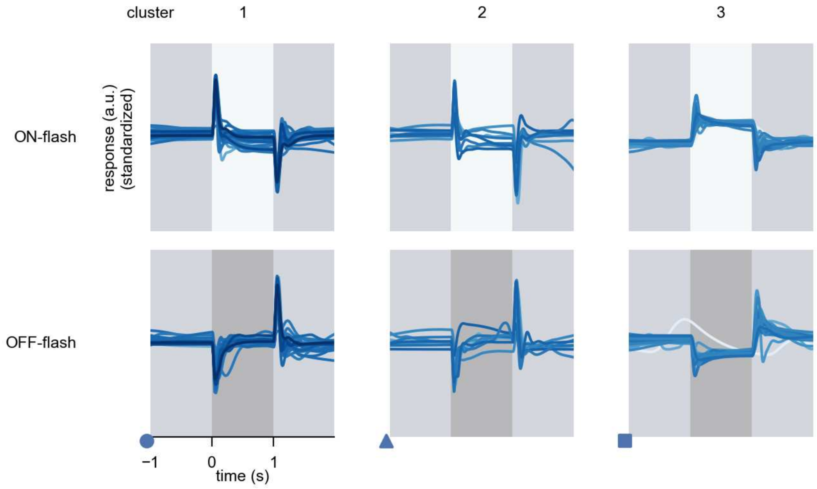

T4a - Figure 4: **Responses to flashes.** The top row shows responses to ON-flashes, the bottom row shows responses to OFF-flashes. The responses from the different models in the different clusters (columns) overlay and are colored in the respective task error of the model. The circular flashes occur at second zero, cover 6 ommatidia in radius, and last for one second. Before and after, a grey-stimulus leads to a stationary state of the network. The traces are standardized.

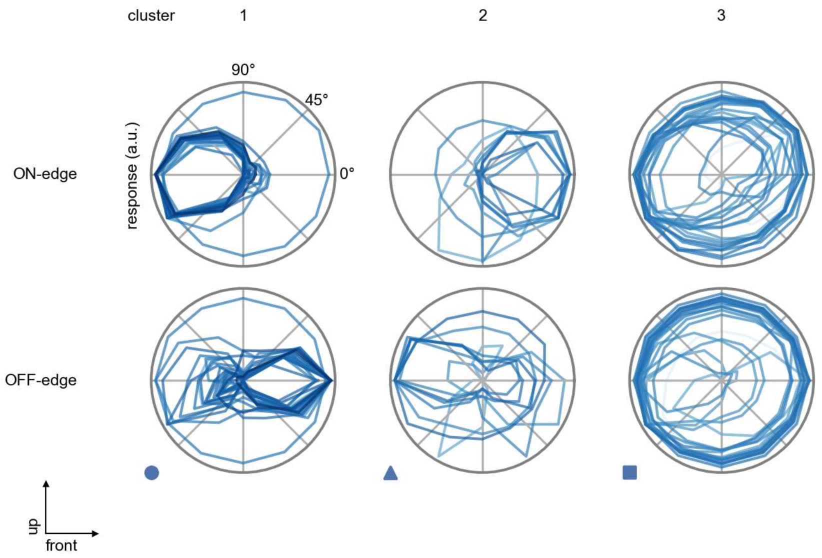

T4a - Figure 5: **Peak responses to moving edges.** The top row shows normalized peak responses to moving ON-edges, the bottom row shows normalized peak responses to moving OFF-edges. The edge-stimuli move in different directions from 0 to 360 degrees. Before, a grey-stimulus led the network to a stationary state. The responses from the different models in the different clusters (columns) overlay and are colored in the respective task error of the model. The peak responses are averaged over all assessed edge-speeds.

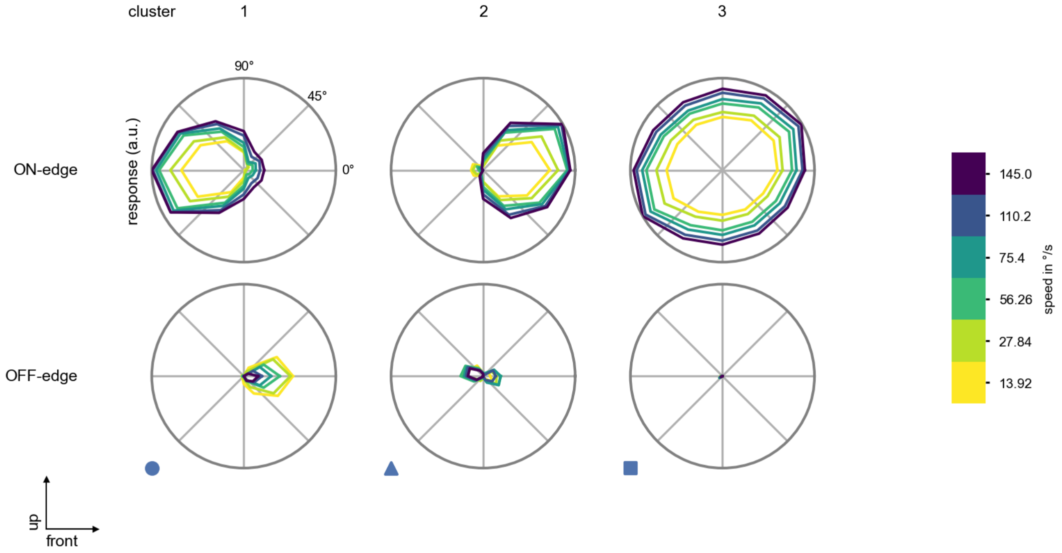

T4a - Figure 6: **Peak responses to moving edges from best-performing models.** The top row shows peak responses to moving ON-edges, the bottom row shows peak responses to moving OFF-edges. The edge-stimuli move in different directions from 0 to 360 degrees and at different speeds. Before, a grey-stimulus led the network to a stationary state. The columns show responses from the best-task-performing model in the respective cluster.

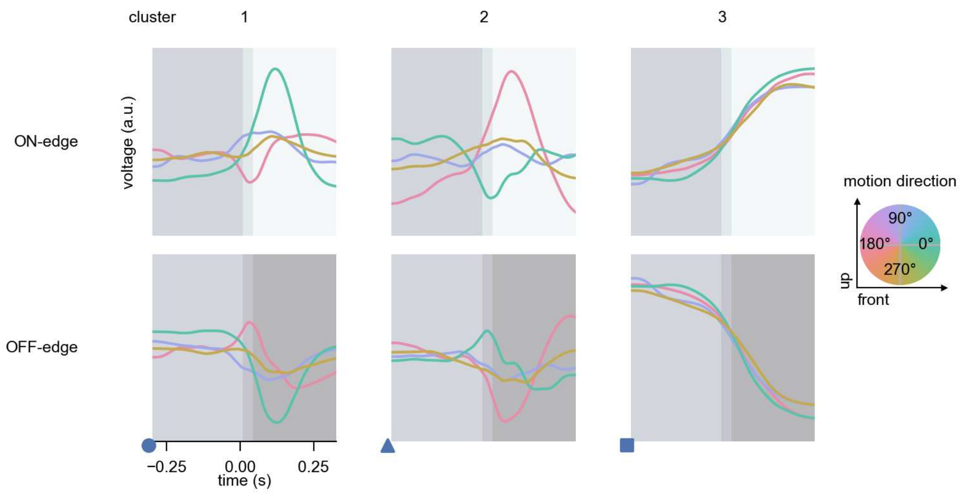

T4a - Figure 7: **Responses to moving edges from best-performing models.** The top row shows responses to moving ON-edges, the bottom row shows peak responses to moving OFF-edges. The edge-stimuli move in different directions from 0 to 360 degrees and at different speeds. Before, a grey-stimulus led the network to a stationary state. The columns show responses from the best-task-performing model in the respective cluster. Example responses to edges moving at 75.4°/s in all cardinal directions (green 0°, blue 90°, red 180°, yellow 270°) from -22.5 to 22.5° visual angle. Background contour indicates the edge, passing by the central column at second zero.

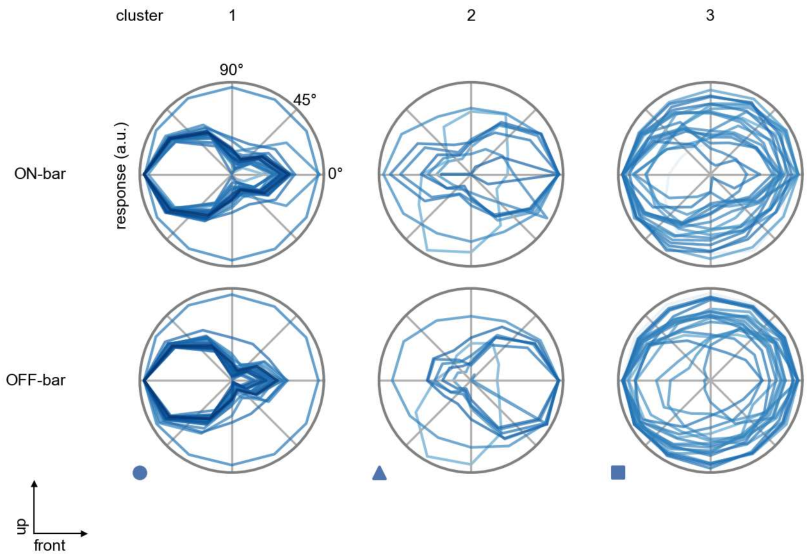

T4a - Figure 8: **Peak responses to moving bars.** The top row shows normalized peak responses to moving ON-bars, the bottom row shows normalized peak responses to moving OFF-bars. The bar-stimuli move in different directions from 0 to 360 degrees. Before, a grey-stimulus led the network to a stationary state. The responses from the different models in the different clusters (columns) overlay and are colored in the respective task error of the model. The peak responses are averaged over all assessed bar-speeds.

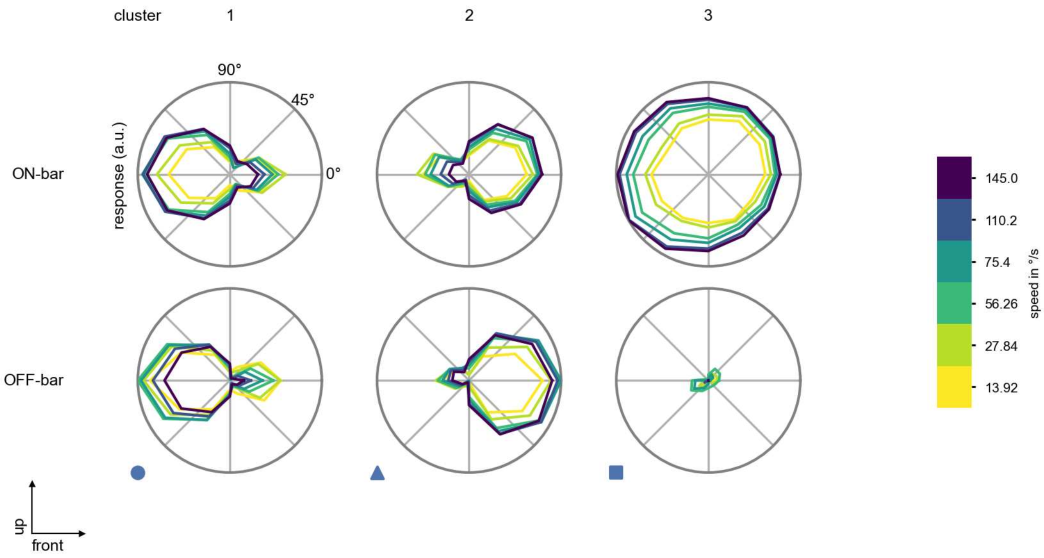

T4a - Figure 9: **Peak responses to moving bars from best-performing models.** The top row shows peak responses to moving ON-bars, the bottom row shows peak responses to moving OFF-bars. The bar-stimuli move in different directions from 0 to 360 degrees and at different speeds. Before, a grey-stimulus led the network to a stationary state. The columns show responses from the best-task-performing model in the respective cluster.

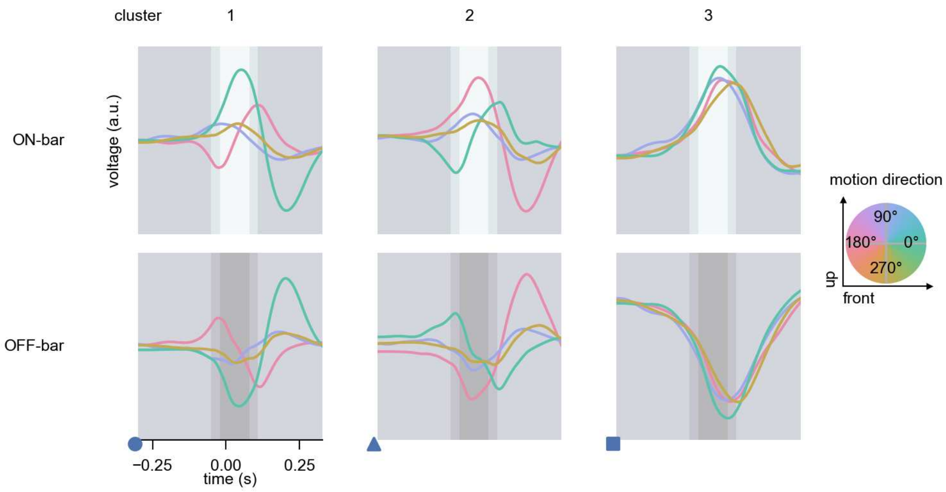

T4a - Figure 10: **Responses to moving bars from best-performing models.** The top row shows responses to moving ON-bars, the bottom row shows peak responses to moving OFF-bars. The bar-stimuli move in different directions from 0 to 360 degrees and at different speeds. Before, a grey-stimulus led the network to a stationary state. The columns show responses from the best-task-performing model in the respective cluster. Example responses to bars moving at 75.4°/s in all cardinal directions (green 0°, blue 90°, red 180°, yellow 270°) from -22.5 to 22.5° visual angle. Background contour indicates the bar, passing by the central column at second zero.

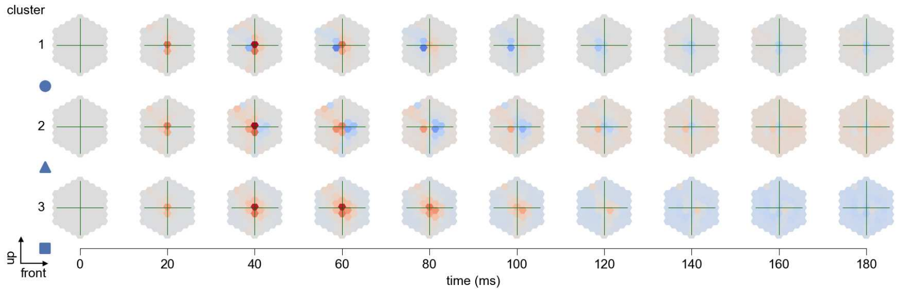

T4a - Figure 11: **Spatio-temporal receptive field.** Responses of the central cell to ON-impulses at single ommatidium impulse locations. The impulse occurs at second zero. Before, a grey-stimulus led the network to a stationary state. Responses are baseline-subtracted. Each row illustrates responses from the best-performing model of the respective cluster. Red indicated depolarization, blue indicates hyperpolarization.

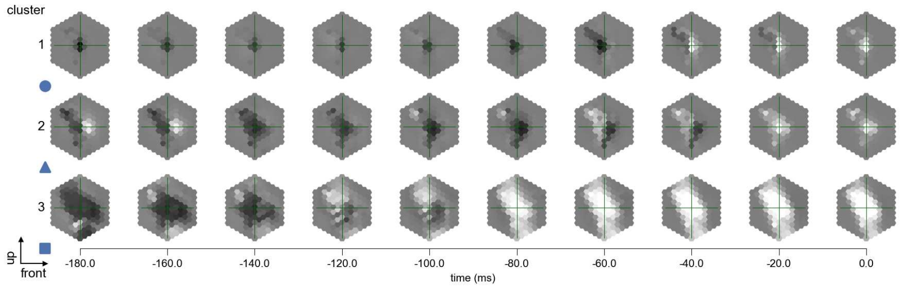

T4a - Figure 12: **Maximally excitatory stimuli.** Each row presents the regularized naturalistic-stimulus from the Sintel dataset that maximizes the cell type's central column response at second zero in the best-performing model of the respective cluster.

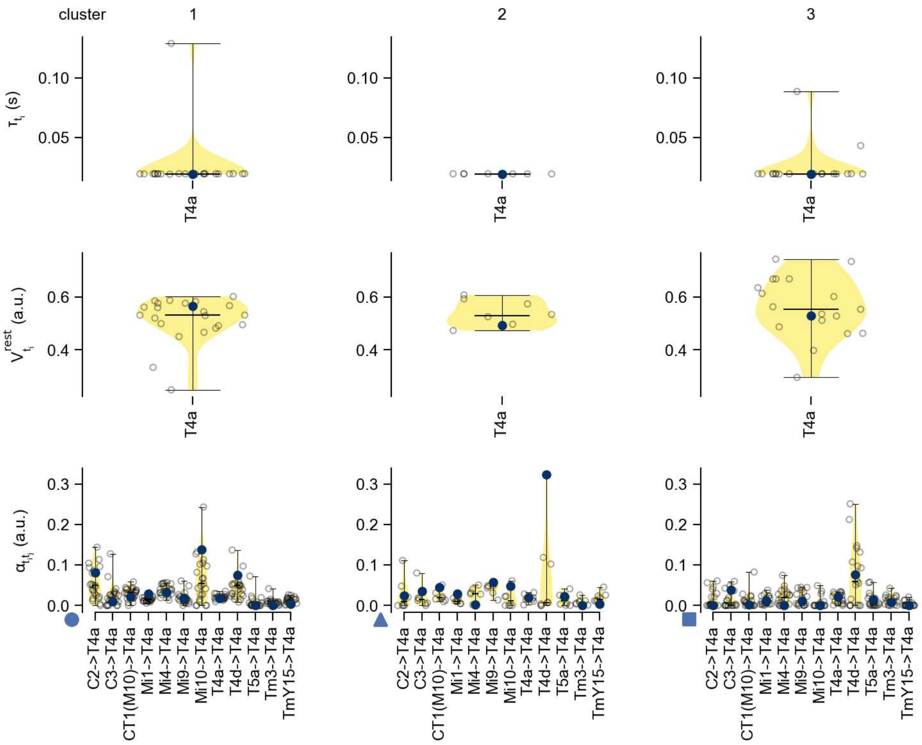

T4a - Figure 13: **Task-constrained parameters.** Each column shows the parameters inferred within the respective cluster. First row: learned time constants of the cell type. Second row: resting potentials of the cell type. Third row: scaling factors for the convolutional filters. The blue scatter represents the parameters from the best-performing model within the cluster.

# 37 T4b

← Cell types

#### Figures

1 Anatomical receptive fields. . . . . 243
2 Anatomical projective fields. . . . . 243
3 Clustering of the responses to naturalistic stimuli. . . . . 244
4 Responses to flashes. . . . . 244
5 Peak responses to moving edges. . . . . 245
6 Peak responses to moving edges from best-performing models. . . . . 245
7 Responses to moving edges from best-performing models. . . . . 246
8 Peak responses to moving bars. . . . . 246
9 Peak responses to moving bars from best-performing models. . . . . 247
10 Responses to moving bars from best-performing models. . . . . 247
11 Spatio-temporal receptive field. . . . . 248
12 Maximally excitatory stimuli. . . . . 248
13 Task-constrained parameters. . . . . 249

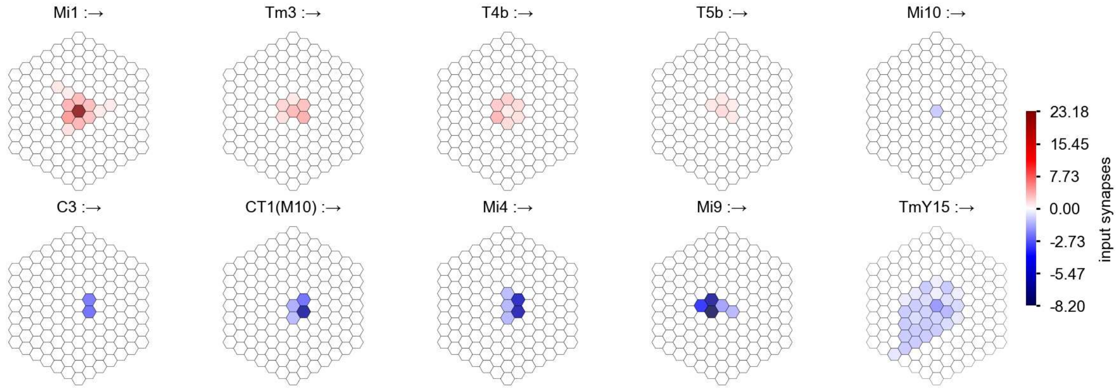

T4b - Figure 1: **Anatomical receptive fields.** Each colored hexagon is a projection, with the projection strength characterized by the average number of synapses that we count from the EM reconstruction. Red indicates excitatory synapses, blue indicates inhibitory synapses from inferred signs. Filters in the order of their total number of synapses.

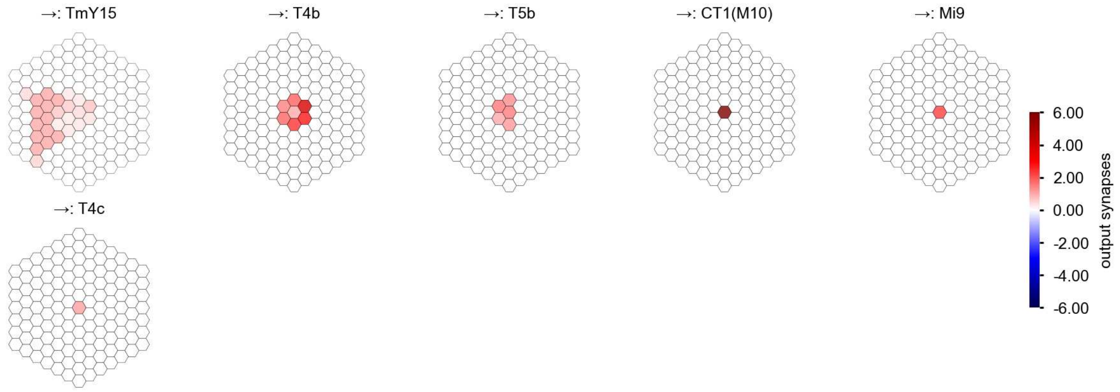

T4b - Figure 2: **Anatomical projective fields.** Each colored hexagon is a projection, with the projection strength characterized by the average number of synapses that we count from the EM reconstruction. Red indicates excitatory synapses, blue indicates inhibitory synapses from inferred signs. Filters in the order of their total number of synapses.

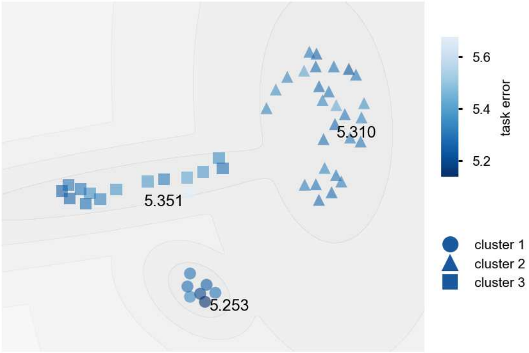

T4b - Figure 3: **Clustering of the responses to naturalistic stimuli.** From responses to naturalistic scenes from Sintel. Scatterpoints represent individual models colored by their task error.

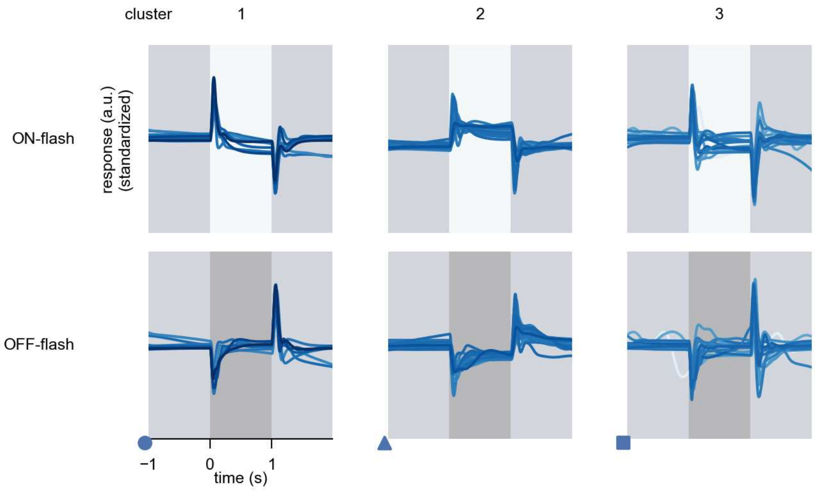

T4b - Figure 4: **Responses to flashes.** The top row shows responses to ON-flashes, the bottom row shows responses to OFF-flashes. The responses from the different models in the different clusters (columns) overlay and are colored in the respective task error of the model. The circular flashes occur at second zero, cover 6 ommatidia in radius, and last for one second. Before and after, a grey-stimulus leads to a stationary state of the network. The traces are standardized.

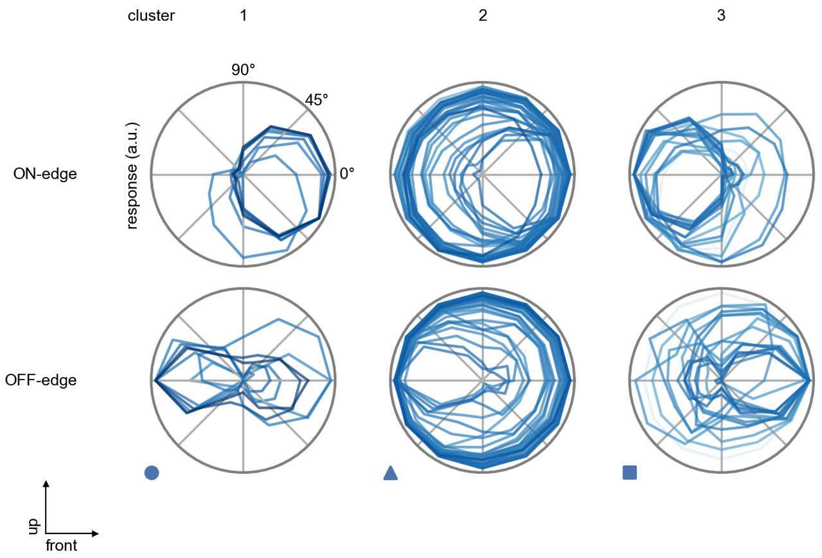

T4b - Figure 5: **Peak responses to moving edges.** The top row shows normalized peak responses to moving ON-edges, the bottom row shows normalized peak responses to moving OFF-edges. The edge-stimuli move in different directions from 0 to 360 degrees. Before, a grey-stimulus led the network to a stationary state. The responses from the different models in the different clusters (columns) overlay and are colored in the respective task error of the model. The peak responses are averaged over all assessed edge-speeds.

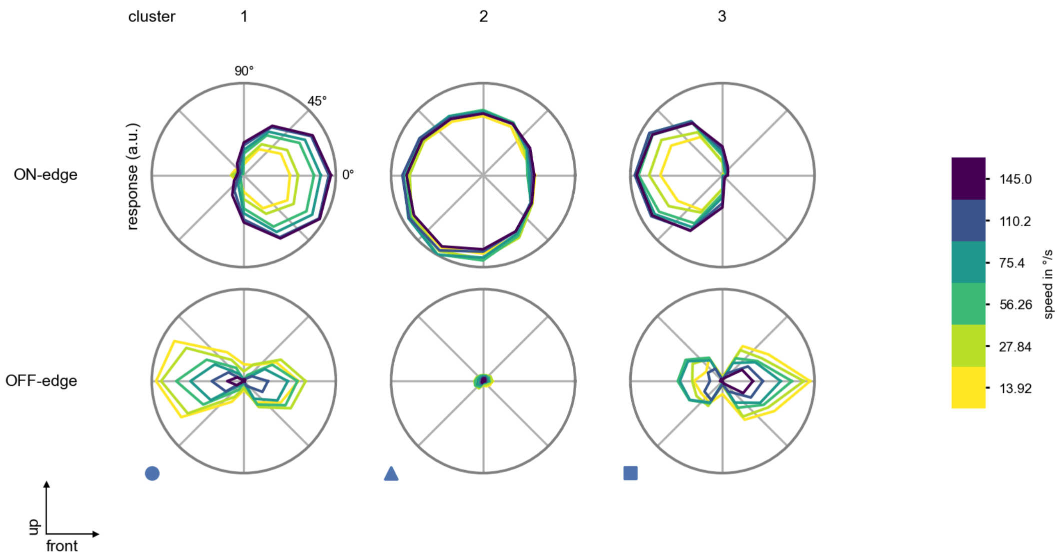

T4b - Figure 6: **Peak responses to moving edges from best-performing models.** The top row shows peak responses to moving ON-edges, the bottom row shows peak responses to moving OFF-edges. The edge-stimuli move in different directions from 0 to 360 degrees and at different speeds. Before, a grey-stimulus led the network to a stationary state. The columns show responses from the best-task-performing model in the respective cluster.

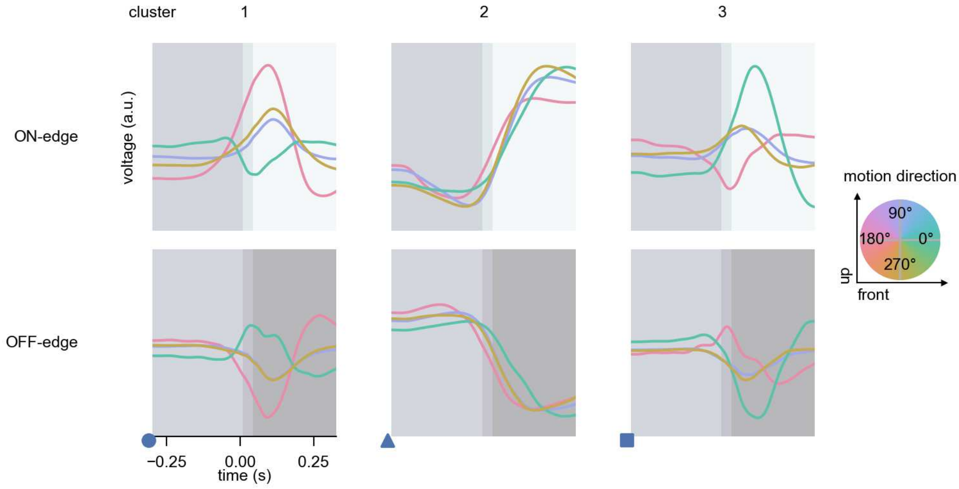

T4b - Figure 7: **Responses to moving edges from best-performing models.** The top row shows responses to moving ON-edges, the bottom row shows peak responses to moving OFF-edges. The edge-stimuli move in different directions from 0 to 360 degrees and at different speeds. Before, a grey-stimulus led the network to a stationary state. The columns show responses from the best-task-performing model in the respective cluster. Example responses to edges moving at 75.4°/s in all cardinal directions (green 0°, blue 90°, red 180°, yellow 270°) from -22.5 to 22.5° visual angle. Background contour indicates the edge, passing by the central column at second zero.

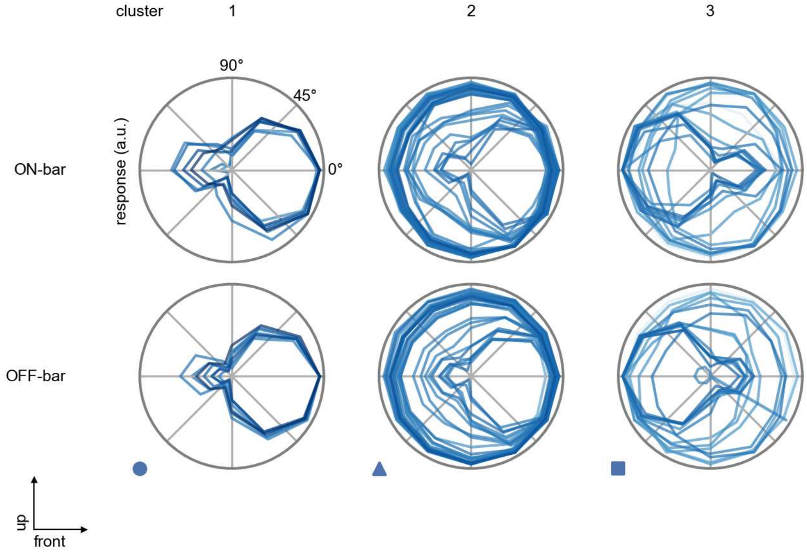

T4b - Figure 8: **Peak responses to moving bars.** The top row shows normalized peak responses to moving ON-bars, the bottom row shows normalized peak responses to moving OFF-bars. The bar-stimuli move in different directions from 0 to 360 degrees. Before, a grey-stimulus led the network to a stationary state. The responses from the different models in the different clusters (columns) overlay and are colored in the respective task error of the model. The peak responses are averaged over all assessed bar-speeds.

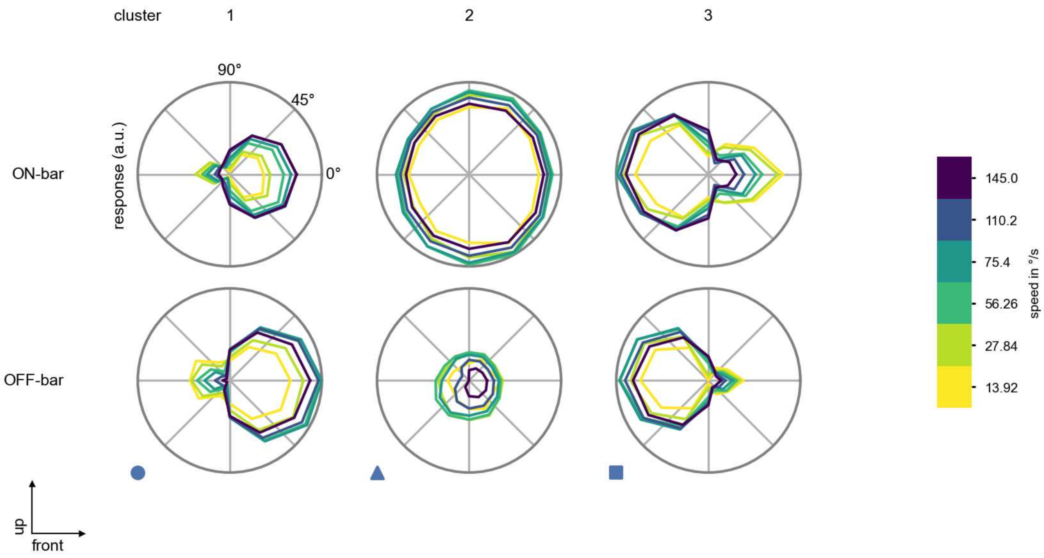

T4b - Figure 9: **Peak responses to moving bars from best-performing models.** The top row shows peak responses to moving ON-bars, the bottom row shows peak responses to moving OFF-bars. The bar-stimuli move in different directions from 0 to 360 degrees and at different speeds. Before, a grey-stimulus led the network to a stationary state. The columns show responses from the best-task-performing model in the respective cluster.

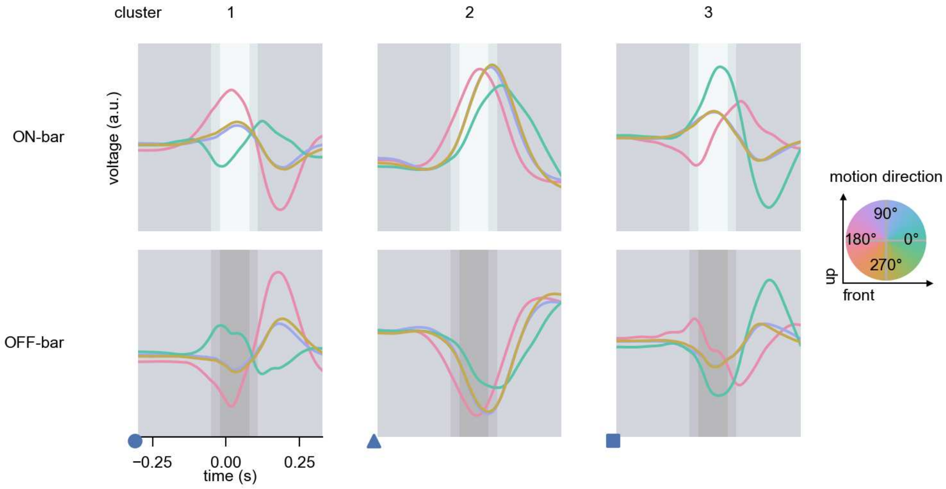

T4b - Figure 10: **Responses to moving bars from best-performing models.** The top row shows responses to moving ON-bars, the bottom row shows peak responses to moving OFF-bars. The bar-stimuli move in different directions from 0 to 360 degrees and at different speeds. Before, a grey-stimulus led the network to a stationary state. The columns show responses from the best-task-performing model in the respective cluster. Example responses to bars moving at 75.4°/s in all cardinal directions (green 0°, blue 90°, red 180°, yellow 270°) from -22.5 to 22.5° visual angle. Background contour indicates the bar, passing by the central column at second zero.

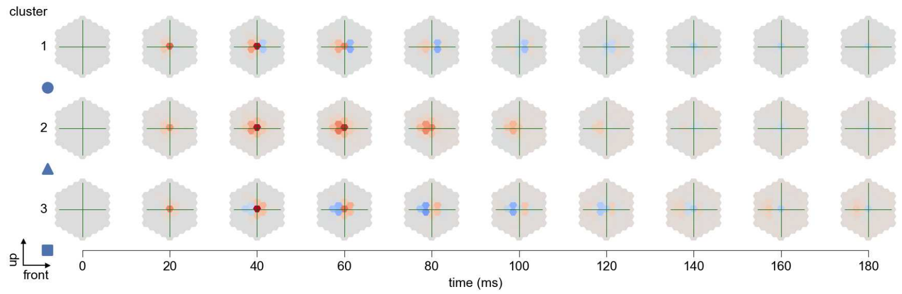

T4b - Figure 11: **Spatio-temporal receptive field.** Responses of the central cell to ON-impulses at single ommatidium impulse locations. The impulse occurs at second zero. Before, a grey-stimulus led the network to a stationary state. Responses are baseline-subtracted. Each row illustrates responses from the best-performing model of the respective cluster. Red indicated depolarization, blue indicates hyperpolarization.

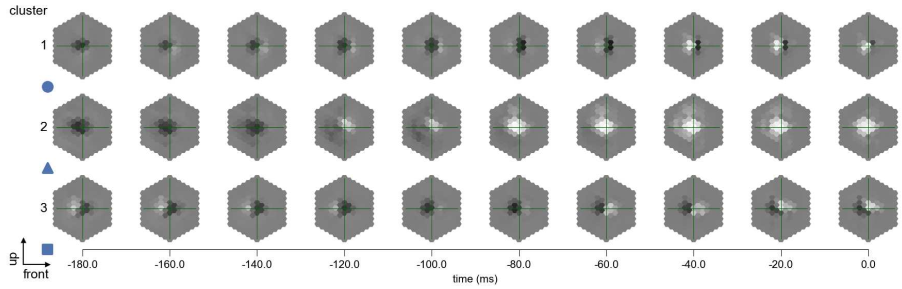

T4b - Figure 12: **Maximally excitatory stimuli.** Each row presents the regularized naturalistic-stimulus from the Sintel dataset that maximizes the cell type's central column response at second zero in the best-performing model of the respective cluster.

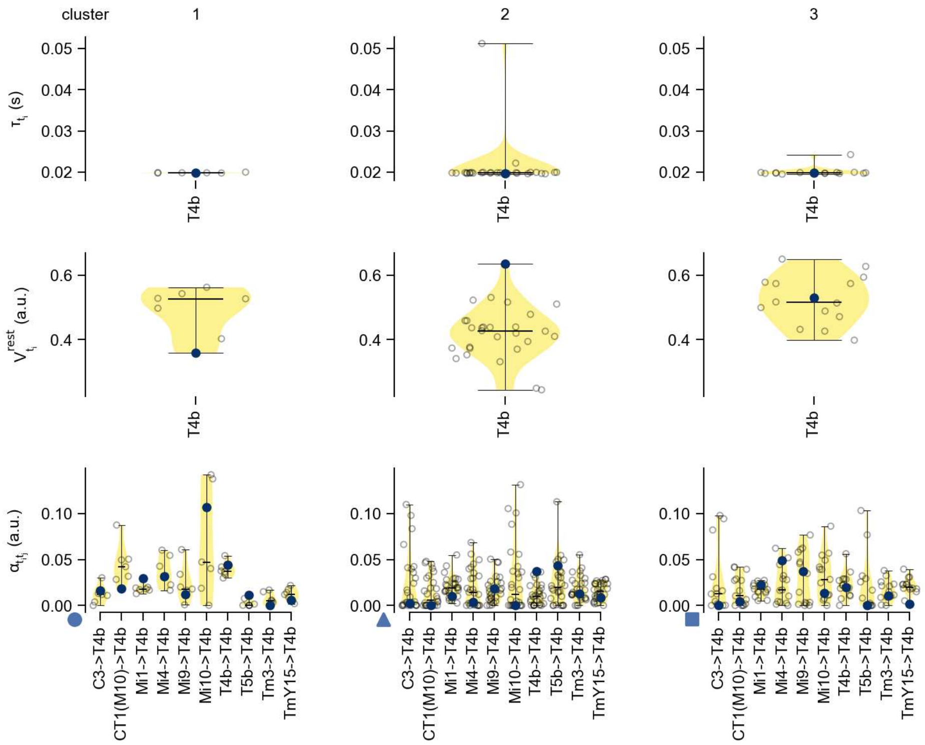

T4b - Figure 13: **Task-constrained parameters.** Each column shows the parameters inferred within the respective cluster. First row: learned time constants of the cell type. Second row: resting potentials of the cell type. Third row: scaling factors for the convolutional filters. The blue scatter represents the parameters from the best-performing model within the cluster.

# 38 T4c

← Cell types

#### Figures

1 Anatomical receptive fields. . . . . 250
2 Anatomical projective fields. . . . . 251
3 Clustering of the responses to naturalistic stimuli. . . . . 251
4 Responses to flashes. . . . . 251
5 Peak responses to moving edges. . . . . 252
6 Peak responses to moving edges from best-performing models. . . . . 252
7 Responses to moving edges from best-performing models. . . . . 253
8 Peak responses to moving bars. . . . . 253
9 Peak responses to moving bars from best-performing models. . . . . 254
10 Responses to moving bars from best-performing models. . . . . 254
11 Spatio-temporal receptive field. . . . . 255
12 Maximally excitatory stimuli. . . . . 255
13 Task-constrained parameters. . . . . 256

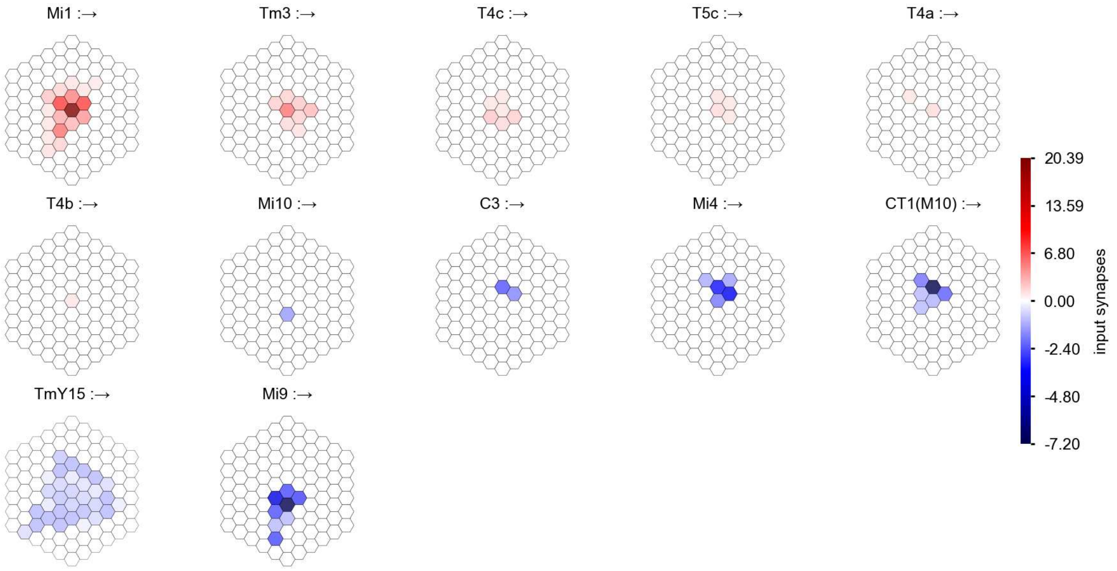

T4c - Figure 1: **Anatomical receptive fields.** Each colored hexagon is a projection, with the projection strength characterized by the average number of synapses that we count from the EM reconstruction. Red indicates excitatory synapses, blue indicates inhibitory synapses from inferred signs. Filters in the order of their total number of synapses.

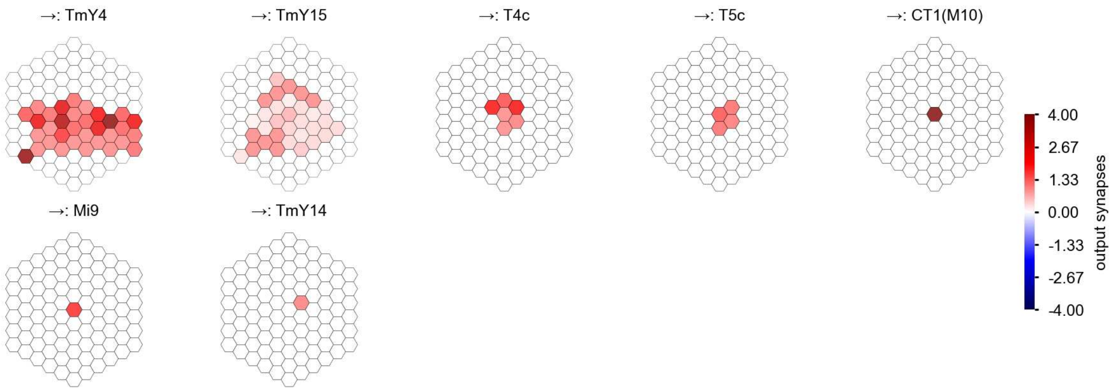

T4c - Figure 2: **Anatomical projective fields.** Each colored hexagon is a projection, with the projection strength characterized by the average number of synapses that we count from the EM reconstruction. Red indicates excitatory synapses, blue indicates inhibitory synapses from inferred signs. Filters in the order of their total number of synapses.

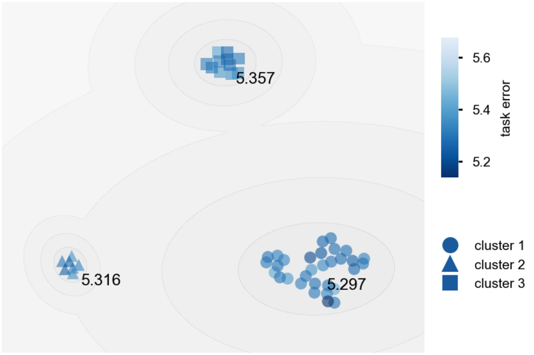

T4c - Figure 3: **Clustering of the responses to naturalistic stimuli.** From responses to naturalistic scenes from Sintel. Scatterpoints represent individual models colored by their task error.

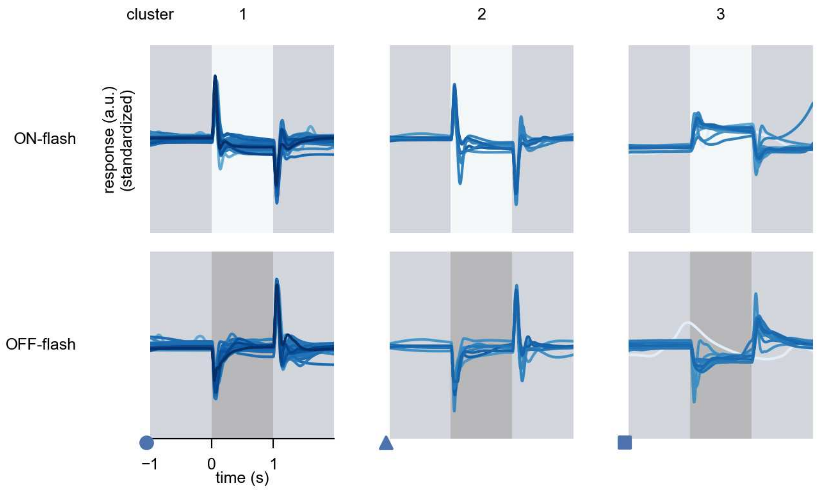

T4c - Figure 4: **Responses to flashes.** The top row shows responses to ON-flashes, the bottom row shows responses to OFF-flashes. The responses from the different models in the different clusters (columns) overlay and are colored in the respective task error of the model. The circular flashes occur at second zero, cover 6 ommatidia in radius, and last for one second. Before and after, a grey-stimulus leads to a stationary state of the network. The traces are standardized.

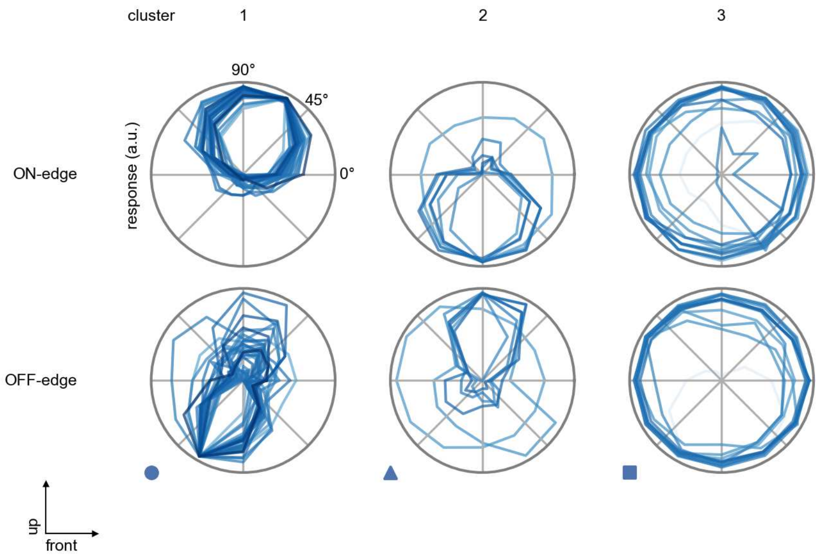

T4c - Figure 5: **Peak responses to moving edges.** The top row shows normalized peak responses to moving ON-edges, the bottom row shows normalized peak responses to moving OFF-edges. The edge-stimuli move in different directions from 0 to 360 degrees. Before, a grey-stimulus led the network to a stationary state. The responses from the different models in the different clusters (columns) overlay and are colored in the respective task error of the model. The peak responses are averaged over all assessed edge-speeds.

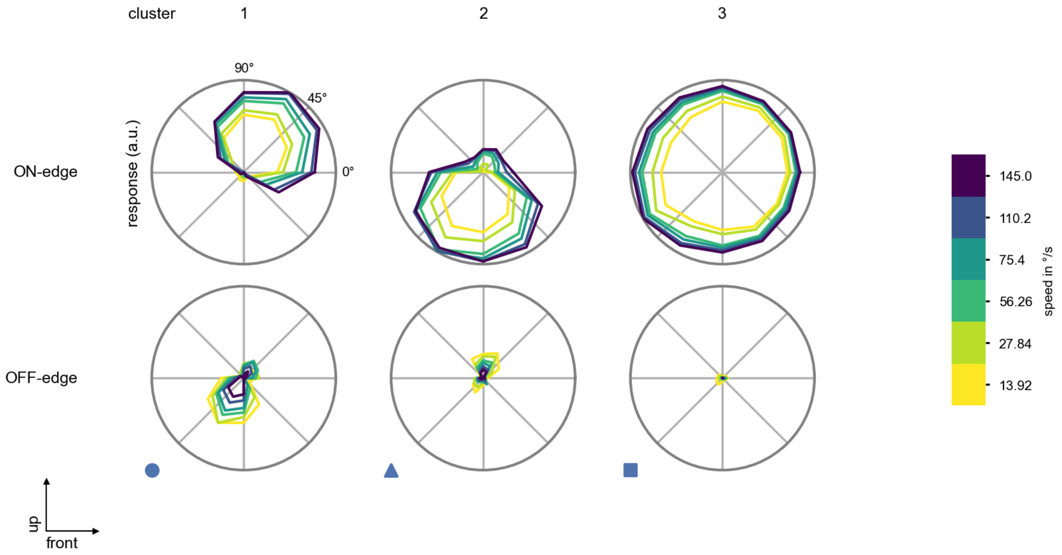

T4c - Figure 6: **Peak responses to moving edges from best-performing models.** The top row shows peak responses to moving ON-edges, the bottom row shows peak responses to moving OFF-edges. The edge-stimuli move in different directions from 0 to 360 degrees and at different speeds. Before, a grey-stimulus led the network to a stationary state. The columns show responses from the best-task-performing model in the respective cluster.

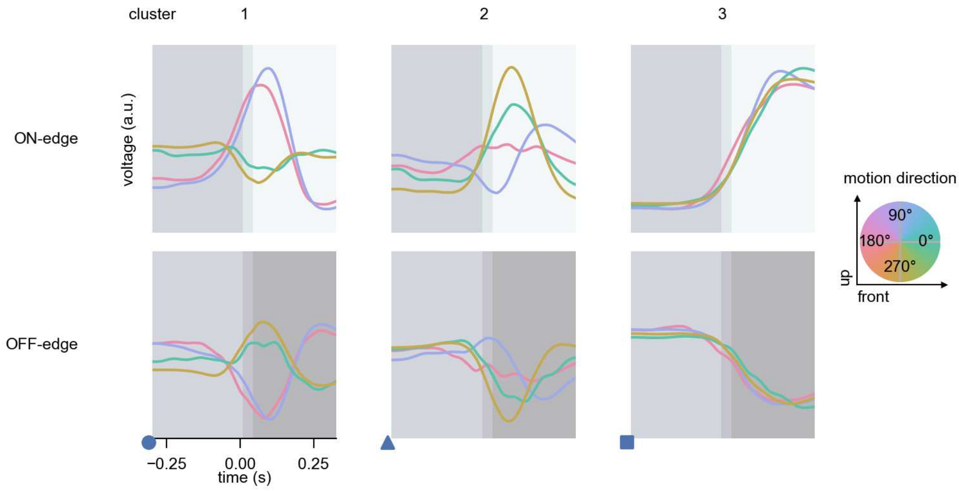

T4c - Figure 7: **Responses to moving edges from best-performing models.** The top row shows responses to moving ON-edges, the bottom row shows peak responses to moving OFF-edges. The edge-stimuli move in different directions from 0 to 360 degrees and at different speeds. Before, a grey-stimulus led the network to a stationary state. The columns show responses from the best-task-performing model in the respective cluster. Example responses to edges moving at 75.4°/s in all cardinal directions (green 0°, blue 90°, red 180°, yellow 270°) from -22.5 to 22.5° visual angle. Background contour indicates the edge, passing by the central column at second zero.

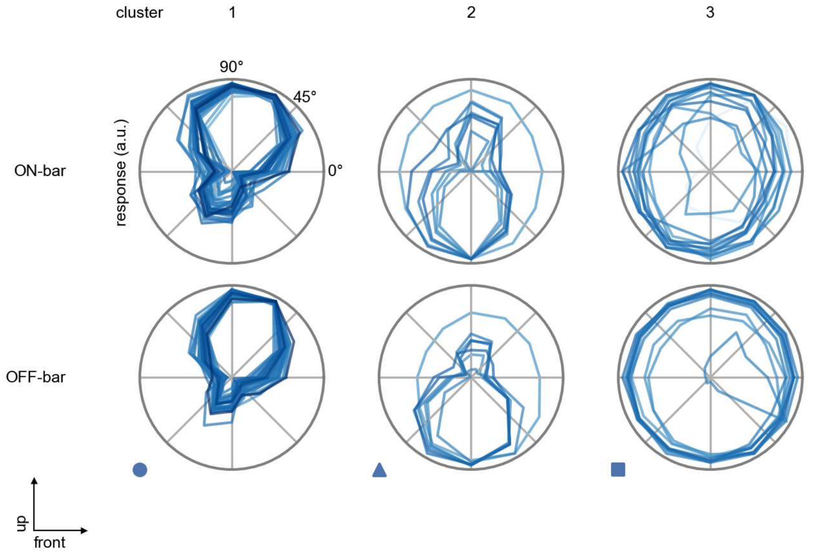

T4c - Figure 8: **Peak responses to moving bars.** The top row shows normalized peak responses to moving ON-bars, the bottom row shows normalized peak responses to moving OFF-bars. The bar-stimuli move in different directions from 0 to 360 degrees. Before, a grey-stimulus led the network to a stationary state. The responses from the different models in the different clusters (columns) overlay and are colored in the respective task error of the model. The peak responses are averaged over all assessed bar-speeds.

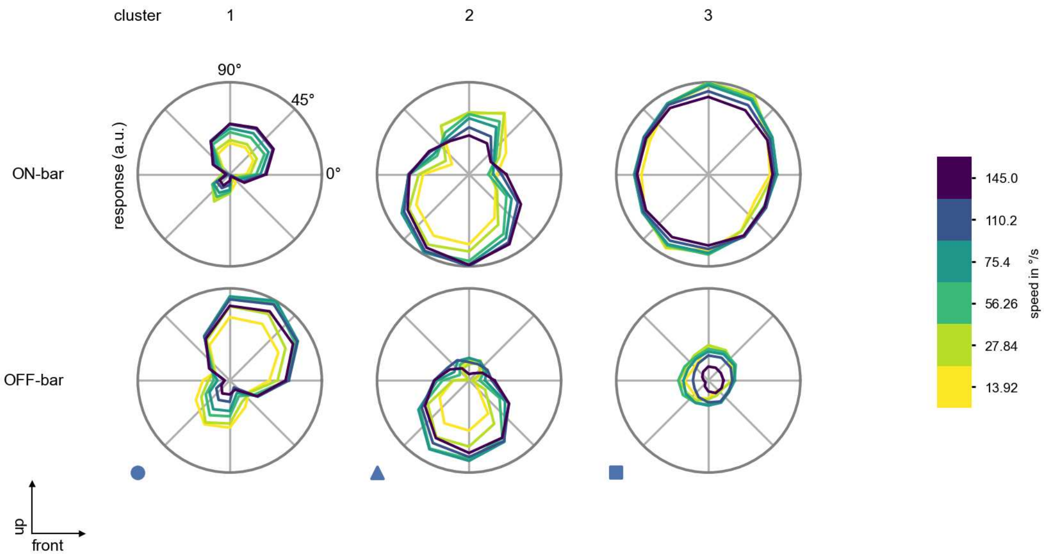

T4c - Figure 9: **Peak responses to moving bars from best-performing models.** The top row shows peak responses to moving ON-bars, the bottom row shows peak responses to moving OFF-bars. The bar-stimuli move in different directions from 0 to 360 degrees and at different speeds. Before, a grey-stimulus led the network to a stationary state. The columns show responses from the best-task-performing model in the respective cluster.

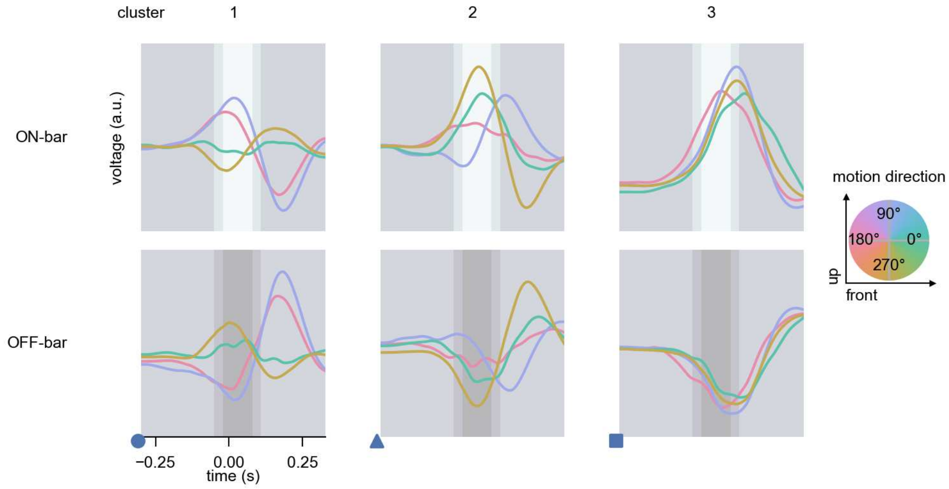

T4c - Figure 10: **Responses to moving bars from best-performing models.** The top row shows responses to moving ON-bars, the bottom row shows peak responses to moving OFF-bars. The bar-stimuli move in different directions from 0 to 360 degrees and at different speeds. Before, a grey-stimulus led the network to a stationary state. The columns show responses from the best-task-performing model in the respective cluster. Example responses to bars moving at 75.4°/s in all cardinal directions (green 0°, blue 90°, red 180°, yellow 270°) from -22.5 to 22.5° visual angle. Background contour indicates the bar, passing by the central column at second zero.

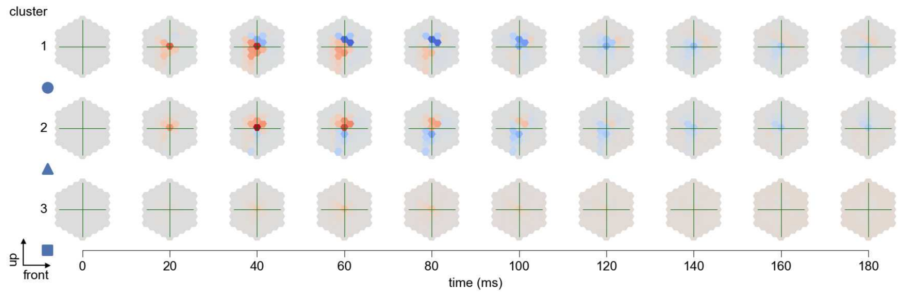

T4c - Figure 11: **Spatio-temporal receptive field.** Responses of the central cell to ON-impulses at single ommatidium impulse locations. The impulse occurs at second zero. Before, a grey-stimulus led the network to a stationary state. Responses are baseline-subtracted. Each row illustrates responses from the best-performing model of the respective cluster. Red indicated depolarization, blue indicates hyperpolarization.

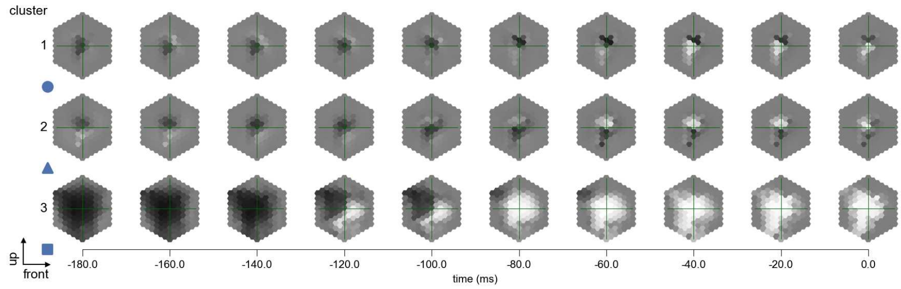

T4c - Figure 12: **Maximally excitatory stimuli.** Each row presents the regularized naturalistic-stimulus from the Sintel dataset that maximizes the cell type's central column response at second zero in the best-performing model of the respective cluster.

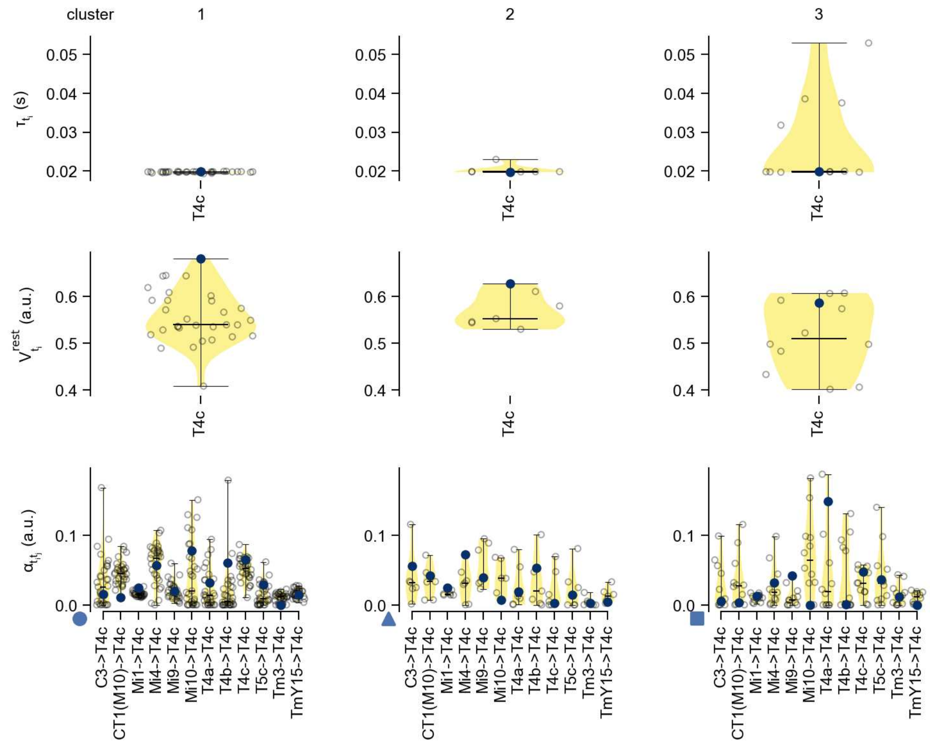

T4c - Figure 13: **Task-constrained parameters.** Each column shows the parameters inferred within the respective cluster. First row: learned time constants of the cell type. Second row: resting potentials of the cell type. Third row: scaling factors for the convolutional filters. The blue scatter represents the parameters from the best-performing model within the cluster.

# 39 T4d

← Cell types

#### Figures

1 Anatomical receptive fields. . . . 257
2 Anatomical projective fields. . . . 257
3 Clustering of the responses to naturalistic stimuli. . . . 258
4 Responses to flashes. . . . 258
5 Peak responses to moving edges. . . . 259
6 Peak responses to moving edges from best-performing models. . . . 259
7 Responses to moving edges from best-performing models. . . . 260
8 Peak responses to moving bars. . . . 260
9 Peak responses to moving bars from best-performing models. . . . 261
10 Responses to moving bars from best-performing models. . . . 261
11 Spatio-temporal receptive field. . . . 262
12 Maximally excitatory stimuli. . . . 262
13 Task-constrained parameters. . . . 263

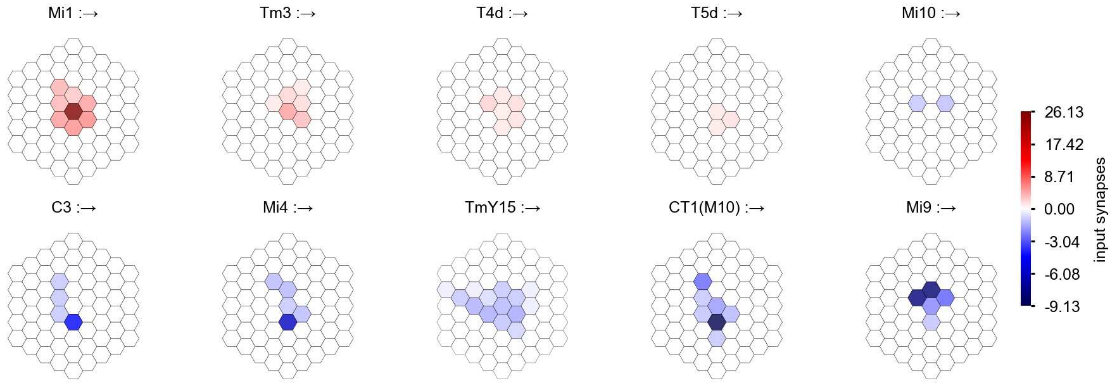

T4d - Figure 1: **Anatomical receptive fields.** Each colored hexagon is a projection, with the projection strength characterized by the average number of synapses that we count from the EM reconstruction. Red indicates excitatory synapses, blue indicates inhibitory synapses from inferred signs. Filters in the order of their total number of synapses.

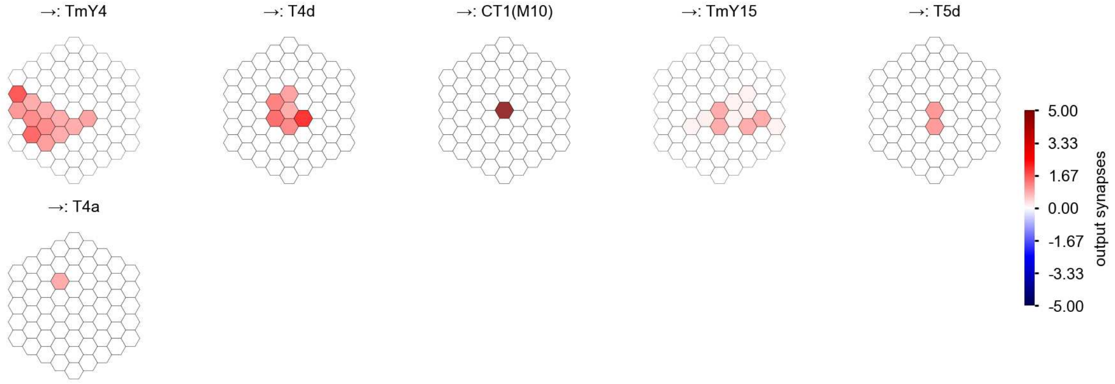

T4d - Figure 2: **Anatomical projective fields.** Each colored hexagon is a projection, with the projection strength characterized by the average number of synapses that we count from the EM reconstruction. Red indicates excitatory synapses, blue indicates inhibitory synapses from inferred signs. Filters in the order of their total number of synapses.

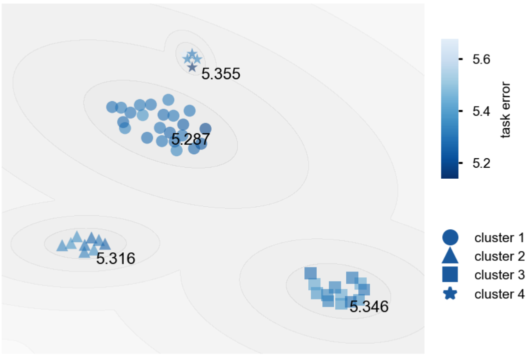

T4d - Figure 3: **Clustering of the responses to naturalistic stimuli.** From responses to naturalistic scenes from Sintel. Scatterpoints represent individual models colored by their task error.

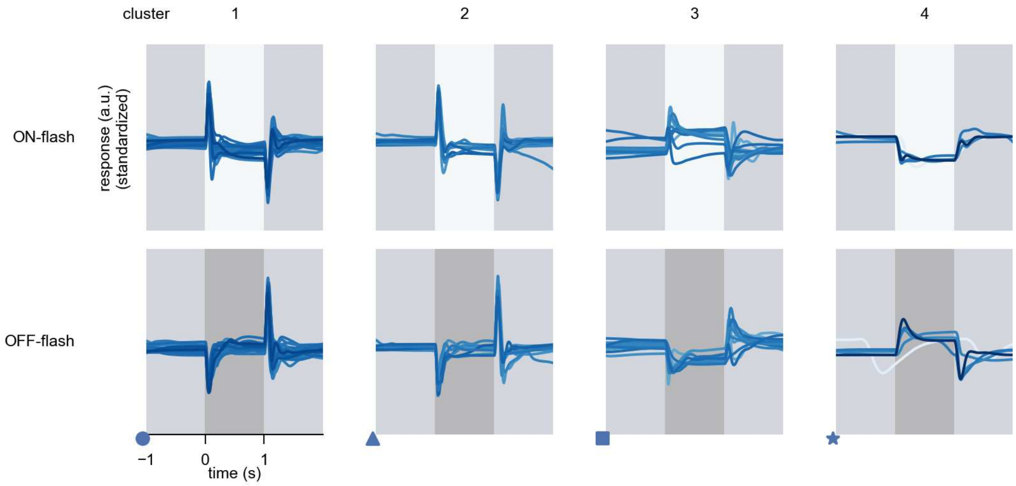

T4d - Figure 4: **Responses to flashes.** The top row shows responses to ON-flashes, the bottom row shows responses to OFF-flashes. The responses from the different models in the different clusters (columns) overlay and are colored in the respective task error of the model. The circular flashes occur at second zero, cover 6 ommatidia in radius, and last for one second. Before and after, a grey-stimulus leads to a stationary state of the network. The traces are standardized.

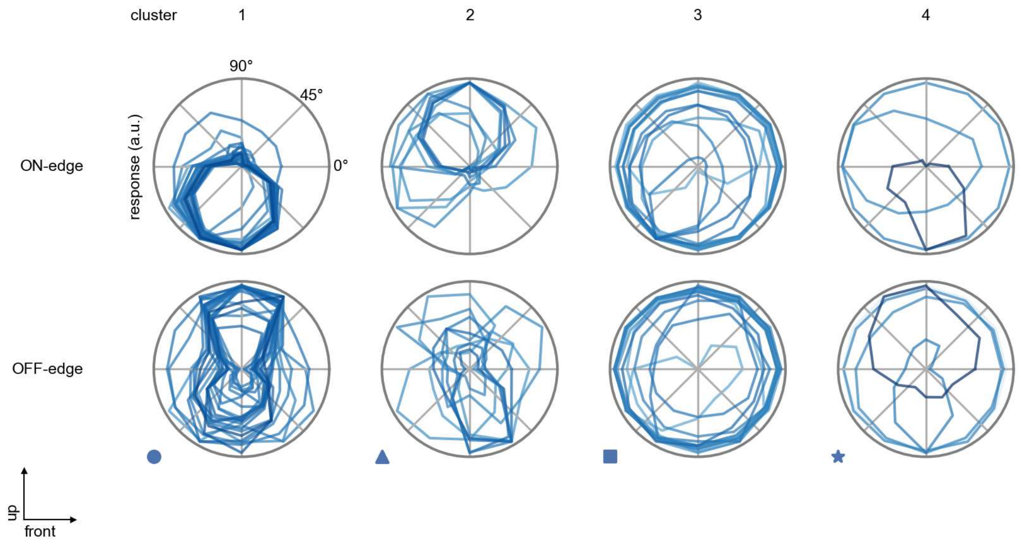

T4d - Figure 5: **Peak responses to moving edges.** The top row shows normalized peak responses to moving ON-edges, the bottom row shows normalized peak responses to moving OFF-edges. The edge-stimuli move in different directions from 0 to 360 degrees. Before, a grey-stimulus led the network to a stationary state. The responses from the different models in the different clusters (columns) overlay and are colored in the respective task error of the model. The peak responses are averaged over all assessed edge-speeds.

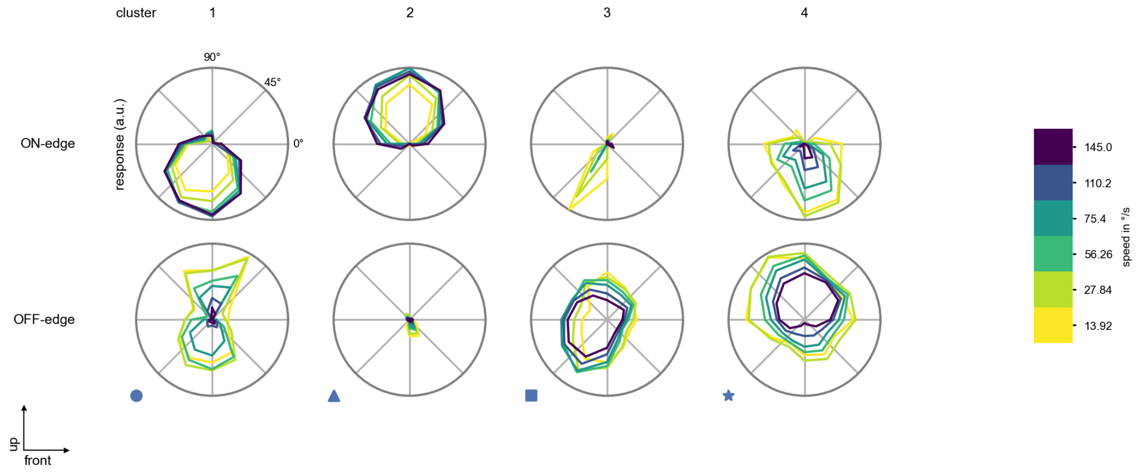

T4d - Figure 6: **Peak responses to moving edges from best-performing models.** The top row shows peak responses to moving ON-edges, the bottom row shows peak responses to moving OFF-edges. The edge-stimuli move in different directions from 0 to 360 degrees and at different speeds. Before, a grey-stimulus led the network to a stationary state. The columns show responses from the best-task-performing model in the respective cluster.

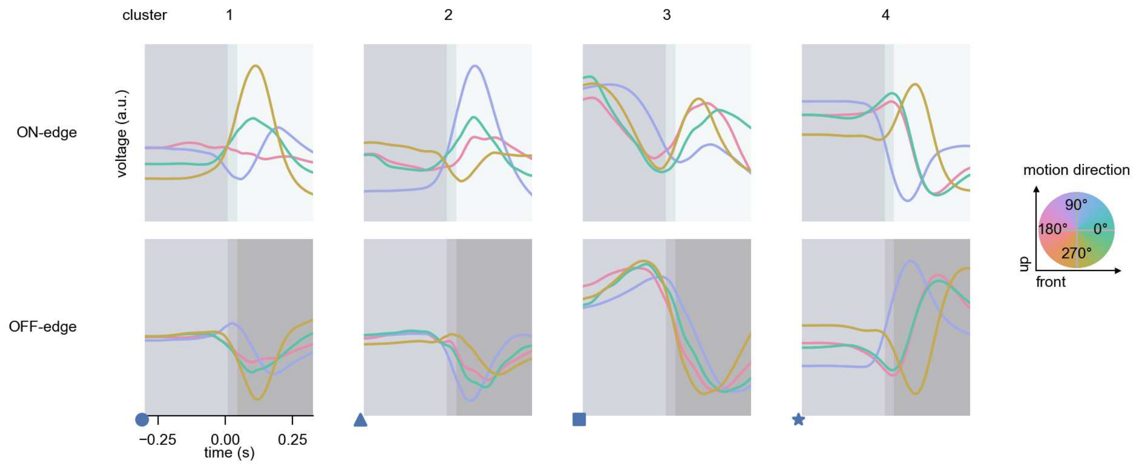

T4d - Figure 7: **Responses to moving edges from best-performing models.** The top row shows responses to moving ON-edges, the bottom row shows peak responses to moving OFF-edges. The edge-stimuli move in different directions from 0 to 360 degrees and at different speeds. Before, a grey-stimulus led the network to a stationary state. The columns show responses from the best-task-performing model in the respective cluster. Example responses to edges moving at 75.4°/s in all cardinal directions (green 0°, blue 90°, red 180°, yellow 270°) from -22.5 to 22.5° visual angle. Background contour indicates the edge, passing by the central column at second zero.

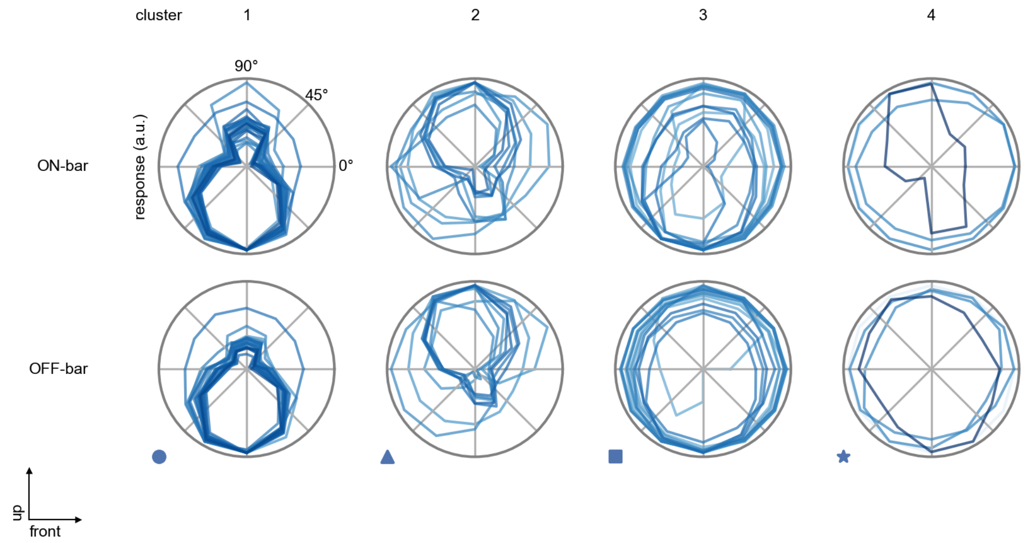

T4d - Figure 8: **Peak responses to moving bars.** The top row shows normalized peak responses to moving ON-bars, the bottom row shows normalized peak responses to moving OFF-bars. The bar-stimuli move in different directions from 0 to 360 degrees. Before, a grey-stimulus led the network to a stationary state. The responses from the different models in the different clusters (columns) overlay and are colored in the respective task error of the model. The peak responses are averaged over all assessed bar-speeds.

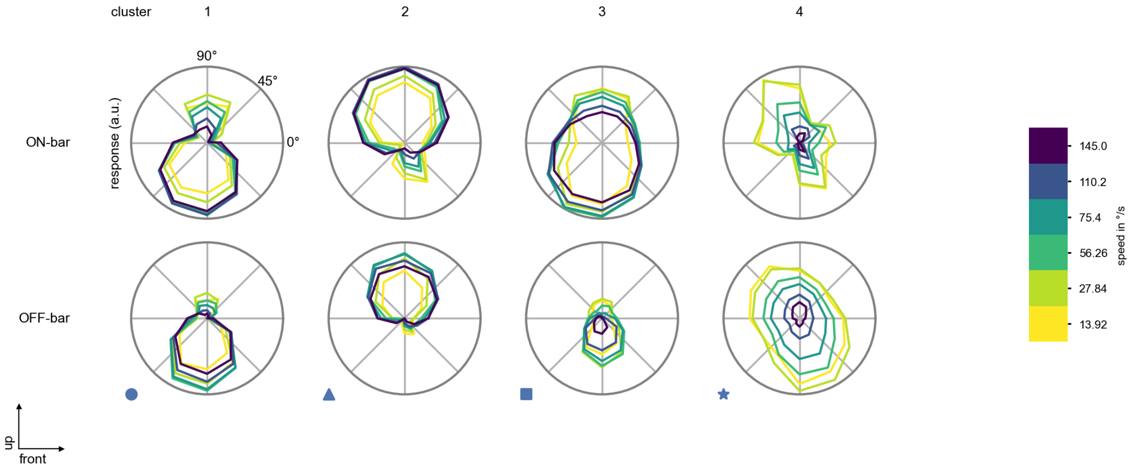

T4d - Figure 9: **Peak responses to moving bars from best-performing models.** The top row shows peak responses to moving ON-bars, the bottom row shows peak responses to moving OFF-bars. The bar-stimuli move in different directions from 0 to 360 degrees and at different speeds. Before, a grey-stimulus led the network to a stationary state. The columns show responses from the best-task-performing model in the respective cluster.

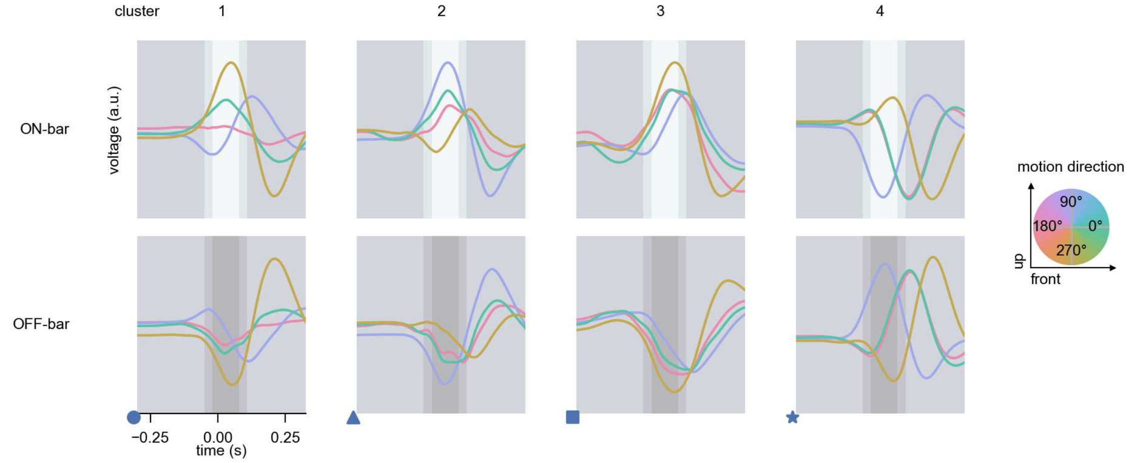

T4d - Figure 10: **Responses to moving bars from best-performing models.** The top row shows responses to moving ON-bars, the bottom row shows peak responses to moving OFF-bars. The bar-stimuli move in different directions from 0 to 360 degrees and at different speeds. Before, a grey-stimulus led the network to a stationary state. The columns show responses from the best-task-performing model in the respective cluster. Example responses to bars moving at 75.4°/s in all cardinal directions (green 0°, blue 90°, red 180°, yellow 270°) from -22.5 to 22.5° visual angle. Background contour indicates the bar, passing by the central column at second zero.

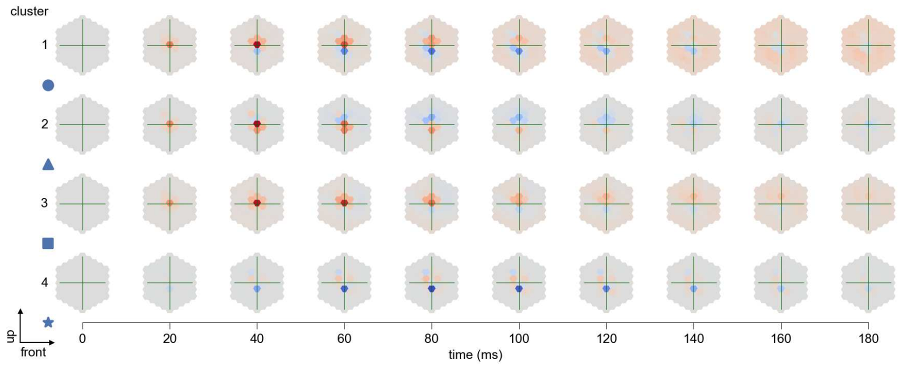

T4d - Figure 11: **Spatio-temporal receptive field.** Responses of the central cell to ON-impulses at single ommatidium impulse locations. The impulse occurs at second zero. Before, a grey-stimulus led the network to a stationary state. Responses are baseline-subtracted. Each row illustrates responses from the best-performing model of the respective cluster. Red indicated depolarization, blue indicates hyperpolarization.

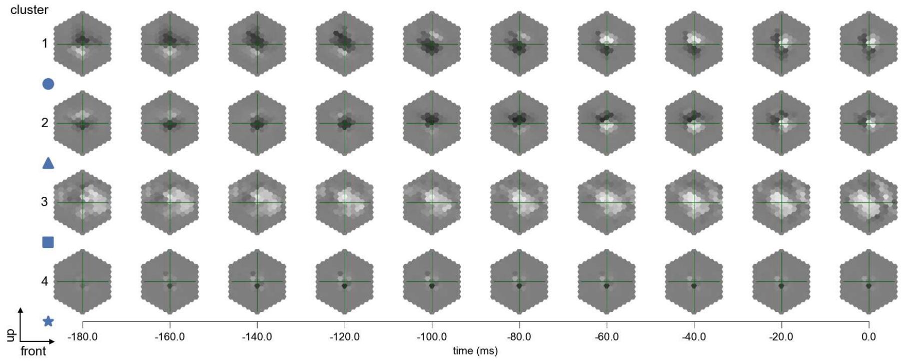

T4d - Figure 12: **Maximally excitatory stimuli.** Each row presents the regularized naturalistic-stimulus from the Sintel dataset that maximizes the cell type's central column response at second zero in the best-performing model of the respective cluster.

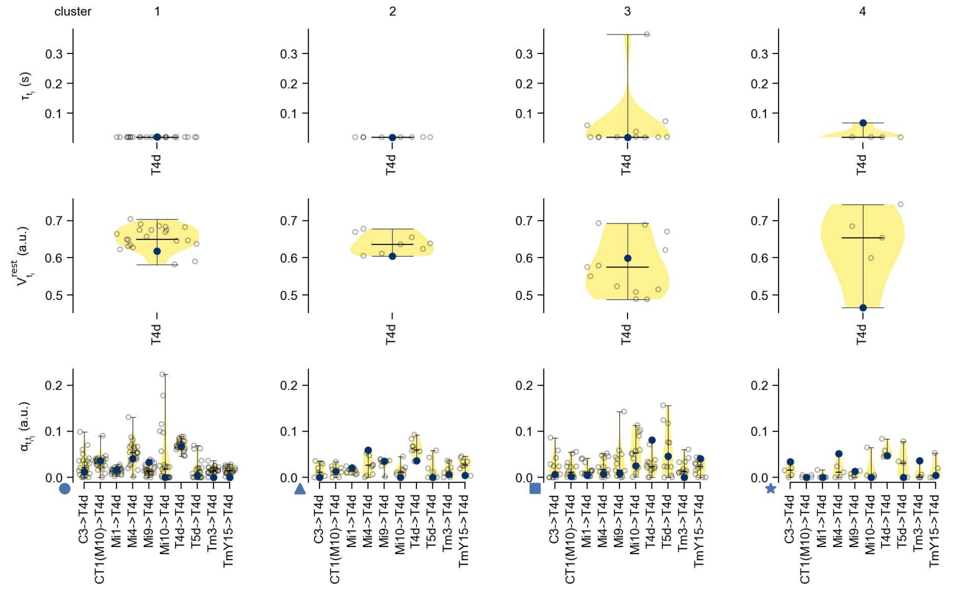

T4d - Figure 13: **Task-constrained parameters.** Each column shows the parameters inferred within the respective cluster. First row: learned time constants of the cell type. Second row: resting potentials of the cell type. Third row: scaling factors for the convolutional filters. The blue scatter represents the parameters from the best-performing model within the cluster.

# 40 T5a

← Cell types

#### Figures

1 Anatomical receptive fields. . . . 264
2 Anatomical projective fields. . . . 264
3 Clustering of the responses to naturalistic stimuli. . . . 265
4 Responses to flashes. . . . 265
5 Peak responses to moving edges. . . . 266
6 Peak responses to moving edges from best-performing models. . . . 266
7 Responses to moving edges from best-performing models. . . . 267
8 Peak responses to moving bars. . . . 267
9 Peak responses to moving bars from best-performing models. . . . 268
10 Responses to moving bars from best-performing models. . . . 268
11 Spatio-temporal receptive field. . . . 269
12 Maximally excitatory stimuli. . . . 269
13 Task-constrained parameters. . . . 269

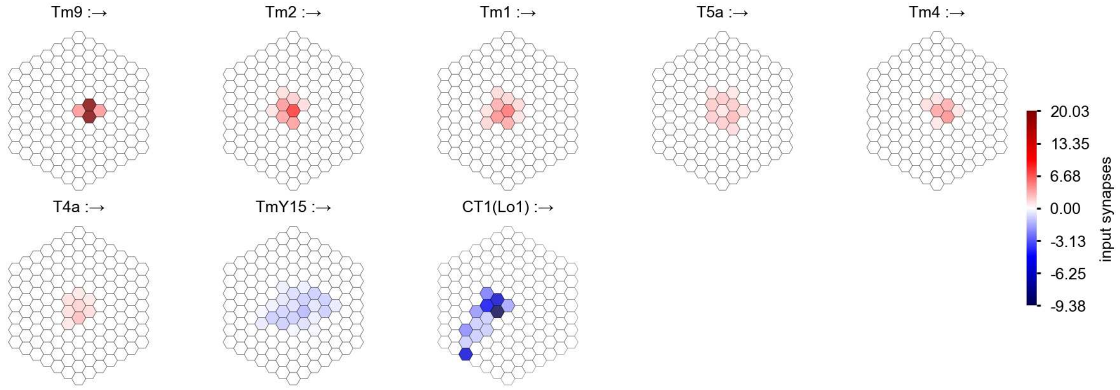

T5a - Figure 1: **Anatomical receptive fields.** Each colored hexagon is a projection, with the projection strength characterized by the average number of synapses that we count from the EM reconstruction. Red indicates excitatory synapses, blue indicates inhibitory synapses from inferred signs. Filters in the order of their total number of synapses.

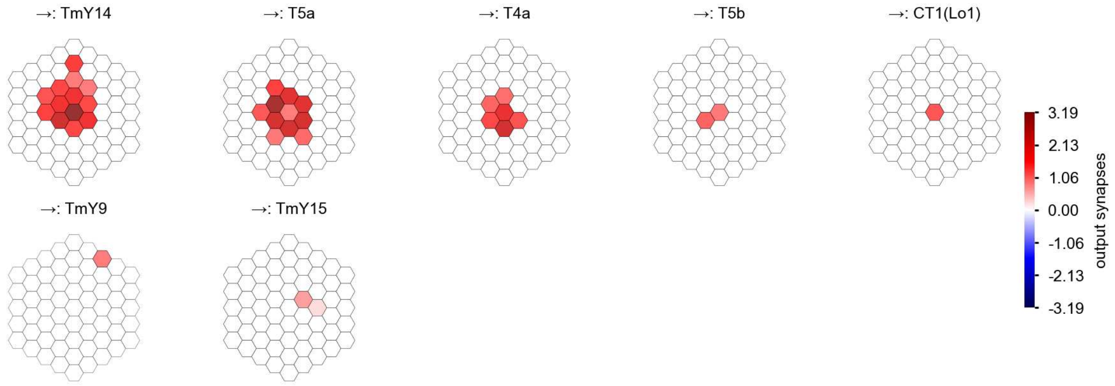

T5a - Figure 2: **Anatomical projective fields.** Each colored hexagon is a projection, with the projection strength characterized by the average number of synapses that we count from the EM reconstruction. Red indicates excitatory synapses, blue indicates inhibitory synapses from inferred signs. Filters in the order of their total number of synapses.

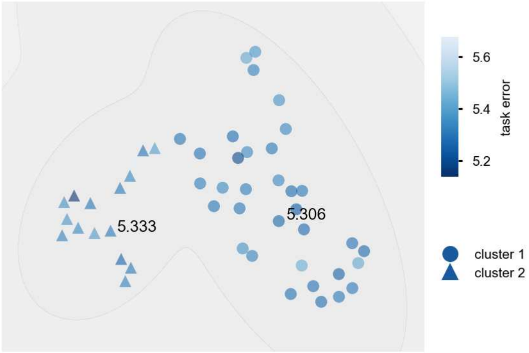

T5a - Figure 3: **Clustering of the responses to naturalistic stimuli.** From responses to naturalistic scenes from Sintel. Scatterpoints represent individual models colored by their task error.

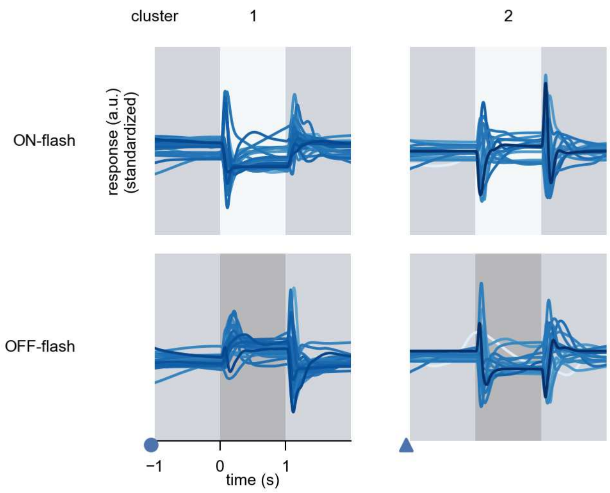

T5a - Figure 4: **Responses to flashes.** The top row shows responses to ON-flashes, the bottom row shows responses to OFF-flashes. The responses from the different models in the different clusters (columns) overlay and are colored in the respective task error of the model. The circular flashes occur at second zero, cover 6 ommatidia in radius, and last for one second. Before and after, a grey-stimulus leads to a stationary state of the network. The traces are standardized.

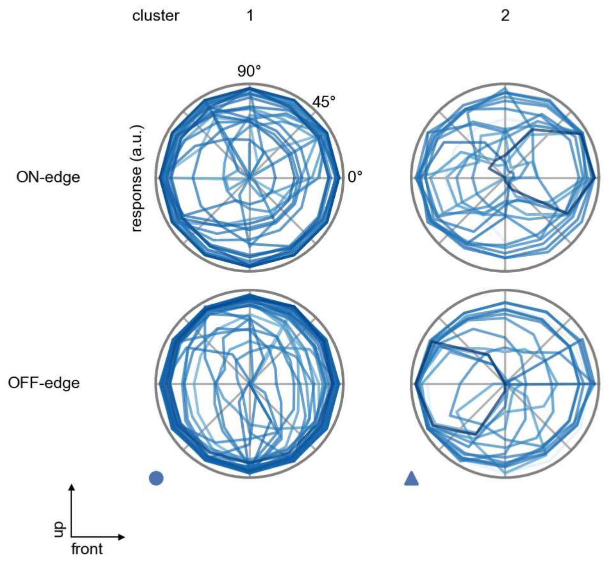

T5a - Figure 5: **Peak responses to moving edges.** The top row shows normalized peak responses to moving ON-edges, the bottom row shows normalized peak responses to moving OFF-edges. The edge-stimuli move in different directions from 0 to 360 degrees. Before, a grey-stimulus led the network to a stationary state. The responses from the different models in the different clusters (columns) overlay and are colored in the respective task error of the model. The peak responses are averaged over all assessed edge-speeds.

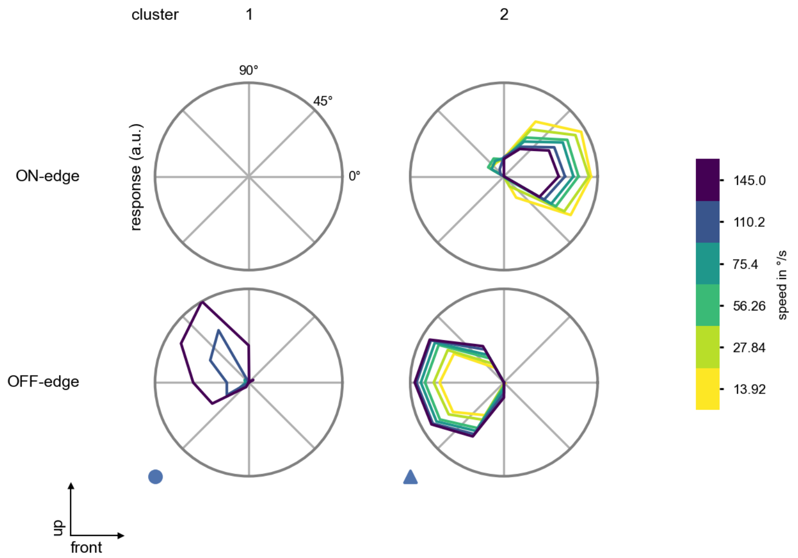

T5a - Figure 6: **Peak responses to moving edges from best-performing models.** The top row shows peak responses to moving ON-edges, the bottom row shows peak responses to moving OFF-edges. The edge-stimuli move in different directions from 0 to 360 degrees and at different speeds. Before, a grey-stimulus led the network to a stationary state. The columns show responses from the best-task-performing model in the respective cluster.

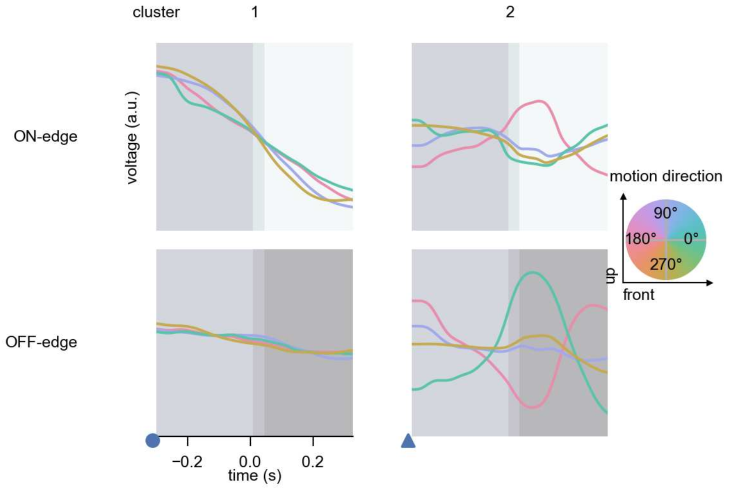

T5a - Figure 7: **Responses to moving edges from best-performing models.** The top row shows responses to moving ON-edges, the bottom row shows peak responses to moving OFF-edges. The edge-stimuli move in different directions from 0 to 360 degrees and at different speeds. Before, a grey-stimulus led the network to a stationary state. The columns show responses from the best-task-performing model in the respective cluster. Example responses to edges moving at 75.4°/s in all cardinal directions (green 0°, blue 90°, red 180°, yellow 270°) from -22.5 to 22.5° visual angle. Background contour indicates the edge, passing by the central column at second zero.

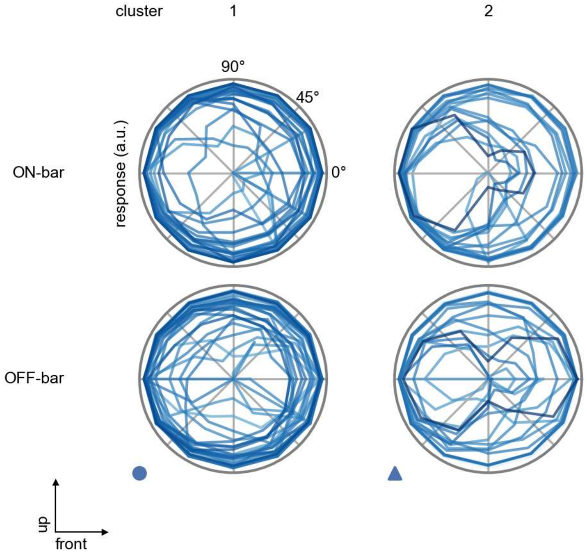

T5a - Figure 8: **Peak responses to moving bars.** The top row shows normalized peak responses to moving ON-bars, the bottom row shows normalized peak responses to moving OFF-bars. The bar-stimuli move in different directions from 0 to 360 degrees. Before, a grey-stimulus led the network to a stationary state. The responses from the different models in the different clusters (columns) overlay and are colored in the respective task error of the model. The peak responses are averaged over all assessed bar-speeds.

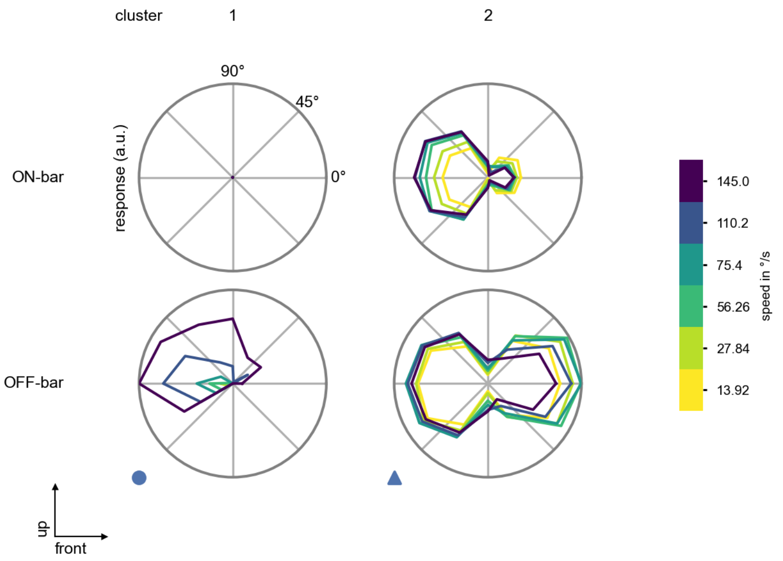

T5a - Figure 9: **Peak responses to moving bars from best-performing models.** The top row shows peak responses to moving ON-bars, the bottom row shows peak responses to moving OFF-bars. The bar-stimuli move in different directions from 0 to 360 degrees and at different speeds. Before, a grey-stimulus led the network to a stationary state. The columns show responses from the best-task-performing model in the respective cluster.

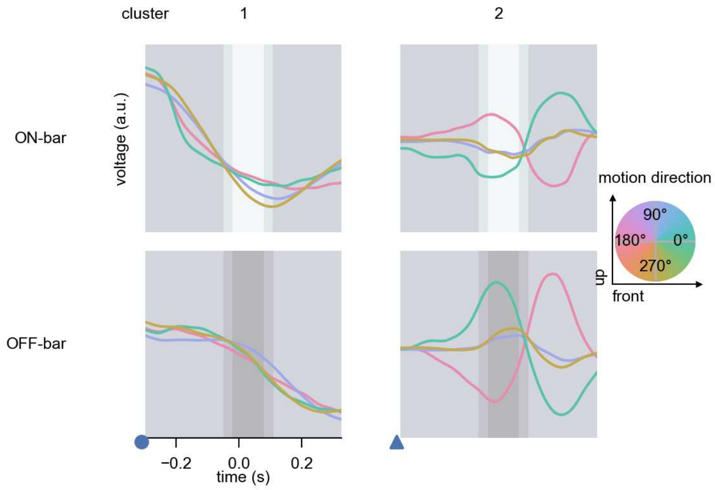

T5a - Figure 10: **Responses to moving bars from best-performing models.** The top row shows responses to moving ON-bars, the bottom row shows peak responses to moving OFF-bars. The bar-stimuli move in different directions from 0 to 360 degrees and at different speeds. Before, a grey-stimulus led the network to a stationary state. The columns show responses from the best-task-performing model in the respective cluster. Example responses to bars moving at 75.4°/s in all cardinal directions (green 0°, blue 90°, red 180°, yellow 270°) from -22.5 to 22.5° visual angle. Background contour indicates the bar, passing by the central column at second zero.

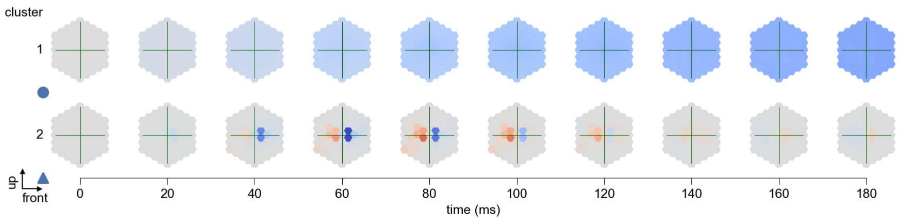

T5a - Figure 11: **Spatio-temporal receptive field.** Responses of the central cell to ON-impulses at single ommatidium impulse locations. The impulse occurs at second zero. Before, a grey-stimulus led the network to a stationary state. Responses are baseline-subtracted. Each row illustrates responses from the best-performing model of the respective cluster. Red indicated depolarization, blue indicates hyperpolarization.

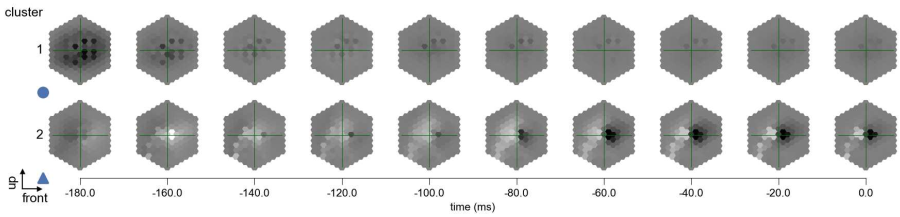

T5a - Figure 12: **Maximally excitatory stimuli.** Each row presents the regularized naturalistic-stimulus from the Sintel dataset that maximizes the cell type's central column response at second zero in the best-performing model of the respective cluster.

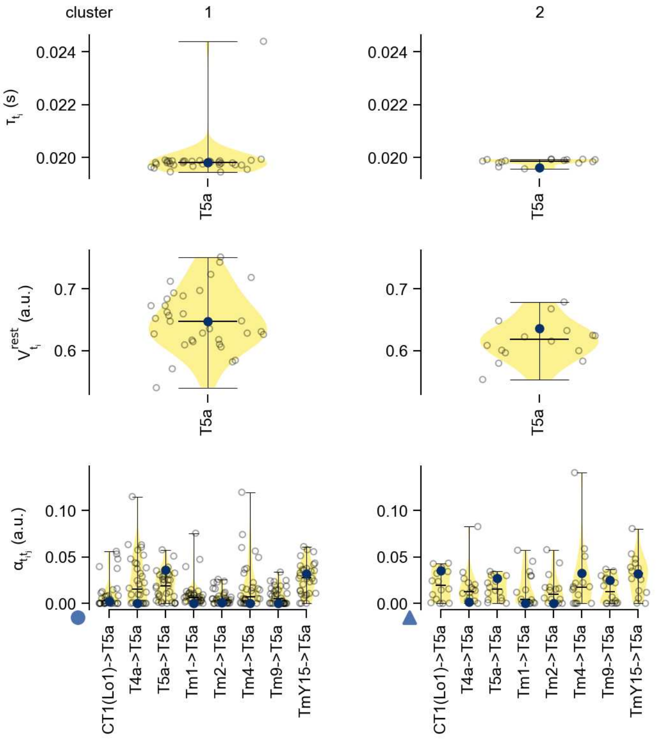

T5a - Figure 13: **Task-constrained parameters.** Each column shows the parameters inferred within the respective cluster. First row: learned time constants of the cell type. Second row: resting potentials of the cell type. Third row: scaling factors for the convolutional filters. The blue scatter represents the parameters from the best-performing model within the cluster.

# 41 T5b

← Cell types

#### Figures

1 Anatomical receptive fields. . . . 270
2 Anatomical projective fields. . . . 270
3 Clustering of the responses to naturalistic stimuli. . . . 271
4 Responses to flashes. . . . 271
5 Peak responses to moving edges. . . . 271
6 Peak responses to moving edges from best-performing models. . . . 272
7 Responses to moving edges from best-performing models. . . . 272
8 Peak responses to moving bars. . . . 273
9 Peak responses to moving bars from best-performing models. . . . 273
10 Responses to moving bars from best-performing models. . . . 274
11 Spatio-temporal receptive field. . . . 274
12 Maximally excitatory stimuli. . . . 275
13 Task-constrained parameters. . . . 275

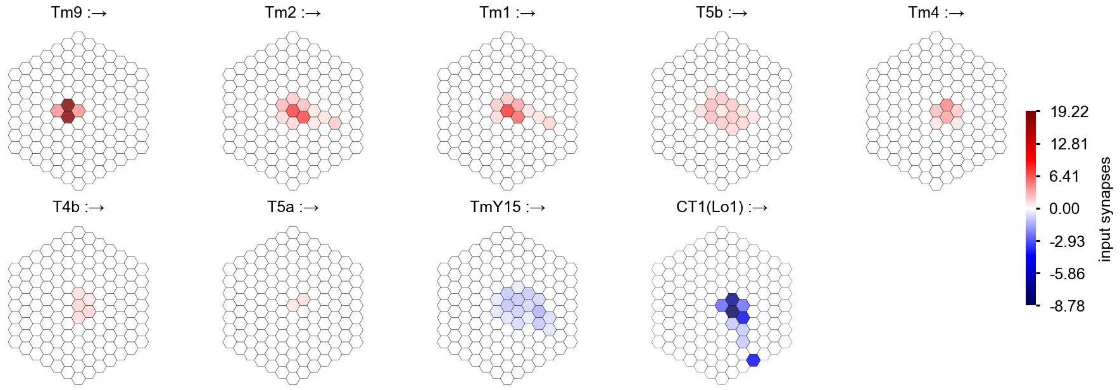

T5b - Figure 1: **Anatomical receptive fields.** Each colored hexagon is a projection, with the projection strength characterized by the average number of synapses that we count from the EM reconstruction. Red indicates excitatory synapses, blue indicates inhibitory synapses from inferred signs. Filters in the order of their total number of synapses.

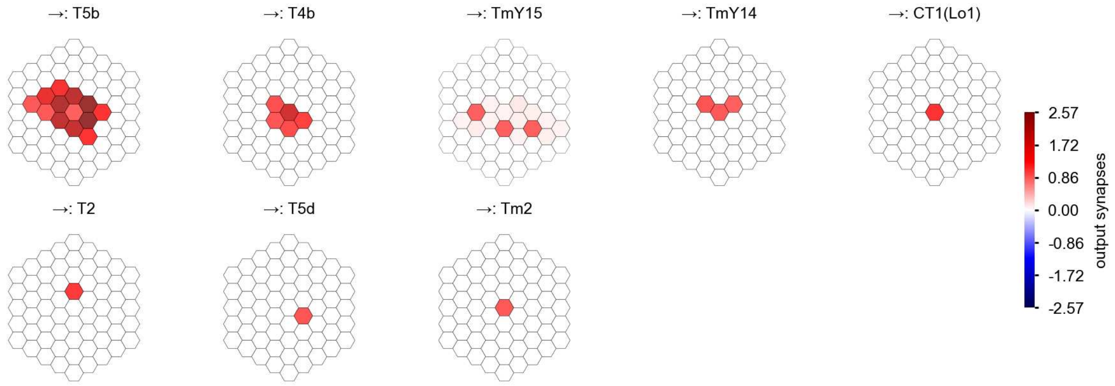

T5b - Figure 2: **Anatomical projective fields.** Each colored hexagon is a projection, with the projection strength characterized by the average number of synapses that we count from the EM reconstruction. Red indicates excitatory synapses, blue indicates inhibitory synapses from inferred signs. Filters in the order of their total number of synapses.

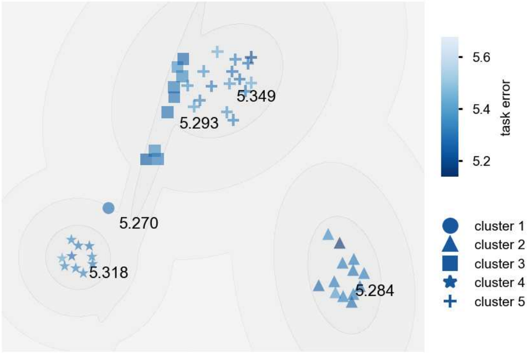

T5b - Figure 3: **Clustering of the responses to naturalistic stimuli.** From responses to naturalistic scenes from Sintel. Scatterpoints represent individual models colored by their task error.

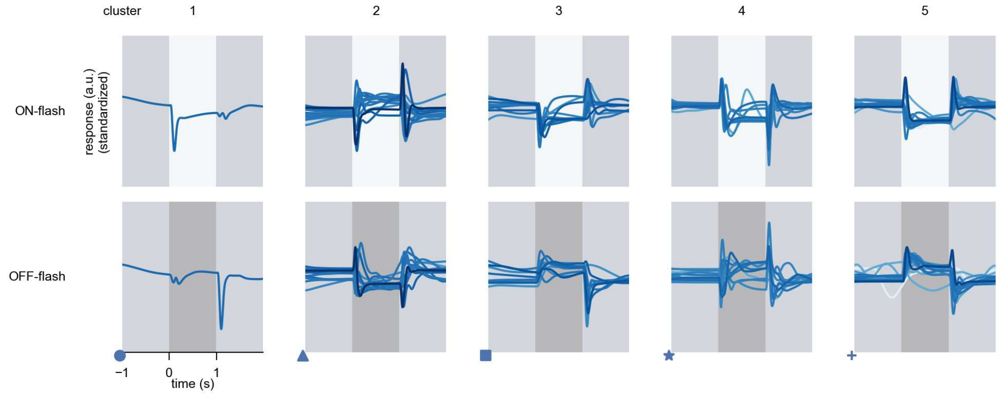

T5b - Figure 4: **Responses to flashes.** The top row shows responses to ON-flashes, the bottom row shows responses to OFF-flashes. The responses from the different models in the different clusters (columns) overlay and are colored in the respective task error of the model. The circular flashes occur at second zero, cover 6 ommatidia in radius, and last for one second. Before and after, a grey-stimulus leads to a stationary state of the network. The traces are standardized.

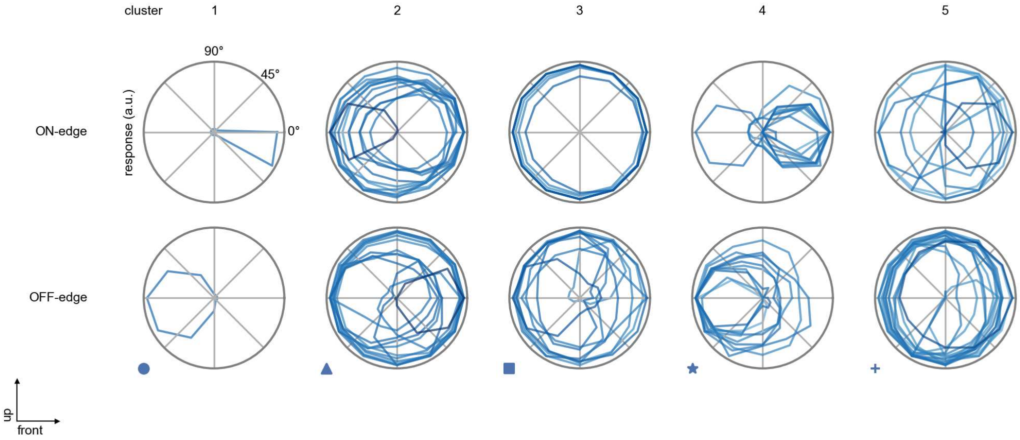

T5b - Figure 5: **Peak responses to moving edges.** The top row shows normalized peak responses to moving ON-edges, the bottom row shows normalized peak responses to moving OFF-edges. The edge-stimuli move in different directions from 0 to 360 degrees. Before, a grey-stimulus led the network to a stationary state. The responses from the different models in the different clusters (columns) overlay and are colored in the respective task error of the model. The peak responses are averaged over all assessed edge-speeds.

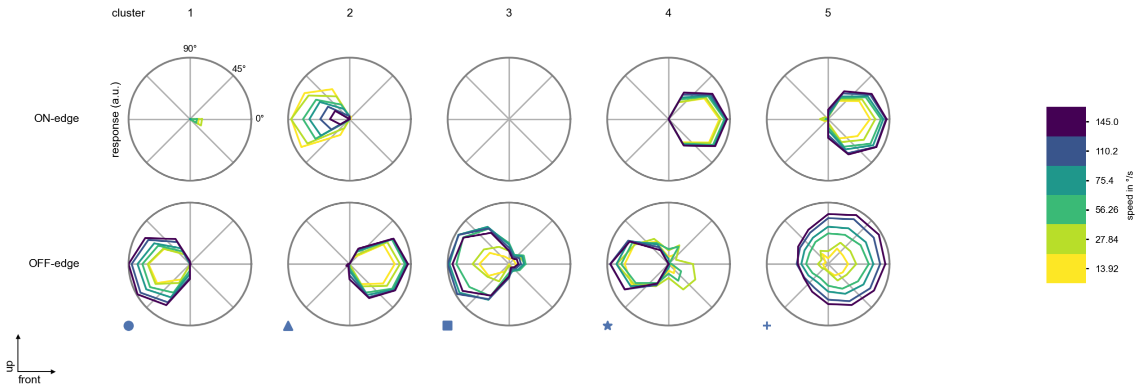

T5b - Figure 6: **Peak responses to moving edges from best-performing models.** The top row shows peak responses to moving ON-edges, the bottom row shows peak responses to moving OFF-edges. The edge-stimuli move in different directions from 0 to 360 degrees and at different speeds. Before, a grey-stimulus led the network to a stationary state. The columns show responses from the best-task-performing model in the respective cluster.

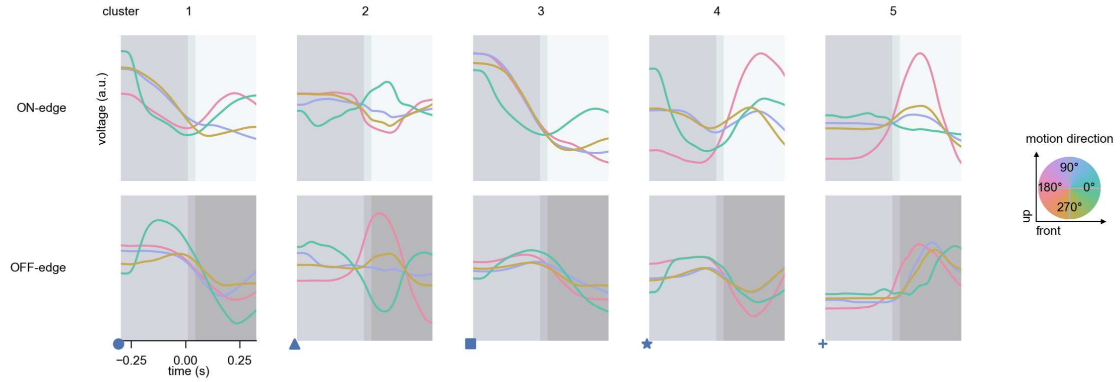

T5b - Figure 7: **Responses to moving edges from best-performing models.** The top row shows responses to moving ON-edges, the bottom row shows peak responses to moving OFF-edges. The edge-stimuli move in different directions from 0 to 360 degrees and at different speeds. Before, a grey-stimulus led the network to a stationary state. The columns show responses from the best-task-performing model in the respective cluster. Example responses to edges moving at 75.4°/s in all cardinal directions (green 0°, blue 90°, red 180°, yellow 270°) from -22.5 to 22.5° visual angle. Background contour indicates the edge, passing by the central column at second zero.

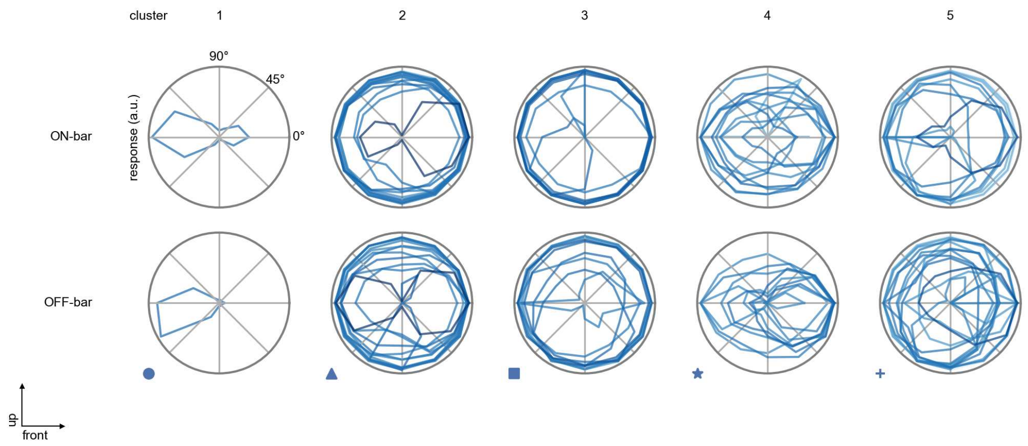

T5b - Figure 8: **Peak responses to moving bars.** The top row shows normalized peak responses to moving ON-bars, the bottom row shows normalized peak responses to moving OFF-bars. The bar-stimuli move in different directions from 0 to 360 degrees. Before, a grey-stimulus led the network to a stationary state. The responses from the different models in the different clusters (columns) overlay and are colored in the respective task error of the model. The peak responses are averaged over all assessed bar-speeds.

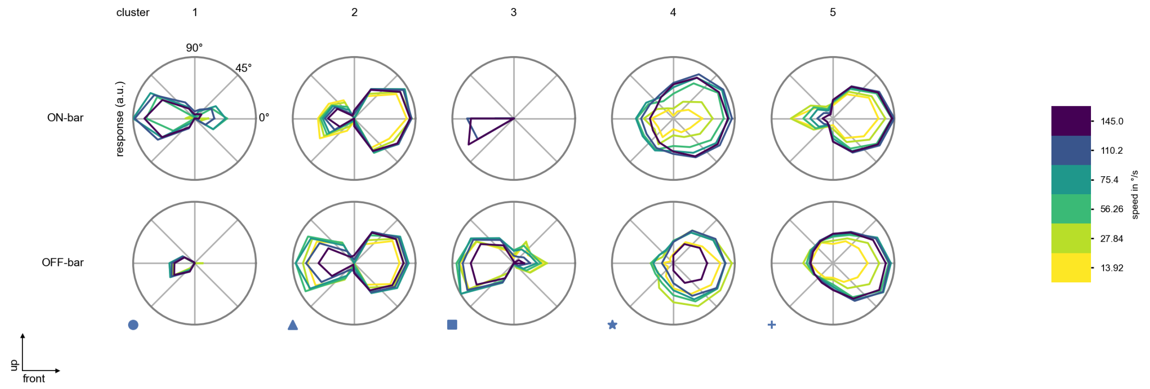

T5b - Figure 9: **Peak responses to moving bars from best-performing models.** The top row shows peak responses to moving ON-bars, the bottom row shows peak responses to moving OFF-bars. The bar-stimuli move in different directions from 0 to 360 degrees and at different speeds. Before, a grey-stimulus led the network to a stationary state. The columns show responses from the best-task-performing model in the respective cluster.

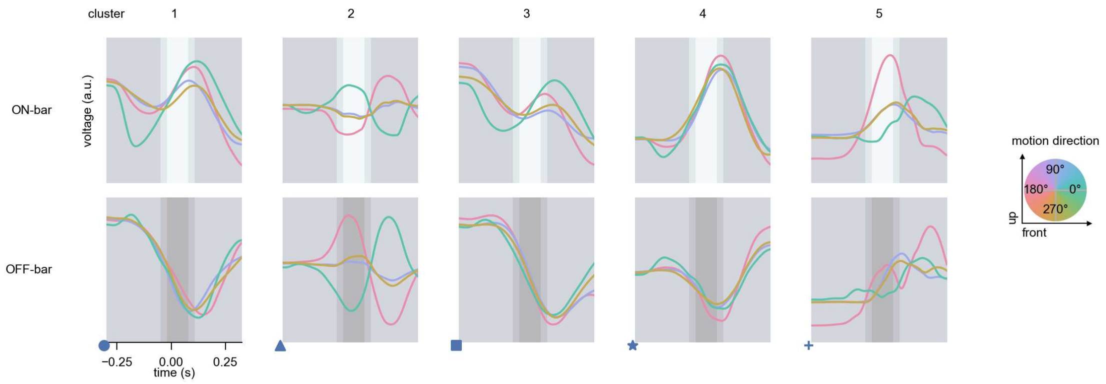

T5b - Figure 10: **Responses to moving bars from best-performing models.** The top row shows responses to moving ON-bars, the bottom row shows peak responses to moving OFF-bars. The bar-stimuli move in different directions from 0 to 360 degrees and at different speeds. Before, a grey-stimulus led the network to a stationary state. The columns show responses from the best-task-performing model in the respective cluster. Example responses to bars moving at 75.4°/s in all cardinal directions (green 0°, blue 90°, red 180°, yellow 270°) from -22.5 to 22.5° visual angle. Background contour indicates the bar, passing by the central column at second zero.

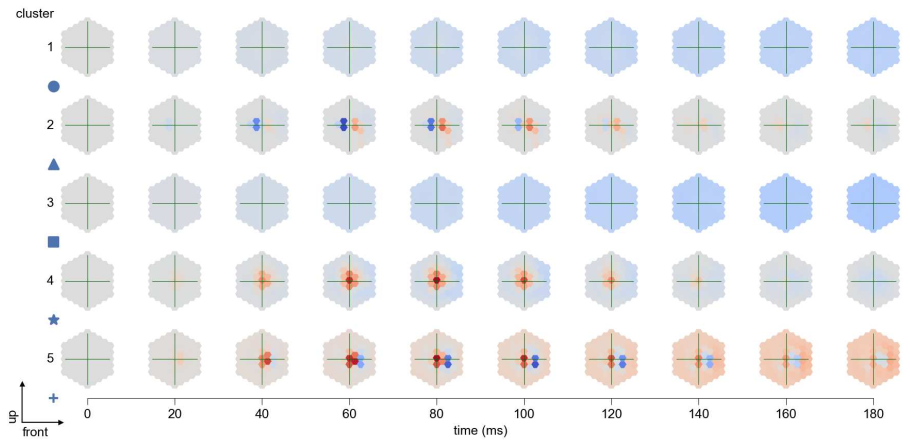

T5b - Figure 11: **Spatio-temporal receptive field.** Responses of the central cell to ON-impulses at single ommatidium impulse locations. The impulse occurs at second zero. Before, a grey-stimulus led the network to a stationary state. Responses are baseline-subtracted. Each row illustrates responses from the best-performing model of the respective cluster. Red indicated depolarization, blue indicates hyperpolarization.

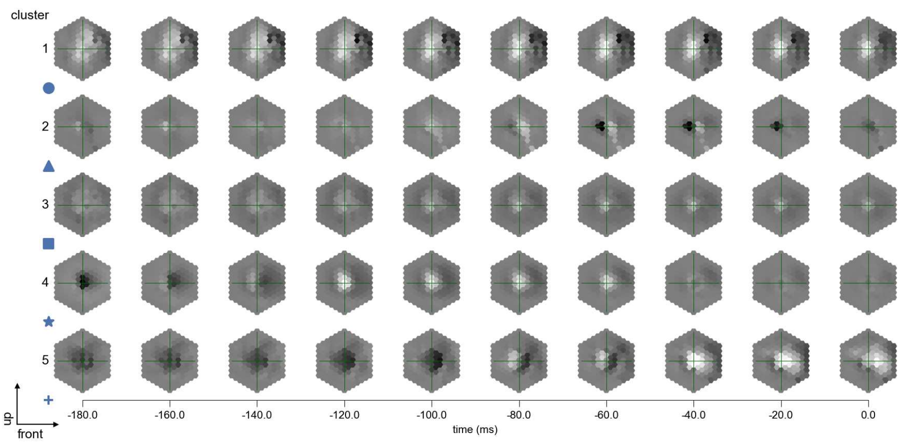

T5b - Figure 12: **Maximally excitatory stimuli.** Each row presents the regularized naturalistic-stimulus from the Sintel dataset that maximizes the cell type's central column response at second zero in the best-performing model of the respective cluster.

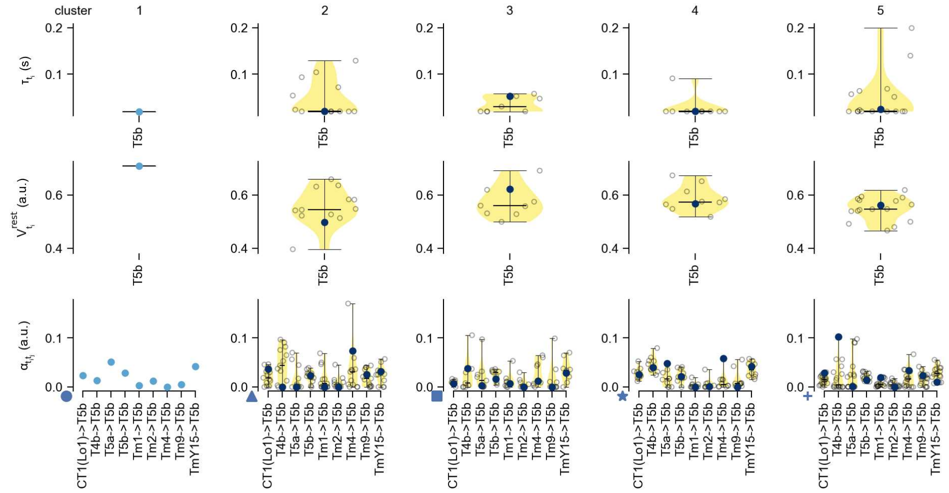

T5b - Figure 13: **Task-constrained parameters.** Each column shows the parameters inferred within the respective cluster. First row: learned time constants of the cell type. Second row: resting potentials of the cell type. Third row: scaling factors for the convolutional filters. The blue scatter represents the parameters from the best-performing model within the cluster.

# 42 T5c

← Cell types

#### Figures

1 Anatomical receptive fields. . . . . 276
2 Anatomical projective fields. . . . . 276
3 Clustering of the responses to naturalistic stimuli. . . . . 277
4 Responses to flashes. . . . . 277
5 Peak responses to moving edges. . . . . 278
6 Peak responses to moving edges from best-performing models. . . . . 278
7 Responses to moving edges from best-performing models. . . . . 279
8 Peak responses to moving bars. . . . . 279
9 Peak responses to moving bars from best-performing models. . . . . 280
10 Responses to moving bars from best-performing models. . . . . 280
11 Spatio-temporal receptive field. . . . . 281
12 Maximally excitatory stimuli. . . . . 281
13 Task-constrained parameters. . . . . 282

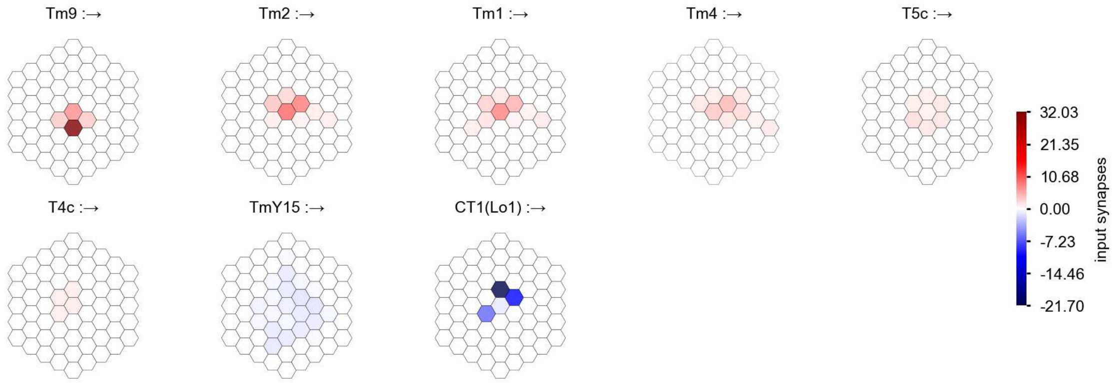

T5c - Figure 1: **Anatomical receptive fields.** Each colored hexagon is a projection, with the projection strength characterized by the average number of synapses that we count from the EM reconstruction. Red indicates excitatory synapses, blue indicates inhibitory synapses from inferred signs. Filters in the order of their total number of synapses.

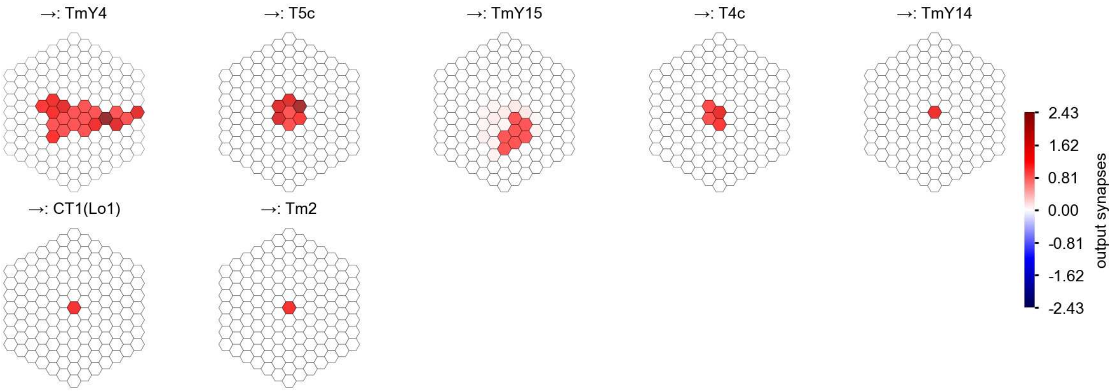

T5c - Figure 2: **Anatomical projective fields.** Each colored hexagon is a projection, with the projection strength characterized by the average number of synapses that we count from the EM reconstruction. Red indicates excitatory synapses, blue indicates inhibitory synapses from inferred signs. Filters in the order of their total number of synapses.

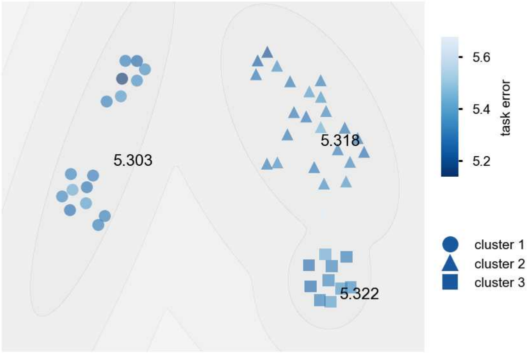

T5c - Figure 3: **Clustering of the responses to naturalistic stimuli.** From responses to naturalistic scenes from Sintel. Scatterpoints represent individual models colored by their task error.

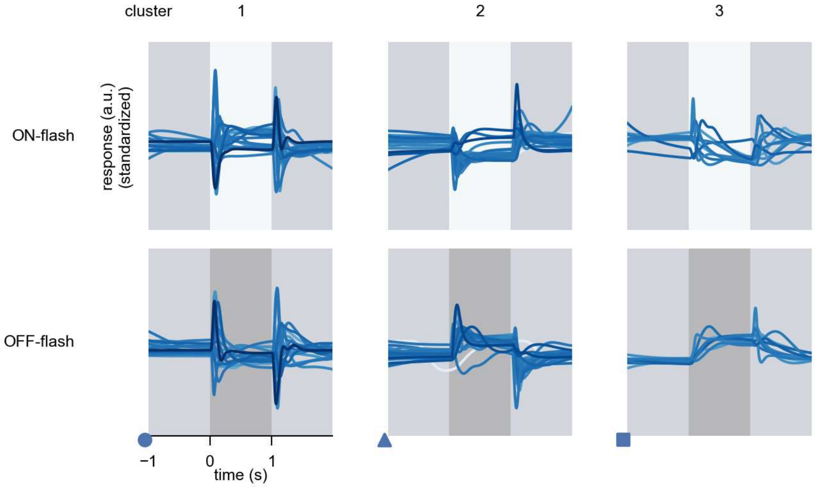

T5c - Figure 4: **Responses to flashes.** The top row shows responses to ON-flashes, the bottom row shows responses to OFF-flashes. The responses from the different models in the different clusters (columns) overlay and are colored in the respective task error of the model. The circular flashes occur at second zero, cover 6 ommatidia in radius, and last for one second. Before and after, a grey-stimulus leads to a stationary state of the network. The traces are standardized.

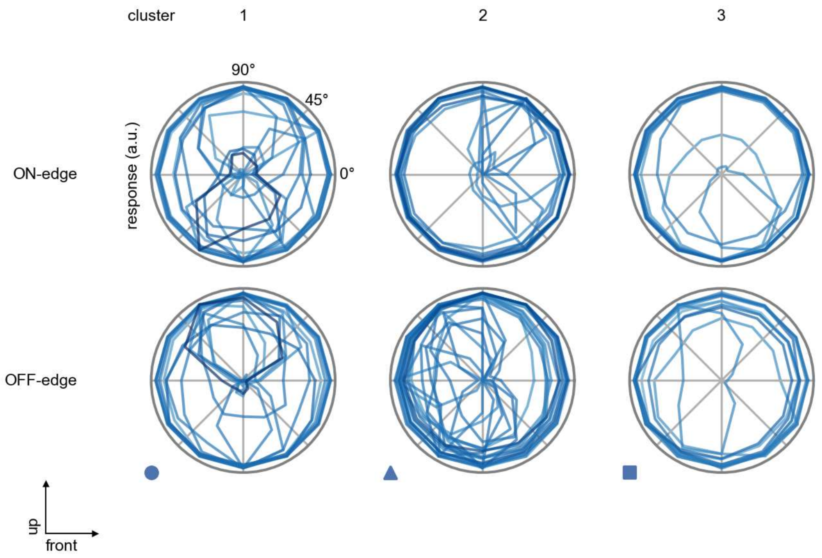

T5c - Figure 5: **Peak responses to moving edges.** The top row shows normalized peak responses to moving ON-edges, the bottom row shows normalized peak responses to moving OFF-edges. The edge-stimuli move in different directions from 0 to 360 degrees. Before, a grey-stimulus led the network to a stationary state. The responses from the different models in the different clusters (columns) overlay and are colored in the respective task error of the model. The peak responses are averaged over all assessed edge-speeds.

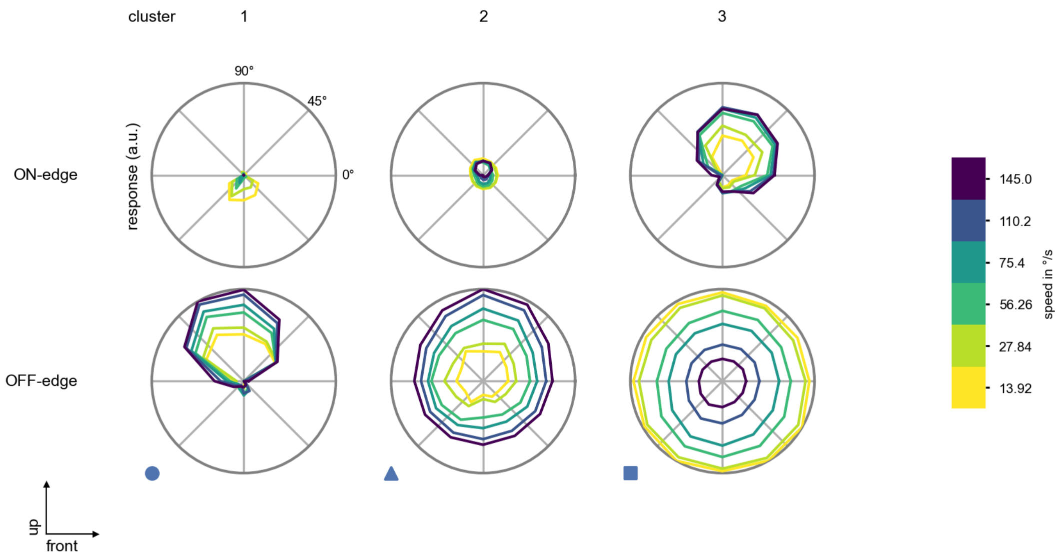

T5c - Figure 6: **Peak responses to moving edges from best-performing models.** The top row shows peak responses to moving ON-edges, the bottom row shows peak responses to moving OFF-edges. The edge-stimuli move in different directions from 0 to 360 degrees and at different speeds. Before, a grey-stimulus led the network to a stationary state. The columns show responses from the best-task-performing model in the respective cluster.

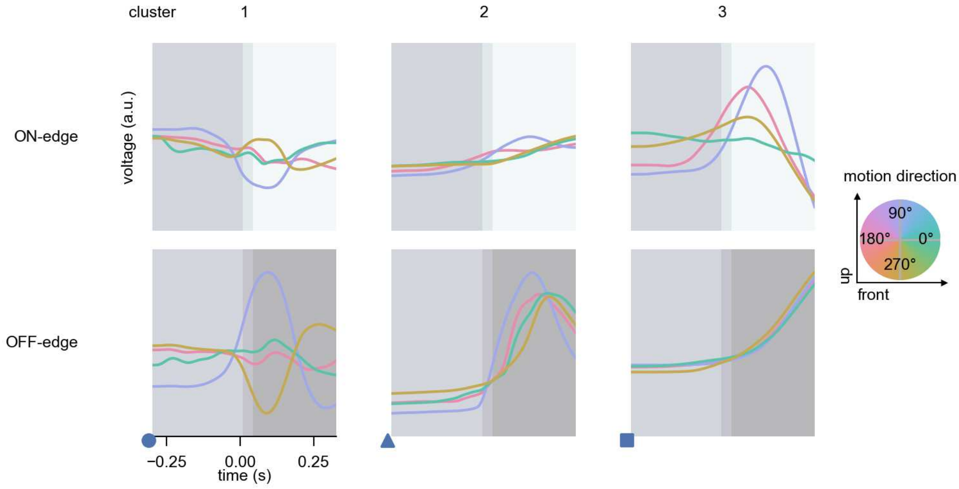

T5c - Figure 7: **Responses to moving edges from best-performing models.** The top row shows responses to moving ON-edges, the bottom row shows peak responses to moving OFF-edges. The edge-stimuli move in different directions from 0 to 360 degrees and at different speeds. Before, a grey-stimulus led the network to a stationary state. The columns show responses from the best-task-performing model in the respective cluster. Example responses to edges moving at 75.4°/s in all cardinal directions (green 0°, blue 90°, red 180°, yellow 270°) from -22.5 to 22.5° visual angle. Background contour indicates the edge, passing by the central column at second zero.

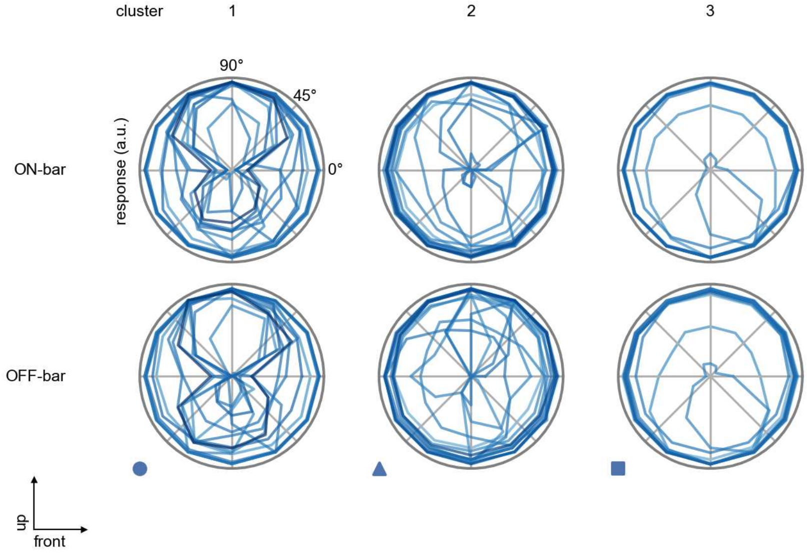

T5c - Figure 8: **Peak responses to moving bars.** The top row shows normalized peak responses to moving ON-bars, the bottom row shows normalized peak responses to moving OFF-bars. The bar-stimuli move in different directions from 0 to 360 degrees. Before, a grey-stimulus led the network to a stationary state. The responses from the different models in the different clusters (columns) overlay and are colored in the respective task error of the model. The peak responses are averaged over all assessed bar-speeds.

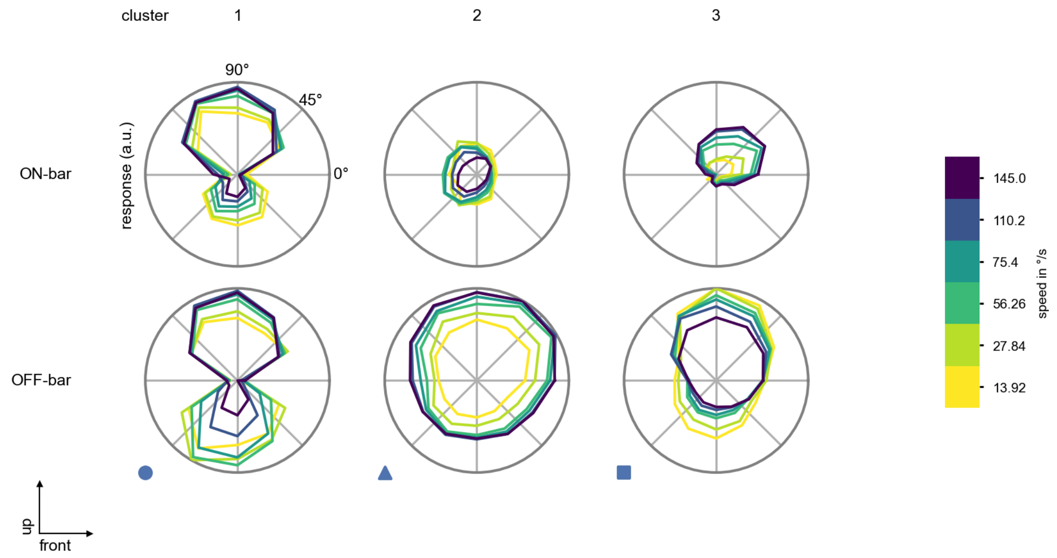

T5c - Figure 9: **Peak responses to moving bars from best-performing models.** The top row shows peak responses to moving ON-bars, the bottom row shows peak responses to moving OFF-bars. The bar-stimuli move in different directions from 0 to 360 degrees and at different speeds. Before, a grey-stimulus led the network to a stationary state. The columns show responses from the best-task-performing model in the respective cluster.

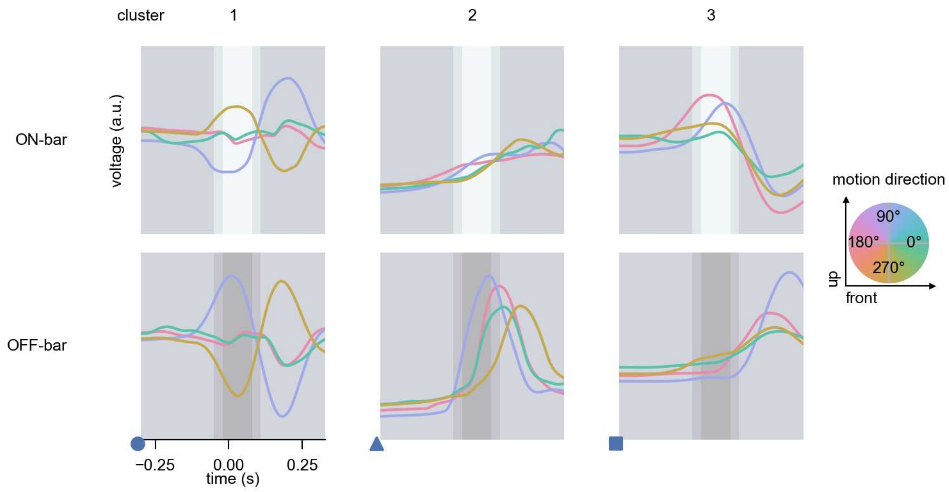

T5c - Figure 10: **Responses to moving bars from best-performing models.** The top row shows responses to moving ON-bars, the bottom row shows peak responses to moving OFF-bars. The bar-stimuli move in different directions from 0 to 360 degrees and at different speeds. Before, a grey-stimulus led the network to a stationary state. The columns show responses from the best-task-performing model in the respective cluster. Example responses to bars moving at 75.4°/s in all cardinal directions (green 0°, blue 90°, red 180°, yellow 270°) from -22.5 to 22.5° visual angle. Background contour indicates the bar, passing by the central column at second zero.

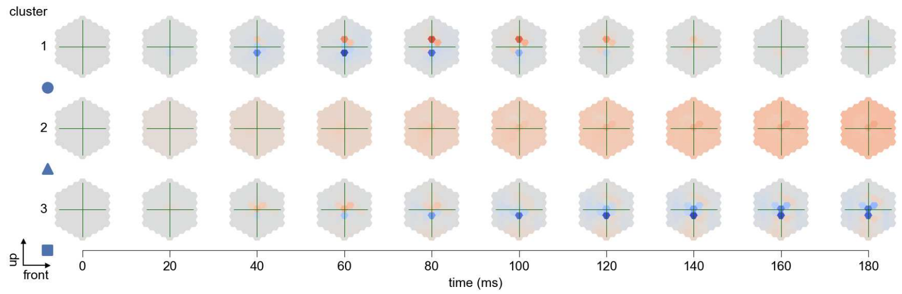

T5c - Figure 11: **Spatio-temporal receptive field.** Responses of the central cell to ON-impulses at single ommatidium impulse locations. The impulse occurs at second zero. Before, a grey-stimulus led the network to a stationary state. Responses are baseline-subtracted. Each row illustrates responses from the best-performing model of the respective cluster. Red indicated depolarization, blue indicates hyperpolarization.

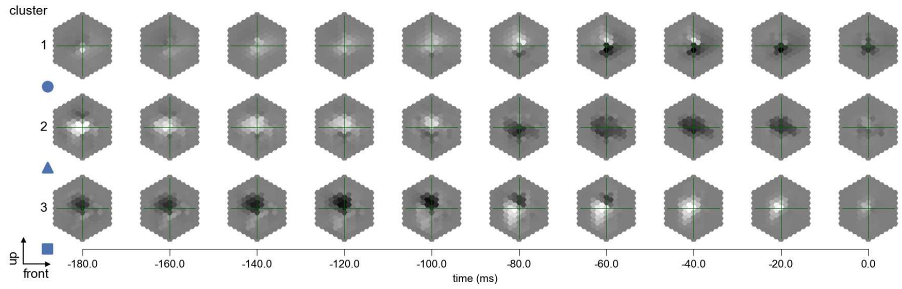

T5c - Figure 12: **Maximally excitatory stimuli.** Each row presents the regularized naturalistic-stimulus from the Sintel dataset that maximizes the cell type's central column response at second zero in the best-performing model of the respective cluster.

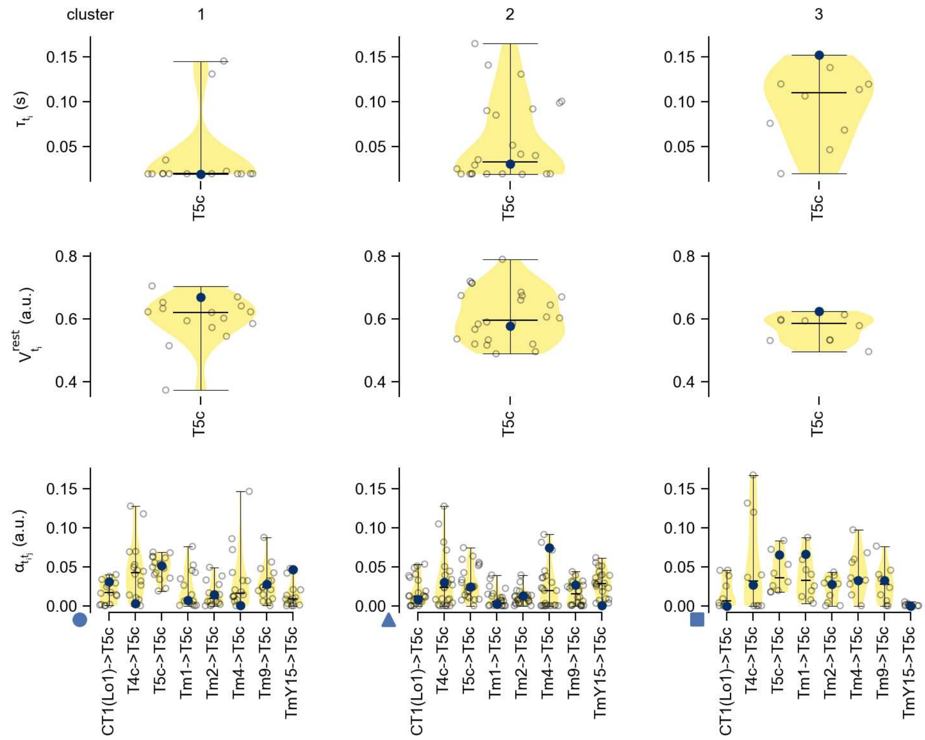

T5c - Figure 13: **Task-constrained parameters.** Each column shows the parameters inferred within the respective cluster. First row: learned time constants of the cell type. Second row: resting potentials of the cell type. Third row: scaling factors for the convolutional filters. The blue scatter represents the parameters from the best-performing model within the cluster.

# 43 T5d

← Cell types

#### Figures

1 Anatomical receptive fields. . . . 283
2 Anatomical projective fields. . . . 283
3 Clustering of the responses to naturalistic stimuli. . . . 284
4 Responses to flashes. . . . 284
5 Peak responses to moving edges. . . . 285
6 Peak responses to moving edges from best-performing models. . . . 285
7 Responses to moving edges from best-performing models. . . . 286
8 Peak responses to moving bars. . . . 286
9 Peak responses to moving bars from best-performing models. . . . 287
10 Responses to moving bars from best-performing models. . . . 287
11 Spatio-temporal receptive field. . . . 288
12 Maximally excitatory stimuli. . . . 288
13 Task-constrained parameters. . . . 289

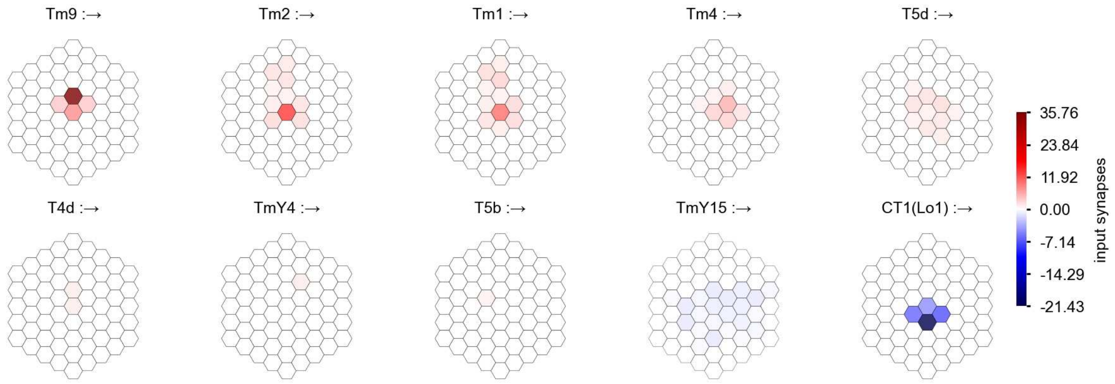

T5d - Figure 1: **Anatomical receptive fields.** Each colored hexagon is a projection, with the projection strength characterized by the average number of synapses that we count from the EM reconstruction. Red indicates excitatory synapses, blue indicates inhibitory synapses from inferred signs. Filters in the order of their total number of synapses.

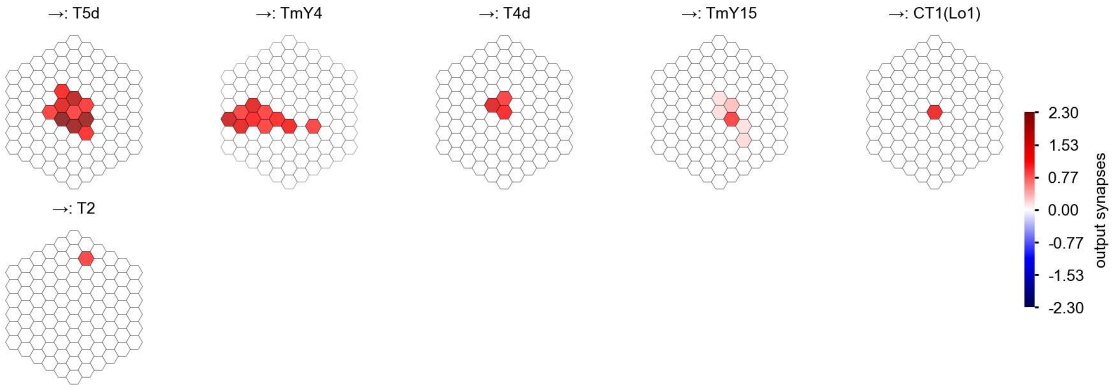

T5d - Figure 2: **Anatomical projective fields.** Each colored hexagon is a projection, with the projection strength characterized by the average number of synapses that we count from the EM reconstruction. Red indicates excitatory synapses, blue indicates inhibitory synapses from inferred signs. Filters in the order of their total number of synapses.

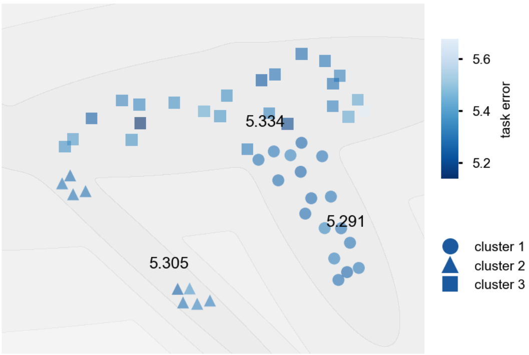

T5d - Figure 3: **Clustering of the responses to naturalistic stimuli.** From responses to naturalistic scenes from Sintel. Scatterpoints represent individual models colored by their task error.

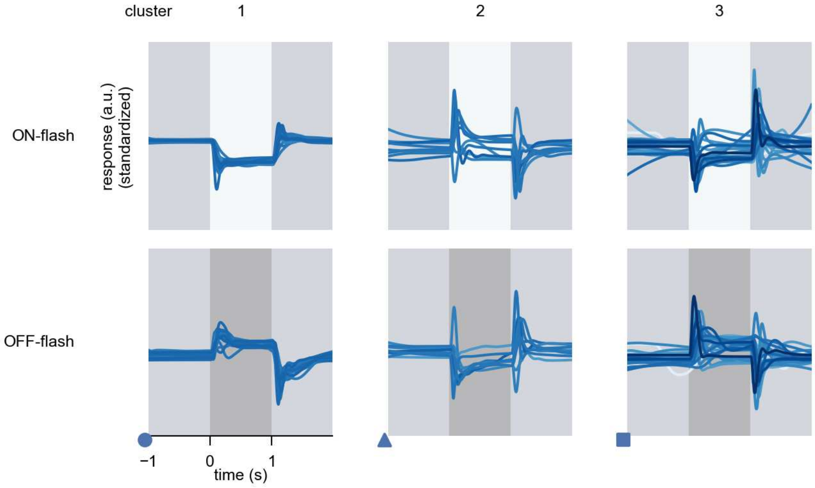

T5d - Figure 4: **Responses to flashes.** The top row shows responses to ON-flashes, the bottom row shows responses to OFF-flashes. The responses from the different models in the different clusters (columns) overlay and are colored in the respective task error of the model. The circular flashes occur at second zero, cover 6 ommatidia in radius, and last for one second. Before and after, a grey-stimulus leads to a stationary state of the network. The traces are standardized.

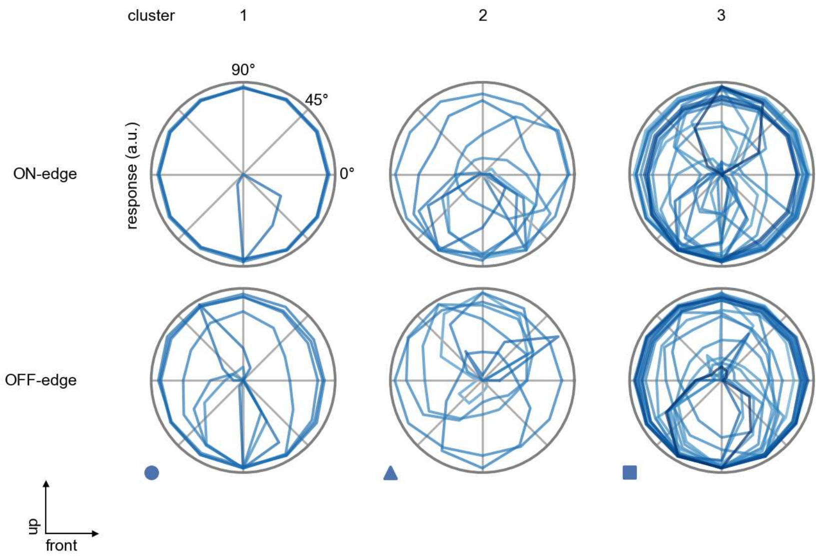

T5d - Figure 5: **Peak responses to moving edges.** The top row shows normalized peak responses to moving ON-edges, the bottom row shows normalized peak responses to moving OFF-edges. The edge-stimuli move in different directions from 0 to 360 degrees. Before, a grey-stimulus led the network to a stationary state. The responses from the different models in the different clusters (columns) overlay and are colored in the respective task error of the model. The peak responses are averaged over all assessed edge-speeds.

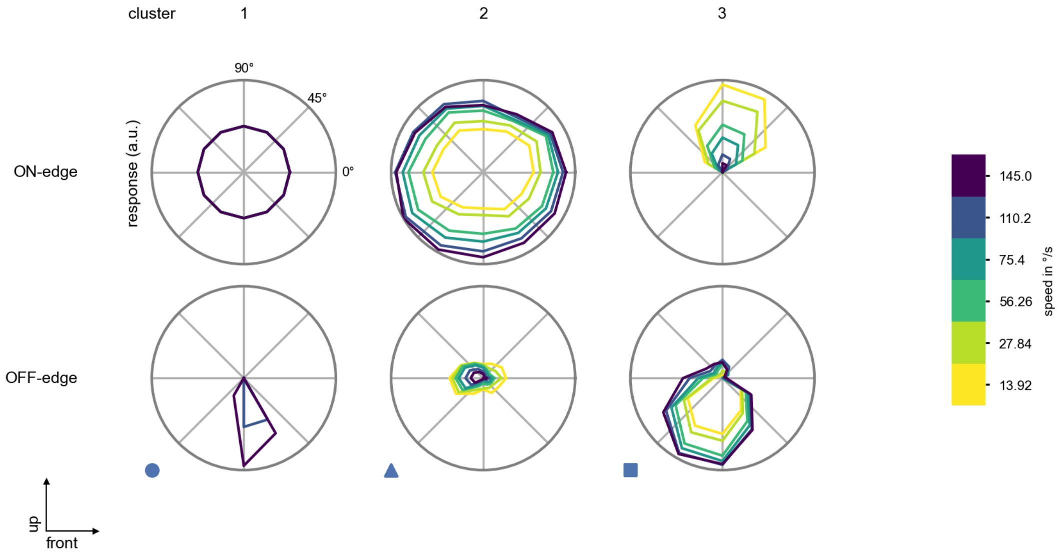

T5d - Figure 6: **Peak responses to moving edges from best-performing models.** The top row shows peak responses to moving ON-edges, the bottom row shows peak responses to moving OFF-edges. The edge-stimuli move in different directions from 0 to 360 degrees and at different speeds. Before, a grey-stimulus led the network to a stationary state. The columns show responses from the best-task-performing model in the respective cluster.

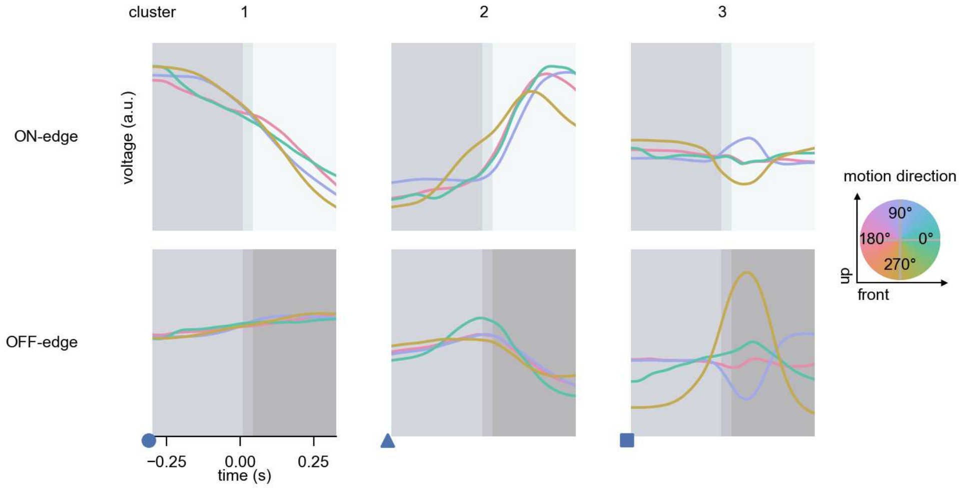

T5d - Figure 7: **Responses to moving edges from best-performing models.** The top row shows responses to moving ON-edges, the bottom row shows peak responses to moving OFF-edges. The edge-stimuli move in different directions from 0 to 360 degrees and at different speeds. Before, a grey-stimulus led the network to a stationary state. The columns show responses from the best-task-performing model in the respective cluster. Example responses to edges moving at 75.4°/s in all cardinal directions (green 0°, blue 90°, red 180°, yellow 270°) from -22.5 to 22.5° visual angle. Background contour indicates the edge, passing by the central column at second zero.

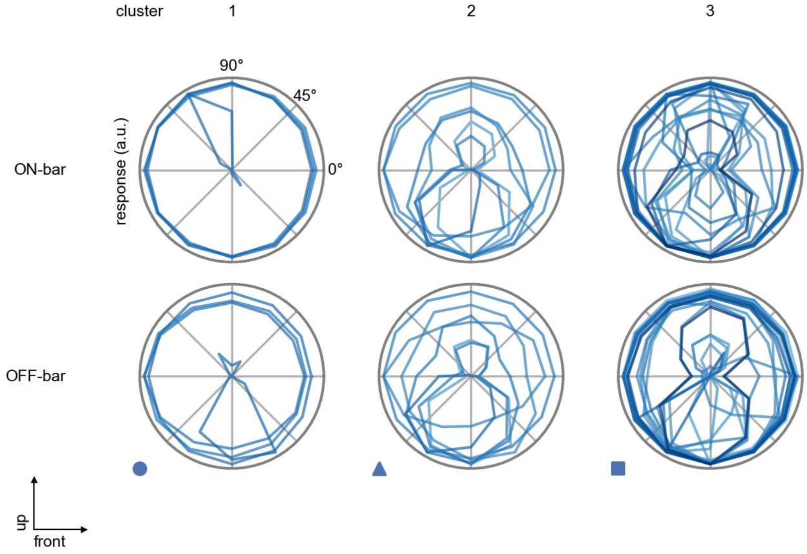

T5d - Figure 8: **Peak responses to moving bars.** The top row shows normalized peak responses to moving ON-bars, the bottom row shows normalized peak responses to moving OFF-bars. The bar-stimuli move in different directions from 0 to 360 degrees. Before, a grey-stimulus led the network to a stationary state. The responses from the different models in the different clusters (columns) overlay and are colored in the respective task error of the model. The peak responses are averaged over all assessed bar-speeds.

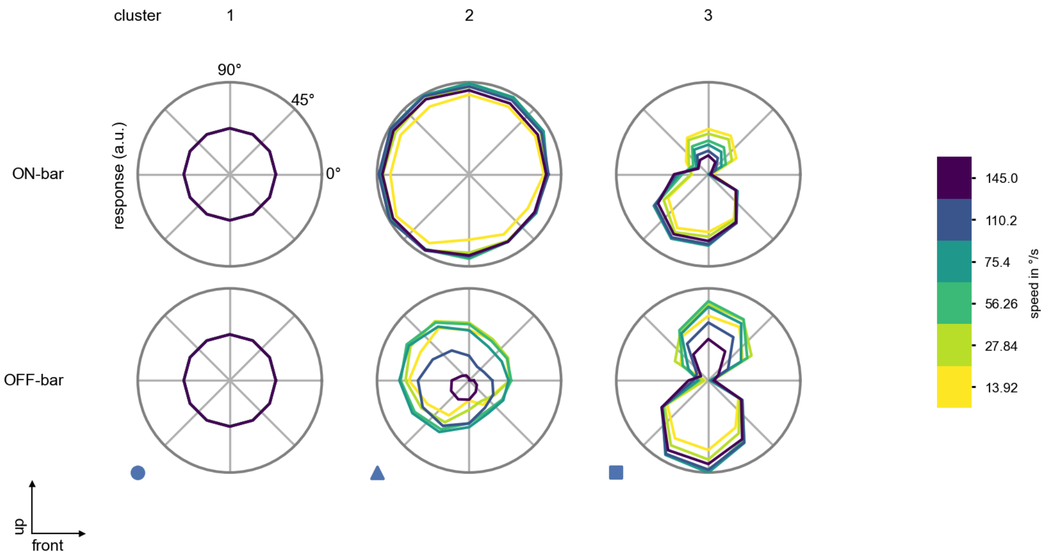

T5d - Figure 9: **Peak responses to moving bars from best-performing models.** The top row shows peak responses to moving ON-bars, the bottom row shows peak responses to moving OFF-bars. The bar-stimuli move in different directions from 0 to 360 degrees and at different speeds. Before, a grey-stimulus led the network to a stationary state. The columns show responses from the best-task-performing model in the respective cluster.

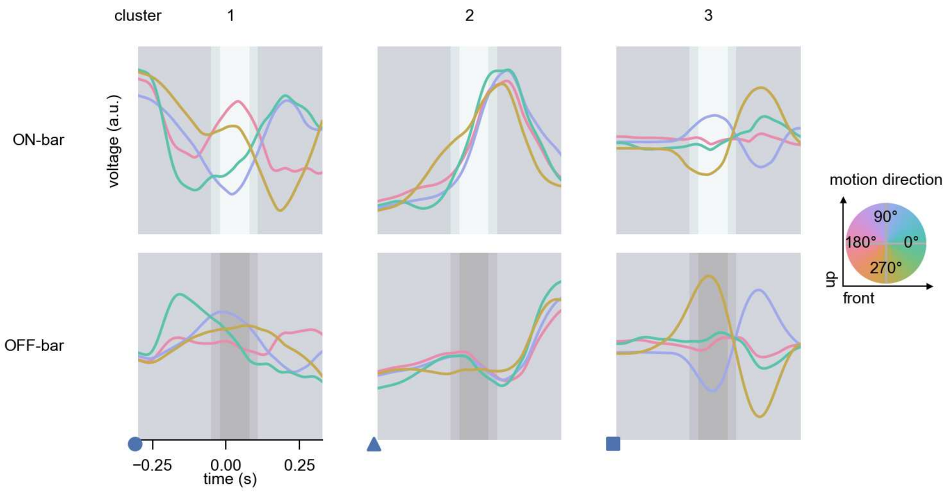

T5d - Figure 10: **Responses to moving bars from best-performing models.** The top row shows responses to moving ON-bars, the bottom row shows peak responses to moving OFF-bars. The bar-stimuli move in different directions from 0 to 360 degrees and at different speeds. Before, a grey-stimulus led the network to a stationary state. The columns show responses from the best-task-performing model in the respective cluster. Example responses to bars moving at 75.4°/s in all cardinal directions (green 0°, blue 90°, red 180°, yellow 270°) from -22.5 to 22.5° visual angle. Background contour indicates the bar, passing by the central column at second zero.

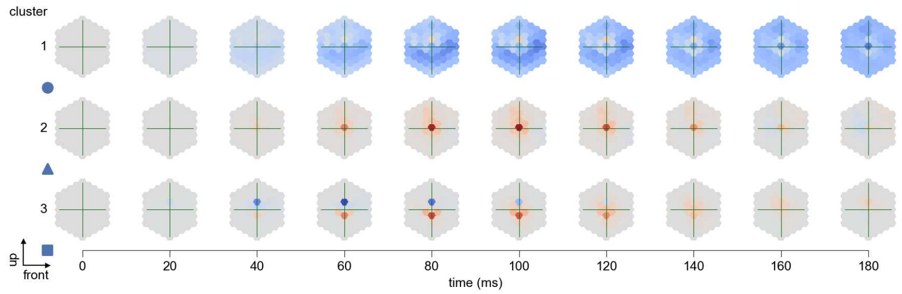

T5d - Figure 11: **Spatio-temporal receptive field.** Responses of the central cell to ON-impulses at single ommatidium impulse locations. The impulse occurs at second zero. Before, a grey-stimulus led the network to a stationary state. Responses are baseline-subtracted. Each row illustrates responses from the best-performing model of the respective cluster. Red indicated depolarization, blue indicates hyperpolarization.

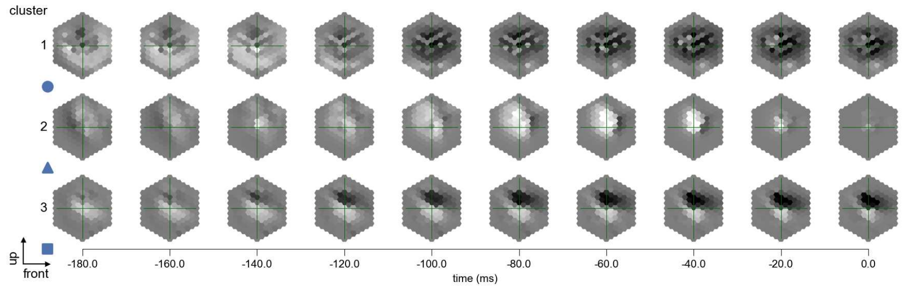

T5d - Figure 12: **Maximally excitatory stimuli.** Each row presents the regularized naturalistic-stimulus from the Sintel dataset that maximizes the cell type's central column response at second zero in the best-performing model of the respective cluster.

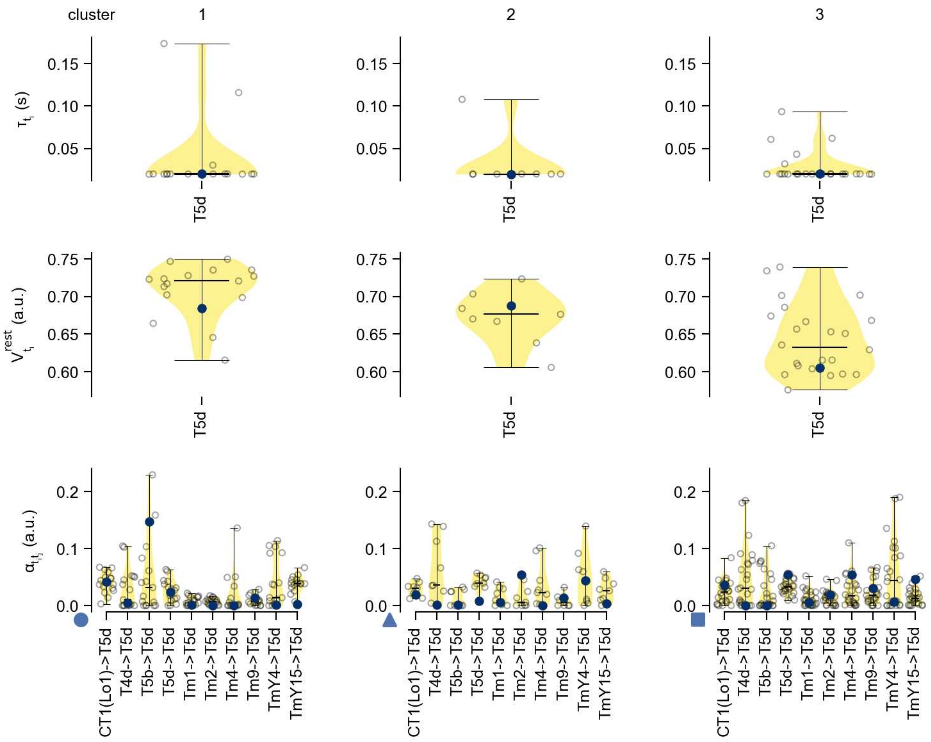

T5d - Figure 13: **Task-constrained parameters.** Each column shows the parameters inferred within the respective cluster. First row: learned time constants of the cell type. Second row: resting potentials of the cell type. Third row: scaling factors for the convolutional filters. The blue scatter represents the parameters from the best-performing model within the cluster.

# 44 Tm1

← Cell types

#### Figures

1 Anatomical receptive fields. . . . 290
2 Anatomical projective fields. . . . 291
3 Clustering of the responses to naturalistic stimuli. . . . 291
4 Responses to flashes. . . . 292
5 Peak responses to moving edges. . . . 292
6 Peak responses to moving edges from best-performing models. . . . 293
7 Responses to moving edges from best-performing models. . . . 293
8 Peak responses to moving bars. . . . 294
9 Peak responses to moving bars from best-performing models. . . . 294
10 Responses to moving bars from best-performing models. . . . 295
11 Spatio-temporal receptive field. . . . 295
12 Maximally excitatory stimuli. . . . 295
13 Task-constrained parameters. . . . 296

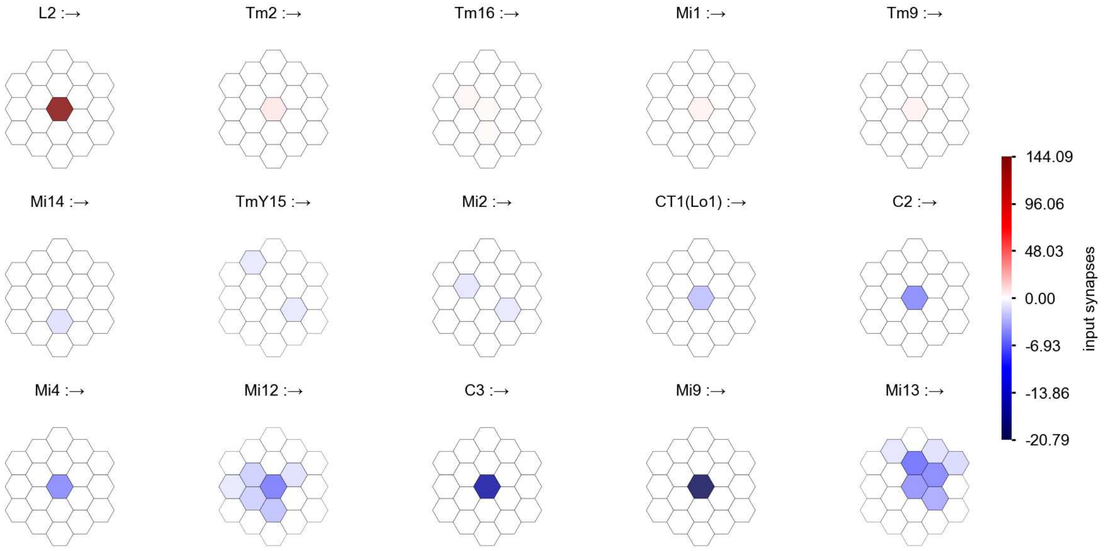

Tm1 - Figure 1: **Anatomical receptive fields.** Each colored hexagon is a projection, with the projection strength characterized by the average number of synapses that we count from the EM reconstruction. Red indicates excitatory synapses, blue indicates inhibitory synapses from inferred signs. Filters in the order of their total number of synapses.

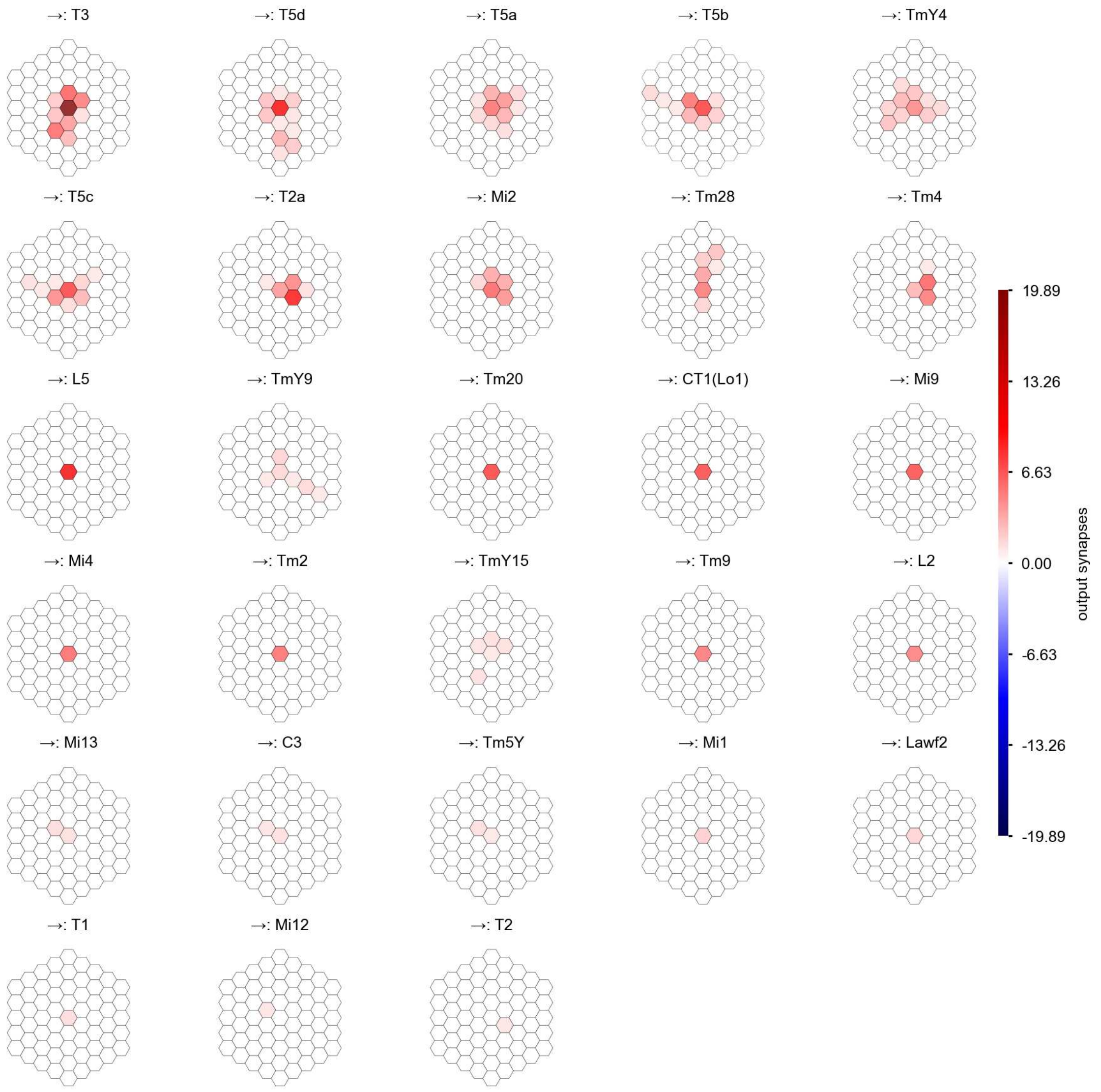

Tm1 - Figure 2: **Anatomical projective fields.** Each colored hexagon is a projection, with the projection strength characterized by the average number of synapses that we count from the EM reconstruction. Red indicates excitatory synapses, blue indicates inhibitory synapses from inferred signs. Filters in the order of their total number of synapses.

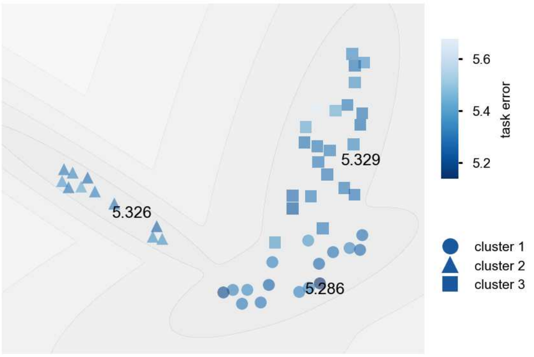

Tm1 - Figure 3: **Clustering of the responses to naturalistic stimuli.** From responses to naturalistic scenes from Sintel. Scatterpoints represent individual models colored by their task error.

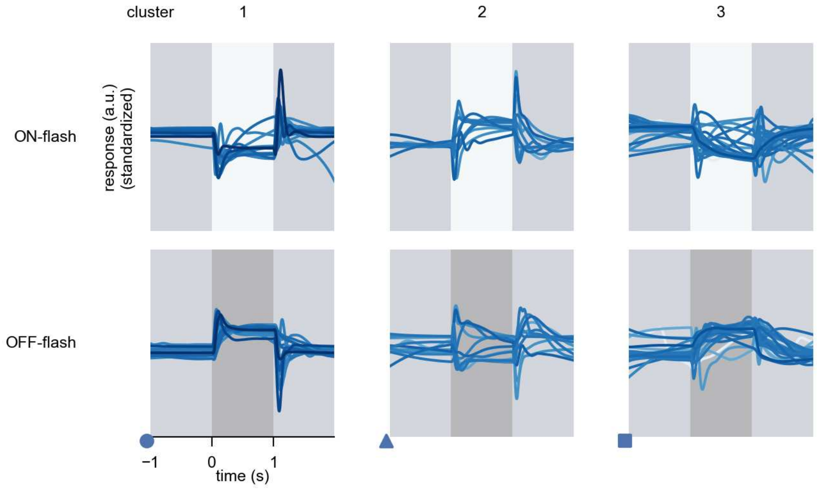

Tm1 - Figure 4: **Responses to flashes.** The top row shows responses to ON-flashes, the bottom row shows responses to OFF-flashes. The responses from the different models in the different clusters (columns) overlay and are colored in the respective task error of the model. The circular flashes occur at second zero, cover 6 ommatidia in radius, and last for one second. Before and after, a grey-stimulus leads to a stationary state of the network. The traces are standardized.

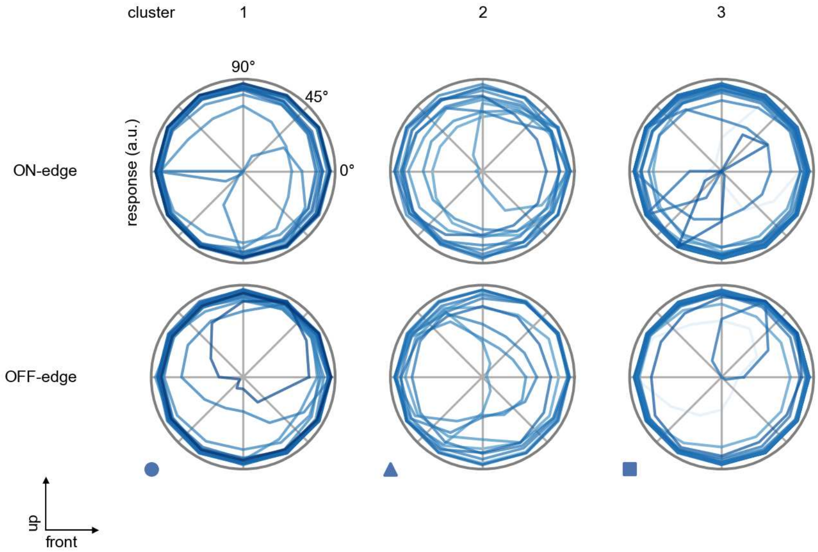

Tm1 - Figure 5: **Peak responses to moving edges.** The top row shows normalized peak responses to moving ON-edges, the bottom row shows normalized peak responses to moving OFF-edges. The edge-stimuli move in different directions from 0 to 360 degrees. Before, a grey-stimulus led the network to a stationary state. The responses from the different models in the different clusters (columns) overlay and are colored in the respective task error of the model. The peak responses are averaged over all assessed edge-speeds.

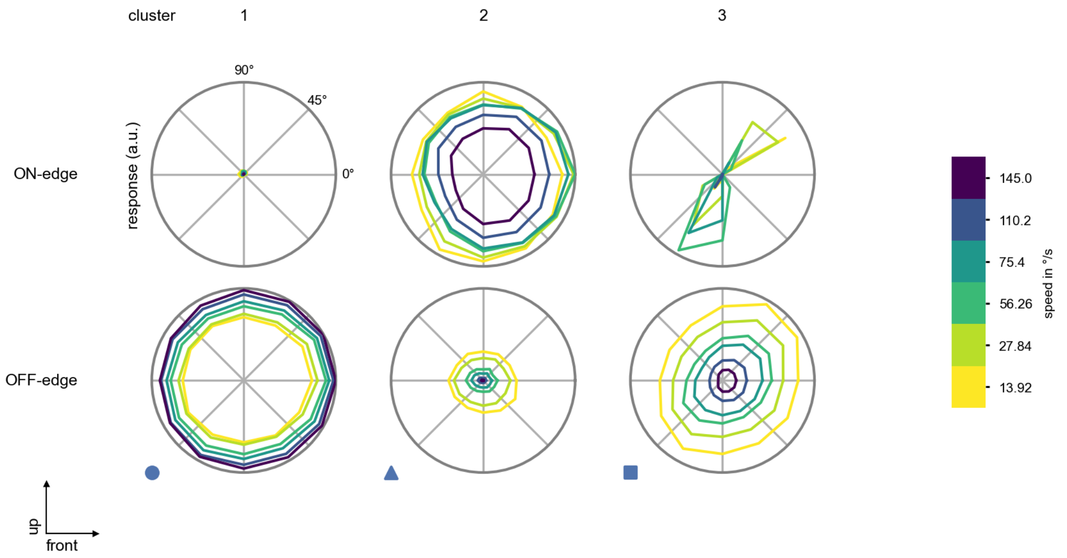

Tm1 - Figure 6: **Peak responses to moving edges from best-performing models.** The top row shows peak responses to moving ON-edges, the bottom row shows peak responses to moving OFF-edges. The edge-stimuli move in different directions from 0 to 360 degrees and at different speeds. Before, a grey-stimulus led the network to a stationary state. The columns show responses from the best-task-performing model in the respective cluster.

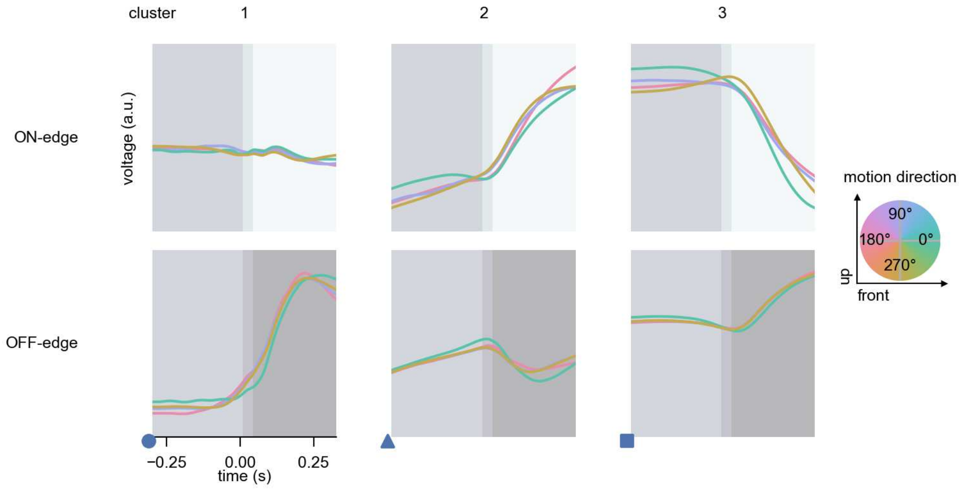

Tm1 - Figure 7: **Responses to moving edges from best-performing models.** The top row shows responses to moving ON-edges, the bottom row shows peak responses to moving OFF-edges. The edge-stimuli move in different directions from 0 to 360 degrees and at different speeds. Before, a grey-stimulus led the network to a stationary state. The columns show responses from the best-task-performing model in the respective cluster. Example responses to edges moving at 75.4°/s in all cardinal directions (green 0°, blue 90°, red 180°, yellow 270°) from -22.5 to 22.5° visual angle. Background contour indicates the edge, passing by the central column at second zero.

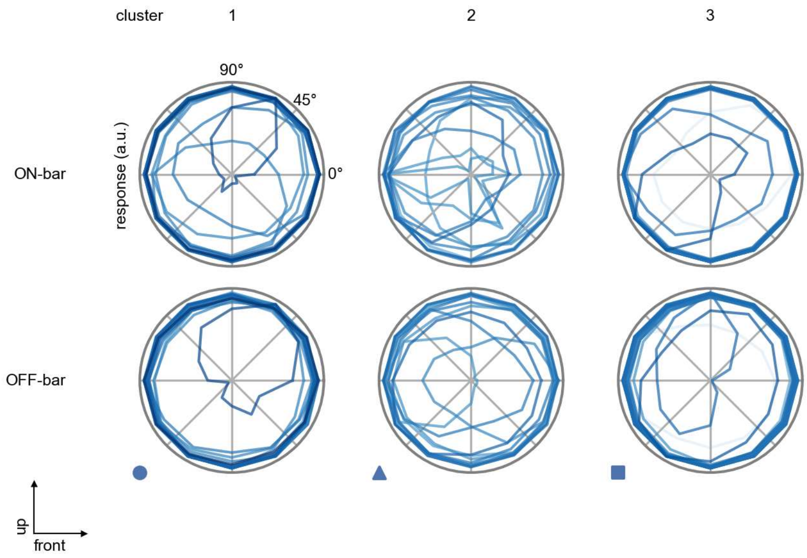

Tm1 - Figure 8: **Peak responses to moving bars.** The top row shows normalized peak responses to moving ON-bars, the bottom row shows normalized peak responses to moving OFF-bars. The bar-stimuli move in different directions from 0 to 360 degrees. Before, a grey-stimulus led the network to a stationary state. The responses from the different models in the different clusters (columns) overlay and are colored in the respective task error of the model. The peak responses are averaged over all assessed bar-speeds.

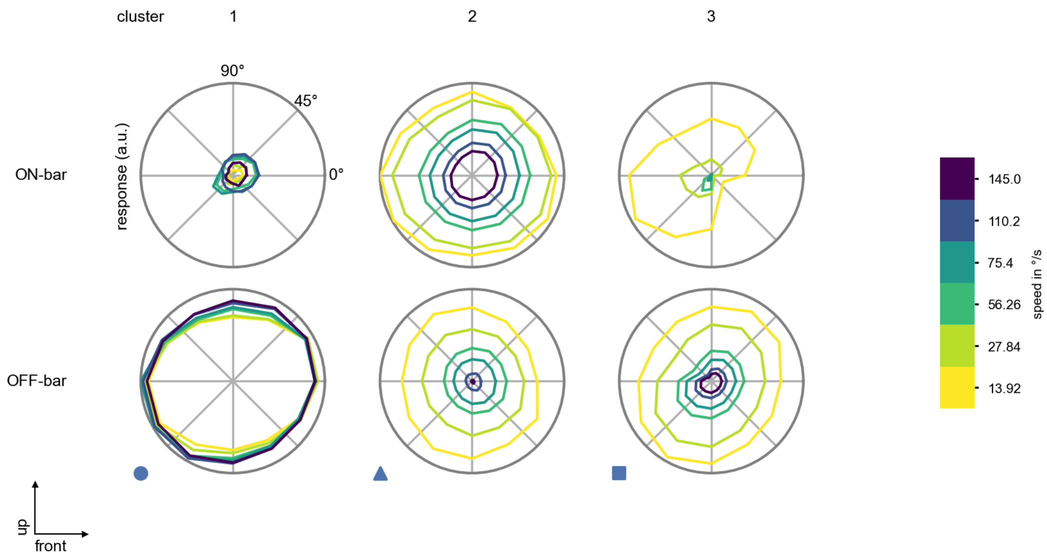

Tm1 - Figure 9: **Peak responses to moving bars from best-performing models.** The top row shows peak responses to moving ON-bars, the bottom row shows peak responses to moving OFF-bars. The bar-stimuli move in different directions from 0 to 360 degrees and at different speeds. Before, a grey-stimulus led the network to a stationary state. The columns show responses from the best-task-performing model in the respective cluster.

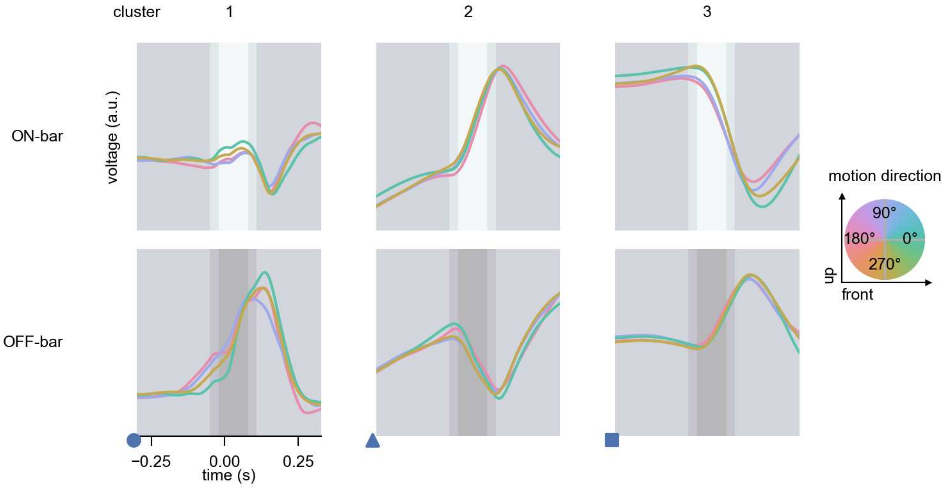

Tm1 - Figure 10: **Responses to moving bars from best-performing models.** The top row shows responses to moving ON-bars, the bottom row shows peak responses to moving OFF-bars. The bar-stimuli move in different directions from 0 to 360 degrees and at different speeds. Before, a grey-stimulus led the network to a stationary state. The columns show responses from the best-task-performing model in the respective cluster. Example responses to bars moving at 75.4°/s in all cardinal directions (green 0°, blue 90°, red 180°, yellow 270°) from -22.5 to 22.5° visual angle. Background contour indicates the bar, passing by the central column at second zero.

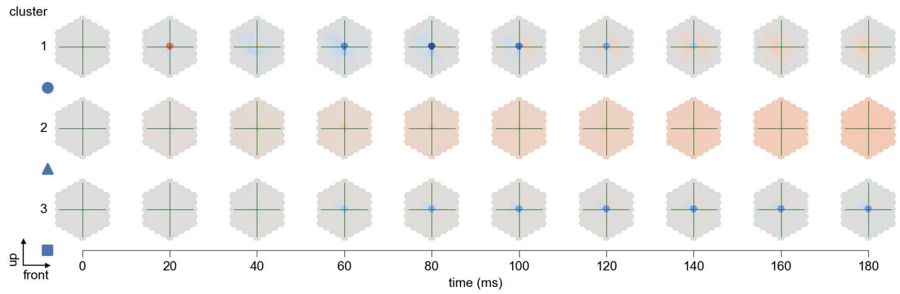

Tm1 - Figure 11: **Spatio-temporal receptive field.** Responses of the central cell to ON-impulses at single ommatidium impulse locations. The impulse occurs at second zero. Before, a grey-stimulus led the network to a stationary state. Responses are baseline-subtracted. Each row illustrates responses from the best-performing model of the respective cluster. Red indicated depolarization, blue indicates hyperpolarization.

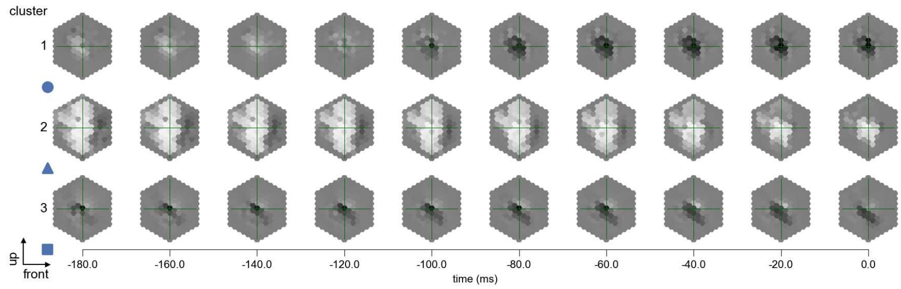

Tm1 - Figure 12: **Maximally excitatory stimuli.** Each row presents the regularized naturalistic-stimulus from the Sintel dataset that maximizes the cell type's central column response at second zero in the best-performing model of the respective cluster.

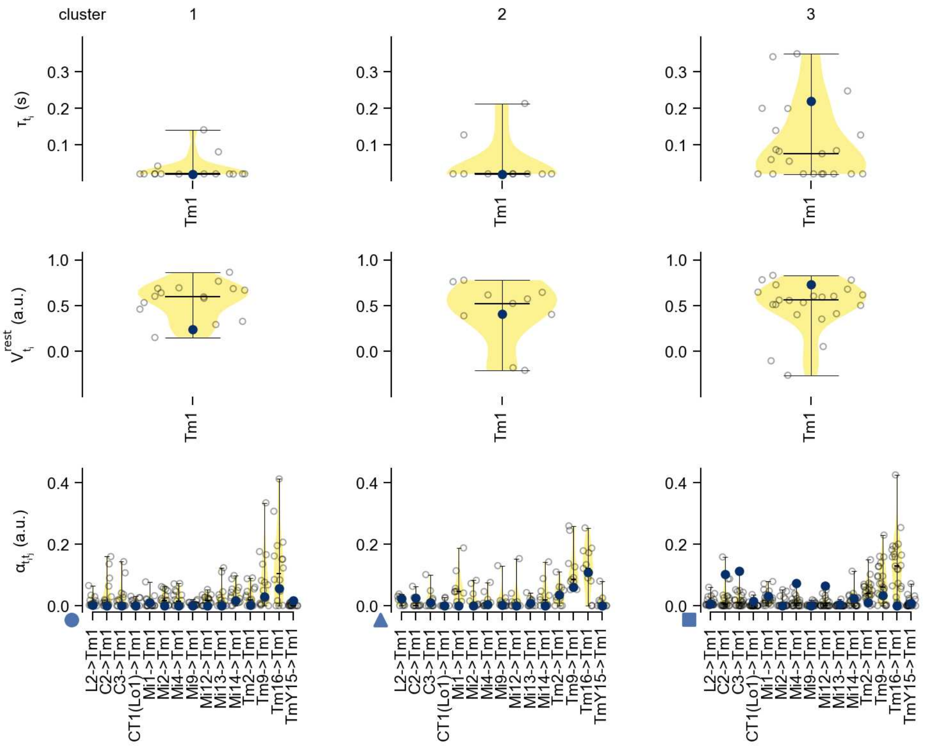

Tm1 - Figure 13: **Task-constrained parameters.** Each column shows the parameters inferred within the respective cluster. First row: learned time constants of the cell type. Second row: resting potentials of the cell type. Third row: scaling factors for the convolutional filters. The blue scatter represents the parameters from the best-performing model within the cluster.

# 45 Tm2

← Cell types

#### Figures

1 Anatomical receptive fields. . . . 297
2 Anatomical projective fields. . . . 298
3 Clustering of the responses to naturalistic stimuli. . . . 298
4 Responses to flashes. . . . 299
5 Peak responses to moving edges. . . . 299
6 Peak responses to moving edges from best-performing models. . . . 300
7 Responses to moving edges from best-performing models. . . . 300
8 Peak responses to moving bars. . . . 301
9 Peak responses to moving bars from best-performing models. . . . 301
10 Responses to moving bars from best-performing models. . . . 302
11 Spatio-temporal receptive field. . . . 302
12 Maximally excitatory stimuli. . . . 303
13 Task-constrained parameters. . . . 303

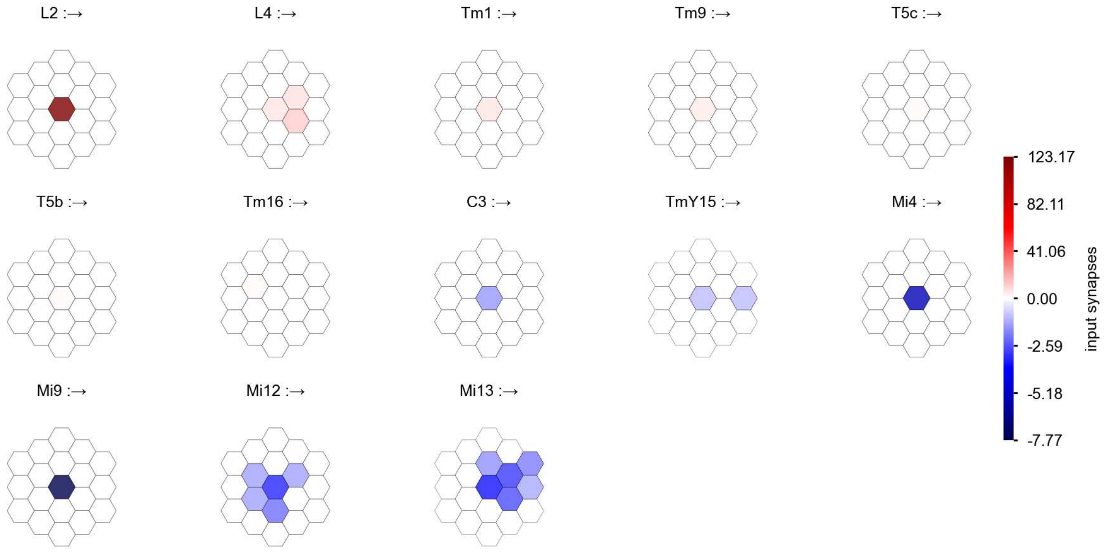

Tm2 - Figure 1: **Anatomical receptive fields.** Each colored hexagon is a projection, with the projection strength characterized by the average number of synapses that we count from the EM reconstruction. Red indicates excitatory synapses, blue indicates inhibitory synapses from inferred signs. Filters in the order of their total number of synapses.

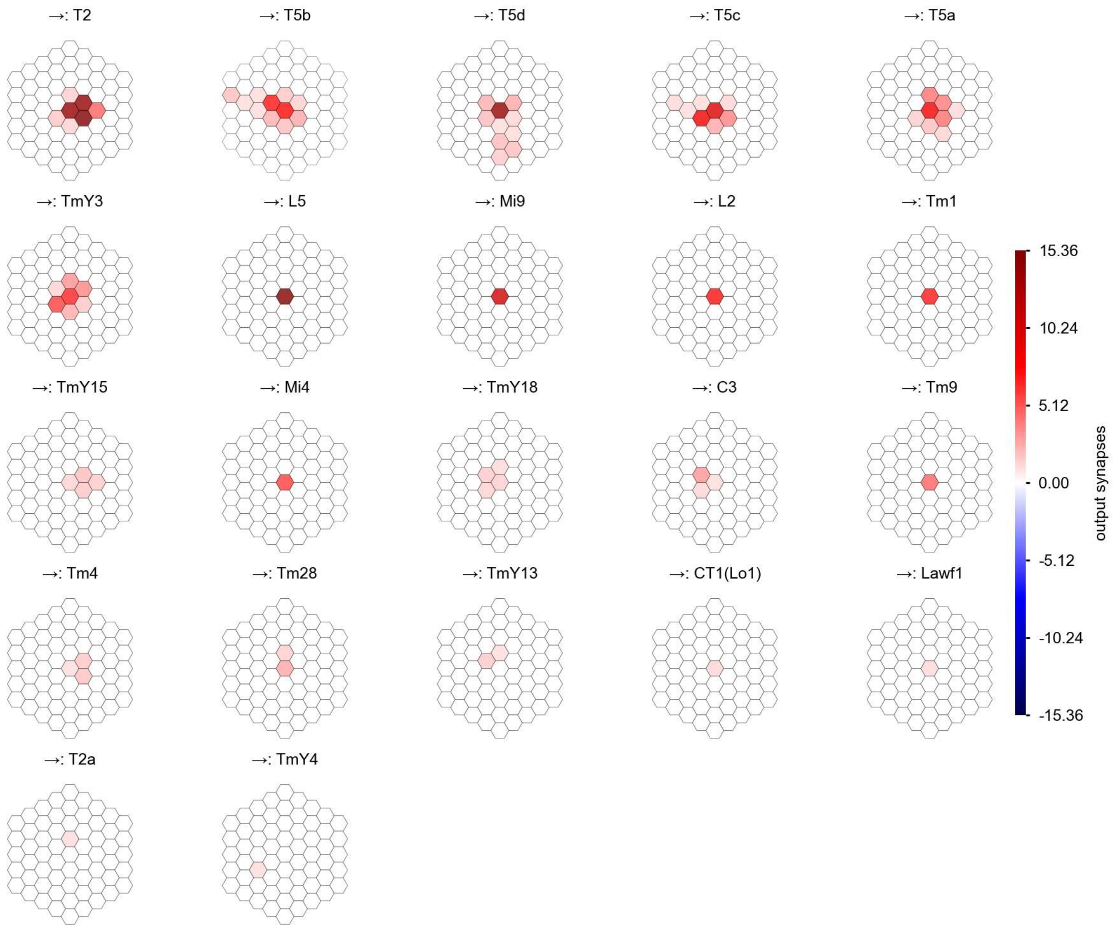

Tm2 - Figure 2: **Anatomical projective fields.** Each colored hexagon is a projection, with the projection strength characterized by the average number of synapses that we count from the EM reconstruction. Red indicates excitatory synapses, blue indicates inhibitory synapses from inferred signs. Filters in the order of their total number of synapses.

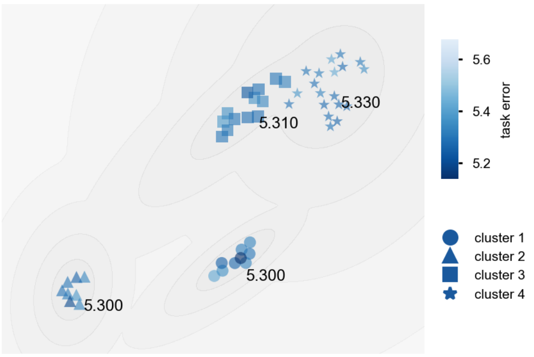

Tm2 - Figure 3: **Clustering of the responses to naturalistic stimuli.** From responses to naturalistic scenes from Sintel. Scatterpoints represent individual models colored by their task error.

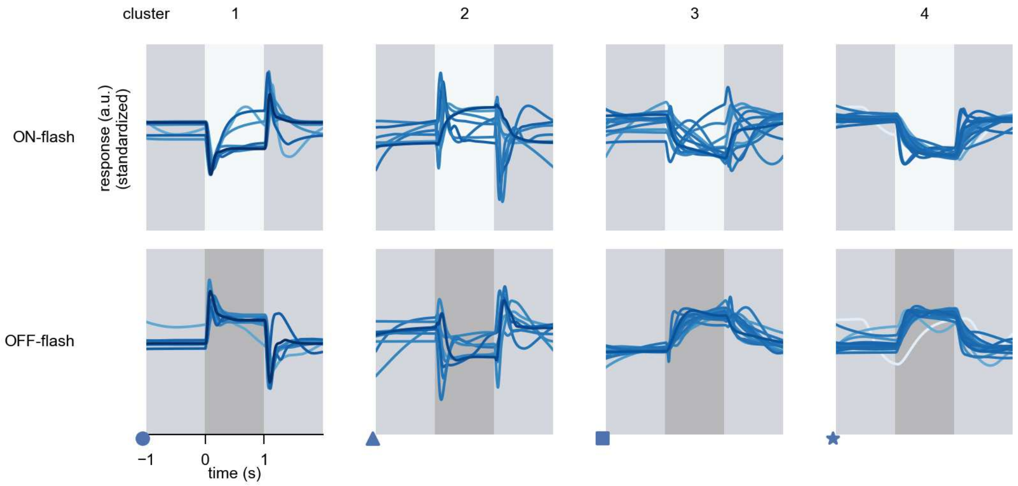

Tm2 - Figure 4: **Responses to flashes.** The top row shows responses to ON-flashes, the bottom row shows responses to OFF-flashes. The responses from the different models in the different clusters (columns) overlay and are colored in the respective task error of the model. The circular flashes occur at second zero, cover 6 ommatidia in radius, and last for one second. Before and after, a grey-stimulus leads to a stationary state of the network. The traces are standardized.

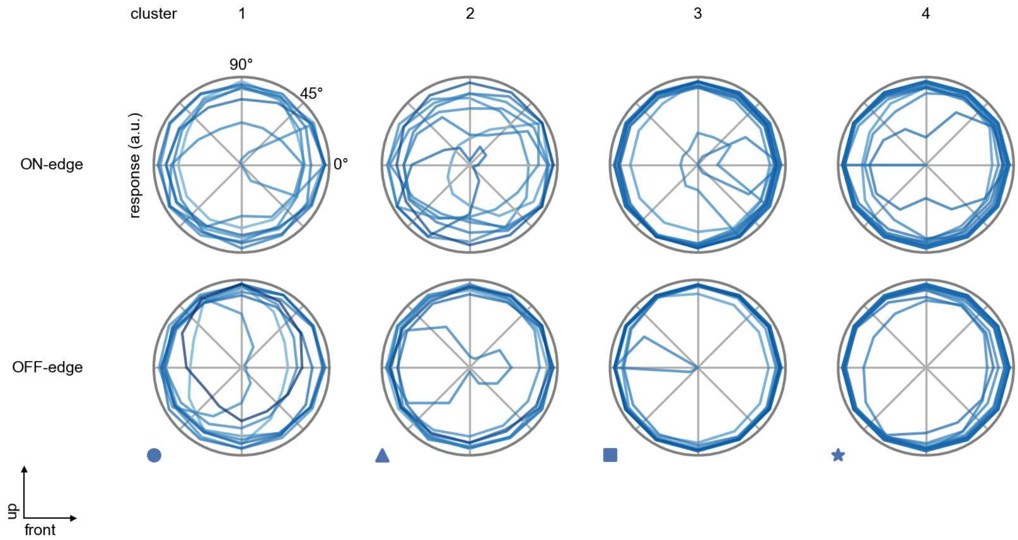

Tm2 - Figure 5: **Peak responses to moving edges.** The top row shows normalized peak responses to moving ON-edges, the bottom row shows normalized peak responses to moving OFF-edges. The edge-stimuli move in different directions from 0 to 360 degrees. Before, a grey-stimulus led the network to a stationary state. The responses from the different models in the different clusters (columns) overlay and are colored in the respective task error of the model. The peak responses are averaged over all assessed edge-speeds.

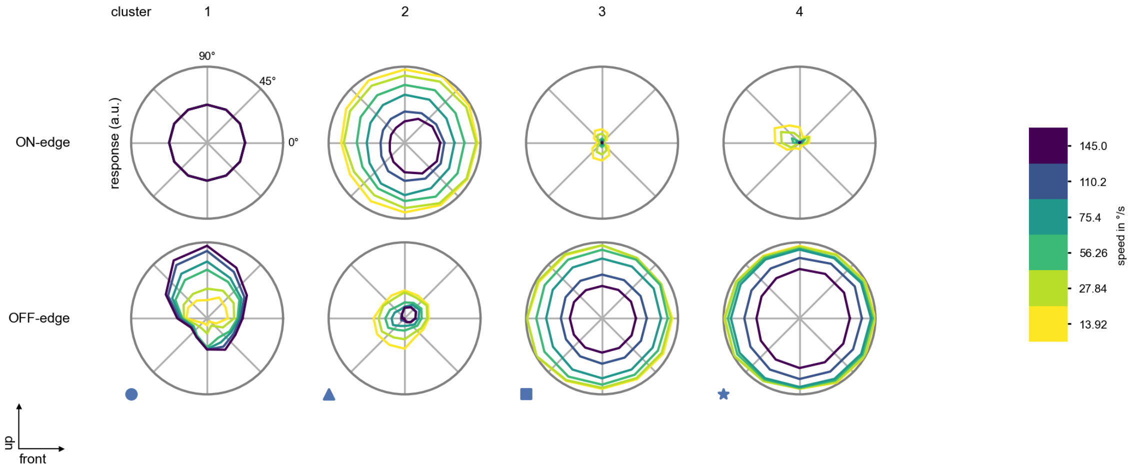

Tm2 - Figure 6: **Peak responses to moving edges from best-performing models.** The top row shows peak responses to moving ON-edges, the bottom row shows peak responses to moving OFF-edges. The edge-stimuli move in different directions from 0 to 360 degrees and at different speeds. Before, a grey-stimulus led the network to a stationary state. The columns show responses from the best-task-performing model in the respective cluster.

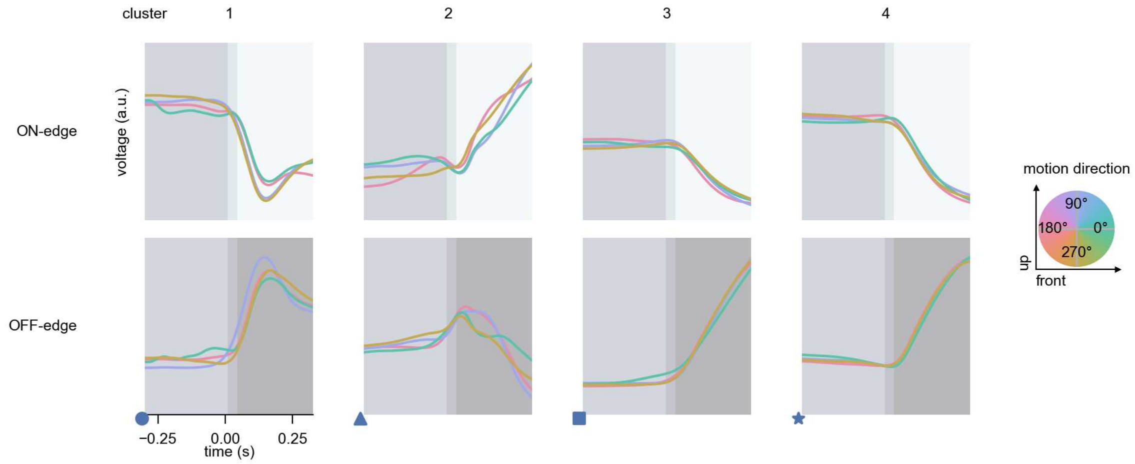

Tm2 - Figure 7: **Responses to moving edges from best-performing models.** The top row shows responses to moving ON-edges, the bottom row shows peak responses to moving OFF-edges. The edge-stimuli move in different directions from 0 to 360 degrees and at different speeds. Before, a grey-stimulus led the network to a stationary state. The columns show responses from the best-task-performing model in the respective cluster. Example responses to edges moving at 75.4°/s in all cardinal directions (green 0°, blue 90°, red 180°, yellow 270°) from -22.5 to 22.5° visual angle. Background contour indicates the edge, passing by the central column at second zero.

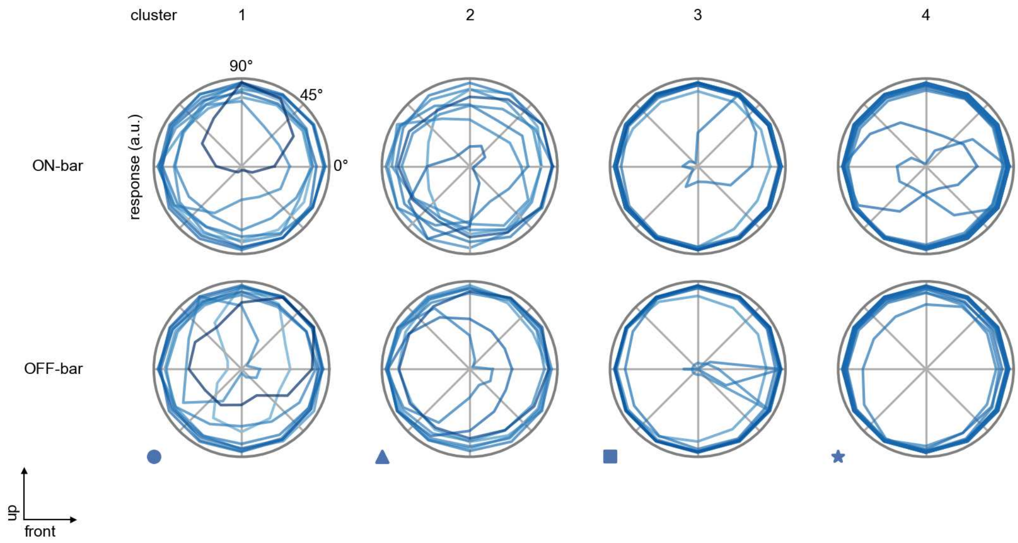

Tm2 - Figure 8: **Peak responses to moving bars.** The top row shows normalized peak responses to moving ON-bars, the bottom row shows normalized peak responses to moving OFF-bars. The bar-stimuli move in different directions from 0 to 360 degrees. Before, a grey-stimulus led the network to a stationary state. The responses from the different models in the different clusters (columns) overlay and are colored in the respective task error of the model. The peak responses are averaged over all assessed bar-speeds.

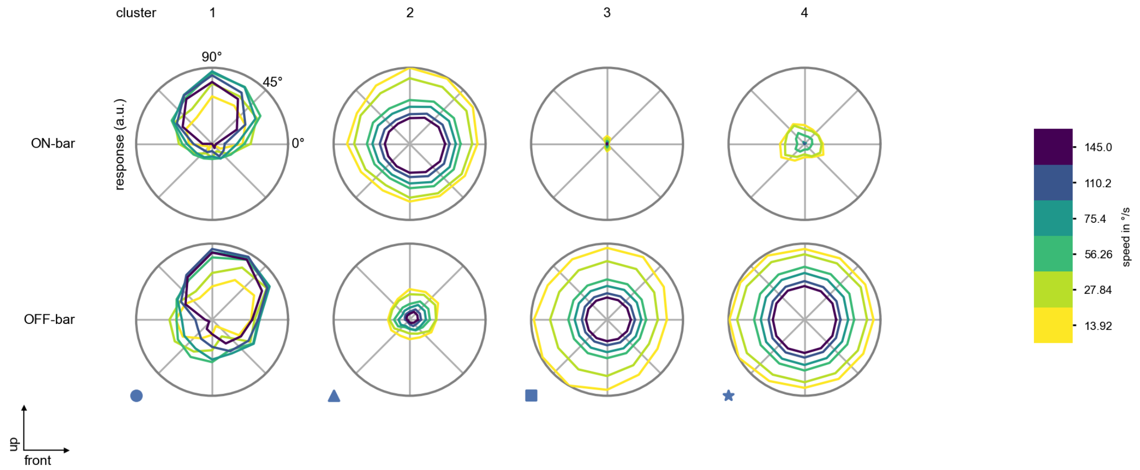

Tm2 - Figure 9: **Peak responses to moving bars from best-performing models.** The top row shows peak responses to moving ON-bars, the bottom row shows peak responses to moving OFF-bars. The bar-stimuli move in different directions from 0 to 360 degrees and at different speeds. Before, a grey-stimulus led the network to a stationary state. The columns show responses from the best-task-performing model in the respective cluster.

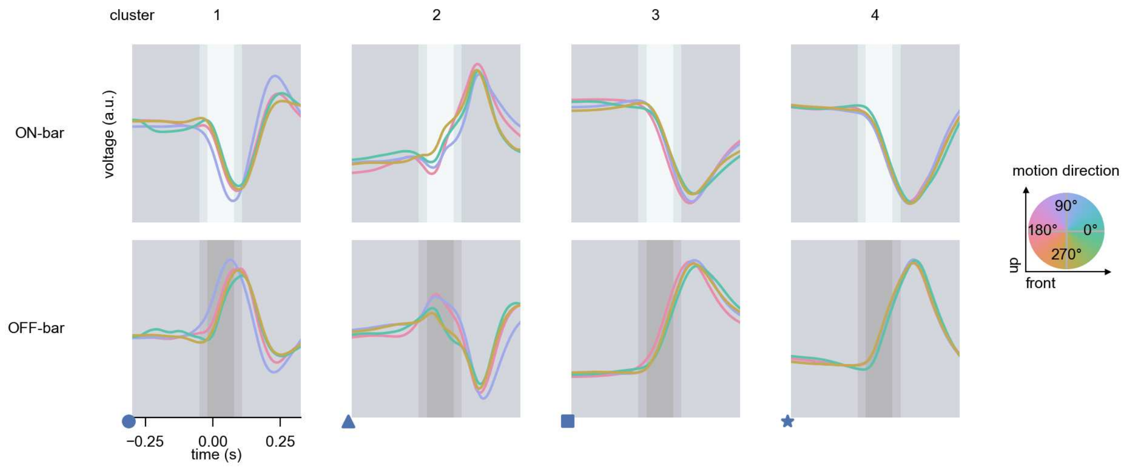

Tm2 - Figure 10: **Responses to moving bars from best-performing models.** The top row shows responses to moving ON-bars, the bottom row shows peak responses to moving OFF-bars. The bar-stimuli move in different directions from 0 to 360 degrees and at different speeds. Before, a grey-stimulus led the network to a stationary state. The columns show responses from the best-task-performing model in the respective cluster. Example responses to bars moving at 75.4°/s in all cardinal directions (green 0°, blue 90°, red 180°, yellow 270°) from -22.5 to 22.5° visual angle. Background contour indicates the bar, passing by the central column at second zero.

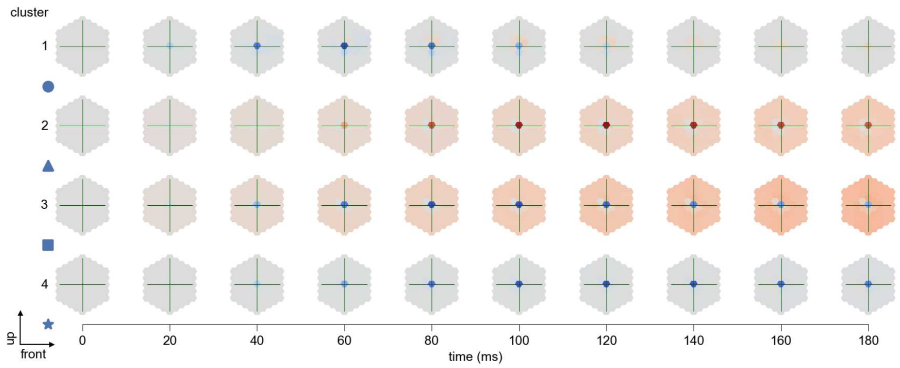

Tm2 - Figure 11: **Spatio-temporal receptive field.** Responses of the central cell to ON-impulses at single ommatidium impulse locations. The impulse occurs at second zero. Before, a grey-stimulus led the network to a stationary state. Responses are baseline-subtracted. Each row illustrates responses from the best-performing model of the respective cluster. Red indicated depolarization, blue indicates hyperpolarization.

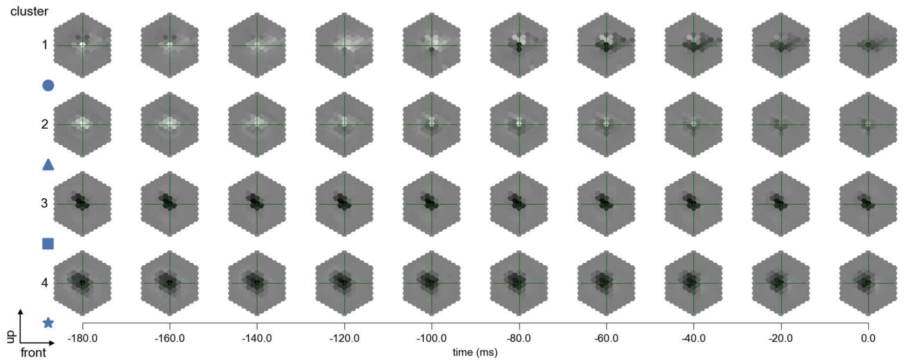

Tm2 - Figure 12: **Maximally excitatory stimuli.** Each row presents the regularized naturalistic-stimulus from the Sintel dataset that maximizes the cell type's central column response at second zero in the best-performing model of the respective cluster.

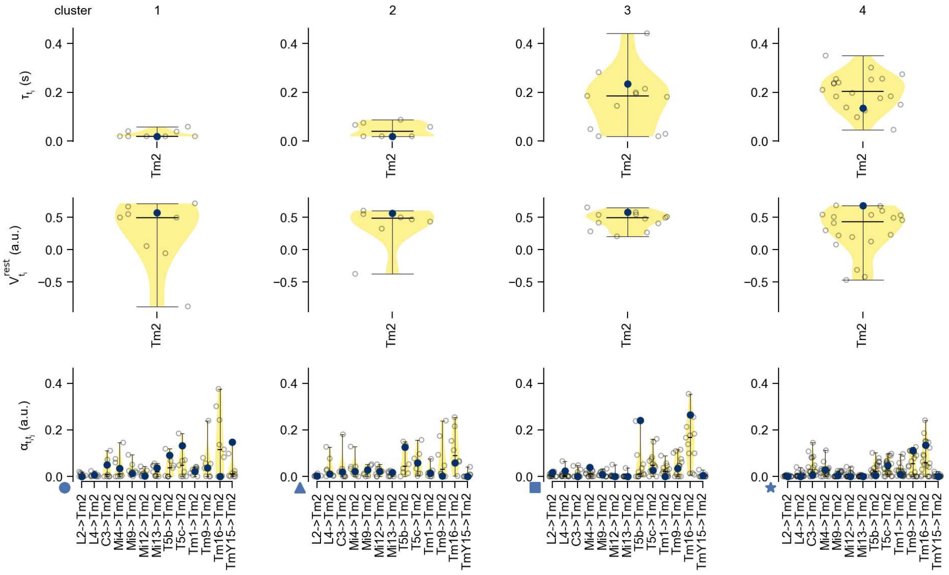

Tm2 - Figure 13: **Task-constrained parameters.** Each column shows the parameters inferred within the respective cluster. First row: learned time constants of the cell type. Second row: resting potentials of the cell type. Third row: scaling factors for the convolutional filters. The blue scatter represents the parameters from the best-performing model within the cluster.

# 46 Tm3

← Cell types

#### Figures

1 Anatomical receptive fields. . . . . 304
2 Anatomical projective fields. . . . . 305
3 Clustering of the responses to naturalistic stimuli. . . . . 305
4 Responses to flashes. . . . . 306
5 Peak responses to moving edges. . . . . 306
6 Peak responses to moving edges from best-performing models. . . . . 307
7 Responses to moving edges from best-performing models. . . . . 307
8 Peak responses to moving bars. . . . . 308
9 Peak responses to moving bars from best-performing models. . . . . 308
10 Responses to moving bars from best-performing models. . . . . 309
11 Spatio-temporal receptive field. . . . . 309
12 Maximally excitatory stimuli. . . . . 310
13 Task-constrained parameters. . . . . 310

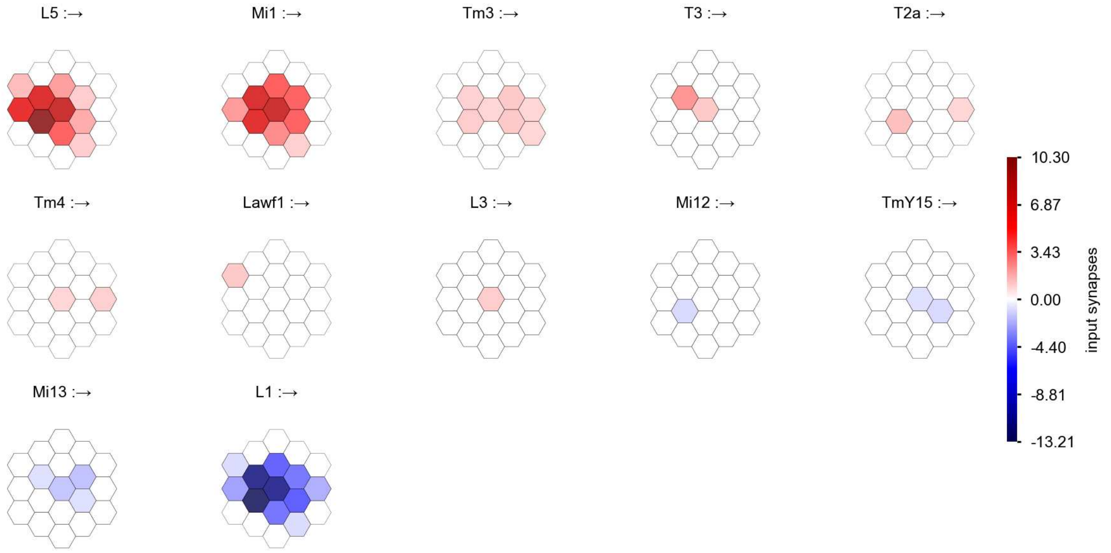

Tm3 - Figure 1: **Anatomical receptive fields.** Each colored hexagon is a projection, with the projection strength characterized by the average number of synapses that we count from the EM reconstruction. Red indicates excitatory synapses, blue indicates inhibitory synapses from inferred signs. Filters in the order of their total number of synapses.

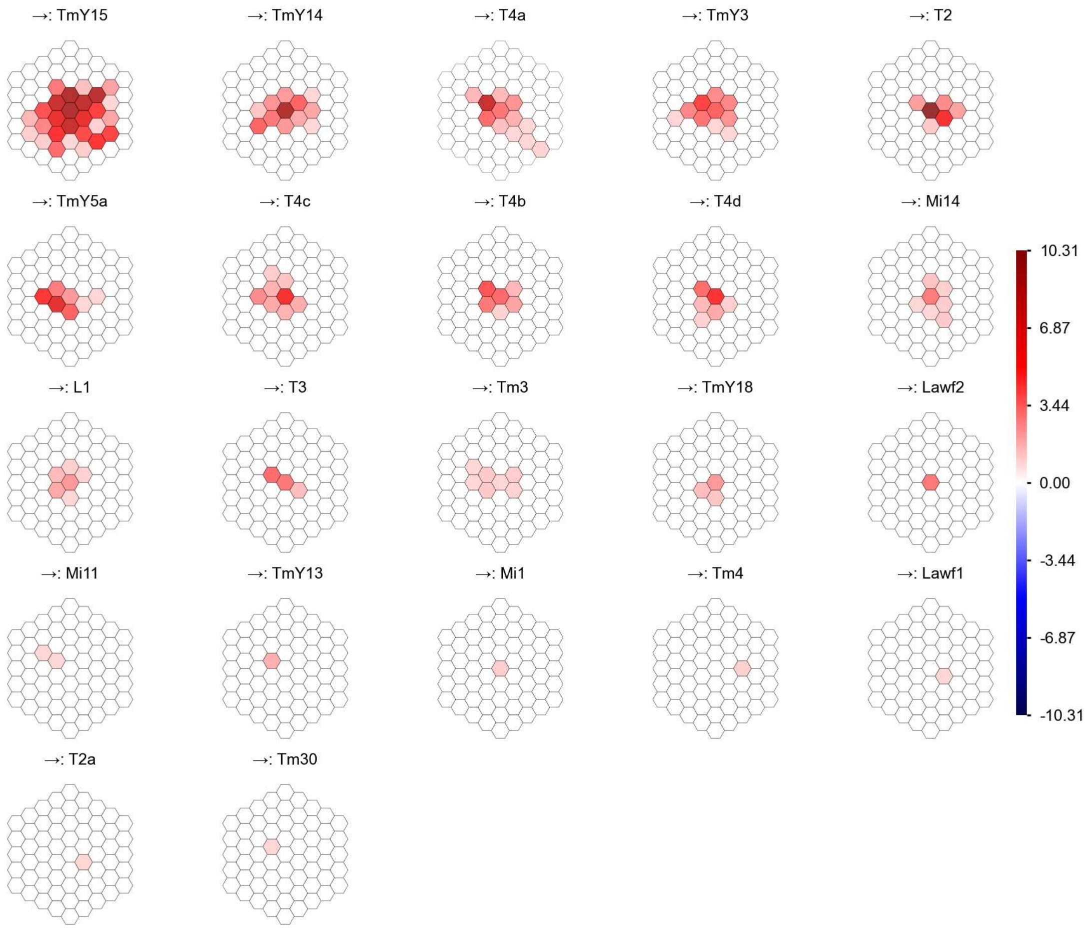

Tm3 - Figure 2: **Anatomical projective fields.** Each colored hexagon is a projection, with the projection strength characterized by the average number of synapses that we count from the EM reconstruction. Red indicates excitatory synapses, blue indicates inhibitory synapses from inferred signs. Filters in the order of their total number of synapses.

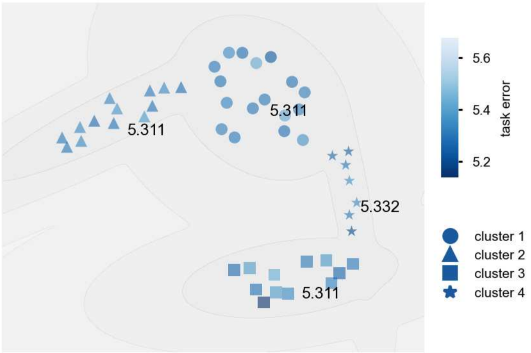

Tm3 - Figure 3: **Clustering of the responses to naturalistic stimuli.** From responses to naturalistic scenes from Sintel. Scatterpoints represent individual models colored by their task error.

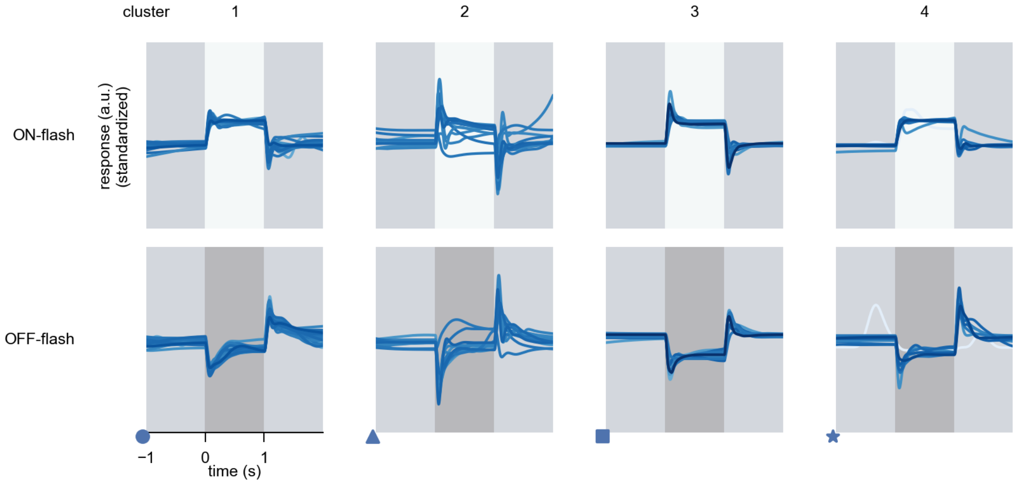

Tm3 - Figure 4: **Responses to flashes.** The top row shows responses to ON-flashes, the bottom row shows responses to OFF-flashes. The responses from the different models in the different clusters (columns) overlay and are colored in the respective task error of the model. The circular flashes occur at second zero, cover 6 ommatidia in radius, and last for one second. Before and after, a grey-stimulus leads to a stationary state of the network. The traces are standardized.

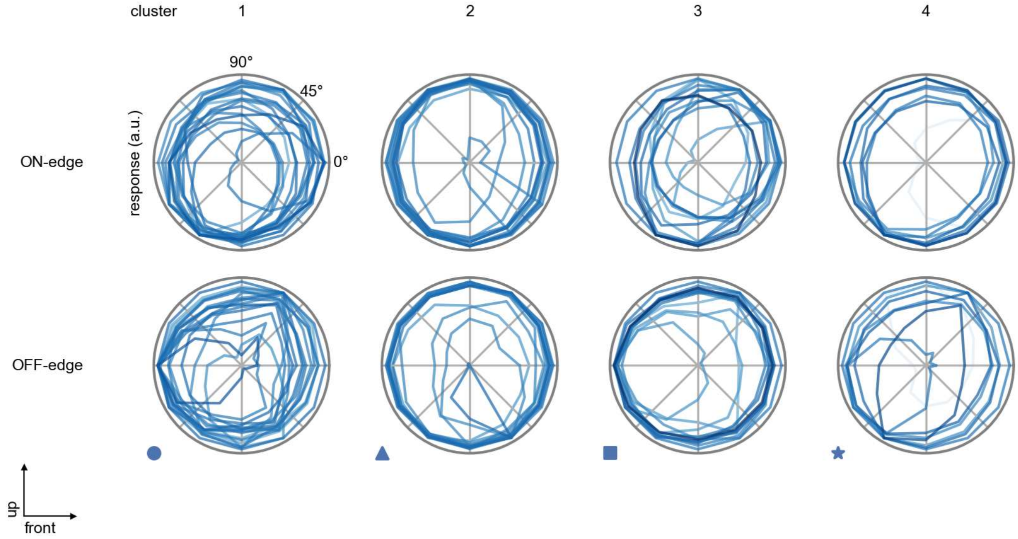

Tm3 - Figure 5: **Peak responses to moving edges.** The top row shows normalized peak responses to moving ON-edges, the bottom row shows normalized peak responses to moving OFF-edges. The edge-stimuli move in different directions from 0 to 360 degrees. Before, a grey-stimulus led the network to a stationary state. The responses from the different models in the different clusters (columns) overlay and are colored in the respective task error of the model. The peak responses are averaged over all assessed edge-speeds.

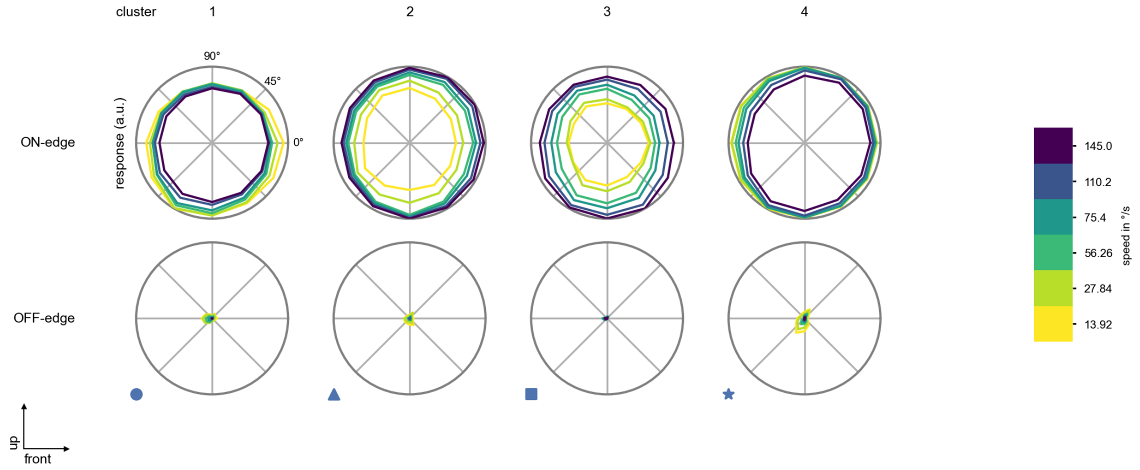

Tm3 - Figure 6: **Peak responses to moving edges from best-performing models.** The top row shows peak responses to moving ON-edges, the bottom row shows peak responses to moving OFF-edges. The edge-stimuli move in different directions from 0 to 360 degrees and at different speeds. Before, a grey-stimulus led the network to a stationary state. The columns show responses from the best-task-performing model in the respective cluster.

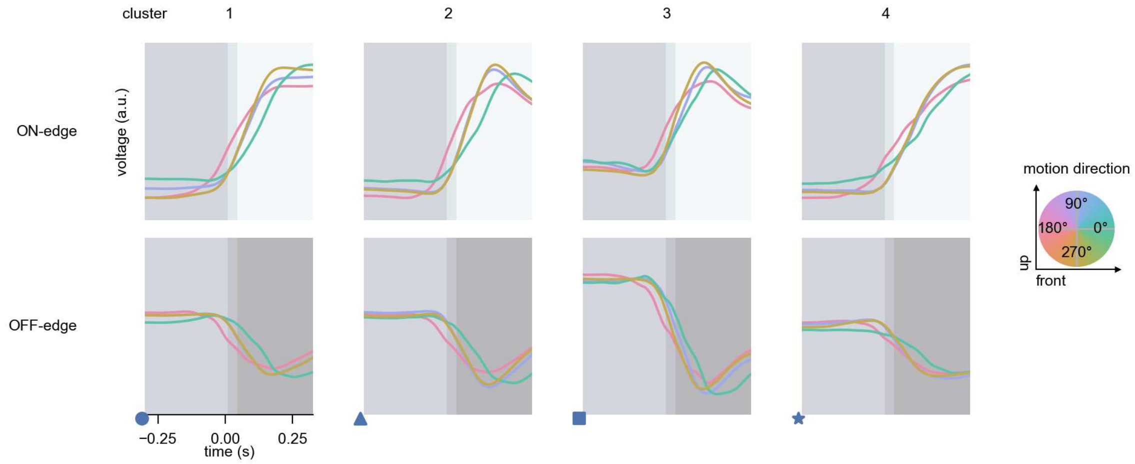

Tm3 - Figure 7: **Responses to moving edges from best-performing models.** The top row shows responses to moving ON-edges, the bottom row shows peak responses to moving OFF-edges. The edge-stimuli move in different directions from 0 to 360 degrees and at different speeds. Before, a grey-stimulus led the network to a stationary state. The columns show responses from the best-task-performing model in the respective cluster. Example responses to edges moving at 75.4°/s in all cardinal directions (green 0°, blue 90°, red 180°, yellow 270°) from -22.5 to 22.5° visual angle. Background contour indicates the edge, passing by the central column at second zero.

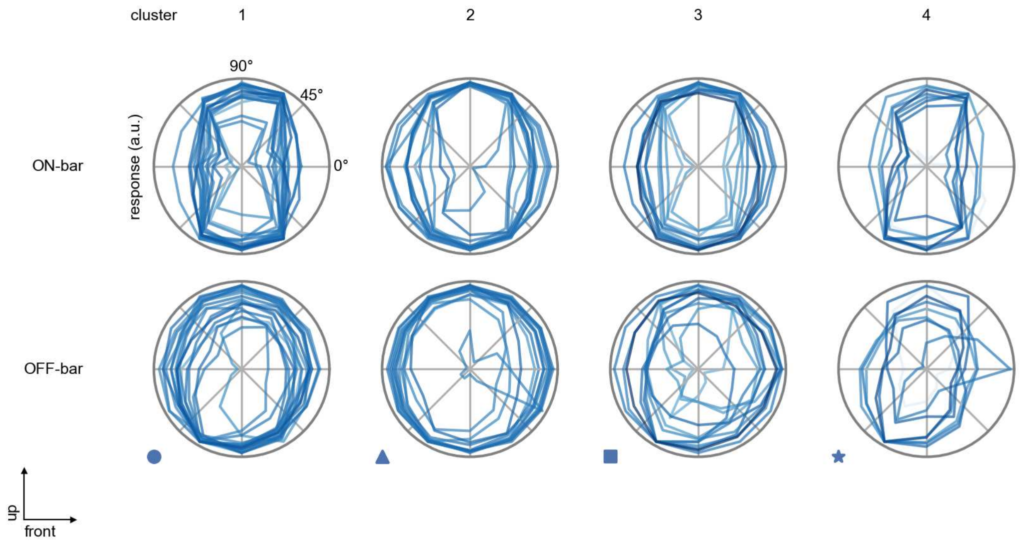

Tm3 - Figure 8: **Peak responses to moving bars.** The top row shows normalized peak responses to moving ON-bars, the bottom row shows normalized peak responses to moving OFF-bars. The bar-stimuli move in different directions from 0 to 360 degrees. Before, a grey-stimulus led the network to a stationary state. The responses from the different models in the different clusters (columns) overlay and are colored in the respective task error of the model. The peak responses are averaged over all assessed bar-speeds.

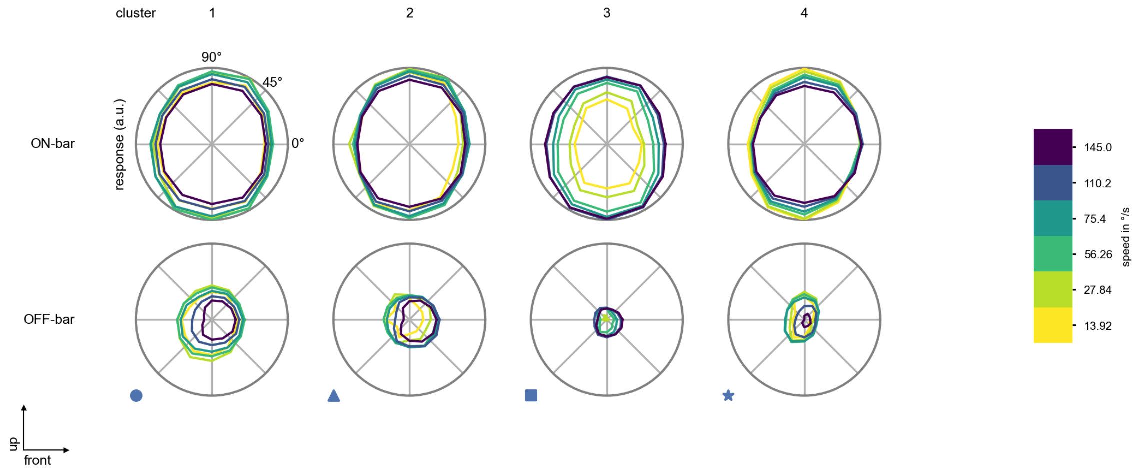

Tm3 - Figure 9: **Peak responses to moving bars from best-performing models.** The top row shows peak responses to moving ON-bars, the bottom row shows peak responses to moving OFF-bars. The bar-stimuli move in different directions from 0 to 360 degrees and at different speeds. Before, a grey-stimulus led the network to a stationary state. The columns show responses from the best-task-performing model in the respective cluster.

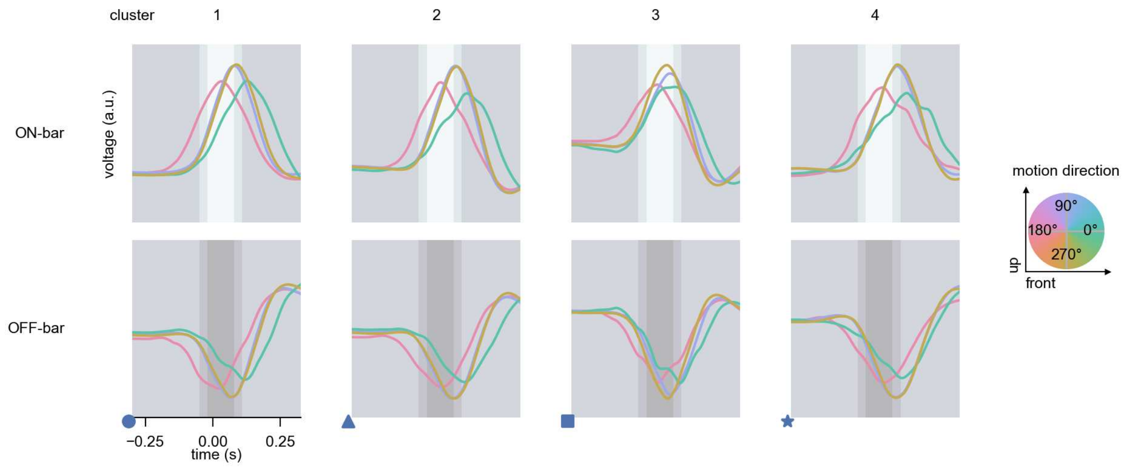

Tm3 - Figure 10: **Responses to moving bars from best-performing models.** The top row shows responses to moving ON-bars, the bottom row shows peak responses to moving OFF-bars. The bar-stimuli move in different directions from 0 to 360 degrees and at different speeds. Before, a grey-stimulus led the network to a stationary state. The columns show responses from the best-task-performing model in the respective cluster. Example responses to bars moving at 75.4°/s in all cardinal directions (green 0°, blue 90°, red 180°, yellow 270°) from -22.5 to 22.5° visual angle. Background contour indicates the bar, passing by the central column at second zero.

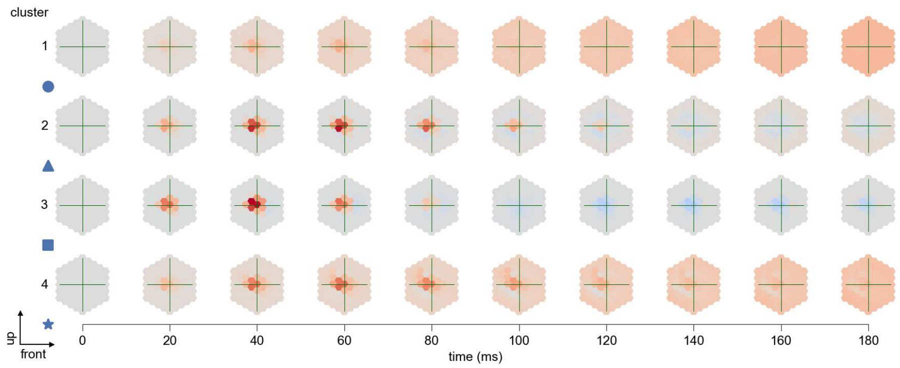

Tm3 - Figure 11: **Spatio-temporal receptive field.** Responses of the central cell to ON-impulses at single ommatidium impulse locations. The impulse occurs at second zero. Before, a grey-stimulus led the network to a stationary state. Responses are baseline-subtracted. Each row illustrates responses from the best-performing model of the respective cluster. Red indicated depolarization, blue indicates hyperpolarization.

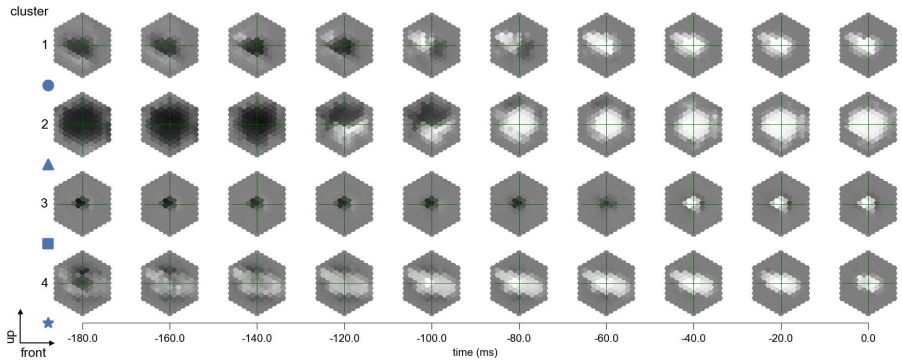

Tm3 - Figure 12: **Maximally excitatory stimuli.** Each row presents the regularized naturalistic-stimulus from the Sintel dataset that maximizes the cell type's central column response at second zero in the best-performing model of the respective cluster.

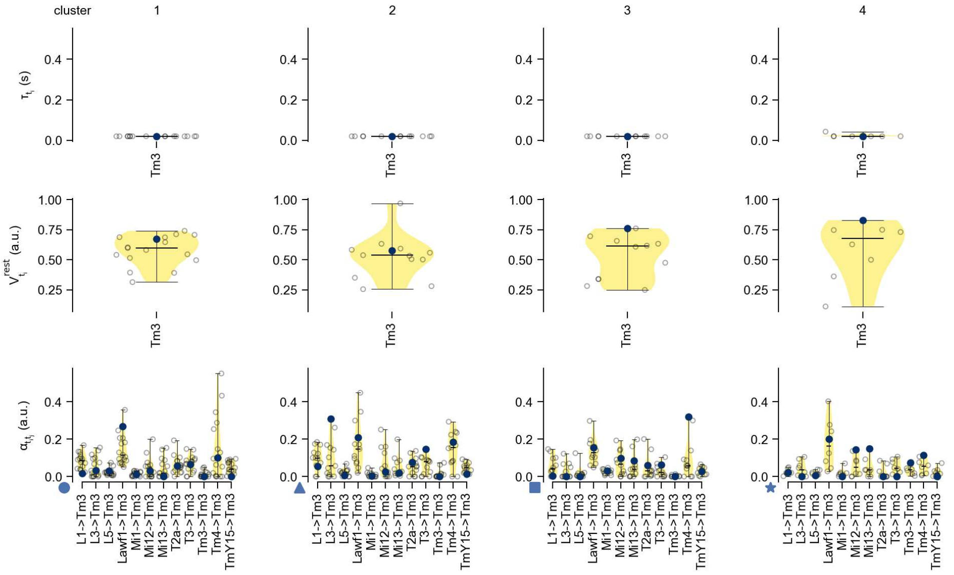

Tm3 - Figure 13: **Task-constrained parameters.** Each column shows the parameters inferred within the respective cluster. First row: learned time constants of the cell type. Second row: resting potentials of the cell type. Third row: scaling factors for the convolutional filters. The blue scatter represents the parameters from the best-performing model within the cluster.

# 47 Tm4

← Cell types

#### Figures

1 Anatomical receptive fields. . . . . 311
2 Anatomical projective fields. . . . . 312
3 Clustering of the responses to naturalistic stimuli. . . . . 312
4 Responses to flashes. . . . . 313
5 Peak responses to moving edges. . . . . 313
6 Peak responses to moving edges from best-performing models. . . . . 314
7 Responses to moving edges from best-performing models. . . . . 314
8 Peak responses to moving bars. . . . . 315
9 Peak responses to moving bars from best-performing models. . . . . 315
10 Responses to moving bars from best-performing models. . . . . 316
11 Spatio-temporal receptive field. . . . . 316
12 Maximally excitatory stimuli. . . . . 316
13 Task-constrained parameters. . . . . 317

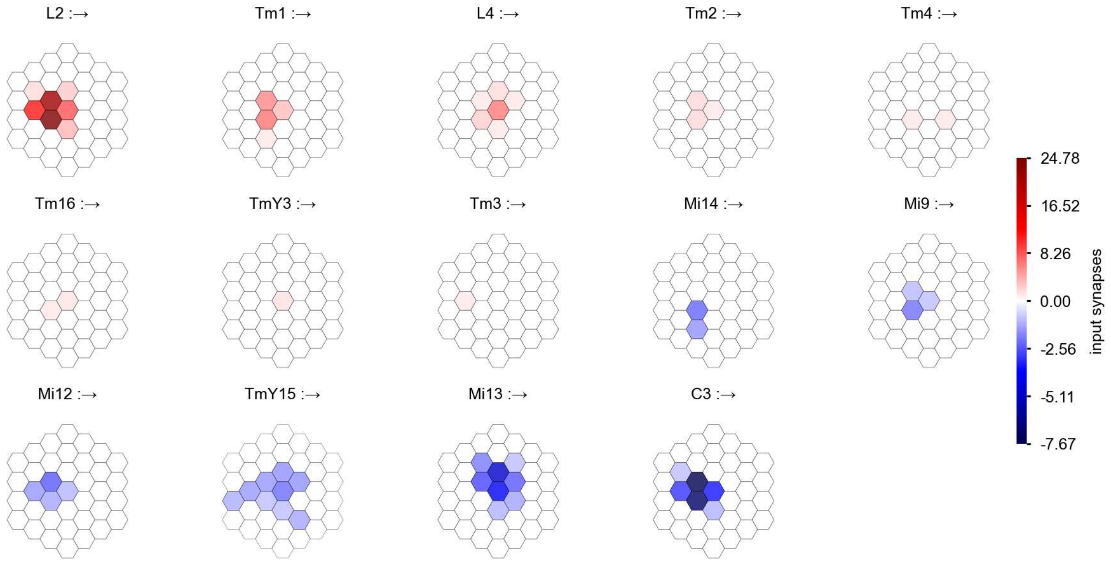

Tm4 - Figure 1: **Anatomical receptive fields.** Each colored hexagon is a projection, with the projection strength characterized by the average number of synapses that we count from the EM reconstruction. Red indicates excitatory synapses, blue indicates inhibitory synapses from inferred signs. Filters in the order of their total number of synapses.

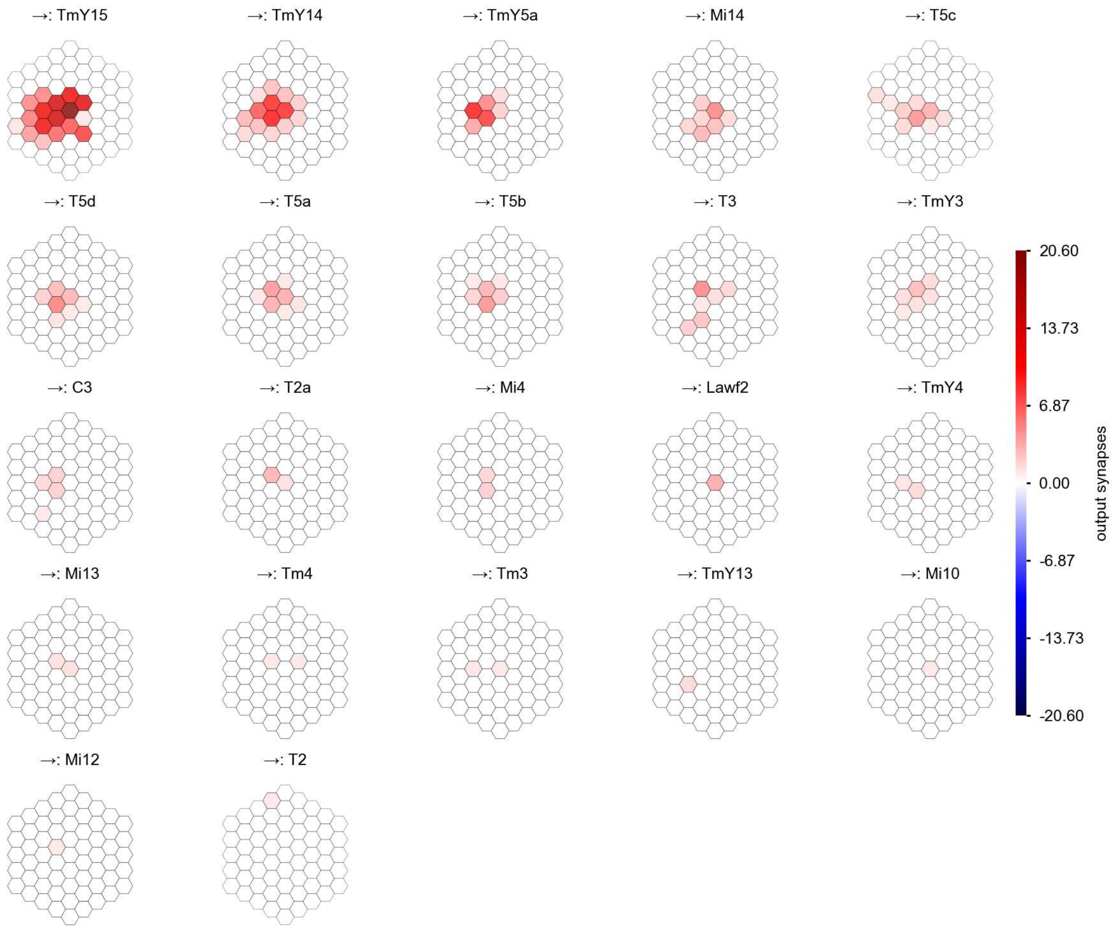

Tm4 - Figure 2: **Anatomical projective fields.** Each colored hexagon is a projection, with the projection strength characterized by the average number of synapses that we count from the EM reconstruction. Red indicates excitatory synapses, blue indicates inhibitory synapses from inferred signs. Filters in the order of their total number of synapses.

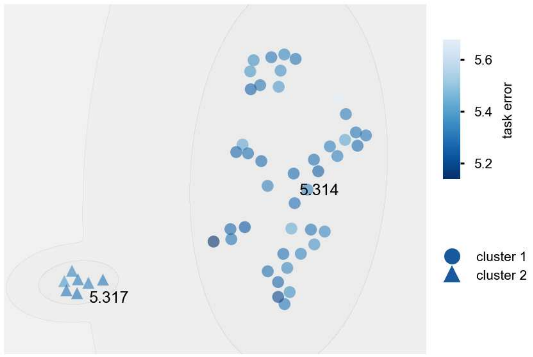

Tm4 - Figure 3: **Clustering of the responses to naturalistic stimuli.** From responses to naturalistic scenes from Sintel. Scatterpoints represent individual models colored by their task error.

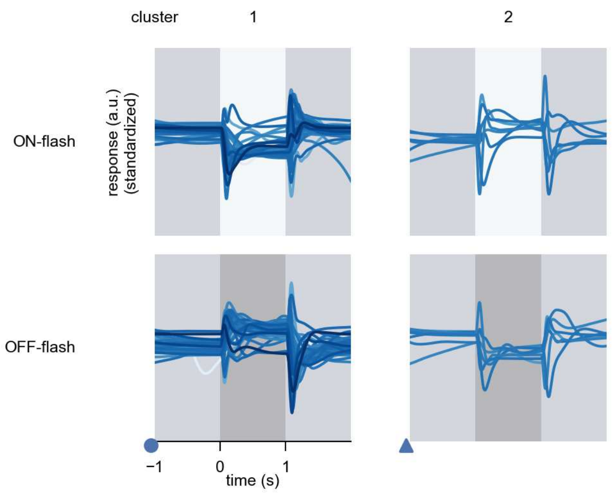

Tm4 - Figure 4: **Responses to flashes.** The top row shows responses to ON-flashes, the bottom row shows responses to OFF-flashes. The responses from the different models in the different clusters (columns) overlay and are colored in the respective task error of the model. The circular flashes occur at second zero, cover 6 ommatidia in radius, and last for one second. Before and after, a grey-stimulus leads to a stationary state of the network. The traces are standardized.

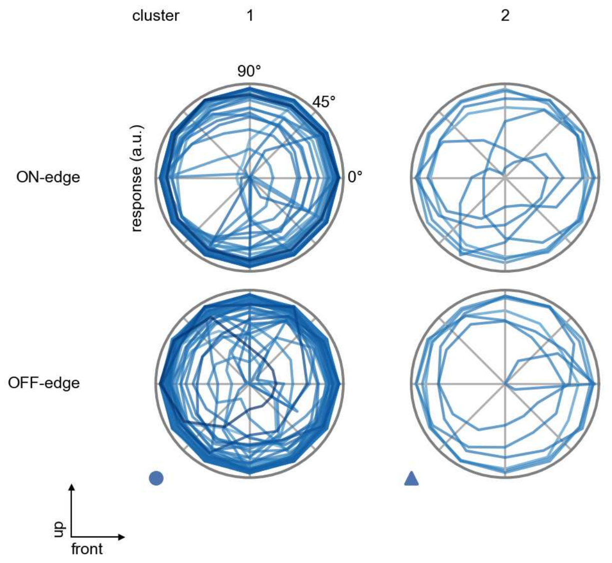

Tm4 - Figure 5: **Peak responses to moving edges.** The top row shows normalized peak responses to moving ON-edges, the bottom row shows normalized peak responses to moving OFF-edges. The edge-stimuli move in different directions from 0 to 360 degrees. Before, a grey-stimulus led the network to a stationary state. The responses from the different models in the different clusters (columns) overlay and are colored in the respective task error of the model. The peak responses are averaged over all assessed edge-speeds.

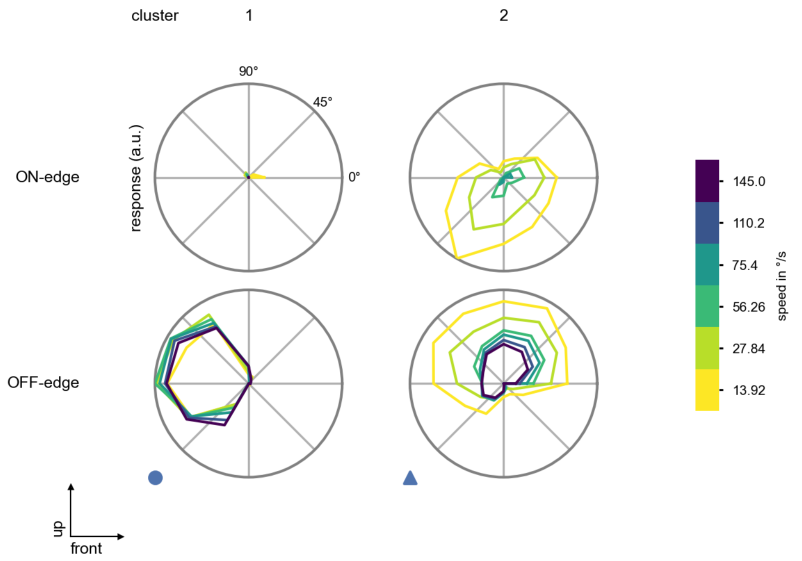

Tm4 - Figure 6: **Peak responses to moving edges from best-performing models.** The top row shows peak responses to moving ON-edges, the bottom row shows peak responses to moving OFF-edges. The edge-stimuli move in different directions from 0 to 360 degrees and at different speeds. Before, a grey-stimulus led the network to a stationary state. The columns show responses from the best-task-performing model in the respective cluster.

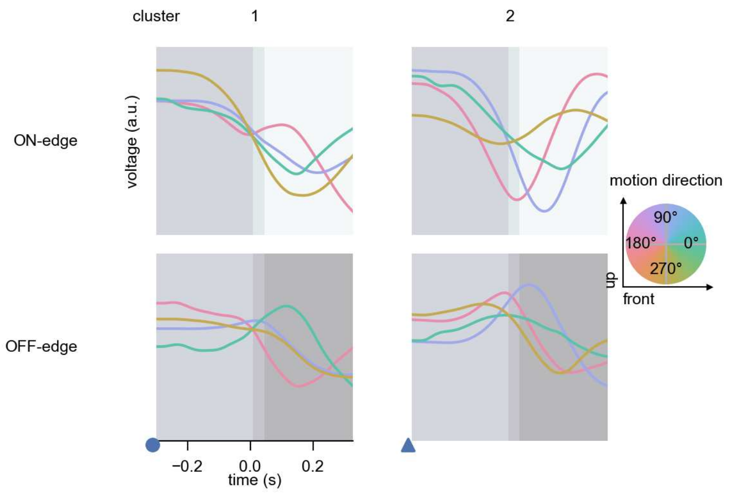

Tm4 - Figure 7: **Responses to moving edges from best-performing models.** The top row shows responses to moving ON-edges, the bottom row shows peak responses to moving OFF-edges. The edge-stimuli move in different directions from 0 to 360 degrees and at different speeds. Before, a grey-stimulus led the network to a stationary state. The columns show responses from the best-task-performing model in the respective cluster. Example responses to edges moving at 75.4°/s in all cardinal directions (green 0°, blue 90°, red 180°, yellow 270°) from -22.5 to 22.5° visual angle. Background contour indicates the edge, passing by the central column at second zero.

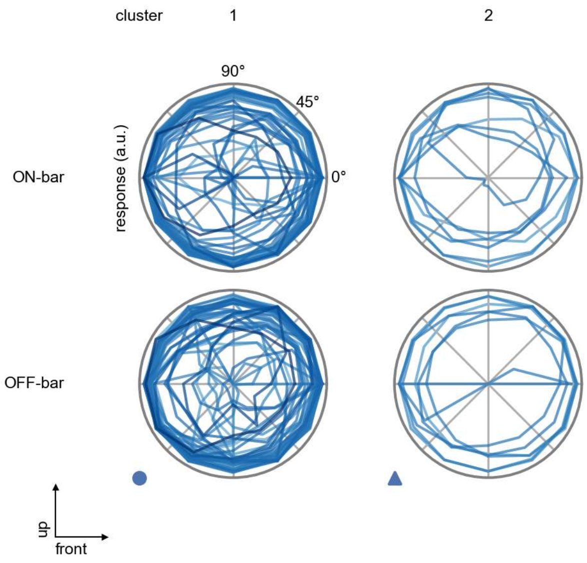

Tm4 - Figure 8: **Peak responses to moving bars.** The top row shows normalized peak responses to moving ON-bars, the bottom row shows normalized peak responses to moving OFF-bars. The bar-stimuli move in different directions from 0 to 360 degrees. Before, a grey-stimulus led the network to a stationary state. The responses from the different models in the different clusters (columns) overlay and are colored in the respective task error of the model. The peak responses are averaged over all assessed bar-speeds.

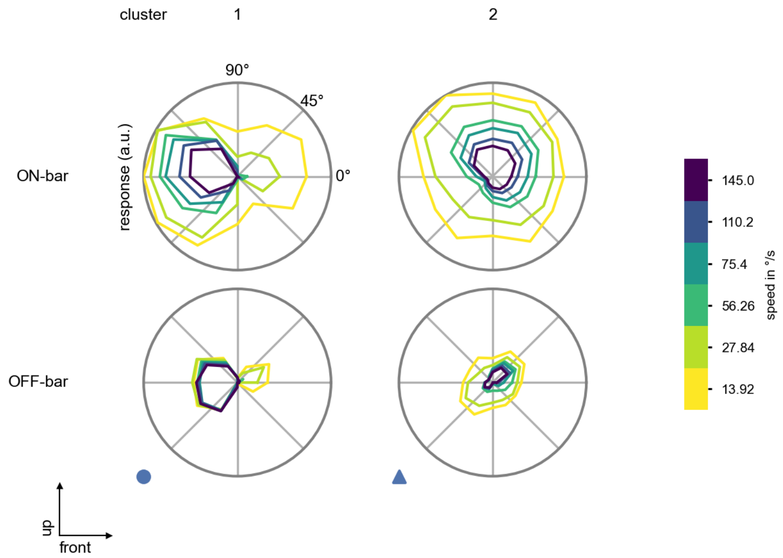

Tm4 - Figure 9: **Peak responses to moving bars from best-performing models.** The top row shows peak responses to moving ON-bars, the bottom row shows peak responses to moving OFF-bars. The bar-stimuli move in different directions from 0 to 360 degrees and at different speeds. Before, a grey-stimulus led the network to a stationary state. The columns show responses from the best-task-performing model in the respective cluster.

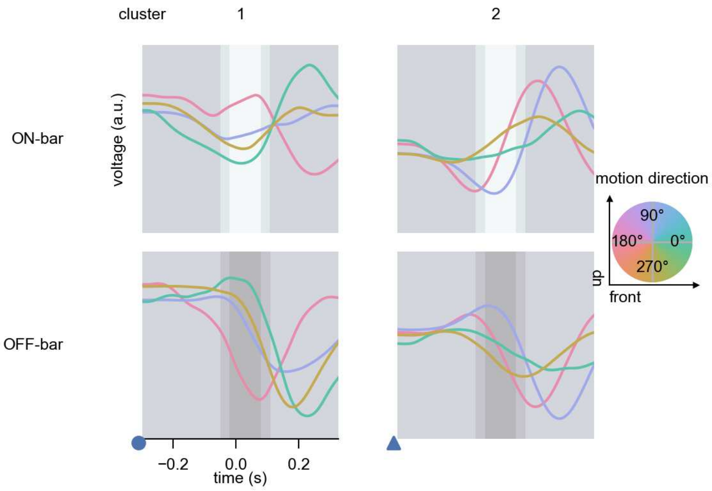

Tm4 - Figure 10: **Responses to moving bars from best-performing models.** The top row shows responses to moving ON-bars, the bottom row shows peak responses to moving OFF-bars. The bar-stimuli move in different directions from 0 to 360 degrees and at different speeds. Before, a grey-stimulus led the network to a stationary state. The columns show responses from the best-task-performing model in the respective cluster. Example responses to bars moving at 75.4°/s in all cardinal directions (green 0°, blue 90°, red 180°, yellow 270°) from -22.5 to 22.5° visual angle. Background contour indicates the bar, passing by the central column at second zero.

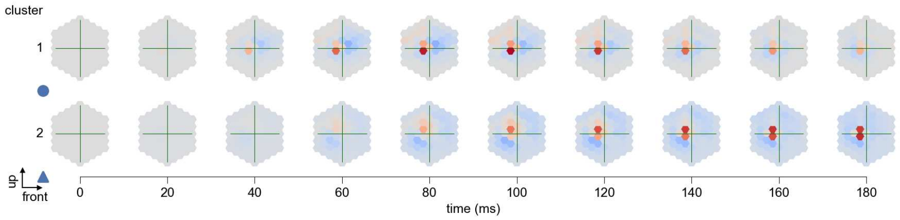

Tm4 - Figure 11: **Spatio-temporal receptive field.** Responses of the central cell to ON-impulses at single ommatidium impulse locations. The impulse occurs at second zero. Before, a grey-stimulus led the network to a stationary state. Responses are baseline-subtracted. Each row illustrates responses from the best-performing model of the respective cluster. Red indicated depolarization, blue indicates hyperpolarization.

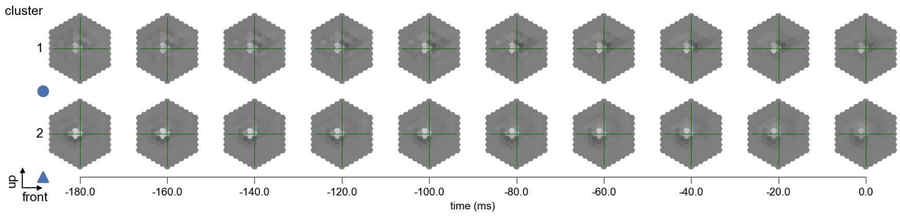

Tm4 - Figure 12: **Maximally excitatory stimuli.** Each row presents the regularized naturalistic-stimulus from the Sintel dataset that maximizes the cell type's central column response at second zero in the best-performing model of the respective cluster.

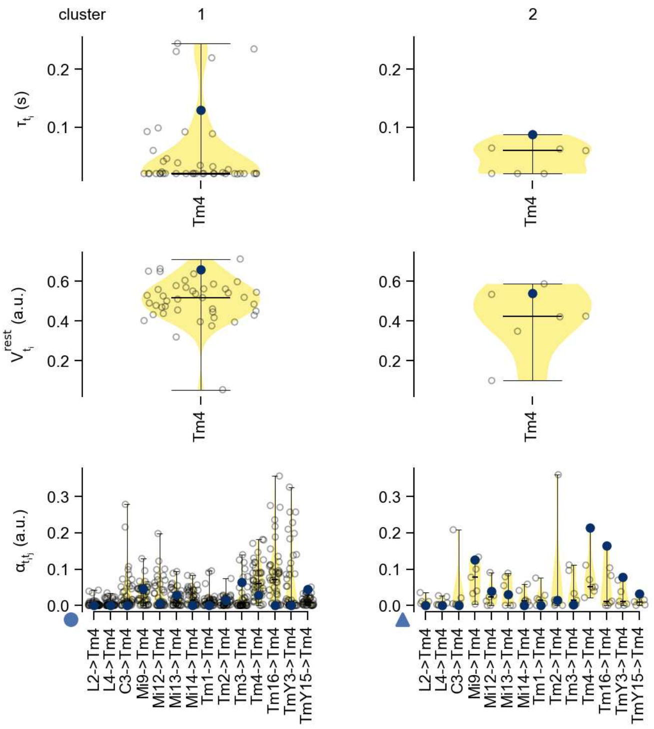

Tm4 - Figure 13: **Task-constrained parameters.** Each column shows the parameters inferred within the respective cluster. First row: learned time constants of the cell type. Second row: resting potentials of the cell type. Third row: scaling factors for the convolutional filters. The blue scatter represents the parameters from the best-performing model within the cluster.

# 48 Tm5Y

← Cell types

#### Figures

1 Anatomical receptive fields. . . . 318
2 Anatomical projective fields. . . . 318
3 Clustering of the responses to naturalistic stimuli. . . . 319
4 Responses to flashes. . . . 319
5 Peak responses to moving edges. . . . 320
6 Peak responses to moving edges from best-performing models. . . . 320
7 Responses to moving edges from best-performing models. . . . 321
8 Peak responses to moving bars. . . . 321
9 Peak responses to moving bars from best-performing models. . . . 322
10 Responses to moving bars from best-performing models. . . . 322
11 Spatio-temporal receptive field. . . . 323
12 Maximally excitatory stimuli. . . . 323
13 Task-constrained parameters. . . . 323

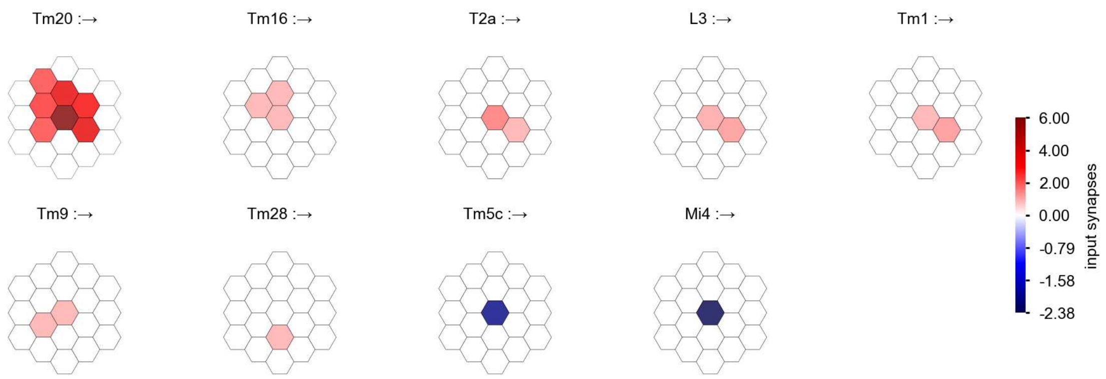

Tm5Y - Figure 1: **Anatomical receptive fields.** Each colored hexagon is a projection, with the projection strength characterized by the average number of synapses that we count from the EM reconstruction. Red indicates excitatory synapses, blue indicates inhibitory synapses from inferred signs. Filters in the order of their total number of synapses.

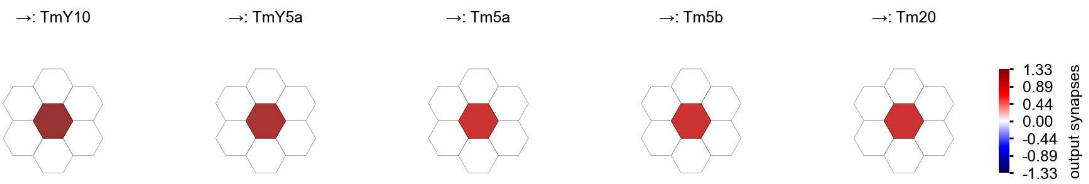

Tm5Y - Figure 2: **Anatomical projective fields.** Each colored hexagon is a projection, with the projection strength characterized by the average number of synapses that we count from the EM reconstruction. Red indicates excitatory synapses, blue indicates inhibitory synapses from inferred signs. Filters in the order of their total number of synapses.

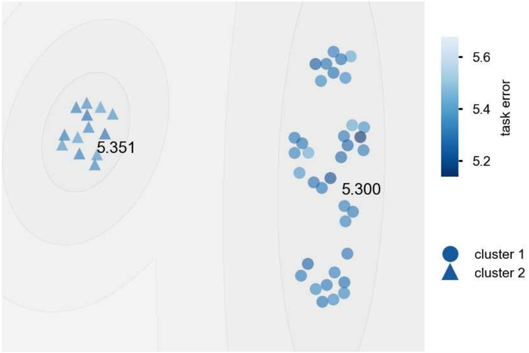

Tm5Y - Figure 3: **Clustering of the responses to naturalistic stimuli.** From responses to naturalistic scenes from Sintel. Scatterpoints represent individual models colored by their task error.

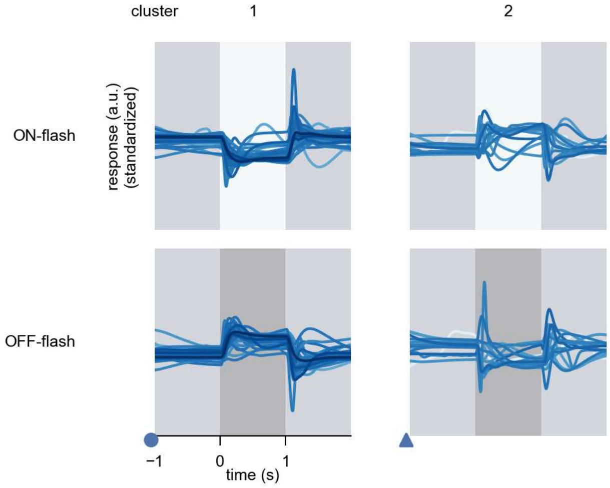

Tm5Y - Figure 4: **Responses to flashes.** The top row shows responses to ON-flashes, the bottom row shows responses to OFF-flashes. The responses from the different models in the different clusters (columns) overlay and are colored in the respective task error of the model. The circular flashes occur at second zero, cover 6 ommatidia in radius, and last for one second. Before and after, a grey-stimulus leads to a stationary state of the network. The traces are standardized.

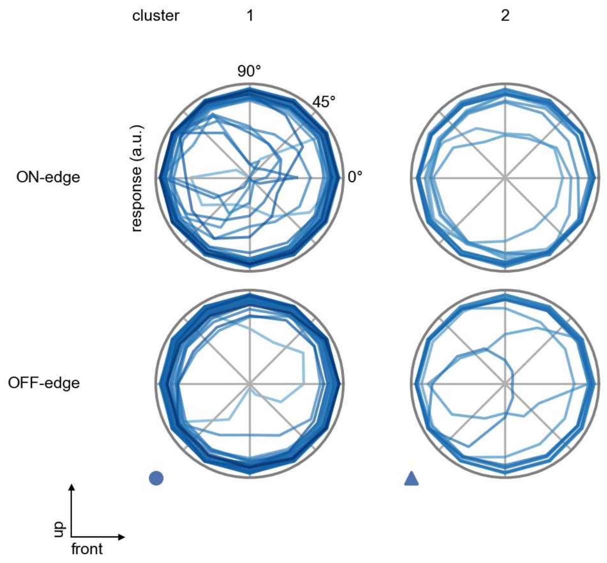

Tm5Y - Figure 5: **Peak responses to moving edges.** The top row shows normalized peak responses to moving ON-edges, the bottom row shows normalized peak responses to moving OFF-edges. The edge-stimuli move in different directions from 0 to 360 degrees. Before, a grey-stimulus led the network to a stationary state. The responses from the different models in the different clusters (columns) overlay and are colored in the respective task error of the model. The peak responses are averaged over all assessed edge-speeds.

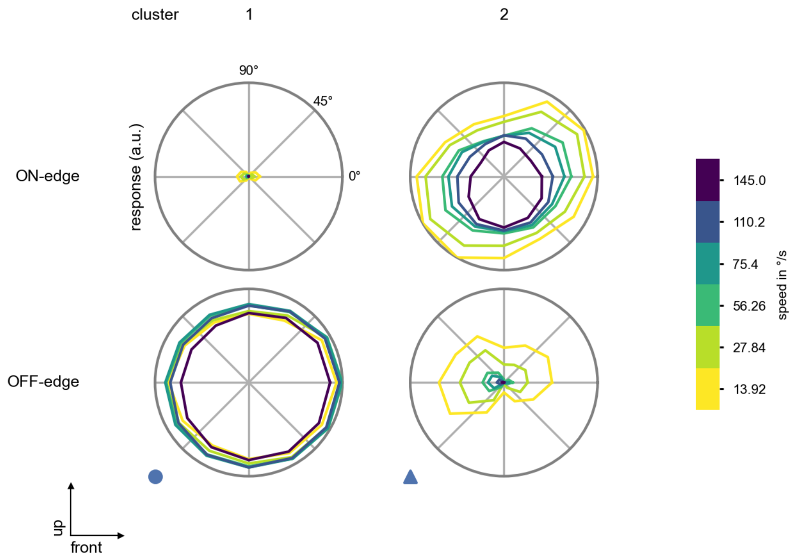

Tm5Y - Figure 6: **Peak responses to moving edges from best-performing models.** The top row shows peak responses to moving ON-edges, the bottom row shows peak responses to moving OFF-edges. The edge-stimuli move in different directions from 0 to 360 degrees and at different speeds. Before, a grey-stimulus led the network to a stationary state. The columns show responses from the best-task-performing model in the respective cluster.

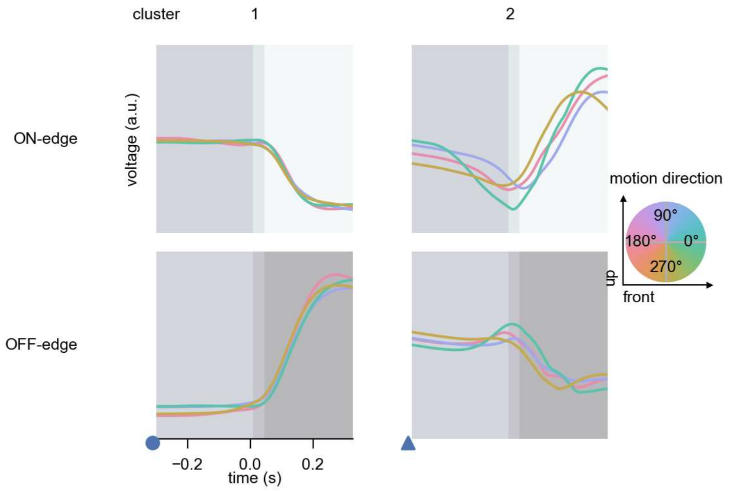

Tm5Y - Figure 7: **Responses to moving edges from best-performing models.** The top row shows responses to moving ON-edges, the bottom row shows peak responses to moving OFF-edges. The edge-stimuli move in different directions from 0 to 360 degrees and at different speeds. Before, a grey-stimulus led the network to a stationary state. The columns show responses from the best-task-performing model in the respective cluster. Example responses to edges moving at 75.4°/s in all cardinal directions (green 0°, blue 90°, red 180°, yellow 270°) from -22.5 to 22.5° visual angle. Background contour indicates the edge, passing by the central column at second zero.

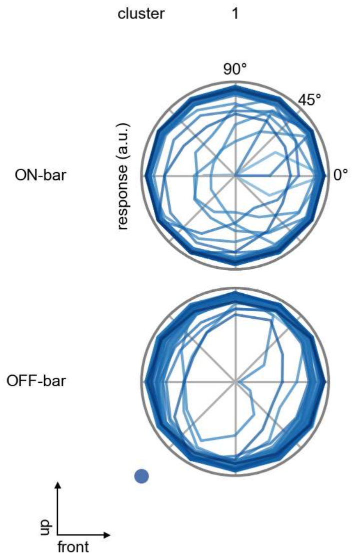

Tm5Y - Figure 8: **Peak responses to moving bars.** The top row shows normalized peak responses to moving ON-bars, the bottom row shows normalized peak responses to moving OFF-bars. The bar-stimuli move in different directions from 0 to 360 degrees. Before, a grey-stimulus led the network to a stationary state. The responses from the different models in the different clusters (columns) overlay and are colored in the respective task error of the model. The peak responses are averaged over all assessed bar-speeds.

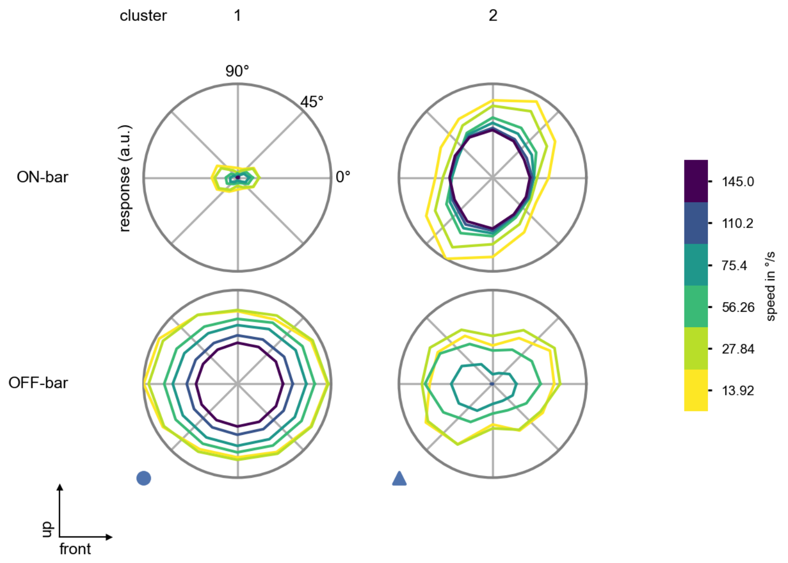

Tm5Y - Figure 9: **Peak responses to moving bars from best-performing models.** The top row shows peak responses to moving ON-bars, the bottom row shows peak responses to moving OFF-bars. The bar-stimuli move in different directions from 0 to 360 degrees and at different speeds. Before, a grey-stimulus led the network to a stationary state. The columns show responses from the best-task-performing model in the respective cluster.

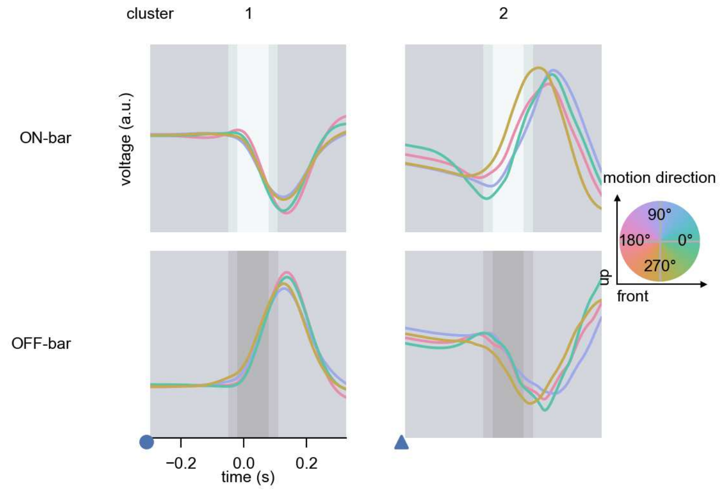

Tm5Y - Figure 10: **Responses to moving bars from best-performing models.** The top row shows responses to moving ON-bars, the bottom row shows peak responses to moving OFF-bars. The bar-stimuli move in different directions from 0 to 360 degrees and at different speeds. Before, a grey-stimulus led the network to a stationary state. The columns show responses from the best-task-performing model in the respective cluster. Example responses to bars moving at 75.4°/s in all cardinal directions (green 0°, blue 90°, red 180°, yellow 270°) from -22.5 to 22.5° visual angle. Background contour indicates the bar, passing by the central column at second zero.

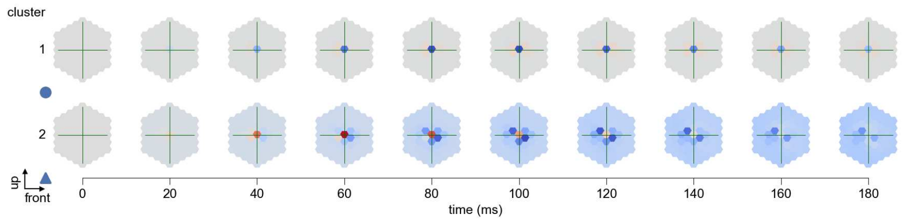

Tm5Y - Figure 11: **Spatio-temporal receptive field.** Responses of the central cell to ON-impulses at single ommatidium impulse locations. The impulse occurs at second zero. Before, a grey-stimulus led the network to a stationary state. Responses are baseline-subtracted. Each row illustrates responses from the best-performing model of the respective cluster. Red indicated depolarization, blue indicates hyperpolarization.

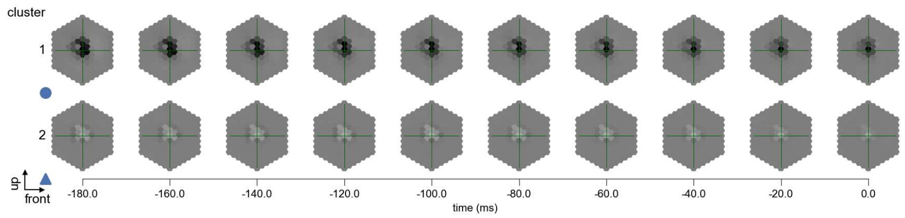

Tm5Y - Figure 12: **Maximally excitatory stimuli.** Each row presents the regularized naturalistic-stimulus from the Sintel dataset that maximizes the cell type's central column response at second zero in the best-performing model of the respective cluster.

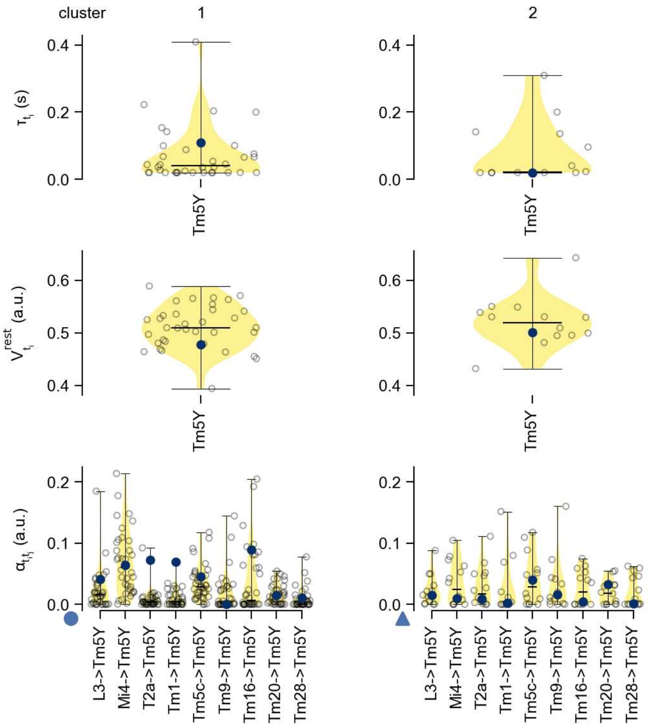

Tm5Y - Figure 13: **Task-constrained parameters.** Each column shows the parameters inferred within the respective cluster. First row: learned time constants of the cell type. Second row: resting potentials of the cell type. Third row: scaling factors for the convolutional filters. The blue scatter represents the parameters from the best-performing model within the cluster.

# 49 Tm5a

← Cell types

#### Figures

1 Anatomical receptive fields. . . . 324
2 Anatomical projective fields. . . . 324
3 Clustering of the responses to naturalistic stimuli. . . . 325
4 Responses to flashes. . . . 325
5 Peak responses to moving edges. . . . 325
6 Peak responses to moving edges from best-performing models. . . . 326
7 Responses to moving edges from best-performing models. . . . 326
8 Peak responses to moving bars. . . . 327
9 Peak responses to moving bars from best-performing models. . . . 327
10 Responses to moving bars from best-performing models. . . . 328
11 Spatio-temporal receptive field. . . . 328
12 Maximally excitatory stimuli. . . . 329
13 Task-constrained parameters. . . . 329

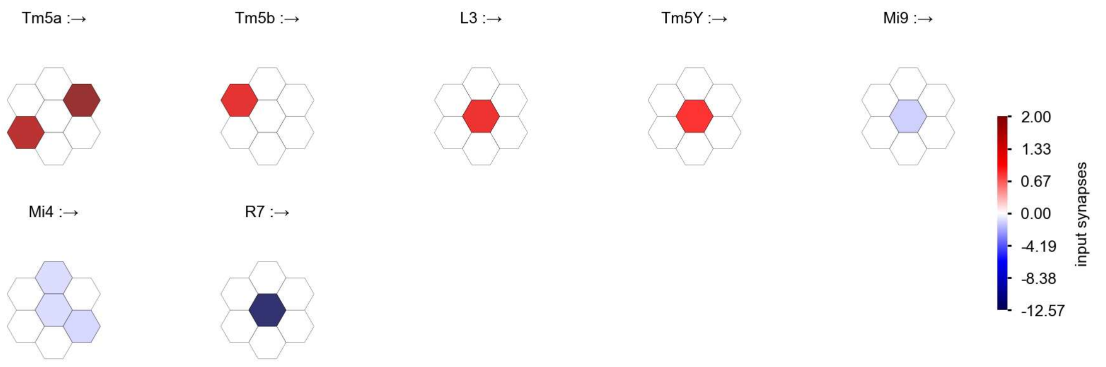

Tm5a - Figure 1: **Anatomical receptive fields.** Each colored hexagon is a projection, with the projection strength characterized by the average number of synapses that we count from the EM reconstruction. Red indicates excitatory synapses, blue indicates inhibitory synapses from inferred signs. Filters in the order of their total number of synapses.

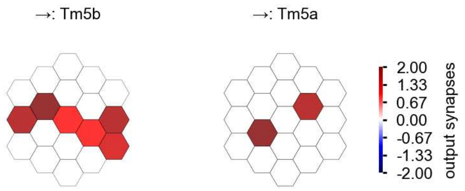

Tm5a - Figure 2: **Anatomical projective fields.** Each colored hexagon is a projection, with the projection strength characterized by the average number of synapses that we count from the EM reconstruction. Red indicates excitatory synapses, blue indicates inhibitory synapses from inferred signs. Filters in the order of their total number of synapses.

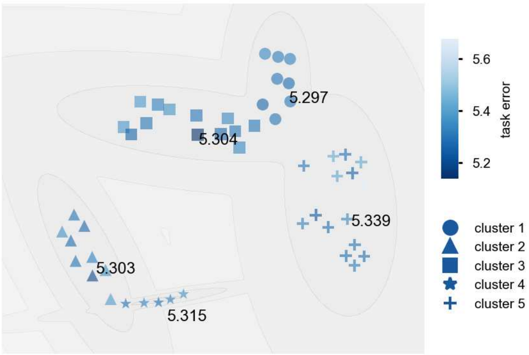

Tm5a - Figure 3: **Clustering of the responses to naturalistic stimuli.** From responses to naturalistic scenes from Sintel. Scatterpoints represent individual models colored by their task error.

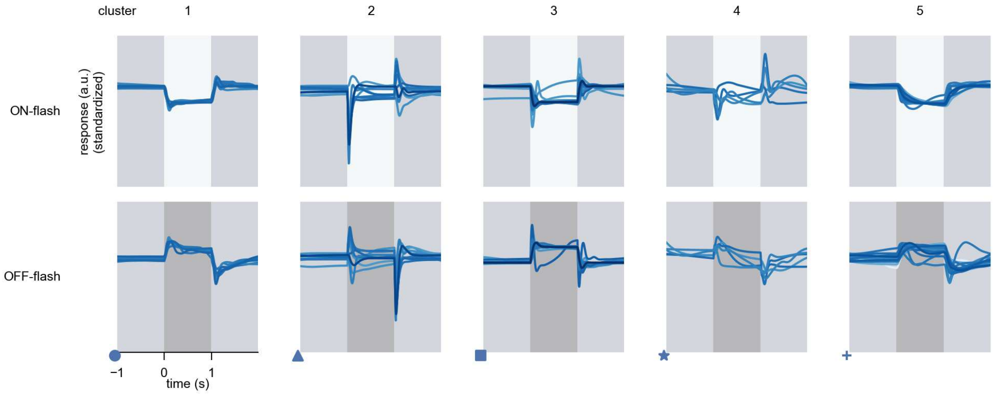

Tm5a - Figure 4: **Responses to flashes.** The top row shows responses to ON-flashes, the bottom row shows responses to OFF-flashes. The responses from the different models in the different clusters (columns) overlay and are colored in the respective task error of the model. The circular flashes occur at second zero, cover 6 ommatidia in radius, and last for one second. Before and after, a grey-stimulus leads to a stationary state of the network. The traces are standardized.

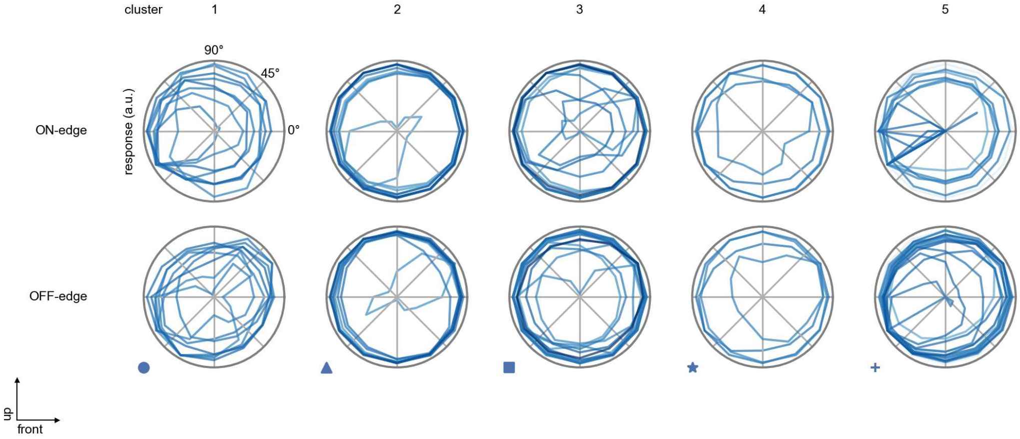

Tm5a - Figure 5: **Peak responses to moving edges.** The top row shows normalized peak responses to moving ON-edges, the bottom row shows normalized peak responses to moving OFF-edges. The edge-stimuli move in different directions from 0 to 360 degrees. Before, a grey-stimulus led the network to a stationary state. The responses from the different models in the different clusters (columns) overlay and are colored in the respective task error of the model. The peak responses are averaged over all assessed edge-speeds.

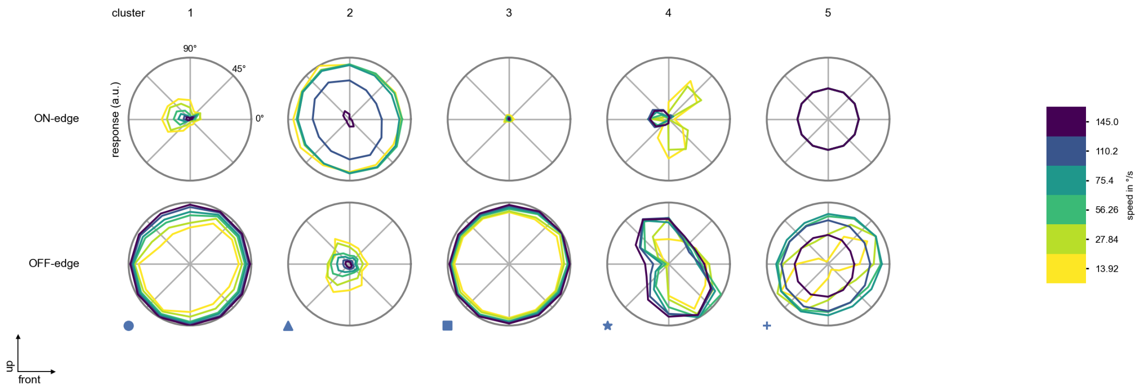

Tm5a - Figure 6: **Peak responses to moving edges from best-performing models.** The top row shows peak responses to moving ON-edges, the bottom row shows peak responses to moving OFF-edges. The edge-stimuli move in different directions from 0 to 360 degrees and at different speeds. Before, a grey-stimulus led the network to a stationary state. The columns show responses from the best-task-performing model in the respective cluster.

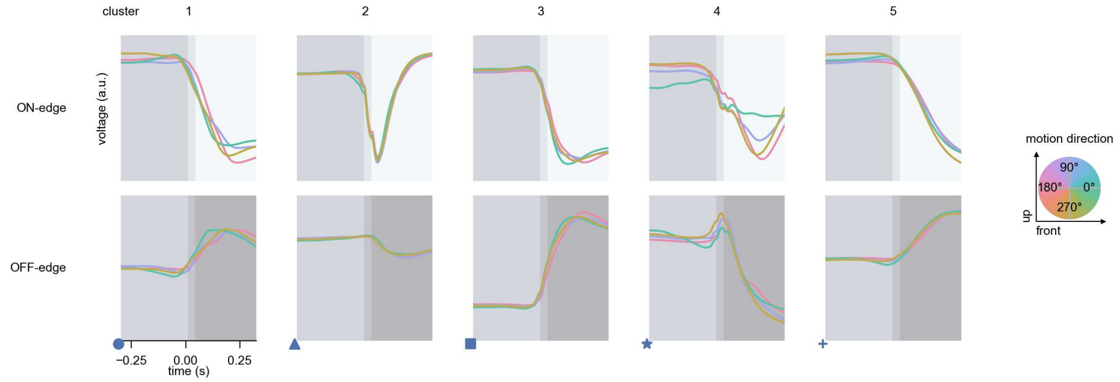

Tm5a - Figure 7: **Responses to moving edges from best-performing models.** The top row shows responses to moving ON-edges, the bottom row shows peak responses to moving OFF-edges. The edge-stimuli move in different directions from 0 to 360 degrees and at different speeds. Before, a grey-stimulus led the network to a stationary state. The columns show responses from the best-task-performing model in the respective cluster. Example responses to edges moving at 75.4°/s in all cardinal directions (green 0°, blue 90°, red 180°, yellow 270°) from -22.5 to 22.5° visual angle. Background contour indicates the edge, passing by the central column at second zero.

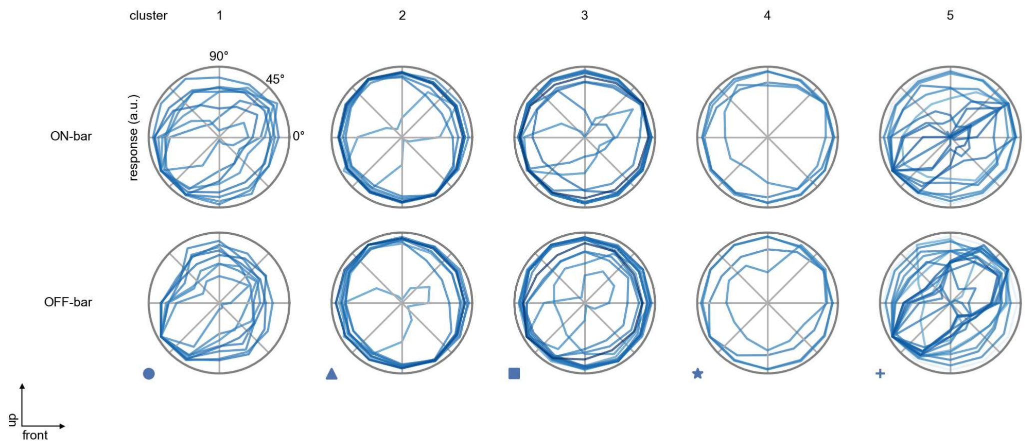

Tm5a - Figure 8: **Peak responses to moving bars.** The top row shows normalized peak responses to moving ON-bars, the bottom row shows normalized peak responses to moving OFF-bars. The bar-stimuli move in different directions from 0 to 360 degrees. Before, a grey-stimulus led the network to a stationary state. The responses from the different models in the different clusters (columns) overlay and are colored in the respective task error of the model. The peak responses are averaged over all assessed bar-speeds.

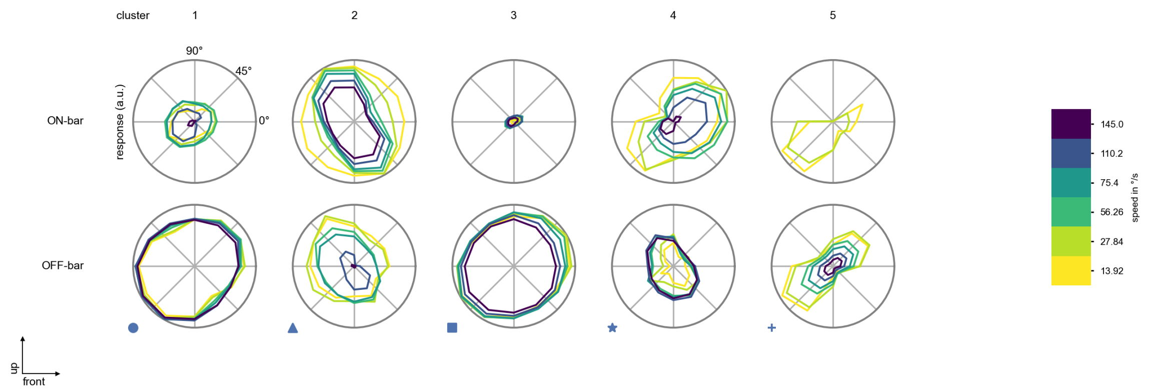

Tm5a - Figure 9: **Peak responses to moving bars from best-performing models.** The top row shows peak responses to moving ON-bars, the bottom row shows peak responses to moving OFF-bars. The bar-stimuli move in different directions from 0 to 360 degrees and at different speeds. Before, a grey-stimulus led the network to a stationary state. The columns show responses from the best-task-performing model in the respective cluster.

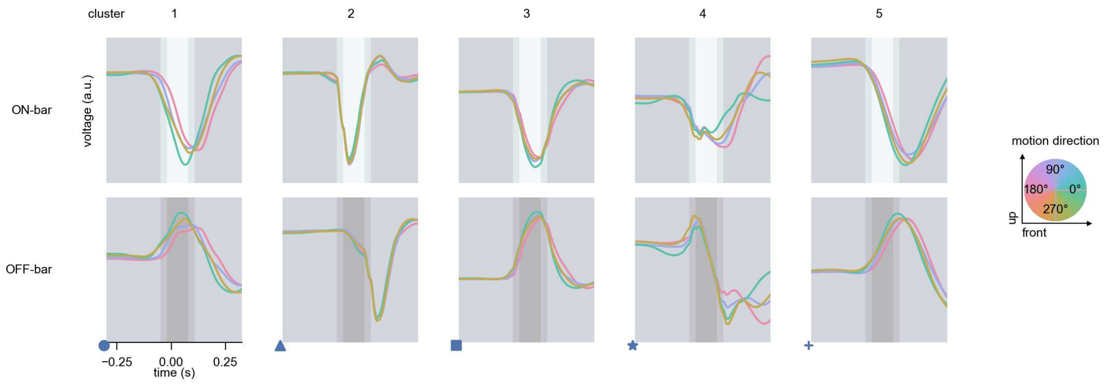

Tm5a - Figure 10: **Responses to moving bars from best-performing models.** The top row shows responses to moving ON-bars, the bottom row shows peak responses to moving OFF-bars. The bar-stimuli move in different directions from 0 to 360 degrees and at different speeds. Before, a grey-stimulus led the network to a stationary state. The columns show responses from the best-task-performing model in the respective cluster. Example responses to bars moving at 75.4°/s in all cardinal directions (green 0°, blue 90°, red 180°, yellow 270°) from -22.5 to 22.5° visual angle. Background contour indicates the bar, passing by the central column at second zero.

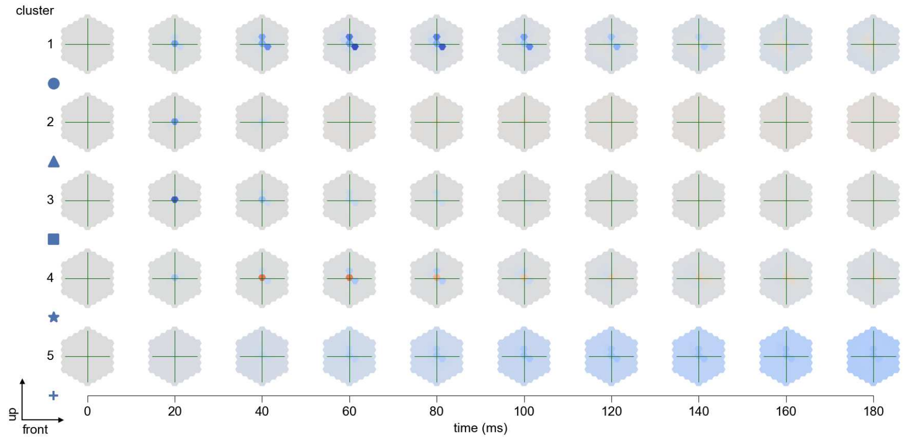

Tm5a - Figure 11: **Spatio-temporal receptive field.** Responses of the central cell to ON-impulses at single ommatidium impulse locations. The impulse occurs at second zero. Before, a grey-stimulus led the network to a stationary state. Responses are baseline-subtracted. Each row illustrates responses from the best-performing model of the respective cluster. Red indicated depolarization, blue indicates hyperpolarization.

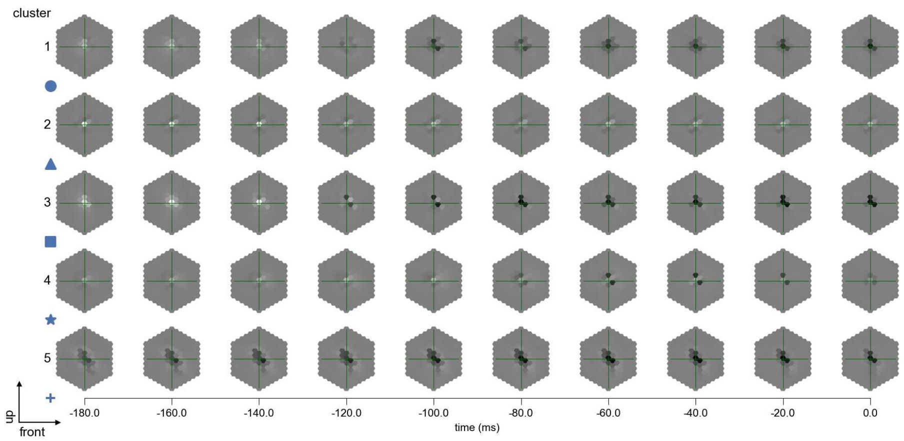

Tm5a - Figure 12: **Maximally excitatory stimuli.** Each row presents the regularized naturalistic-stimulus from the Sintel dataset that maximizes the cell type's central column response at second zero in the best-performing model of the respective cluster.

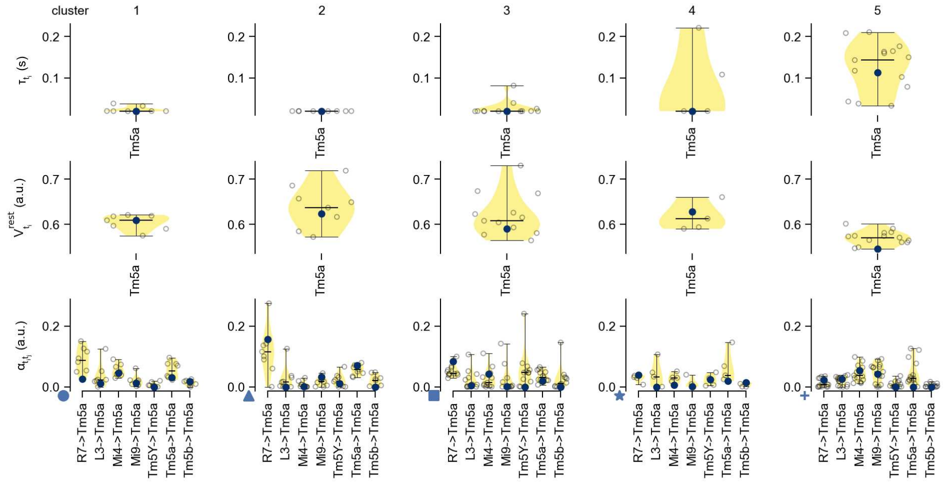

Tm5a - Figure 13: **Task-constrained parameters.** Each column shows the parameters inferred within the respective cluster. First row: learned time constants of the cell type. Second row: resting potentials of the cell type. Third row: scaling factors for the convolutional filters. The blue scatter represents the parameters from the best-performing model within the cluster.

# 50 Tm5b

← Cell types

#### Figures

1 Anatomical receptive fields. . . . 330
2 Anatomical projective fields. . . . 330
3 Clustering of the responses to naturalistic stimuli. . . . 331
4 Responses to flashes. . . . 331
5 Peak responses to moving edges. . . . 332
6 Peak responses to moving edges from best-performing models. . . . 332
7 Responses to moving edges from best-performing models. . . . 333
8 Peak responses to moving bars. . . . 333
9 Peak responses to moving bars from best-performing models. . . . 334
10 Responses to moving bars from best-performing models. . . . 334
11 Spatio-temporal receptive field. . . . 335
12 Maximally excitatory stimuli. . . . 335
13 Task-constrained parameters. . . . 335

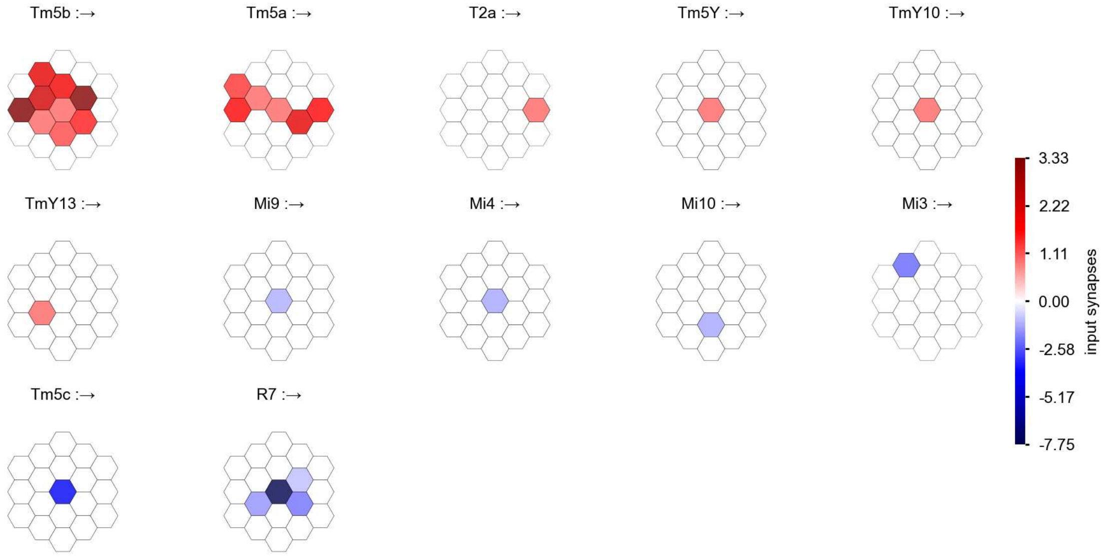

Tm5b - Figure 1: **Anatomical receptive fields.** Each colored hexagon is a projection, with the projection strength characterized by the average number of synapses that we count from the EM reconstruction. Red indicates excitatory synapses, blue indicates inhibitory synapses from inferred signs. Filters in the order of their total number of synapses.

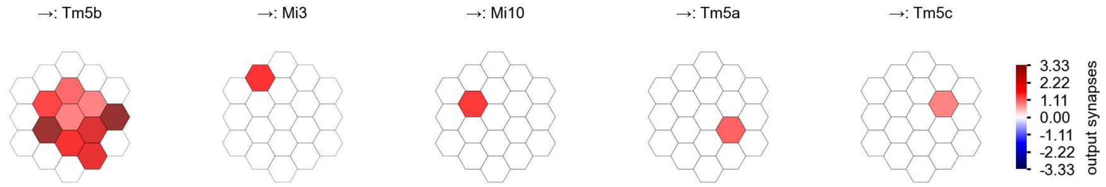

Tm5b - Figure 2: **Anatomical projective fields.** Each colored hexagon is a projection, with the projection strength characterized by the average number of synapses that we count from the EM reconstruction. Red indicates excitatory synapses, blue indicates inhibitory synapses from inferred signs. Filters in the order of their total number of synapses.

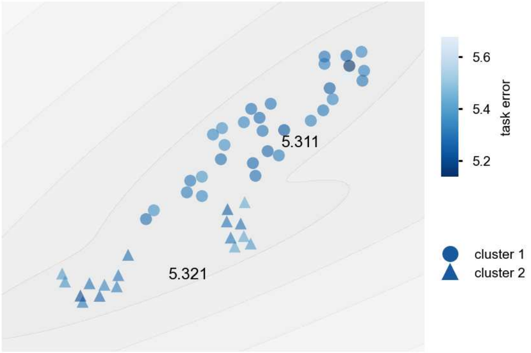

Tm5b - Figure 3: **Clustering of the responses to naturalistic stimuli.** From responses to naturalistic scenes from Sintel. Scatterpoints represent individual models colored by their task error.

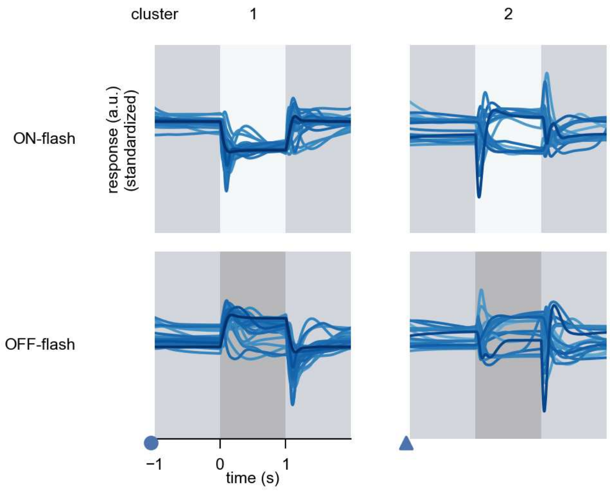

Tm5b - Figure 4: **Responses to flashes.** The top row shows responses to ON-flashes, the bottom row shows responses to OFF-flashes. The responses from the different models in the different clusters (columns) overlay and are colored in the respective task error of the model. The circular flashes occur at second zero, cover 6 ommatidia in radius, and last for one second. Before and after, a grey-stimulus leads to a stationary state of the network. The traces are standardized.

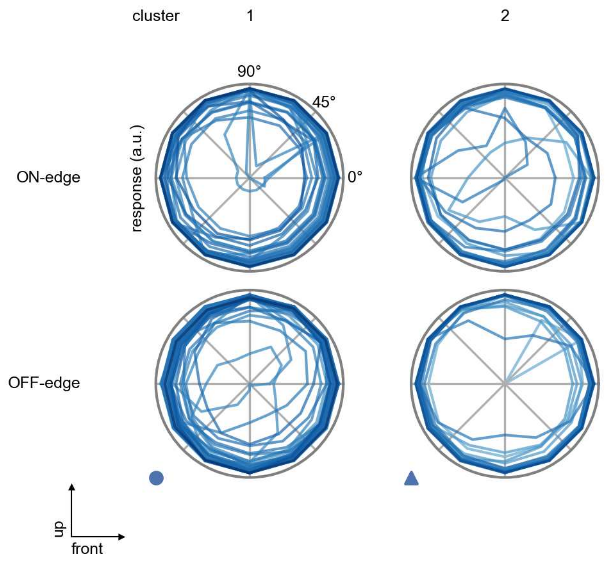

Tm5b - Figure 5: **Peak responses to moving edges.** The top row shows normalized peak responses to moving ON-edges, the bottom row shows normalized peak responses to moving OFF-edges. The edge-stimuli move in different directions from 0 to 360 degrees. Before, a grey-stimulus led the network to a stationary state. The responses from the different models in the different clusters (columns) overlay and are colored in the respective task error of the model. The peak responses are averaged over all assessed edge-speeds.

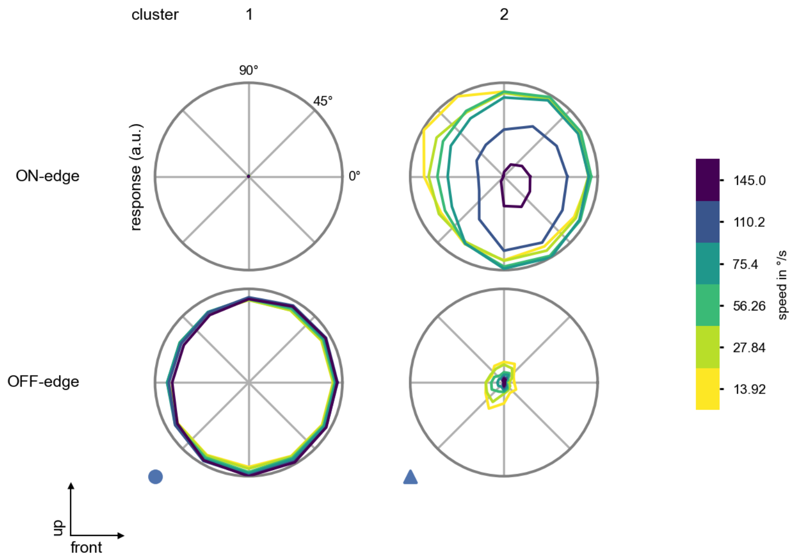

Tm5b - Figure 6: **Peak responses to moving edges from best-performing models.** The top row shows peak responses to moving ON-edges, the bottom row shows peak responses to moving OFF-edges. The edge-stimuli move in different directions from 0 to 360 degrees and at different speeds. Before, a grey-stimulus led the network to a stationary state. The columns show responses from the best-task-performing model in the respective cluster.

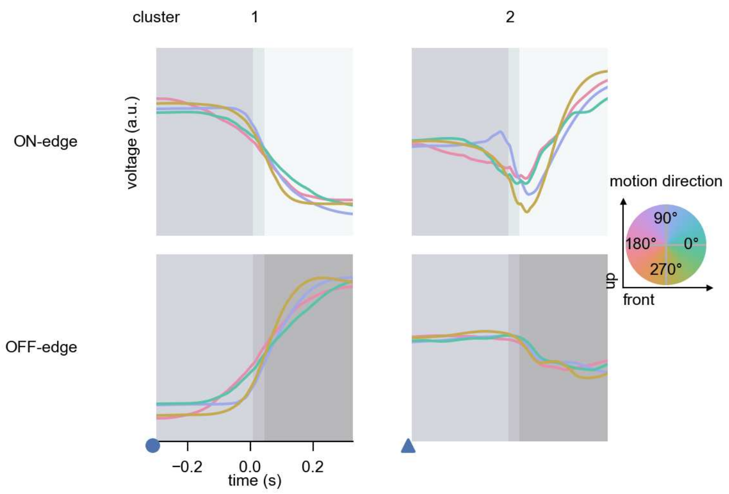

Tm5b - Figure 7: **Responses to moving edges from best-performing models.** The top row shows responses to moving ON-edges, the bottom row shows peak responses to moving OFF-edges. The edge-stimuli move in different directions from 0 to 360 degrees and at different speeds. Before, a grey-stimulus led the network to a stationary state. The columns show responses from the best-task-performing model in the respective cluster. Example responses to edges moving at 75.4°/s in all cardinal directions (green 0°, blue 90°, red 180°, yellow 270°) from -22.5 to 22.5° visual angle. Background contour indicates the edge, passing by the central column at second zero.

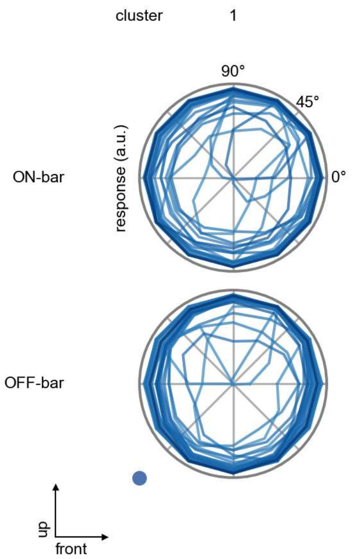

Tm5b - Figure 8: **Peak responses to moving bars.** The top row shows normalized peak responses to moving ON-bars, the bottom row shows normalized peak responses to moving OFF-bars. The bar-stimuli move in different directions from 0 to 360 degrees. Before, a grey-stimulus led the network to a stationary state. The responses from the different models in the different clusters (columns) overlay and are colored in the respective task error of the model. The peak responses are averaged over all assessed bar-speeds.

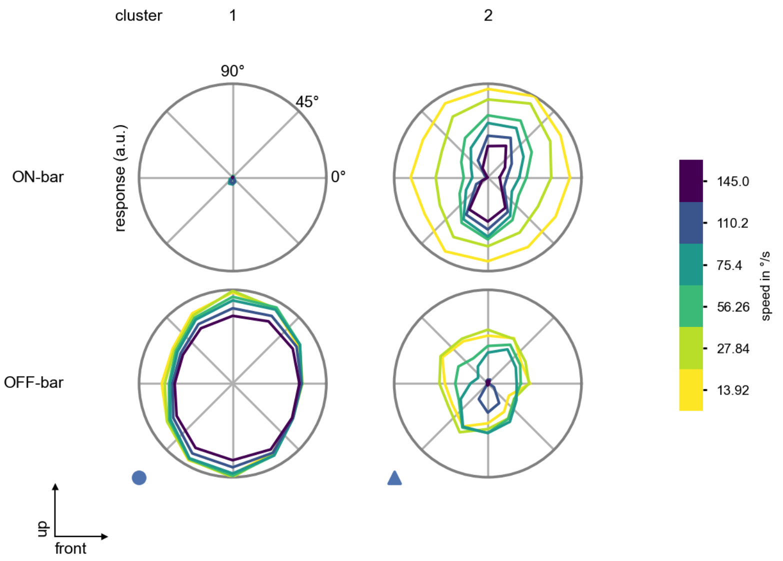

Tm5b - Figure 9: **Peak responses to moving bars from best-performing models.** The top row shows peak responses to moving ON-bars, the bottom row shows peak responses to moving OFF-bars. The bar-stimuli move in different directions from 0 to 360 degrees and at different speeds. Before, a grey-stimulus led the network to a stationary state. The columns show responses from the best-task-performing model in the respective cluster.

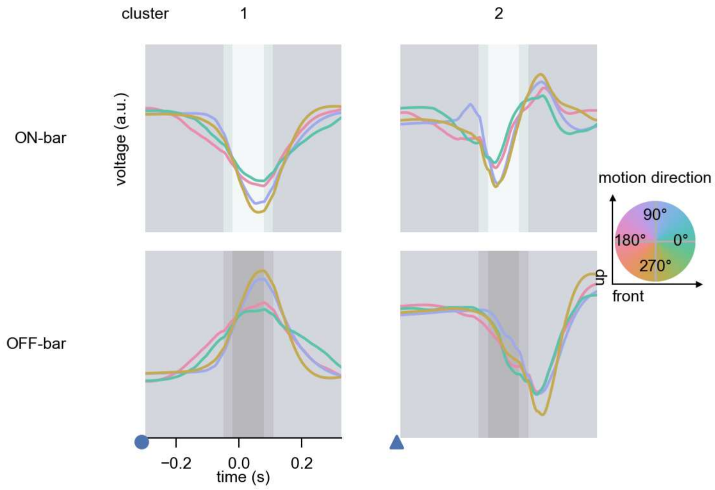

Tm5b - Figure 10: **Responses to moving bars from best-performing models.** The top row shows responses to moving ON-bars, the bottom row shows peak responses to moving OFF-bars. The bar-stimuli move in different directions from 0 to 360 degrees and at different speeds. Before, a grey-stimulus led the network to a stationary state. The columns show responses from the best-task-performing model in the respective cluster. Example responses to bars moving at 75.4°/s in all cardinal directions (green 0°, blue 90°, red 180°, yellow 270°) from -22.5 to 22.5° visual angle. Background contour indicates the bar, passing by the central column at second zero.

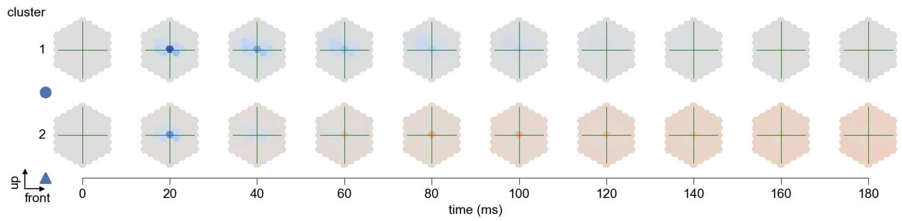

Tm5b - Figure 11: **Spatio-temporal receptive field.** Responses of the central cell to ON-impulses at single ommatidium impulse locations. The impulse occurs at second zero. Before, a grey-stimulus led the network to a stationary state. Responses are baseline-subtracted. Each row illustrates responses from the best-performing model of the respective cluster. Red indicated depolarization, blue indicates hyperpolarization.

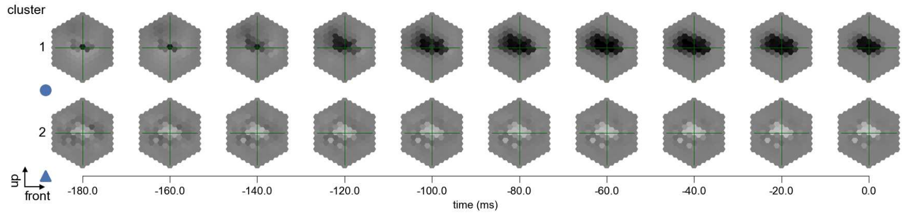

Tm5b - Figure 12: **Maximally excitatory stimuli.** Each row presents the regularized naturalistic-stimulus from the Sintel dataset that maximizes the cell type's central column response at second zero in the best-performing model of the respective cluster.

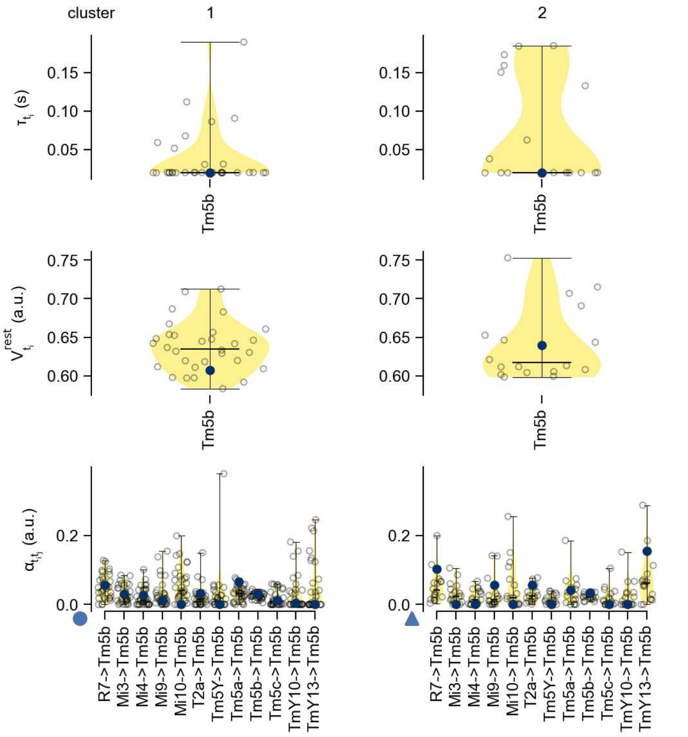

Tm5b - Figure 13: **Task-constrained parameters.** Each column shows the parameters inferred within the respective cluster. First row: learned time constants of the cell type. Second row: resting potentials of the cell type. Third row: scaling factors for the convolutional filters. The blue scatter represents the parameters from the best-performing model within the cluster.

# 51 Tm5c

← Cell types

#### Figures

1 Anatomical receptive fields. . . . 336
2 Anatomical projective fields. . . . 336
3 Clustering of the responses to naturalistic stimuli. . . . 337
4 Responses to flashes. . . . 337
5 Peak responses to moving edges. . . . 338
6 Peak responses to moving edges from best-performing models. . . . 338
7 Responses to moving edges from best-performing models. . . . 339
8 Peak responses to moving bars. . . . 339
9 Peak responses to moving bars from best-performing models. . . . 340
10 Responses to moving bars from best-performing models. . . . 340
11 Spatio-temporal receptive field. . . . 341
12 Maximally excitatory stimuli. . . . 341
13 Task-constrained parameters. . . . 342

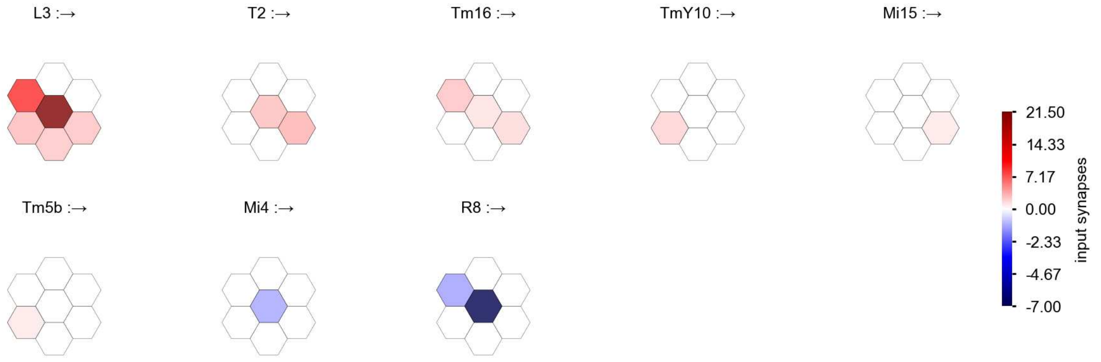

Tm5c - Figure 1: **Anatomical receptive fields.** Each colored hexagon is a projection, with the projection strength characterized by the average number of synapses that we count from the EM reconstruction. Red indicates excitatory synapses, blue indicates inhibitory synapses from inferred signs. Filters in the order of their total number of synapses.

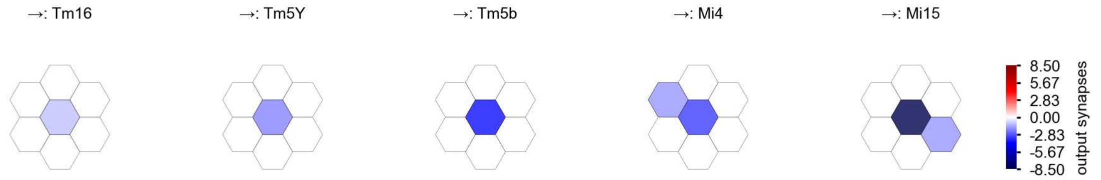

Tm5c - Figure 2: **Anatomical projective fields.** Each colored hexagon is a projection, with the projection strength characterized by the average number of synapses that we count from the EM reconstruction. Red indicates excitatory synapses, blue indicates inhibitory synapses from inferred signs. Filters in the order of their total number of synapses.

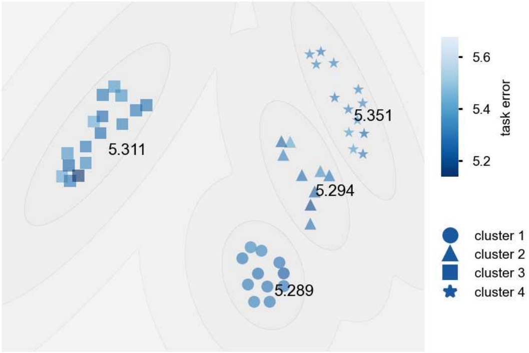

Tm5c - Figure 3: **Clustering of the responses to naturalistic stimuli.** From responses to naturalistic scenes from Sintel. Scatterpoints represent individual models colored by their task error.

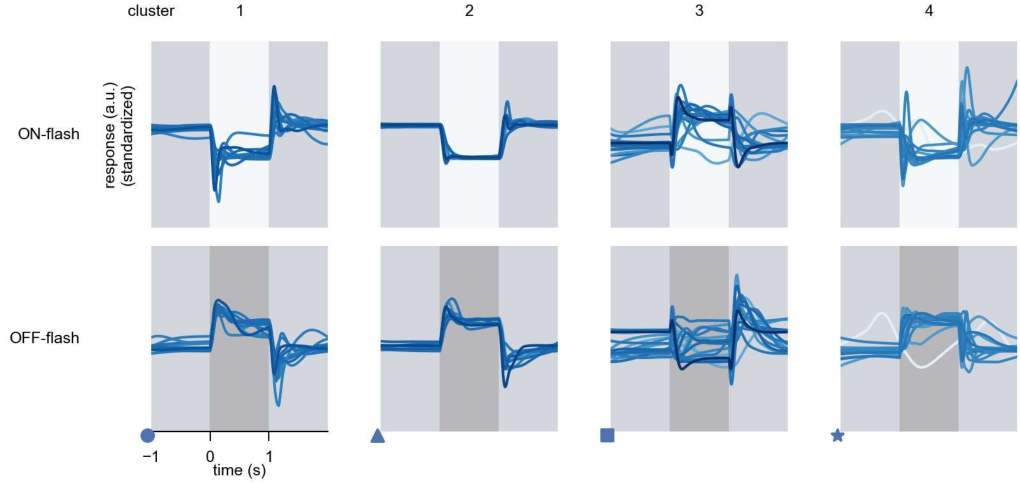

Tm5c - Figure 4: **Responses to flashes.** The top row shows responses to ON-flashes, the bottom row shows responses to OFF-flashes. The responses from the different models in the different clusters (columns) overlay and are colored in the respective task error of the model. The circular flashes occur at second zero, cover 6 ommatidia in radius, and last for one second. Before and after, a grey-stimulus leads to a stationary state of the network. The traces are standardized.

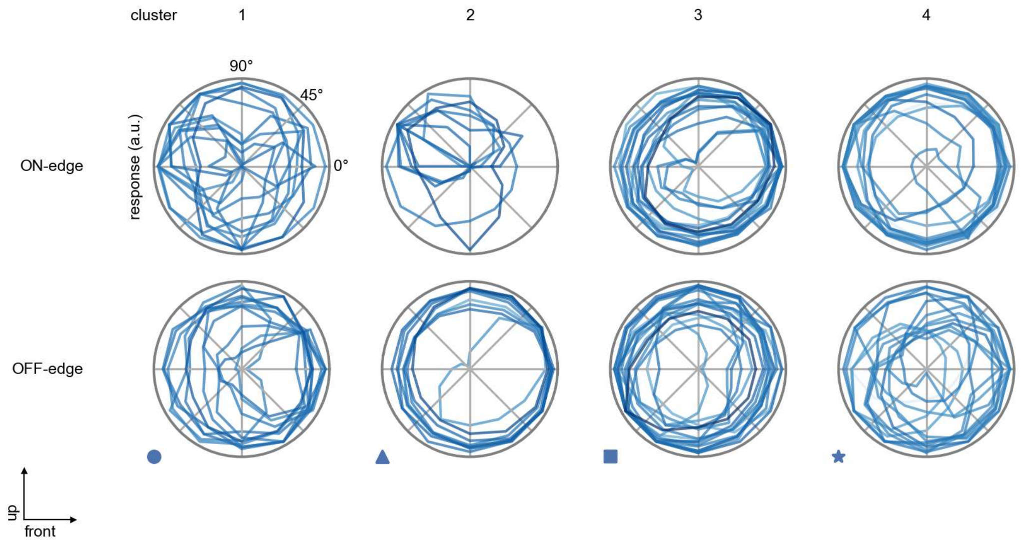

Tm5c - Figure 5: **Peak responses to moving edges.** The top row shows normalized peak responses to moving ON-edges, the bottom row shows normalized peak responses to moving OFF-edges. The edge-stimuli move in different directions from 0 to 360 degrees. Before, a grey-stimulus led the network to a stationary state. The responses from the different models in the different clusters (columns) overlay and are colored in the respective task error of the model. The peak responses are averaged over all assessed edge-speeds.

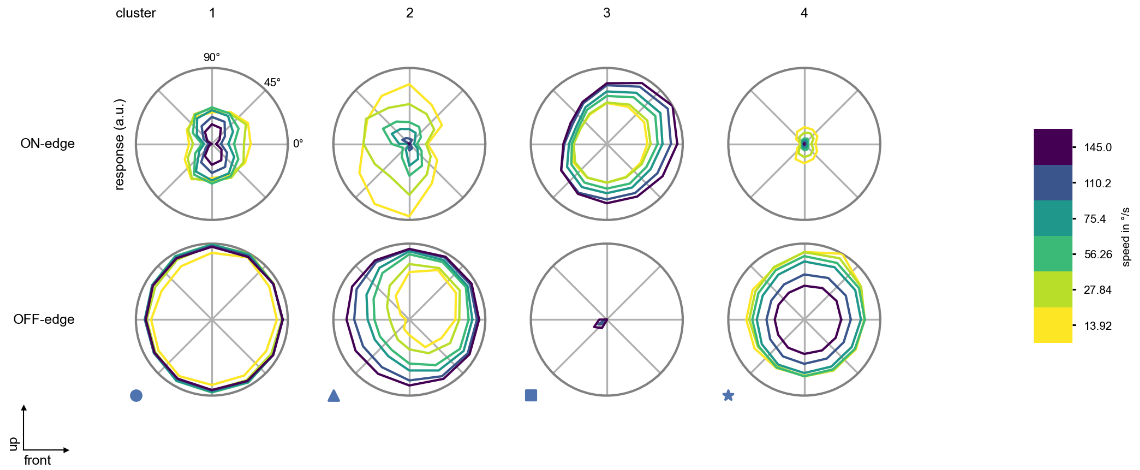

Tm5c - Figure 6: **Peak responses to moving edges from best-performing models.** The top row shows peak responses to moving ON-edges, the bottom row shows peak responses to moving OFF-edges. The edge-stimuli move in different directions from 0 to 360 degrees and at different speeds. Before, a grey-stimulus led the network to a stationary state. The columns show responses from the best-task-performing model in the respective cluster.

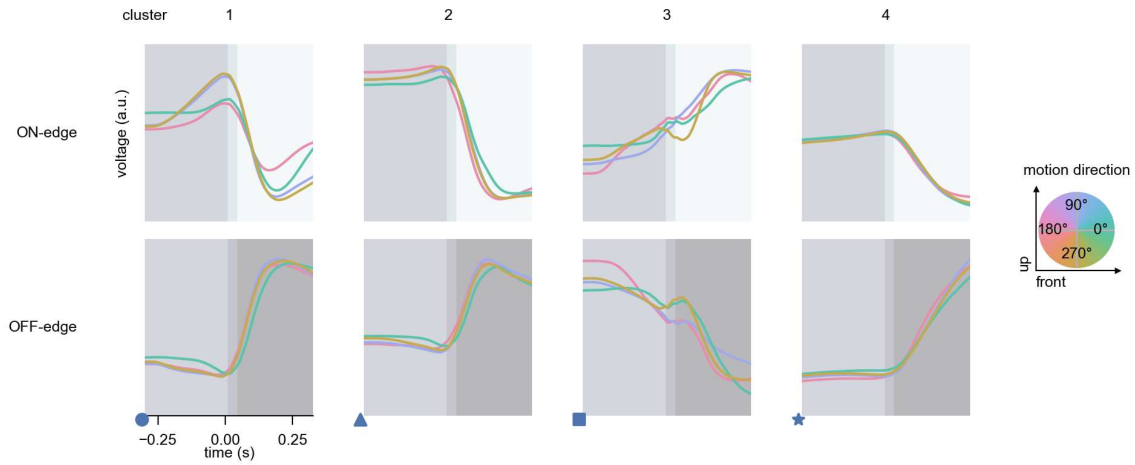

Tm5c - Figure 7: **Responses to moving edges from best-performing models.** The top row shows responses to moving ON-edges, the bottom row shows peak responses to moving OFF-edges. The edge-stimuli move in different directions from 0 to 360 degrees and at different speeds. Before, a grey-stimulus led the network to a stationary state. The columns show responses from the best-task-performing model in the respective cluster. Example responses to edges moving at 75.4°/s in all cardinal directions (green 0°, blue 90°, red 180°, yellow 270°) from -22.5 to 22.5° visual angle. Background contour indicates the edge, passing by the central column at second zero.

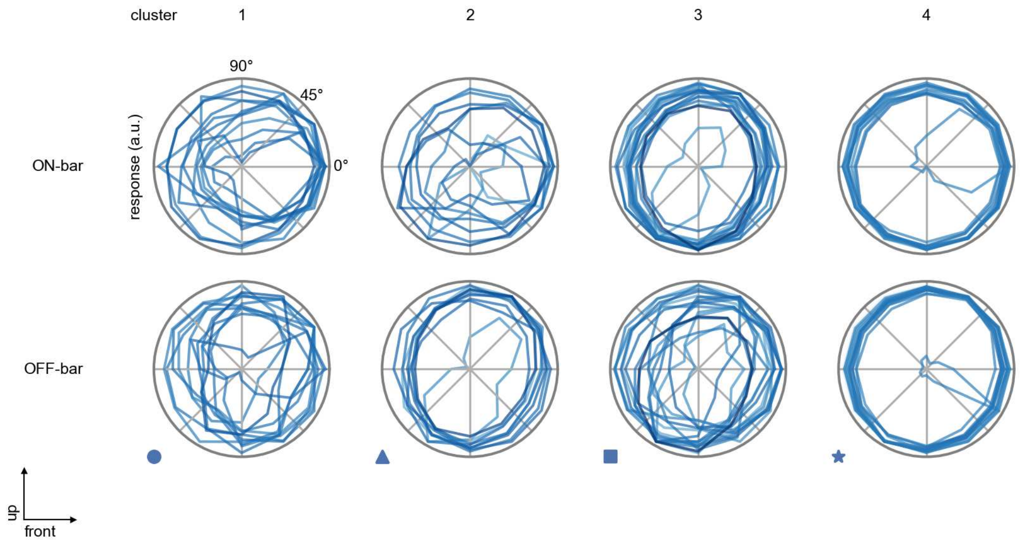

Tm5c - Figure 8: **Peak responses to moving bars.** The top row shows normalized peak responses to moving ON-bars, the bottom row shows normalized peak responses to moving OFF-bars. The bar-stimuli move in different directions from 0 to 360 degrees. Before, a grey-stimulus led the network to a stationary state. The responses from the different models in the different clusters (columns) overlay and are colored in the respective task error of the model. The peak responses are averaged over all assessed bar-speeds.

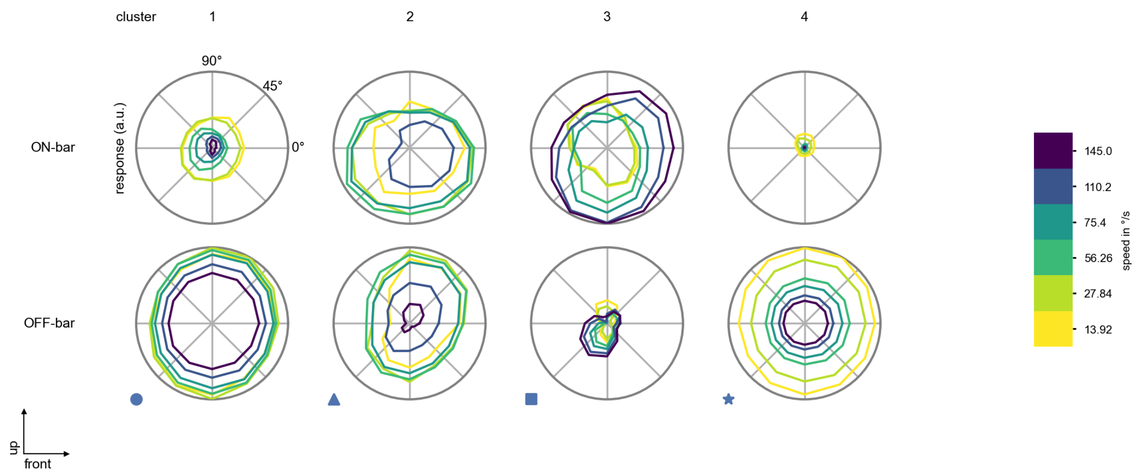

Tm5c - Figure 9: **Peak responses to moving bars from best-performing models.** The top row shows peak responses to moving ON-bars, the bottom row shows peak responses to moving OFF-bars. The bar-stimuli move in different directions from 0 to 360 degrees and at different speeds. Before, a grey-stimulus led the network to a stationary state. The columns show responses from the best-task-performing model in the respective cluster.

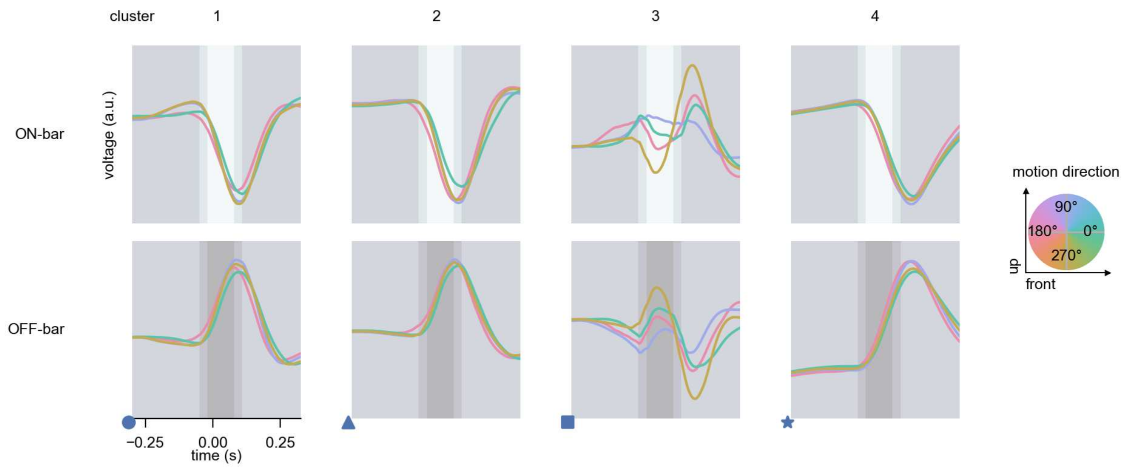

Tm5c - Figure 10: **Responses to moving bars from best-performing models.** The top row shows responses to moving ON-bars, the bottom row shows peak responses to moving OFF-bars. The bar-stimuli move in different directions from 0 to 360 degrees and at different speeds. Before, a grey-stimulus led the network to a stationary state. The columns show responses from the best-task-performing model in the respective cluster. Example responses to bars moving at 75.4°/s in all cardinal directions (green 0°, blue 90°, red 180°, yellow 270°) from -22.5 to 22.5° visual angle. Background contour indicates the bar, passing by the central column at second zero.

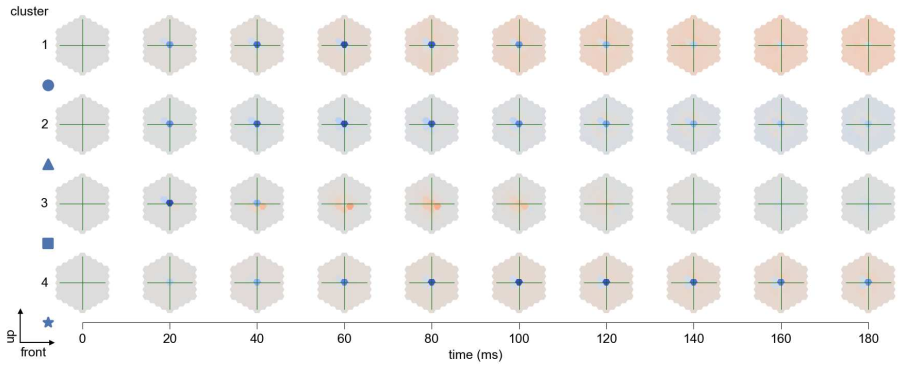

Tm5c - Figure 11: **Spatio-temporal receptive field.** Responses of the central cell to ON-impulses at single ommatidium impulse locations. The impulse occurs at second zero. Before, a grey-stimulus led the network to a stationary state. Responses are baseline-subtracted. Each row illustrates responses from the best-performing model of the respective cluster. Red indicated depolarization, blue indicates hyperpolarization.

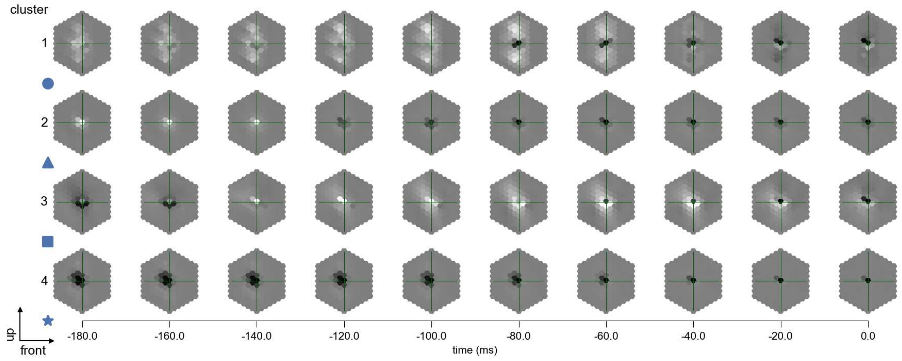

Tm5c - Figure 12: **Maximally excitatory stimuli.** Each row presents the regularized naturalistic-stimulus from the Sintel dataset that maximizes the cell type's central column response at second zero in the best-performing model of the respective cluster.

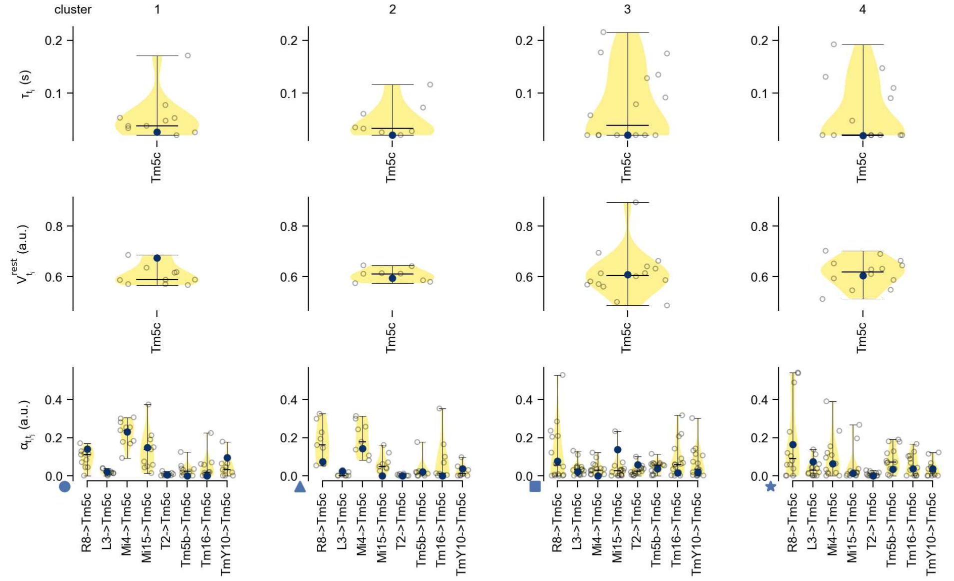

Tm5c - Figure 13: **Task-constrained parameters.** Each column shows the parameters inferred within the respective cluster. First row: learned time constants of the cell type. Second row: resting potentials of the cell type. Third row: scaling factors for the convolutional filters. The blue scatter represents the parameters from the best-performing model within the cluster.

# 52 Tm9

← Cell types

#### Figures

1 Anatomical receptive fields. . . . . 343
2 Anatomical projective fields. . . . . 344
3 Clustering of the responses to naturalistic stimuli. . . . . 344
4 Responses to flashes. . . . . 345
5 Peak responses to moving edges. . . . . 345
6 Peak responses to moving edges from best-performing models. . . . . 346
7 Responses to moving edges from best-performing models. . . . . 346
8 Peak responses to moving bars. . . . . 347
9 Peak responses to moving bars from best-performing models. . . . . 347
10 Responses to moving bars from best-performing models. . . . . 348
11 Spatio-temporal receptive field. . . . . 348
12 Maximally excitatory stimuli. . . . . 348
13 Task-constrained parameters. . . . . 349

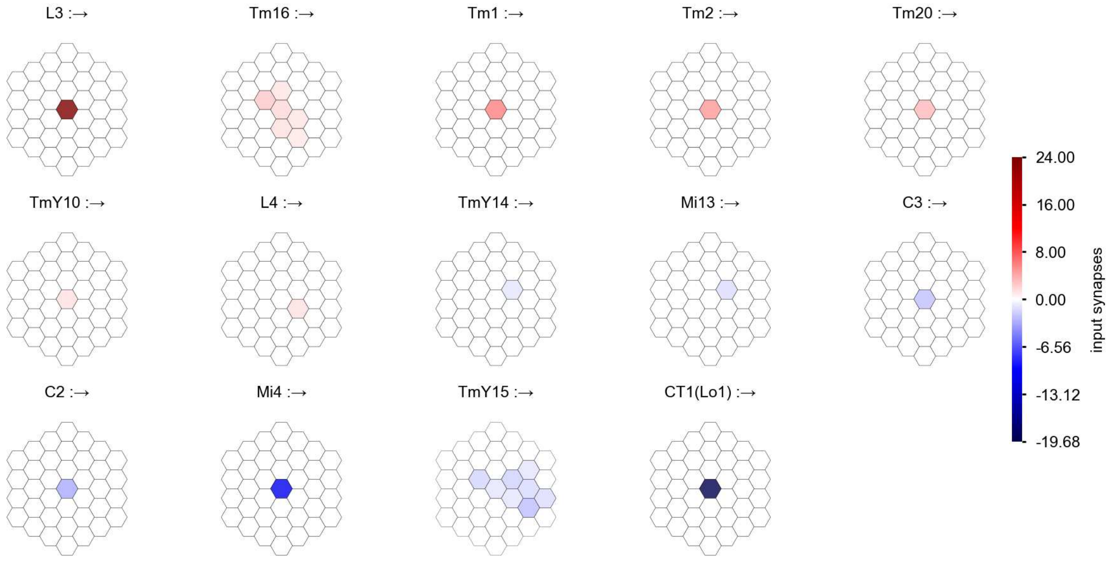

Tm9 - Figure 1: **Anatomical receptive fields.** Each colored hexagon is a projection, with the projection strength characterized by the average number of synapses that we count from the EM reconstruction. Red indicates excitatory synapses, blue indicates inhibitory synapses from inferred signs. Filters in the order of their total number of synapses.

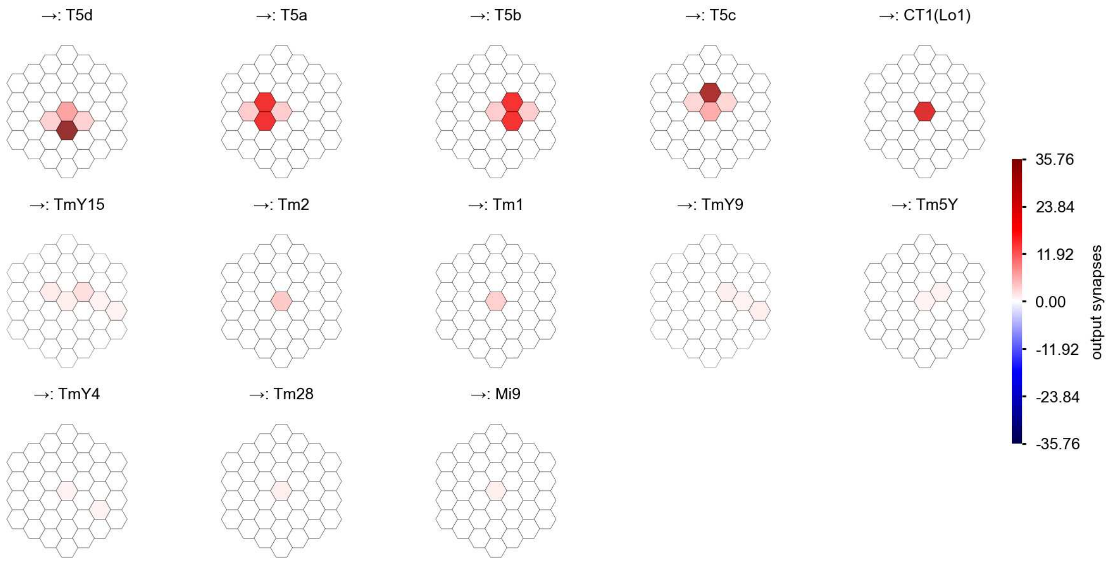

Tm9 - Figure 2: **Anatomical projective fields.** Each colored hexagon is a projection, with the projection strength characterized by the average number of synapses that we count from the EM reconstruction. Red indicates excitatory synapses, blue indicates inhibitory synapses from inferred signs. Filters in the order of their total number of synapses.

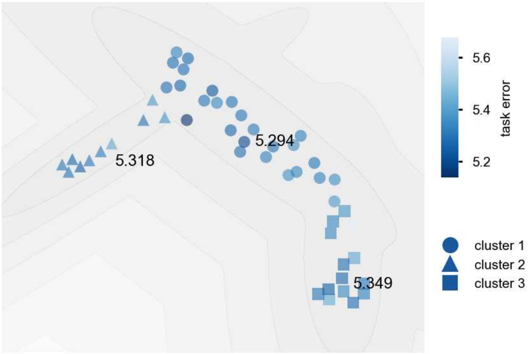

Tm9 - Figure 3: **Clustering of the responses to naturalistic stimuli.** From responses to naturalistic scenes from Sintel. Scatterpoints represent individual models colored by their task error.

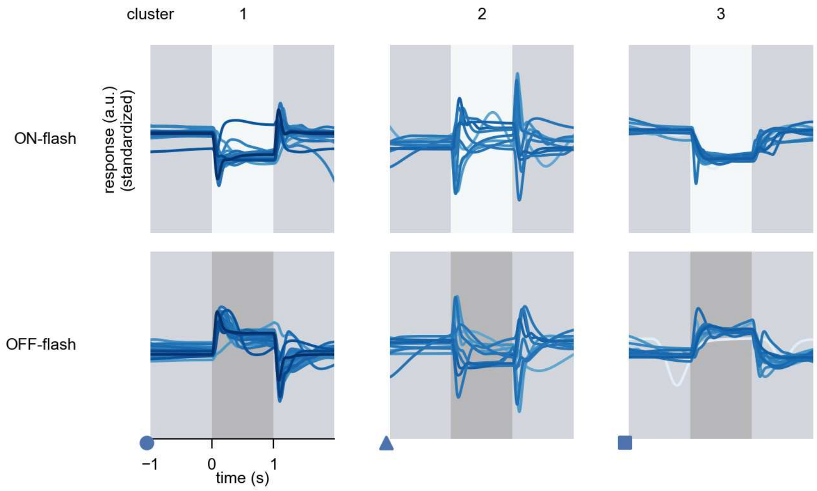

Tm9 - Figure 4: **Responses to flashes.** The top row shows responses to ON-flashes, the bottom row shows responses to OFF-flashes. The responses from the different models in the different clusters (columns) overlay and are colored in the respective task error of the model. The circular flashes occur at second zero, cover 6 ommatidia in radius, and last for one second. Before and after, a grey-stimulus leads to a stationary state of the network. The traces are standardized.

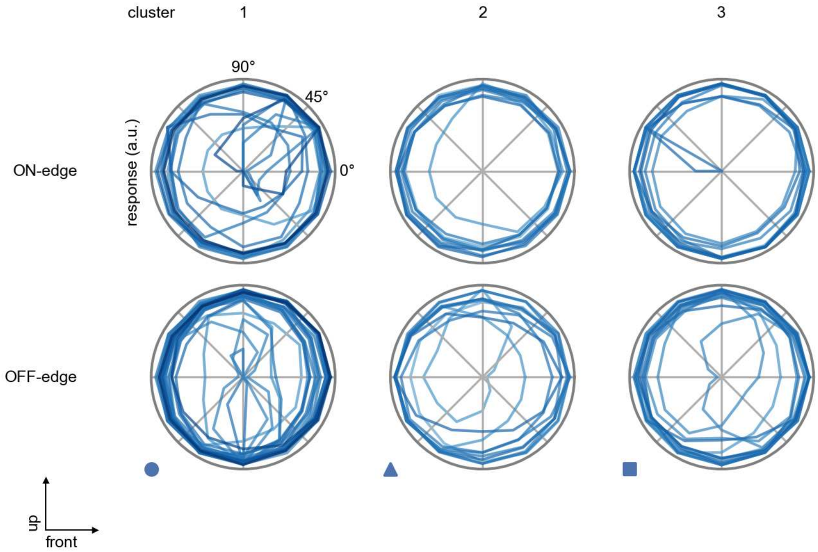

Tm9 - Figure 5: **Peak responses to moving edges.** The top row shows normalized peak responses to moving ON-edges, the bottom row shows normalized peak responses to moving OFF-edges. The edge-stimuli move in different directions from 0 to 360 degrees. Before, a grey-stimulus led the network to a stationary state. The responses from the different models in the different clusters (columns) overlay and are colored in the respective task error of the model. The peak responses are averaged over all assessed edge-speeds.

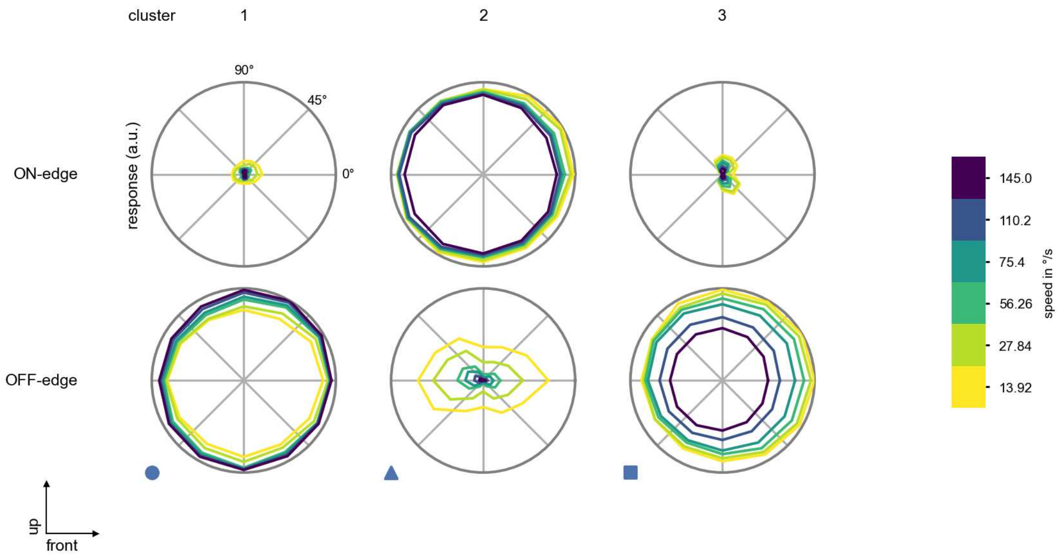

Tm9 - Figure 6: **Peak responses to moving edges from best-performing models.** The top row shows peak responses to moving ON-edges, the bottom row shows peak responses to moving OFF-edges. The edge-stimuli move in different directions from 0 to 360 degrees and at different speeds. Before, a grey-stimulus led the network to a stationary state. The columns show responses from the best-task-performing model in the respective cluster.

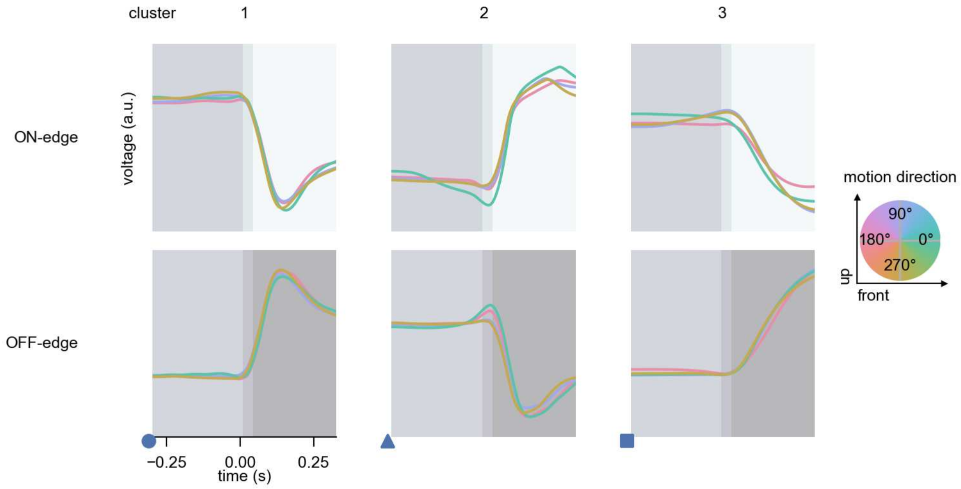

Tm9 - Figure 7: **Responses to moving edges from best-performing models.** The top row shows responses to moving ON-edges, the bottom row shows peak responses to moving OFF-edges. The edge-stimuli move in different directions from 0 to 360 degrees and at different speeds. Before, a grey-stimulus led the network to a stationary state. The columns show responses from the best-task-performing model in the respective cluster. Example responses to edges moving at 75.4°/s in all cardinal directions (green 0°, blue 90°, red 180°, yellow 270°) from -22.5 to 22.5° visual angle. Background contour indicates the edge, passing by the central column at second zero.

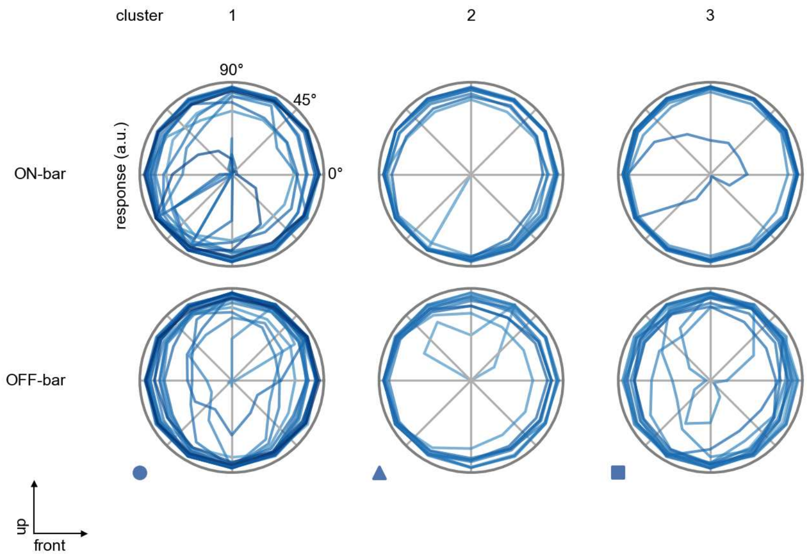

Tm9 - Figure 8: **Peak responses to moving bars.** The top row shows normalized peak responses to moving ON-bars, the bottom row shows normalized peak responses to moving OFF-bars. The bar-stimuli move in different directions from 0 to 360 degrees. Before, a grey-stimulus led the network to a stationary state. The responses from the different models in the different clusters (columns) overlay and are colored in the respective task error of the model. The peak responses are averaged over all assessed bar-speeds.

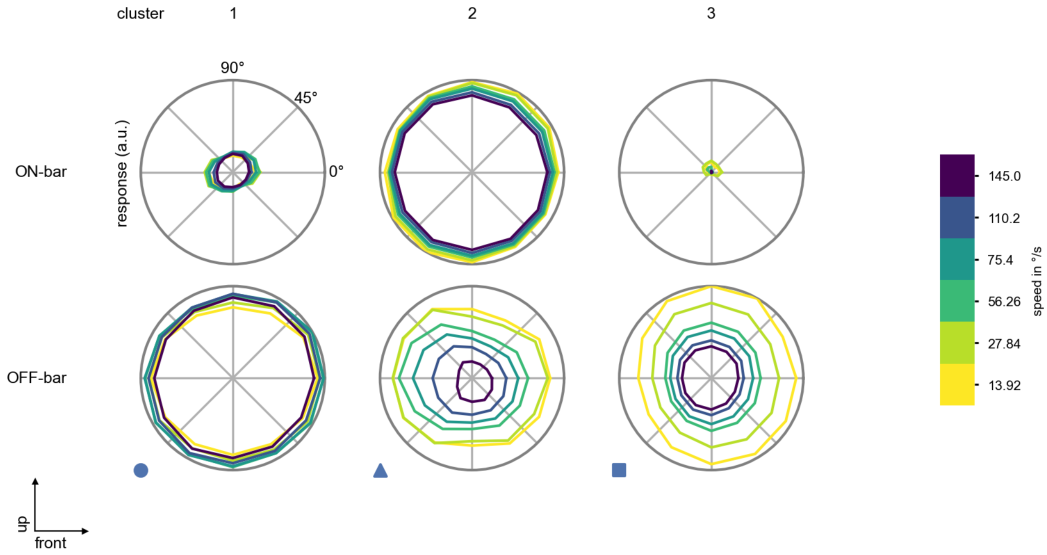

Tm9 - Figure 9: **Peak responses to moving bars from best-performing models.** The top row shows peak responses to moving ON-bars, the bottom row shows peak responses to moving OFF-bars. The bar-stimuli move in different directions from 0 to 360 degrees and at different speeds. Before, a grey-stimulus led the network to a stationary state. The columns show responses from the best-task-performing model in the respective cluster.

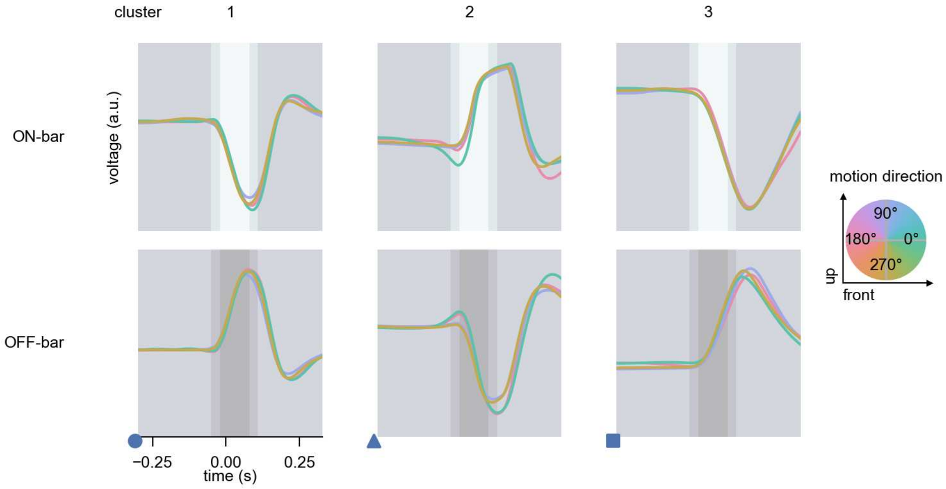

Tm9 - Figure 10: **Responses to moving bars from best-performing models.** The top row shows responses to moving ON-bars, the bottom row shows peak responses to moving OFF-bars. The bar-stimuli move in different directions from 0 to 360 degrees and at different speeds. Before, a grey-stimulus led the network to a stationary state. The columns show responses from the best-task-performing model in the respective cluster. Example responses to bars moving at 75.4°/s in all cardinal directions (green 0°, blue 90°, red 180°, yellow 270°) from -22.5 to 22.5° visual angle. Background contour indicates the bar, passing by the central column at second zero.

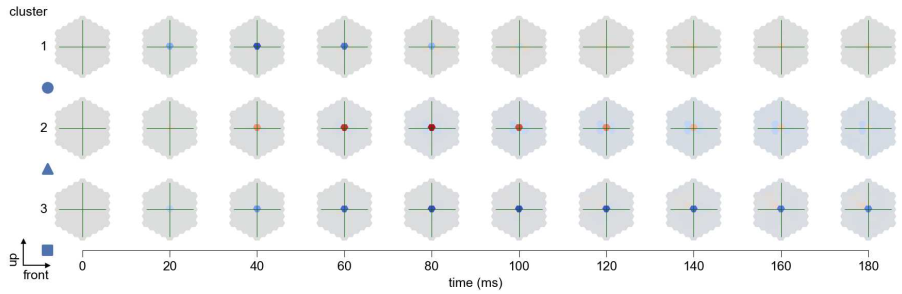

Tm9 - Figure 11: **Spatio-temporal receptive field.** Responses of the central cell to ON-impulses at single ommatidium impulse locations. The impulse occurs at second zero. Before, a grey-stimulus led the network to a stationary state. Responses are baseline-subtracted. Each row illustrates responses from the best-performing model of the respective cluster. Red indicated depolarization, blue indicates hyperpolarization.

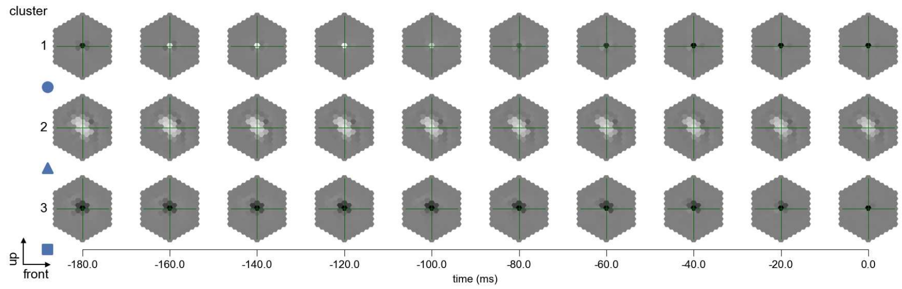

Tm9 - Figure 12: **Maximally excitatory stimuli.** Each row presents the regularized naturalistic-stimulus from the Sintel dataset that maximizes the cell type's central column response at second zero in the best-performing model of the respective cluster.

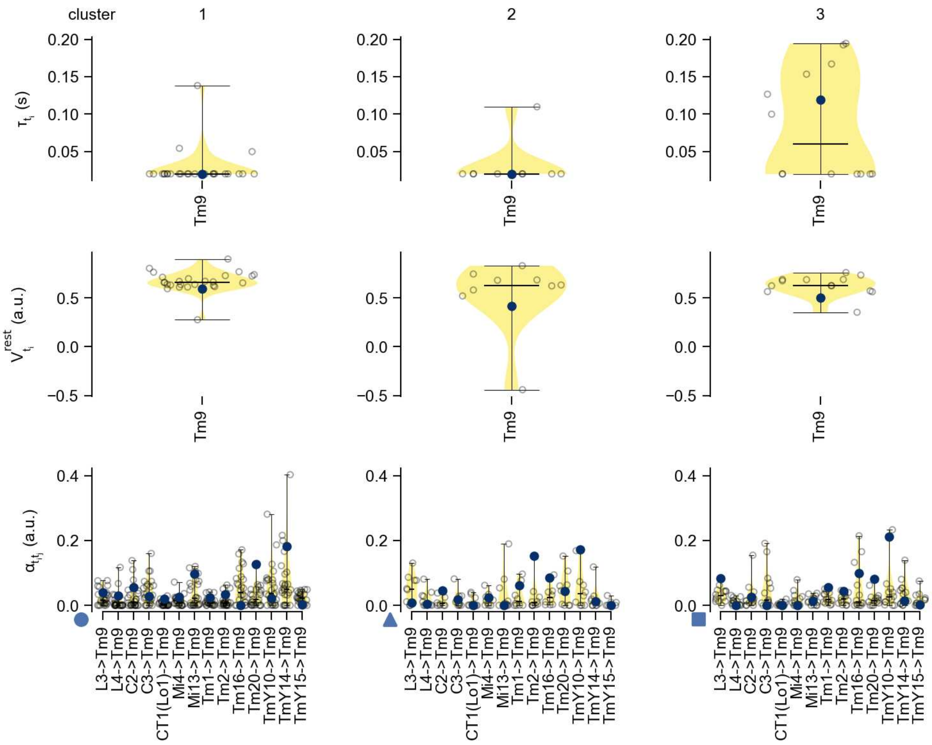

Tm9 - Figure 13: **Task-constrained parameters.** Each column shows the parameters inferred within the respective cluster. First row: learned time constants of the cell type. Second row: resting potentials of the cell type. Third row: scaling factors for the convolutional filters. The blue scatter represents the parameters from the best-performing model within the cluster.

# 53 Tm16

← Cell types

#### Figures

1 Anatomical receptive fields. . . . . 350
2 Anatomical projective fields. . . . . 351
3 Clustering of the responses to naturalistic stimuli. . . . . 351
4 Responses to flashes. . . . . 352
5 Peak responses to moving edges. . . . . 352
6 Peak responses to moving edges from best-performing models. . . . . 353
7 Responses to moving edges from best-performing models. . . . . 353
8 Peak responses to moving bars. . . . . 354
9 Peak responses to moving bars from best-performing models. . . . . 354
10 Responses to moving bars from best-performing models. . . . . 355
11 Spatio-temporal receptive field. . . . . 355
12 Maximally excitatory stimuli. . . . . 355
13 Task-constrained parameters. . . . . 356

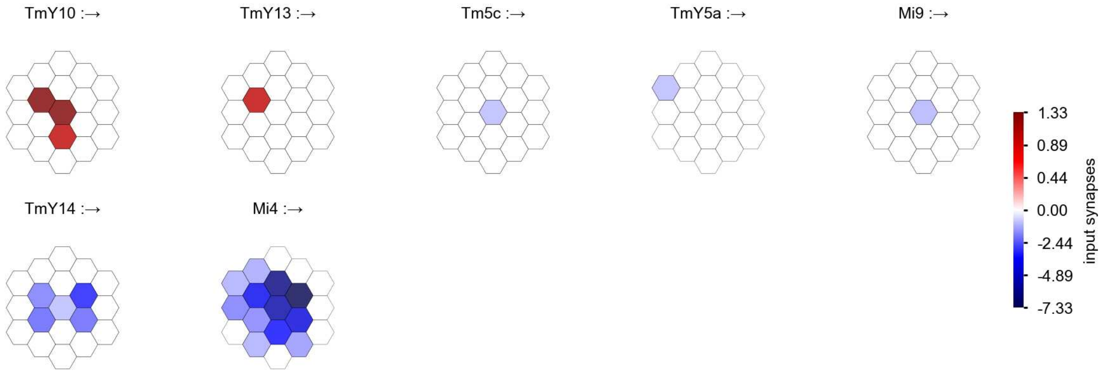

Tm16 - Figure 1: **Anatomical receptive fields.** Each colored hexagon is a projection, with the projection strength characterized by the average number of synapses that we count from the EM reconstruction. Red indicates excitatory synapses, blue indicates inhibitory synapses from inferred signs. Filters in the order of their total number of synapses.

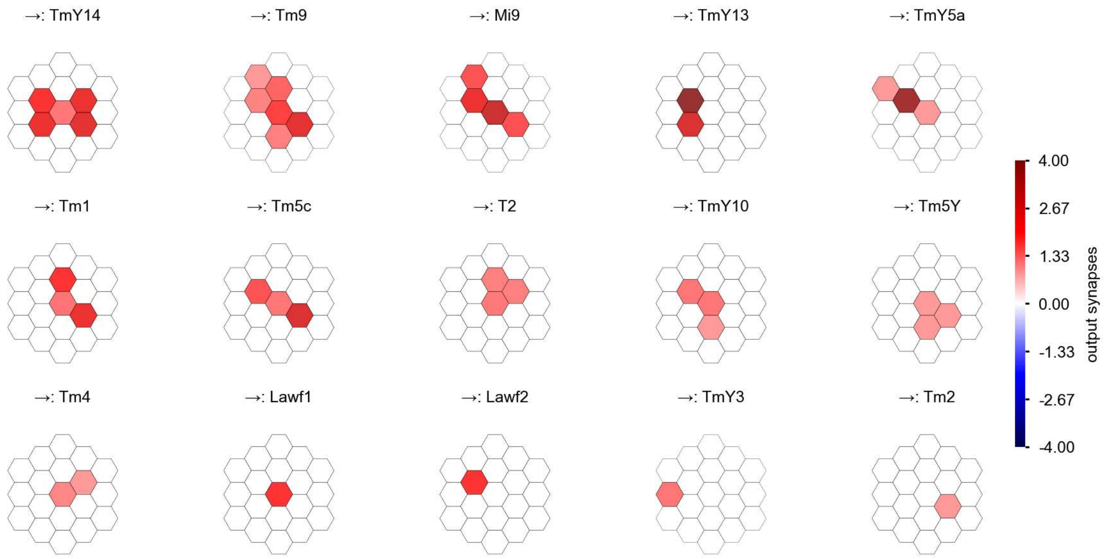

Tm16 - Figure 2: **Anatomical projective fields.** Each colored hexagon is a projection, with the projection strength characterized by the average number of synapses that we count from the EM reconstruction. Red indicates excitatory synapses, blue indicates inhibitory synapses from inferred signs. Filters in the order of their total number of synapses.

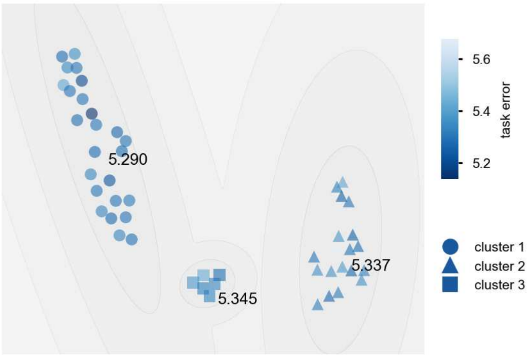

Tm16 - Figure 3: **Clustering of the responses to naturalistic stimuli.** From responses to naturalistic scenes from Sintel. Scatterpoints represent individual models colored by their task error.

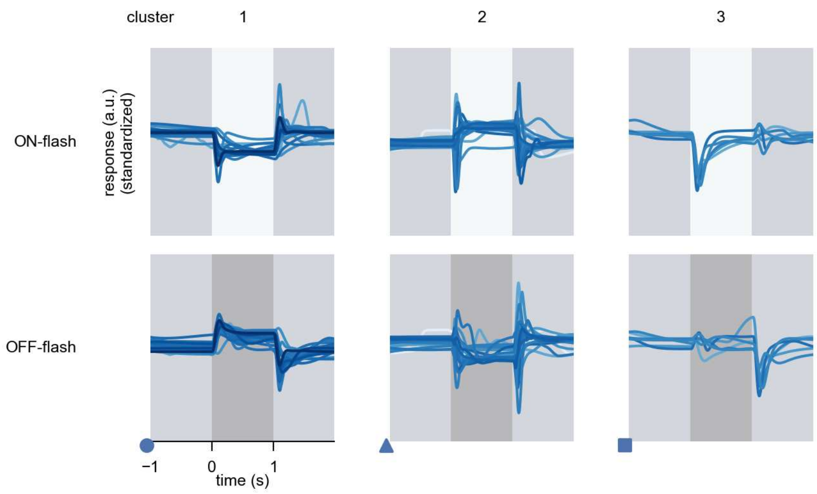

Tm16 - Figure 4: **Responses to flashes.** The top row shows responses to ON-flashes, the bottom row shows responses to OFF-flashes. The responses from the different models in the different clusters (columns) overlay and are colored in the respective task error of the model. The circular flashes occur at second zero, cover 6 ommatidia in radius, and last for one second. Before and after, a grey-stimulus leads to a stationary state of the network. The traces are standardized.

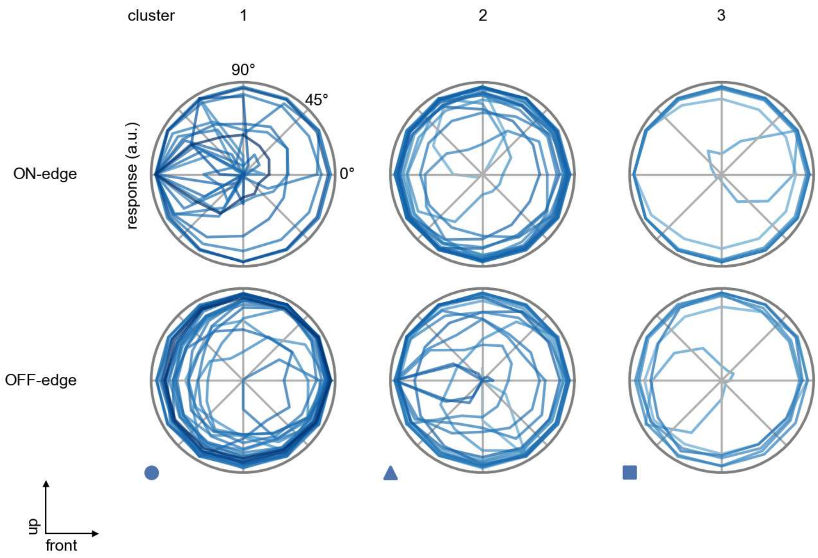

Tm16 - Figure 5: **Peak responses to moving edges.** The top row shows normalized peak responses to moving ON-edges, the bottom row shows normalized peak responses to moving OFF-edges. The edge-stimuli move in different directions from 0 to 360 degrees. Before, a grey-stimulus led the network to a stationary state. The responses from the different models in the different clusters (columns) overlay and are colored in the respective task error of the model. The peak responses are averaged over all assessed edge-speeds.

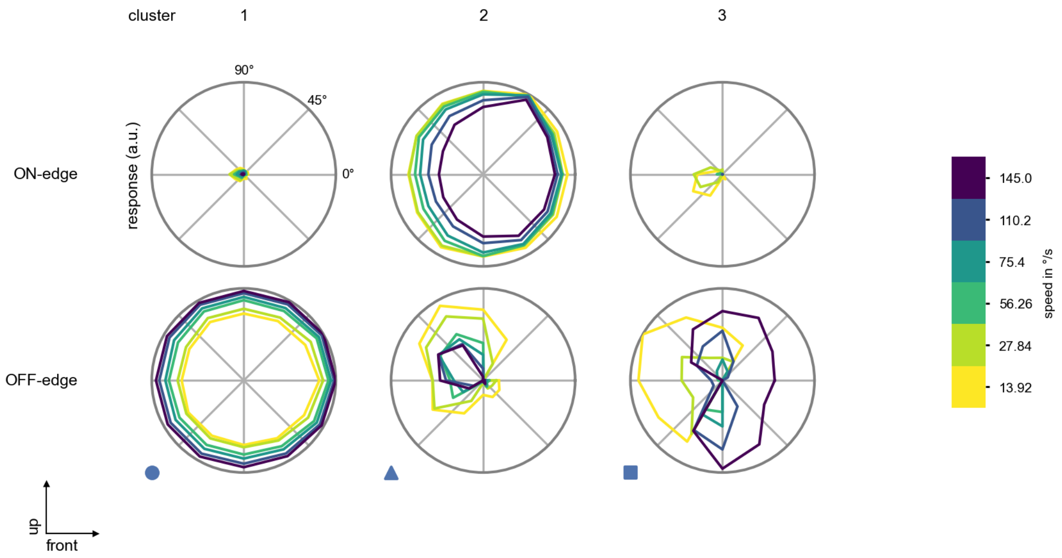

Tm16 - Figure 6: **Peak responses to moving edges from best-performing models.** The top row shows peak responses to moving ON-edges, the bottom row shows peak responses to moving OFF-edges. The edge-stimuli move in different directions from 0 to 360 degrees and at different speeds. Before, a grey-stimulus led the network to a stationary state. The columns show responses from the best-task-performing model in the respective cluster.

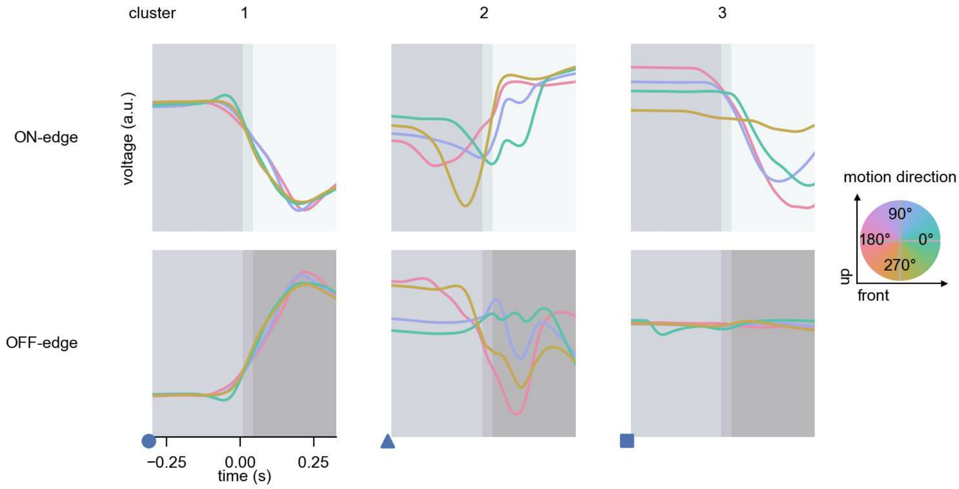

Tm16 - Figure 7: **Responses to moving edges from best-performing models.** The top row shows responses to moving ON-edges, the bottom row shows peak responses to moving OFF-edges. The edge-stimuli move in different directions from 0 to 360 degrees and at different speeds. Before, a grey-stimulus led the network to a stationary state. The columns show responses from the best-task-performing model in the respective cluster. Example responses to edges moving at 75.4°/s in all cardinal directions (green 0°, blue 90°, red 180°, yellow 270°) from -22.5 to 22.5° visual angle. Background contour indicates the edge, passing by the central column at second zero.

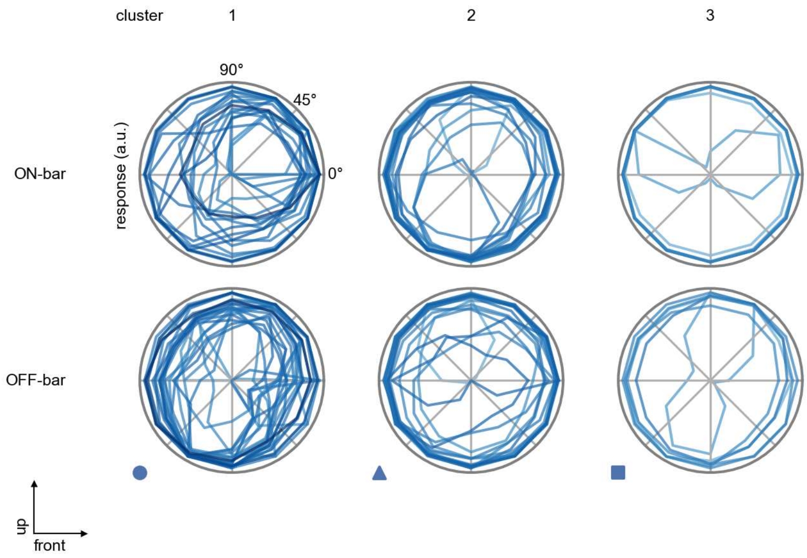

Tm16 - Figure 8: **Peak responses to moving bars.** The top row shows normalized peak responses to moving ON-bars, the bottom row shows normalized peak responses to moving OFF-bars. The bar-stimuli move in different directions from 0 to 360 degrees. Before, a grey-stimulus led the network to a stationary state. The responses from the different models in the different clusters (columns) overlay and are colored in the respective task error of the model. The peak responses are averaged over all assessed bar-speeds.

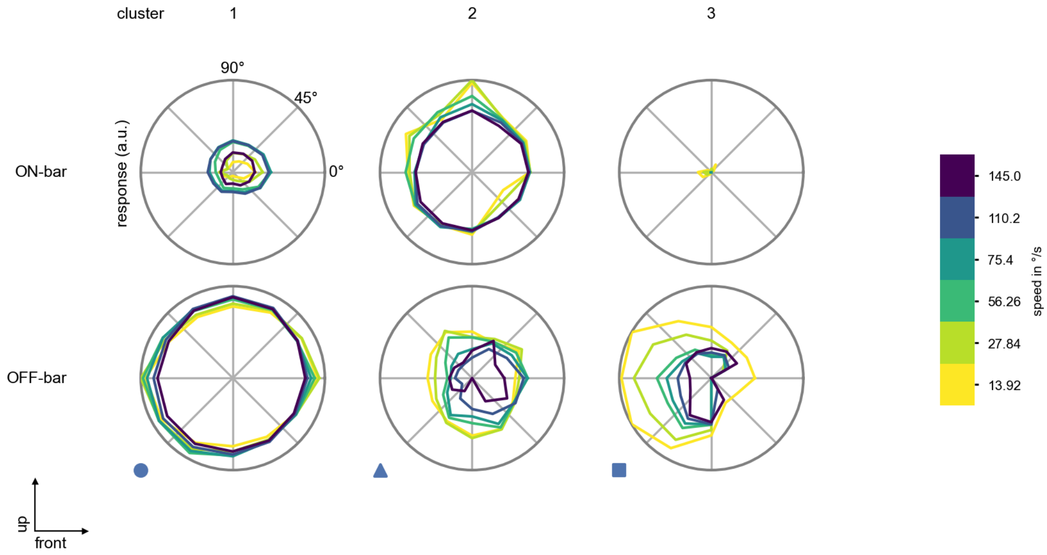

Tm16 - Figure 9: **Peak responses to moving bars from best-performing models.** The top row shows peak responses to moving ON-bars, the bottom row shows peak responses to moving OFF-bars. The bar-stimuli move in different directions from 0 to 360 degrees and at different speeds. Before, a grey-stimulus led the network to a stationary state. The columns show responses from the best-task-performing model in the respective cluster.

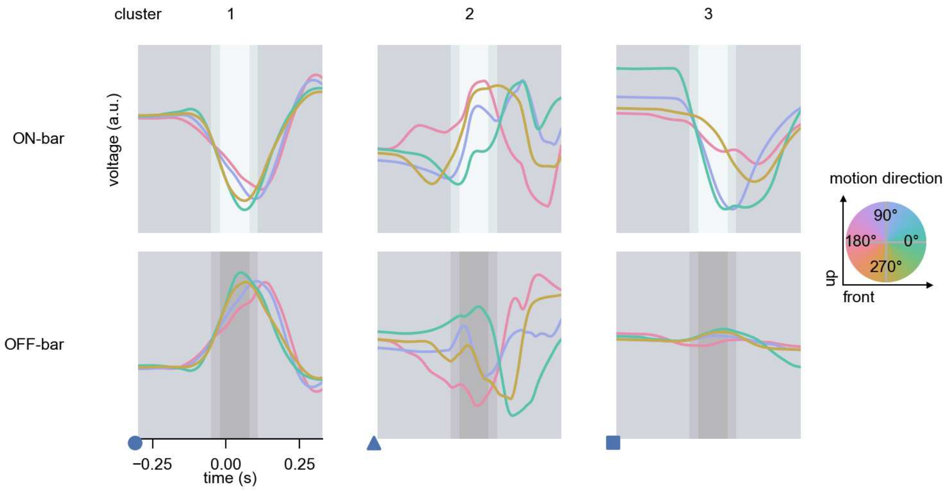

Tm16 - Figure 10: **Responses to moving bars from best-performing models.** The top row shows responses to moving ON-bars, the bottom row shows peak responses to moving OFF-bars. The bar-stimuli move in different directions from 0 to 360 degrees and at different speeds. Before, a grey-stimulus led the network to a stationary state. The columns show responses from the best-task-performing model in the respective cluster. Example responses to bars moving at 75.4°/s in all cardinal directions (green 0°, blue 90°, red 180°, yellow 270°) from -22.5 to 22.5° visual angle. Background contour indicates the bar, passing by the central column at second zero.

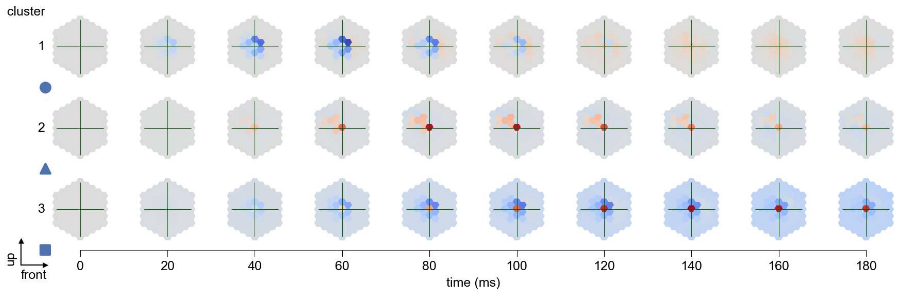

Tm16 - Figure 11: **Spatio-temporal receptive field.** Responses of the central cell to ON-impulses at single ommatidium impulse locations. The impulse occurs at second zero. Before, a grey-stimulus led the network to a stationary state. Responses are baseline-subtracted. Each row illustrates responses from the best-performing model of the respective cluster. Red indicated depolarization, blue indicates hyperpolarization.

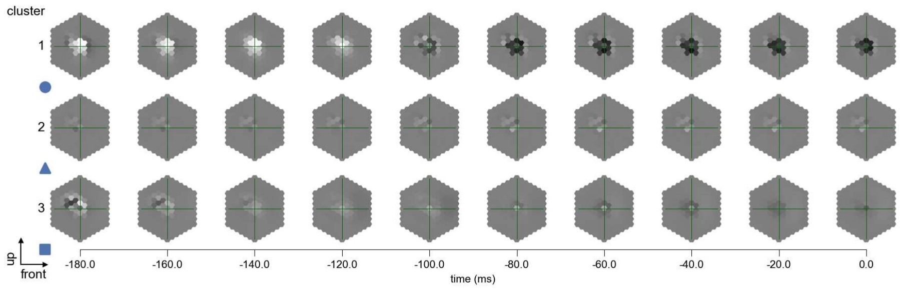

Tm16 - Figure 12: **Maximally excitatory stimuli.** Each row presents the regularized naturalistic-stimulus from the Sintel dataset that maximizes the cell type's central column response at second zero in the best-performing model of the respective cluster.

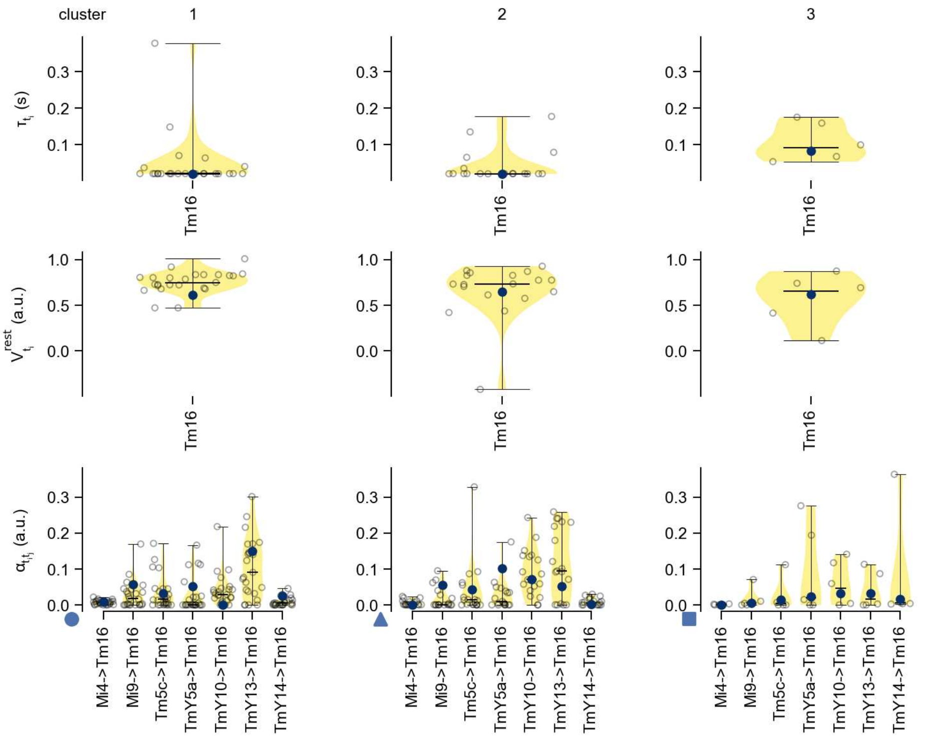

Tm16 - Figure 13: **Task-constrained parameters.** Each column shows the parameters inferred within the respective cluster. First row: learned time constants of the cell type. Second row: resting potentials of the cell type. Third row: scaling factors for the convolutional filters. The blue scatter represents the parameters from the best-performing model within the cluster.

# 54 Tm20

← Cell types

#### Figures

1 Anatomical receptive fields. . . . . 357
2 Anatomical projective fields. . . . . 357
3 Clustering of the responses to naturalistic stimuli. . . . . 358
4 Responses to flashes. . . . . 358
5 Peak responses to moving edges. . . . . 358
6 Peak responses to moving edges from best-performing models. . . . . 359
7 Responses to moving edges from best-performing models. . . . . 359
8 Peak responses to moving bars. . . . . 360
9 Peak responses to moving bars from best-performing models. . . . . 360
10 Responses to moving bars from best-performing models. . . . . 361
11 Spatio-temporal receptive field. . . . . 361
12 Maximally excitatory stimuli. . . . . 362
13 Task-constrained parameters. . . . . 362

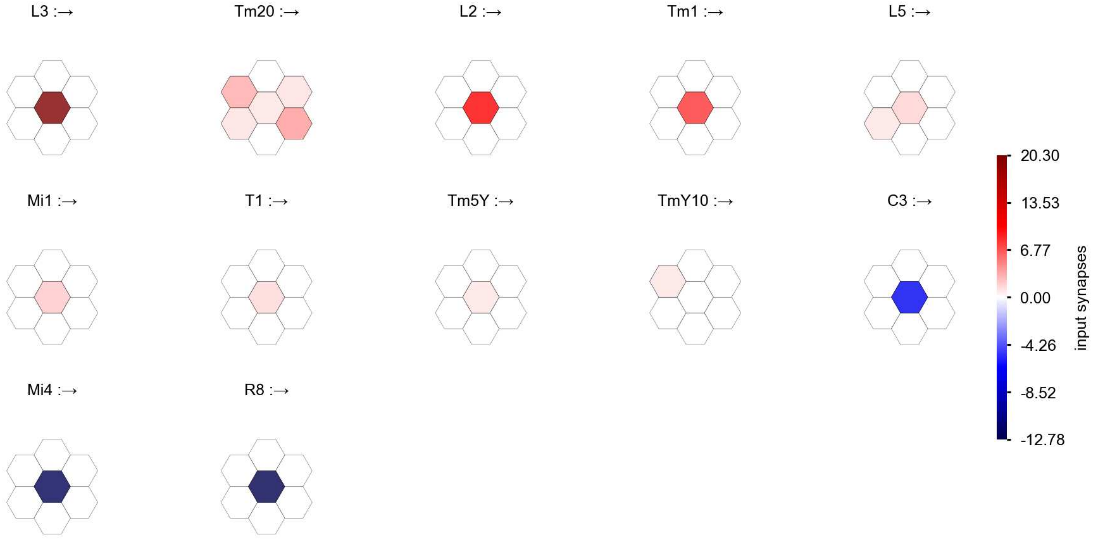

Tm20 - Figure 1: **Anatomical receptive fields.** Each colored hexagon is a projection, with the projection strength characterized by the average number of synapses that we count from the EM reconstruction. Red indicates excitatory synapses, blue indicates inhibitory synapses from inferred signs. Filters in the order of their total number of synapses.

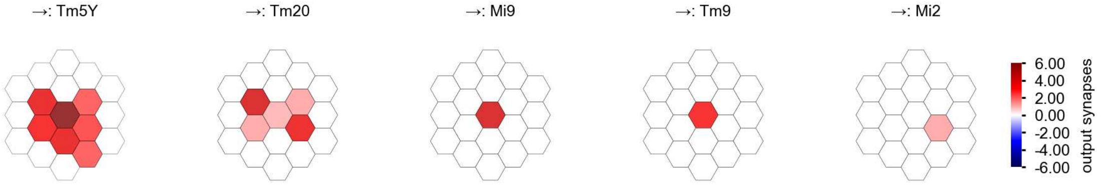

Tm20 - Figure 2: **Anatomical projective fields.** Each colored hexagon is a projection, with the projection strength characterized by the average number of synapses that we count from the EM reconstruction. Red indicates excitatory synapses, blue indicates inhibitory synapses from inferred signs. Filters in the order of their total number of synapses.

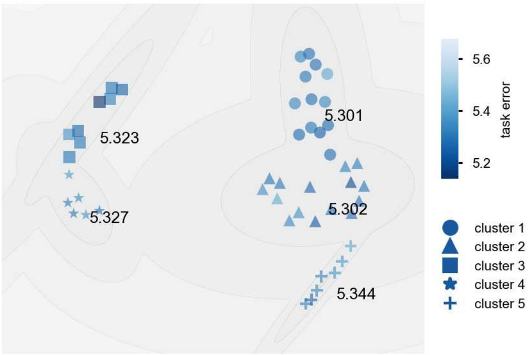

Tm20 - Figure 3: **Clustering of the responses to naturalistic stimuli.** From responses to naturalistic scenes from Sintel. Scatterpoints represent individual models colored by their task error.

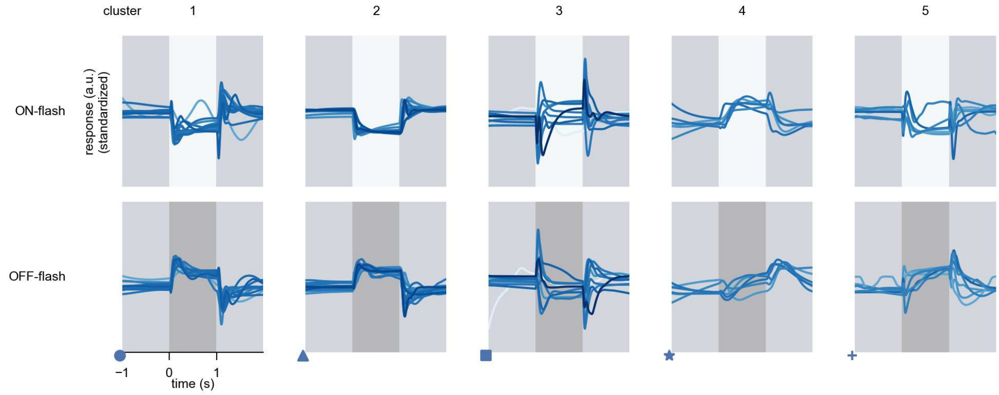

Tm20 - Figure 4: **Responses to flashes.** The top row shows responses to ON-flashes, the bottom row shows responses to OFF-flashes. The responses from the different models in the different clusters (columns) overlay and are colored in the respective task error of the model. The circular flashes occur at second zero, cover 6 ommatidia in radius, and last for one second. Before and after, a grey-stimulus leads to a stationary state of the network. The traces are standardized.

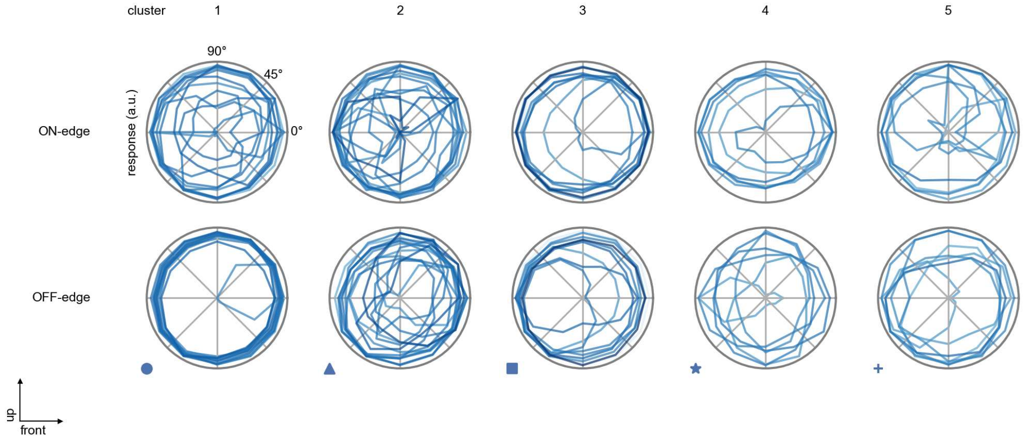

Tm20 - Figure 5: **Peak responses to moving edges.** The top row shows normalized peak responses to moving ON-edges, the bottom row shows normalized peak responses to moving OFF-edges. The edge-stimuli move in different directions from 0 to 360 degrees. Before, a grey-stimulus led the network to a stationary state. The responses from the different models in the different clusters (columns) overlay and are colored in the respective task error of the model. The peak responses are averaged over all assessed edge-speeds.

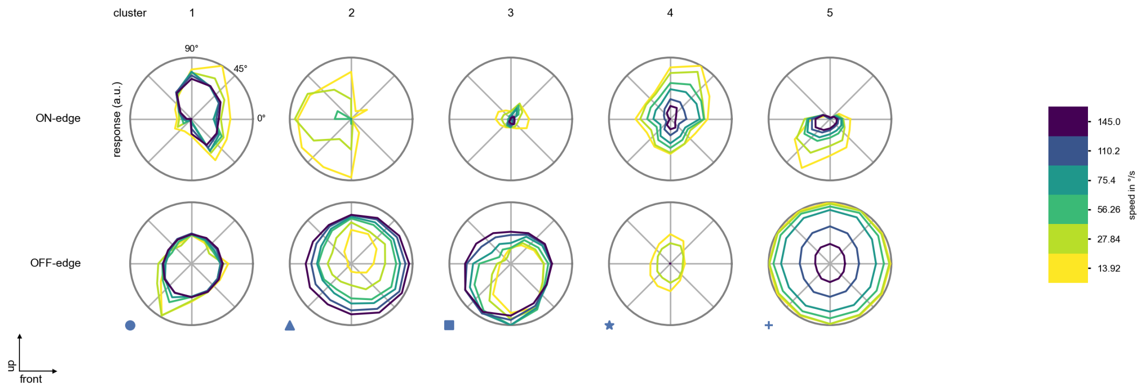

Tm20 - Figure 6: **Peak responses to moving edges from best-performing models.** The top row shows peak responses to moving ON-edges, the bottom row shows peak responses to moving OFF-edges. The edge-stimuli move in different directions from 0 to 360 degrees and at different speeds. Before, a grey-stimulus led the network to a stationary state. The columns show responses from the best-task-performing model in the respective cluster.

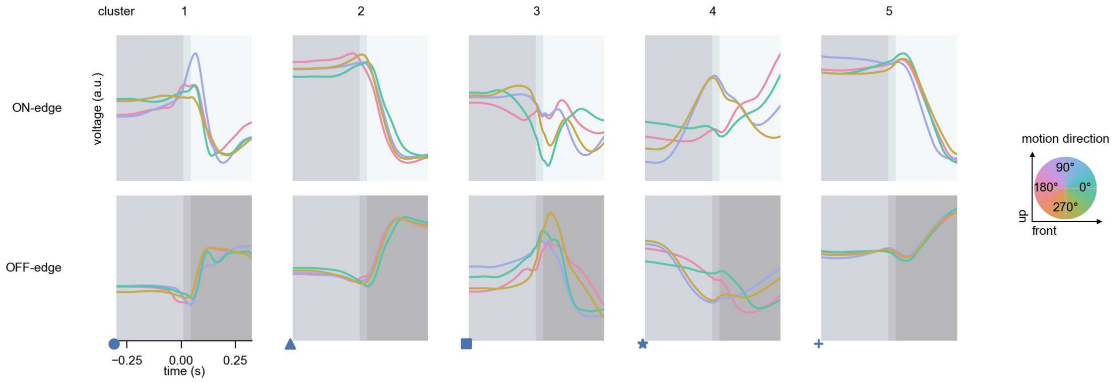

Tm20 - Figure 7: **Responses to moving edges from best-performing models.** The top row shows responses to moving ON-edges, the bottom row shows peak responses to moving OFF-edges. The edge-stimuli move in different directions from 0 to 360 degrees and at different speeds. Before, a grey-stimulus led the network to a stationary state. The columns show responses from the best-task-performing model in the respective cluster. Example responses to edges moving at 75.4°/s in all cardinal directions (green 0°, blue 90°, red 180°, yellow 270°) from -22.5 to 22.5° visual angle. Background contour indicates the edge, passing by the central column at second zero.

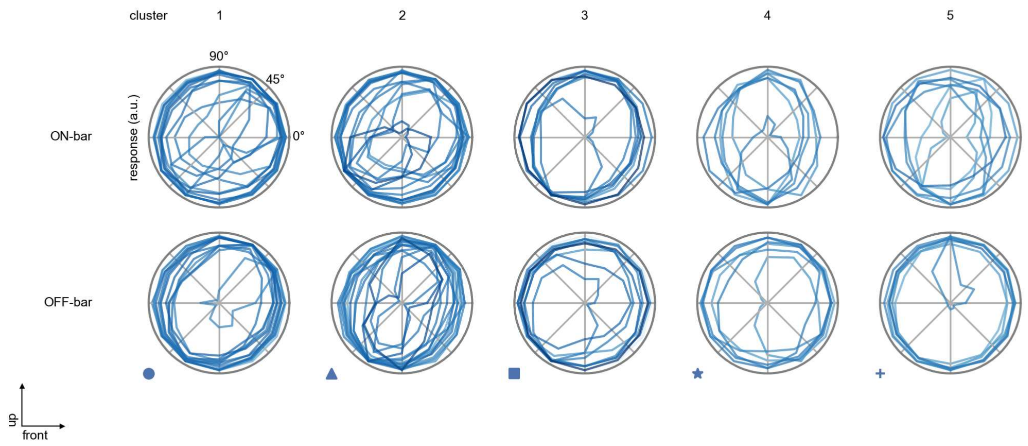

Tm20 - Figure 8: **Peak responses to moving bars.** The top row shows normalized peak responses to moving ON-bars, the bottom row shows normalized peak responses to moving OFF-bars. The bar-stimuli move in different directions from 0 to 360 degrees. Before, a grey-stimulus led the network to a stationary state. The responses from the different models in the different clusters (columns) overlay and are colored in the respective task error of the model. The peak responses are averaged over all assessed bar-speeds.

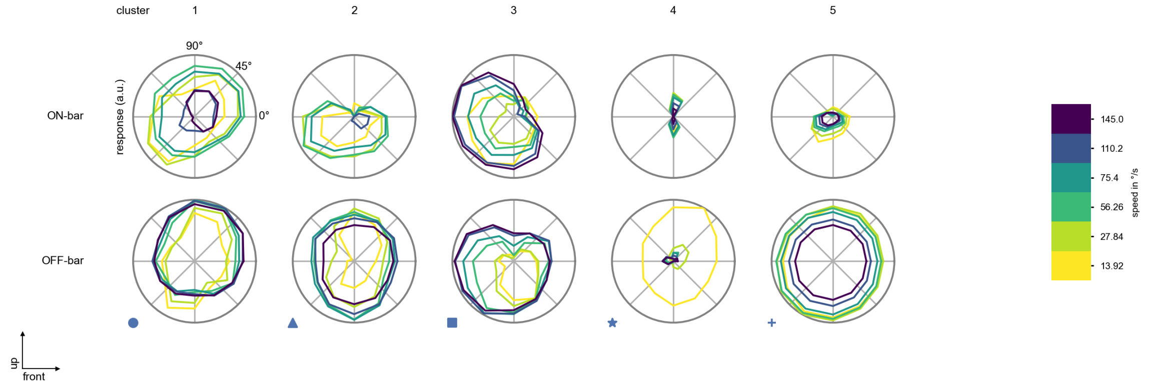

Tm20 - Figure 9: **Peak responses to moving bars from best-performing models.** The top row shows peak responses to moving ON-bars, the bottom row shows peak responses to moving OFF-bars. The bar-stimuli move in different directions from 0 to 360 degrees and at different speeds. Before, a grey-stimulus led the network to a stationary state. The columns show responses from the best-task-performing model in the respective cluster.

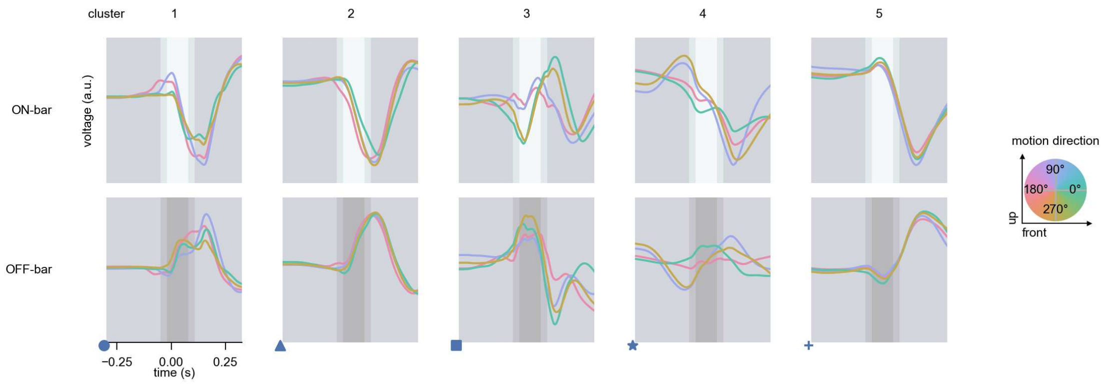

Tm20 - Figure 10: **Responses to moving bars from best-performing models.** The top row shows responses to moving ON-bars, the bottom row shows peak responses to moving OFF-bars. The bar-stimuli move in different directions from 0 to 360 degrees and at different speeds. Before, a grey-stimulus led the network to a stationary state. The columns show responses from the best-task-performing model in the respective cluster. Example responses to bars moving at 75.4°/s in all cardinal directions (green 0°, blue 90°, red 180°, yellow 270°) from -22.5 to 22.5° visual angle. Background contour indicates the bar, passing by the central column at second zero.

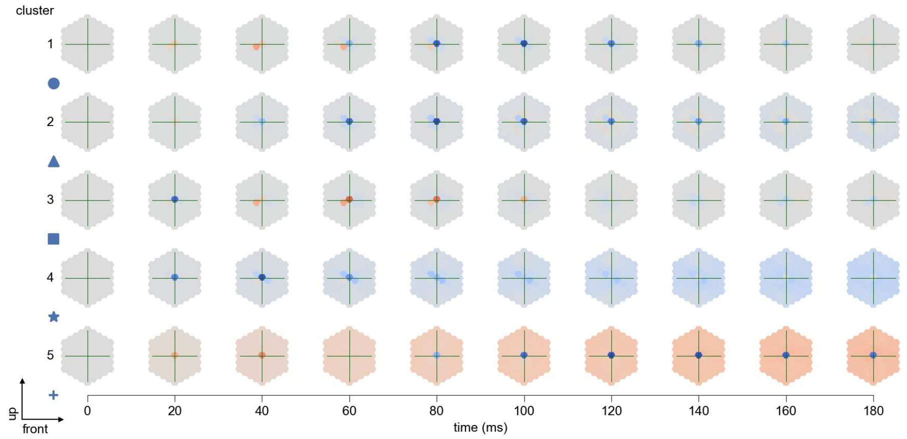

Tm20 - Figure 11: **Spatio-temporal receptive field.** Responses of the central cell to ON-impulses at single ommatidium impulse locations. The impulse occurs at second zero. Before, a grey-stimulus led the network to a stationary state. Responses are baseline-subtracted. Each row illustrates responses from the best-performing model of the respective cluster. Red indicated depolarization, blue indicates hyperpolarization.

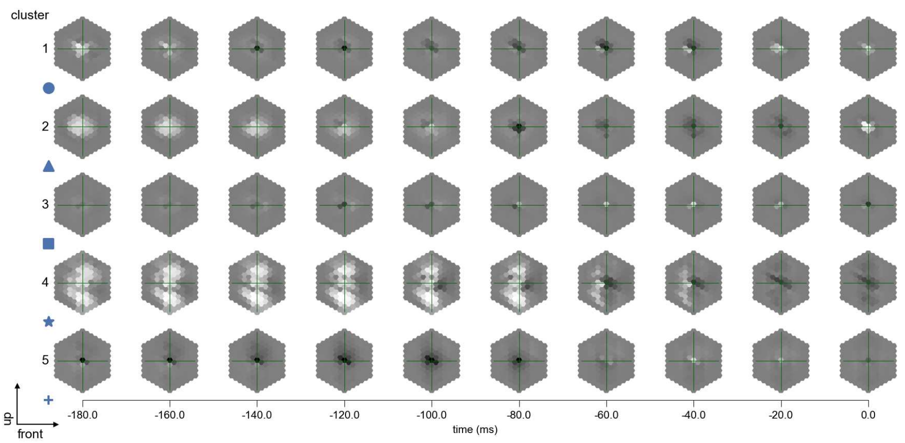

Tm20 - Figure 12: **Maximally excitatory stimuli.** Each row presents the regularized naturalistic-stimulus from the Sintel dataset that maximizes the cell type's central column response at second zero in the best-performing model of the respective cluster.

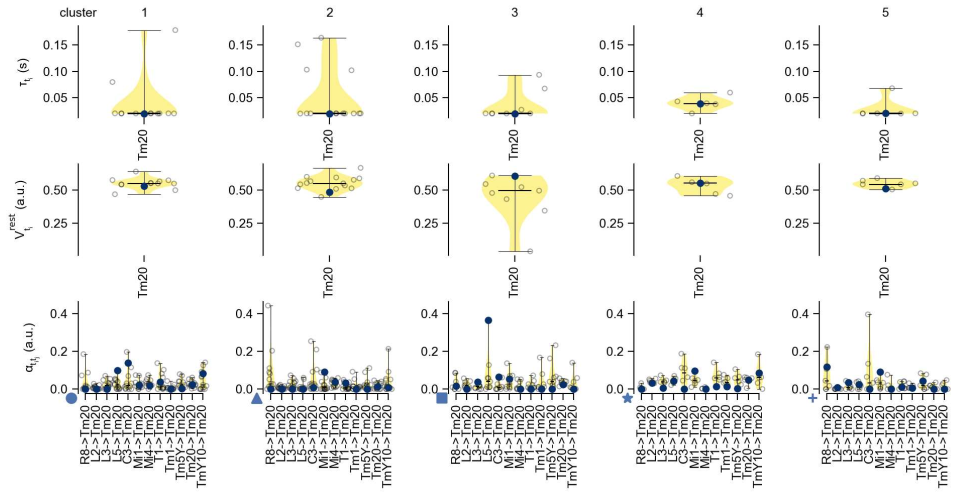

Tm20 - Figure 13: **Task-constrained parameters.** Each column shows the parameters inferred within the respective cluster. First row: learned time constants of the cell type. Second row: resting potentials of the cell type. Third row: scaling factors for the convolutional filters. The blue scatter represents the parameters from the best-performing model within the cluster.

# 55 Tm28

← Cell types

#### Figures

1 Anatomical receptive fields. . . . . 363
2 Anatomical projective fields. . . . . 363
3 Clustering of the responses to naturalistic stimuli. . . . . 364
4 Responses to flashes. . . . . 364
5 Peak responses to moving edges. . . . . 365
6 Peak responses to moving edges from best-performing models. . . . . 365
7 Responses to moving edges from best-performing models. . . . . 366
8 Peak responses to moving bars. . . . . 366
9 Peak responses to moving bars from best-performing models. . . . . 367
10 Responses to moving bars from best-performing models. . . . . 367
11 Spatio-temporal receptive field. . . . . 368
12 Maximally excitatory stimuli. . . . . 368
13 Task-constrained parameters. . . . . 369

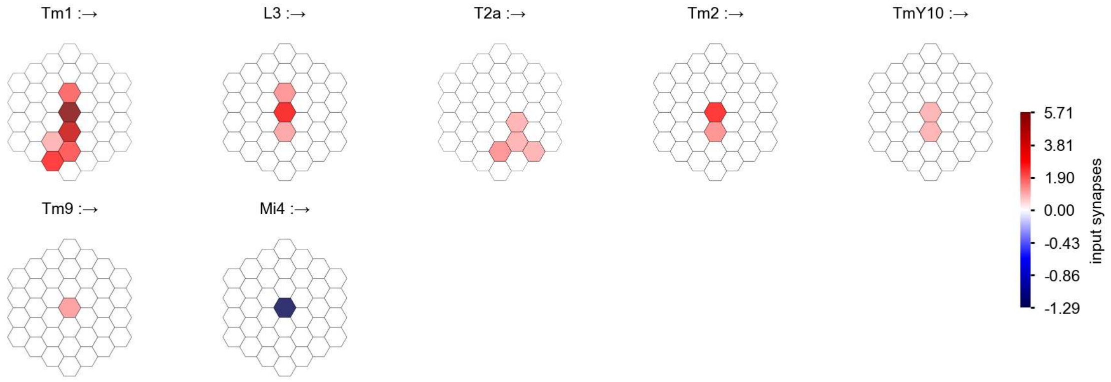

Tm28 - Figure 1: **Anatomical receptive fields.** Each colored hexagon is a projection, with the projection strength characterized by the average number of synapses that we count from the EM reconstruction. Red indicates excitatory synapses, blue indicates inhibitory synapses from inferred signs. Filters in the order of their total number of synapses.

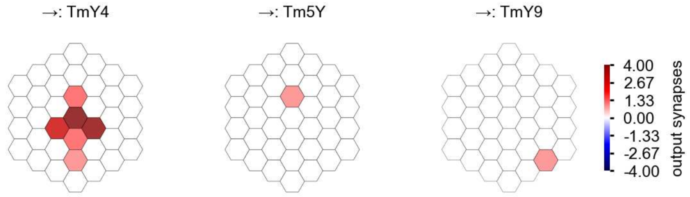

Tm28 - Figure 2: **Anatomical projective fields.** Each colored hexagon is a projection, with the projection strength characterized by the average number of synapses that we count from the EM reconstruction. Red indicates excitatory synapses, blue indicates inhibitory synapses from inferred signs. Filters in the order of their total number of synapses.

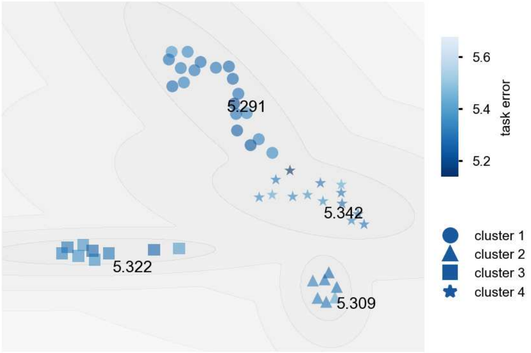

Tm28 - Figure 3: **Clustering of the responses to naturalistic stimuli.** From responses to naturalistic scenes from Sintel. Scatterpoints represent individual models colored by their task error.

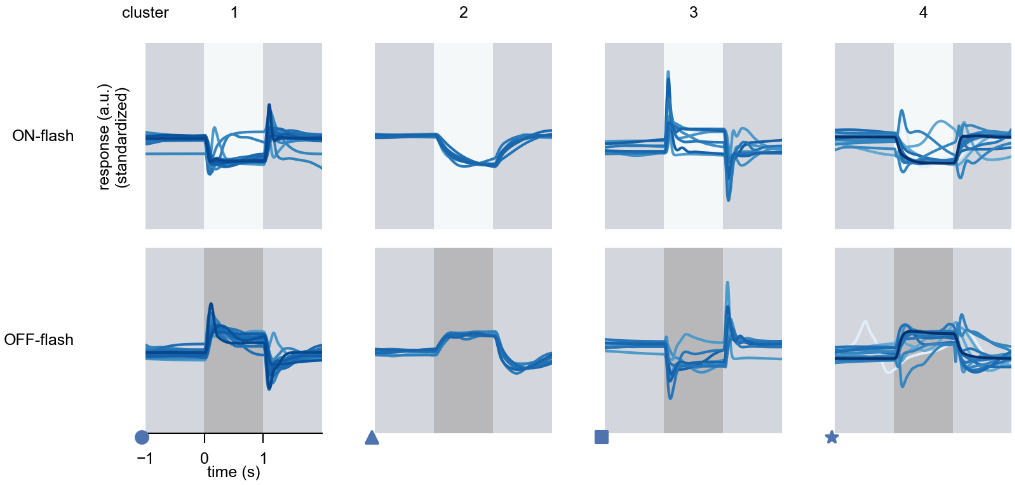

Tm28 - Figure 4: **Responses to flashes.** The top row shows responses to ON-flashes, the bottom row shows responses to OFF-flashes. The responses from the different models in the different clusters (columns) overlay and are colored in the respective task error of the model. The circular flashes occur at second zero, cover 6 ommatidia in radius, and last for one second. Before and after, a grey-stimulus leads to a stationary state of the network. The traces are standardized.

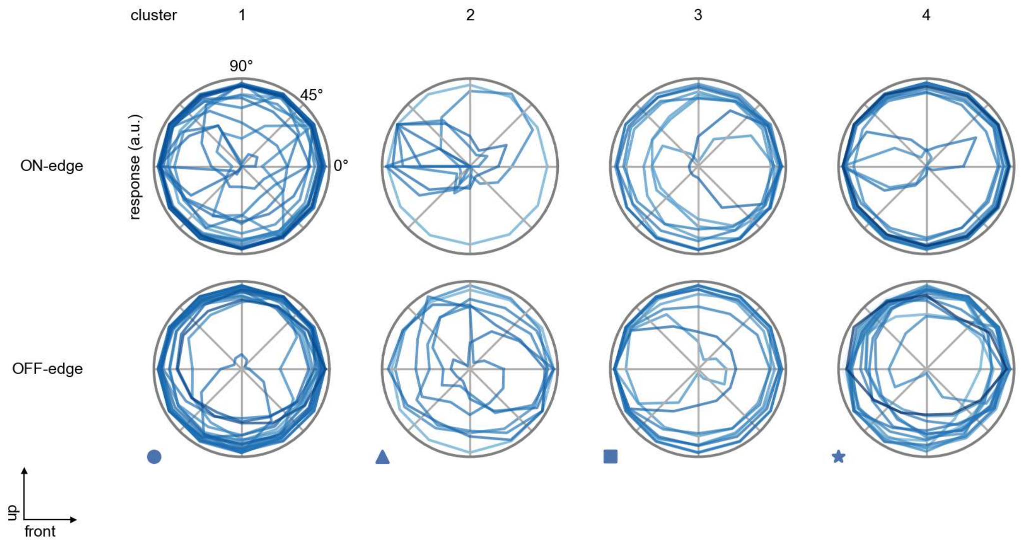

Tm28 - Figure 5: **Peak responses to moving edges.** The top row shows normalized peak responses to moving ON-edges, the bottom row shows normalized peak responses to moving OFF-edges. The edge-stimuli move in different directions from 0 to 360 degrees. Before, a grey-stimulus led the network to a stationary state. The responses from the different models in the different clusters (columns) overlay and are colored in the respective task error of the model. The peak responses are averaged over all assessed edge-speeds.

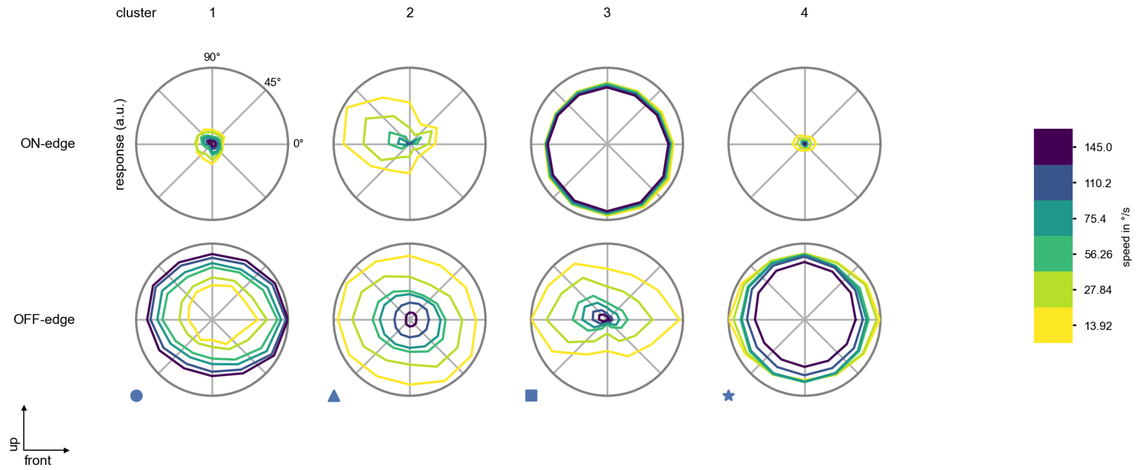

Tm28 - Figure 6: **Peak responses to moving edges from best-performing models.** The top row shows peak responses to moving ON-edges, the bottom row shows peak responses to moving OFF-edges. The edge-stimuli move in different directions from 0 to 360 degrees and at different speeds. Before, a grey-stimulus led the network to a stationary state. The columns show responses from the best-task-performing model in the respective cluster.

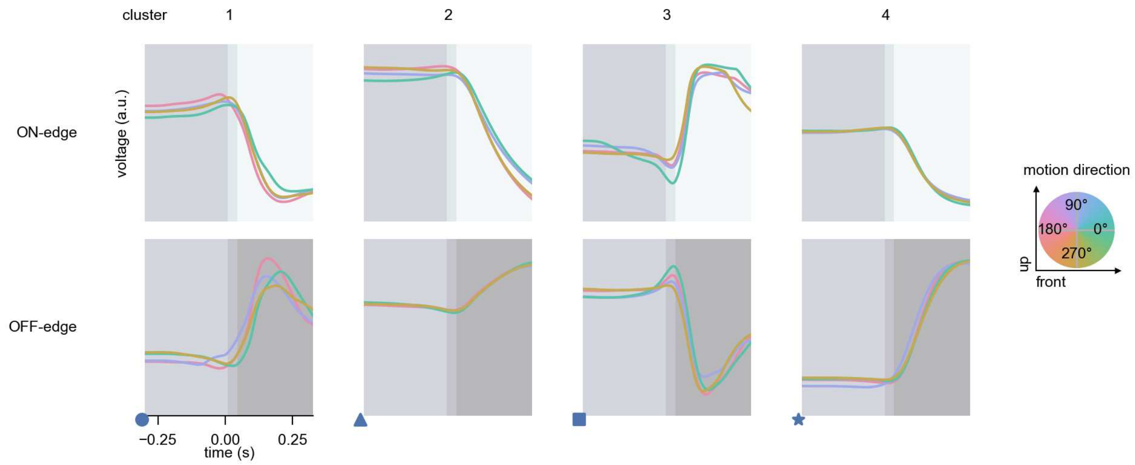

Tm28 - Figure 7: **Responses to moving edges from best-performing models.** The top row shows responses to moving ON-edges, the bottom row shows peak responses to moving OFF-edges. The edge-stimuli move in different directions from 0 to 360 degrees and at different speeds. Before, a grey-stimulus led the network to a stationary state. The columns show responses from the best-task-performing model in the respective cluster. Example responses to edges moving at 75.4°/s in all cardinal directions (green 0°, blue 90°, red 180°, yellow 270°) from -22.5 to 22.5° visual angle. Background contour indicates the edge, passing by the central column at second zero.

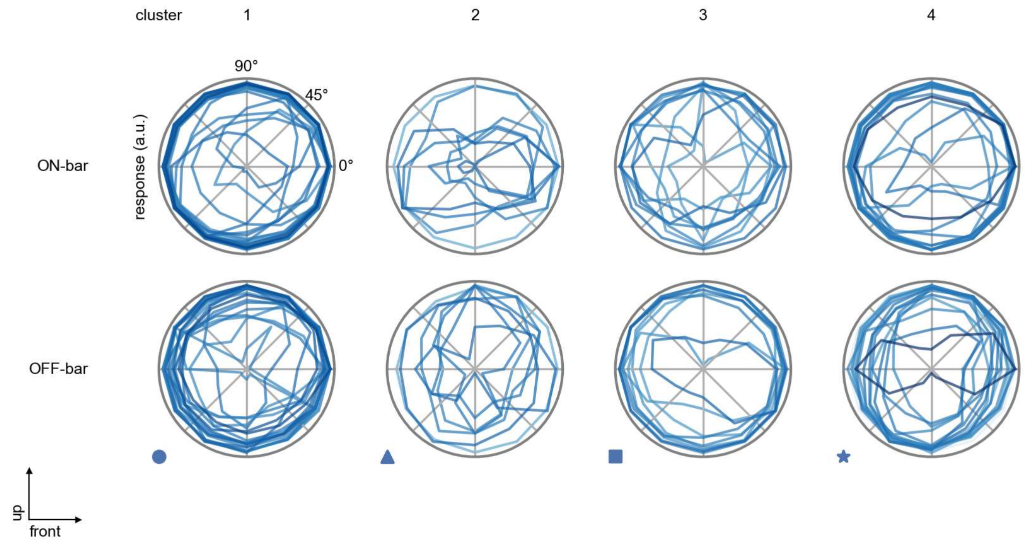

Tm28 - Figure 8: **Peak responses to moving bars.** The top row shows normalized peak responses to moving ON-bars, the bottom row shows normalized peak responses to moving OFF-bars. The bar-stimuli move in different directions from 0 to 360 degrees. Before, a grey-stimulus led the network to a stationary state. The responses from the different models in the different clusters (columns) overlay and are colored in the respective task error of the model. The peak responses are averaged over all assessed bar-speeds.

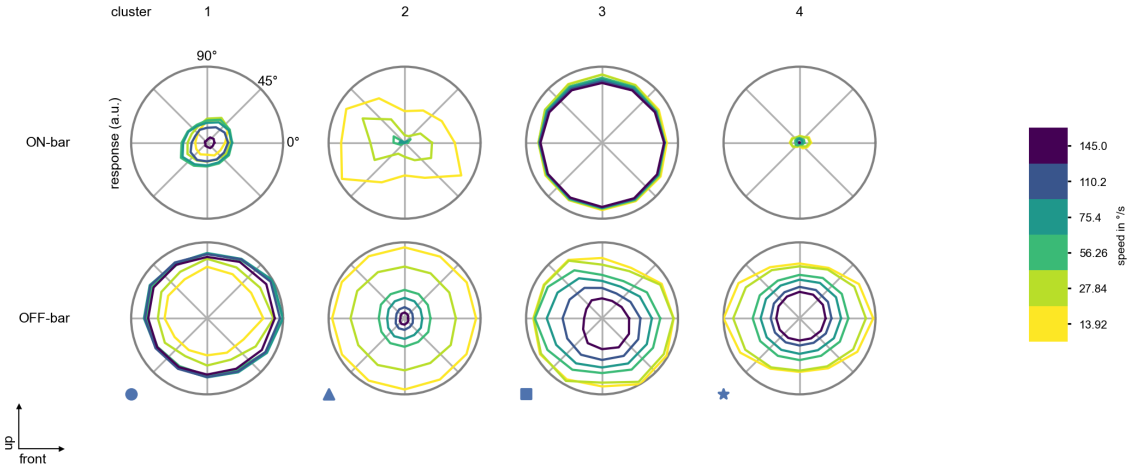

Tm28 - Figure 9: **Peak responses to moving bars from best-performing models.** The top row shows peak responses to moving ON-bars, the bottom row shows peak responses to moving OFF-bars. The bar-stimuli move in different directions from 0 to 360 degrees and at different speeds. Before, a grey-stimulus led the network to a stationary state. The columns show responses from the best-task-performing model in the respective cluster.

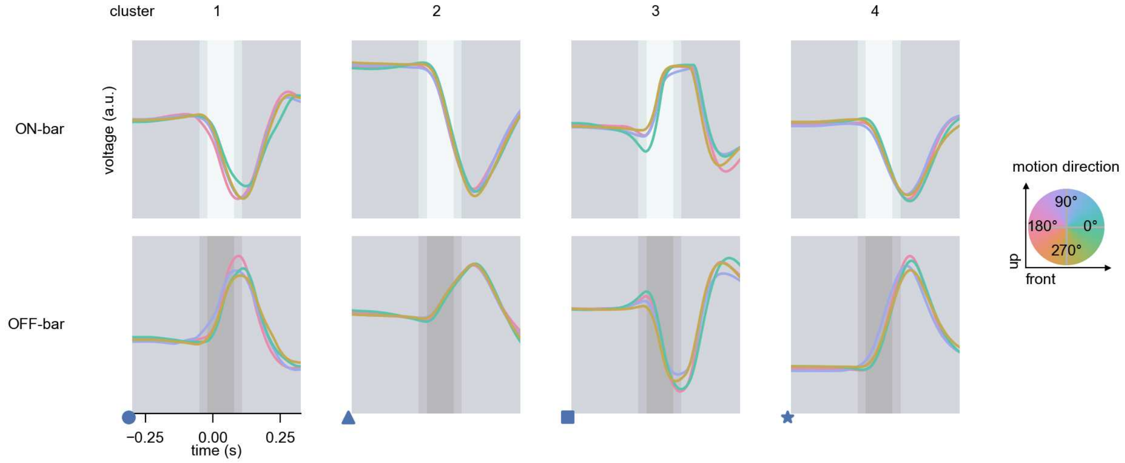

Tm28 - Figure 10: **Responses to moving bars from best-performing models.** The top row shows responses to moving ON-bars, the bottom row shows peak responses to moving OFF-bars. The bar-stimuli move in different directions from 0 to 360 degrees and at different speeds. Before, a grey-stimulus led the network to a stationary state. The columns show responses from the best-task-performing model in the respective cluster. Example responses to bars moving at 75.4°/s in all cardinal directions (green 0°, blue 90°, red 180°, yellow 270°) from -22.5 to 22.5° visual angle. Background contour indicates the bar, passing by the central column at second zero.

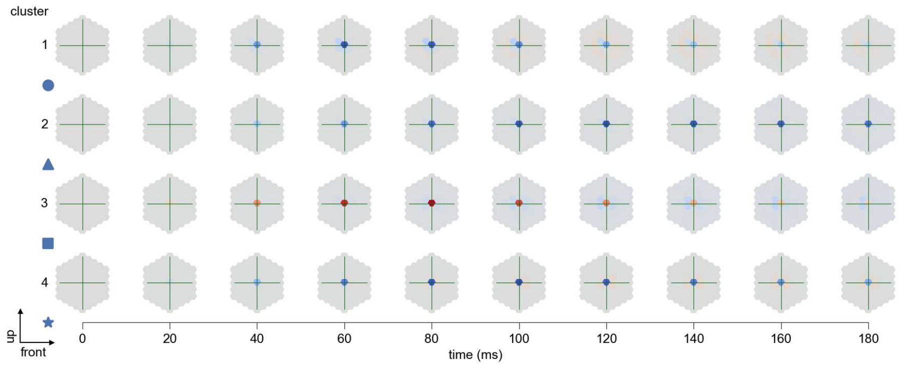

Tm28 - Figure 11: **Spatio-temporal receptive field.** Responses of the central cell to ON-impulses at single ommatidium impulse locations. The impulse occurs at second zero. Before, a grey-stimulus led the network to a stationary state. Responses are baseline-subtracted. Each row illustrates responses from the best-performing model of the respective cluster. Red indicated depolarization, blue indicates hyperpolarization.

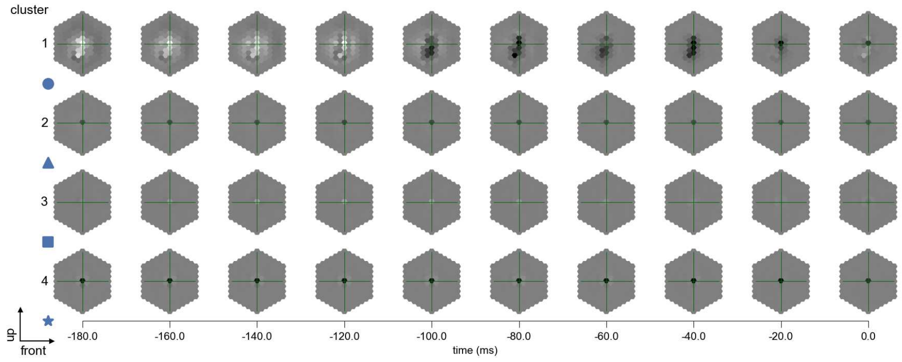

Tm28 - Figure 12: **Maximally excitatory stimuli.** Each row presents the regularized naturalistic-stimulus from the Sintel dataset that maximizes the cell type's central column response at second zero in the best-performing model of the respective cluster.

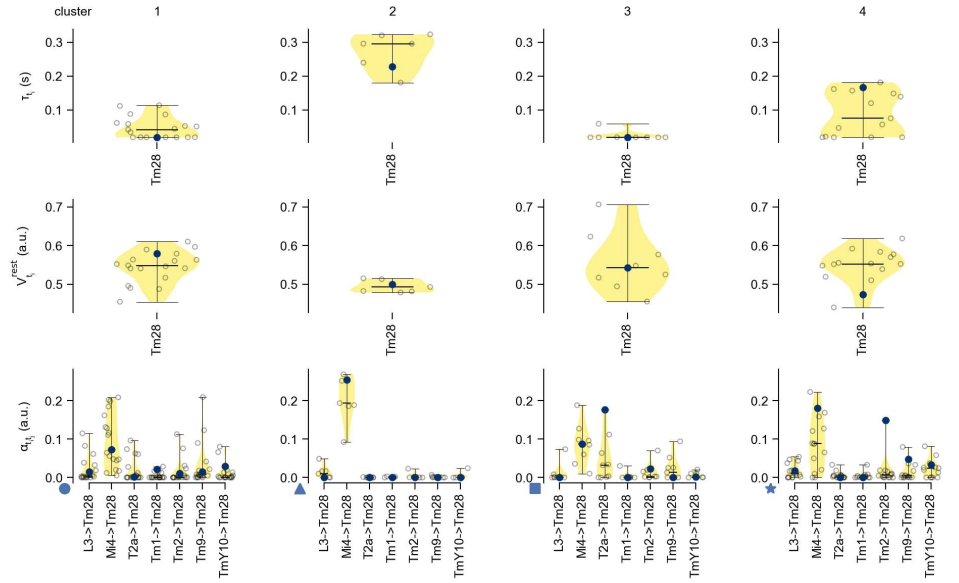

Tm28 - Figure 13: **Task-constrained parameters.** Each column shows the parameters inferred within the respective cluster. First row: learned time constants of the cell type. Second row: resting potentials of the cell type. Third row: scaling factors for the convolutional filters. The blue scatter represents the parameters from the best-performing model within the cluster.

# 56 Tm30

← Cell types

#### Figures

1 Anatomical receptive fields. . . . 370
2 Clustering of the responses to naturalistic stimuli. . . . 370
3 Responses to flashes. . . . 371
4 Peak responses to moving edges. . . . 371
5 Peak responses to moving edges from best-performing models. . . . 372
6 Responses to moving edges from best-performing models. . . . 372
7 Peak responses to moving bars. . . . 373
8 Peak responses to moving bars from best-performing models. . . . 373
9 Responses to moving bars from best-performing models. . . . 374
10 Spatio-temporal receptive field. . . . 374
11 Maximally excitatory stimuli. . . . 374
12 Task-constrained parameters. . . . 375

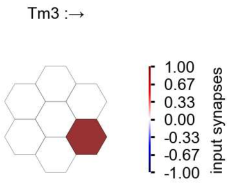

Tm30 - Figure 1: **Anatomical receptive fields.** Each colored hexagon is a projection, with the projection strength characterized by the average number of synapses that we count from the EM reconstruction. Red indicates excitatory synapses, blue indicates inhibitory synapses from inferred signs. Filters in the order of their total number of synapses.

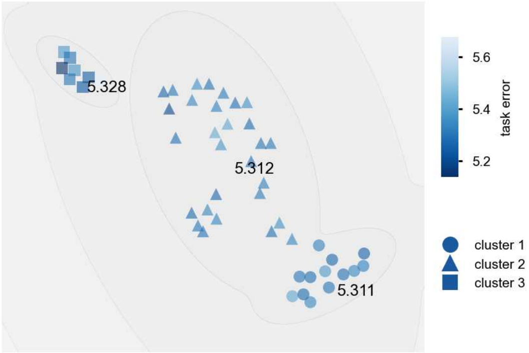

Tm30 - Figure 2: **Clustering of the responses to naturalistic stimuli.** From responses to naturalistic scenes from Sintel. Scatterpoints represent individual models colored by their task error.

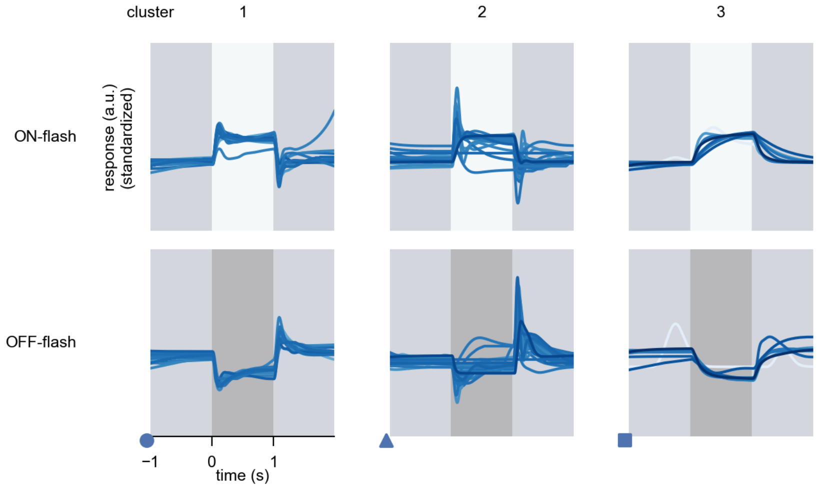

Tm30 - Figure 3: **Responses to flashes.** The top row shows responses to ON-flashes, the bottom row shows responses to OFF-flashes. The responses from the different models in the different clusters (columns) overlay and are colored in the respective task error of the model. The circular flashes occur at second zero, cover 6 ommatidia in radius, and last for one second. Before and after, a grey-stimulus leads to a stationary state of the network. The traces are standardized.

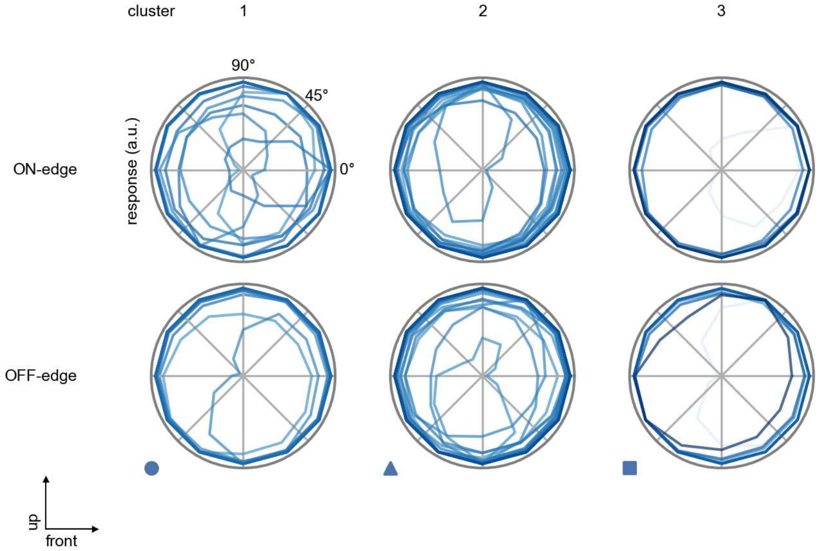

Tm30 - Figure 4: **Peak responses to moving edges.** The top row shows normalized peak responses to moving ON-edges, the bottom row shows normalized peak responses to moving OFF-edges. The edge-stimuli move in different directions from 0 to 360 degrees. Before, a grey-stimulus led the network to a stationary state. The responses from the different models in the different clusters (columns) overlay and are colored in the respective task error of the model. The peak responses are averaged over all assessed edge-speeds.

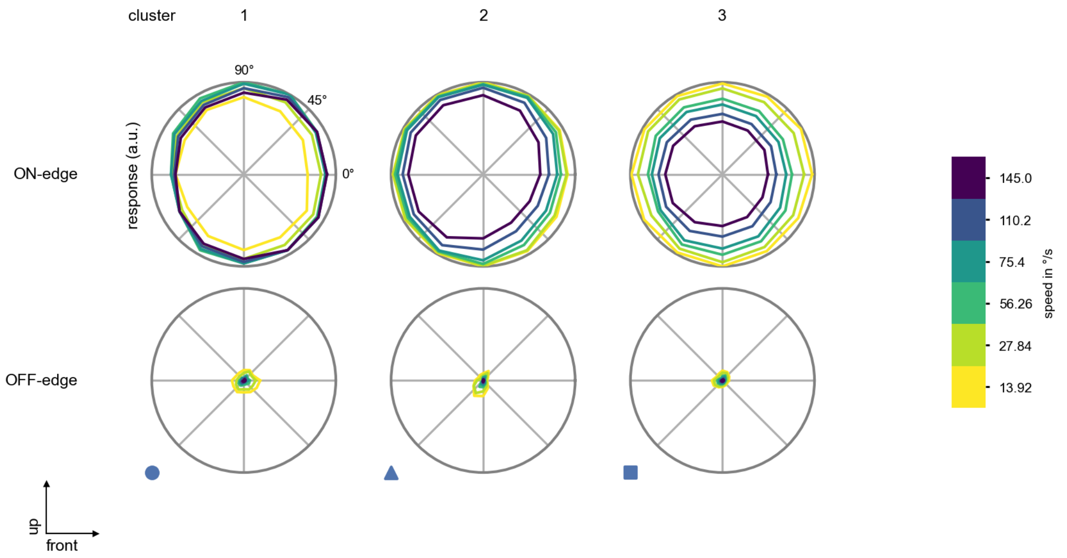

Tm30 - Figure 5: **Peak responses to moving edges from best-performing models.** The top row shows peak responses to moving ON-edges, the bottom row shows peak responses to moving OFF-edges. The edge-stimuli move in different directions from 0 to 360 degrees and at different speeds. Before, a grey-stimulus led the network to a stationary state. The columns show responses from the best-task-performing model in the respective cluster.

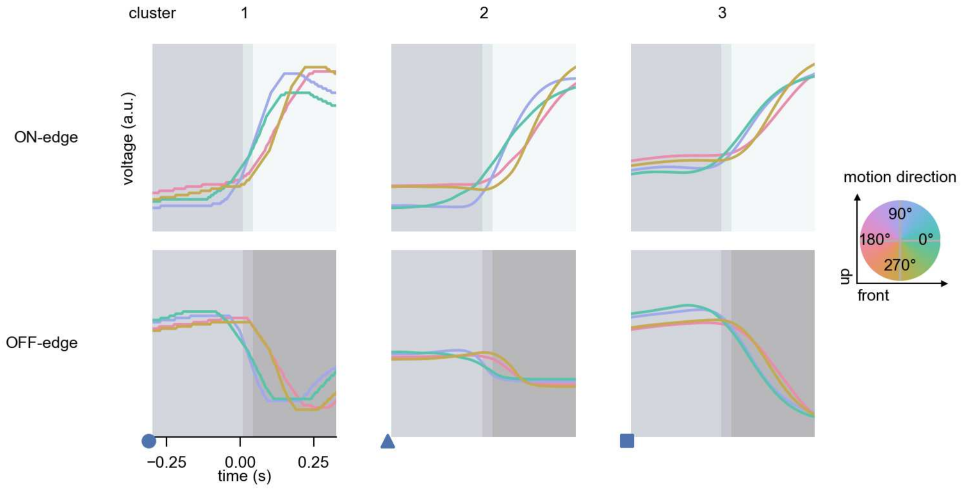

Tm30 - Figure 6: **Responses to moving edges from best-performing models.** The top row shows responses to moving ON-edges, the bottom row shows peak responses to moving OFF-edges. The edge-stimuli move in different directions from 0 to 360 degrees and at different speeds. Before, a grey-stimulus led the network to a stationary state. The columns show responses from the best-task-performing model in the respective cluster. Example responses to edges moving at 75.4°/s in all cardinal directions (green 0°, blue 90°, red 180°, yellow 270°) from -22.5 to 22.5° visual angle. Background contour indicates the edge, passing by the central column at second zero.

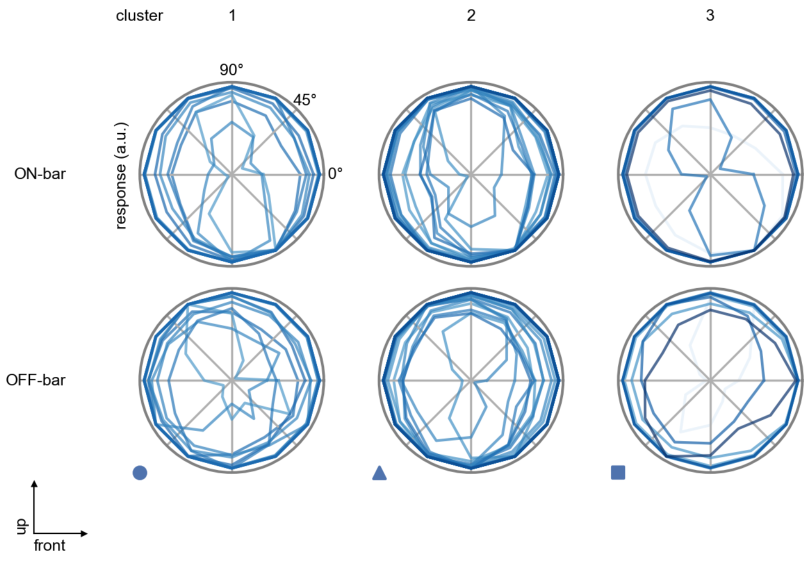

Tm30 - Figure 7: **Peak responses to moving bars.** The top row shows normalized peak responses to moving ON-bars, the bottom row shows normalized peak responses to moving OFF-bars. The bar-stimuli move in different directions from 0 to 360 degrees. Before, a grey-stimulus led the network to a stationary state. The responses from the different models in the different clusters (columns) overlay and are colored in the respective task error of the model. The peak responses are averaged over all assessed bar-speeds.

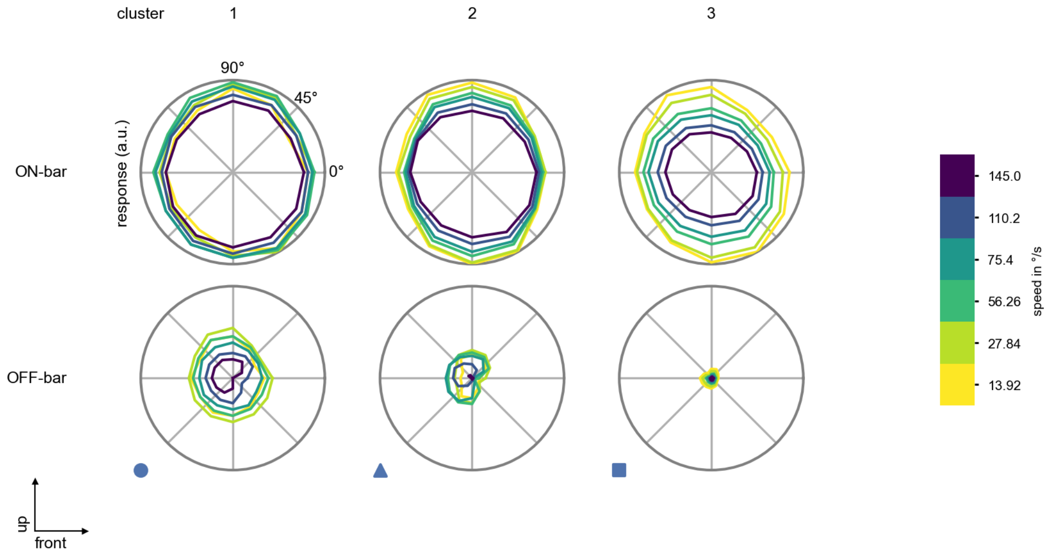

Tm30 - Figure 8: **Peak responses to moving bars from best-performing models.** The top row shows peak responses to moving ON-bars, the bottom row shows peak responses to moving OFF-bars. The bar-stimuli move in different directions from 0 to 360 degrees and at different speeds. Before, a grey-stimulus led the network to a stationary state. The columns show responses from the best-task-performing model in the respective cluster.

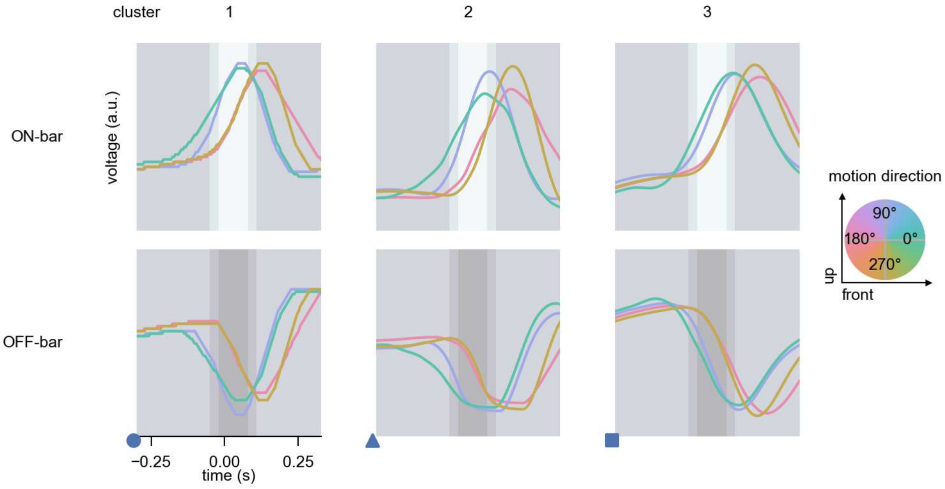

Tm30 - Figure 9: **Responses to moving bars from best-performing models.** The top row shows responses to moving ON-bars, the bottom row shows peak responses to moving OFF-bars. The bar-stimuli move in different directions from 0 to 360 degrees and at different speeds. Before, a grey-stimulus led the network to a stationary state. The columns show responses from the best-task-performing model in the respective cluster. Example responses to bars moving at 75.4°/s in all cardinal directions (green 0°, blue 90°, red 180°, yellow 270°) from -22.5 to 22.5° visual angle. Background contour indicates the bar, passing by the central column at second zero.

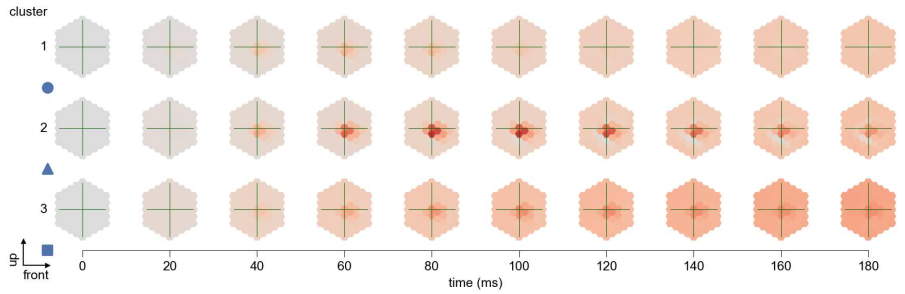

Tm30 - Figure 10: **Spatio-temporal receptive field.** Responses of the central cell to ON-impulses at single ommatidium impulse locations. The impulse occurs at second zero. Before, a grey-stimulus led the network to a stationary state. Responses are baseline-subtracted. Each row illustrates responses from the best-performing model of the respective cluster. Red indicated depolarization, blue indicates hyperpolarization.

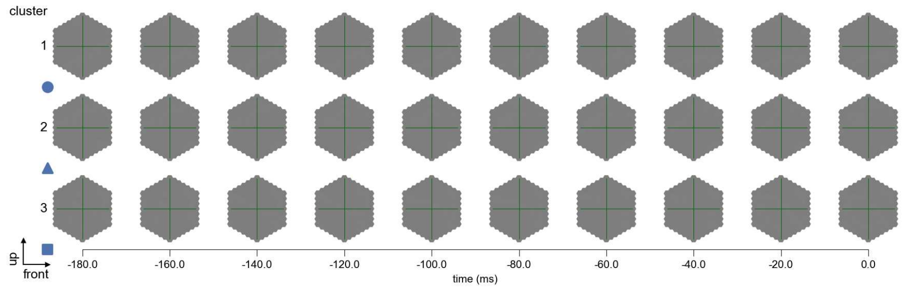

Tm30 - Figure 11: **Maximally excitatory stimuli.** Each row presents the regularized naturalistic-stimulus from the Sintel dataset that maximizes the cell type's central column response at second zero in the best-performing model of the respective cluster.

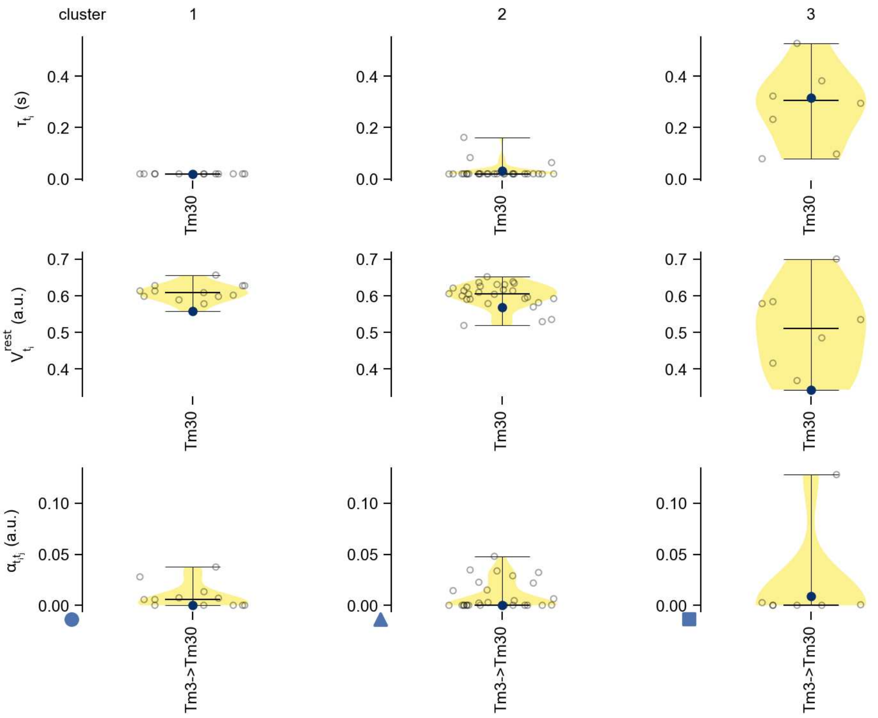

Tm30 - Figure 12: **Task-constrained parameters.** Each column shows the parameters inferred within the respective cluster. First row: learned time constants of the cell type. Second row: resting potentials of the cell type. Third row: scaling factors for the convolutional filters. The blue scatter represents the parameters from the best-performing model within the cluster.

### 57 TmY3

← Cell types

#### Figures

1 Anatomical receptive fields. . . . . 376
2 Anatomical projective fields. . . . . 377
3 Clustering of the responses to naturalistic stimuli. . . . . 377
4 Responses to flashes. . . . . 377
5 Peak responses to moving edges. . . . . 378
6 Peak responses to moving edges from best-performing models. . . . . 378
7 Responses to moving edges from best-performing models. . . . . 379
8 Peak responses to moving bars. . . . . 379
9 Peak responses to moving bars from best-performing models. . . . . 380
10 Responses to moving bars from best-performing models. . . . . 380
11 Spatio-temporal receptive field. . . . . 381
12 Maximally excitatory stimuli. . . . . 381
13 Task-constrained parameters. . . . . 382

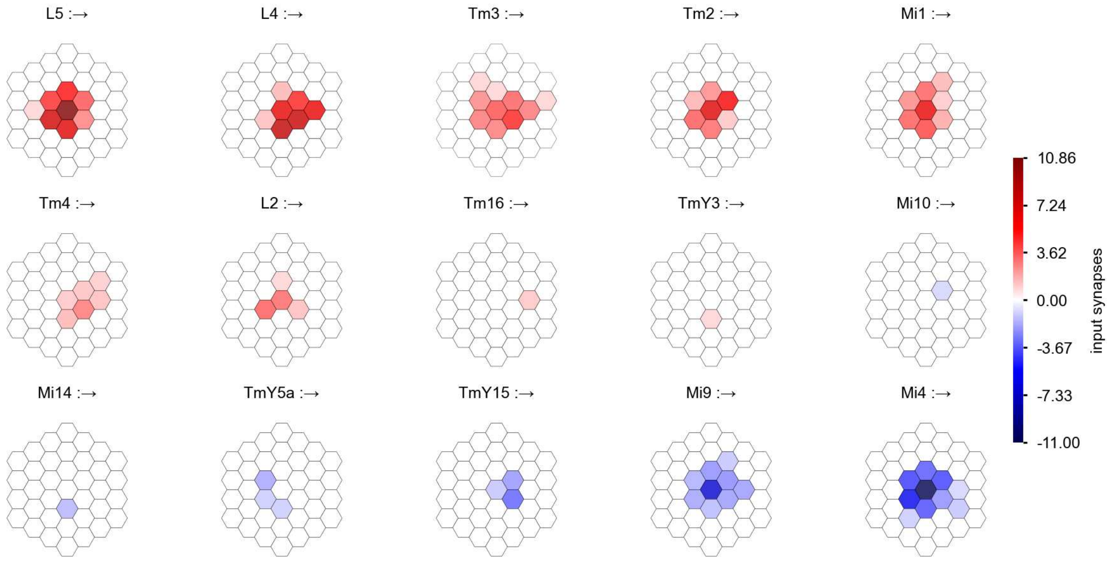

TmY3 - Figure 1: **Anatomical receptive fields.** Each colored hexagon is a projection, with the projection strength characterized by the average number of synapses that we count from the EM reconstruction. Red indicates excitatory synapses, blue indicates inhibitory synapses from inferred signs. Filters in the order of their total number of synapses.

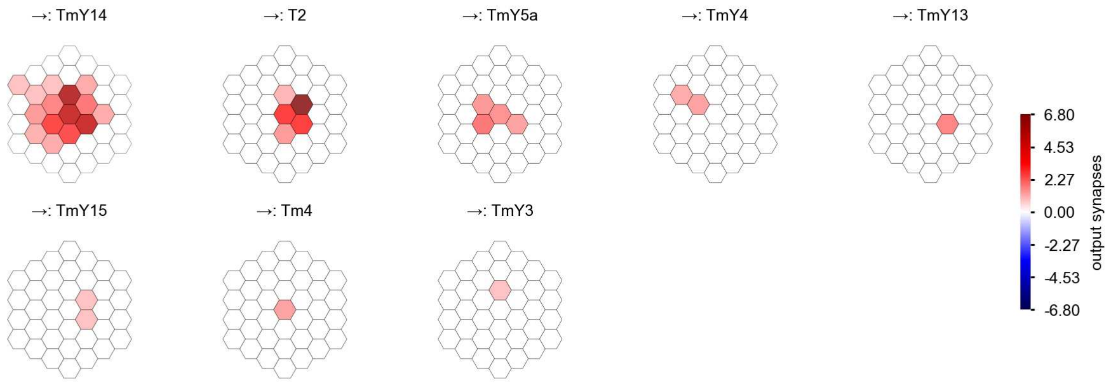

TmY3 - Figure 2: **Anatomical projective fields.** Each colored hexagon is a projection, with the projection strength characterized by the average number of synapses that we count from the EM reconstruction. Red indicates excitatory synapses, blue indicates inhibitory synapses from inferred signs. Filters in the order of their total number of synapses.

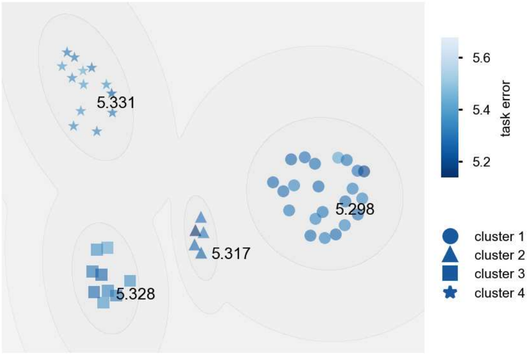

TmY3 - Figure 3: **Clustering of the responses to naturalistic stimuli.** From responses to naturalistic scenes from Sintel. Scatterpoints represent individual models colored by their task error.

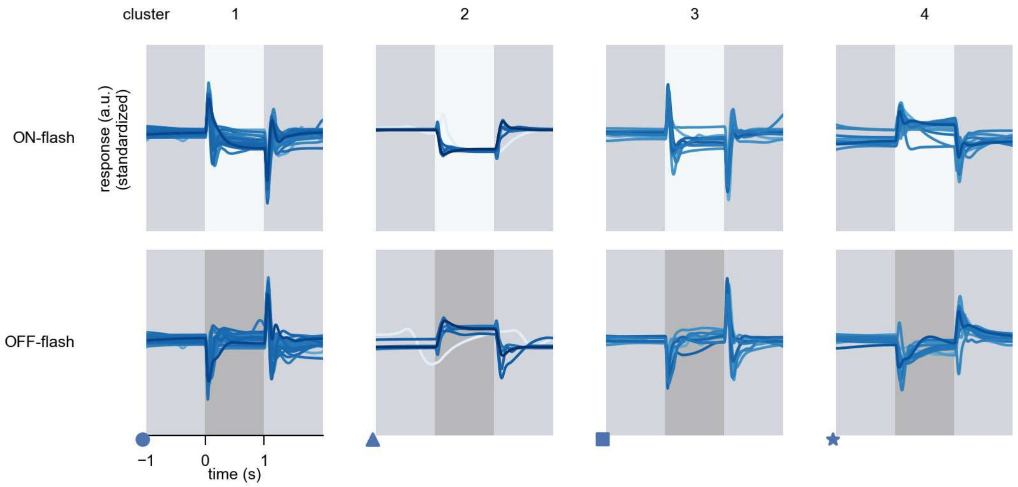

TmY3 - Figure 4: **Responses to flashes.** The top row shows responses to ON-flashes, the bottom row shows responses to OFF-flashes. The responses from the different models in the different clusters (columns) overlay and are colored in the respective task error of the model. The circular flashes occur at second zero, cover 6 ommatidia in radius, and last for one second. Before and after, a grey-stimulus leads to a stationary state of the network. The traces are standardized.

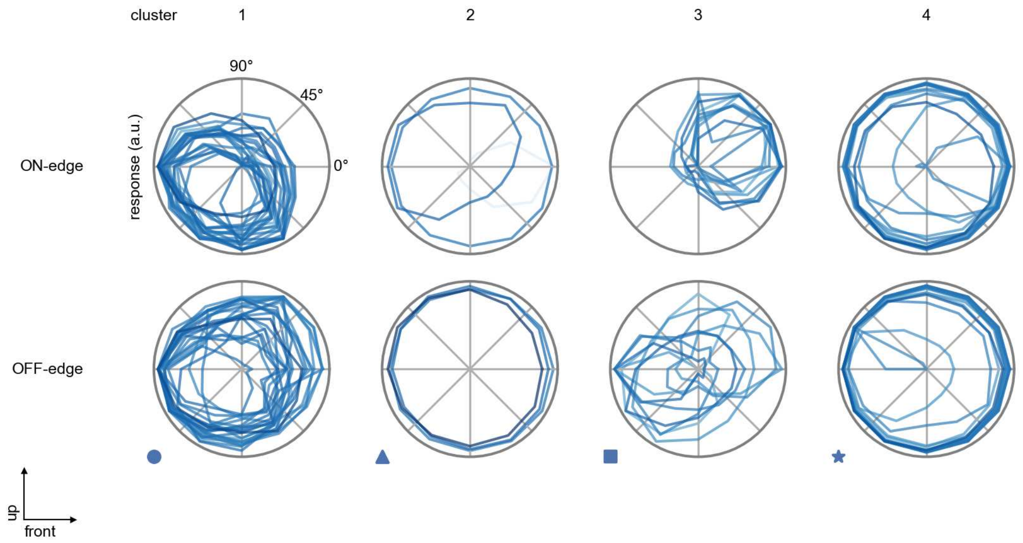

TmY3 - Figure 5: **Peak responses to moving edges.** The top row shows normalized peak responses to moving ON-edges, the bottom row shows normalized peak responses to moving OFF-edges. The edge-stimuli move in different directions from 0 to 360 degrees. Before, a grey-stimulus led the network to a stationary state. The responses from the different models in the different clusters (columns) overlay and are colored in the respective task error of the model. The peak responses are averaged over all assessed edge-speeds.

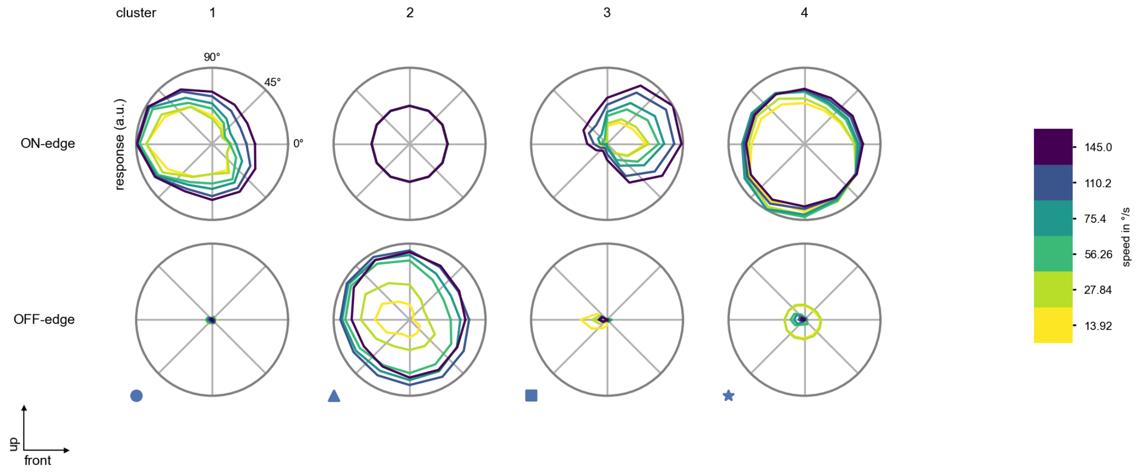

TmY3 - Figure 6: **Peak responses to moving edges from best-performing models.** The top row shows peak responses to moving ON-edges, the bottom row shows peak responses to moving OFF-edges. The edge-stimuli move in different directions from 0 to 360 degrees and at different speeds. Before, a grey-stimulus led the network to a stationary state. The columns show responses from the best-task-performing model in the respective cluster.

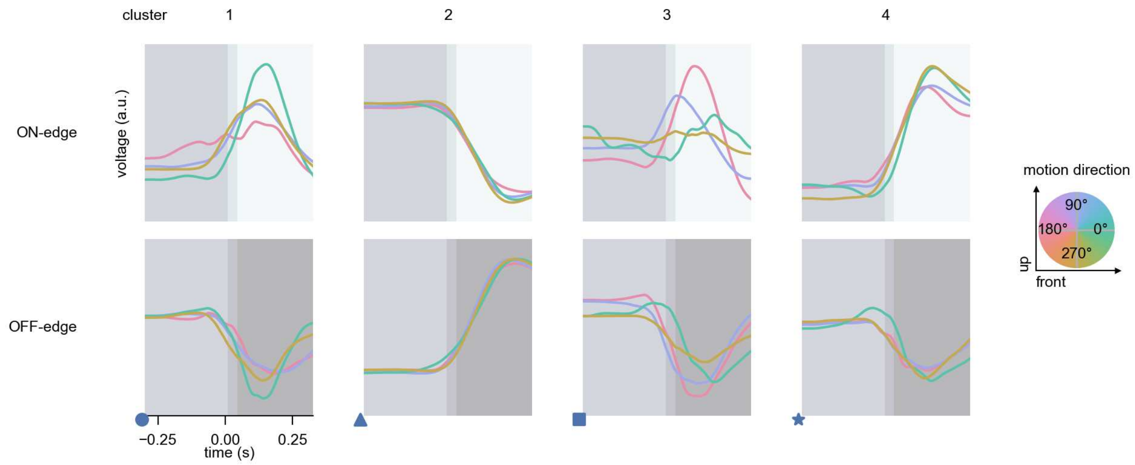

TmY3 - Figure 7: **Responses to moving edges from best-performing models.** The top row shows responses to moving ON-edges, the bottom row shows peak responses to moving OFF-edges. The edge-stimuli move in different directions from 0 to 360 degrees and at different speeds. Before, a grey-stimulus led the network to a stationary state. The columns show responses from the best-task-performing model in the respective cluster. Example responses to edges moving at 75.4°/s in all cardinal directions (green 0°, blue 90°, red 180°, yellow 270°) from -22.5 to 22.5° visual angle. Background contour indicates the edge, passing by the central column at second zero.

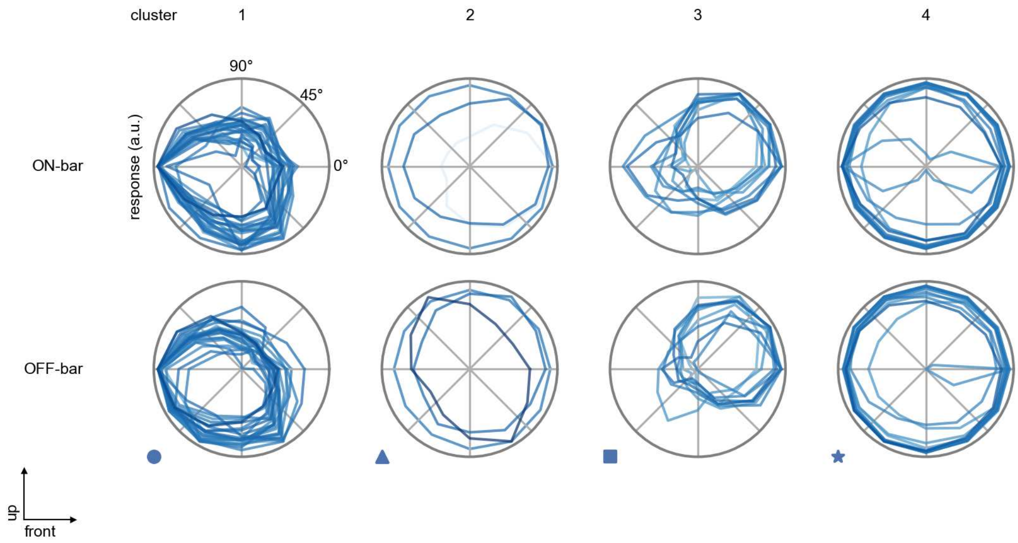

TmY3 - Figure 8: **Peak responses to moving bars.** The top row shows normalized peak responses to moving ON-bars, the bottom row shows normalized peak responses to moving OFF-bars. The bar-stimuli move in different directions from 0 to 360 degrees. Before, a grey-stimulus led the network to a stationary state. The responses from the different models in the different clusters (columns) overlay and are colored in the respective task error of the model. The peak responses are averaged over all assessed bar-speeds.

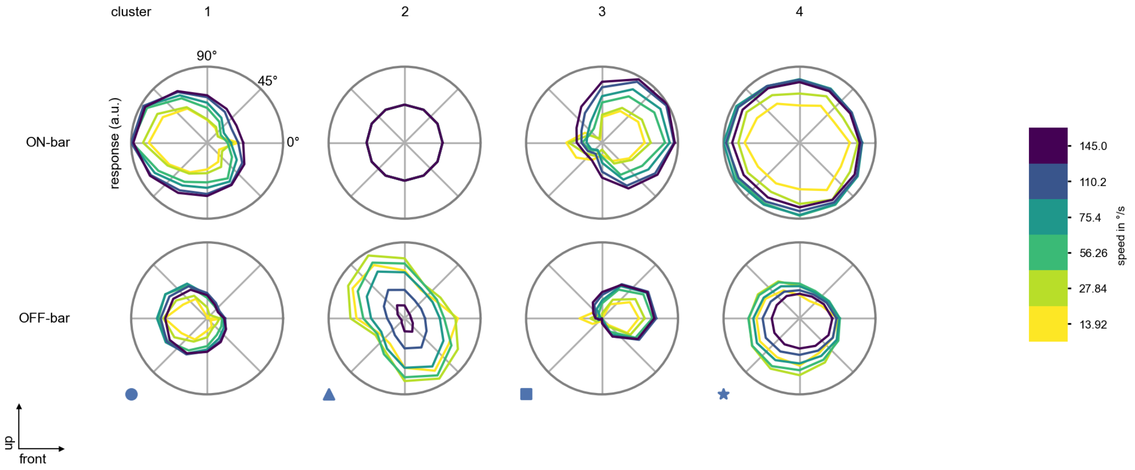

TmY3 - Figure 9: **Peak responses to moving bars from best-performing models.** The top row shows peak responses to moving ON-bars, the bottom row shows peak responses to moving OFF-bars. The bar-stimuli move in different directions from 0 to 360 degrees and at different speeds. Before, a grey-stimulus led the network to a stationary state. The columns show responses from the best-task-performing model in the respective cluster.

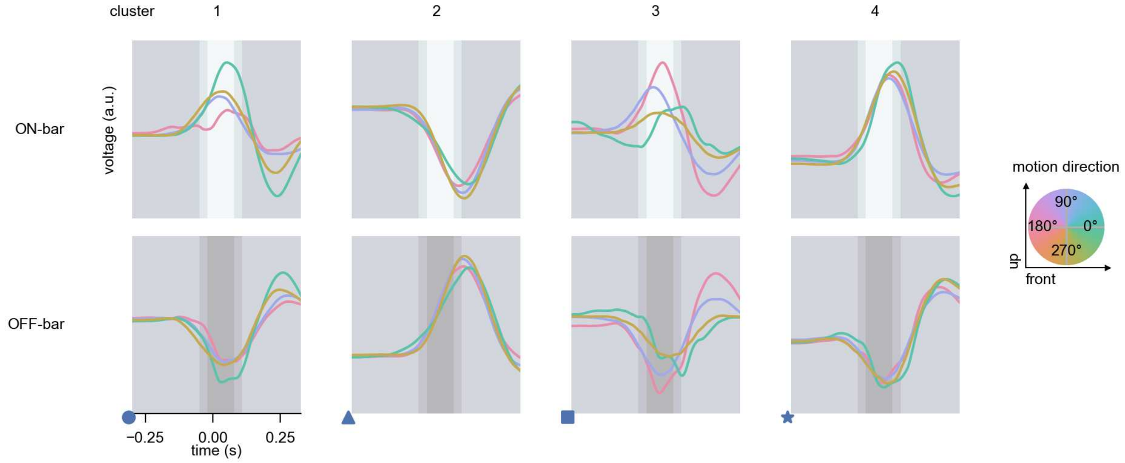

TmY3 - Figure 10: **Responses to moving bars from best-performing models.** The top row shows responses to moving ON-bars, the bottom row shows peak responses to moving OFF-bars. The bar-stimuli move in different directions from 0 to 360 degrees and at different speeds. Before, a grey-stimulus led the network to a stationary state. The columns show responses from the best-task-performing model in the respective cluster. Example responses to bars moving at 75.4°/s in all cardinal directions (green 0°, blue 90°, red 180°, yellow 270°) from -22.5 to 22.5° visual angle. Background contour indicates the bar, passing by the central column at second zero.

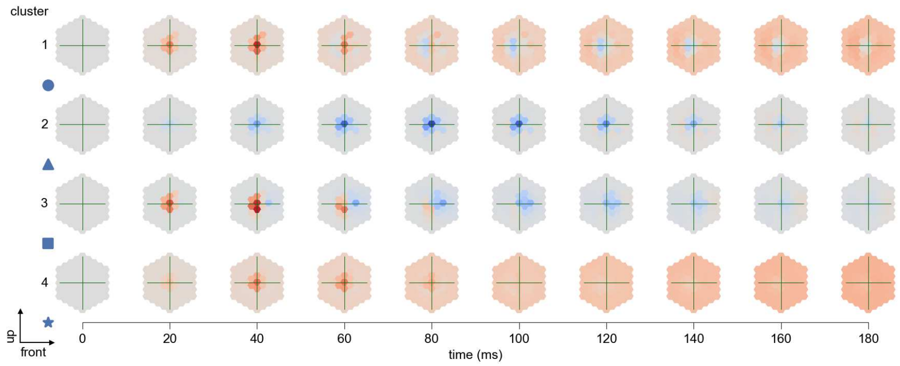

TmY3 - Figure 11: **Spatio-temporal receptive field.** Responses of the central cell to ON-impulses at single ommatidium impulse locations. The impulse occurs at second zero. Before, a grey-stimulus led the network to a stationary state. Responses are baseline-subtracted. Each row illustrates responses from the best-performing model of the respective cluster. Red indicated depolarization, blue indicates hyperpolarization.

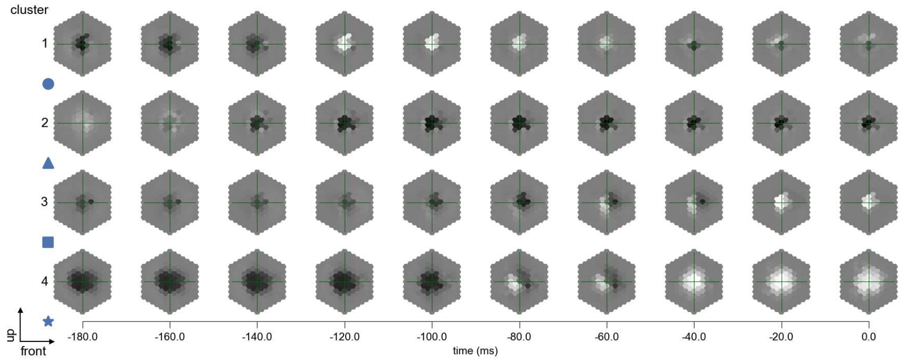

TmY3 - Figure 12: **Maximally excitatory stimuli.** Each row presents the regularized naturalistic-stimulus from the Sintel dataset that maximizes the cell type's central column response at second zero in the best-performing model of the respective cluster.

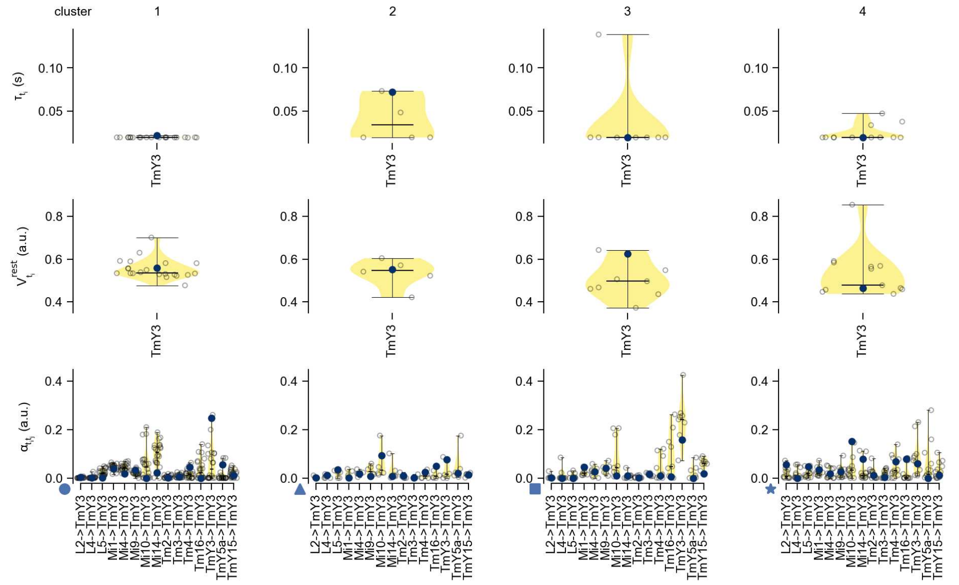

TmY3 - Figure 13: **Task-constrained parameters.** Each column shows the parameters inferred within the respective cluster. First row: learned time constants of the cell type. Second row: resting potentials of the cell type. Third row: scaling factors for the convolutional filters. The blue scatter represents the parameters from the best-performing model within the cluster.

### 58 TmY4

← Cell types

#### Figures

1 Anatomical receptive fields. . . . . 383
2 Anatomical projective fields. . . . . 384
3 Clustering of the responses to naturalistic stimuli. . . . . 384
4 Responses to flashes. . . . . 384
5 Peak responses to moving edges. . . . . 385
6 Peak responses to moving edges from best-performing models. . . . . 385
7 Responses to moving edges from best-performing models. . . . . 386
8 Peak responses to moving bars. . . . . 386
9 Peak responses to moving bars from best-performing models. . . . . 387
10 Responses to moving bars from best-performing models. . . . . 387
11 Spatio-temporal receptive field. . . . . 388
12 Maximally excitatory stimuli. . . . . 388
13 Task-constrained parameters. . . . . 389

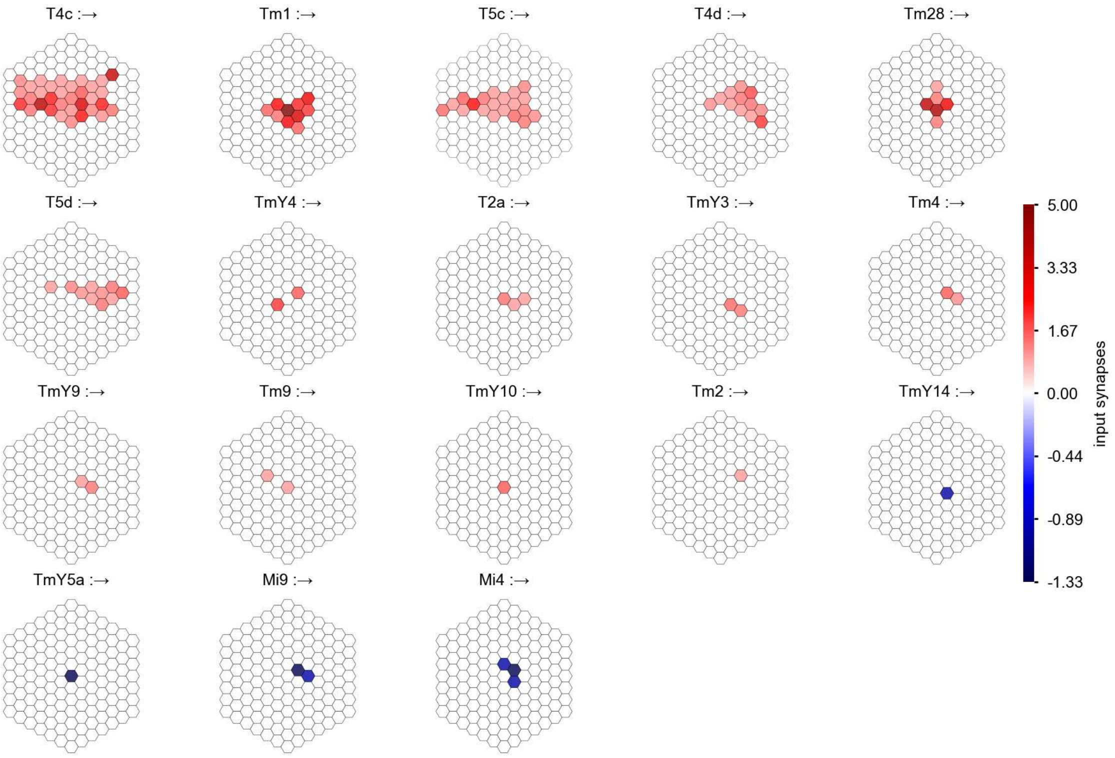

TmY4 - Figure 1: **Anatomical receptive fields.** Each colored hexagon is a projection, with the projection strength characterized by the average number of synapses that we count from the EM reconstruction. Red indicates excitatory synapses, blue indicates inhibitory synapses from inferred signs. Filters in the order of their total number of synapses.

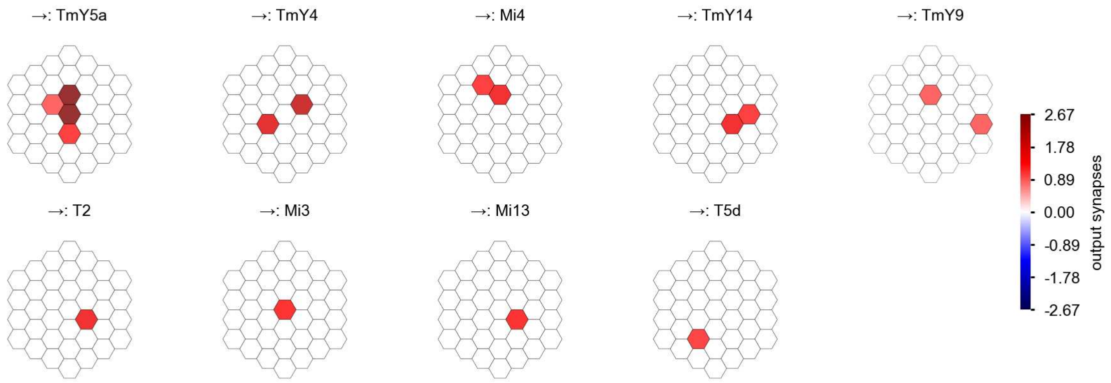

TmY4 - Figure 2: **Anatomical projective fields.** Each colored hexagon is a projection, with the projection strength characterized by the average number of synapses that we count from the EM reconstruction. Red indicates excitatory synapses, blue indicates inhibitory synapses from inferred signs. Filters in the order of their total number of synapses.

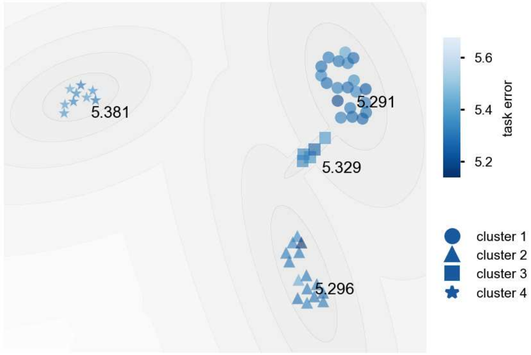

TmY4 - Figure 3: **Clustering of the responses to naturalistic stimuli.** From responses to naturalistic scenes from Sintel. Scatterpoints represent individual models colored by their task error.

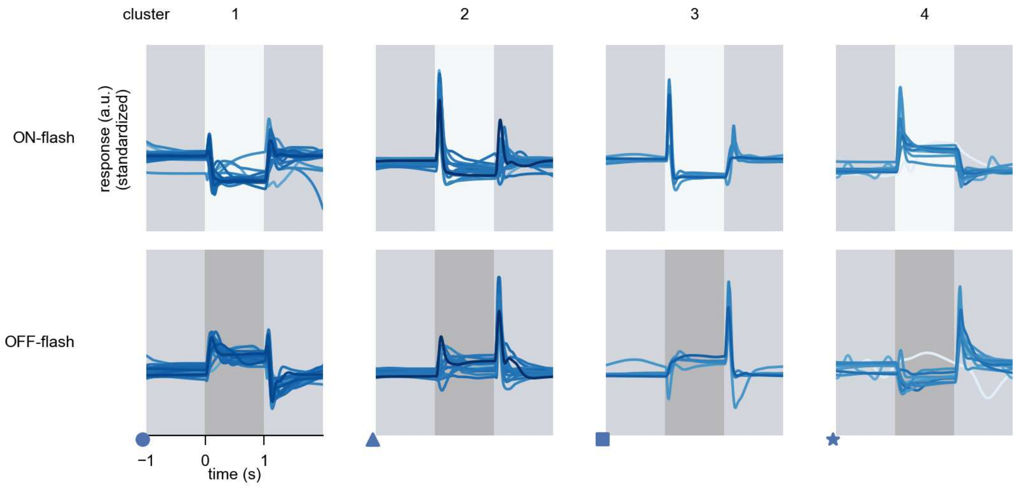

TmY4 - Figure 4: **Responses to flashes.** The top row shows responses to ON-flashes, the bottom row shows responses to OFF-flashes. The responses from the different models in the different clusters (columns) overlay and are colored in the respective task error of the model. The circular flashes occur at second zero, cover 6 ommatidia in radius, and last for one second. Before and after, a grey-stimulus leads to a stationary state of the network. The traces are standardized.

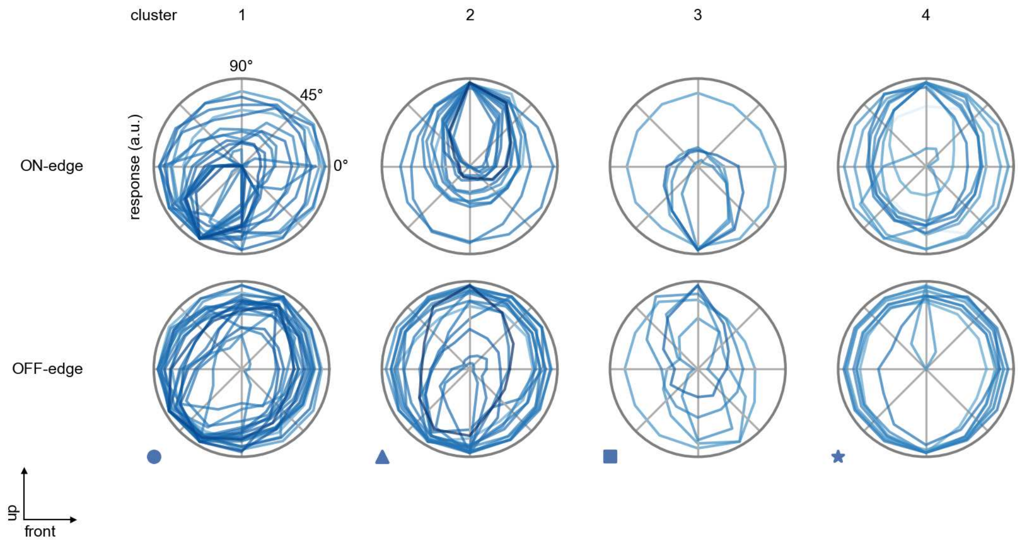

TmY4 - Figure 5: **Peak responses to moving edges.** The top row shows normalized peak responses to moving ON-edges, the bottom row shows normalized peak responses to moving OFF-edges. The edge-stimuli move in different directions from 0 to 360 degrees. Before, a grey-stimulus led the network to a stationary state. The responses from the different models in the different clusters (columns) overlay and are colored in the respective task error of the model. The peak responses are averaged over all assessed edge-speeds.

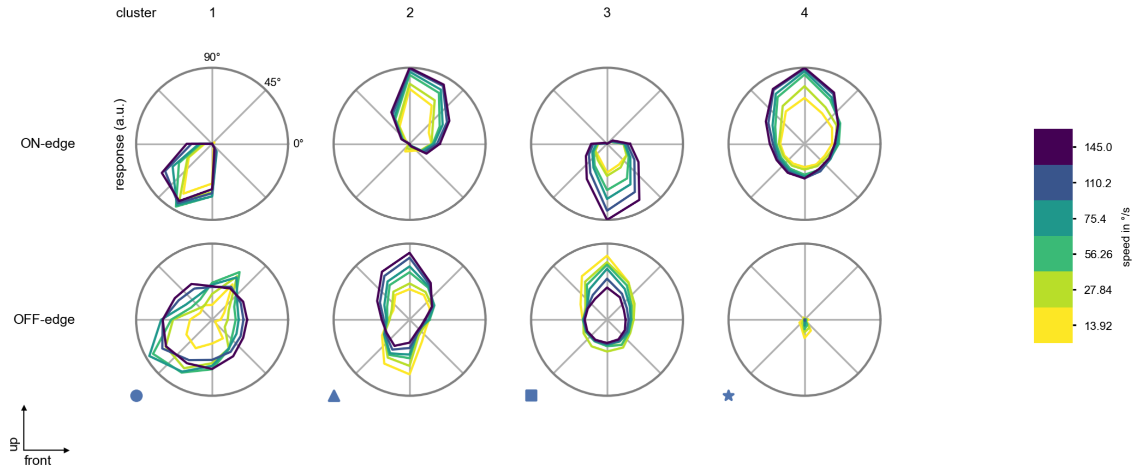

TmY4 - Figure 6: **Peak responses to moving edges from best-performing models.** The top row shows peak responses to moving ON-edges, the bottom row shows peak responses to moving OFF-edges. The edge-stimuli move in different directions from 0 to 360 degrees and at different speeds. Before, a grey-stimulus led the network to a stationary state. The columns show responses from the best-task-performing model in the respective cluster.

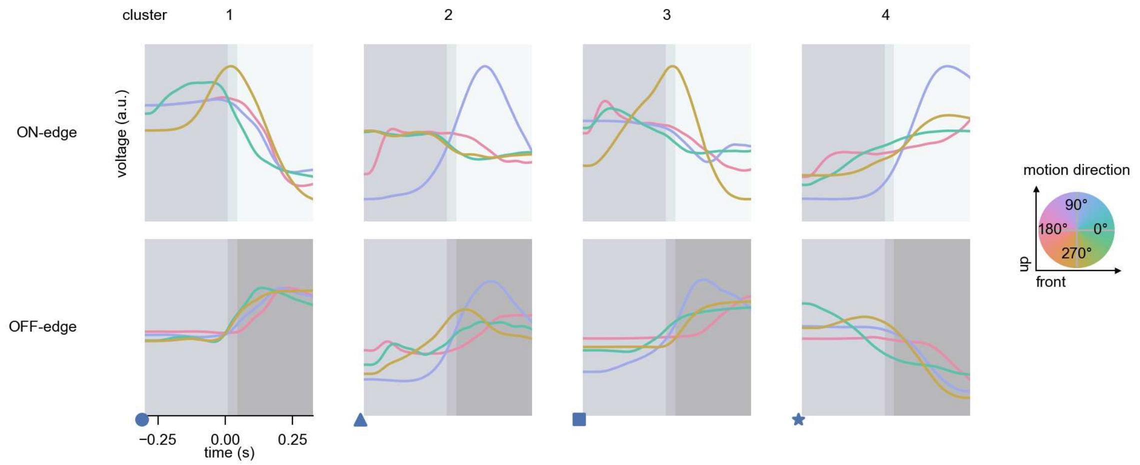

TmY4 - Figure 7: **Responses to moving edges from best-performing models.** The top row shows responses to moving ON-edges, the bottom row shows peak responses to moving OFF-edges. The edge-stimuli move in different directions from 0 to 360 degrees and at different speeds. Before, a grey-stimulus led the network to a stationary state. The columns show responses from the best-task-performing model in the respective cluster. Example responses to edges moving at 75.4°/s in all cardinal directions (green 0°, blue 90°, red 180°, yellow 270°) from -22.5 to 22.5° visual angle. Background contour indicates the edge, passing by the central column at second zero.

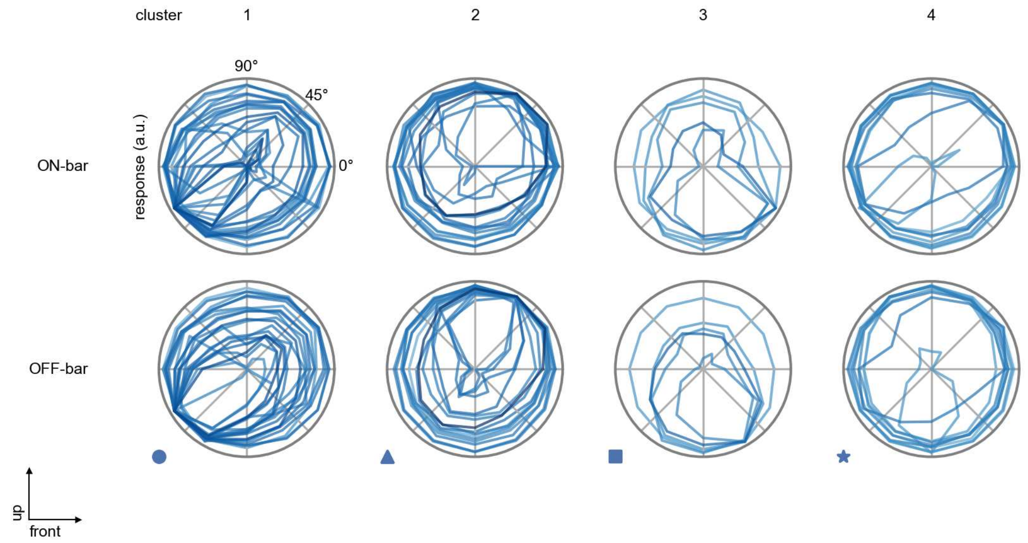

TmY4 - Figure 8: **Peak responses to moving bars.** The top row shows normalized peak responses to moving ON-bars, the bottom row shows normalized peak responses to moving OFF-bars. The bar-stimuli move in different directions from 0 to 360 degrees. Before, a grey-stimulus led the network to a stationary state. The responses from the different models in the different clusters (columns) overlay and are colored in the respective task error of the model. The peak responses are averaged over all assessed bar-speeds.

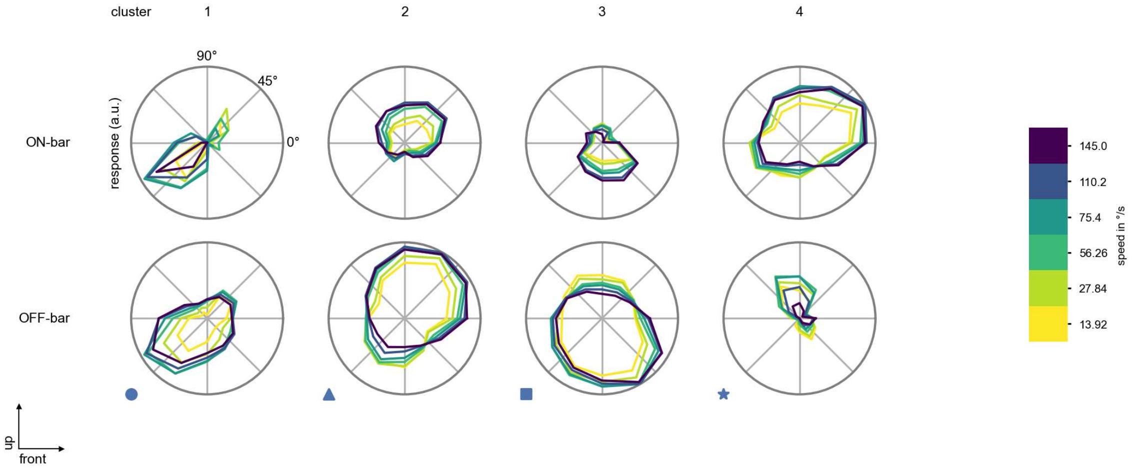

TmY4 - Figure 9: **Peak responses to moving bars from best-performing models.** The top row shows peak responses to moving ON-bars, the bottom row shows peak responses to moving OFF-bars. The bar-stimuli move in different directions from 0 to 360 degrees and at different speeds. Before, a grey-stimulus led the network to a stationary state. The columns show responses from the best-task-performing model in the respective cluster.

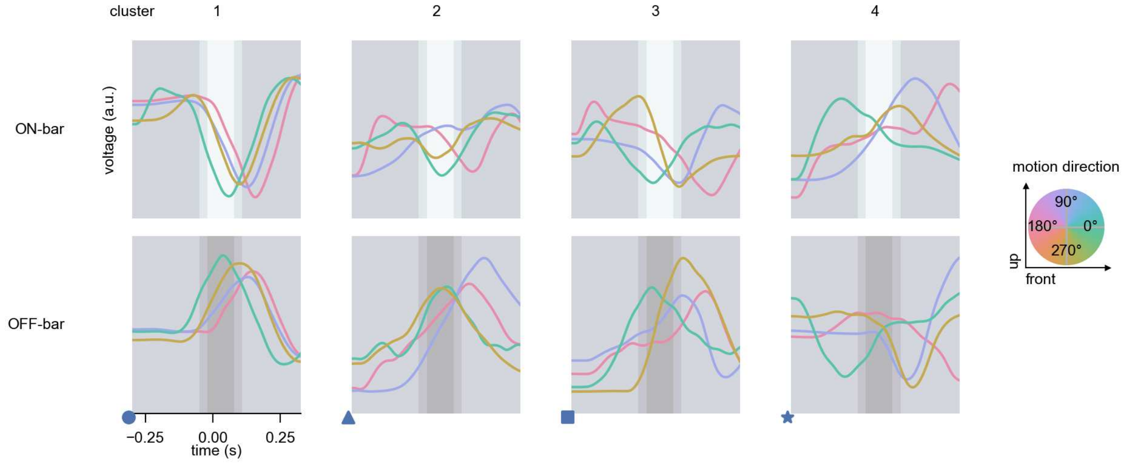

TmY4 - Figure 10: **Responses to moving bars from best-performing models.** The top row shows responses to moving ON-bars, the bottom row shows peak responses to moving OFF-bars. The bar-stimuli move in different directions from 0 to 360 degrees and at different speeds. Before, a grey-stimulus led the network to a stationary state. The columns show responses from the best-task-performing model in the respective cluster. Example responses to bars moving at 75.4°/s in all cardinal directions (green 0°, blue 90°, red 180°, yellow 270°) from -22.5 to 22.5° visual angle. Background contour indicates the bar, passing by the central column at second zero.

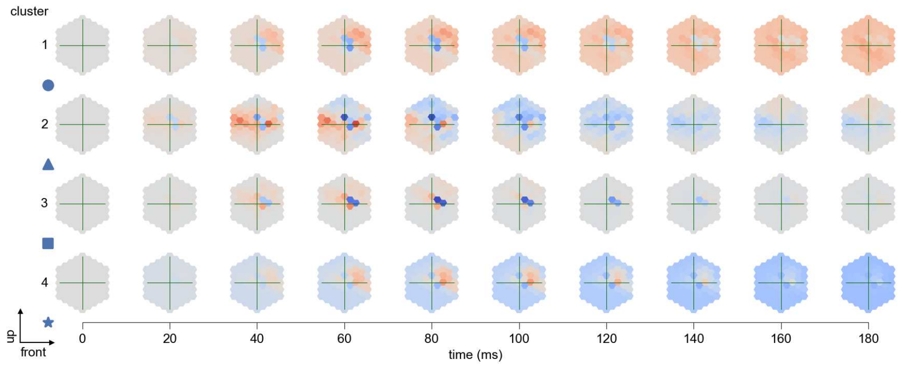

TmY4 - Figure 11: **Spatio-temporal receptive field.** Responses of the central cell to ON-impulses at single ommatidium impulse locations. The impulse occurs at second zero. Before, a grey-stimulus led the network to a stationary state. Responses are baseline-subtracted. Each row illustrates responses from the best-performing model of the respective cluster. Red indicated depolarization, blue indicates hyperpolarization.

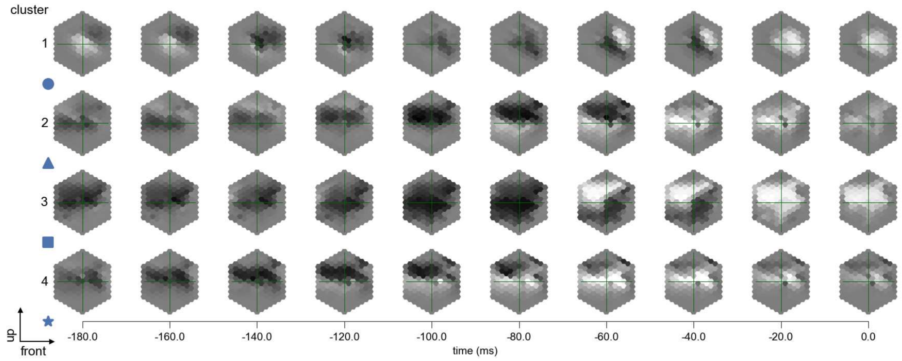

TmY4 - Figure 12: **Maximally excitatory stimuli.** Each row presents the regularized naturalistic-stimulus from the Sintel dataset that maximizes the cell type's central column response at second zero in the best-performing model of the respective cluster.

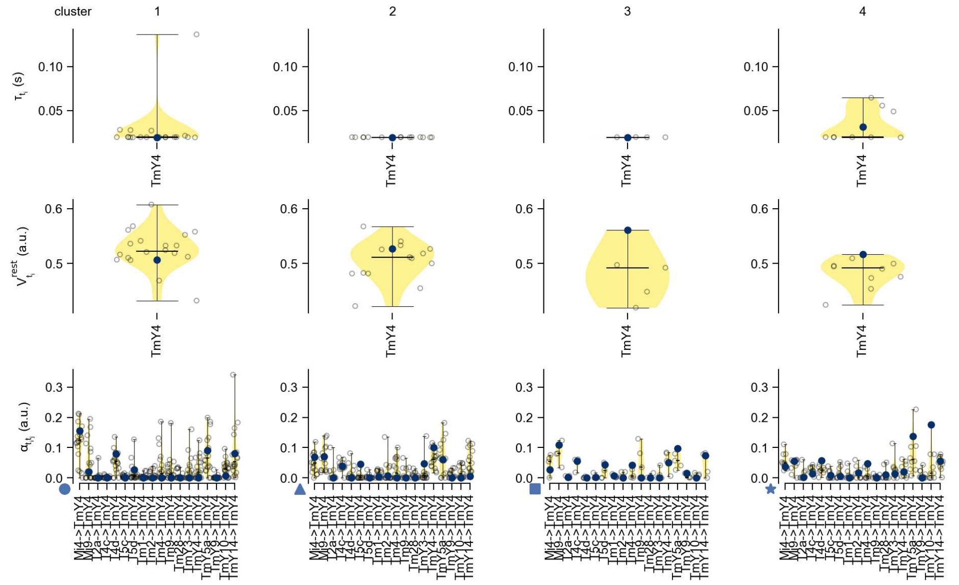

TmY4 - Figure 13: **Task-constrained parameters.** Each column shows the parameters inferred within the respective cluster. First row: learned time constants of the cell type. Second row: resting potentials of the cell type. Third row: scaling factors for the convolutional filters. The blue scatter represents the parameters from the best-performing model within the cluster.

### 59 TmY5a

← Cell types

#### Figures

1 Anatomical receptive fields. . . . 390
2 Anatomical projective fields. . . . 391
3 Clustering of the responses to naturalistic stimuli. . . . 391
4 Responses to flashes. . . . 391
5 Peak responses to moving edges. . . . 392
6 Peak responses to moving edges from best-performing models. . . . 392
7 Responses to moving edges from best-performing models. . . . 393
8 Peak responses to moving bars. . . . 393
9 Peak responses to moving bars from best-performing models. . . . 394
10 Responses to moving bars from best-performing models. . . . 394
11 Spatio-temporal receptive field. . . . 395
12 Maximally excitatory stimuli. . . . 395
13 Task-constrained parameters. . . . 396

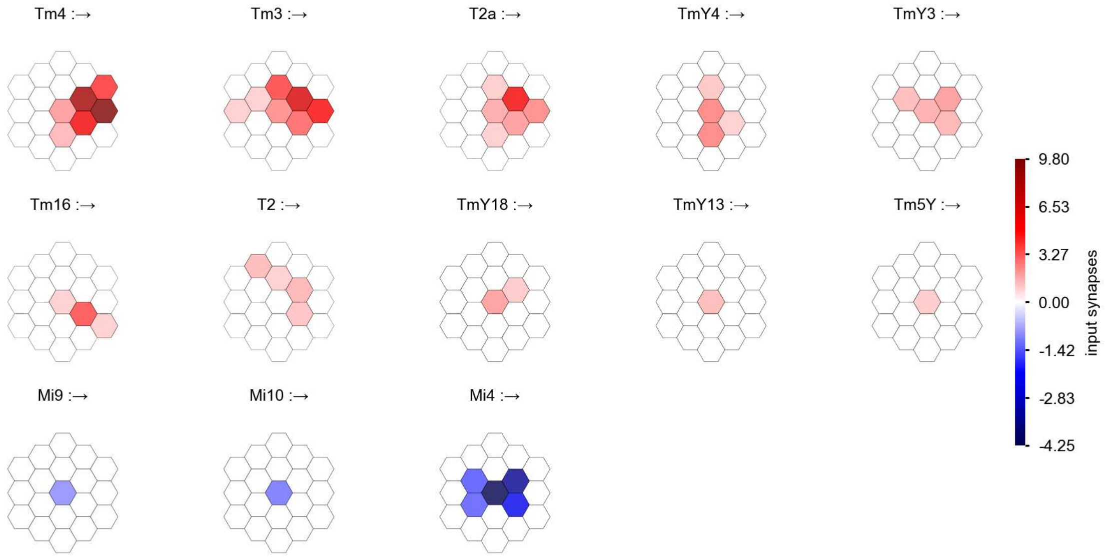

TmY5a - Figure 1: **Anatomical receptive fields.** Each colored hexagon is a projection, with the projection strength characterized by the average number of synapses that we count from the EM reconstruction. Red indicates excitatory synapses, blue indicates inhibitory synapses from inferred signs. Filters in the order of their total number of synapses.

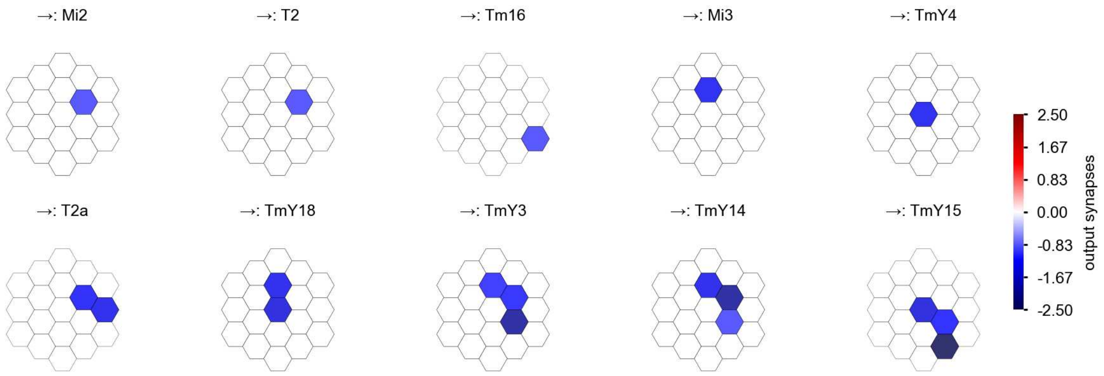

TmY5a - Figure 2: **Anatomical projective fields.** Each colored hexagon is a projection, with the projection strength characterized by the average number of synapses that we count from the EM reconstruction. Red indicates excitatory synapses, blue indicates inhibitory synapses from inferred signs. Filters in the order of their total number of synapses.

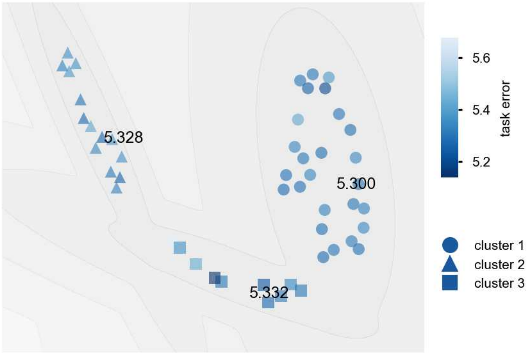

TmY5a - Figure 3: **Clustering of the responses to naturalistic stimuli.** From responses to naturalistic scenes from Sintel. Scatterpoints represent individual models colored by their task error.

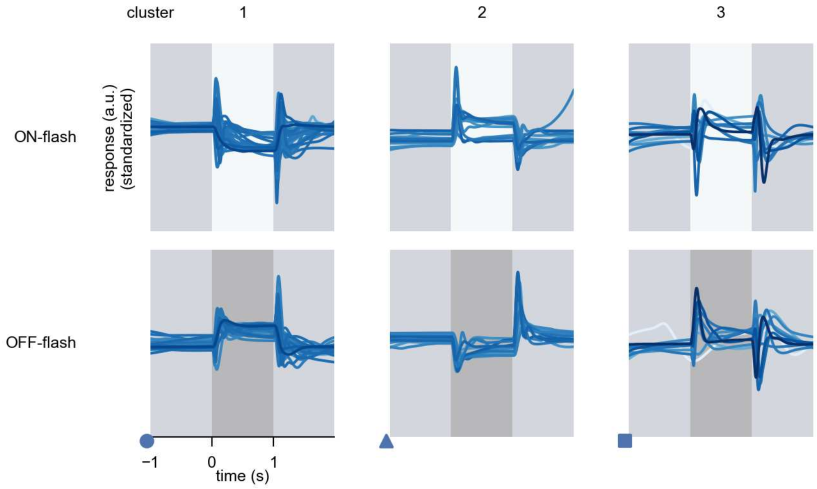

TmY5a - Figure 4: **Responses to flashes.** The top row shows responses to ON-flashes, the bottom row shows responses to OFF-flashes. The responses from the different models in the different clusters (columns) overlay and are colored in the respective task error of the model. The circular flashes occur at second zero, cover 6 ommatidia in radius, and last for one second. Before and after, a grey-stimulus leads to a stationary state of the network. The traces are standardized.

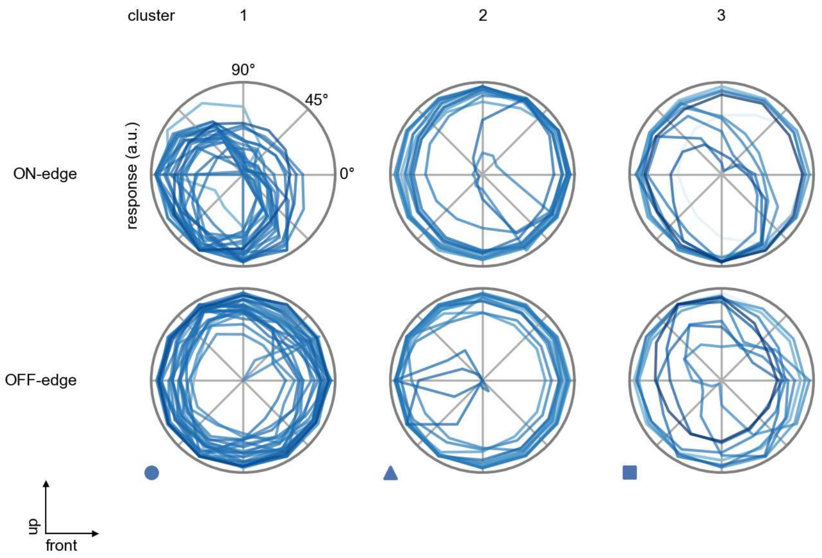

TmY5a - Figure 5: **Peak responses to moving edges.** The top row shows normalized peak responses to moving ON-edges, the bottom row shows normalized peak responses to moving OFF-edges. The edge-stimuli move in different directions from 0 to 360 degrees. Before, a grey-stimulus led the network to a stationary state. The responses from the different models in the different clusters (columns) overlay and are colored in the respective task error of the model. The peak responses are averaged over all assessed edge-speeds.

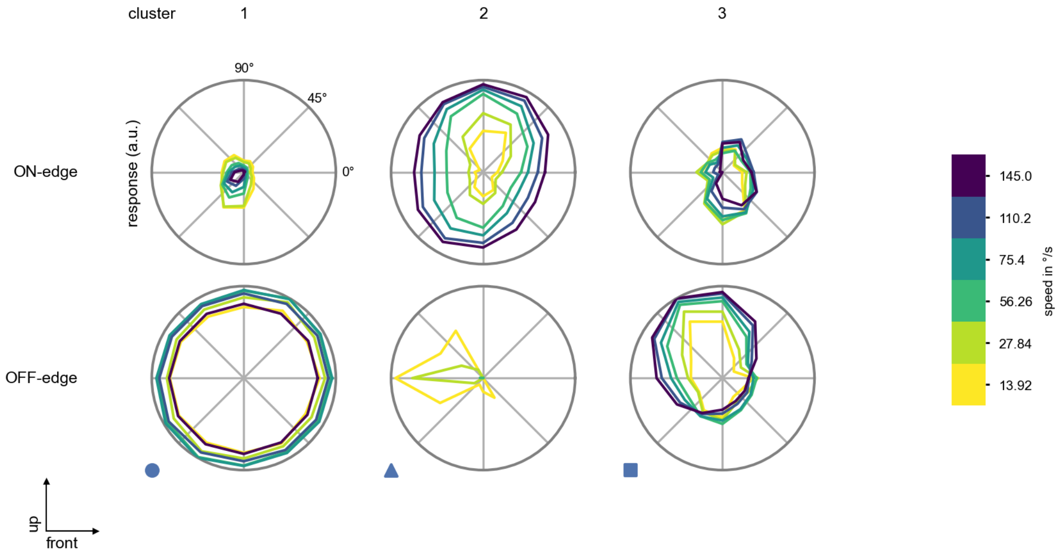

TmY5a - Figure 6: **Peak responses to moving edges from best-performing models.** The top row shows peak responses to moving ON-edges, the bottom row shows peak responses to moving OFF-edges. The edge-stimuli move in different directions from 0 to 360 degrees and at different speeds. Before, a grey-stimulus led the network to a stationary state. The columns show responses from the best-task-performing model in the respective cluster.

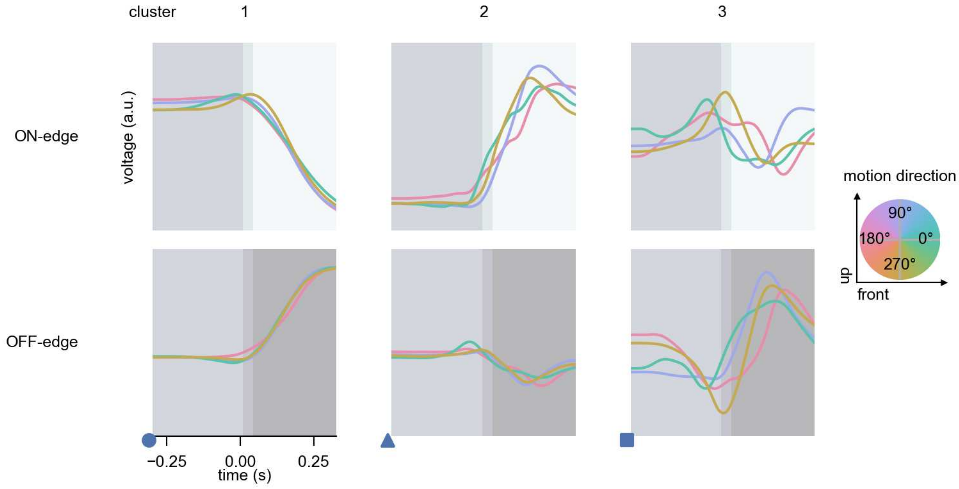

TmY5a - Figure 7: **Responses to moving edges from best-performing models.** The top row shows responses to moving ON-edges, the bottom row shows peak responses to moving OFF-edges. The edge-stimuli move in different directions from 0 to 360 degrees and at different speeds. Before, a grey-stimulus led the network to a stationary state. The columns show responses from the best-task-performing model in the respective cluster. Example responses to edges moving at 75.4°/s in all cardinal directions (green 0°, blue 90°, red 180°, yellow 270°) from -22.5 to 22.5° visual angle. Background contour indicates the edge, passing by the central column at second zero.

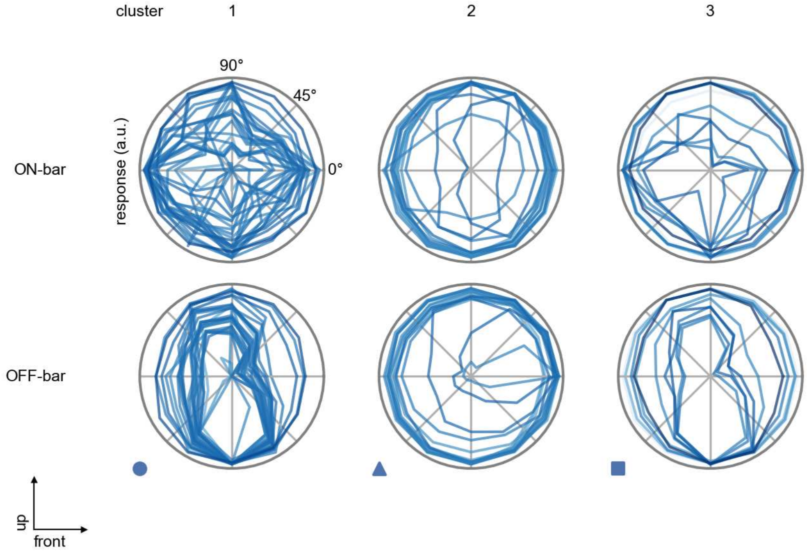

TmY5a - Figure 8: **Peak responses to moving bars.** The top row shows normalized peak responses to moving ON-bars, the bottom row shows normalized peak responses to moving OFF-bars. The bar-stimuli move in different directions from 0 to 360 degrees. Before, a grey-stimulus led the network to a stationary state. The responses from the different models in the different clusters (columns) overlay and are colored in the respective task error of the model. The peak responses are averaged over all assessed bar-speeds.

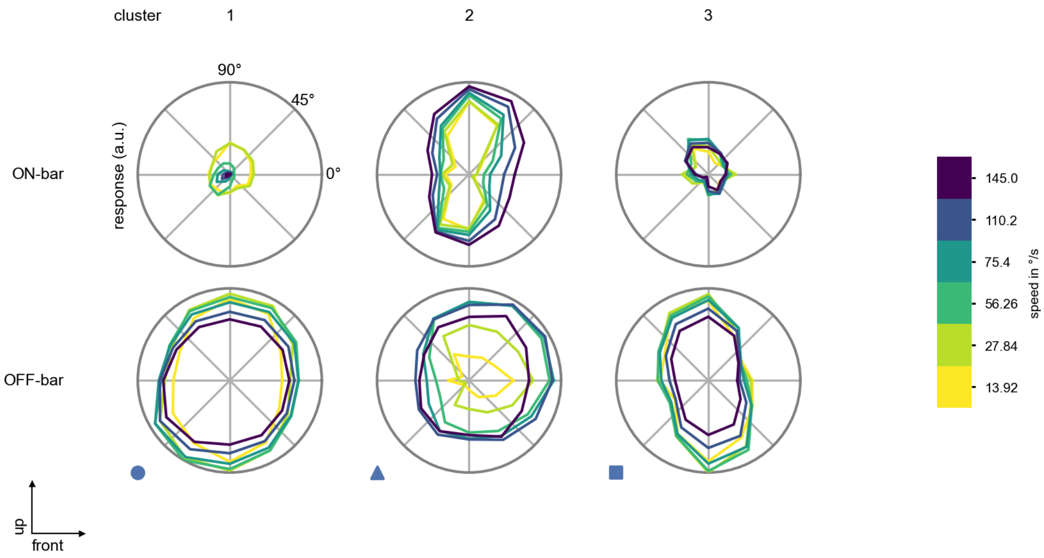

TmY5a - Figure 9: **Peak responses to moving bars from best-performing models.** The top row shows peak responses to moving ON-bars, the bottom row shows peak responses to moving OFF-bars. The bar-stimuli move in different directions from 0 to 360 degrees and at different speeds. Before, a grey-stimulus led the network to a stationary state. The columns show responses from the best-task-performing model in the respective cluster.

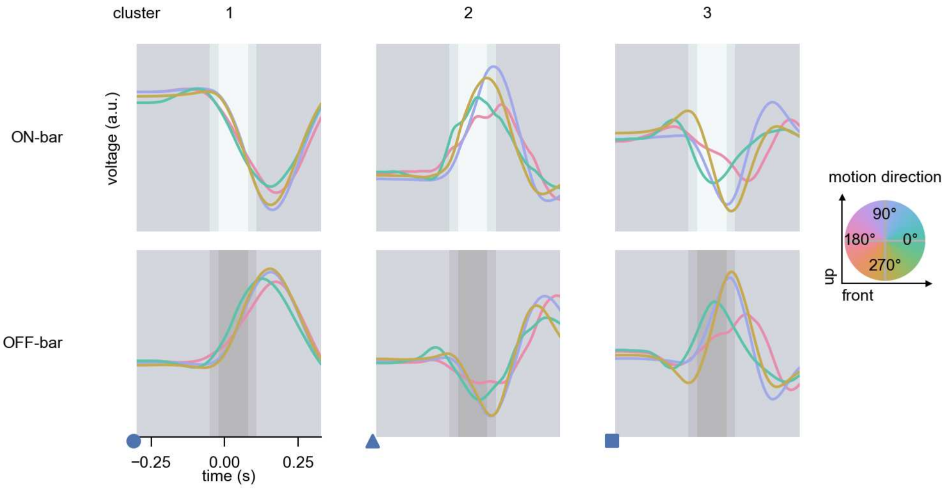

TmY5a - Figure 10: **Responses to moving bars from best-performing models.** The top row shows responses to moving ON-bars, the bottom row shows peak responses to moving OFF-bars. The bar-stimuli move in different directions from 0 to 360 degrees and at different speeds. Before, a grey-stimulus led the network to a stationary state. The columns show responses from the best-task-performing model in the respective cluster. Example responses to bars moving at 75.4°/s in all cardinal directions (green 0°, blue 90°, red 180°, yellow 270°) from -22.5 to 22.5° visual angle. Background contour indicates the bar, passing by the central column at second zero.

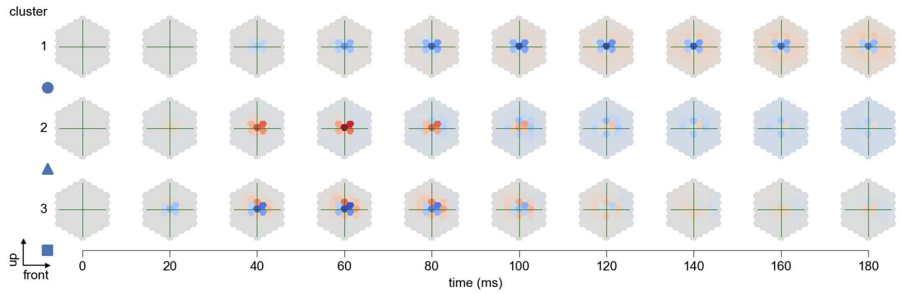

TmY5a - Figure 11: **Spatio-temporal receptive field.** Responses of the central cell to ON-impulses at single ommatidium impulse locations. The impulse occurs at second zero. Before, a grey-stimulus led the network to a stationary state. Responses are baseline-subtracted. Each row illustrates responses from the best-performing model of the respective cluster. Red indicated depolarization, blue indicates hyperpolarization.

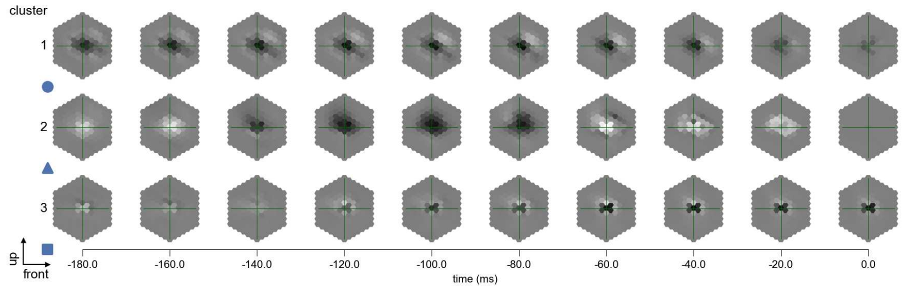

TmY5a - Figure 12: **Maximally excitatory stimuli.** Each row presents the regularized naturalistic-stimulus from the Sintel dataset that maximizes the cell type's central column response at second zero in the best-performing model of the respective cluster.

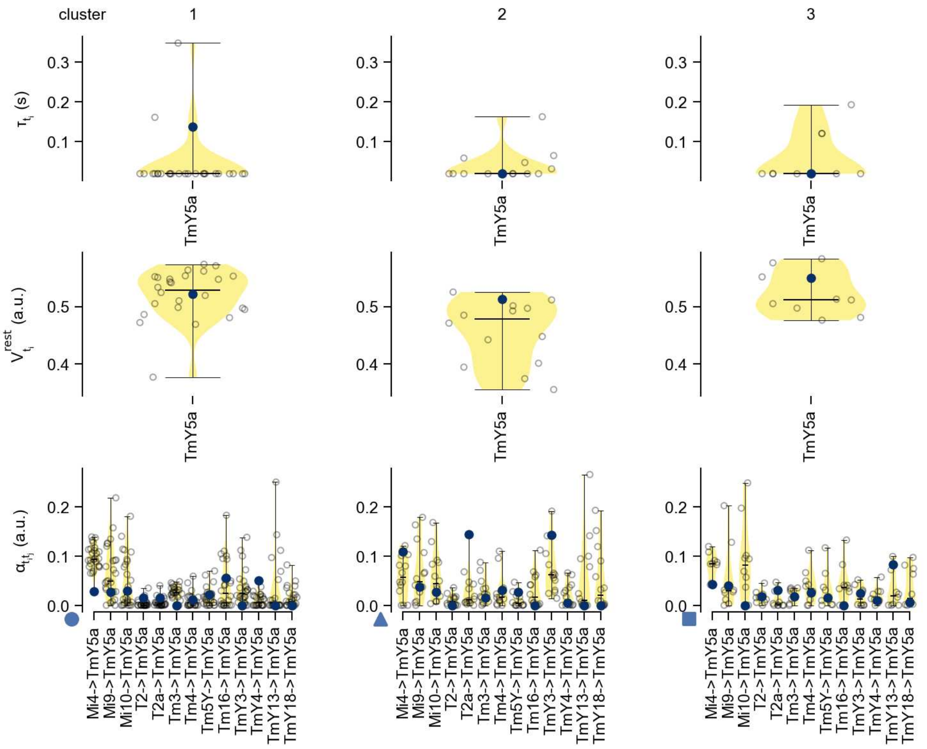

TmY5a - Figure 13: **Task-constrained parameters.** Each column shows the parameters inferred within the respective cluster. First row: learned time constants of the cell type. Second row: resting potentials of the cell type. Third row: scaling factors for the convolutional filters. The blue scatter represents the parameters from the best-performing model within the cluster.

### 60 TmY9

← Cell types

#### Figures

1 Anatomical receptive fields. . . . 397
2 Anatomical projective fields. . . . 397
3 Clustering of the responses to naturalistic stimuli. . . . 398
4 Responses to flashes. . . . 398
5 Peak responses to moving edges. . . . 399
6 Peak responses to moving edges from best-performing models. . . . 399
7 Responses to moving edges from best-performing models. . . . 400
8 Peak responses to moving bars. . . . 400
9 Peak responses to moving bars from best-performing models. . . . 401
10 Responses to moving bars from best-performing models. . . . 401
11 Spatio-temporal receptive field. . . . 402
12 Maximally excitatory stimuli. . . . 402
13 Task-constrained parameters. . . . 402

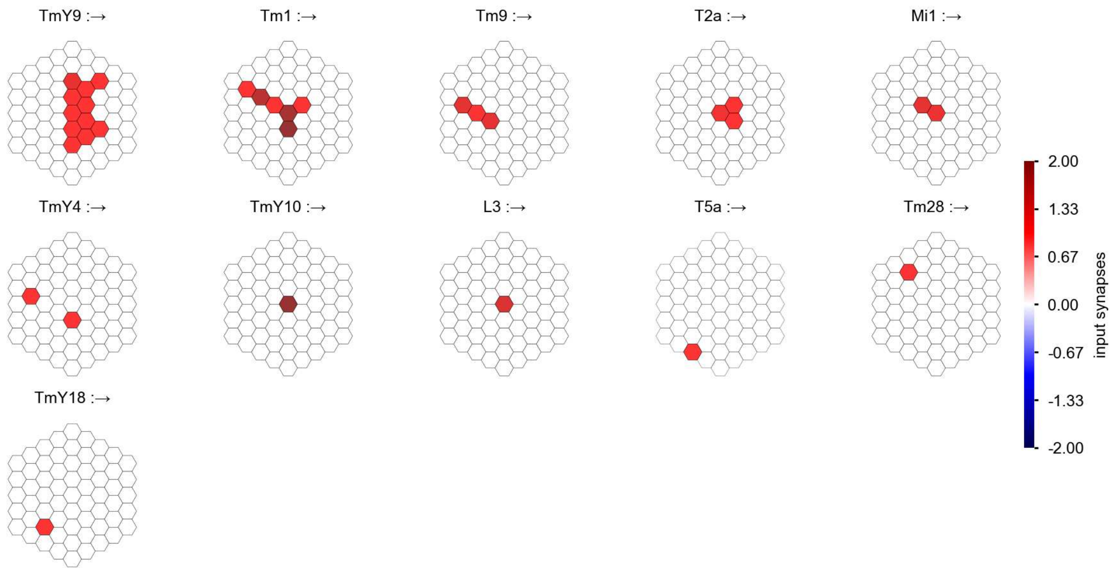

TmY9 - Figure 1: **Anatomical receptive fields.** Each colored hexagon is a projection, with the projection strength characterized by the average number of synapses that we count from the EM reconstruction. Red indicates excitatory synapses, blue indicates inhibitory synapses from inferred signs. Filters in the order of their total number of synapses.

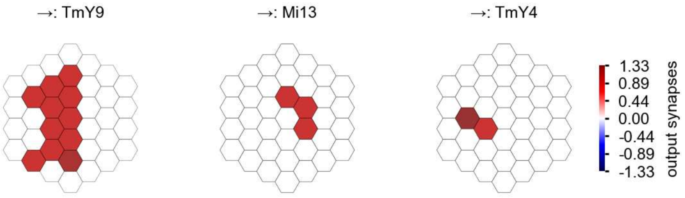

TmY9 - Figure 2: **Anatomical projective fields.** Each colored hexagon is a projection, with the projection strength characterized by the average number of synapses that we count from the EM reconstruction. Red indicates excitatory synapses, blue indicates inhibitory synapses from inferred signs. Filters in the order of their total number of synapses.

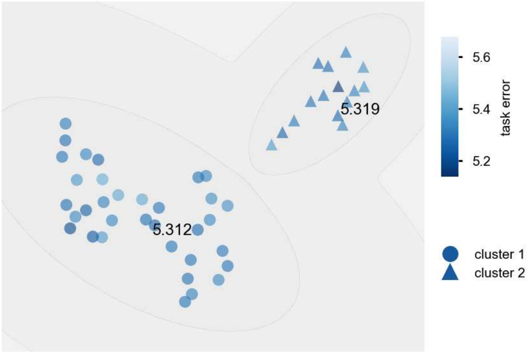

TmY9 - Figure 3: **Clustering of the responses to naturalistic stimuli.** From responses to naturalistic scenes from Sintel. Scatterpoints represent individual models colored by their task error.

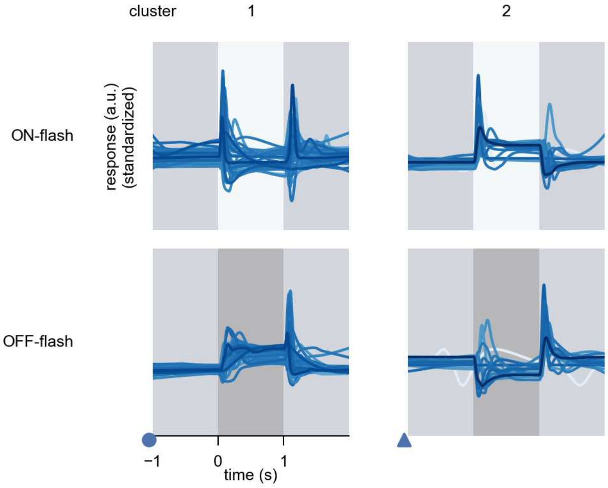

TmY9 - Figure 4: **Responses to flashes.** The top row shows responses to ON-flashes, the bottom row shows responses to OFF-flashes. The responses from the different models in the different clusters (columns) overlay and are colored in the respective task error of the model. The circular flashes occur at second zero, cover 6 ommatidia in radius, and last for one second. Before and after, a grey-stimulus leads to a stationary state of the network. The traces are standardized.

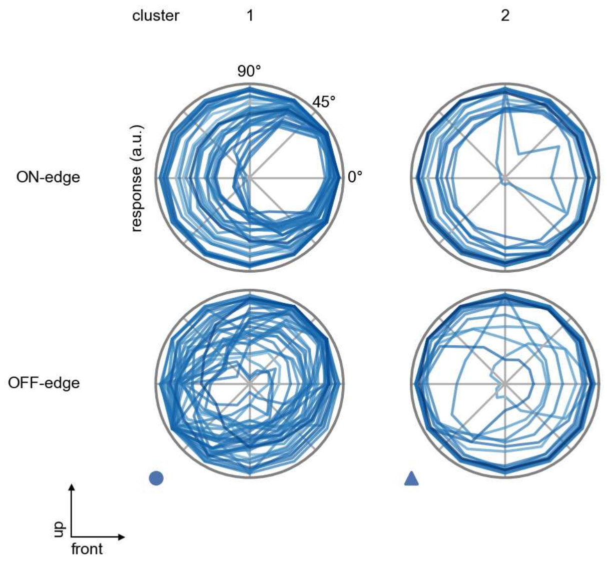

TmY9 - Figure 5: **Peak responses to moving edges.** The top row shows normalized peak responses to moving ON-edges, the bottom row shows normalized peak responses to moving OFF-edges. The edge-stimuli move in different directions from 0 to 360 degrees. Before, a grey-stimulus led the network to a stationary state. The responses from the different models in the different clusters (columns) overlay and are colored in the respective task error of the model. The peak responses are averaged over all assessed edge-speeds.

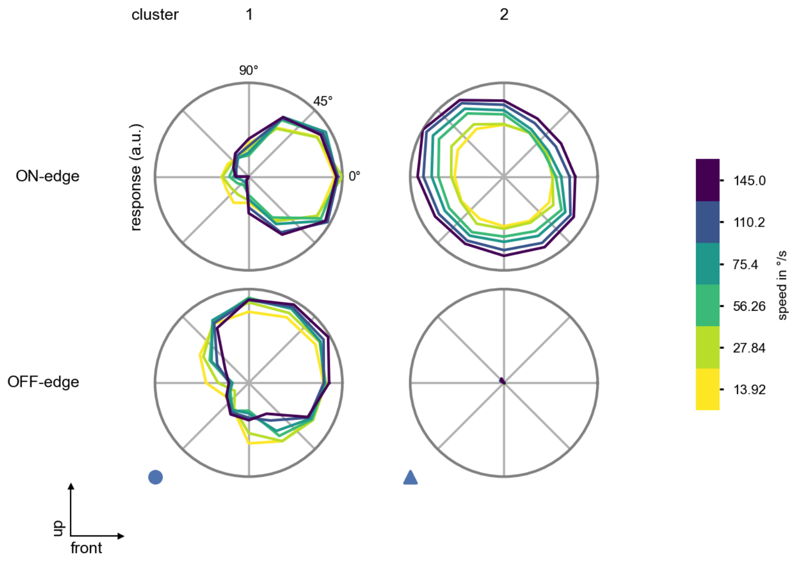

TmY9 - Figure 6: **Peak responses to moving edges from best-performing models.** The top row shows peak responses to moving ON-edges, the bottom row shows peak responses to moving OFF-edges. The edge-stimuli move in different directions from 0 to 360 degrees and at different speeds. Before, a grey-stimulus led the network to a stationary state. The columns show responses from the best-task-performing model in the respective cluster.

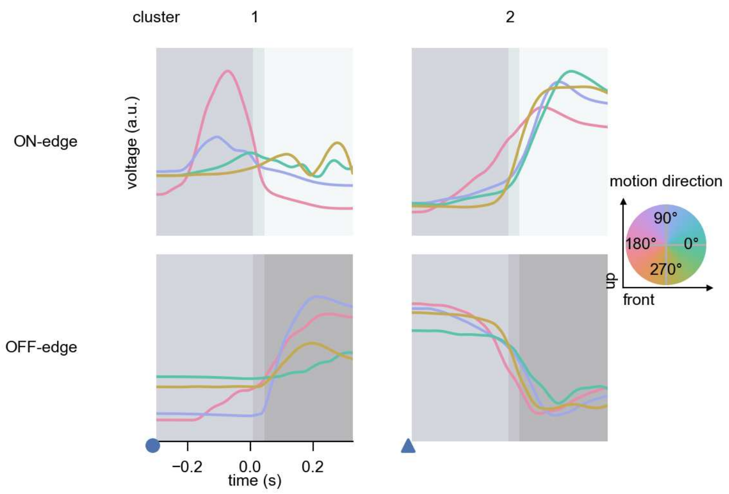

TmY9 - Figure 7: **Responses to moving edges from best-performing models.** The top row shows responses to moving ON-edges, the bottom row shows peak responses to moving OFF-edges. The edge-stimuli move in different directions from 0 to 360 degrees and at different speeds. Before, a grey-stimulus led the network to a stationary state. The columns show responses from the best-task-performing model in the respective cluster. Example responses to edges moving at 75.4°/s in all cardinal directions (green 0°, blue 90°, red 180°, yellow 270°) from -22.5 to 22.5° visual angle. Background contour indicates the edge, passing by the central column at second zero.

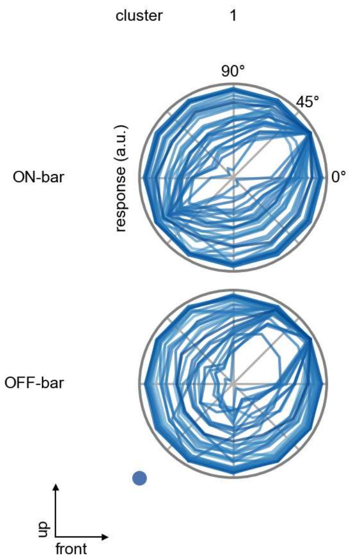

TmY9 - Figure 8: **Peak responses to moving bars.** The top row shows normalized peak responses to moving ON-bars, the bottom row shows normalized peak responses to moving OFF-bars. The bar-stimuli move in different directions from 0 to 360 degrees. Before, a grey-stimulus led the network to a stationary state. The responses from the different models in the different clusters (columns) overlay and are colored in the respective task error of the model. The peak responses are averaged over all assessed bar-speeds.

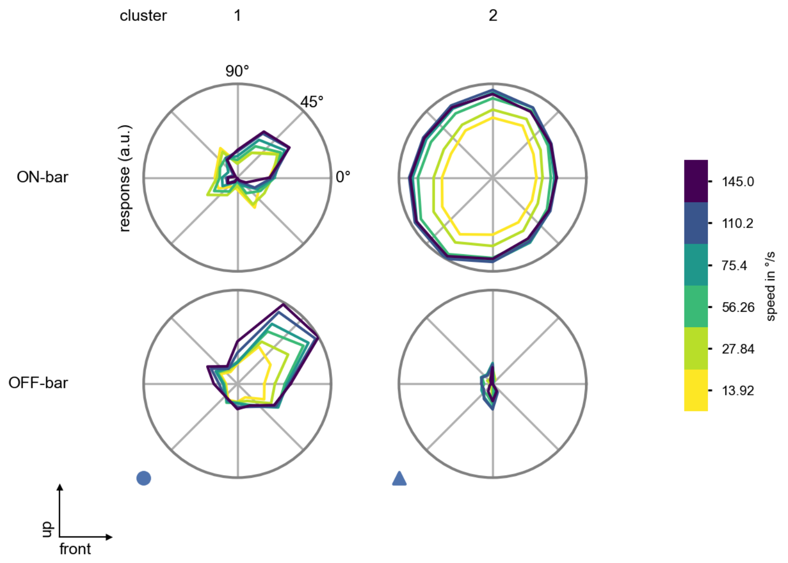

TmY9 - Figure 9: **Peak responses to moving bars from best-performing models.** The top row shows peak responses to moving ON-bars, the bottom row shows peak responses to moving OFF-bars. The bar-stimuli move in different directions from 0 to 360 degrees and at different speeds. Before, a grey-stimulus led the network to a stationary state. The columns show responses from the best-task-performing model in the respective cluster.

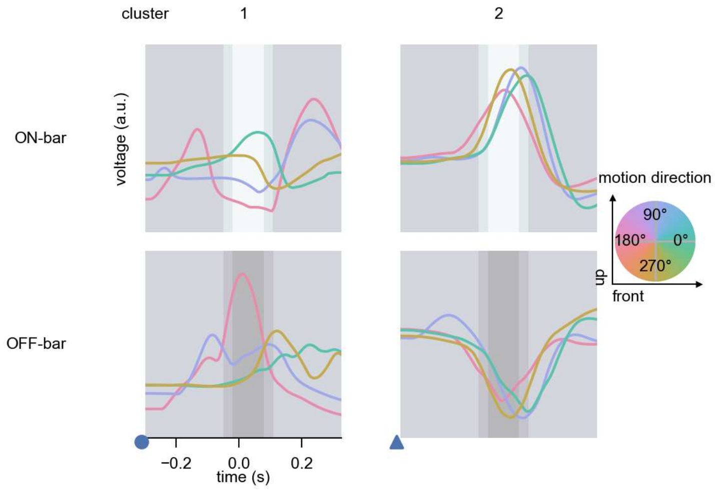

TmY9 - Figure 10: **Responses to moving bars from best-performing models.** The top row shows responses to moving ON-bars, the bottom row shows peak responses to moving OFF-bars. The bar-stimuli move in different directions from 0 to 360 degrees and at different speeds. Before, a grey-stimulus led the network to a stationary state. The columns show responses from the best-task-performing model in the respective cluster. Example responses to bars moving at 75.4°/s in all cardinal directions (green 0°, blue 90°, red 180°, yellow 270°) from -22.5 to 22.5° visual angle. Background contour indicates the bar, passing by the central column at second zero.

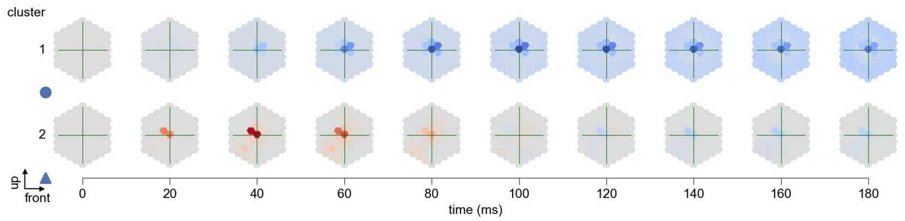

TmY9 - Figure 11: **Spatio-temporal receptive field.** Responses of the central cell to ON-impulses at single ommatidium impulse locations. The impulse occurs at second zero. Before, a grey-stimulus led the network to a stationary state. Responses are baseline-subtracted. Each row illustrates responses from the best-performing model of the respective cluster. Red indicated depolarization, blue indicates hyperpolarization.

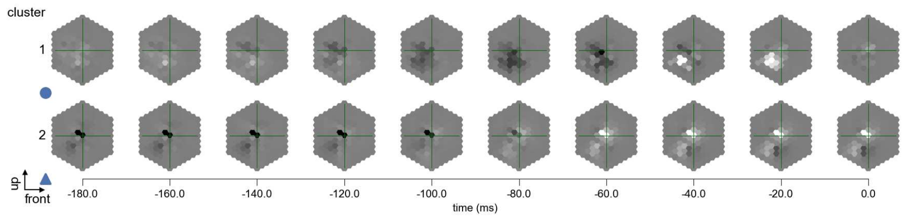

TmY9 - Figure 12: **Maximally excitatory stimuli.** Each row presents the regularized naturalistic-stimulus from the Sintel dataset that maximizes the cell type's central column response at second zero in the best-performing model of the respective cluster.

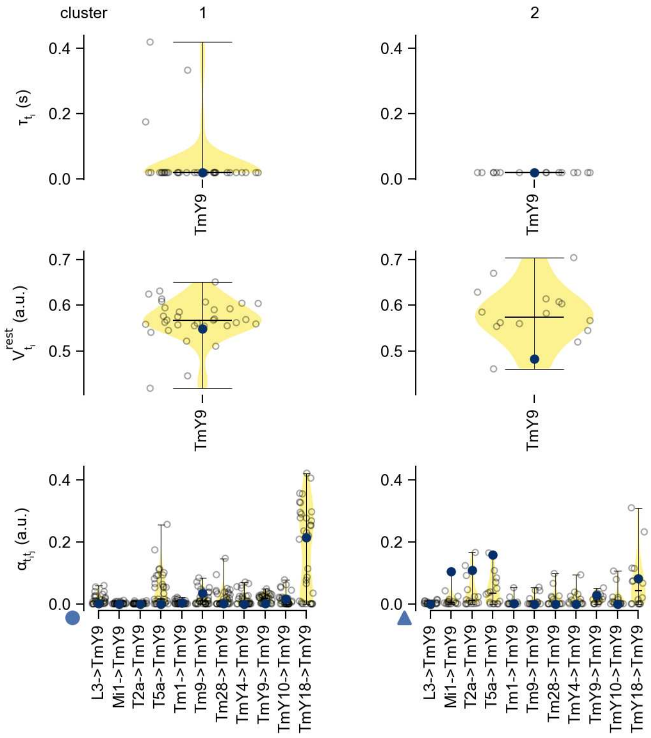

TmY9 - Figure 13: **Task-constrained parameters.** Each column shows the parameters inferred within the respective cluster. First row: learned time constants of the cell type. Second row: resting potentials of the cell type. Third row: scaling factors for the convolutional filters. The blue scatter represents the parameters from the best-performing model within the cluster.

### 61 TmY10

← Cell types

#### Figures

1 Anatomical receptive fields. . . . 403
2 Anatomical projective fields. . . . 403
3 Clustering of the responses to naturalistic stimuli. . . . 404
4 Responses to flashes. . . . 404
5 Peak responses to moving edges. . . . 405
6 Peak responses to moving edges from best-performing models. . . . 405
7 Responses to moving edges from best-performing models. . . . 406
8 Peak responses to moving bars. . . . 406
9 Peak responses to moving bars from best-performing models. . . . 407
10 Responses to moving bars from best-performing models. . . . 407
11 Spatio-temporal receptive field. . . . 408
12 Maximally excitatory stimuli. . . . 408
13 Task-constrained parameters. . . . 408

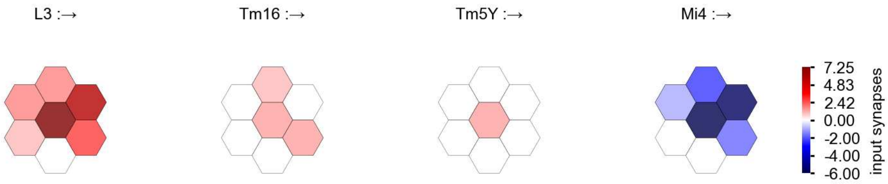

TmY10 - Figure 1: **Anatomical receptive fields.** Each colored hexagon is a projection, with the projection strength characterized by the average number of synapses that we count from the EM reconstruction. Red indicates excitatory synapses, blue indicates inhibitory synapses from inferred signs. Filters in the order of their total number of synapses.

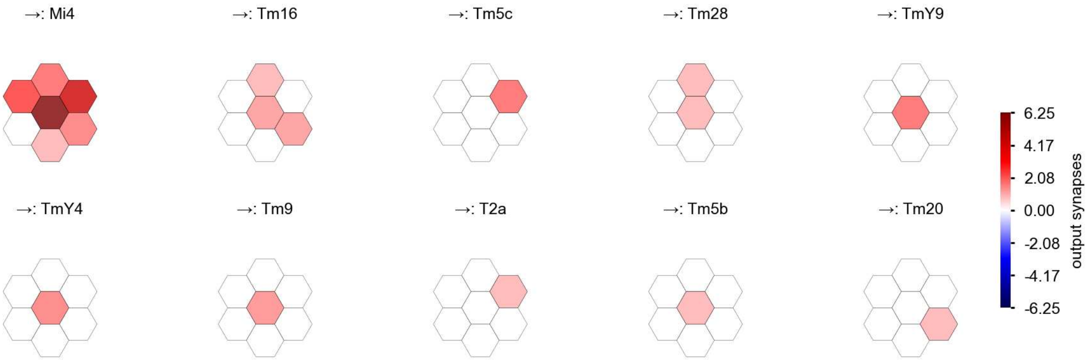

TmY10 - Figure 2: **Anatomical projective fields.** Each colored hexagon is a projection, with the projection strength characterized by the average number of synapses that we count from the EM reconstruction. Red indicates excitatory synapses, blue indicates inhibitory synapses from inferred signs. Filters in the order of their total number of synapses.

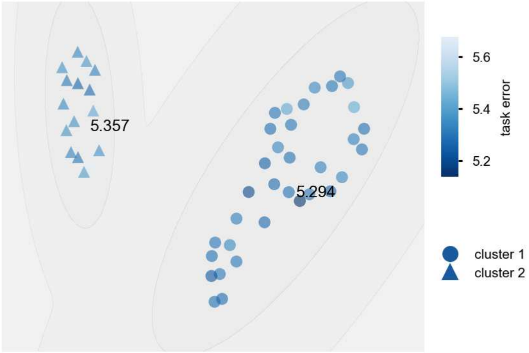

TmY10 - Figure 3: **Clustering of the responses to naturalistic stimuli.** From responses to naturalistic scenes from Sintel. Scatterpoints represent individual models colored by their task error.

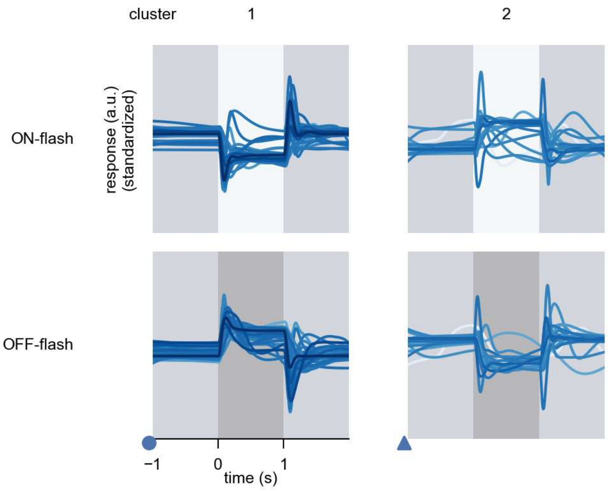

TmY10 - Figure 4: **Responses to flashes.** The top row shows responses to ON-flashes, the bottom row shows responses to OFF-flashes. The responses from the different models in the different clusters (columns) overlay and are colored in the respective task error of the model. The circular flashes occur at second zero, cover 6 ommatidia in radius, and last for one second. Before and after, a grey-stimulus leads to a stationary state of the network. The traces are standardized.

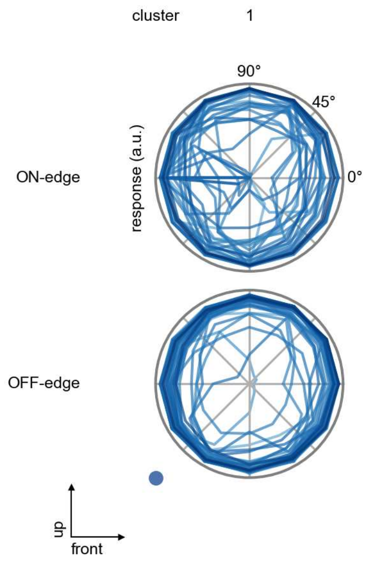

TmY10 - Figure 5: **Peak responses to moving edges.** The top row shows normalized peak responses to moving ON-edges, the bottom row shows normalized peak responses to moving OFF-edges. The edge-stimuli move in different directions from 0 to 360 degrees. Before, a grey-stimulus led the network to a stationary state. The responses from the different models in the different clusters (columns) overlay and are colored in the respective task error of the model. The peak responses are averaged over all assessed edge-speeds.

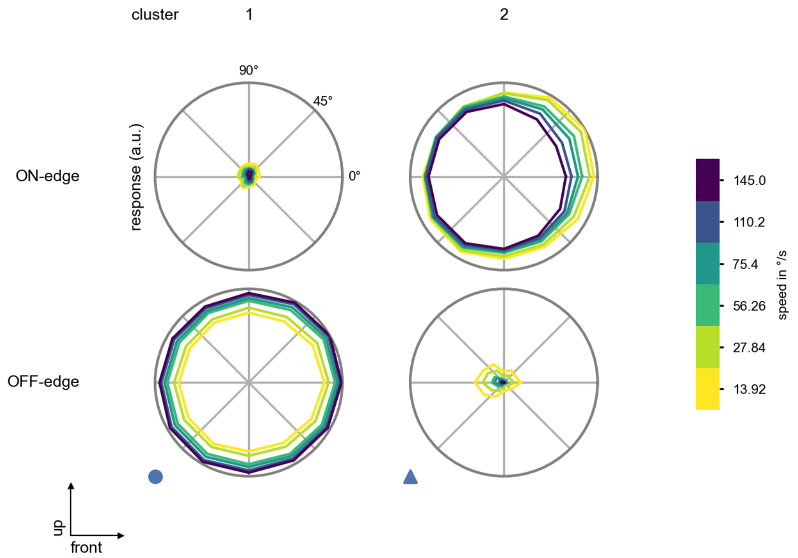

TmY10 - Figure 6: **Peak responses to moving edges from best-performing models.** The top row shows peak responses to moving ON-edges, the bottom row shows peak responses to moving OFF-edges. The edge-stimuli move in different directions from 0 to 360 degrees and at different speeds. Before, a grey-stimulus led the network to a stationary state. The columns show responses from the best-task-performing model in the respective cluster.

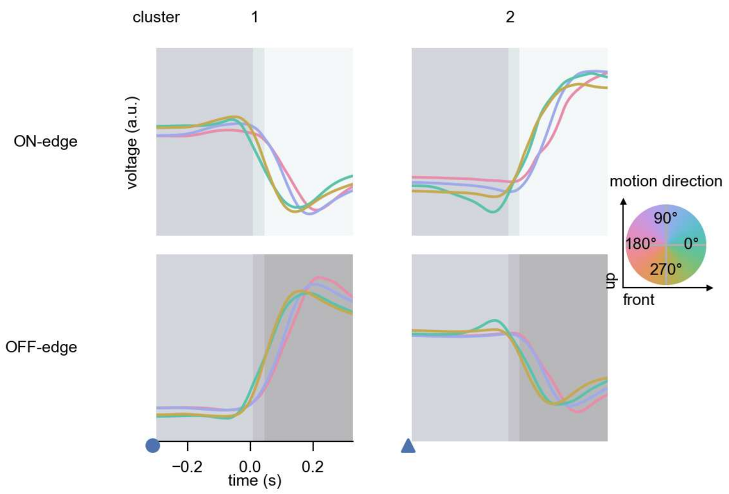

TmY10 - Figure 7: **Responses to moving edges from best-performing models.** The top row shows responses to moving ON-edges, the bottom row shows peak responses to moving OFF-edges. The edge-stimuli move in different directions from 0 to 360 degrees and at different speeds. Before, a grey-stimulus led the network to a stationary state. The columns show responses from the best-task-performing model in the respective cluster. Example responses to edges moving at 75.4°/s in all cardinal directions (green 0°, blue 90°, red 180°, yellow 270°) from -22.5 to 22.5° visual angle. Background contour indicates the edge, passing by the central column at second zero.

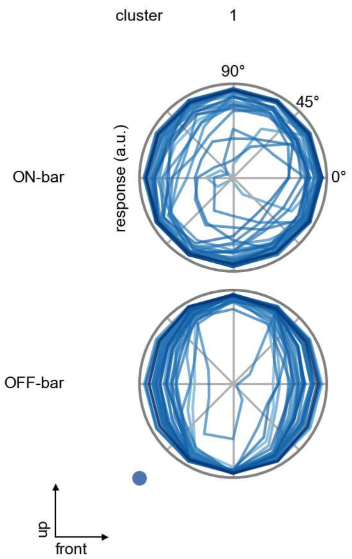

TmY10 - Figure 8: **Peak responses to moving bars.** The top row shows normalized peak responses to moving ON-bars, the bottom row shows normalized peak responses to moving OFF-bars. The bar-stimuli move in different directions from 0 to 360 degrees. Before, a grey-stimulus led the network to a stationary state. The responses from the different models in the different clusters (columns) overlay and are colored in the respective task error of the model. The peak responses are averaged over all assessed bar-speeds.

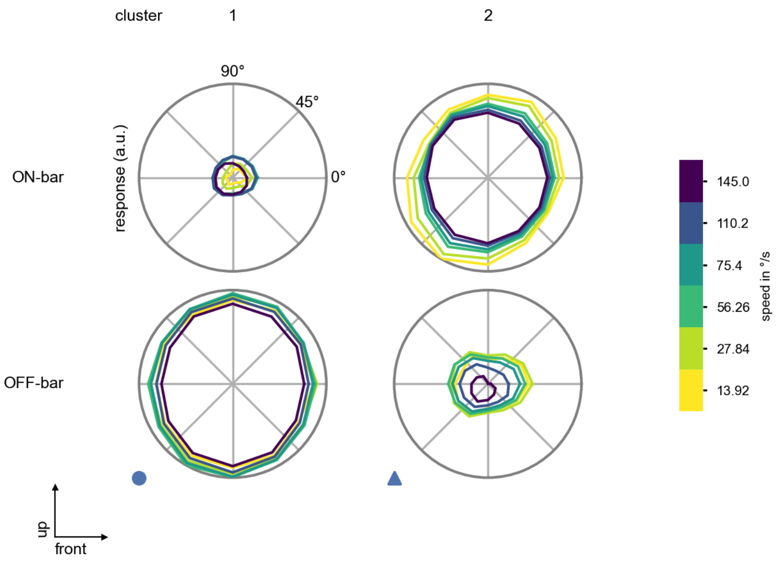

TmY10 - Figure 9: **Peak responses to moving bars from best-performing models.** The top row shows peak responses to moving ON-bars, the bottom row shows peak responses to moving OFF-bars. The bar-stimuli move in different directions from 0 to 360 degrees and at different speeds. Before, a grey-stimulus led the network to a stationary state. The columns show responses from the best-task-performing model in the respective cluster.

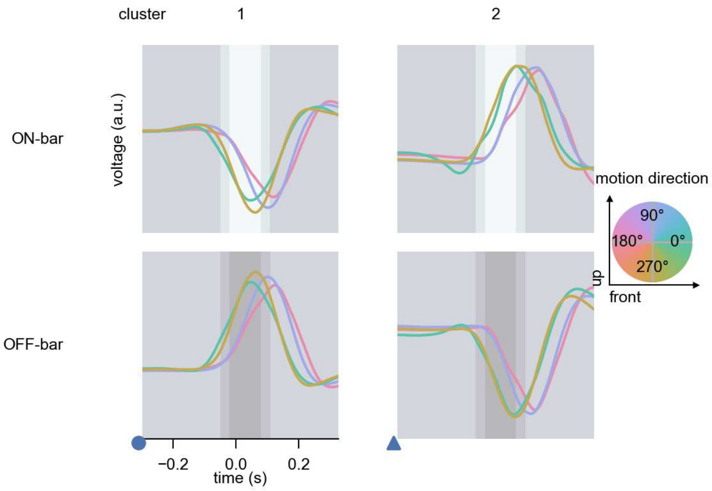

TmY10 - Figure 10: **Responses to moving bars from best-performing models.** The top row shows responses to moving ON-bars, the bottom row shows peak responses to moving OFF-bars. The bar-stimuli move in different directions from 0 to 360 degrees and at different speeds. Before, a grey-stimulus led the network to a stationary state. The columns show responses from the best-task-performing model in the respective cluster. Example responses to bars moving at 75.4°/s in all cardinal directions (green 0°, blue 90°, red 180°, yellow 270°) from -22.5 to 22.5° visual angle. Background contour indicates the bar, passing by the central column at second zero.

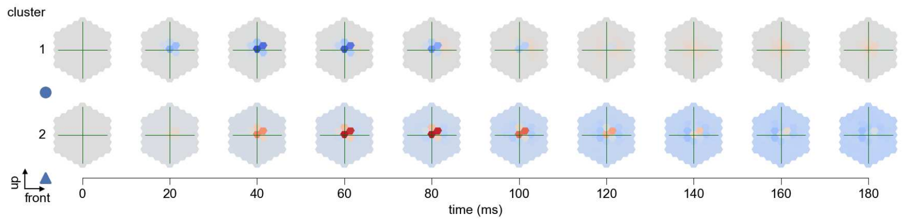

TmY10 - Figure 11: **Spatio-temporal receptive field.** Responses of the central cell to ON-impulses at single ommatidium impulse locations. The impulse occurs at second zero. Before, a grey-stimulus led the network to a stationary state. Responses are baseline-subtracted. Each row illustrates responses from the best-performing model of the respective cluster. Red indicated depolarization, blue indicates hyperpolarization.

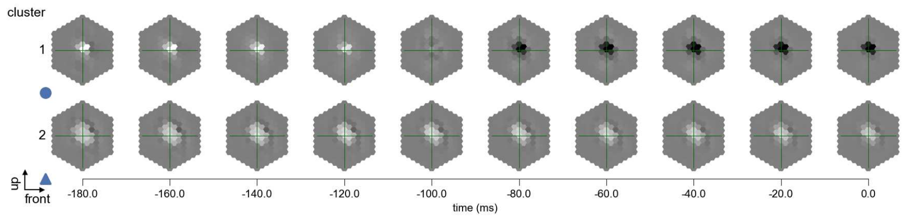

TmY10 - Figure 12: **Maximally excitatory stimuli.** Each row presents the regularized naturalistic-stimulus from the Sintel dataset that maximizes the cell type's central column response at second zero in the best-performing model of the respective cluster.

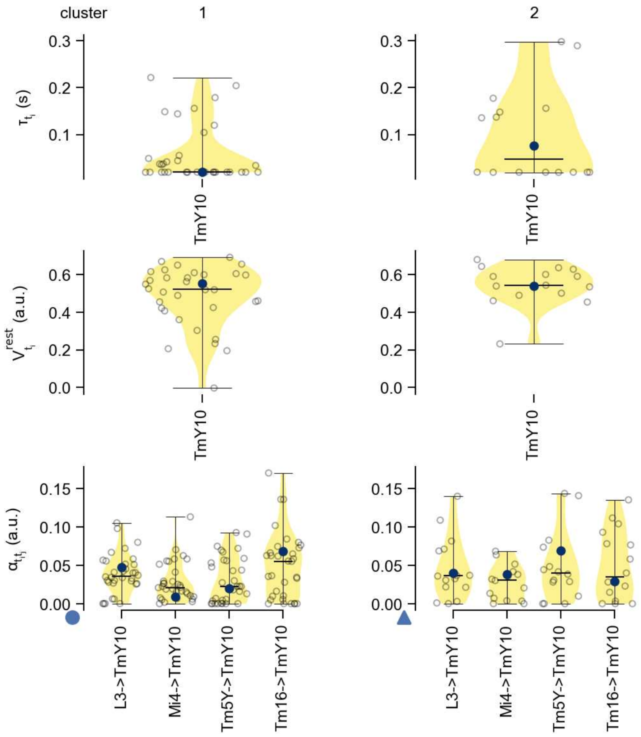

TmY10 - Figure 13: **Task-constrained parameters.** Each column shows the parameters inferred within the respective cluster. First row: learned time constants of the cell type. Second row: resting potentials of the cell type. Third row: scaling factors for the convolutional filters. The blue scatter represents the parameters from the best-performing model within the cluster.

### 62 TmY13

← Cell types

#### Figures

1 Anatomical receptive fields. . . . 409
2 Anatomical projective fields. . . . 409
3 Clustering of the responses to naturalistic stimuli. . . . 410
4 Responses to flashes. . . . 410
5 Peak responses to moving edges. . . . 411
6 Peak responses to moving edges from best-performing models. . . . 411
7 Responses to moving edges from best-performing models. . . . 412
8 Peak responses to moving bars. . . . 412
9 Peak responses to moving bars from best-performing models. . . . 413
10 Responses to moving bars from best-performing models. . . . 413
11 Spatio-temporal receptive field. . . . 414
12 Maximally excitatory stimuli. . . . 414
13 Task-constrained parameters. . . . 415

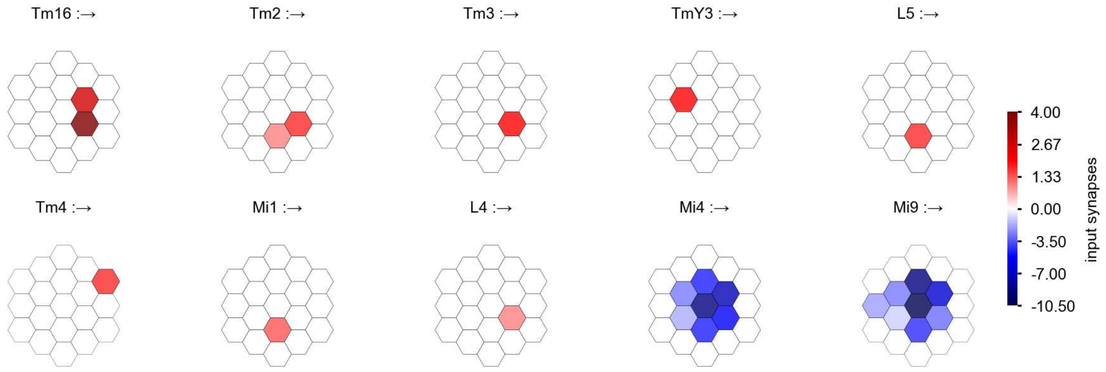

TmY13 - Figure 1: **Anatomical receptive fields.** Each colored hexagon is a projection, with the projection strength characterized by the average number of synapses that we count from the EM reconstruction. Red indicates excitatory synapses, blue indicates inhibitory synapses from inferred signs. Filters in the order of their total number of synapses.

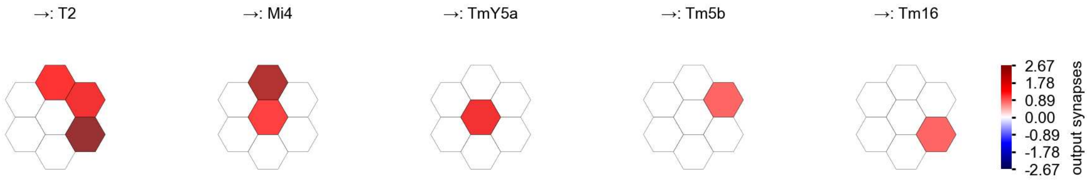

TmY13 - Figure 2: **Anatomical projective fields.** Each colored hexagon is a projection, with the projection strength characterized by the average number of synapses that we count from the EM reconstruction. Red indicates excitatory synapses, blue indicates inhibitory synapses from inferred signs. Filters in the order of their total number of synapses.

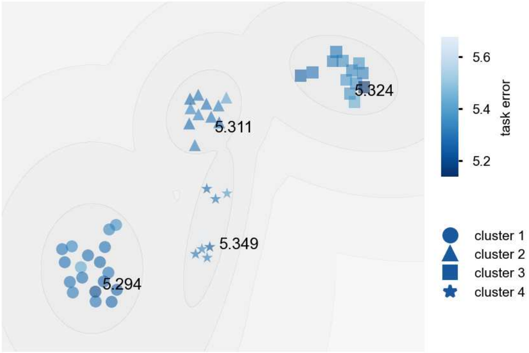

TmY13 - Figure 3: **Clustering of the responses to naturalistic stimuli.** From responses to naturalistic scenes from Sintel. Scatterpoints represent individual models colored by their task error.

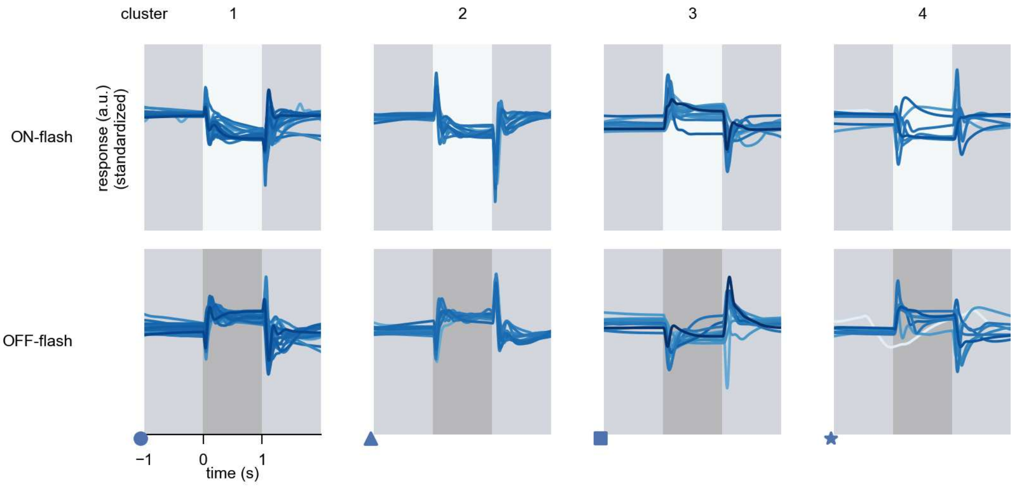

TmY13 - Figure 4: **Responses to flashes.** The top row shows responses to ON-flashes, the bottom row shows responses to OFF-flashes. The responses from the different models in the different clusters (columns) overlay and are colored in the respective task error of the model. The circular flashes occur at second zero, cover 6 ommatidia in radius, and last for one second. Before and after, a grey-stimulus leads to a stationary state of the network. The traces are standardized.

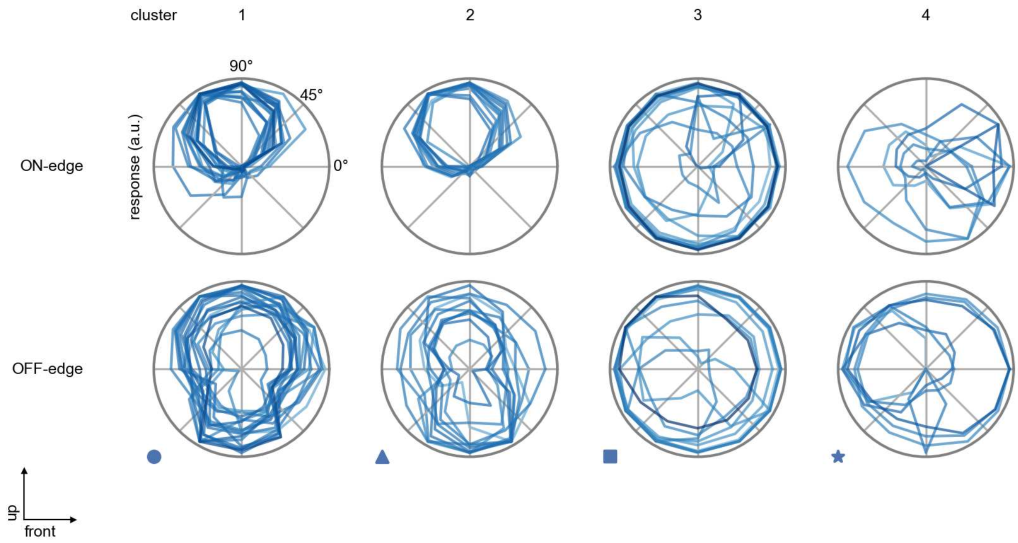

TmY13 - Figure 5: **Peak responses to moving edges.** The top row shows normalized peak responses to moving ON-edges, the bottom row shows normalized peak responses to moving OFF-edges. The edge-stimuli move in different directions from 0 to 360 degrees. Before, a grey-stimulus led the network to a stationary state. The responses from the different models in the different clusters (columns) overlay and are colored in the respective task error of the model. The peak responses are averaged over all assessed edge-speeds.

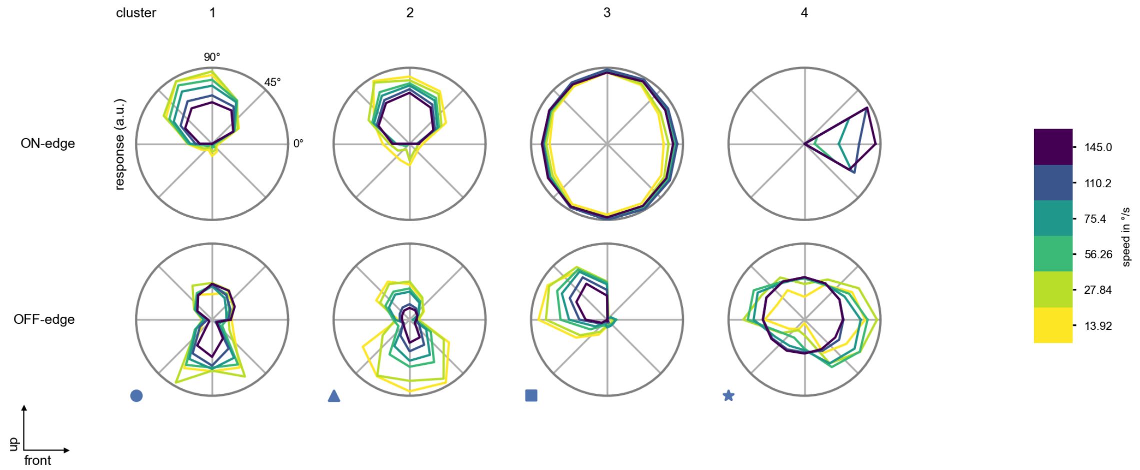

TmY13 - Figure 6: **Peak responses to moving edges from best-performing models.** The top row shows peak responses to moving ON-edges, the bottom row shows peak responses to moving OFF-edges. The edge-stimuli move in different directions from 0 to 360 degrees and at different speeds. Before, a grey-stimulus led the network to a stationary state. The columns show responses from the best-task-performing model in the respective cluster.

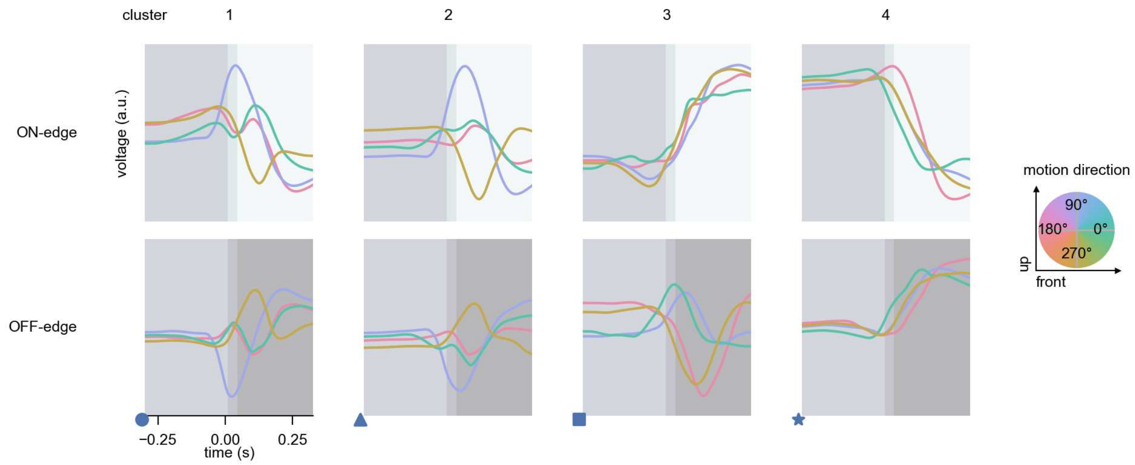

TmY13 - Figure 7: **Responses to moving edges from best-performing models.** The top row shows responses to moving ON-edges, the bottom row shows peak responses to moving OFF-edges. The edge-stimuli move in different directions from 0 to 360 degrees and at different speeds. Before, a grey-stimulus led the network to a stationary state. The columns show responses from the best-task-performing model in the respective cluster. Example responses to edges moving at 75.4°/s in all cardinal directions (green 0°, blue 90°, red 180°, yellow 270°) from -22.5 to 22.5° visual angle. Background contour indicates the edge, passing by the central column at second zero.

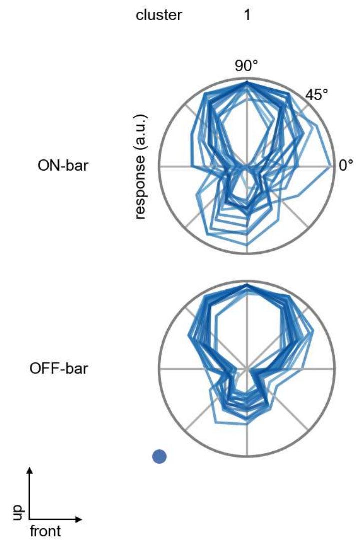

TmY13 - Figure 8: **Peak responses to moving bars.** The top row shows normalized peak responses to moving ON-bars, the bottom row shows normalized peak responses to moving OFF-bars. The bar-stimuli move in different directions from 0 to 360 degrees. Before, a grey-stimulus led the network to a stationary state. The responses from the different models in the different clusters (columns) overlay and are colored in the respective task error of the model. The peak responses are averaged over all assessed bar-speeds.

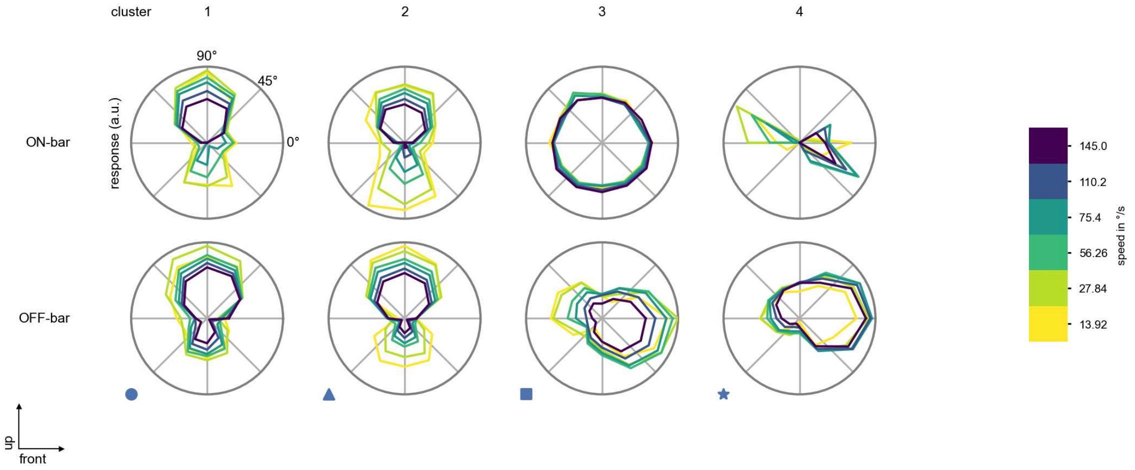

TmY13 - Figure 9: **Peak responses to moving bars from best-performing models.** The top row shows peak responses to moving ON-bars, the bottom row shows peak responses to moving OFF-bars. The bar-stimuli move in different directions from 0 to 360 degrees and at different speeds. Before, a grey-stimulus led the network to a stationary state. The columns show responses from the best-task-performing model in the respective cluster.

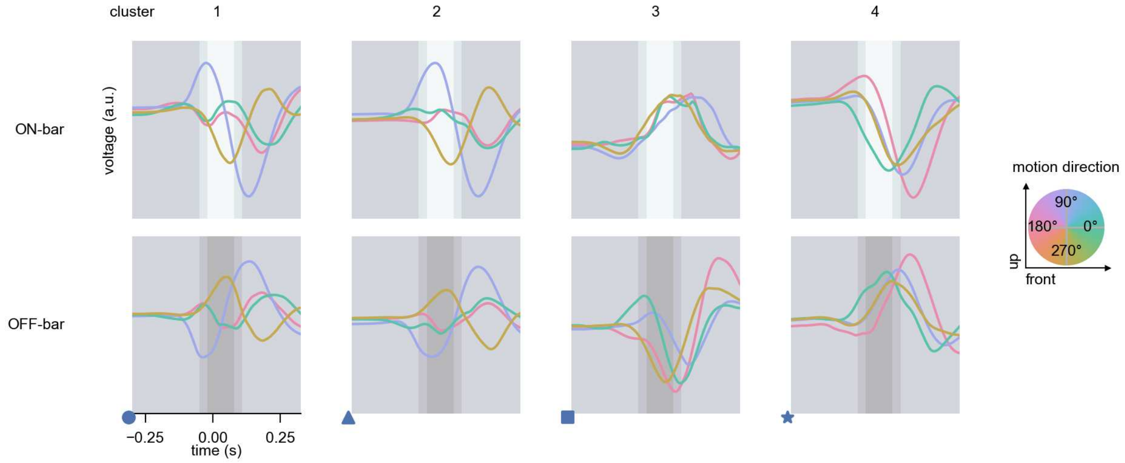

TmY13 - Figure 10: **Responses to moving bars from best-performing models.** The top row shows responses to moving ON-bars, the bottom row shows peak responses to moving OFF-bars. The bar-stimuli move in different directions from 0 to 360 degrees and at different speeds. Before, a grey-stimulus led the network to a stationary state. The columns show responses from the best-task-performing model in the respective cluster. Example responses to bars moving at 75.4°/s in all cardinal directions (green 0°, blue 90°, red 180°, yellow 270°) from -22.5 to 22.5° visual angle. Background contour indicates the bar, passing by the central column at second zero.

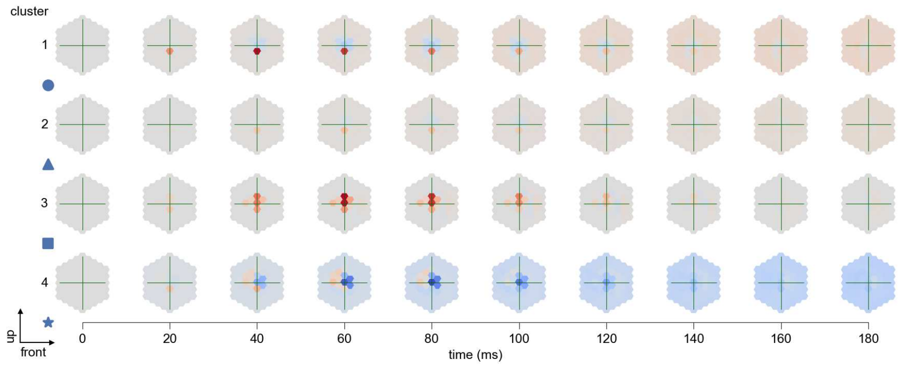

TmY13 - Figure 11: **Spatio-temporal receptive field.** Responses of the central cell to ON-impulses at single ommatidium impulse locations. The impulse occurs at second zero. Before, a grey-stimulus led the network to a stationary state. Responses are baseline-subtracted. Each row illustrates responses from the best-performing model of the respective cluster. Red indicated depolarization, blue indicates hyperpolarization.

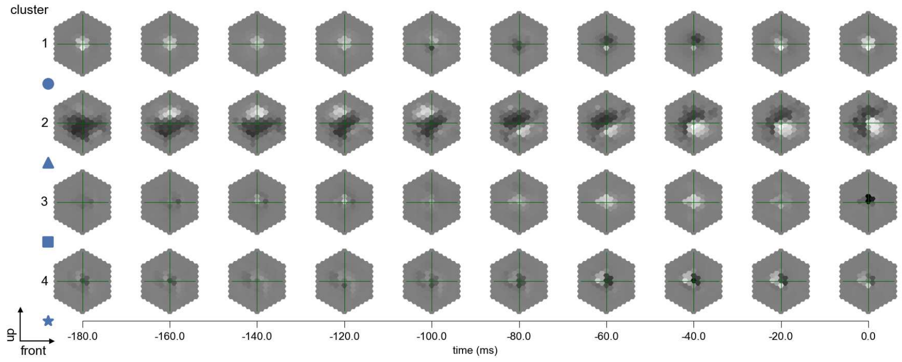

TmY13 - Figure 12: **Maximally excitatory stimuli.** Each row presents the regularized naturalistic-stimulus from the Sintel dataset that maximizes the cell type's central column response at second zero in the best-performing model of the respective cluster.

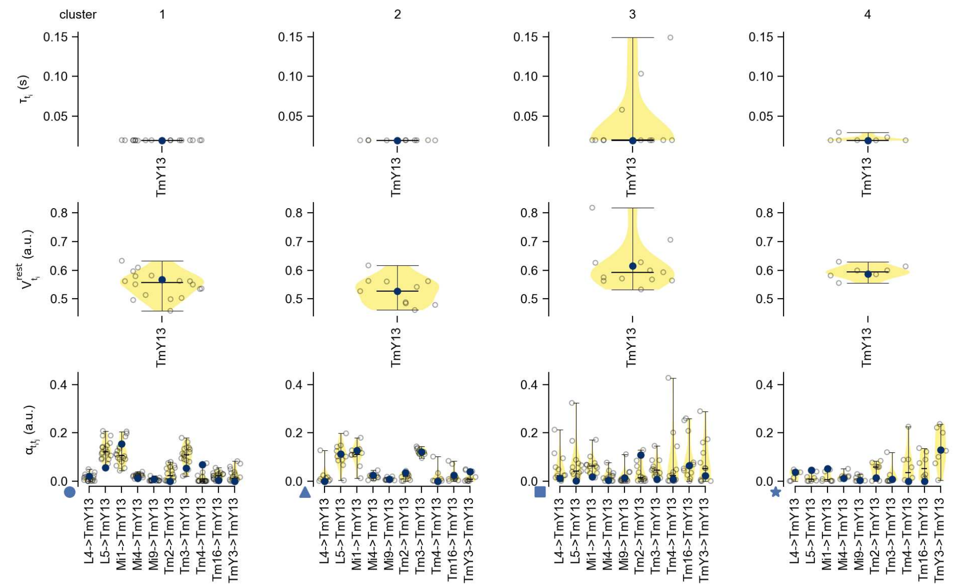

TmY13 - Figure 13: **Task-constrained parameters.** Each column shows the parameters inferred within the respective cluster. First row: learned time constants of the cell type. Second row: resting potentials of the cell type. Third row: scaling factors for the convolutional filters. The blue scatter represents the parameters from the best-performing model within the cluster.

### 63 TmY14

← Cell types

#### Figures

1 Anatomical receptive fields. . . . . 416
2 Anatomical projective fields. . . . . 417
3 Clustering of the responses to naturalistic stimuli. . . . . 417
4 Responses to flashes. . . . . 417
5 Peak responses to moving edges. . . . . 418
6 Peak responses to moving edges from best-performing models. . . . . 418
7 Responses to moving edges from best-performing models. . . . . 419
8 Peak responses to moving bars. . . . . 419
9 Peak responses to moving bars from best-performing models. . . . . 420
10 Responses to moving bars from best-performing models. . . . . 420
11 Spatio-temporal receptive field. . . . . 421
12 Maximally excitatory stimuli. . . . . 421
13 Task-constrained parameters. . . . . 422

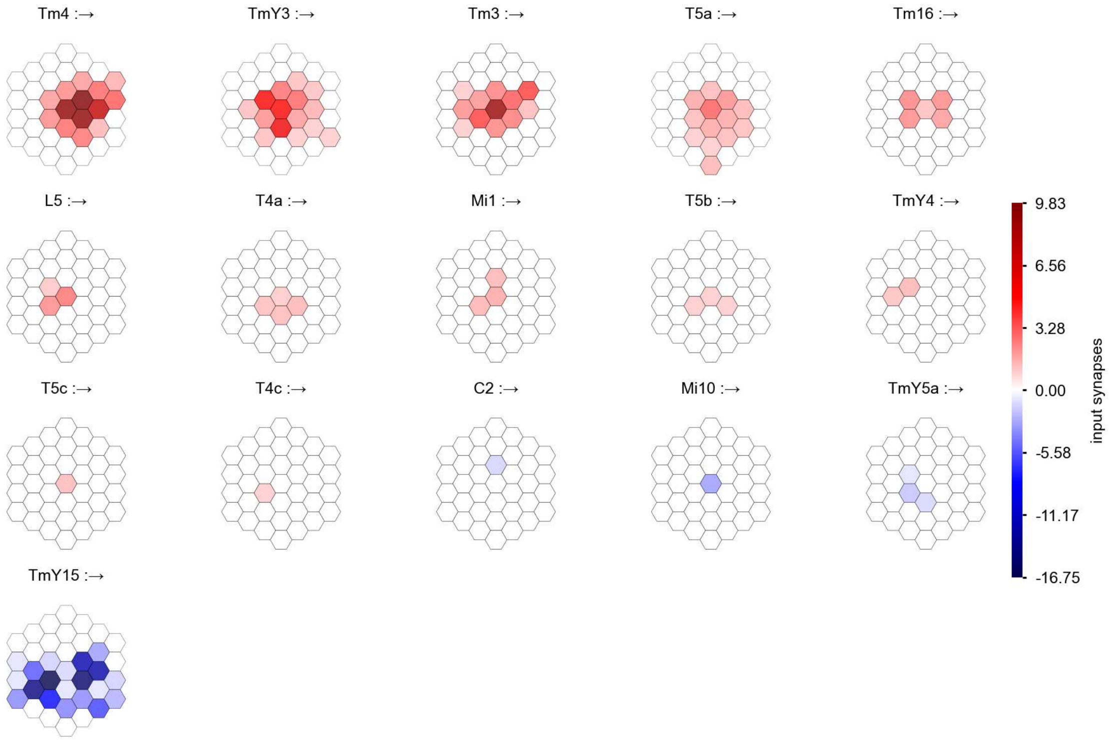

TmY14 - Figure 1: **Anatomical receptive fields.** Each colored hexagon is a projection, with the projection strength characterized by the average number of synapses that we count from the EM reconstruction. Red indicates excitatory synapses, blue indicates inhibitory synapses from inferred signs. Filters in the order of their total number of synapses.

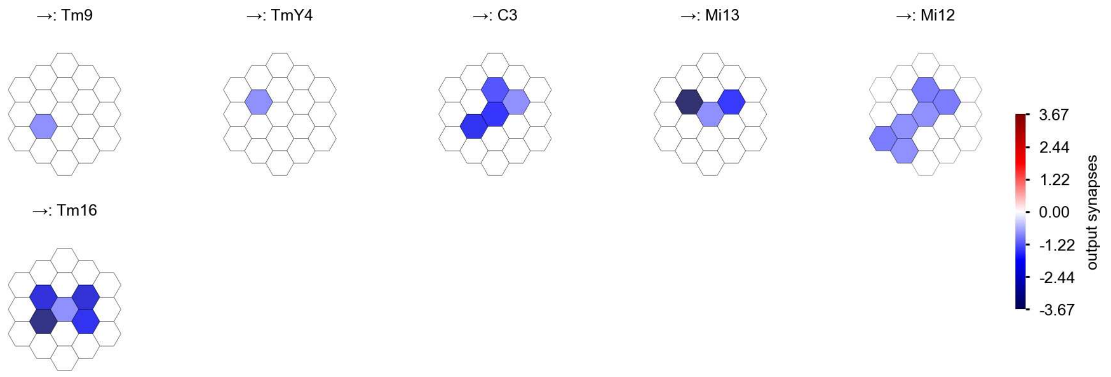

TmY14 - Figure 2: **Anatomical projective fields.** Each colored hexagon is a projection, with the projection strength characterized by the average number of synapses that we count from the EM reconstruction. Red indicates excitatory synapses, blue indicates inhibitory synapses from inferred signs. Filters in the order of their total number of synapses.

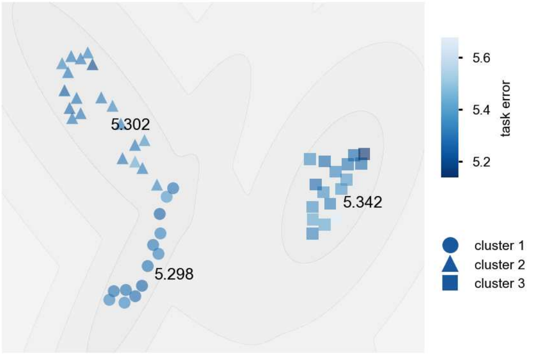

TmY14 - Figure 3: **Clustering of the responses to naturalistic stimuli.** From responses to naturalistic scenes from Sintel. Scatterpoints represent individual models colored by their task error.

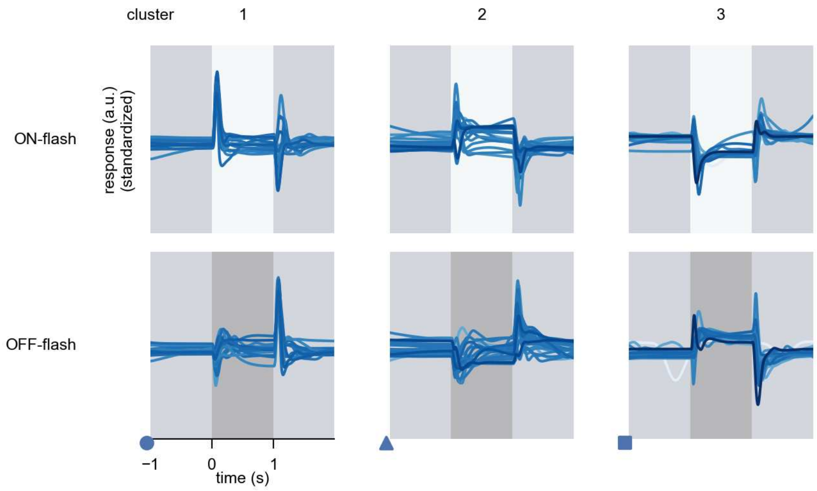

TmY14 - Figure 4: **Responses to flashes.** The top row shows responses to ON-flashes, the bottom row shows responses to OFF-flashes. The responses from the different models in the different clusters (columns) overlay and are colored in the respective task error of the model. The circular flashes occur at second zero, cover 6 ommatidia in radius, and last for one second. Before and after, a grey-stimulus leads to a stationary state of the network. The traces are standardized.

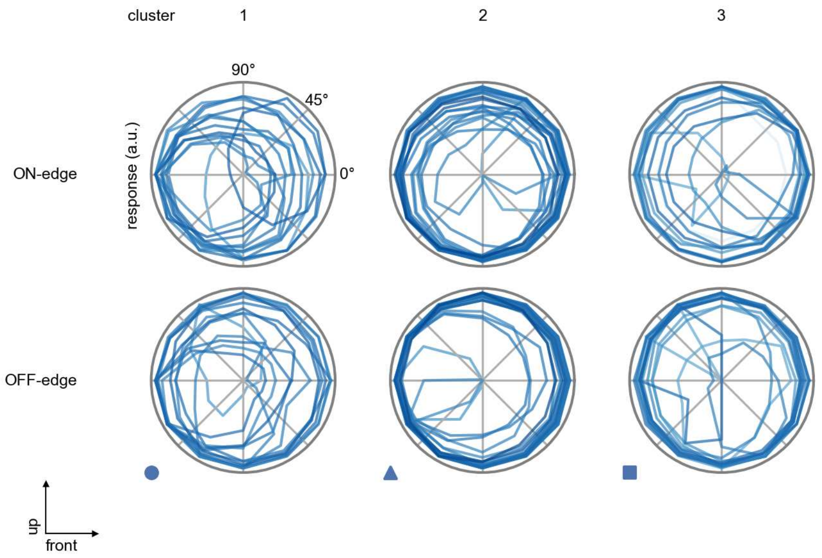

TmY14 - Figure 5: **Peak responses to moving edges.** The top row shows normalized peak responses to moving ON-edges, the bottom row shows normalized peak responses to moving OFF-edges. The edge-stimuli move in different directions from 0 to 360 degrees. Before, a grey-stimulus led the network to a stationary state. The responses from the different models in the different clusters (columns) overlay and are colored in the respective task error of the model. The peak responses are averaged over all assessed edge-speeds.

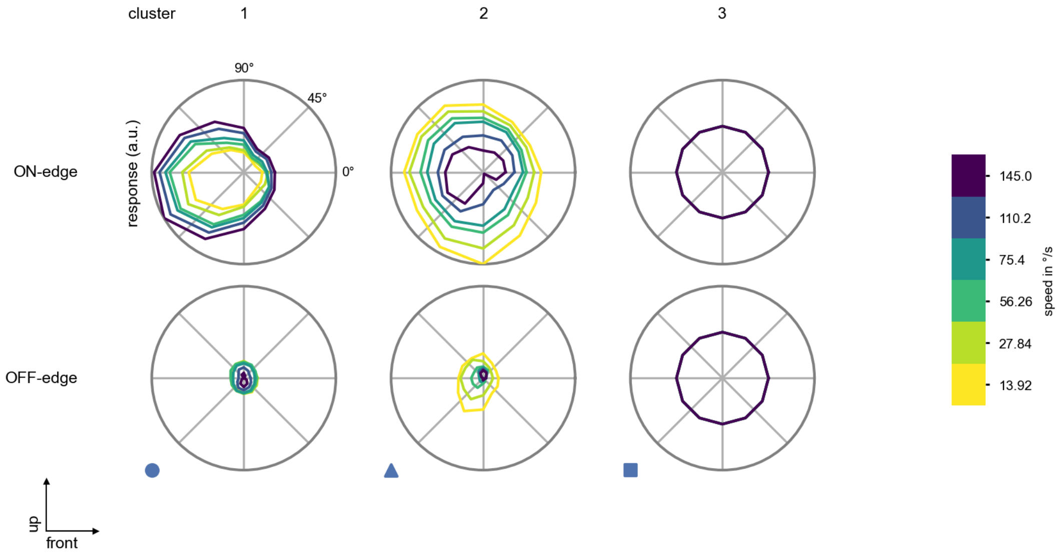

TmY14 - Figure 6: **Peak responses to moving edges from best-performing models.** The top row shows peak responses to moving ON-edges, the bottom row shows peak responses to moving OFF-edges. The edge-stimuli move in different directions from 0 to 360 degrees and at different speeds. Before, a grey-stimulus led the network to a stationary state. The columns show responses from the best-task-performing model in the respective cluster.

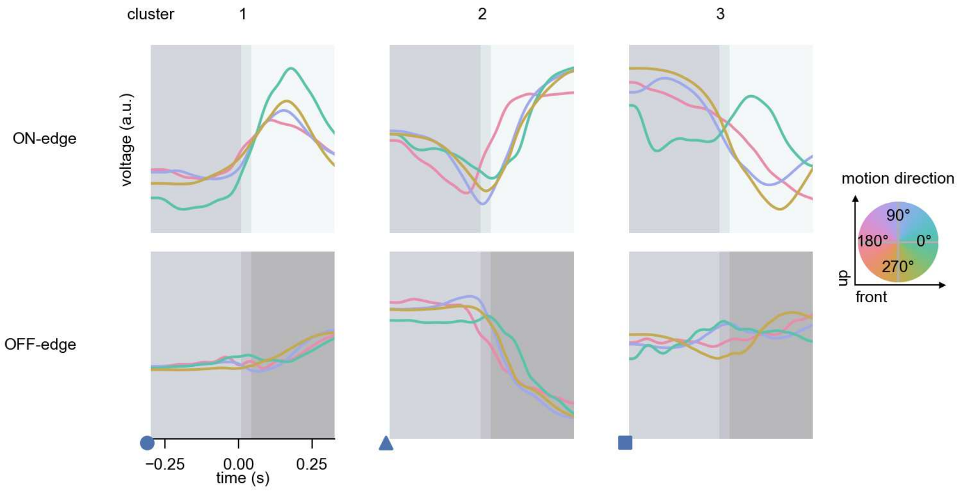

TmY14 - Figure 7: **Responses to moving edges from best-performing models.** The top row shows responses to moving ON-edges, the bottom row shows peak responses to moving OFF-edges. The edge-stimuli move in different directions from 0 to 360 degrees and at different speeds. Before, a grey-stimulus led the network to a stationary state. The columns show responses from the best-task-performing model in the respective cluster. Example responses to edges moving at 75.4°/s in all cardinal directions (green 0°, blue 90°, red 180°, yellow 270°) from -22.5 to 22.5° visual angle. Background contour indicates the edge, passing by the central column at second zero.

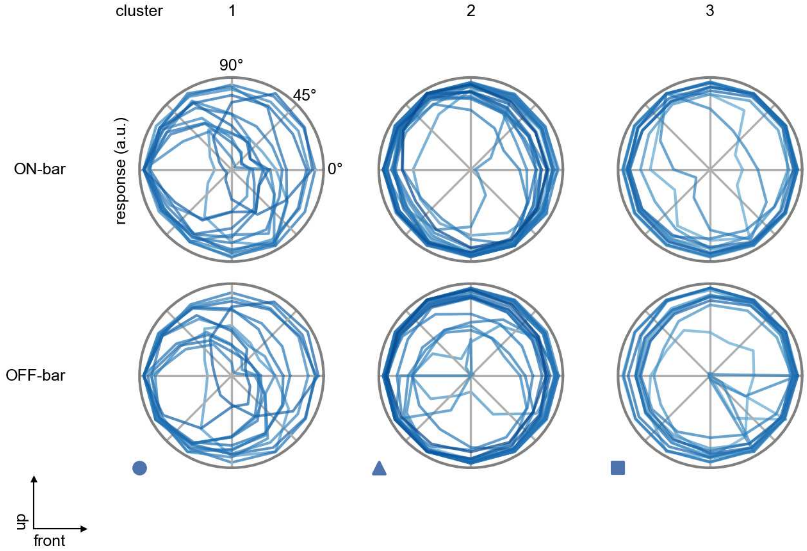

TmY14 - Figure 8: **Peak responses to moving bars.** The top row shows normalized peak responses to moving ON-bars, the bottom row shows normalized peak responses to moving OFF-bars. The bar-stimuli move in different directions from 0 to 360 degrees. Before, a grey-stimulus led the network to a stationary state. The responses from the different models in the different clusters (columns) overlay and are colored in the respective task error of the model. The peak responses are averaged over all assessed bar-speeds.

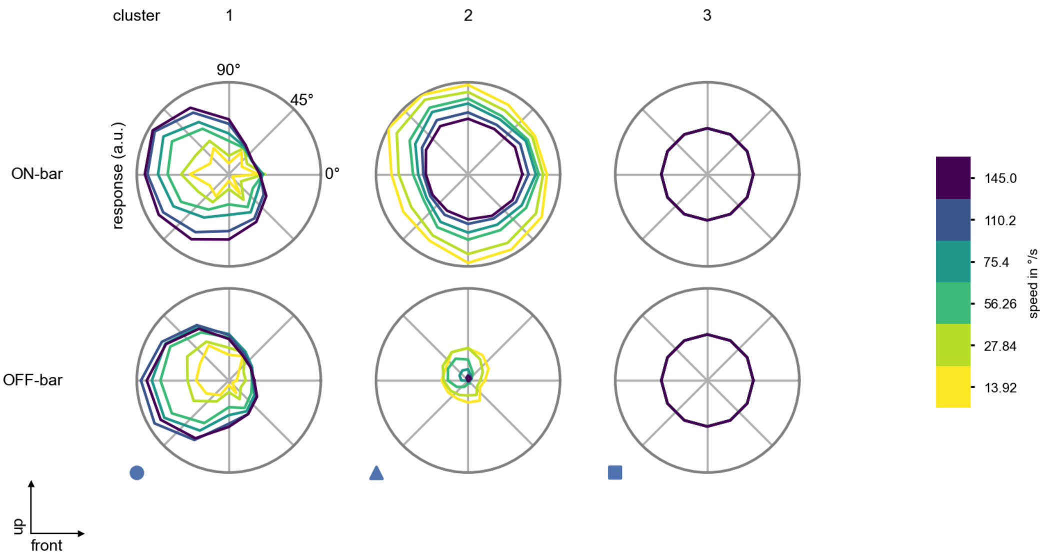

TmY14 - Figure 9: **Peak responses to moving bars from best-performing models.** The top row shows peak responses to moving ON-bars, the bottom row shows peak responses to moving OFF-bars. The bar-stimuli move in different directions from 0 to 360 degrees and at different speeds. Before, a grey-stimulus led the network to a stationary state. The columns show responses from the best-task-performing model in the respective cluster.

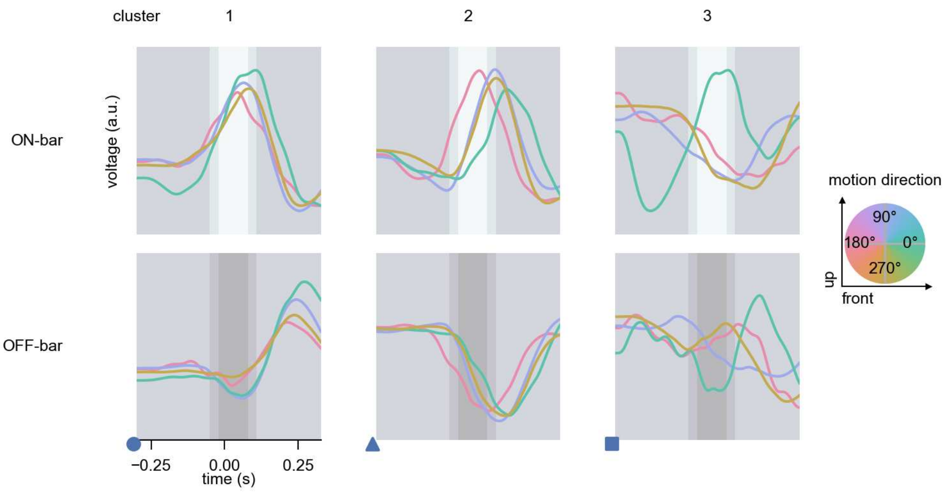

TmY14 - Figure 10: **Responses to moving bars from best-performing models.** The top row shows responses to moving ON-bars, the bottom row shows peak responses to moving OFF-bars. The bar-stimuli move in different directions from 0 to 360 degrees and at different speeds. Before, a grey-stimulus led the network to a stationary state. The columns show responses from the best-task-performing model in the respective cluster. Example responses to bars moving at 75.4°/s in all cardinal directions (green 0°, blue 90°, red 180°, yellow 270°) from -22.5 to 22.5° visual angle. Background contour indicates the bar, passing by the central column at second zero.

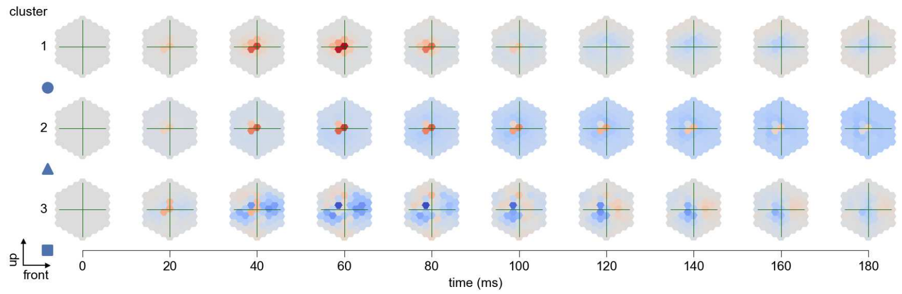

TmY14 - Figure 11: **Spatio-temporal receptive field.** Responses of the central cell to ON-impulses at single ommatidium impulse locations. The impulse occurs at second zero. Before, a grey-stimulus led the network to a stationary state. Responses are baseline-subtracted. Each row illustrates responses from the best-performing model of the respective cluster. Red indicated depolarization, blue indicates hyperpolarization.

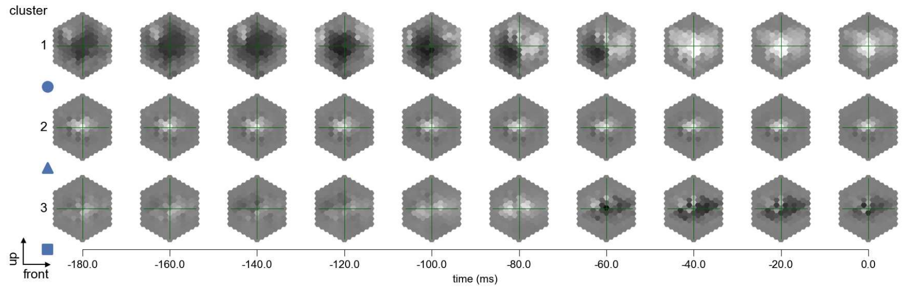

TmY14 - Figure 12: **Maximally excitatory stimuli.** Each row presents the regularized naturalistic-stimulus from the Sintel dataset that maximizes the cell type's central column response at second zero in the best-performing model of the respective cluster.

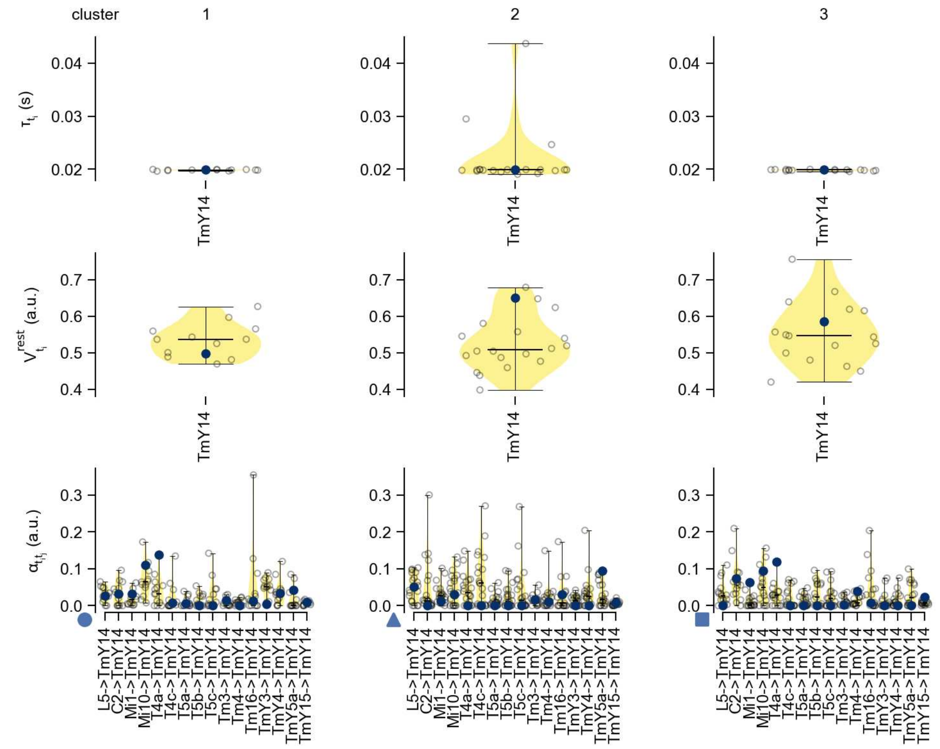

TmY14 - Figure 13: **Task-constrained parameters.** Each column shows the parameters inferred within the respective cluster. First row: learned time constants of the cell type. Second row: resting potentials of the cell type. Third row: scaling factors for the convolutional filters. The blue scatter represents the parameters from the best-performing model within the cluster.

### 64 TmY15

← Cell types

#### Figures

1 Anatomical receptive fields. . . . . 423
2 Anatomical projective fields. . . . . 424
3 Clustering of the responses to naturalistic stimuli. . . . . 424
4 Responses to flashes. . . . . 425
5 Peak responses to moving edges. . . . . 425
6 Peak responses to moving edges from best-performing models. . . . . 426
7 Responses to moving edges from best-performing models. . . . . 426
8 Peak responses to moving bars. . . . . 427
9 Peak responses to moving bars from best-performing models. . . . . 427
10 Responses to moving bars from best-performing models. . . . . 428
11 Spatio-temporal receptive field. . . . . 428
12 Maximally excitatory stimuli. . . . . 428
13 Task-constrained parameters. . . . . 429

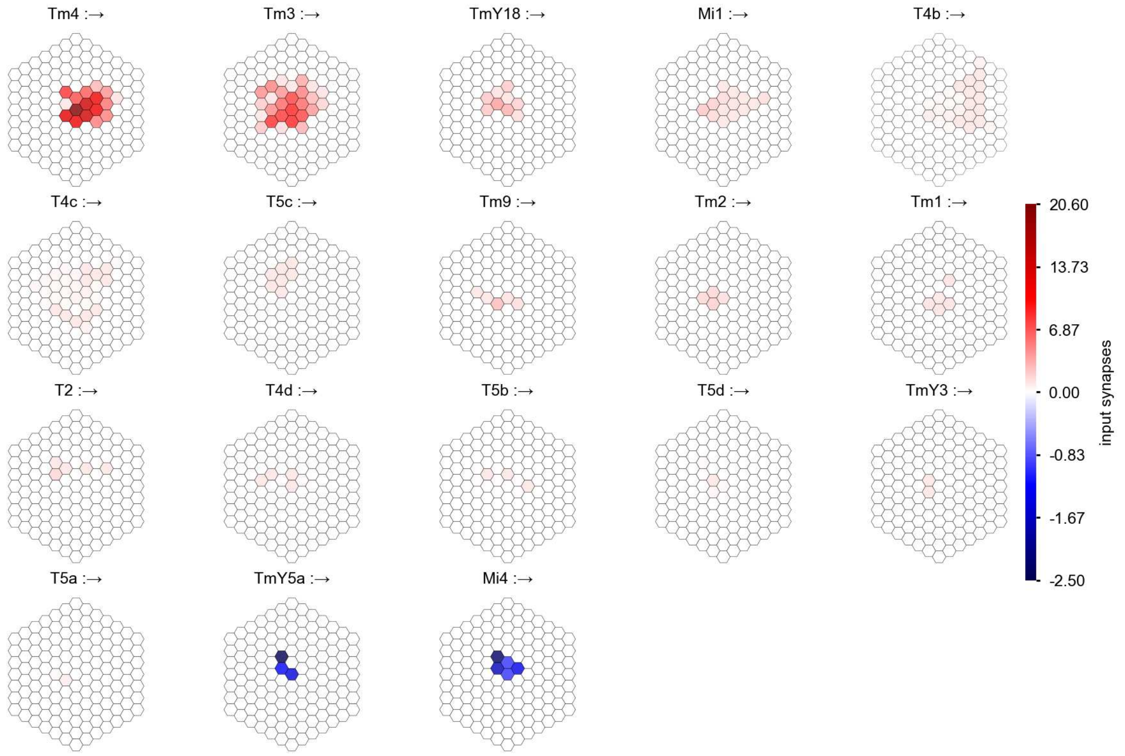

TmY15 - Figure 1: **Anatomical receptive fields.** Each colored hexagon is a projection, with the projection strength characterized by the average number of synapses that we count from the EM reconstruction. Red indicates excitatory synapses, blue indicates inhibitory synapses from inferred signs. Filters in the order of their total number of synapses.

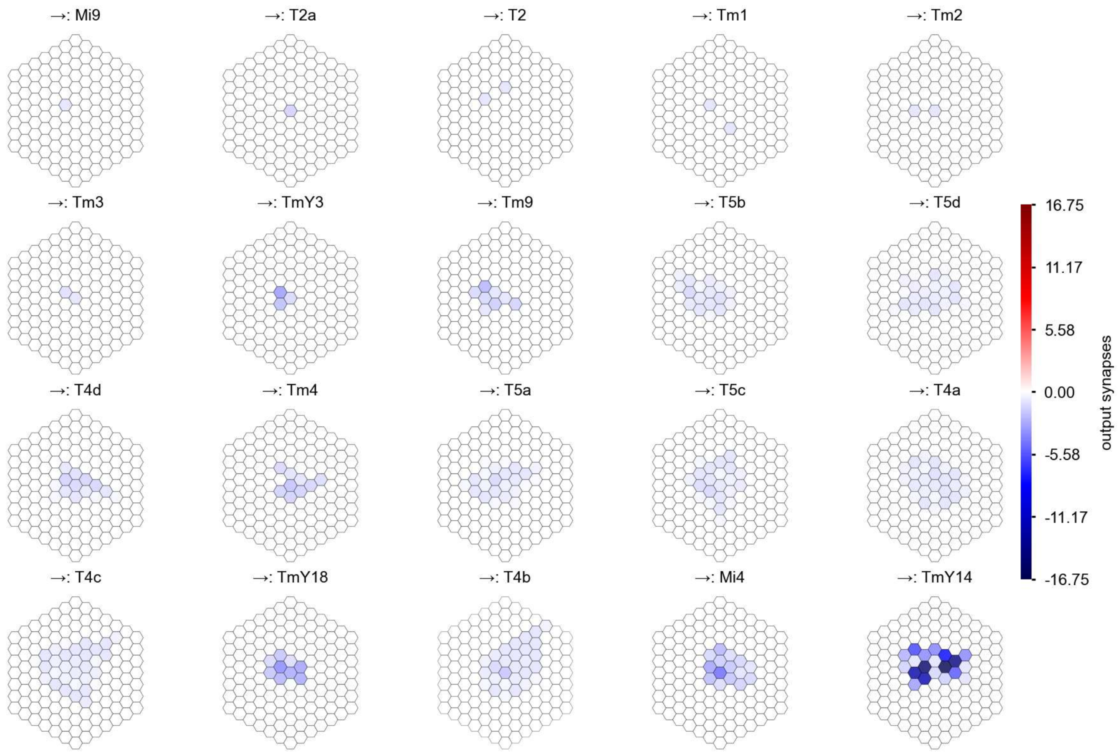

TmY15 - Figure 2: **Anatomical projective fields.** Each colored hexagon is a projection, with the projection strength characterized by the average number of synapses that we count from the EM reconstruction. Red indicates excitatory synapses, blue indicates inhibitory synapses from inferred signs. Filters in the order of their total number of synapses.

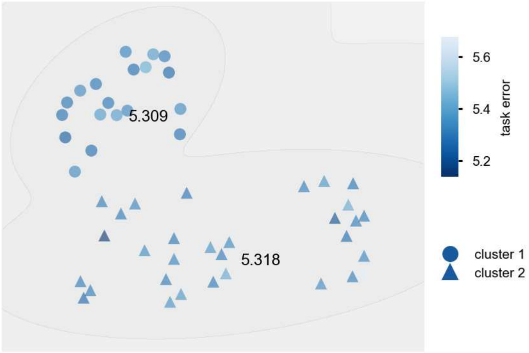

TmY15 - Figure 3: **Clustering of the responses to naturalistic stimuli.** From responses to naturalistic scenes from Sintel. Scatterpoints represent individual models colored by their task error.

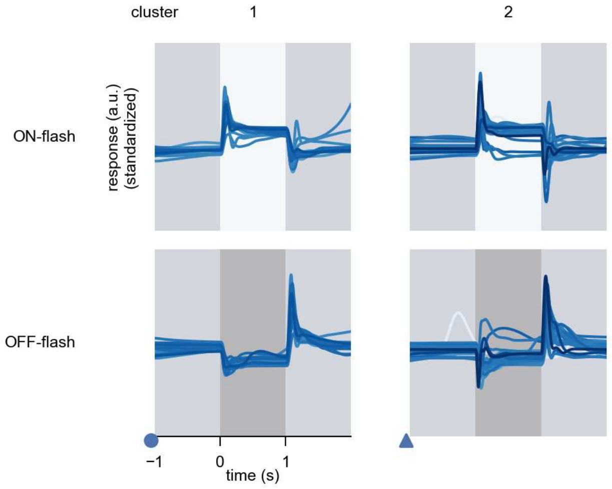

TmY15 - Figure 4: **Responses to flashes.** The top row shows responses to ON-flashes, the bottom row shows responses to OFF-flashes. The responses from the different models in the different clusters (columns) overlay and are colored in the respective task error of the model. The circular flashes occur at second zero, cover 6 ommatidia in radius, and last for one second. Before and after, a grey-stimulus leads to a stationary state of the network. The traces are standardized.

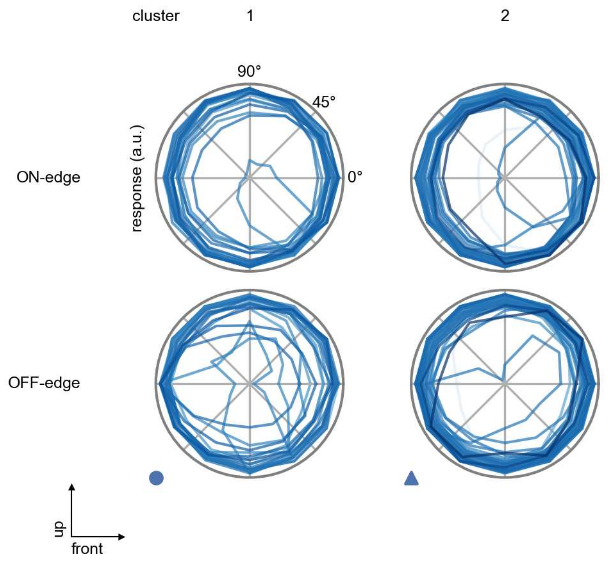

TmY15 - Figure 5: **Peak responses to moving edges.** The top row shows normalized peak responses to moving ON-edges, the bottom row shows normalized peak responses to moving OFF-edges. The edge-stimuli move in different directions from 0 to 360 degrees. Before, a grey-stimulus led the network to a stationary state. The responses from the different models in the different clusters (columns) overlay and are colored in the respective task error of the model. The peak responses are averaged over all assessed edge-speeds.

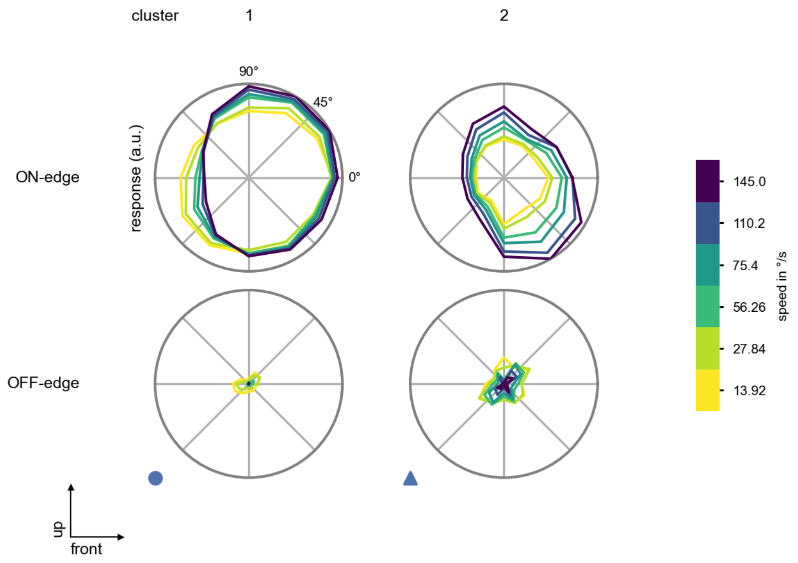

TmY15 - Figure 6: **Peak responses to moving edges from best-performing models.** The top row shows peak responses to moving ON-edges, the bottom row shows peak responses to moving OFF-edges. The edge-stimuli move in different directions from 0 to 360 degrees and at different speeds. Before, a grey-stimulus led the network to a stationary state. The columns show responses from the best-task-performing model in the respective cluster.

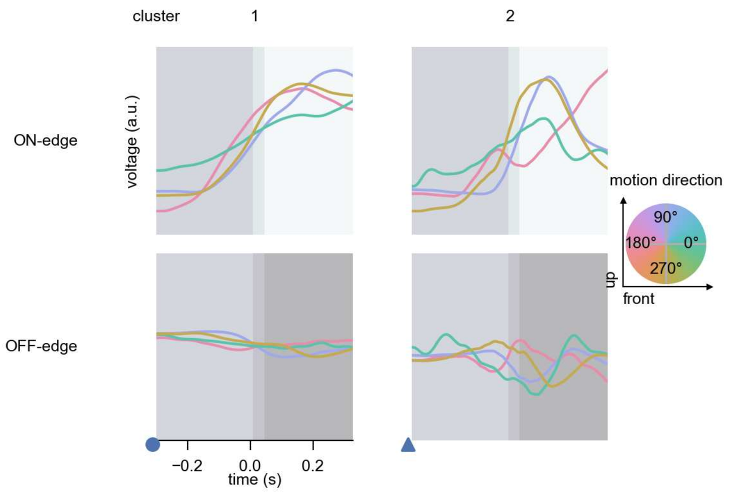

TmY15 - Figure 7: **Responses to moving edges from best-performing models.** The top row shows responses to moving ON-edges, the bottom row shows peak responses to moving OFF-edges. The edge-stimuli move in different directions from 0 to 360 degrees and at different speeds. Before, a grey-stimulus led the network to a stationary state. The columns show responses from the best-task-performing model in the respective cluster. Example responses to edges moving at 75.4°/s in all cardinal directions (green 0°, blue 90°, red 180°, yellow 270°) from -22.5 to 22.5° visual angle. Background contour indicates the edge, passing by the central column at second zero.

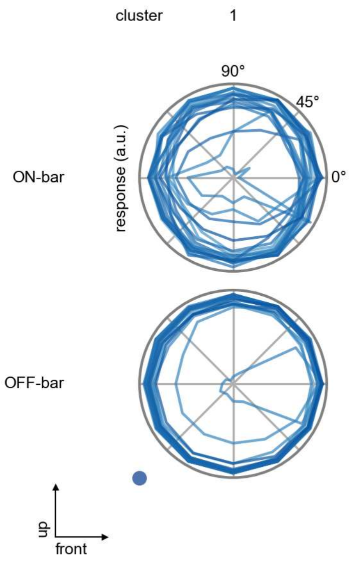

TmY15 - Figure 8: **Peak responses to moving bars.** The top row shows normalized peak responses to moving ON-bars, the bottom row shows normalized peak responses to moving OFF-bars. The bar-stimuli move in different directions from 0 to 360 degrees. Before, a grey-stimulus led the network to a stationary state. The responses from the different models in the different clusters (columns) overlay and are colored in the respective task error of the model. The peak responses are averaged over all assessed bar-speeds.

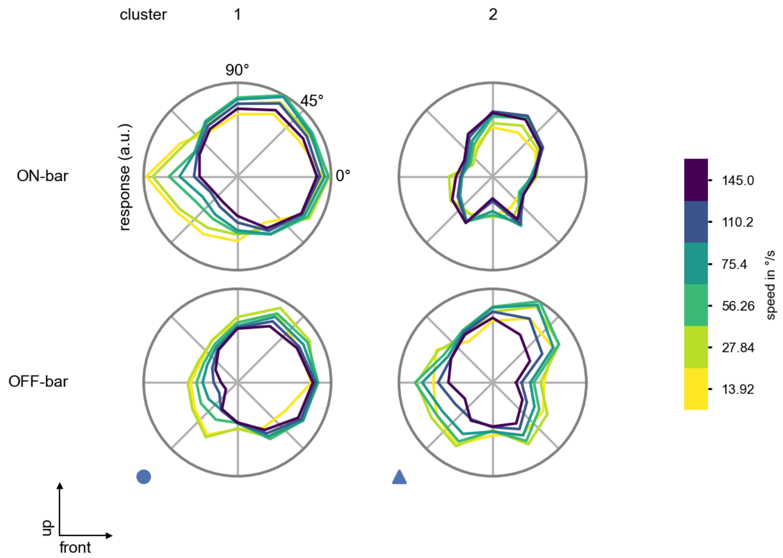

TmY15 - Figure 9: **Peak responses to moving bars from best-performing models.** The top row shows peak responses to moving ON-bars, the bottom row shows peak responses to moving OFF-bars. The bar-stimuli move in different directions from 0 to 360 degrees and at different speeds. Before, a grey-stimulus led the network to a stationary state. The columns show responses from the best-task-performing model in the respective cluster.

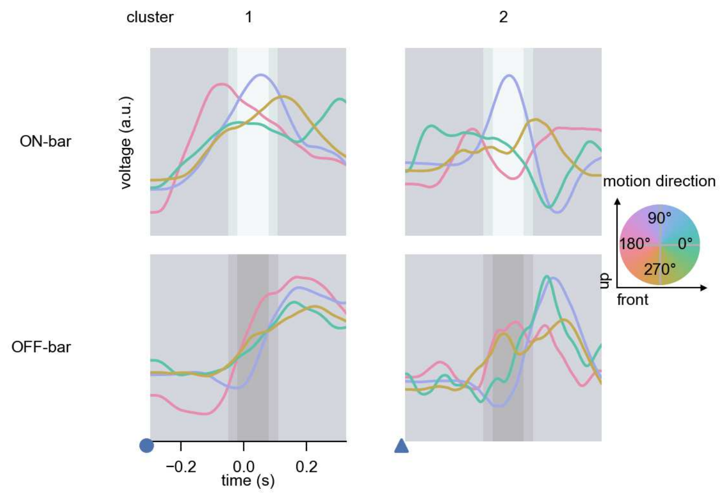

TmY15 - Figure 10: **Responses to moving bars from best-performing models.** The top row shows responses to moving ON-bars, the bottom row shows peak responses to moving OFF-bars. The bar-stimuli move in different directions from 0 to 360 degrees and at different speeds. Before, a grey-stimulus led the network to a stationary state. The columns show responses from the best-task-performing model in the respective cluster. Example responses to bars moving at 75.4°/s in all cardinal directions (green 0°, blue 90°, red 180°, yellow 270°) from -22.5 to 22.5° visual angle. Background contour indicates the bar, passing by the central column at second zero.

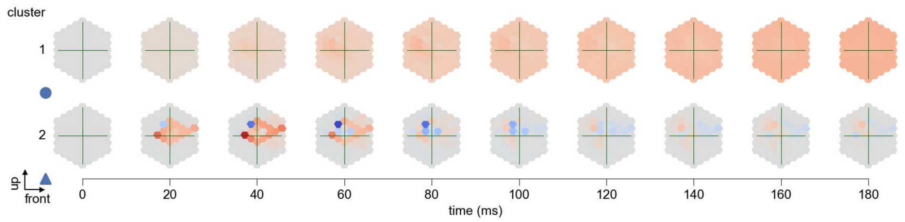

TmY15 - Figure 11: **Spatio-temporal receptive field.** Responses of the central cell to ON-impulses at single ommatidium impulse locations. The impulse occurs at second zero. Before, a grey-stimulus led the network to a stationary state. Responses are baseline-subtracted. Each row illustrates responses from the best-performing model of the respective cluster. Red indicated depolarization, blue indicates hyperpolarization.

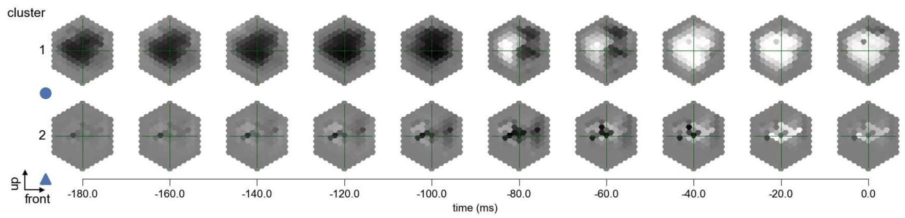

TmY15 - Figure 12: **Maximally excitatory stimuli.** Each row presents the regularized naturalistic-stimulus from the Sintel dataset that maximizes the cell type's central column response at second zero in the best-performing model of the respective cluster.

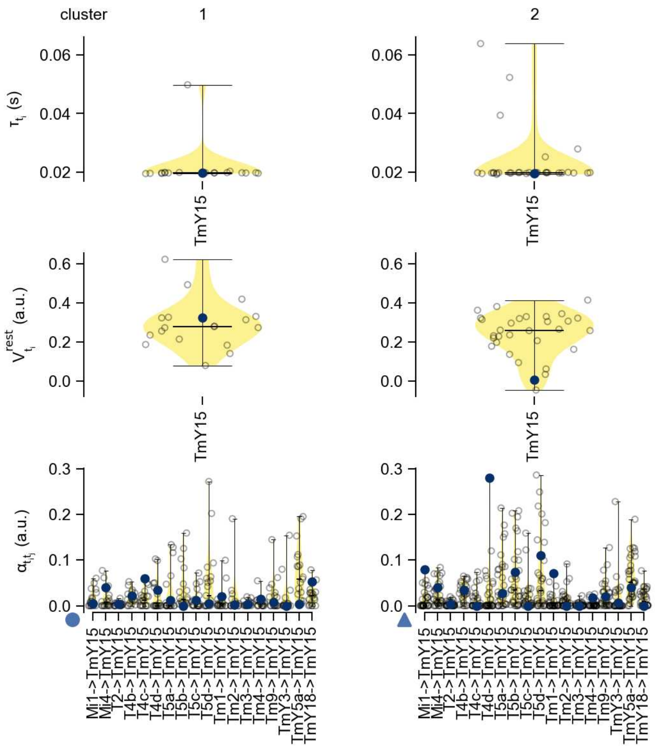

TmY15 - Figure 13: **Task-constrained parameters.** Each column shows the parameters inferred within the respective cluster. First row: learned time constants of the cell type. Second row: resting potentials of the cell type. Third row: scaling factors for the convolutional filters. The blue scatter represents the parameters from the best-performing model within the cluster.

### 65 TmY18

← Cell types

#### Figures

1 Anatomical receptive fields. . . . . 430
2 Anatomical projective fields. . . . . 431
3 Clustering of the responses to naturalistic stimuli. . . . . 431
4 Responses to flashes. . . . . 431
5 Peak responses to moving edges. . . . . 432
6 Peak responses to moving edges from best-performing models. . . . . 432
7 Responses to moving edges from best-performing models. . . . . 433
8 Peak responses to moving bars. . . . . 433
9 Peak responses to moving bars from best-performing models. . . . . 434
10 Responses to moving bars from best-performing models. . . . . 434
11 Spatio-temporal receptive field. . . . . 435
12 Maximally excitatory stimuli. . . . . 435
13 Task-constrained parameters. . . . . 436

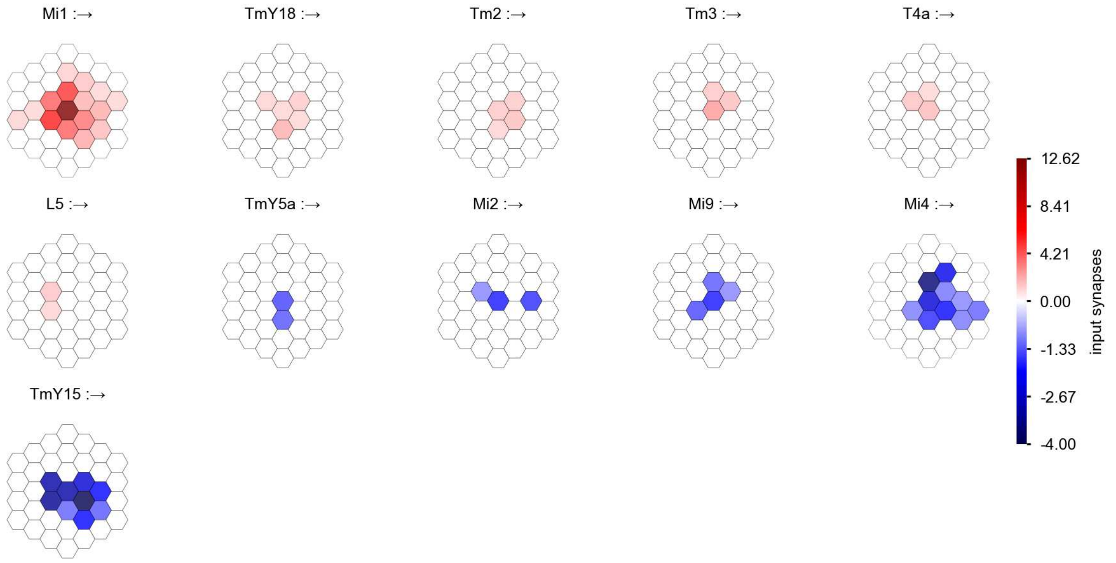

TmY18 - Figure 1: **Anatomical receptive fields.** Each colored hexagon is a projection, with the projection strength characterized by the average number of synapses that we count from the EM reconstruction. Red indicates excitatory synapses, blue indicates inhibitory synapses from inferred signs. Filters in the order of their total number of synapses.

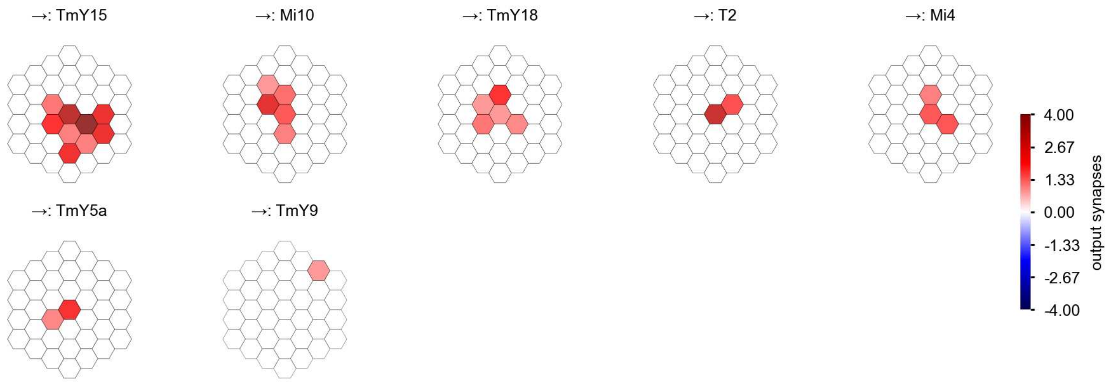

TmY18 - Figure 2: **Anatomical projective fields.** Each colored hexagon is a projection, with the projection strength characterized by the average number of synapses that we count from the EM reconstruction. Red indicates excitatory synapses, blue indicates inhibitory synapses from inferred signs. Filters in the order of their total number of synapses.

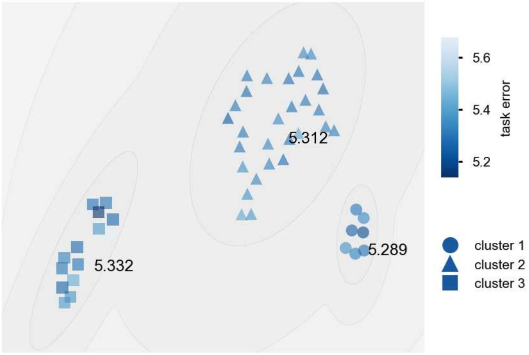

TmY18 - Figure 3: **Clustering of the responses to naturalistic stimuli.** From responses to naturalistic scenes from Sintel. Scatterpoints represent individual models colored by their task error.

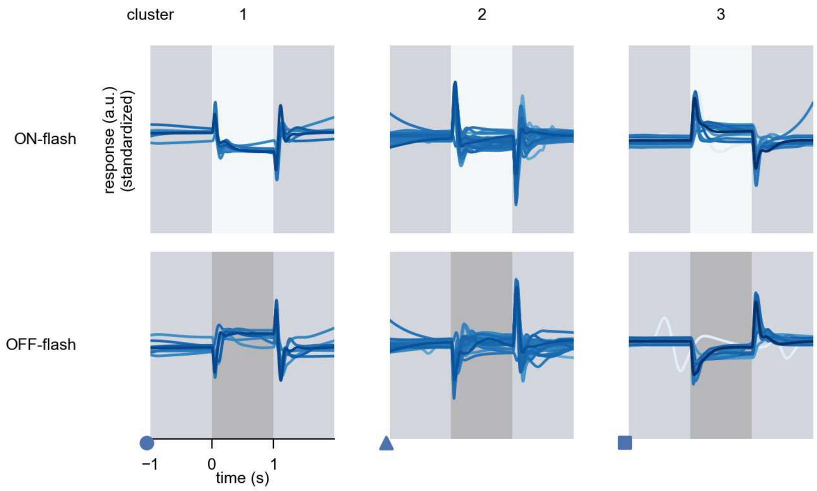

TmY18 - Figure 4: **Responses to flashes.** The top row shows responses to ON-flashes, the bottom row shows responses to OFF-flashes. The responses from the different models in the different clusters (columns) overlay and are colored in the respective task error of the model. The circular flashes occur at second zero, cover 6 ommatidia in radius, and last for one second. Before and after, a grey-stimulus leads to a stationary state of the network. The traces are standardized.

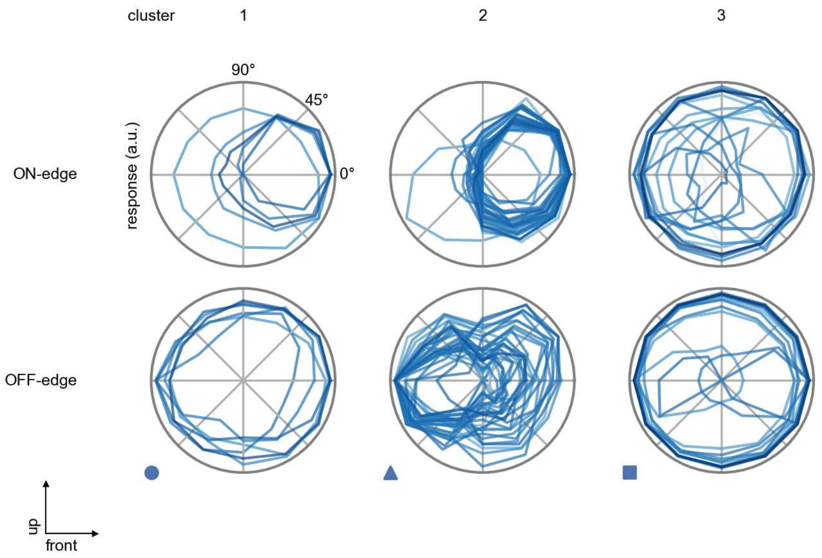

TmY18 - Figure 5: **Peak responses to moving edges.** The top row shows normalized peak responses to moving ON-edges, the bottom row shows normalized peak responses to moving OFF-edges. The edge-stimuli move in different directions from 0 to 360 degrees. Before, a grey-stimulus led the network to a stationary state. The responses from the different models in the different clusters (columns) overlay and are colored in the respective task error of the model. The peak responses are averaged over all assessed edge-speeds.

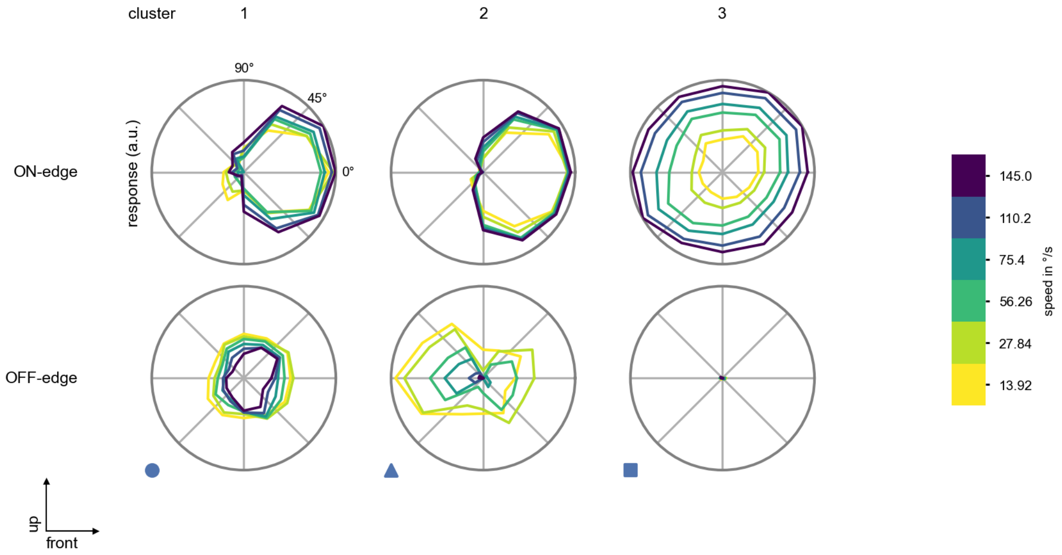

TmY18 - Figure 6: **Peak responses to moving edges from best-performing models.** The top row shows peak responses to moving ON-edges, the bottom row shows peak responses to moving OFF-edges. The edge-stimuli move in different directions from 0 to 360 degrees and at different speeds. Before, a grey-stimulus led the network to a stationary state. The columns show responses from the best-task-performing model in the respective cluster.

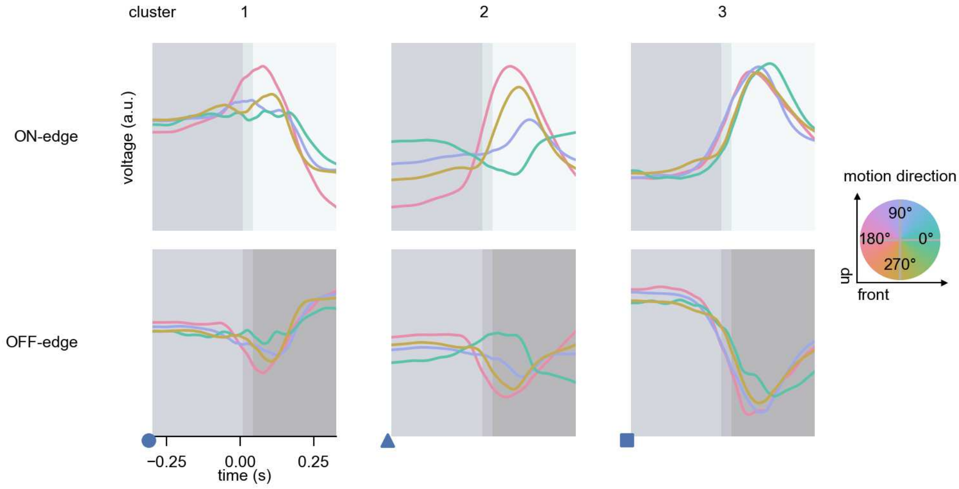

TmY18 - Figure 7: **Responses to moving edges from best-performing models.** The top row shows responses to moving ON-edges, the bottom row shows peak responses to moving OFF-edges. The edge-stimuli move in different directions from 0 to 360 degrees and at different speeds. Before, a grey-stimulus led the network to a stationary state. The columns show responses from the best-task-performing model in the respective cluster. Example responses to edges moving at 75.4°/s in all cardinal directions (green 0°, blue 90°, red 180°, yellow 270°) from -22.5 to 22.5° visual angle. Background contour indicates the edge, passing by the central column at second zero.

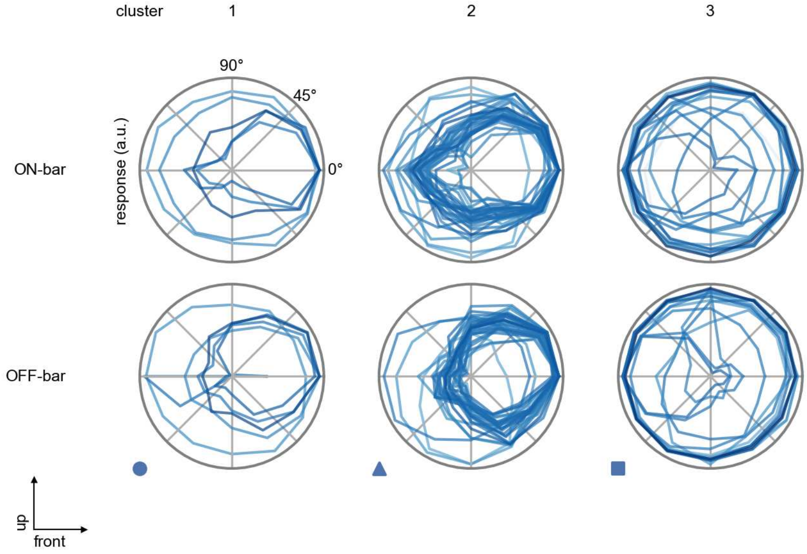

TmY18 - Figure 8: **Peak responses to moving bars.** The top row shows normalized peak responses to moving ON-bars, the bottom row shows normalized peak responses to moving OFF-bars. The bar-stimuli move in different directions from 0 to 360 degrees. Before, a grey-stimulus led the network to a stationary state. The responses from the different models in the different clusters (columns) overlay and are colored in the respective task error of the model. The peak responses are averaged over all assessed bar-speeds.

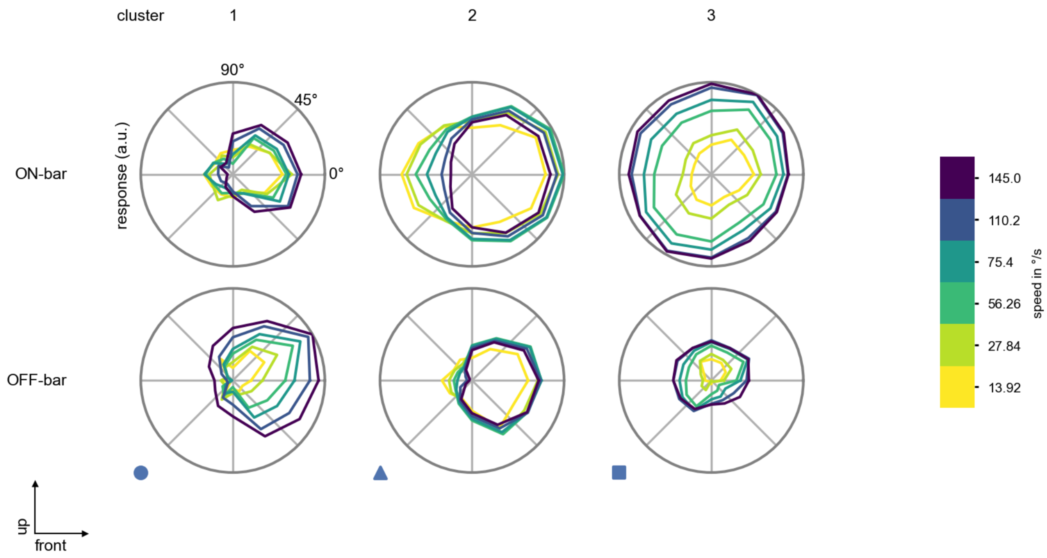

TmY18 - Figure 9: **Peak responses to moving bars from best-performing models.** The top row shows peak responses to moving ON-bars, the bottom row shows peak responses to moving OFF-bars. The bar-stimuli move in different directions from 0 to 360 degrees and at different speeds. Before, a grey-stimulus led the network to a stationary state. The columns show responses from the best-task-performing model in the respective cluster.

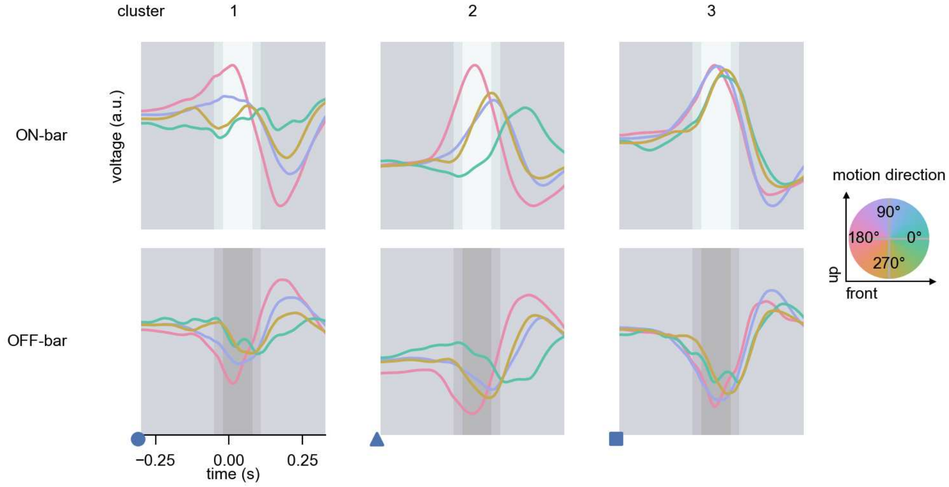

TmY18 - Figure 10: **Responses to moving bars from best-performing models.** The top row shows responses to moving ON-bars, the bottom row shows peak responses to moving OFF-bars. The bar-stimuli move in different directions from 0 to 360 degrees and at different speeds. Before, a grey-stimulus led the network to a stationary state. The columns show responses from the best-task-performing model in the respective cluster. Example responses to bars moving at 75.4°/s in all cardinal directions (green 0°, blue 90°, red 180°, yellow 270°) from -22.5 to 22.5° visual angle. Background contour indicates the bar, passing by the central column at second zero.

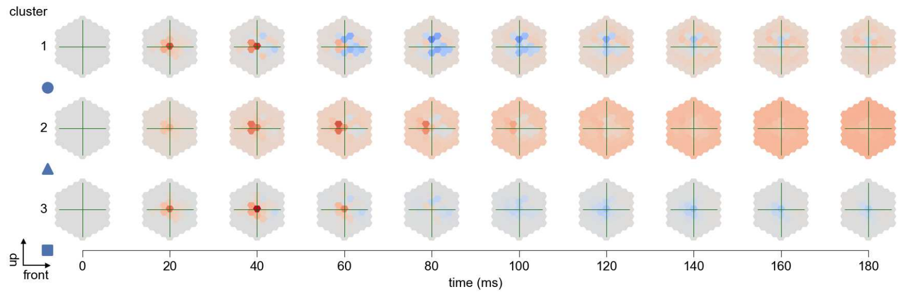

TmY18 - Figure 11: **Spatio-temporal receptive field.** Responses of the central cell to ON-impulses at single ommatidium impulse locations. The impulse occurs at second zero. Before, a grey-stimulus led the network to a stationary state. Responses are baseline-subtracted. Each row illustrates responses from the best-performing model of the respective cluster. Red indicated depolarization, blue indicates hyperpolarization.

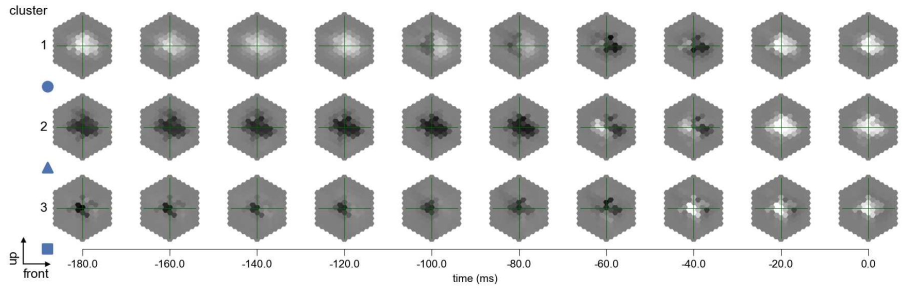

TmY18 - Figure 12: **Maximally excitatory stimuli.** Each row presents the regularized naturalistic-stimulus from the Sintel dataset that maximizes the cell type's central column response at second zero in the best-performing model of the respective cluster.

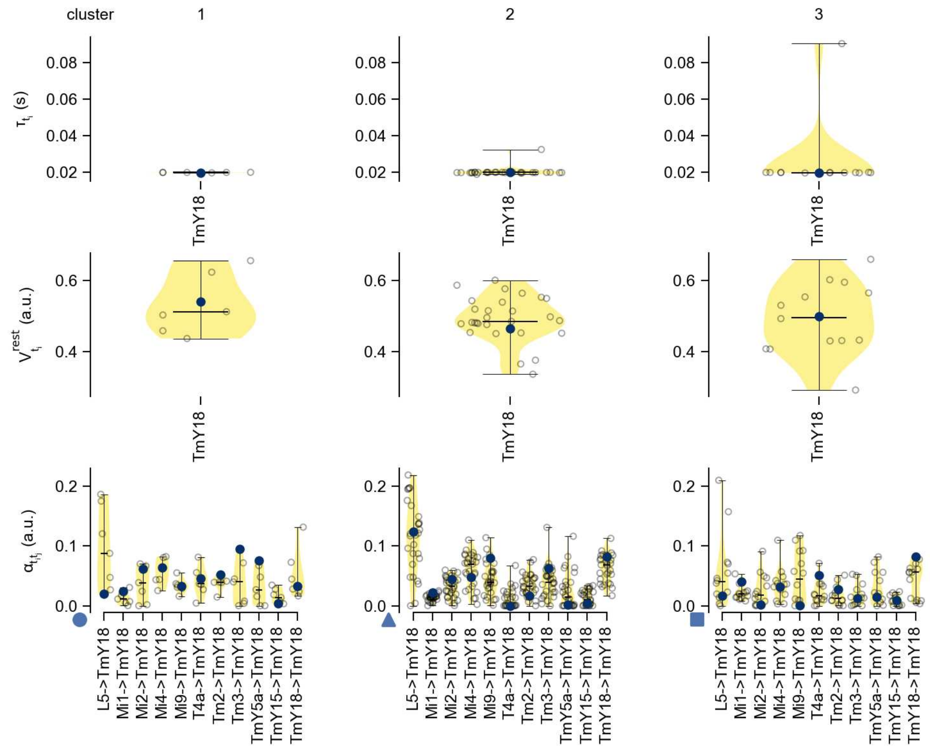

TmY18 - Figure 13: **Task-constrained parameters.** Each column shows the parameters inferred within the respective cluster. First row: learned time constants of the cell type. Second row: resting potentials of the cell type. Third row: scaling factors for the convolutional filters. The blue scatter represents the parameters from the best-performing model within the cluster.
